# Cell Senescence-Independent Ageing of Human Skin

**DOI:** 10.1101/2022.08.15.504045

**Authors:** J. Wordsworth, N. Fullard, C. Welsh, V. Maltman, C. Bascom, R. Tasseff, R. Isfort, L. Costello, R. Scanlan, S. Przyborski, D. Shanley

## Abstract

Skin ageing is defined in part by collagen depletion and fragmentation that leads to a loss of mechanical tension. This is currently believed to reflect, in part, the accumulation of senescent cells. We compared the expression of genes and proteins for components of the extracellular matrix (ECM) as well as their regulators and found that senescent cells produced more matrix metalloproteinases (MMPs) than proliferating cells from adult and neonatal donors. This was consistent with senescent cells contributing to increased matrix degradation with age; however, cells from adult donors proved significantly less capable of producing new collagen than neonatal or senescent cells, and they showed significantly lower myofibroblast activation as determined by the marker α-SMA. Functionally, adult cells also showed slower migration than neonatal cells. We concluded that while increased collagen degradation with age might reflect senescent cell accumulation, the reduced collagen production that prevents the skin from maintaining homeostasis must reflect senescence-independent processes.

## Introduction

Skin is the largest organ in the human body, performing numerous functions beyond that of a barrier to environmental stress. For example, skin is involved in sensory perception, thermoregulation, and immunosurveillance. Like other tissues, skin is subject to intrinsic ageing, where the ability of skin to perform its functions is diminished as a result of intrinsic changes. Phenotypically, old skin is dry, rough, and itchy, with uneven pigmentation, reduced capacity for wound healing, wrinkles, and impaired collagen and elastin networks (Shin et al., 2019). Old skin has diminished hair growth and sebaceous gland function, flatter dermal papillae, reduced melanocyte concentrations and less cellular turnover compared with young skin (Farage et al., 2013; Liao et al., 2014; Velarde et al., 2012). Skin ageing is the most visible aspect of ageing and the resulting phenotypic changes impact both physiological and psychological well-being (Blume-Peytavi et al., 2016).

Skin is a multi-layered tissue composed of an outer epidermis and an underlying dermis. The epidermis is comprised mainly of keratinocytes which are avascular and form a highly organized and stratified structure, with proliferative basal keratinocytes differentiating as they progress towards the skin surface. Beneath the epidermis lies the dermal-epidermal junction which is a thin basement membrane that enables communication between the dermis and epidermis. The dermis is a vascular, cell-sparse tissue composed mostly of a fibrous connective tissue known as the dermal extracellular matrix (ECM). Dermal tissue is essential for providing structural integrity to the skin and nourishing support for the epidermis (Lu et al., 2011).

Dermal fibroblasts are responsible for synthesising ECM components as well as proteins which degrade them; the balance between these two opposing functions regulates the ECM in both healthy skin and damaged or aged skin. Fibroblasts respond to local mechanical forces (Varani et al., 2006) as well as biochemical cues such as stimulation by TGF-β, which at higher levels can induce differentiation into myofibroblasts (Desmoulière, 1995). Myofibroblasts are metabolically active cells which secrete high levels of ECM proteins such as collagens 1 and 3 (Klingberg et al., 2013). The composition of the dermal microenvironment changes as we age. Fibroblasts in young tissue reside in an environment under mechanical tension which arises from physical binding to the ECM (Fisher et al., 2008a), but with age collagen fibrils become increasingly fragmented and fewer in number (Smith et al., 1962), resulting from both decreased production and enhanced degradation by matrix metalloproteinases (MMPs) such as MMP1 (Fisher et al., 2009; Fisher et al., 2008b; Varani et al., 2006; Vedrenne et al., 2012). Currently, little is known about how the capacity of fibroblasts to differentiate into myofibroblasts changes with age, although there is some evidence pointing to reduced hyaluronan synthesis and subsequent loss of CD44/epidermal growth factor (EGF) receptor signalling, resulting in impaired TGF-β1 dependent fibroblast activation (Simpson et al., 2010).

The aged dermal environment also contains an increasing population of senescent fibroblasts (Dimri et al., 1995; Ressler et al., 2006), and it is becoming increasingly accepted that “skin aging is significantly enforced by the accumulation of senescent dermal fibroblasts” (Wlaschek et al., 2021). Here, we compared ageing and senescent fibroblasts to assess their impact on TGF-β signalling and myofibroblast differentiation, and our results indicated that while senescent cells may contribute to increased matrix degradation, they did not affect the depleted response of adult fibroblasts to TGF-β or their reduced myofibroblast differentiation, which seemed instead to reflect an altered TGF-β homeostasis.

## Methods

### Cell Culture

Cultured fibroblast lines included three independent human neonatal dermal fibroblasts (HDFn) labelled A (Caucasian male, catalogue number: C-004-5C, lot number: #1366434), B (Caucasian male, catalogue number: C-004-5C, lot number: #1366356), and C (Caucasian male, catalogue number: C-004-5C, lot number: #1206197), and three independent adult fibroblast lines labelled G (55 years old Caucasian female, catalogue number: C-013-5C, lot number: #1528526), H (60 year old, Caucasian male, catalogue number: A11634, lot number: #1090465), I (65 year old, Caucasian female, catalogue number: A11636, lot number: #200910-901) purchased from Life Technologies, and one adult cell line lablled J (59 year old, Caucasian female, catalogue number: CC-2511, lot number: #693503) purchased from Lonza.

Senescence was induced in cells with 20Gy X-irradiation ten days prior to seeding. Cells were seeded into standard tissue culture treated 12-well dishes at a density of 10,000 cells/cm^2^, proliferative, or 65,000 cells/cm^2^, senescent, in 3.5ml M106 medium (ThermoFisher Scientific, catalogue number: M106500) supplemented with low serum growth supplement (LSGS, ThermoFisher Scientific, catalogue number: S00310) at 37°C, 5%CO_2_ for 4 days.

### Treatment Protocol

Cells from each cell line were serum starved 24h prior to treatment by removing LSGS supplementation from media. Following 24h of incubation at 37°C and 5% CO_2_, cells were assigned to one of three treatments: baseline, control, or TGF-β. Baseline samples were not treated in any way prior to harvesting at experiment start (0h). TGF-β and control samples were treated with media containing 5ng mL^−1^ TGF-β1 reconstituted in 10mM citric acid/0.1% BSA or 10mM citric acid/0.1% BSA vehicle control respectively. In control and TGF-β groups, cells were harvested at 0.5, 1, 2, 3, 4, 8, 12, 24, 48, 72, 96 hours post treatment. All 216 conditions were repeated 6 times resulting in 1296 individual samples. Samples were shipped to Procter and Gamble (P&G), Cincinnati for quantification on a high throughput PCR Smart chip platform by WaferGen.

### 3D cell culture and treatment protocol

Human male neonatal (<14 days), young (<30 years), middle-aged (40-45 years), and old (60+ years) dermal fibroblasts (Life Technologies) were maintained in Dermal Fibroblast Growth Medium (DFGM) comprised of Medium 106 (Thermo Fisher Scientific), supplemented with LSGS (Thermo Fisher Scientific), 10 μg mL^−1^ gentamicin, and 0.25 μg mL^−1^ amphotericin B (Thermo Fisher Scientific) at 37°C in a 5% CO_2_ humidified incubator following the supplier’s instructions.

Dermal models were generated by seeding HDFs (5 × 10^5^ cells) onto inert porous polystyrene membranes (12-well Alvetex^®^ scaffold inserts, Reprocell) and incubating at 37°C in a 5% CO_2_ humidified incubator in DFGM supplemented with 5 ng mL^−1^ TGF-β1 (Thermo Fisher Scientific) and 100 μg mL^−1^ ascorbic acid. Dermal equivalents were maintained for 28 days, then harvested as appropriate for downstream processing. For the 3D culture analysis two additional housekeeping genes ACTB and GAPDH were also used for normalisation, then log fold change (LogFC) was calculated for all aged cells compared to neonatal.

### Normalization

The raw data (containing cycle threshold values) is available for download in Supplementary file 1. Each gene measured in each sample was normalised to peptidylprolyl isomerase A (PPIA) which is stationary throughout treatment with TGF-β. Equation (1) shows the calculation producing the 2^−ΔCT^ for each gene (g) at each timepoint. The 2^−ΔCT^ value allowed comparison of the relative gene expression levels between samples from different conditions. To compare the magnitude of response to the TGF-β signal (the ΔΔCTg), equation 2 further normalised the ΔCTg value by subtracting the value at time zero (CT_0_) from the value at time, t (CT_t_).

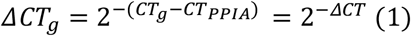

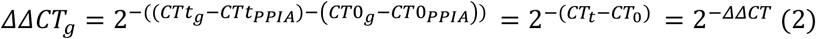

Additionally, because the adult senescent cells were not included in the initial experiment, another experiment was conducted using two of the three previous adult cell lines (H and I) plus a third cell line, J (cell line G was exhausted). To make the data comparable across experiments all data from the latter experiment were normalised by multiplication with factor γ. Factor γ was calculated as the average 2^−ΔCT^ value for experiment 1 divided by the average 2^−ΔCT^ for experiment 2, for each gene. Cell line J used the average value for each gene from cell lines H and I. However, graphs demonstrating the unnormalized 2^−ΔCT^ values for each gene in both experiments are found in Supplementary files 2 and 3.

### Western blots

Fibroblasts were washed with PBS, then lysed in M-PER^™^ Mammalian Protein Extraction Reagent (Thermo Fisher 78501). Samples were transferred to an Eppendorf tube and were heated to 95°C (10 minutes), sonicated, and centrifuged for 30 minutes at 10,000 × g and supernatant removed. Protein (15-30 μg) was resolved on either 8%, 10%, or 12% SDS-polyacrylamide gels, dependent upon molecular weight of target protein. Protein was transferred onto nitrocellulose membrane. Membranes were blocked for 45 minutes at room temperature with 5% non-fat milk powder in Tris-buffered saline–Tween. The following antibodies were used: collagen 1 (1:1000, Southern Biotech, 1310), collagen 3 (1:500; Proteintech, 22734-1-AP), α-SMA (1:25,000 Sigma #A4416), LOXL2 (1:000, AbCam, 179810), LTBP2 (1:250 Santa Cruz, 166199), TAGLN (1:2000, AbCam, 14106), elastin (1:1000, AbCam 77804), MMP2 (1:1000, AbCam, 37150), Smad3 (1:1000, Cell Signaling, 9523), P-Smad3 (1:1000, Cell Signaling, 9520), and β-actin (1:50,000, AbCam 8224), and incubated overnight at 4°C. After washing, membranes were incubated at room temperature for 1 hour with horseradish peroxidase (HRP) conjugated goat anti-mouse (1:10:000 Sigma 4416) or donkey anti-rabbit (1:10,000, Millipore, AP182P) diluted in 5% milk/Tris-buffered saline–Tween. Antigens were visualized using enhanced chemiluminescence (BioRad Clarity ECL; 1705061).

### Data analysis

All data were analysed using R version 3.6.3. Significance values shown in the tile plots were calculated by ANCOVA analysis adjusted with the Bonferroni correction. The R package, LIMMA (Smyth, 2004, 2005; Ritchie et al., 2015) was used to compute differential expression statistics. LIMMA was configured to using a spline matrix with 4 degrees of freedom, as per the LIMMA user guide, to compare either the control time series between adult/senescent and neonatal cell lines or the TGF-β treated cells. To make use of the available cell line replicates, differential expression calculation was repeated so that all adult/senescent cell lines were compared with all neonatal cell lines. The results were aggregated by calculating the percentage of times a gene has a false discovery rate (FDR) corrected p value less than 0.05. This was repeated for both control and TGF-β datasets and for both adult and senescent cell lines, compared with neonatal. Because our data is dynamic, for each comparison LIMMA was configured to evaluate whether two full time series objects were different from one other using a moderated F-statistic (Ritchie et al., 2015; Smyth, 2004, 2005). Since in experiment 1, we had data from 3 cell lines per treatment, we made use of the available data by repeating the LIMMA analysis for all combinations of cell line under a comparison. A total of 10 differential expression analyses were conducted, 5 each to compare TGF-β or negative control time series, two of which were ‘between’ groups comparisons and three ‘within’ groups comparisons. The output from LIMMA is a list of FDR-corrected p-values representing the probability that two groups are equal. Usually, a p-value threshold is chosen and any gene with a p-value lower than that threshold is considered differentially expressed. Since we had 9 equivalent LIMMA analyses for the ‘between groups’ analysis and 3 for the ‘within groups’ analysis, a consensus strategy was used where if a gene was considered differentially expressed in >60% of LIMMA analyses, the gene was considered differentially expressed.

### Fibroblast migration assay

Fibroblasts were seeded into 6 well plates at a density of 60,000 cells/cm^2^, in M106 medium (ThermoFisher Scientific, catalogue number: M106500) supplemented with LSGS (ThermoFisher Scientific, catalogue number: S00310) and incubated at 37°C, 5% CO_2_ for 24h. Fibroblasts were washed with PBS and serum starved for 24h by removing LSGS from media. Cells were then treated with Mitomycin C (Sigma, M0503) at 10μg/ml for 2h and washed in PBS before a scratch was made with a pipette tip and serum free M106 was added per well. Migration of cells into cell free area was monitored over 48h using a Zeiss cell observer.

## Results

To identify differences between neonatal and adult fibroblasts, the activity of 72 genes were measured over 96h in the presence or absence of TGF-β1 for three independent cell lines. Senescent cells were derived from the neonatal cell lines irradiated with 20Gy X-rays 10 days before seeding. The genes were manually selected based on a literature search to be relevant to ECM biology or TGF-β signalling. Of the 72 genes, 55 provided reliable data and were included in further analysis.

Principal component analysis (PCA) was first conducted to gain confidence in the measurements. In the results shown in Figure 1, each point on the PCA plot represents one of the 1296 samples. The high dimensional data is reduced to fewer dimensions whilst maintaining as much of the original variance as possible. Therefore, when two points are close together on the PCA, the underlying data are similar while two distant points have more different gene expression profiles. A scree plot (Figure S1) indicates that the first three principal components account for >80% of the variance. PCA plots for the first three principle components (PCs) were coloured for cell type (neonatal, senescent, or adult, Figure 1 A). Importantly, there was obvious clustering of the three cell types, with PC1 showing separation between the adult and the other two cell types, while PC2 showed separation of the neonatal from the other two cell types, and PC3 robustly separated senescent cells from neonatal and adult cell types. Cells also clustered according to cell line and showed no clustering according to biological replicate indicating no obvious batch effects (Figure S2 and S3). However, as shown in Figure 1 B, samples also clustered by timepoint.

**Figure 1.**
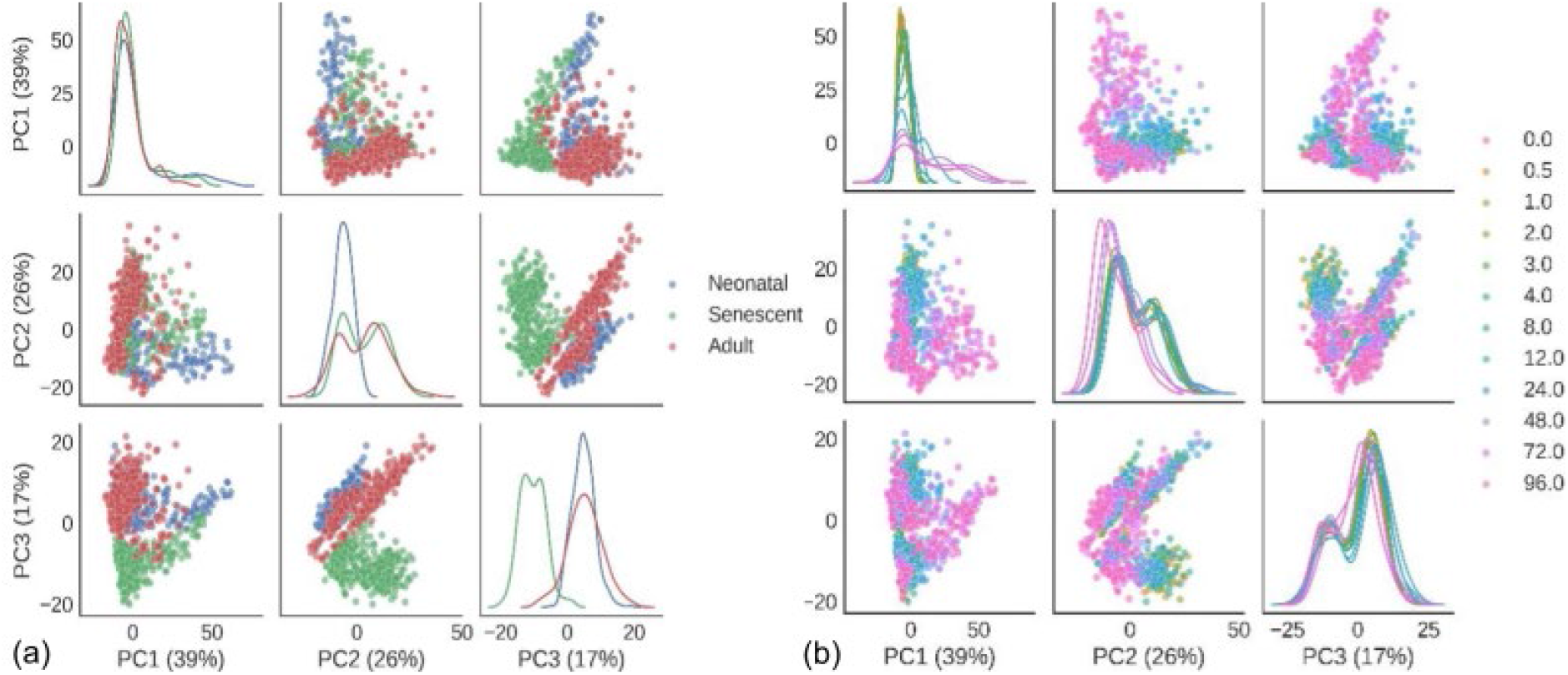
Principle component analysis of gene expression data. Each plot point reflects a summary of all gene expression for a single sample plotted for principle components 1-3 (PC1-3) which combined explain 82% of the variance. Scatter plots indicate the contribution to the variance across each PC. Kernel density plots (along the diagonal) represent the distribution of the relevant principle component. (A) Data is coloured by cell type. (B) Data is coloured by timepoint (0h to 96h).

Differential expression analysis was conducted to investigate whether the time series measurements for each gene was statistically different between or within cell line groups. A detailed summary of the differential expression analysis is presented in using a p-value threshold of 0.001 (Figure S4), while the number of genes that were differentially expressed in >60% of LIMMA analyses were counted and displayed in Table S1.

Both differential expression analysis and PCA analysis indicated that aged cells showed more differences to neonatal cells than did senescent cells. If ageing reflected in large part an increase in senescent cells and their effects (Wlaschek et al., 2021), then this outcome is surprising. While senescence is induced in the neonatal cell lines, they are still three different cell lines, so this variable is unlikely to explain the increased similarity.

To further explore this unpredicted relationship between ageing and senescence, we analysed the data for individual genes. An additional array was conducted for the same genes using three adult cell lines and inducing senescence in these cell lines. These data were then normalised (see methods) to make the data comparable across the two experiments. However, un-normalised data for both experiments are shown for each gene in Supplementary files 2 and 3.

### Ageing results in reduced collagen production independent of cell senescence

One clear change with age was the production of collagen mRNAs COL1A1, COL1A2, COL4A1 and COL5A1 as shown in Figure 2 A, B, C, and D. Importantly, all four mRNAs showed significant decreases in adult cells when compared with both neonatal and neonatal senescent cells. There was also no significant difference in COL4A1 between neonatal and neonatal senescent cells and a significant increase in the same mRNA from adult to adult senescent cells. All other collagen mRNAs showed significant decrease from neonatal to neonatal senescent, but not to the same extent as in adult cells, and no significant change was observed from adult to adult senescent cells. These observations are consistent with the observations of the increased extracellular matrix (ECM) degradation associated with senescent cells (Levi et al., 2020), but inconsistent with the idea that ageing tissues lose ECM through either an increase in senescent cells (Wlaschek et al., 2021) or progression of cells toward a “senescent-like” phenotype.

**Figure 2.**
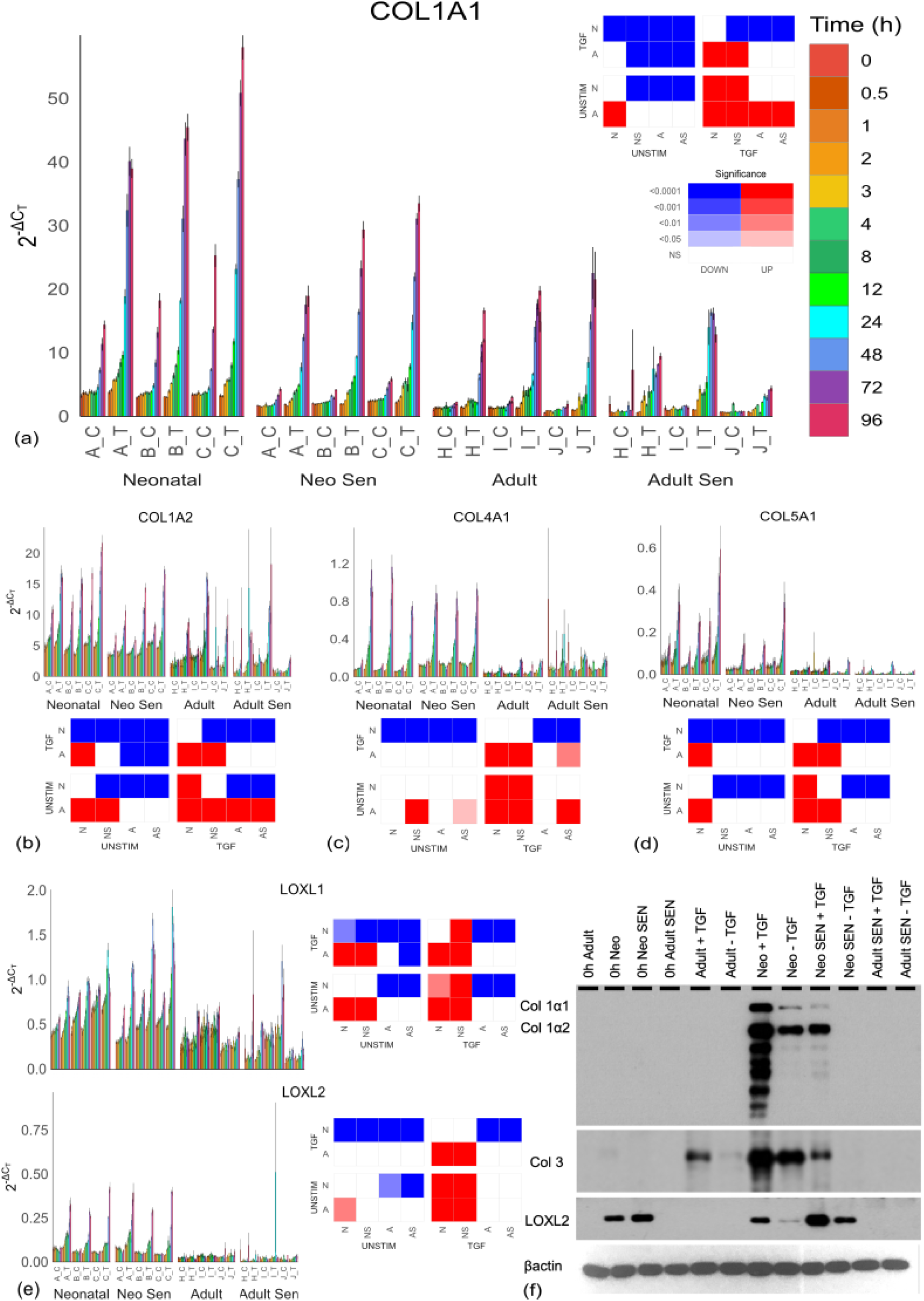
Collagen production and crosslinking by cell type. (A-E) Collagen and LOXL mRNAs for each cell line over the 96-hour timecourse. The x axes show neonatal cell lines A-C which include non-senescent (Neonatal) and senescent (Neo Sen) cells. Adult cell lines H-J were either non-senescent (Adult) or senescent (Adult Sen). After the letter indicating cell line (A-C and H-J), “_T” indicated stimulation with TGF-β, while “_C” indicated the unstimulated controls. Each graph has an accompanying tile plot indicating whether comparison of the changing profile over time was significantly different between cell types as determined by ANCOVA. The cell types: Adult, A; Adult senescent, AS; Neonatal, N; and neonatal senescent, NS were split by either TGF-β stimulation (TGF) or control (UNSTIM). Colour red indicates that the cell type along the x axis showed a significant increase compared to the cell type and condition depicted on the y axis, while blue reflects a significant decrease along the x axis, and white reflects no significant change (NS). Increasing colour intensity reflects increasing significance. (F) Western blot of collagen 1 and 3 and LOXL2 showing protein levels at zero hours and 96 hours with stimulation (+TGF) and without stimulation (−TGF) with TGF-β.

LOXL1 and LOXL2, which are involved in collagen crosslinking (Kagan and Li, 2003), are also stimulated by TGF-β and showed the same significant decline in expression in adult cells compared with neonatal cells (Figure 2 E), but LOXL2 does not decrease with senescence while LOXL1 significantly increased with senescence in neonatal cells. Western blotting of collagens 1 and 3, which are the main collagens in human skin (Naomi et al., 2021; Prockop and Kivirikko, 1995), after 96 hours of TGF-β stimulation (or control) confirmed production of both collagens was stimulated by TGF-β, but the amounts were reduced or absent in adult and adult senescent fibroblasts. We concluded that adult fibroblasts are less fibrotic than neonatal fibroblasts, resulting from a mechanism independent of cell senescence.

### Ageing and senescence increase matrix degradation

Next, we looked at matrix metalloproteinases (MMPs) involved in collagen degradation (Van Doren, 2015). No MMPs were significantly affected by TGF-β stimulation; however, as expected adult senescent cells showed significantly increased expression of MMP1 (Figure 3 A), MMP2 (Figure 3 B), and MMP14 (Supplementary file 4) compared to adult cells. What was less expected was that there was no difference between neonatal senescent and neonatal cells. Thus, the increased ECM degradation we see with age might not reflect the impact of senescent cells on ageing, but ageing on senescent cells. When we looked at the protein level (Figure 3 E), MMP1 showed a similar profile to its mRNA with higher expression in adult senescent cells than any other cell types; however, MMP2 protein was highest in the neonatal senescent cells and showed some evidence of TGF-β stimulation in the neonatal and adult senescent cells. While this complicated the profile seen in the mRNA, it was not inconsistent with the idea that older tissues might undergo more matrix degradation through an increase in senescent cells, as there was an increase in MMP2 expression in adult cells compared to neonatal cells, which was not as large as the increase in the fully senescent population.

**Figure 3.**
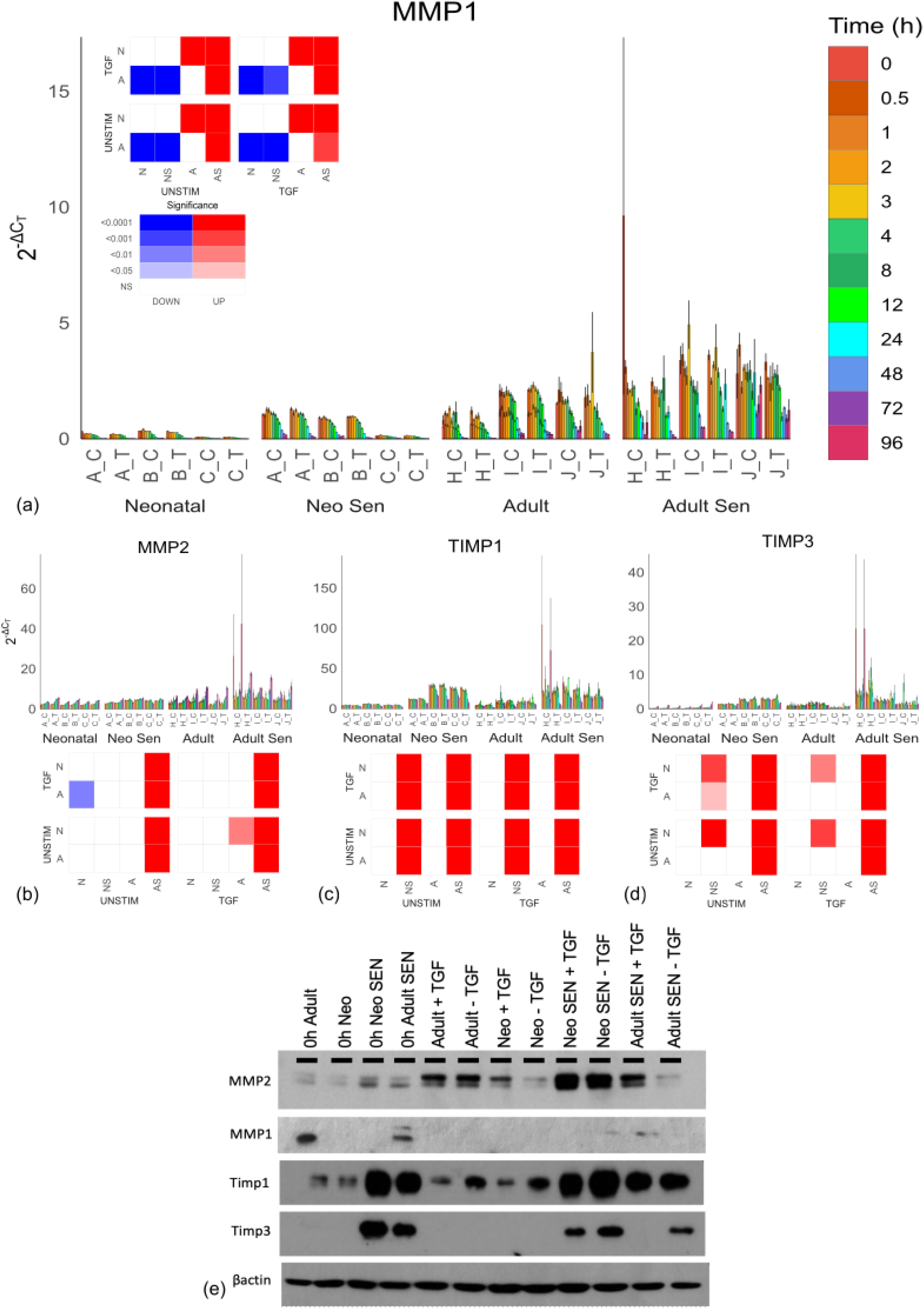
MMP and TIMP production by cell type. (A-D) Collagen and LOXL mRNAs for each cell line over the 96-hour timecourse. The x axes show neonatal cell lines A-C which include non-senescent (Neonatal) and senescent (Neo Sen) cells. Adult cell lines H-J were either non-senescent (Adult) or senescent (Adult Sen). After the letter indicating cell line (A-C and H-J), “_T” indicated stimulation with TGF-β, while “_C” indicated the unstimulated controls. Each graph has an accompanying tile plot indicating whether comparison of the changing profile over time was significantly different between cell types as determined by ANCOVA. The cell types: Adult, A; Adult Senescent, AS; Neonatal, N; and Neonatal Senescent, NS were split by either TGF-β stimulation (TGF) or control (UNSTIM). Colour red indicates that the cell type along the x axis showed a significant increase compared to the cell type and condition depicted on the y axis, while blue reflects a significant decrease along the x axis, and white reflects no significant change (NS). Increasing colour intensity reflects increasing significance. (E) Western blot of MMP1 and 2 and TIMPs 1 and 3 showing protein levels at zero hours and 96 hours with stimulation (+TGF) and without stimulation (−TGF) with TGF-β.

We also looked at the expression of tissue inhibitors of metalloproteinases (TIMPs) (Figure 3 C and D), which play a wide role in metalloproteinase inhibition (Brew and Nagase, 2010). At the mRNA level, both TIMPs showed no significant difference between aged and neonatal cells or TGF-β stimulation, but were significantly higher in senescent cells than non-senescent cells. We confirmed this at the protein level (Figure 3 E). Notably, while the increase in MMP2 in adult tissue might reflect increased senescent cells, it is not compensated for by an increase in TIMP production as senescent cells do.

Combined, these observations suggest two independent but synergistic effects of ageing fibroblasts on collagen in the ECM. Firstly, increased collagen degradation may reflect, in part, the accumulation of senescent cells, while reduced replenishment of the collagen matrix and cross-linking which protects it from degradation appear independent of cell senescence. However, these are in vitro observations from cells in monolayers. To test if ageing and senescent cells behaved similarly under more in vivo-like conditions, we constructed 3D Alvetex^®^ matrices (see methods) populated by cells from neonatal (< 14 days), young (<30 years), middle-aged (40-45 years), and old (60+ years) individuals, as well as senescent cells (induced from neonatal cells). We then carried out an array of similar ECM genes after exposure of the matrix to TGF-β just as for the monolayer (see Supplementary file 5 for all genes). The results for collagen mRNAs are shown in Figure 4 A-E. COL1A1 and COL5A1 showed significant decrease in expression in older cells compared to neonatal cells, while COL1A2, COL3A1, and COL4A1 showed no significant change with age. Even for COL1A1 and COL5A1 there was no evidence of continual decline in expression with age. However, all collagen mRNAs showed increased expression in senescent cells compared to neonatal cells, opposite to the effects of age. Thus, the decreases with age were unlikely to reflect higher numbers of senescent cells. The same was true for the LOXL genes (Figure 4 F-G). Strangely, MMP1 showed no significant change for ageing or senescent cells (Figure 4 H), while MMP2 and MMP14 (Figure 4 I-J) showed significant increases only in senescent cells, as was the case for the TIMP genes. These data were highly consistent with results from 2D culture, indicating that while senescent cells might be responsible for increased matrix degradation with age, reduced replacement of collagen likely reflects a senescence-independent process.

**Figure 4.**
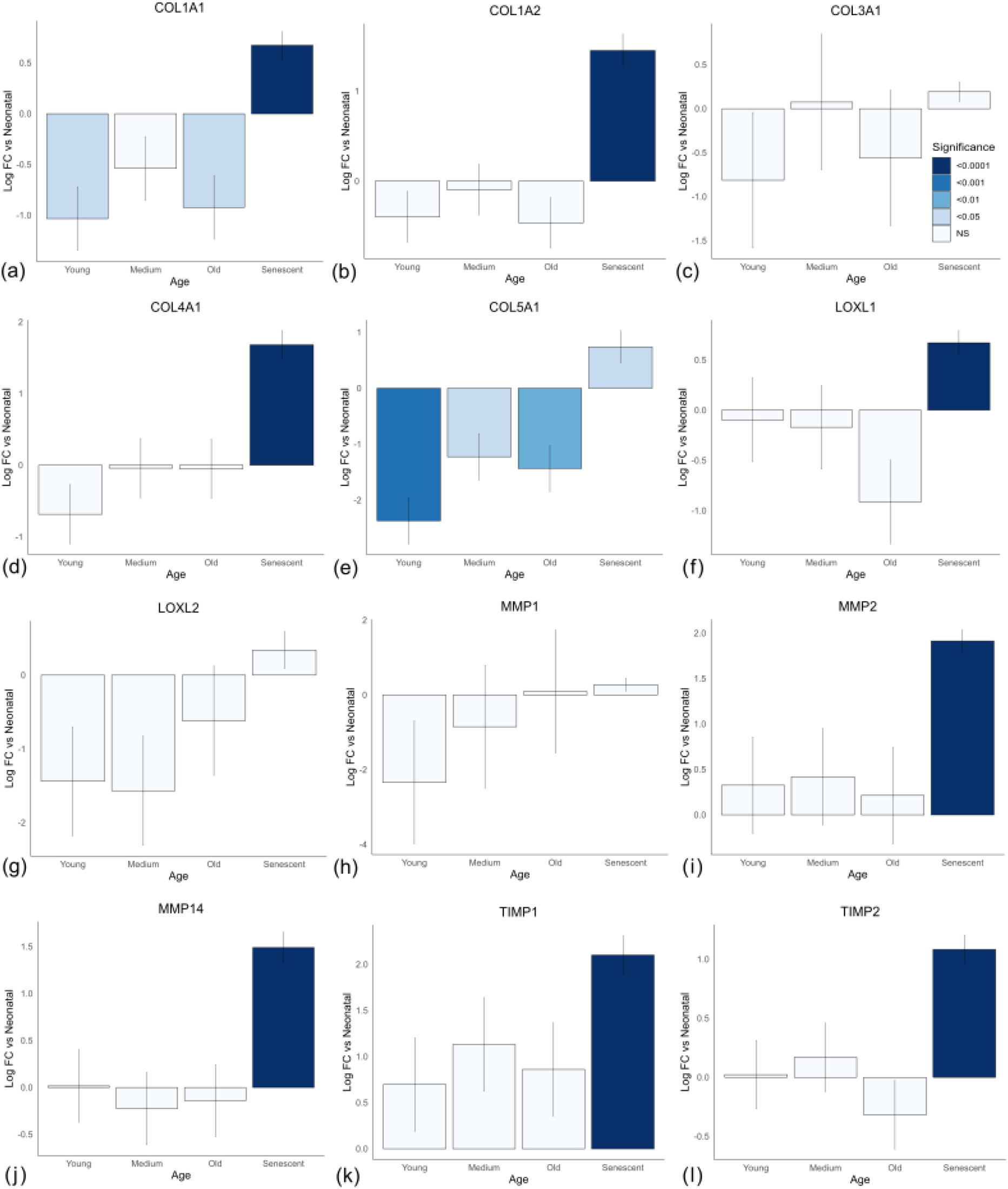
Relative gene expression for cells on 3D Alvetex^®^ matrix from different age donors and senescence. (A-L) Log fold change (Log FC) of gene expression from neonatal reflects the ΔΔCT. Each group reflects the mean of three cell lines. Neonatal cells were from donors <14 days; young cells from donors < 30 years; “medium” reflects donors 40-45 years, and “old” donors were 60+ years. Senescence was induced before seeding on 3D matrix in the neonatal cell lines.

### Ageing cells form fewer myofibroblasts

We next asked why ageing cells might have reduced collagen expression if not resulting from an increase in senescence. We observed that the collagen and LOXL mRNAs which decreased in ageing (but not senescence) were stimulated by TGF-β, whereas the MMP and TIMP genes which were mainly affected by senescence (but not ageing) showed little effect from TGF-β stimulation. We therefore wondered if ageing resulted in some defect in TGF-β signalling that could have a downstream impact on the ECM.

In canonical signalling, TGF-β binds a TGF-β receptor (TGFBR1/2) which triggers Smad2/3 phosphorylation and activation. Smad4 is recruited to the activated Smad2/3 complex and the Smad complex translocates to the nucleus where it binds to various co-factors and transcription factors to induce target gene expression (Zi et al., 2012). One such target gene is Smad7 (Nagarajan et al., 1999) which acts to inhibit canonical TGF-β signalling through dephosphorylation of Smad2/3 (Yan et al., 2016).

Interestingly, TGFBR1 expression showed TGF-β-stimulated increases in all groups but adult cells (Figure 5 A), suggesting that adult cells lack the positive feedback that allows them to better respond to a TGF-β signal. This was unlikely to reflect the amount of senescent cells as both TGFBR1 and TGFBR2 (Supplementary file 4) expression showed significant increase in senescent cells. Smad3 signalling showed a clear (though not-significant) reduction in expression in all cells stimulated by TGF-β compared to unstimulated controls. This decline usually began around 4-8 hours after stimulation and likely corresponded to the increase in Smad7 expression which peaked initially between 2-3 hours in response to TGF-β. However, the Smad7 response was significantly weaker in adult cells compared to all other groups, possibly reflecting a muted TGF-β signal rather than a proportionate reduction in repression, because collagen and other genes presented below suggest adult cells have reduced rather than increased response to TGF-β. Importantly, a muted signal in adult cells is unlikely to reflect a simple reduction in receptor expression as the upregulation of TGFBR1 in all cell lines is a late timepoint event (24hrs+), and is thus likely a consequence rather than a cause of this age-associated effect.

**Figure 5.**
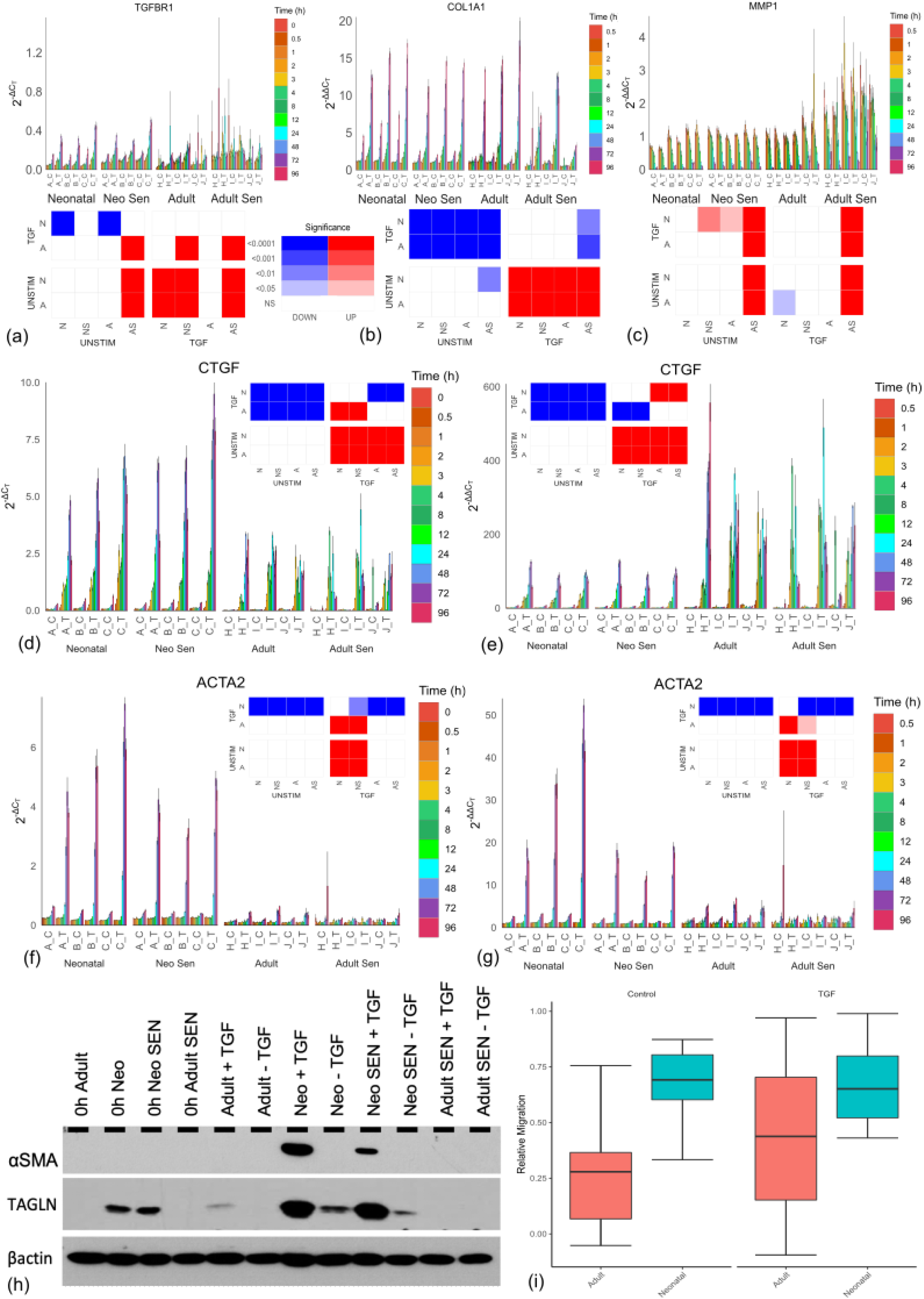
Myofibroblastic activity by cell type. (A-G) Relative mRNA levels for each cell line over the 96-hour timecourse. A, D, and F show 2^−ΔCT^ values comparing gene expression levels between samples. B, C, E, and G show 2^−ΔΔCT^ values comparing level of change of expression from baseline at zero hours for each cell line. The x axes show neonatal cell lines A-C which include non-senescent (Neonatal) and senescent (Neo Sen) cells. Adult cell lines H-J were either non-senescent (Adult) or senescent (Adult Sen). After the letter indicating cell line (A-C and H-J), “_T” indicated stimulation with TGF-β, while “_C” indicated the unstimulated controls. Each graph has an accompanying tile plot indicating whether comparison of the changing profile over time was significantly different between cell types as determined by ANCOVA. The cell types: Adult, A; Adult Senescent, AS; Neonatal, N; and Neonatal Senescent, NS were split by either TGF-β stimulation (TGF) or control (UNSTIM). Colour red indicates that the cell type along the x axis showed a significant increase compared to the cell type and condition depicted on the y axis, while blue reflects a significant decrease along the x axis, and white reflects no significant change (NS). Increasing colour intensity reflects increasing significance. (H) Western blot of myofibroblast markers α-SMA and TAGLN showing protein levels at zero hours and 96 hours with stimulation (+TGF) and without stimulation (−TGF) with TGF-β. (I) Relative migration over 48 hours for neonatal and adult cells stimulated with TGF-β or unstimulated control.

To investigate this possibility further, we normalised the ΔC_T_ values (used hitherto) by further subtracting the baseline value at timepoint zero for each cell line. Thus, instead of 2^−ΔCT^, we used 2^−ΔΔCT^ values indicating the level of change from basal expression for each cell line. Intriguingly, the differences between neonatal and adult cells not only vanished for COL1A1 and COL1A2 (and were greatly reduced for COL5A1), but adult cells showed a trend toward higher increases in mRNAs than neonatal cells (Figure 5 B and Supplementary file 6).

This result was highly unexpected as it suggested that the adult cells could increase collagen production by the same proportion (or more) as neonatal cells. We predicted that defective cells that had accumulated molecular damage would have a proportionately lower response to stimulus, whereas here their response was proportionately (though not significantly) higher. Such an observation is more consistent with a homeostatic change toward reduced TGF-β signalling in adult cells than accumulated damage. Importantly, CTGF, a gene involved in myofibroblast formation (Lipson et al., 2012; Tsai et al., 2018), is significantly reduced in adult cells compared to neonatal cells (as measured by 2^−ΔCT^, Figure 5 D), but the increase in proportion to basal levels (2^−ΔΔCT^) is significantly larger in adult cells compared to neonatal cells (Figure 5 E). While these cells may therefore be perfectly functional in their capacity to respond to TGF-β, they have a reduced basal activity which means even a proportionately larger response still results in a subsequently lower level of total mRNA compared to neonatal cells. And this has important consequences. The most frequent marker of myofibroblast formation, α-SMA, encoded by the ACTA2 gene (Hinz, 2016; Wynn and Ramalingam, 2012), is significantly lower in adult cells compared to neonatal cells (2^−ΔCT^, Figure 5 F), but unlike CTGF the proportionate increase over time is also significantly lower (2^−ΔΔCT^, Figure 5 G). Importantly, western blotting confirmed that differences in mRNA translated into differences in protein levels, including another myofibroblast marker, transgelin (TAGLN) (Assinder et al., 2009; Elsafadi et al., 2016) (Figure 5 H), suggesting that adult cells form fewer myofibroblasts. Potentially, this reflects that formation of myofibroblasts is a threshold event. Unlike CTGF and collagen production which rise by equal proportion in both neonatal and adult cells according to the TGF-β stimulus, because the final amount of CTGF is lower in adult cells, it is less likely to reach the threshold required for myofibroblast differentiation. As myofibroblasts are highly important in wound healing and maintenance of the ECM, this difference, which may reflect no damage or incapacity, but a simple homeostatic change, may have highly detrimental consequences to continued skin health. To test if this also translated to functional differences, we used a migration assay, which showed a decline in the level of migration of adult cell lines compared to neonatal cell lines (Figure 5 I).

Similar to prior observations, the reduced expression of α-SMA in adult cells is unlikely to reflect an increase in senescent cells, as neonatal senescent cells still have robust α-SMA expression at the mRNA and protein levels in response to TGF-β (Figure 5 D and F). We considered one final possible way that senescent cells could be responsible for the age-related changes we see here. We and others have demonstrated a senescent cell bystander effect (Acosta et al., 2013; Nelson et al., 2012) by which the associated secretory phenotype (SASP) and juxtacrine signalling (Hoare et al., 2016) provoke changes in the surrounding tissue. The reason cultures of purely senescent cells would not show the same decline in TGF-β response may therefore reflect that there are no remaining non-senescent cells to undergo the bystander effect. Further research should assess whether the bystander effect could feasibly explain the changes in TGF-β signalling we see with age; however, these data robustly show that changes in collagen production and myofibroblast formation are not directly the result of older cells more frequently becoming senescent.

## Discussion

Here we demonstrate that changes in aged cells (55+ year old donors) for collagen mRNAs and proteins are dramatically reduced compared to neonatal cells. Senescent cells from the neonatal cell lines also showed a reduction in collagen production, but not to the level of adult cells, suggesting that senescence could not be responsible for the lower drop seen in adult cells.

As summarised by Wlaschek et al. (2021), “an emerging hypothesis of skin aging postulates that mainly fibroblast senescence drives skin decline and skin aging due to irreversible proliferation arrest and enhanced release of a senescence-associated secretory phenotype (SASP). SASP, through chemokines and proinflammatory factors, induces chronic inflammation, reduces proliferation by impaired release of essential GFs [growth factors], and enhances the degradation of the ECM (connective tissue) by enhanced activation of proteolytic enzymes, including matrix-degrading metalloproteinases.” Our data are consistent with senescent cells increasing matrix degradation, but increased degradation only threatens homeostasis if it is not accompanied by increased deposition. However, our data suggest that another process independent of senescence appears to be preventing this from occurring. Our data from normalisation of ΔCT by value at time zero suggests that adult cells are still capable of similar scale increases in production of mRNA as neonatal cells; but from a smaller starting value the end result is still lower. This may reflect that similar fold change is easier from lower starting values, and thus ageing cells might still have reduced capacity, but another possibility is that homeostasis has shifted toward reduced ECM production and slower metabolism, consistent with a different model of ageing we have recently described as selective destruction theory (SDT) (Wordsworth et al., 2022). SDT predicts that many cells will undergo a reduction in metabolism that may manifest in lower basal rates of anabolic activity such as ECM production. However, whether this should result in reduced levels after stimulation will depend on the regulating protein network, and requires further modelling. Plausibly, systems with threshold effects as we propose for myofibroblast formation could simply lower the threshold to meet the new level of homeostasis. What is clear from these data is that the reduced fibrotic nature of adult cells does not directly reflect an increase in senescent cells, so another process, be it damage accumulation or selective destruction must be occurring. However, there are limitations to the conclusions drawn here. We demonstrated that adult fibroblasts are not as migratory as neonatal fibroblasts, but this was not affected by TGF-β and thus may reflect different underlying processes. We also did not control for sex differences in the fibroblast cell lines, and because the young cells are neonatal, it is plausible that the differences reflect developmental processes rather than ageing *per se*. Further work should address whether similar changes in homeostasis continue after development ceases; however, the reduction of collagen levels and increase in MMP levels are well established markers of ageing (Levi et al., 2020; Reilly and Lozano, 2021; Varani et al., 2006). These data are therefore good evidence that while senescent cells may play a role in tissue ageing and ECM degradation, cells are also undergoing an ageing process that is independent of their replicative lifespan.

## Author Contributions

Nicola Fullard and Victoria Maltman carried out the lab experiments with the exception of the 3D dermal equivalent work carried out by Lydia Costello; James Wordsworth and Ciaran Welsh analysed the data; Charlie Bascom, Ryan Tasseff, and Bob Isfort designed and carried out the qPCR smart chip experiments and contributed to analysis; Rebekah Scanlan contributed to figure production and edited the paper; the study was designed, developed, and supervised by Daryl Shanley and Stefan Przyborski, with input on study direction from James Wordsworth who also wrote the paper.

## Acknowledgements

We would like to thank NC3Rs, Novo Nordisk Fonden (NNF) and Procter & Gamble for funding the work. Grant numbers: NC3Rs, (NC/S001050/1); Novo Nordisk Fonden Denmark, (NNF17OC0027812).

