## Supplementary file 2 for "Cell Senescence-Independent Ageing of Human Skin"

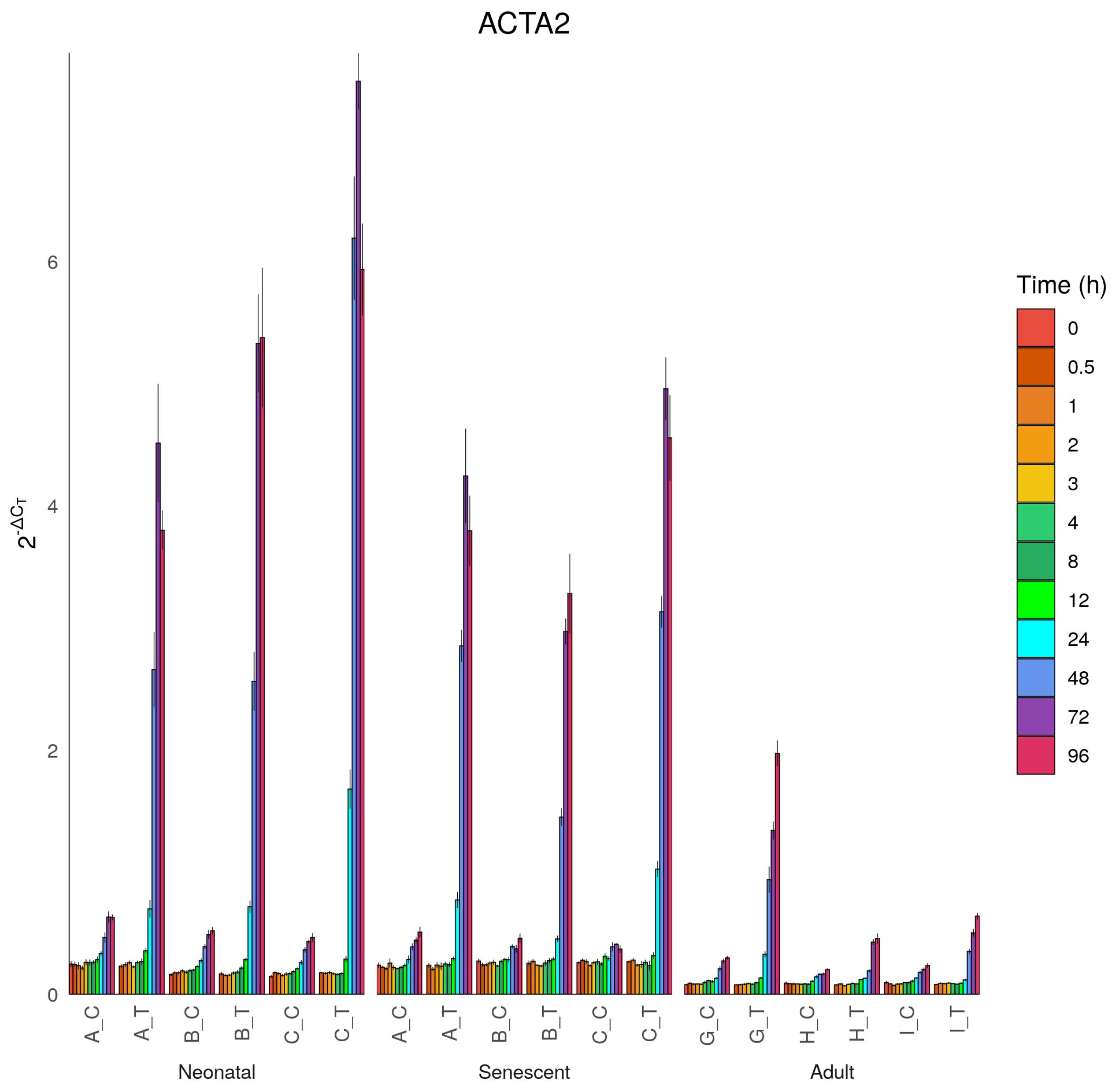

ACTA2
$2^{-\Delta C_T}$
0
2
4
6
A_C
A_T
B_C
B_T
C_C
C_T
A_C
A_T
B_C
B_T
C_C
C_T
G_C
G_T
H_C
H_T
I_C
I_T
Neonatal
Senescent
Adult
Time (h)
0
0.5
1
2
3
4
8
12
24
48
72
96

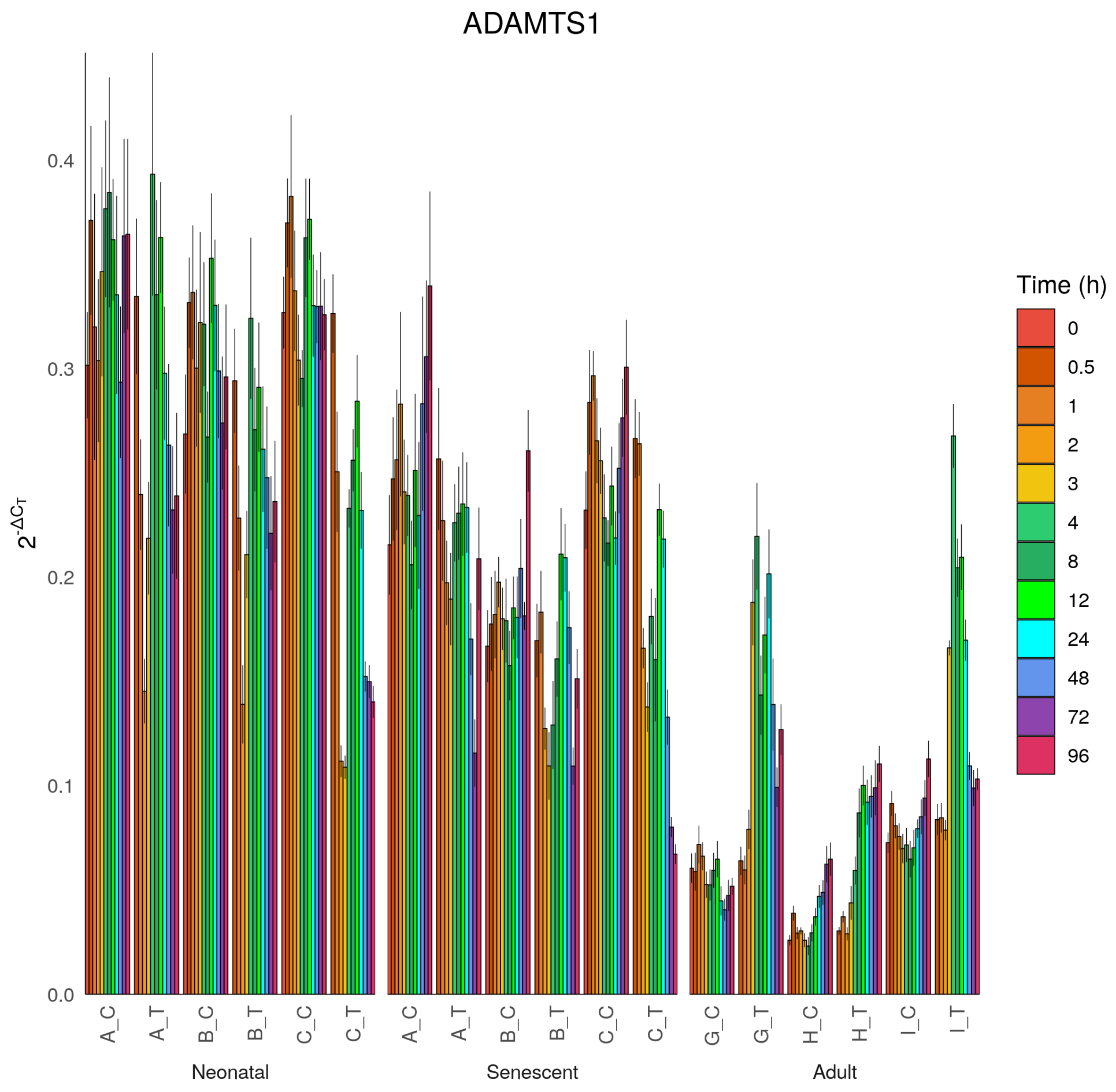

ADAMTS1
$2^{-\Delta C_T}$
0.0
0.1
0.2
0.3
0.4
A_C
A_T
B_C
B_T
C_C
C_T
G_C
G_T
H_C
H_T
I_C
I_T
Neonatal
Senescent
Adult
Time (h)
0
0.5
1
2
3
4
8
12
24
48
72
96

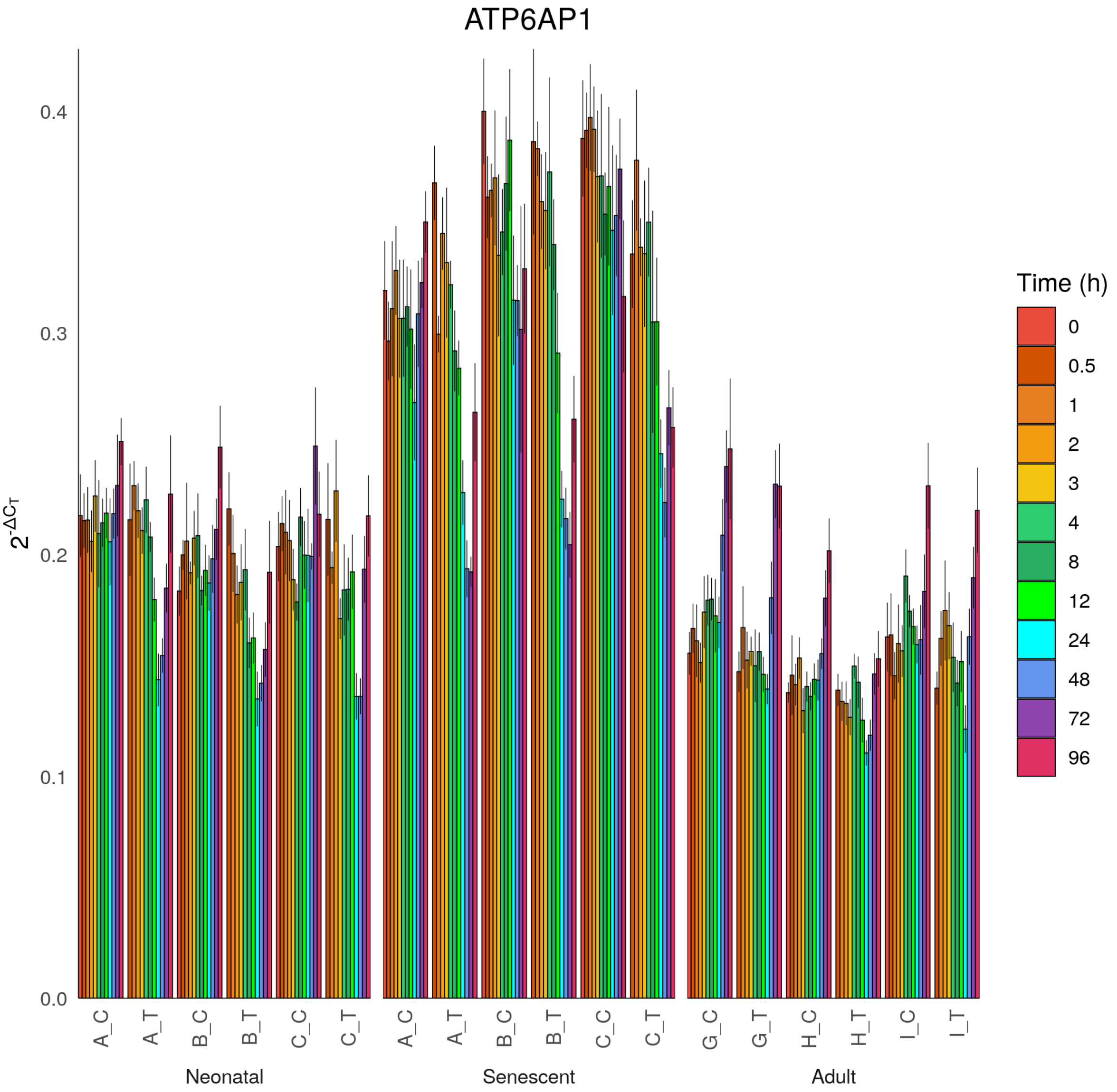

ATP6AP1
$2^{-\Delta C_T}$
0.0
0.1
0.2
0.3
0.4
A_C
A_T
B_C
B_T
C_C
C_T
G_C
G_T
H_C
H_T
I_C
I_T
Neonatal
Senescent
Adult
Time (h)
0
0.5
1
2
3
4
8
12
24
48
72
96

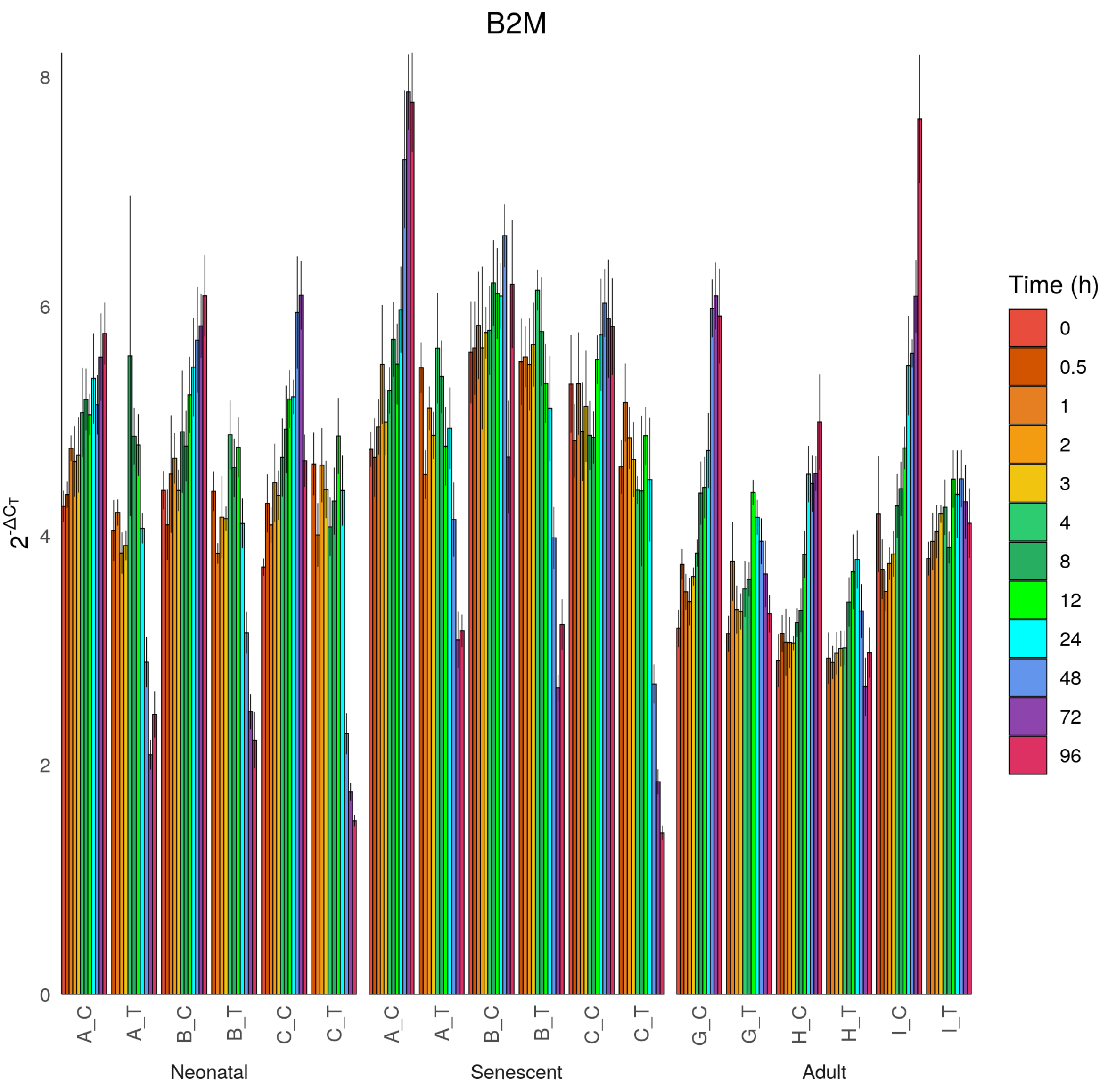

B2M
$2^{-\Delta C_T}$
0
2
4
6
8
A_C
A_T
B_C
B_T
C_C
C_T
Neonatal
A_C
A_T
B_C
B_T
C_C
C_T
Senescent
G_C
G_T
H_C
H_T
I_C
I_T
Adult
Time (h)
0
0.5
1
2
3
4
8
12
24
48
72
96

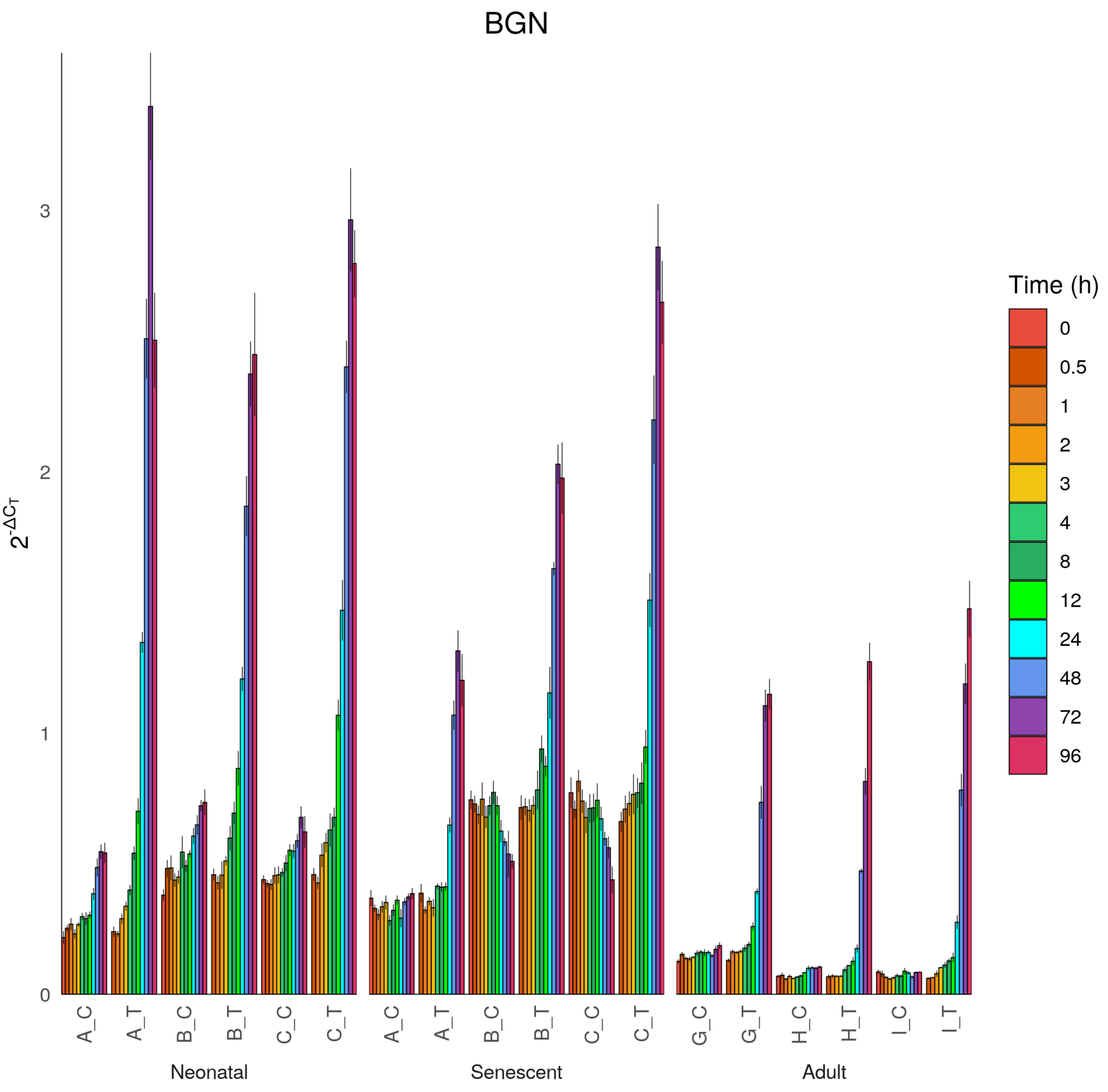

BGN
$2^{-\Delta C_T}$
0
1
2
3
A_C
A_T
B_C
B_T
C_C
C_T
G_C
G_T
H_C
H_T
I_C
I_T
Neonatal
Senescent
Adult
Time (h)
0
0.5
1
2
3
4
8
12
24
48
72
96

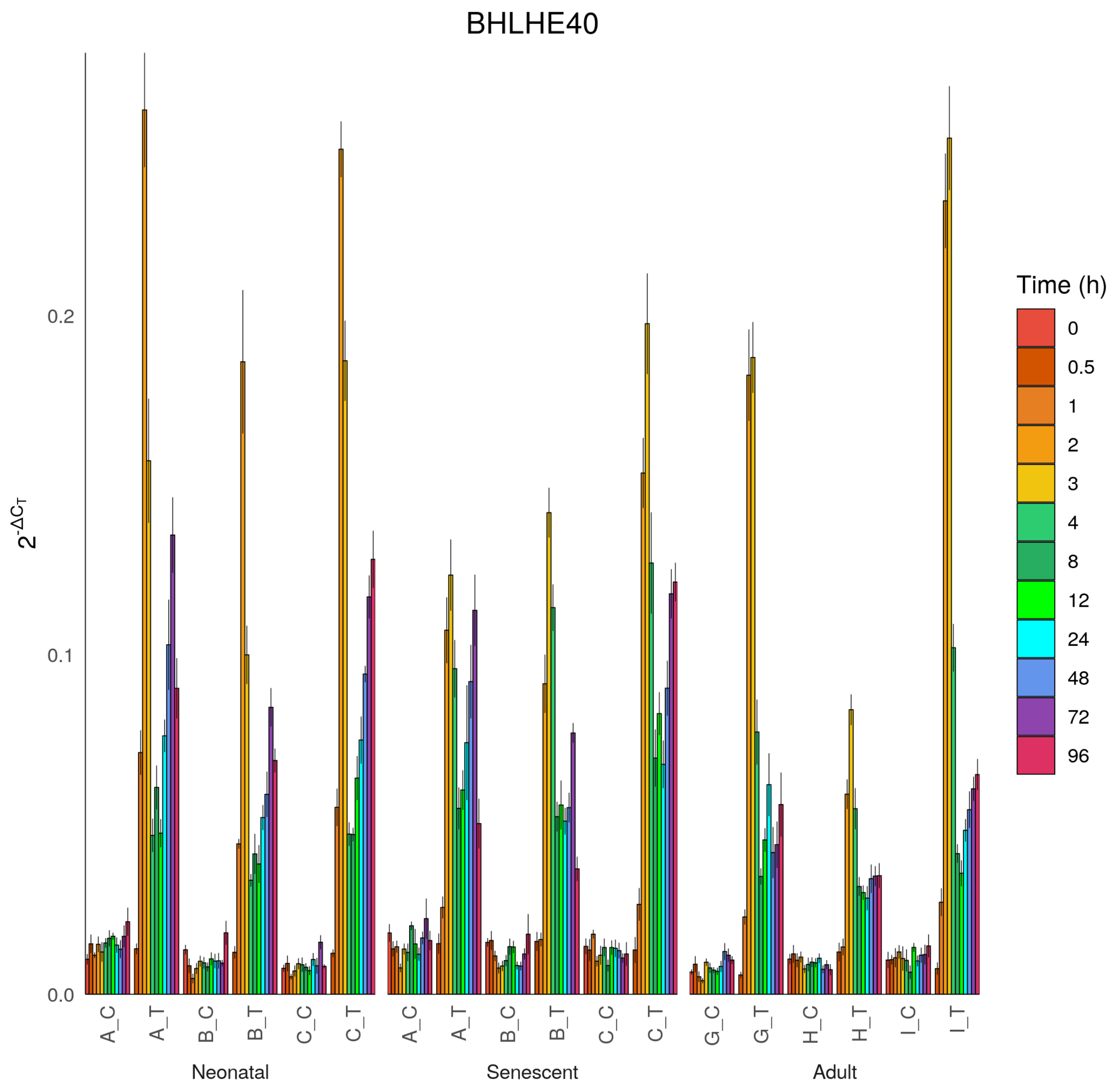
BHLHE40
2^-ΔCT
0.0
0.1
0.2
A_C
A_T
B_C
B_T
C_C
C_T
A_C
A_T
B_C
B_T
C_C
C_T
G_C
G_T
H_C
H_T
I_C
I_T
Neonatal
Senescent
Adult
Time (h)
0
0.5
1
2
3
4
8
12
24
48
72
96

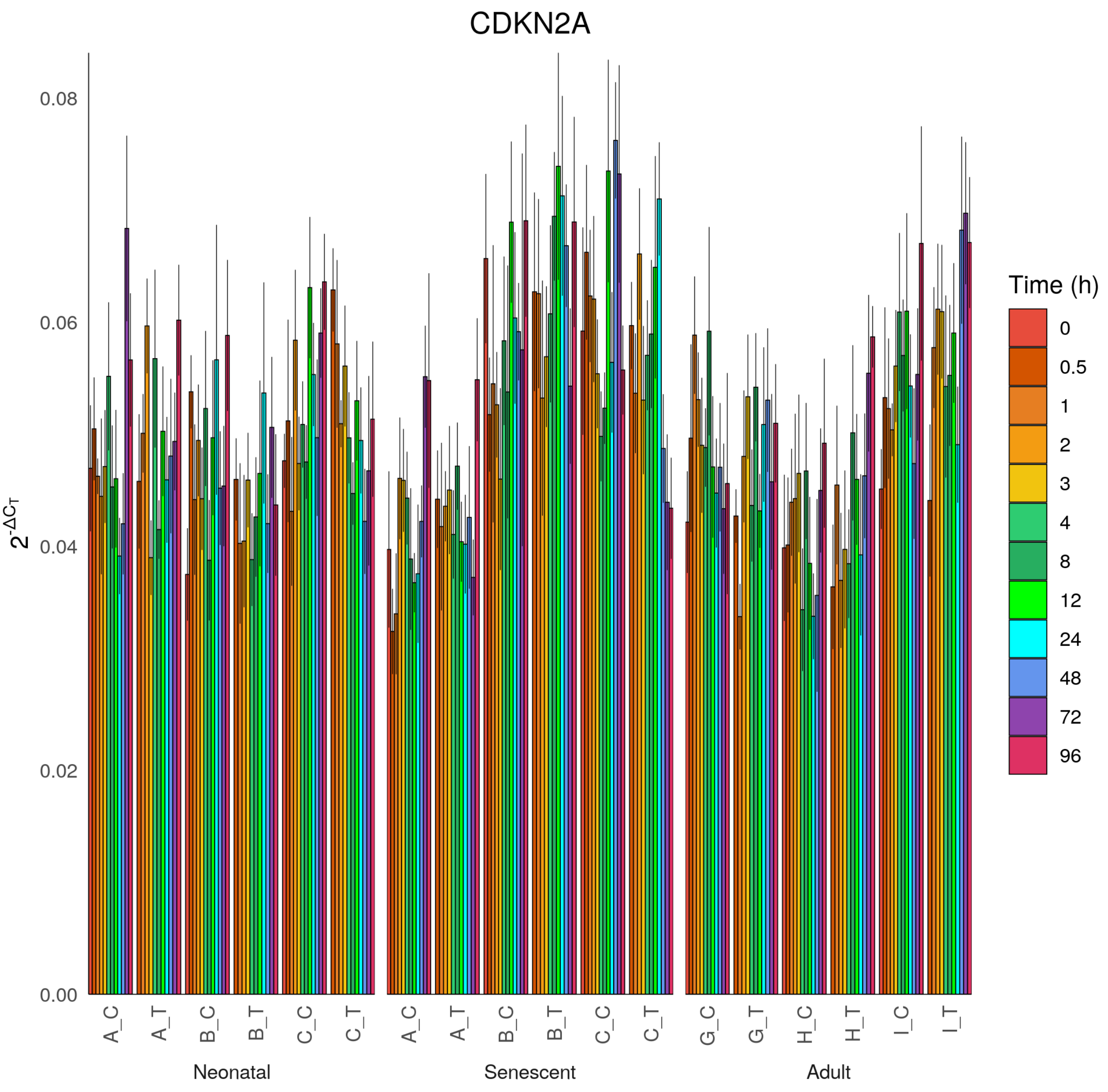

CDKN2A
0.08
0.06
0.04
0.02
0.00
$2^{-\Delta C_T}$
A_C
A_T
B_C
B_T
C_C
C_T
G_C
G_T
H_C
H_T
I_C
I_T
Neonatal
Senescent
Adult
Time (h)
0
0.5
1
2
3
4
8
12
24
48
72
96

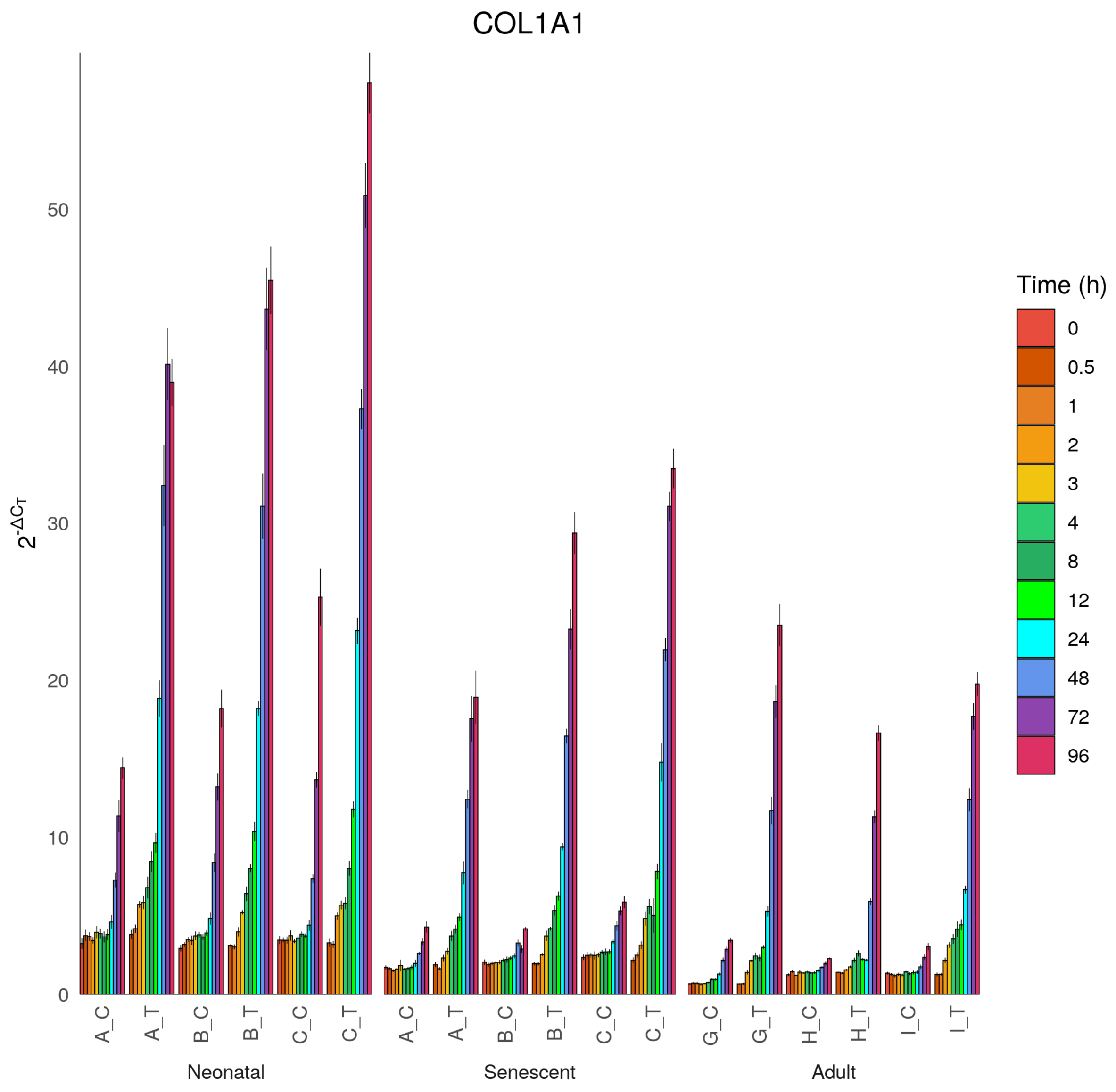

COL1A1
$2^{-\Delta C_T}$
0
10
20
30
40
50
A_C
A_T
B_C
B_T
C_C
C_T
A_C
A_T
B_C
B_T
C_C
C_T
G_C
G_T
H_C
H_T
I_C
I_T
Neonatal
Senescent
Adult
Time (h)
0
0.5
1
2
3
4
8
12
24
48
72
96

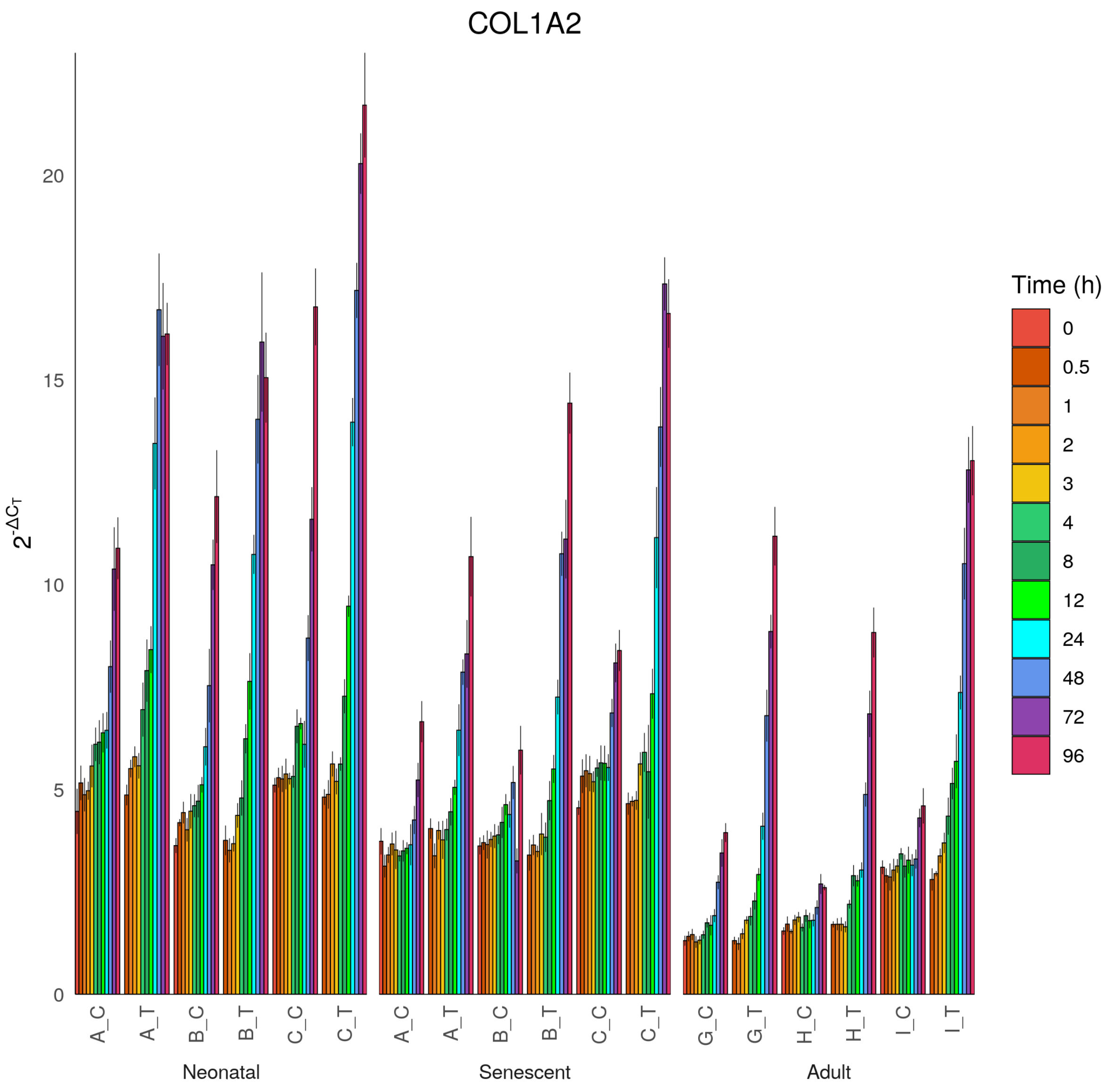
COL1A2
$2^{-\Delta C_T}$
0
5
10
15
20
A_C
A_T
B_C
B_T
C_C
C_T
A_C
A_T
B_C
B_T
C_C
C_T
G_C
G_T
H_C
H_T
I_C
I_T
Neonatal
Senescent
Adult
Time (h)
0
0.5
1
2
3
4
8
12
24
48
72
96

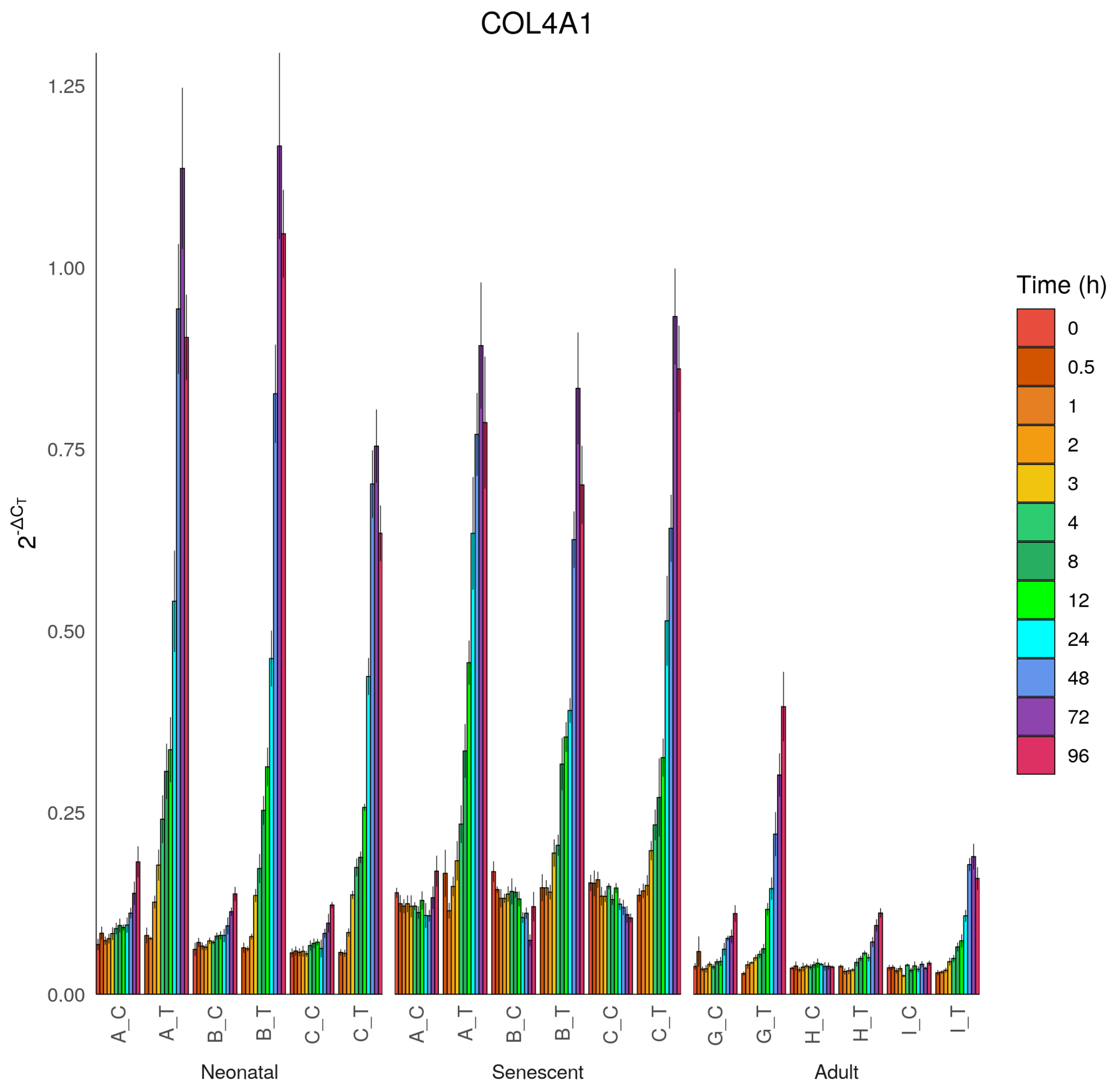

COL4A1
1.25
1.00
0.75
0.50
0.25
0.00
2^-ΔCT
A_C
A_T
B_C
B_T
C_C
C_T
Neonatal
Senescent
G_C
G_T
H_C
H_T
I_C
I_T
Adult
Time (h)
0
0.5
1
2
3
4
8
12
24
48
72
96

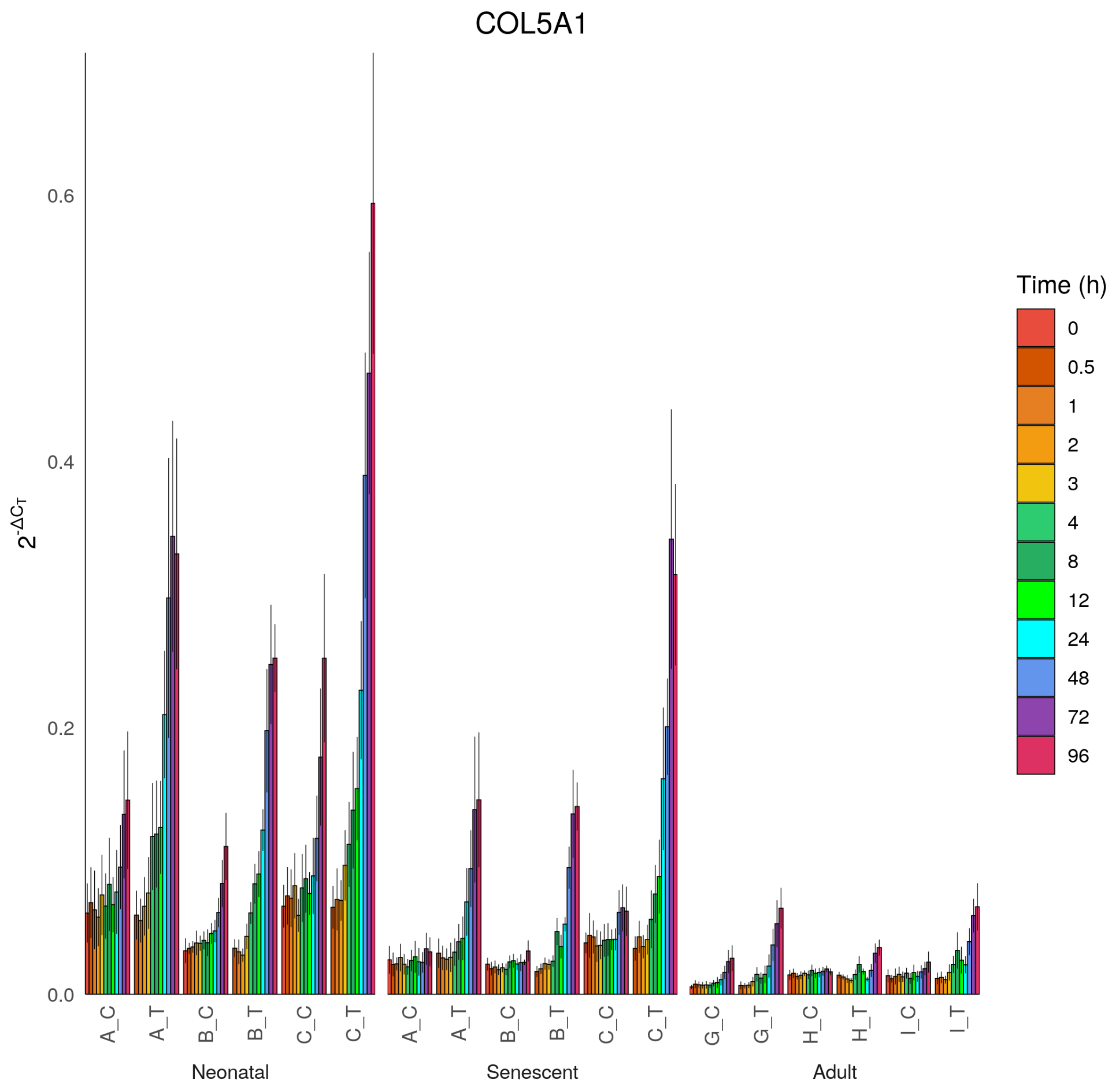

COL5A1
$2^{-\Delta C_T}$
0.0
0.2
0.4
0.6
A_C
A_T
B_C
B_T
C_C
C_T
G_C
G_T
H_C
H_T
I_C
I_T
Neonatal
Senescent
Adult
Time (h)
0
0.5
1
2
3
4
8
12
24
48
72
96

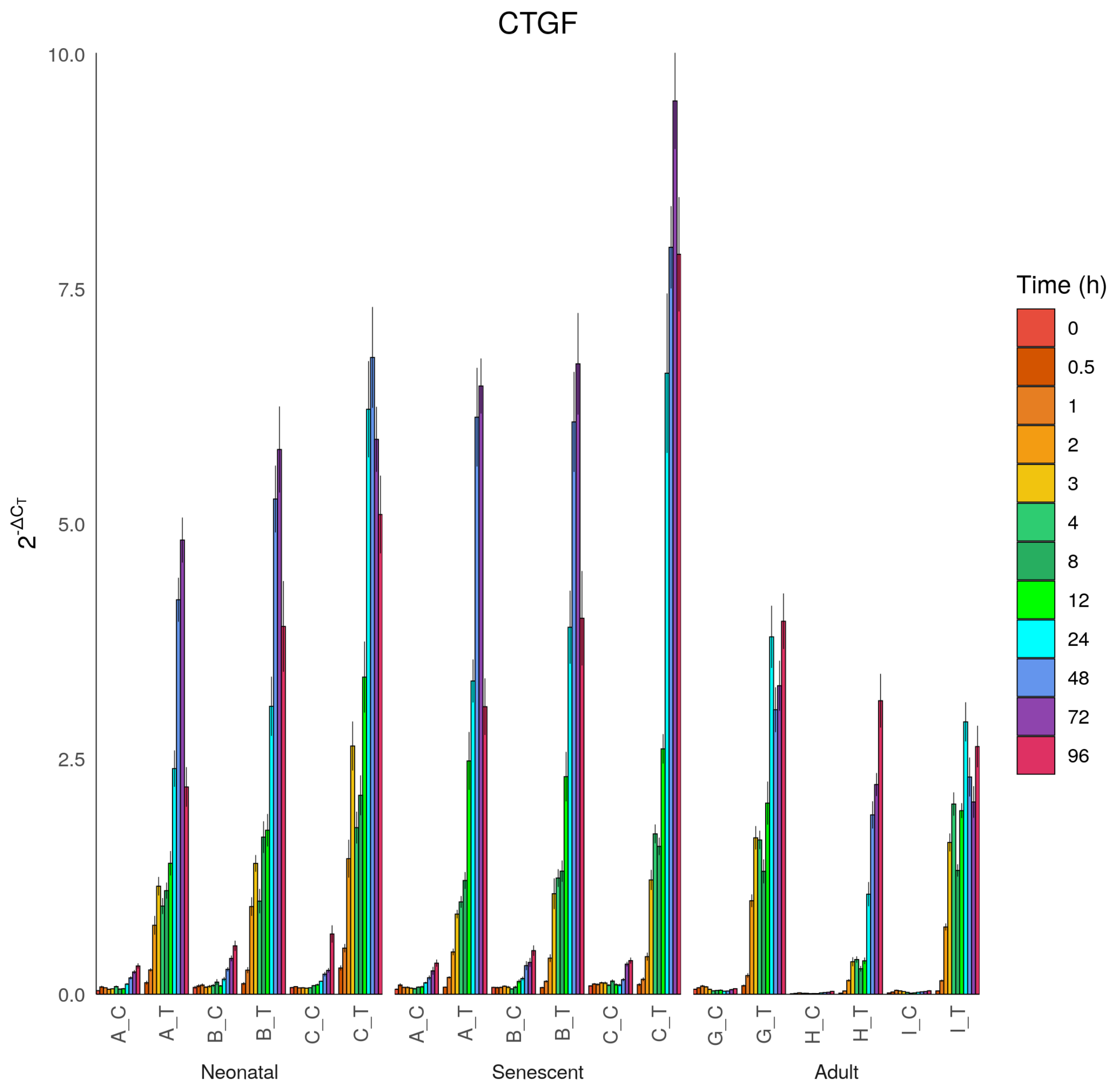

CTGF
10.0
7.5
5.0
2.5
0.0
$2^{-\Delta C_T}$
A_C
A_T
B_C
B_T
C_C
C_T
A_C
A_T
B_C
B_T
C_C
C_T
G_C
G_T
H_C
H_T
I_C
I_T
Neonatal
Senescent
Adult
Time (h)
0
0.5
1
2
3
4
8
12
24
48
72
96

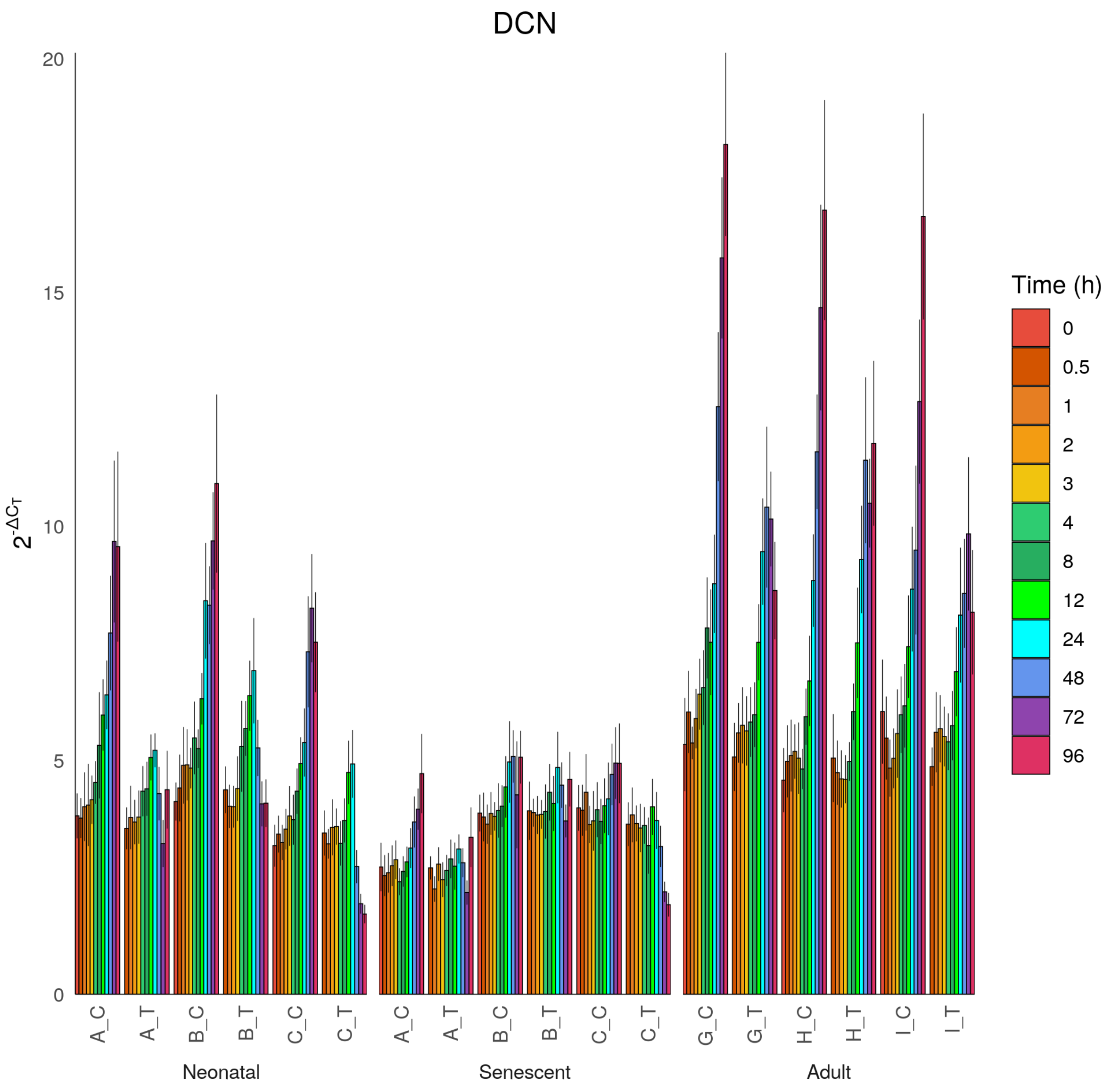

DCN
$2^{-\Delta C_T}$
0
5
10
15
20
A_C
A_T
B_C
B_T
C_C
C_T
Neonatal
Senescent
G_C
G_T
H_C
H_T
I_C
I_T
Adult
Time (h)
0
0.5
1
2
3
4
8
12
24
48
72
96

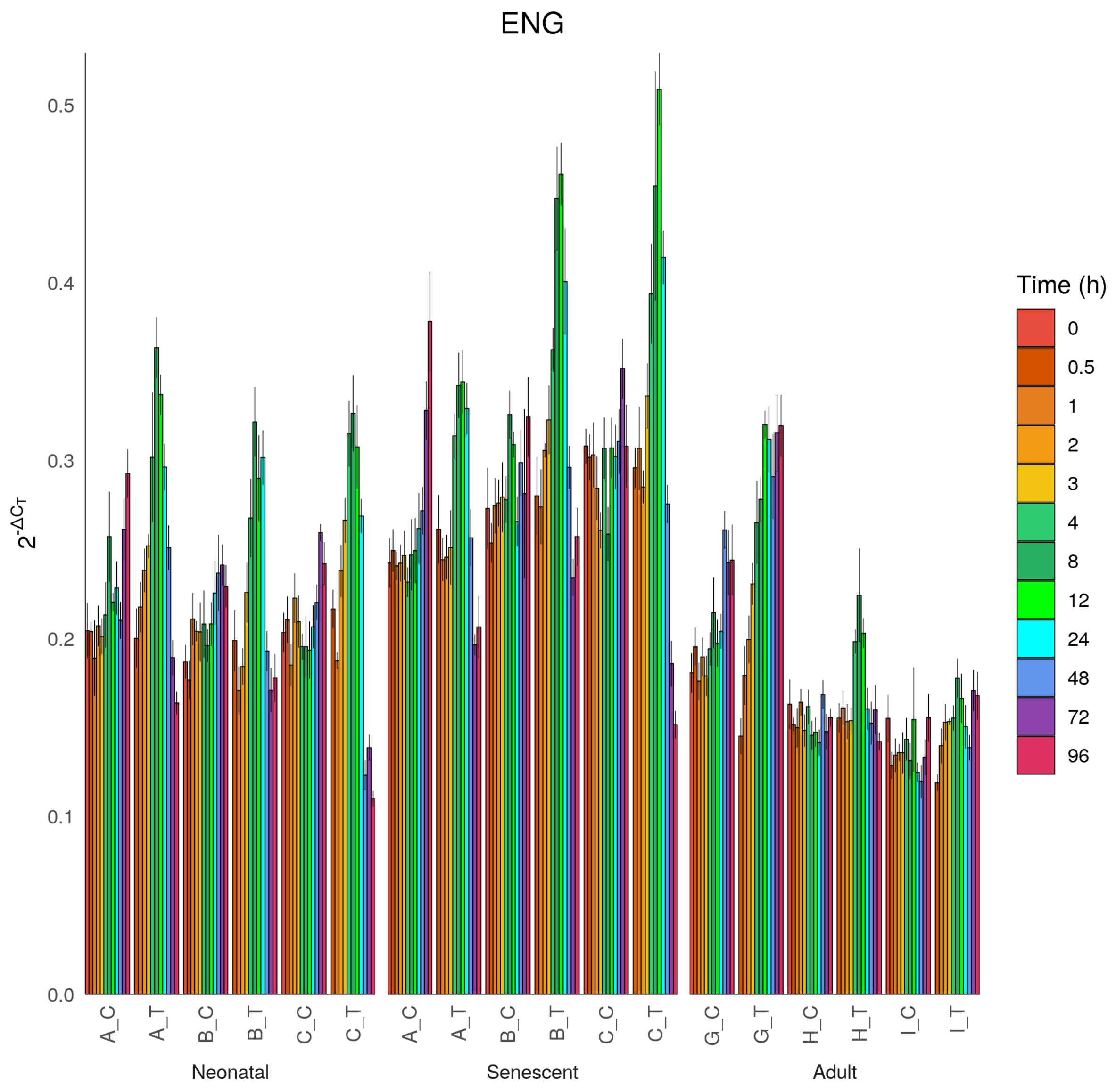
ENG
2^-ΔCT
0.0
0.1
0.2
0.3
0.4
0.5
A_C
A_T
B_C
B_T
C_C
C_T
A_C
A_T
B_C
B_T
C_C
C_T
G_C
G_T
H_C
H_T
I_C
I_T
Neonatal
Senescent
Adult
Time (h)
0
0.5
1
2
3
4
8
12
24
48
72
96

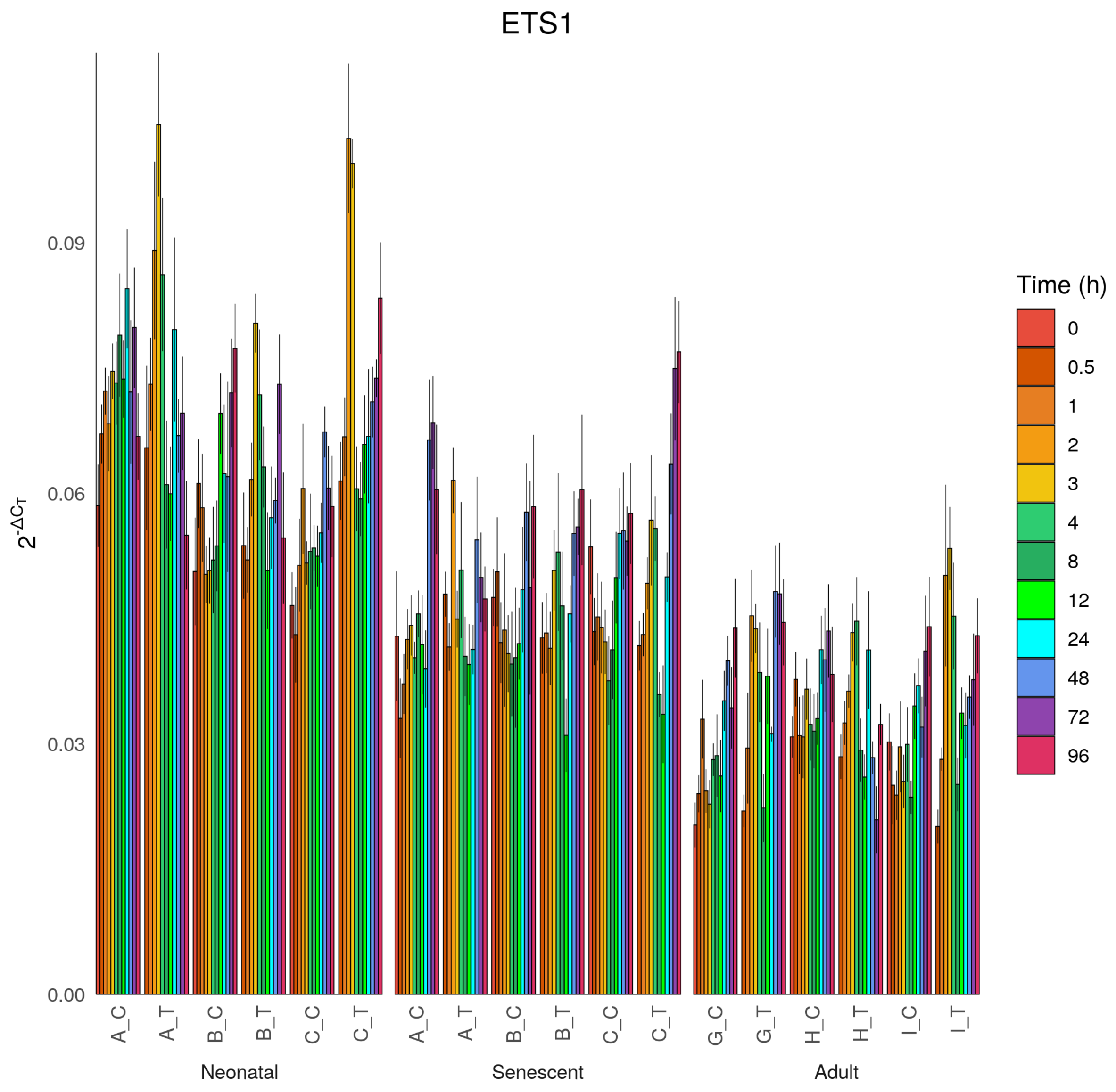

ETS1
$2^{-\Delta C_T}$
0.00
0.03
0.06
0.09
A_C
A_T
B_C
B_T
C_C
C_T
Neonatal
A_C
A_T
B_C
B_T
C_C
C_T
Senescent
G_C
G_T
H_C
H_T
I_C
I_T
Adult
Time (h)
0
0.5
1
2
3
4
8
12
24
48
72
96

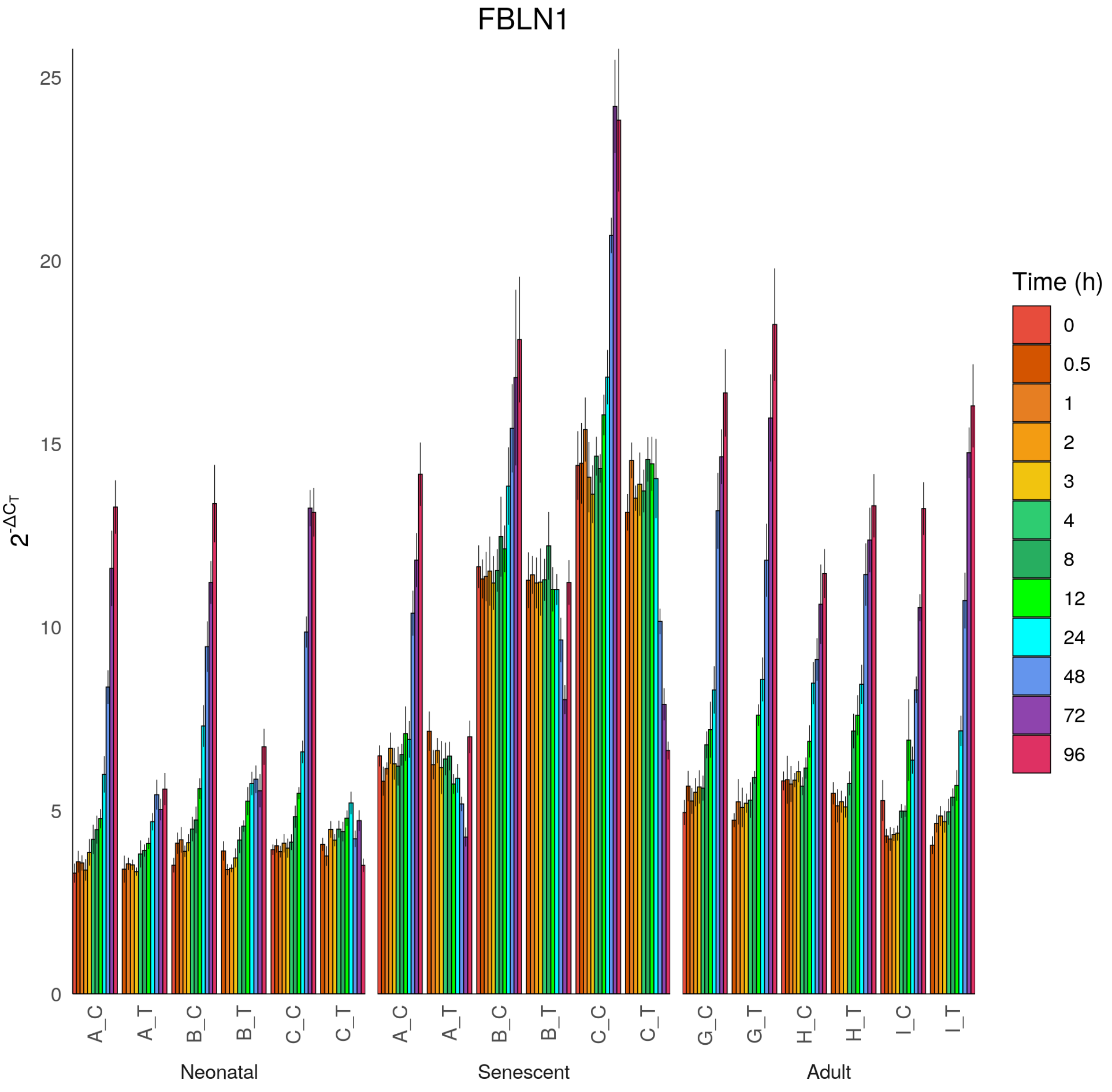
FBLN1
Time (h)
0
0.5
1
2
3
4
8
12
24
48
72
96
$2^{-\Delta C_T}$
0
5
10
15
20
25
A_C
A_T
B_C
B_T
C_C
C_T
G_C
G_T
H_C
H_T
I_C
I_T
Neonatal
Senescent
Adult

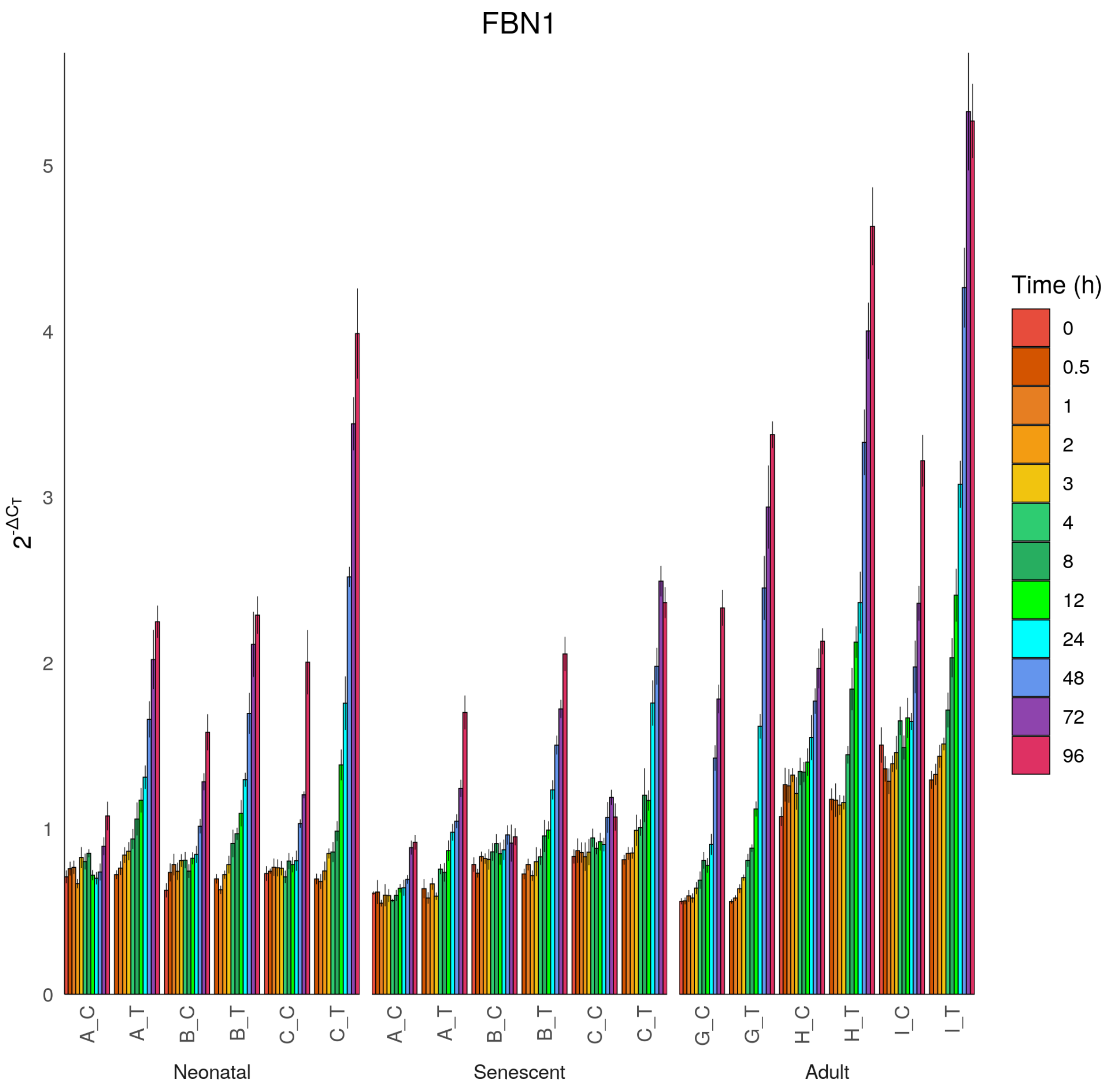
FBN1
$2^{-\Delta C_T}$
0
1
2
3
4
5
A_C
A_T
B_C
B_T
C_C
C_T
Neonatal
A_C
A_T
B_C
B_T
C_C
C_T
Senescent
G_C
G_T
H_C
H_T
I_C
I_T
Adult
Time (h)
0
0.5
1
2
3
4
8
12
24
48
72
96

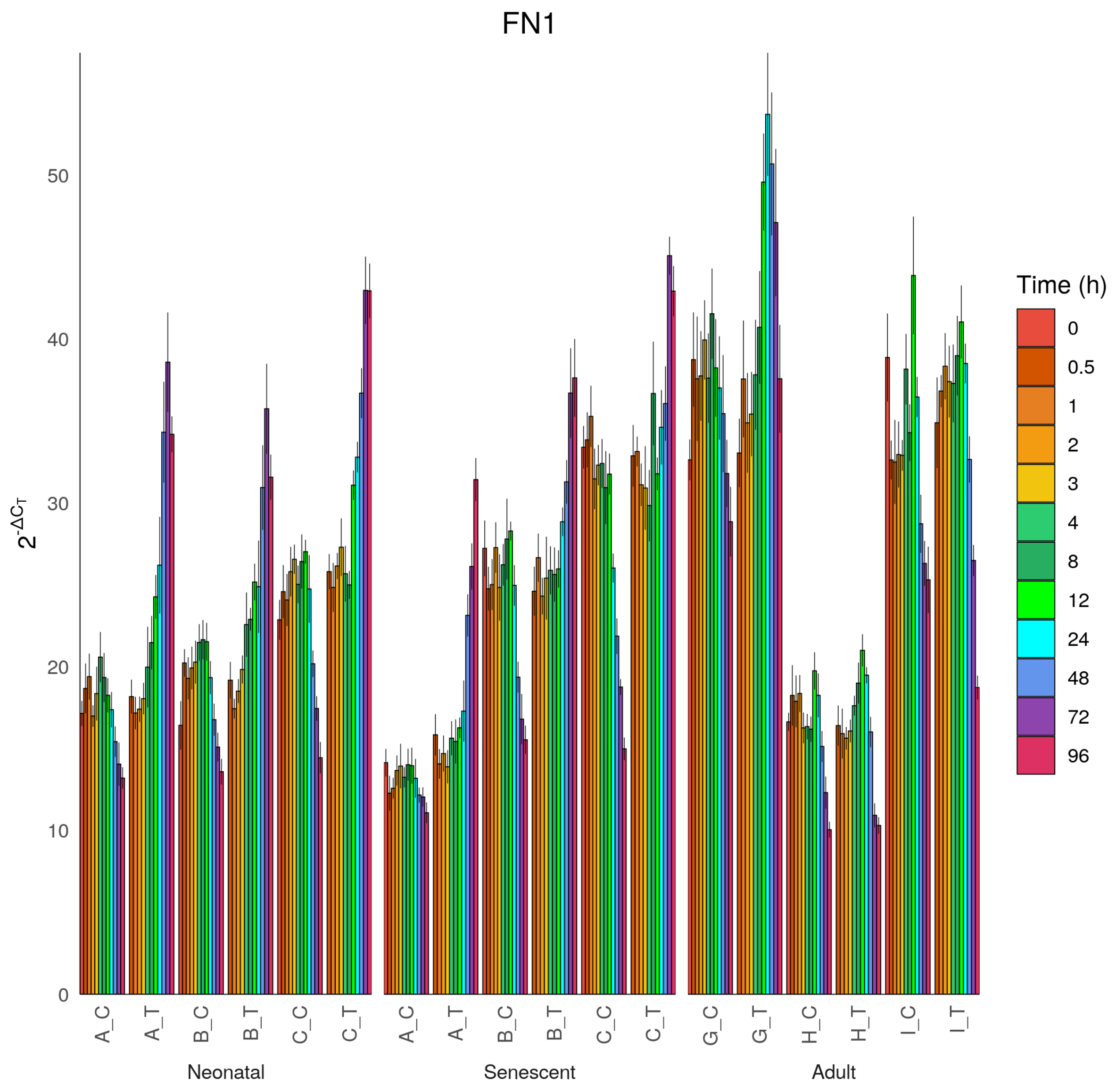

FN1
$2^{-\Delta C_T}$
0
10
20
30
40
50
A_C
A_T
B_C
B_T
C_C
C_T
A_C
A_T
B_C
B_T
C_C
C_T
G_C
G_T
H_C
H_T
I_C
I_T
Neonatal
Senescent
Adult
Time (h)
0
0.5
1
2
3
4
8
12
24
48
72
96

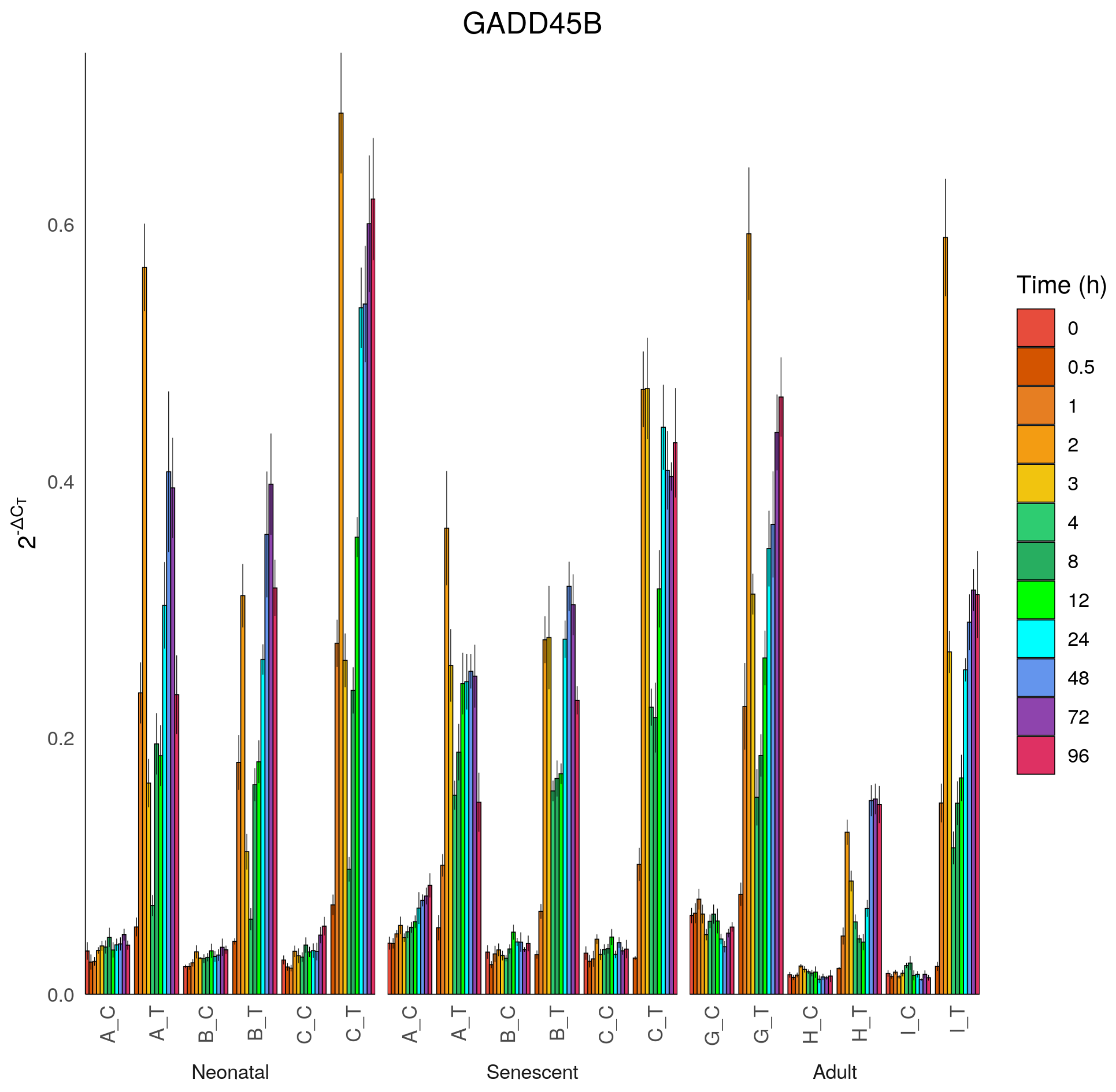
GADD45B
$2^{-\Delta C_T}$
0.0
0.2
0.4
0.6
A_C
A_T
B_C
B_T
C_C
C_T
Neonatal
A_C
A_T
B_C
B_T
C_C
C_T
Senescent
G_C
G_T
H_C
H_T
I_C
I_T
Adult
Time (h)
0
0.5
1
2
3
4
8
12
24
48
72
96

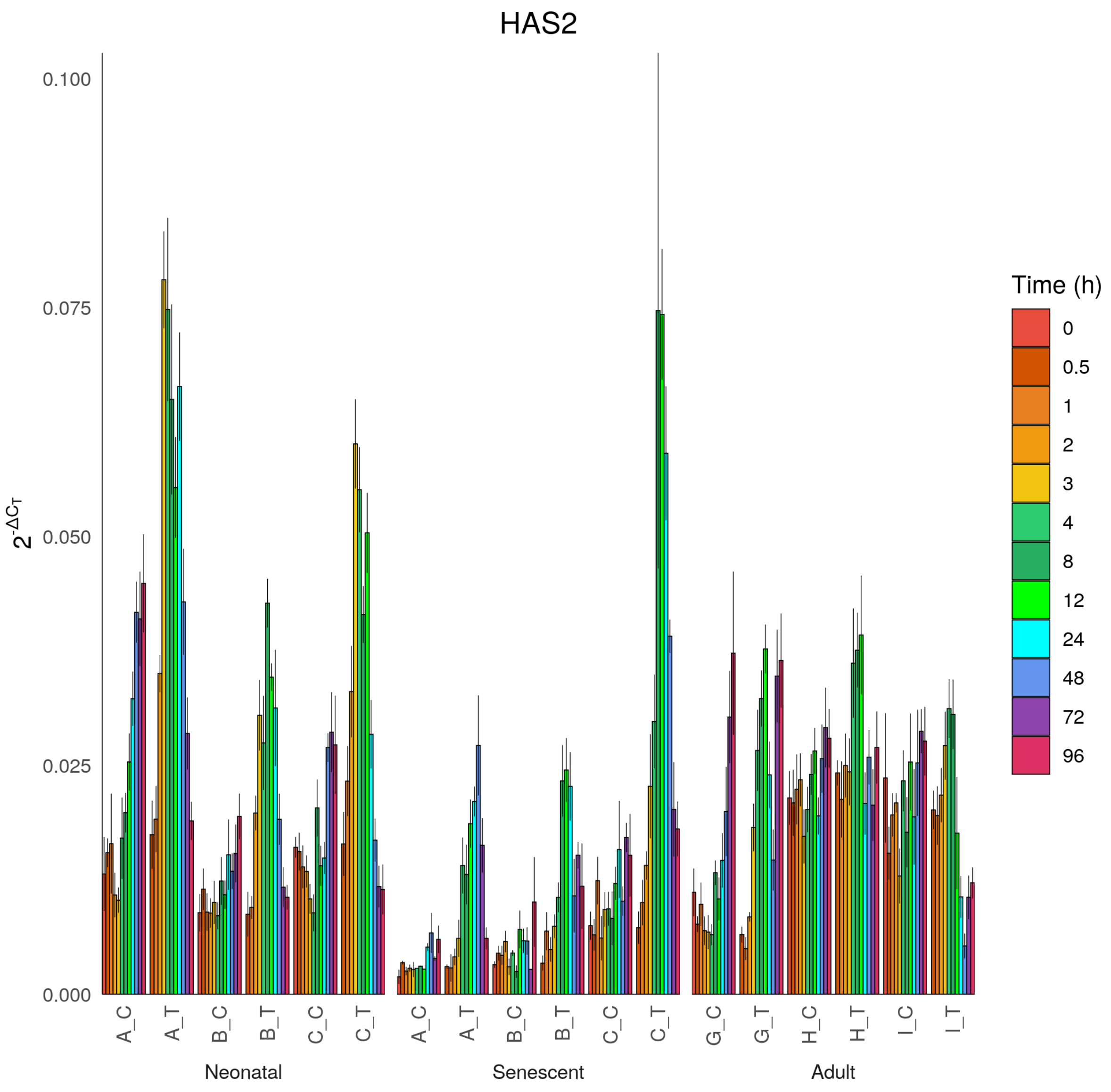
HAS2
$2^{-\Delta C_T}$
0.100
0.075
0.050
0.025
0.000
A_C
A_T
B_C
B_T
C_C
C_T
A_C
A_T
B_C
B_T
C_C
C_T
G_C
G_T
H_C
H_T
I_C
I_T
Neonatal
Senescent
Adult
Time (h)
0
0.5
1
2
3
4
8
12
24
48
72
96

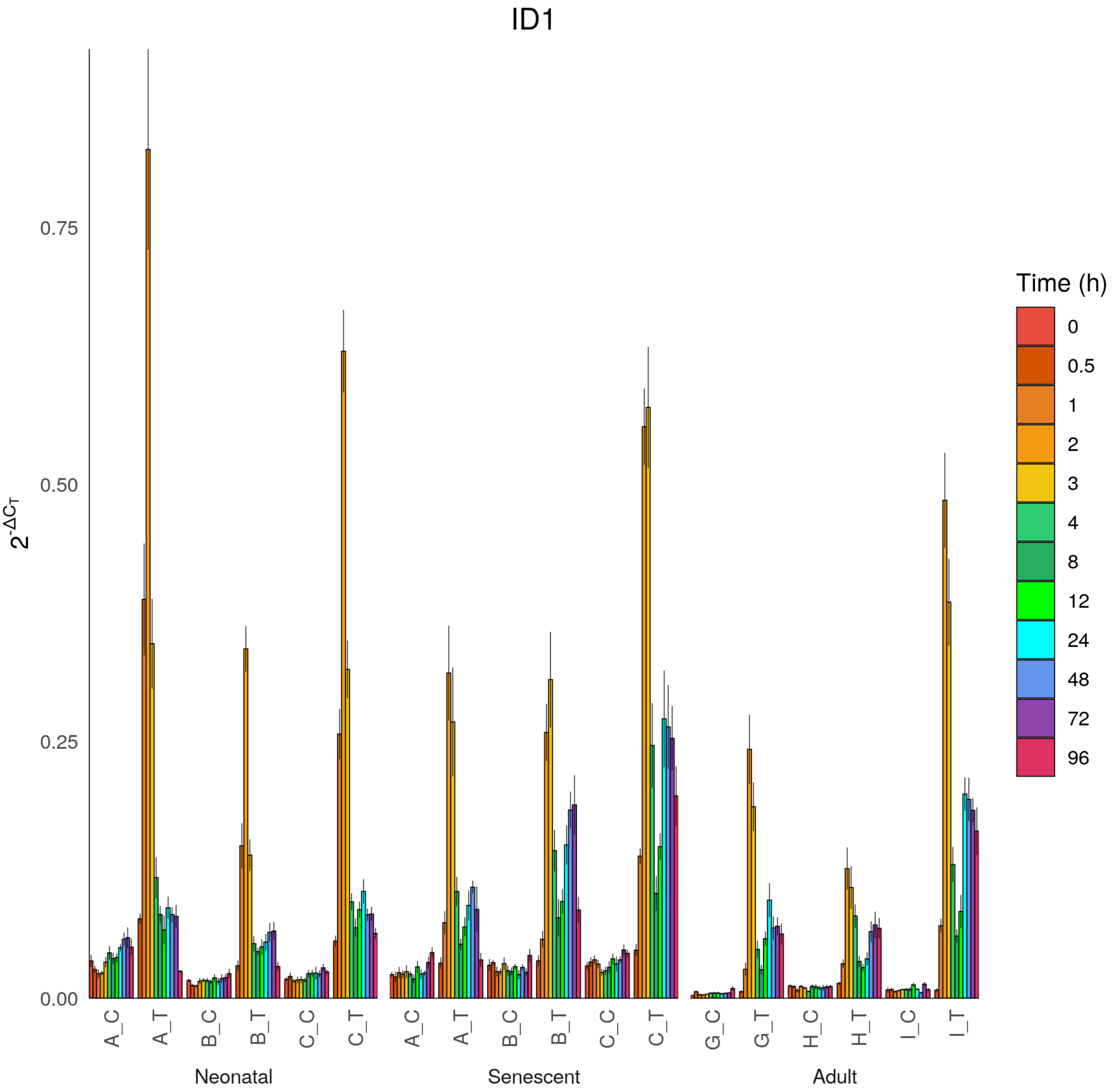

ID1
$2^{-\Delta C_T}$
0.00
0.25
0.50
0.75
A_C
A_T
B_C
B_T
C_C
C_T
A_C
A_T
B_C
B_T
C_C
C_T
G_C
G_T
H_C
H_T
I_C
I_T
Neonatal
Senescent
Adult
Time (h)
0
0.5
1
2
3
4
8
12
24
48
72
96

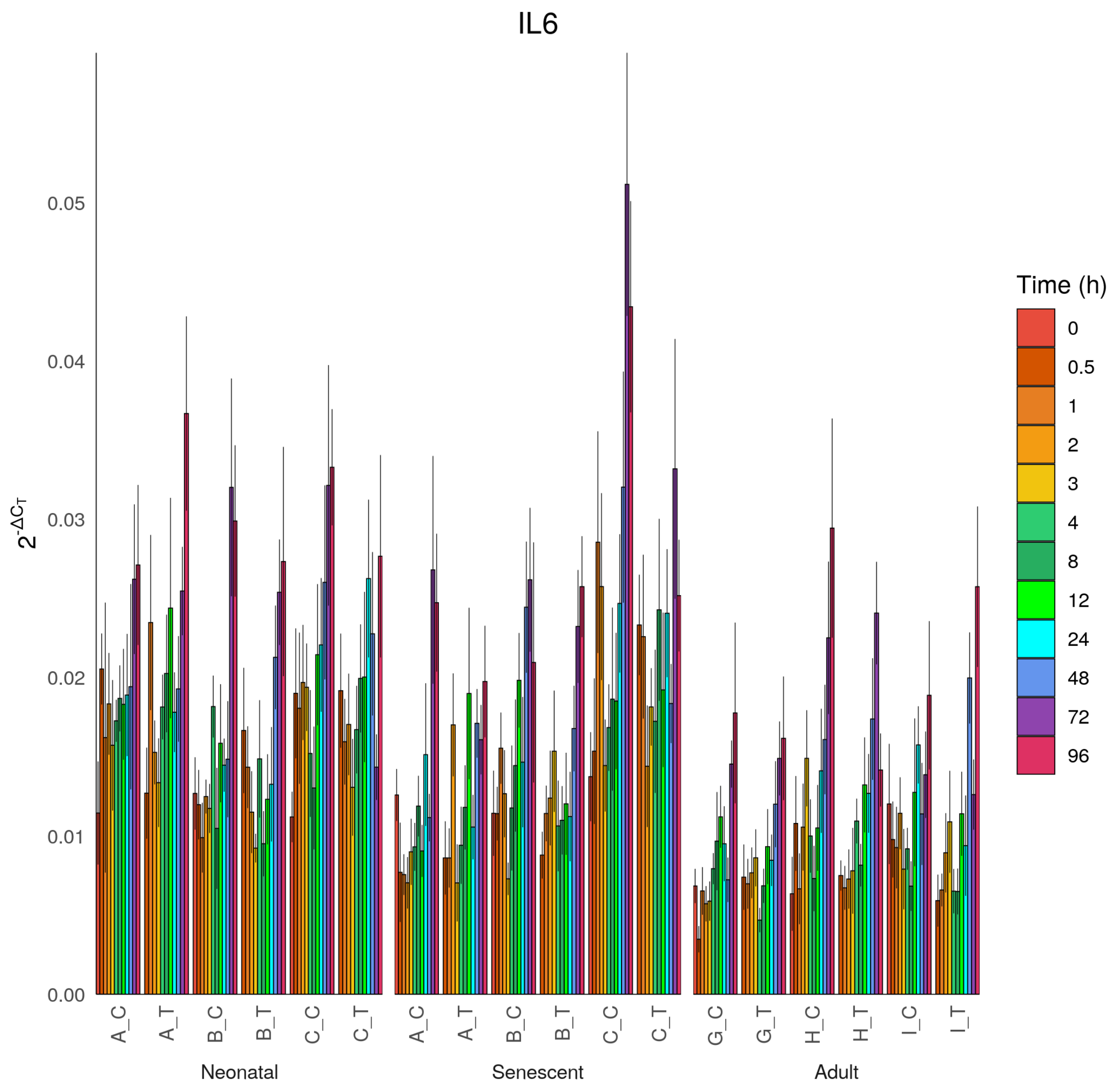

IL6
$2^{-\Delta C_T}$
0.00
0.01
0.02
0.03
0.04
0.05
A_C
A_T
B_C
B_T
C_C
C_T
Neonatal
A_C
A_T
B_C
B_T
C_C
C_T
Senescent
G_C
G_T
H_C
H_T
I_C
I_T
Adult
Time (h)
0
0.5
1
2
3
4
8
12
24
48
72
96

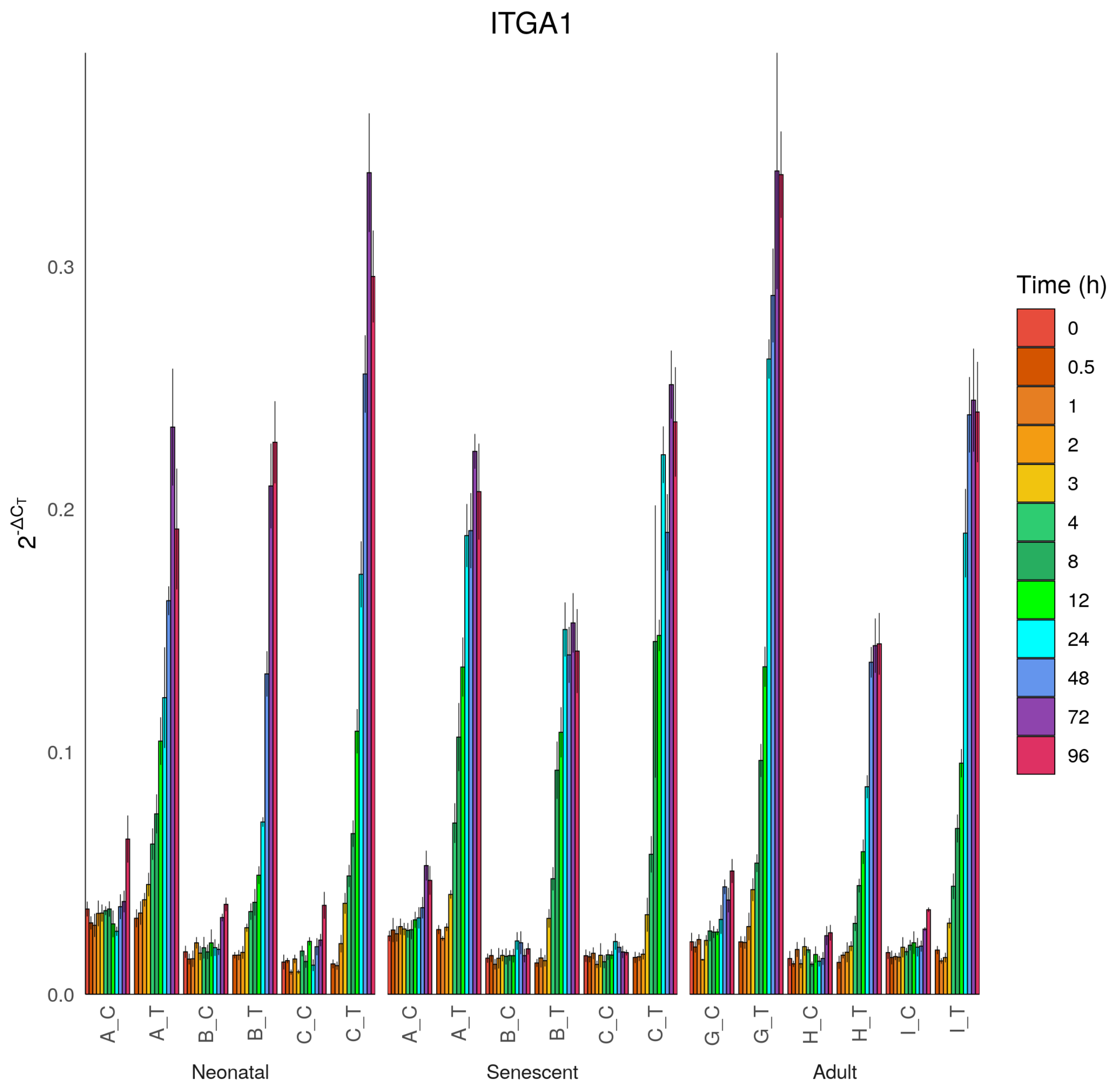

ITGA1
$2^{-\Delta C_T}$
0.0
0.1
0.2
0.3
A_C
A_T
B_C
B_T
C_C
C_T
A_C
A_T
B_C
B_T
C_C
C_T
G_C
G_T
H_C
H_T
I_C
I_T
Neonatal
Senescent
Adult
Time (h)
0
0.5
1
2
3
4
8
12
24
48
72
96

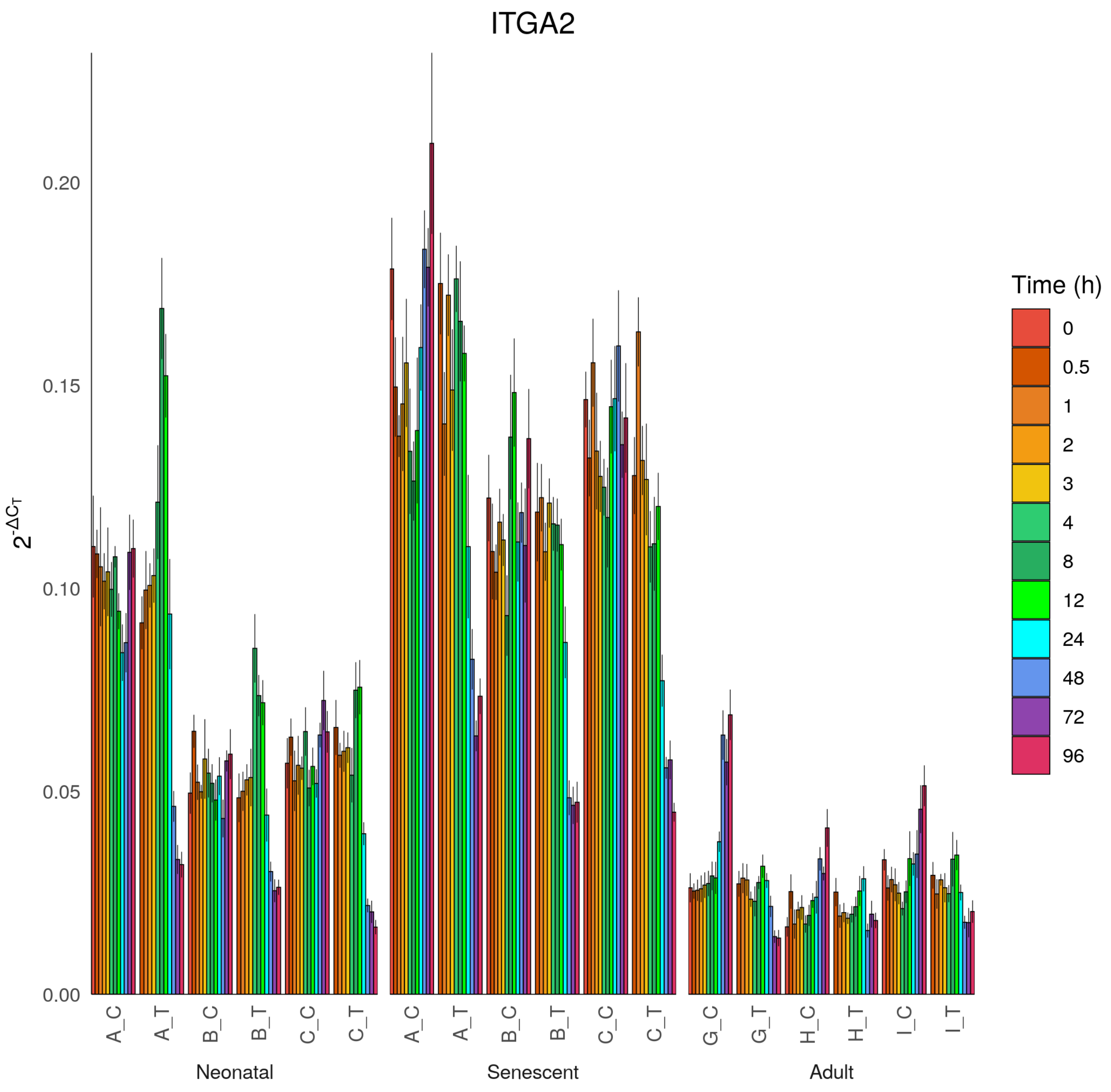

ITGA2
$2^{-\Delta C_T}$
0.00
0.05
0.10
0.15
0.20
A_C
A_T
B_C
B_T
C_C
C_T
G_C
G_T
H_C
H_T
I_C
I_T
Neonatal
Senescent
Adult
Time (h)
0
0.5
1
2
3
4
8
12
24
48
72
96

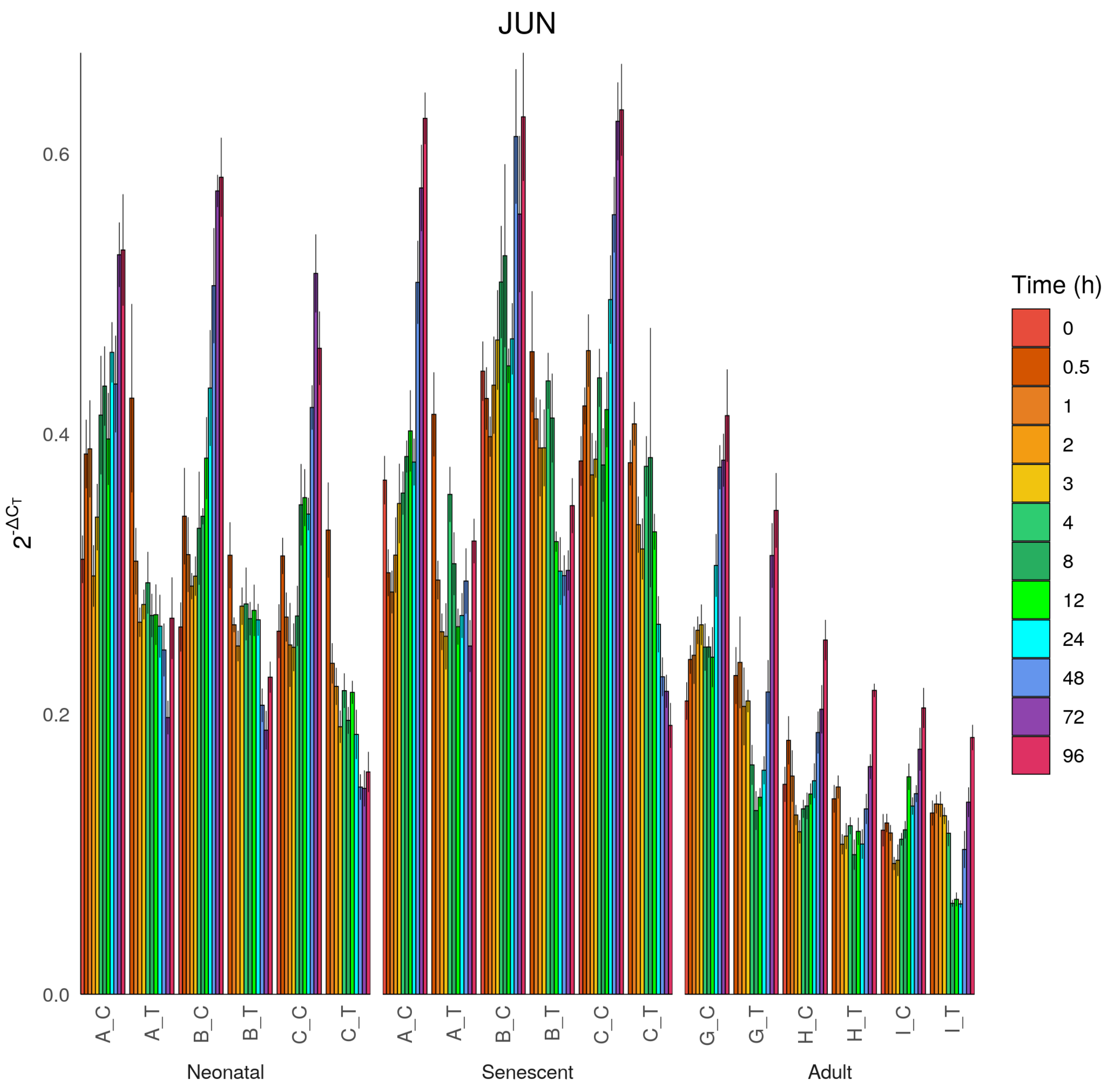
JUN
2^-ΔCT
0.0
0.2
0.4
0.6
A_C
A_T
B_C
B_T
C_C
C_T
Neonatal
A_C
A_T
B_C
B_T
C_C
C_T
Senescent
G_C
G_T
H_C
H_T
I_C
I_T
Adult
Time (h)
0
0.5
1
2
3
4
8
12
24
48
72
96

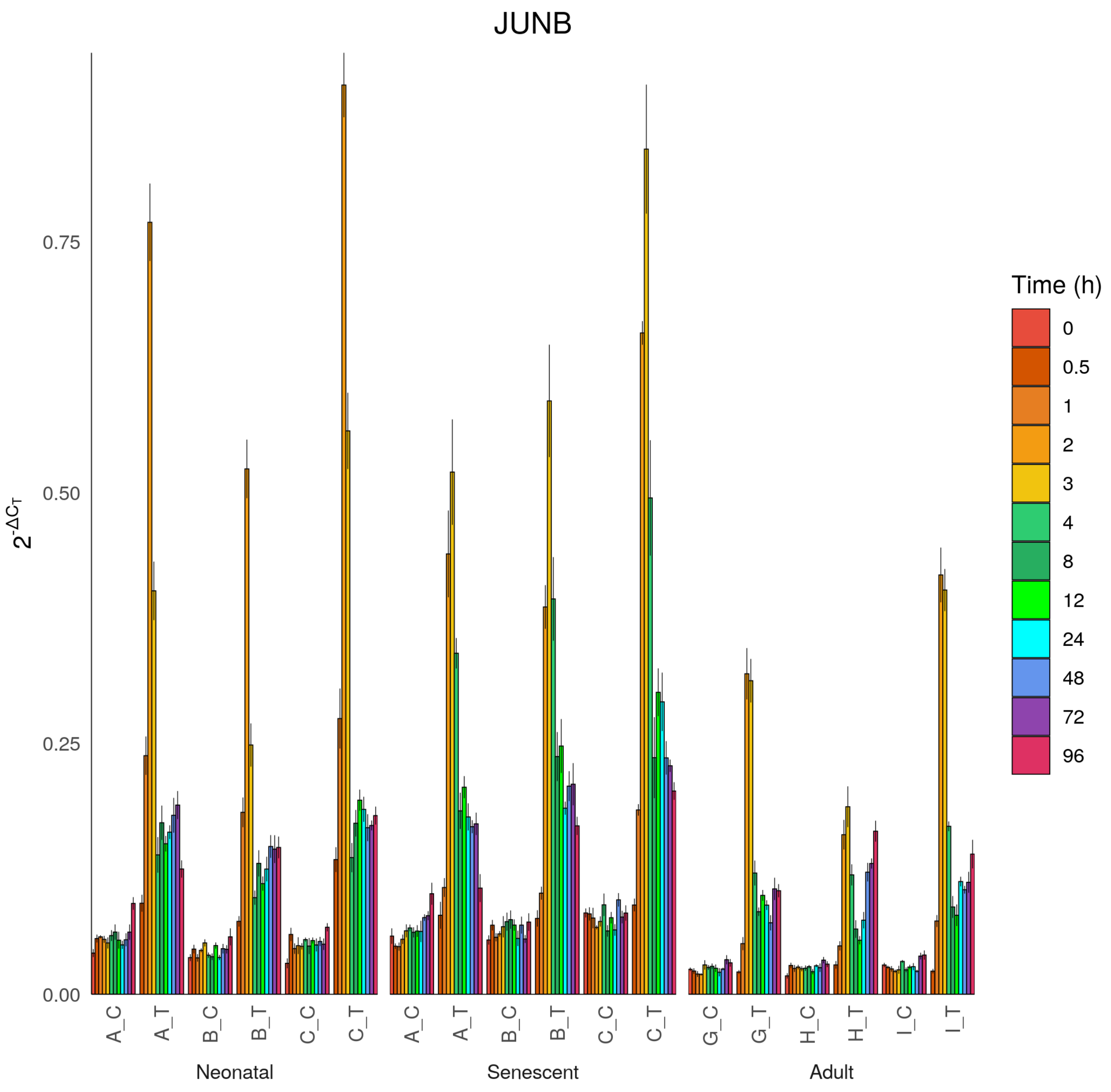

JUNB
$2^{-\Delta C_T}$
0.00
0.25
0.50
0.75
A_C
A_T
B_C
B_T
C_C
C_T
G_C
G_T
H_C
H_T
I_C
I_T
Neonatal
Senescent
Adult
Time (h)
0
0.5
1
2
3
4
8
12
24
48
72
96

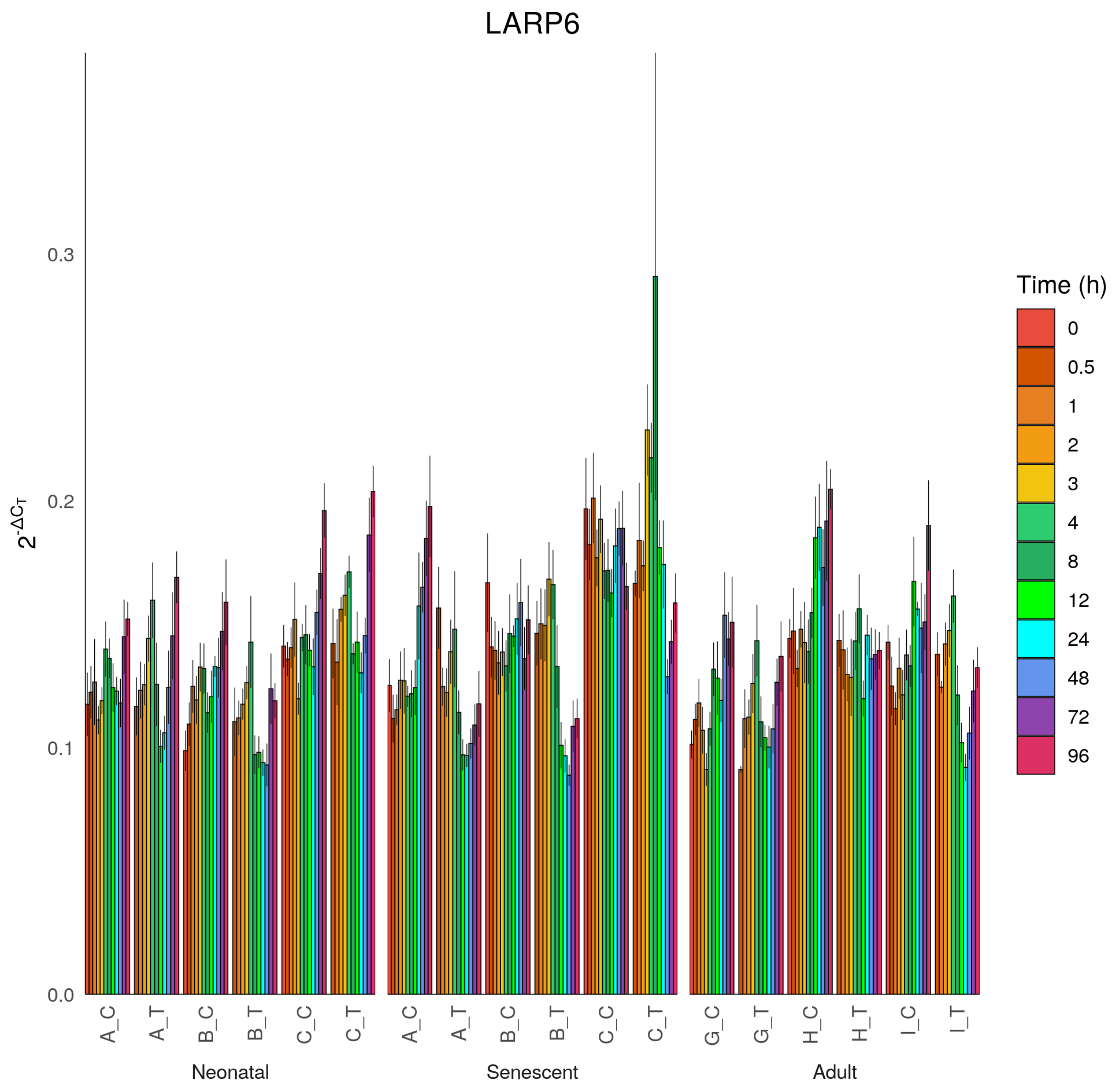

LARP6
2^-ΔCT
0.0
0.1
0.2
0.3
A_C
A_T
B_C
B_T
C_C
C_T
A_C
A_T
B_C
B_T
C_C
C_T
G_C
G_T
H_C
H_T
I_C
I_T
Neonatal
Senescent
Adult
Time (h)
0
0.5
1
2
3
4
8
12
24
48
72
96

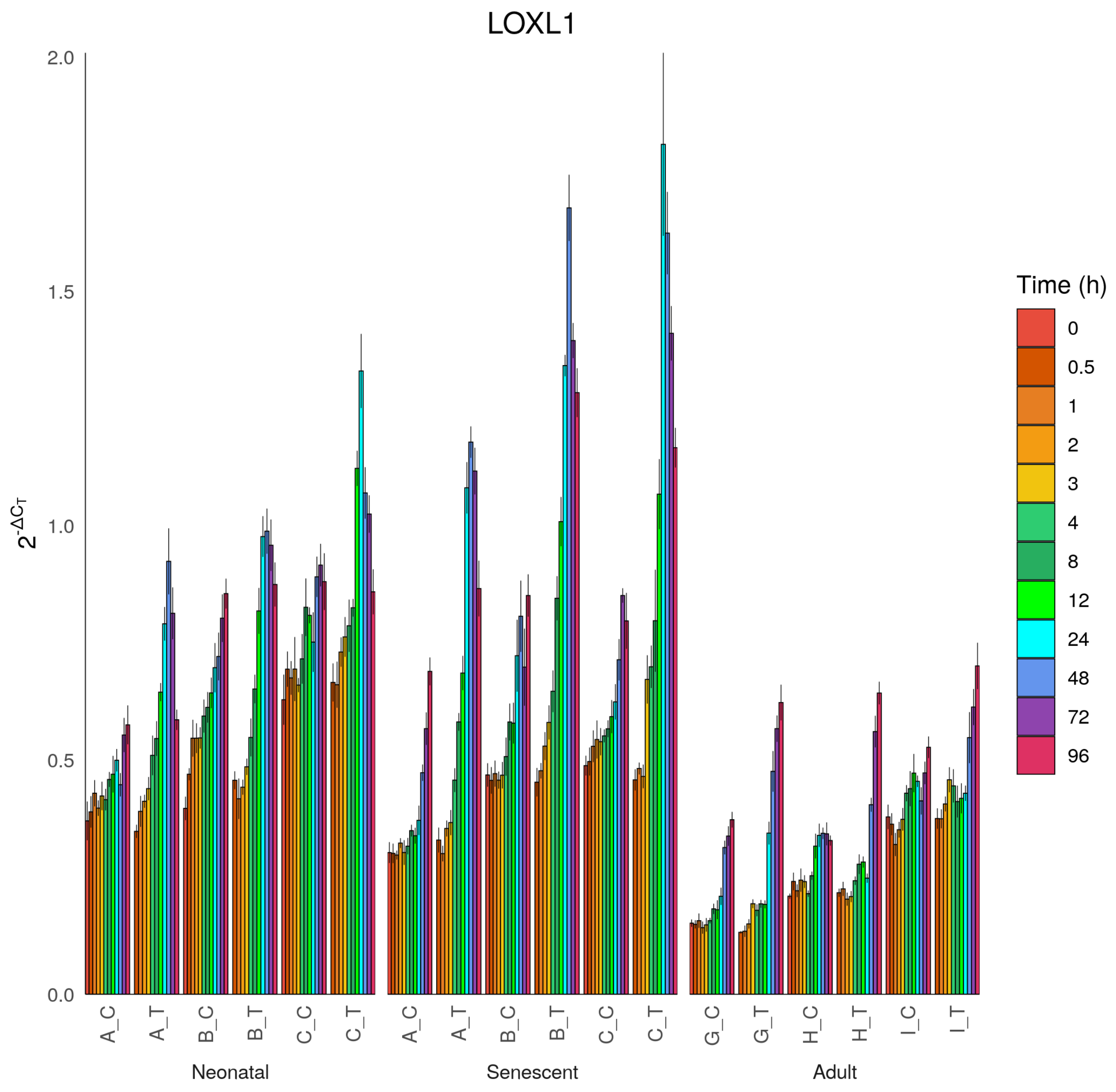

LOXL1
2.0
1.5
1.0
0.5
0.0
$2^{-\Delta C_T}$
A_C
A_T
B_C
B_T
C_C
C_T
Neonatal
A_C
A_T
B_C
B_T
C_C
C_T
Senescent
G_C
G_T
H_C
H_T
I_C
I_T
Adult
Time (h)
0
0.5
1
2
3
4
8
12
24
48
72
96

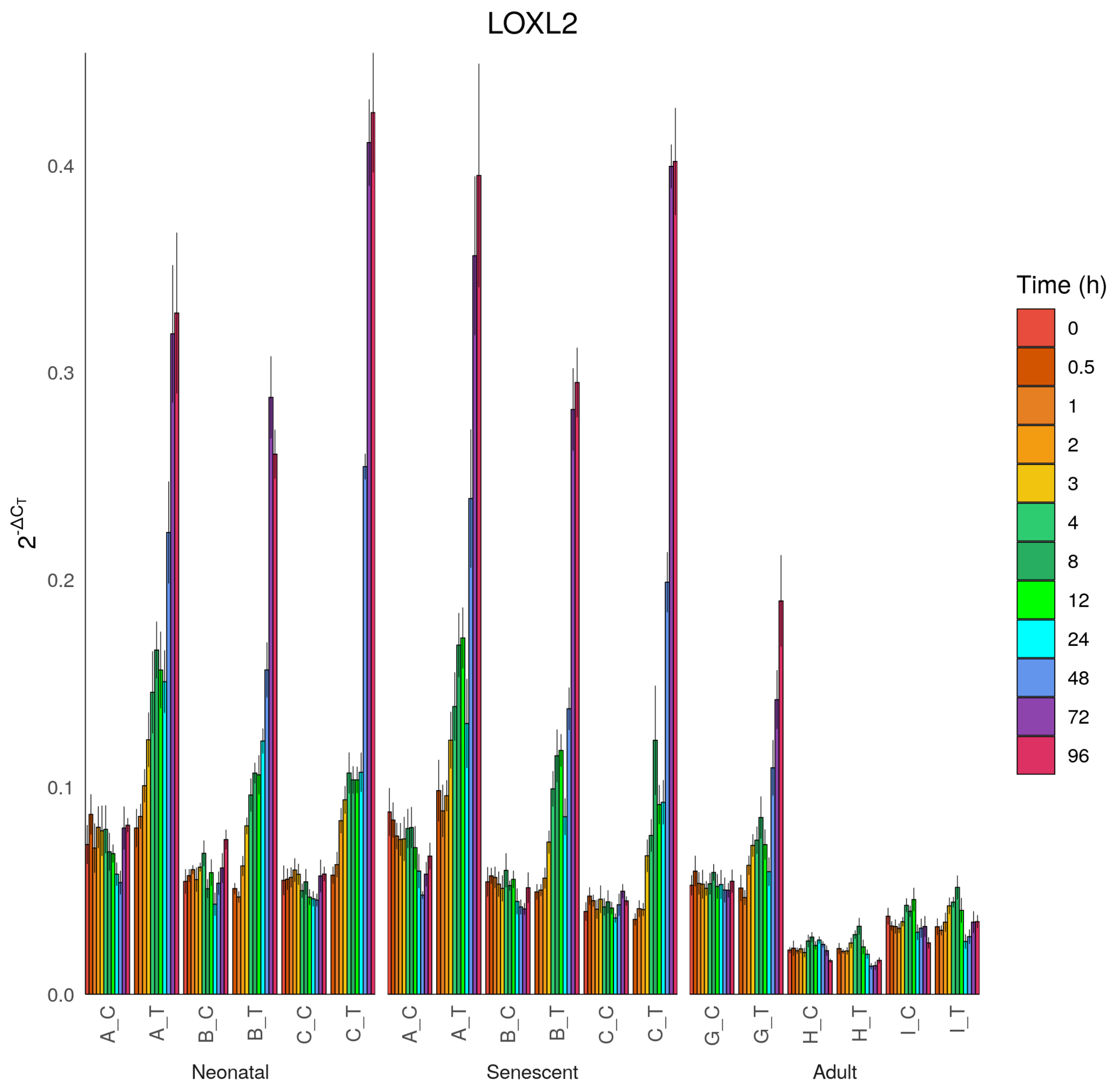

LOXL2
2^-ΔCT
0.0
0.1
0.2
0.3
0.4
A_C
A_T
B_C
B_T
C_C
C_T
G_C
G_T
H_C
H_T
I_C
I_T
Neonatal
Senescent
Adult
Time (h)
0
0.5
1
2
3
4
8
12
24
48
72
96

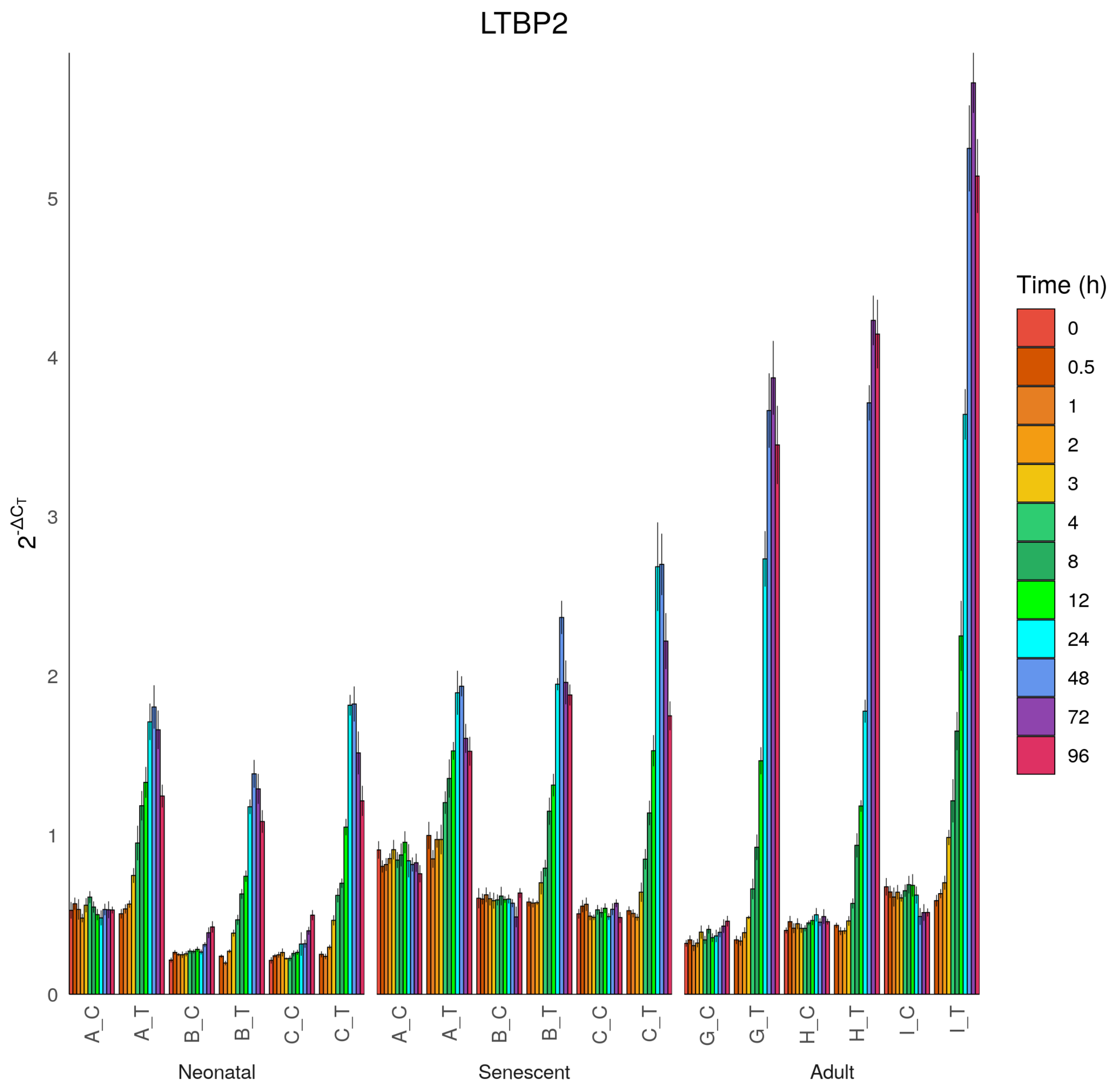
LTBP2
$2^{-\Delta C_T}$
0
1
2
3
4
5
A_C
A_T
B_C
B_T
C_C
C_T
Neonatal
A_C
A_T
B_C
B_T
C_C
C_T
Senescent
G_C
G_T
H_C
H_T
I_C
I_T
Adult
Time (h)
0
0.5
1
2
3
4
8
12
24
48
72
96

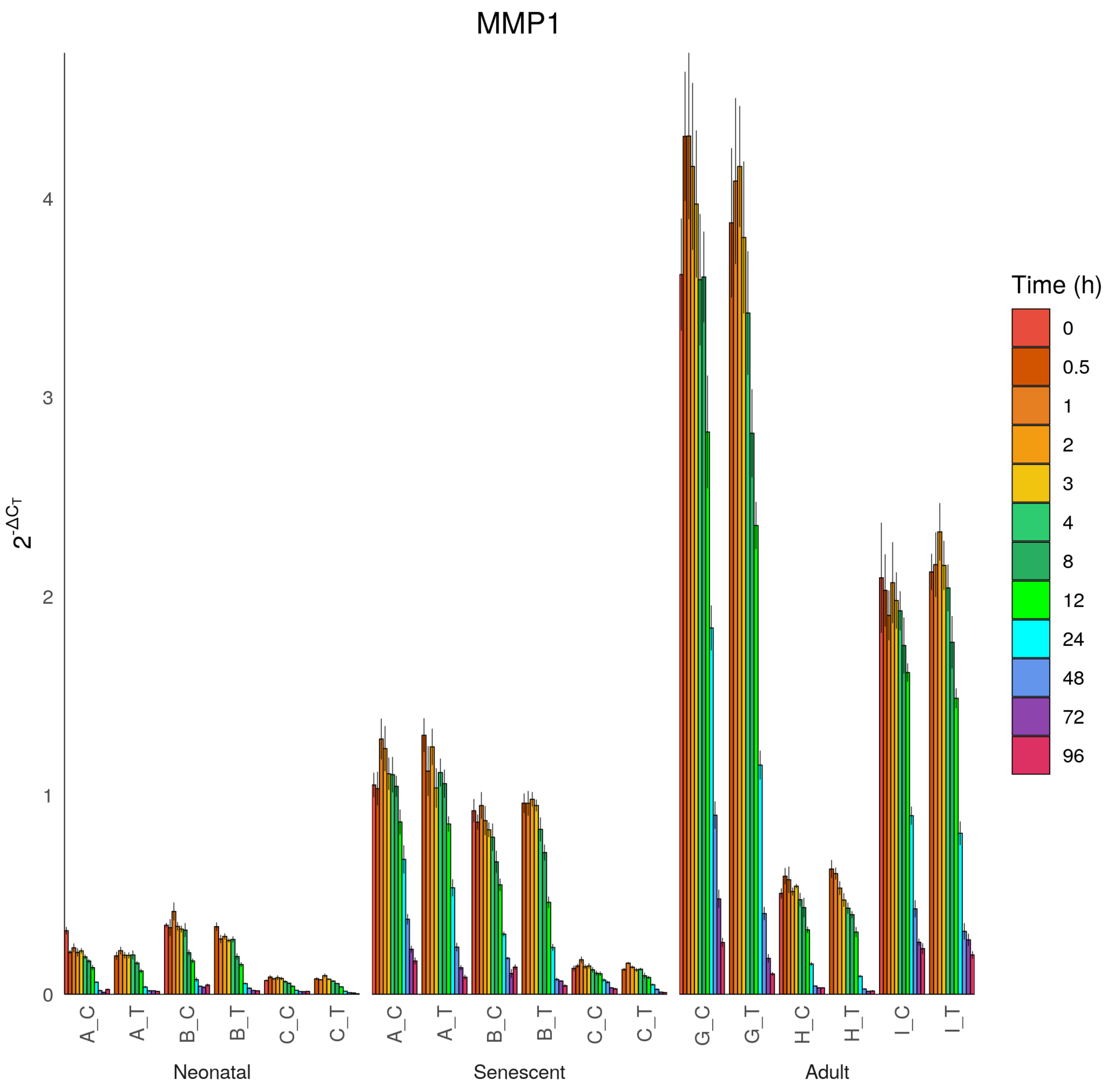

MMP1
$2^{-\Delta C_T}$
0
1
2
3
4
A_C
A_T
B_C
B_T
C_C
C_T
Neonatal
A_C
A_T
B_C
B_T
C_C
C_T
Senescent
G_C
G_T
H_C
H_T
I_C
I_T
Adult
Time (h)
0
0.5
1
2
3
4
8
12
24
48
72
96

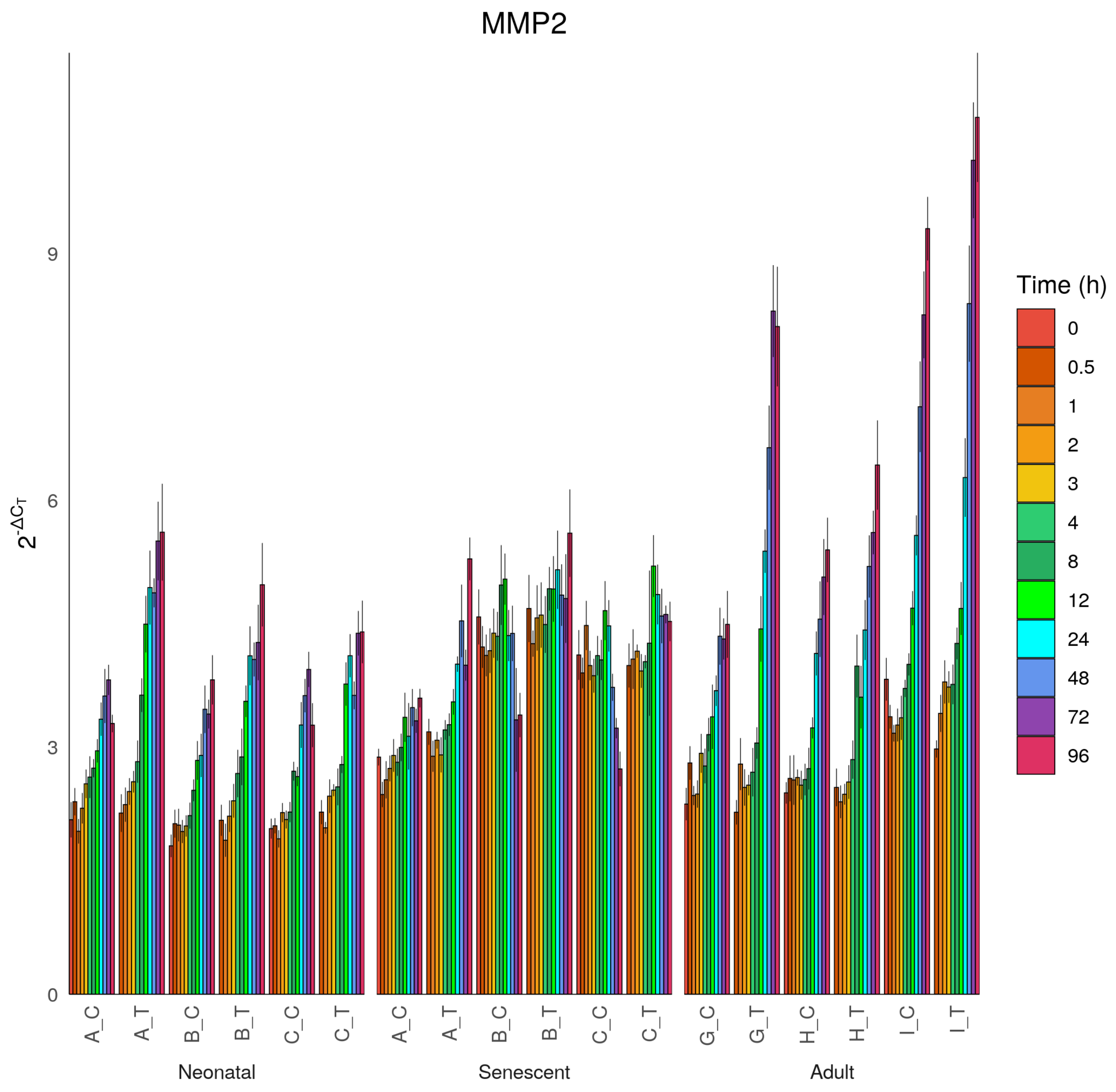

MMP2
$2^{-\Delta C_T}$
0
3
6
9
A_C
A_T
B_C
B_T
C_C
C_T
Neonatal
A_C
A_T
B_C
B_T
C_C
C_T
Senescent
G_C
G_T
H_C
H_T
I_C
I_T
Adult
Time (h)
0
0.5
1
2
3
4
8
12
24
48
72
96

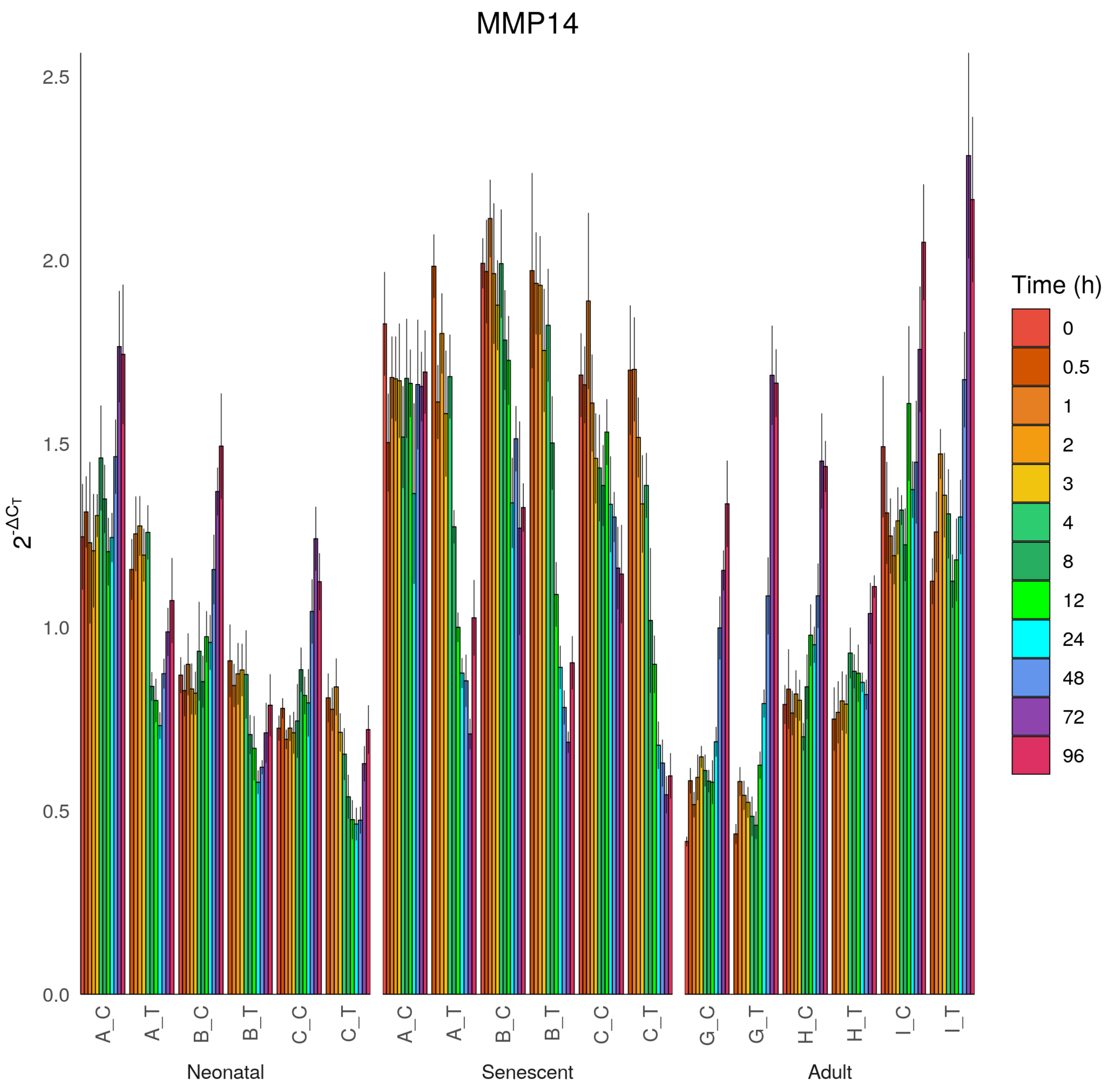
MMP14
2.5
2.0
1.5
1.0
0.5
0.0
$2^{-\Delta C_T}$
A_C
A_T
B_C
B_T
C_C
C_T
Neonatal
A_C
A_T
B_C
B_T
C_C
C_T
Senescent
G_C
G_T
H_C
H_T
I_C
I_T
Adult
Time (h)
0
0.5
1
2
3
4
8
12
24
48
72
96

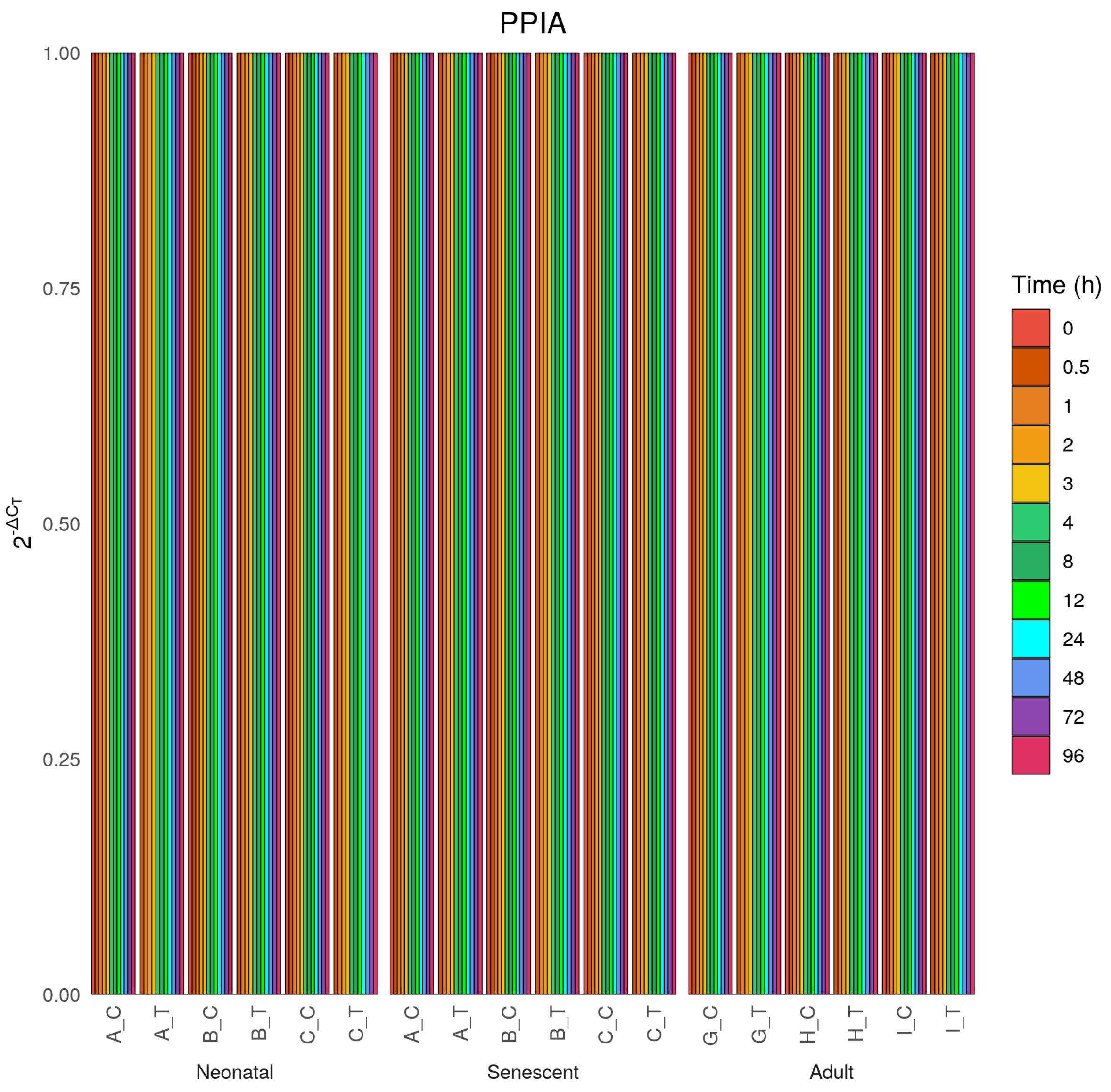

PPIA
$2^{-\Delta C_T}$
1.00
0.75
0.50
0.25
0.00
A_C
A_T
B_C
B_T
C_C
C_T
Neonatal
A_C
A_T
B_C
B_T
C_C
C_T
Senescent
G_C
G_T
H_C
H_T
I_C
I_T
Adult
Time (h)
0
0.5
1
2
3
4
8
12
24
48
72
96

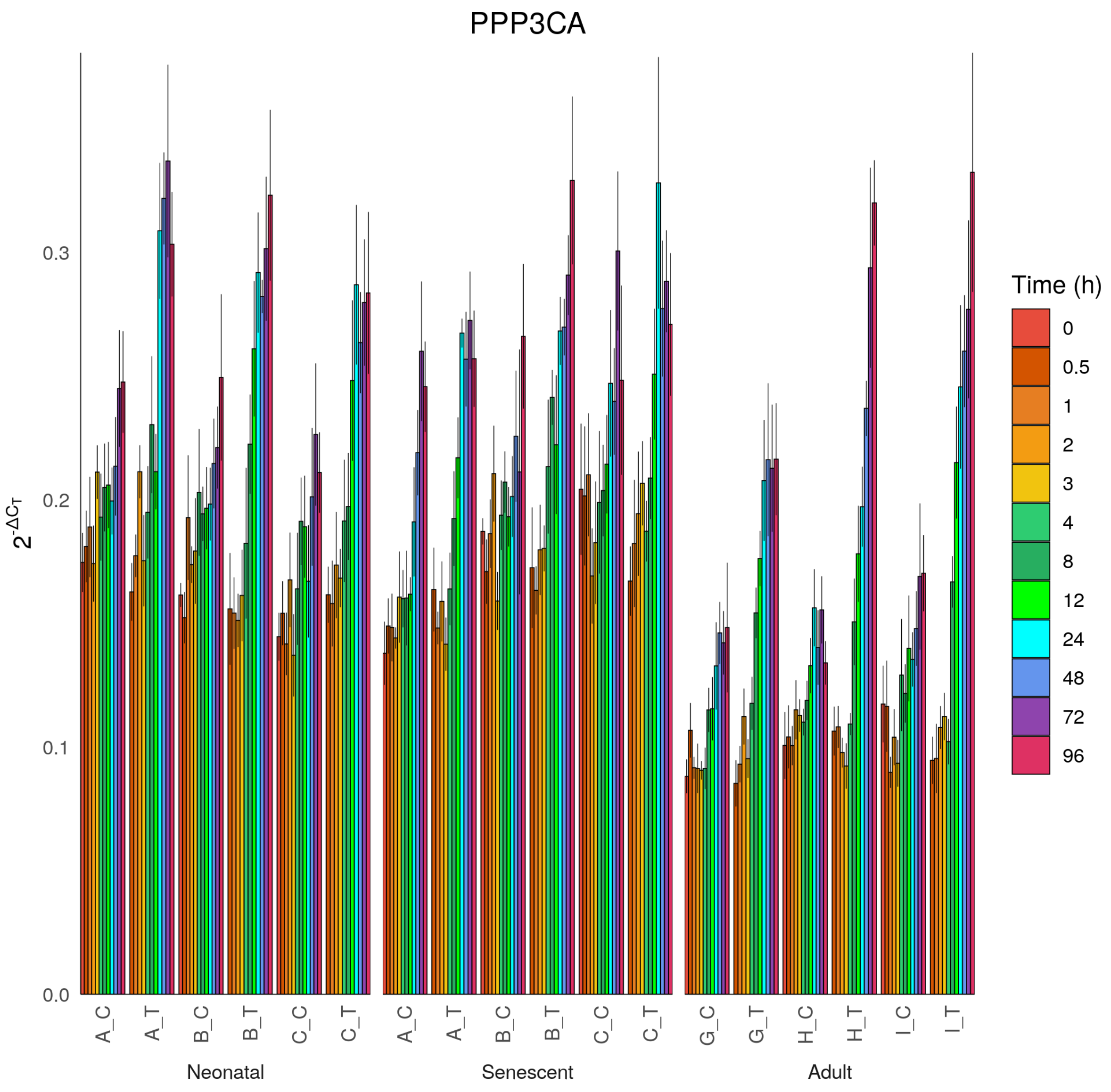

PPP3CA
2^-ΔCT
0.0
0.1
0.2
0.3
A_C
A_T
B_C
B_T
C_C
C_T
G_C
G_T
H_C
H_T
I_C
I_T
Neonatal
Senescent
Adult
Time (h)
0
0.5
1
2
3
4
8
12
24
48
72
96

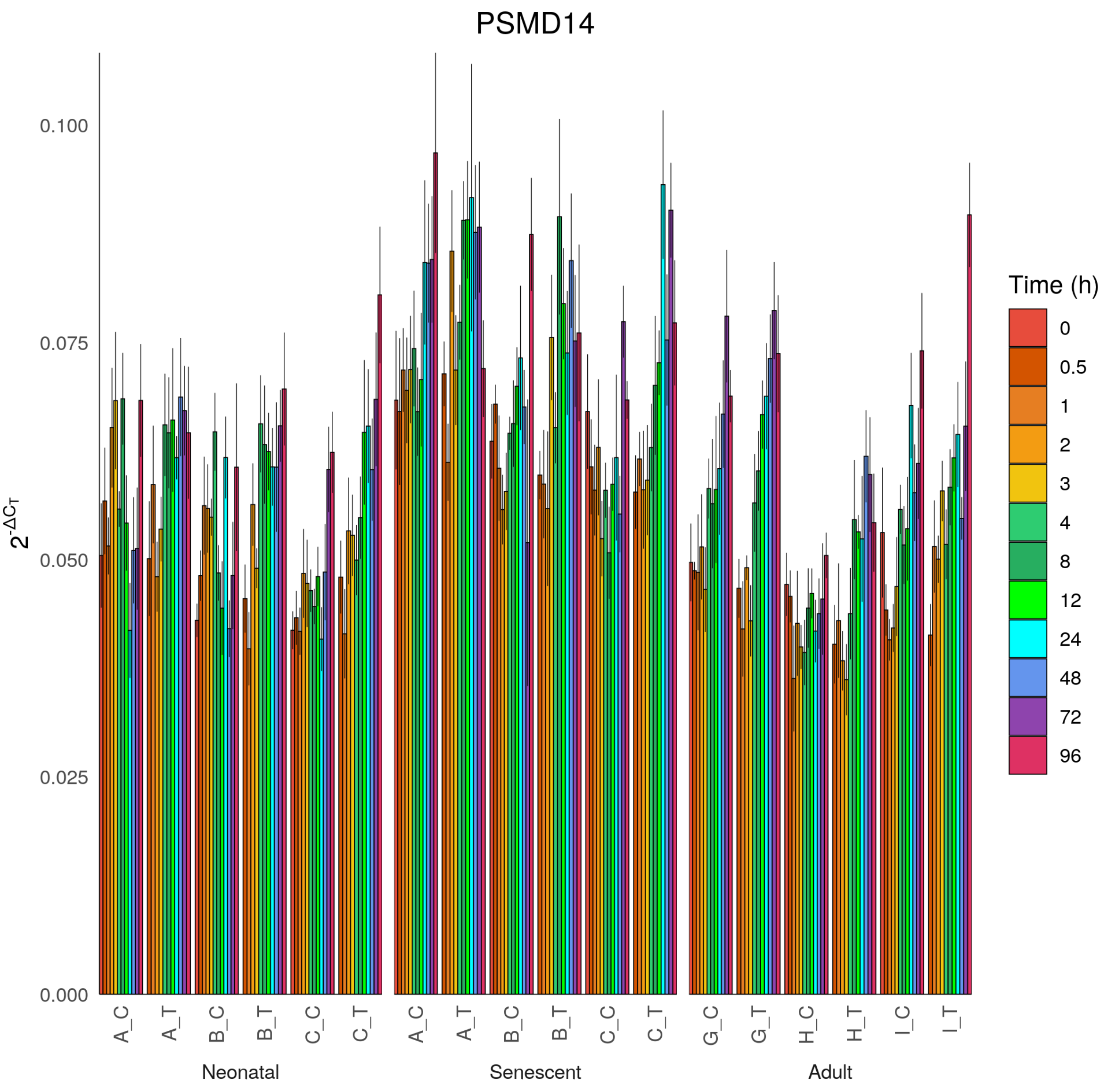
PSMD14
$2^{-\Delta C_T}$
0.100
0.075
0.050
0.025
0.000
A_C
A_T
B_C
B_T
C_C
C_T
G_C
G_T
H_C
H_T
I_C
I_T
Neonatal
Senescent
Adult
Time (h)
0
0.5
1
2
3
4
8
12
24
48
72
96

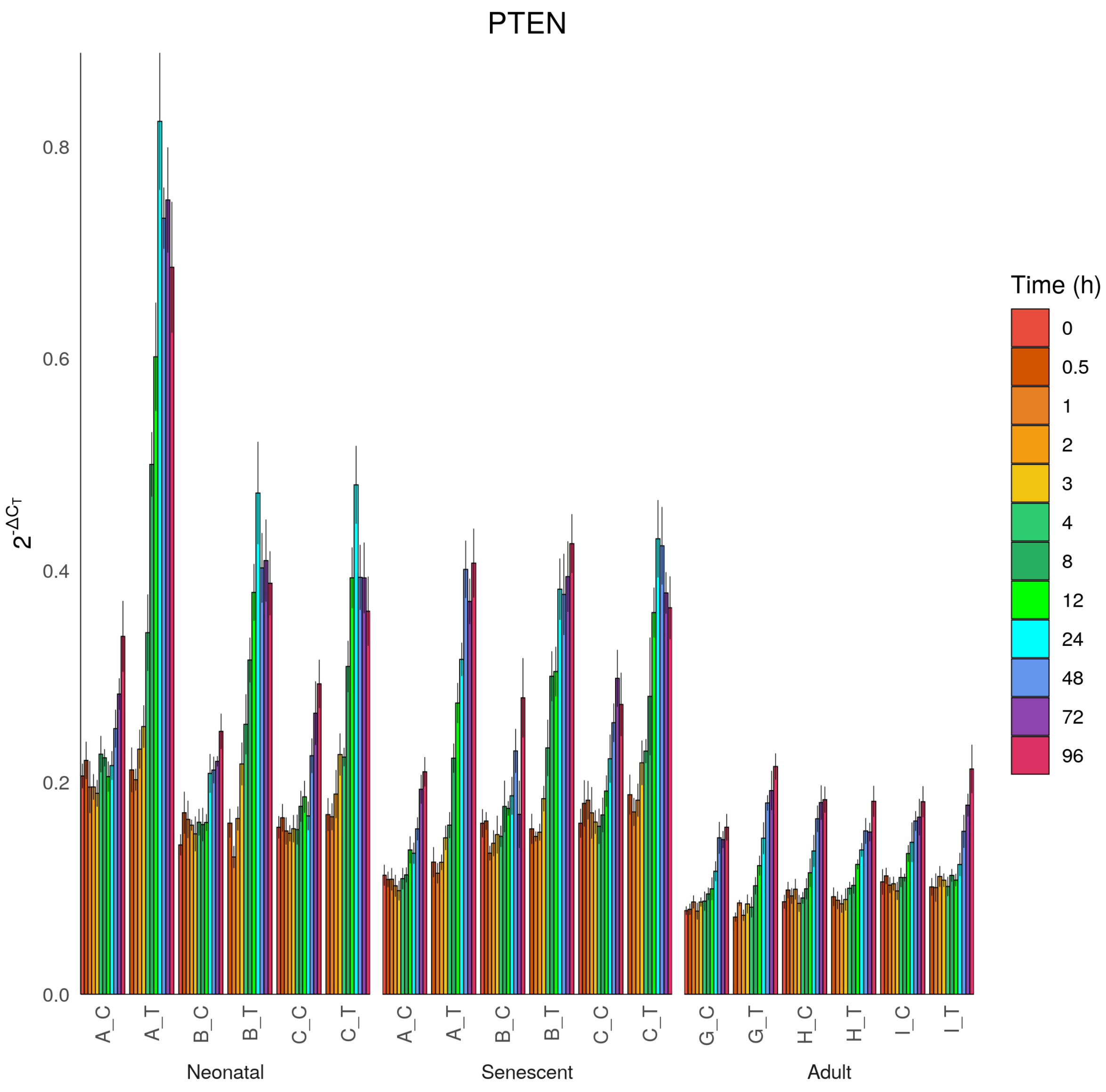
PTEN
$2^{-\Delta C_T}$
0.0
0.2
0.4
0.6
0.8
A_C
A_T
B_C
B_T
C_C
C_T
Neonatal
A_C
A_T
B_C
B_T
C_C
C_T
Senescent
G_C
G_T
H_C
H_T
I_C
I_T
Adult
Time (h)
0
0.5
1
2
3
4
8
12
24
48
72
96

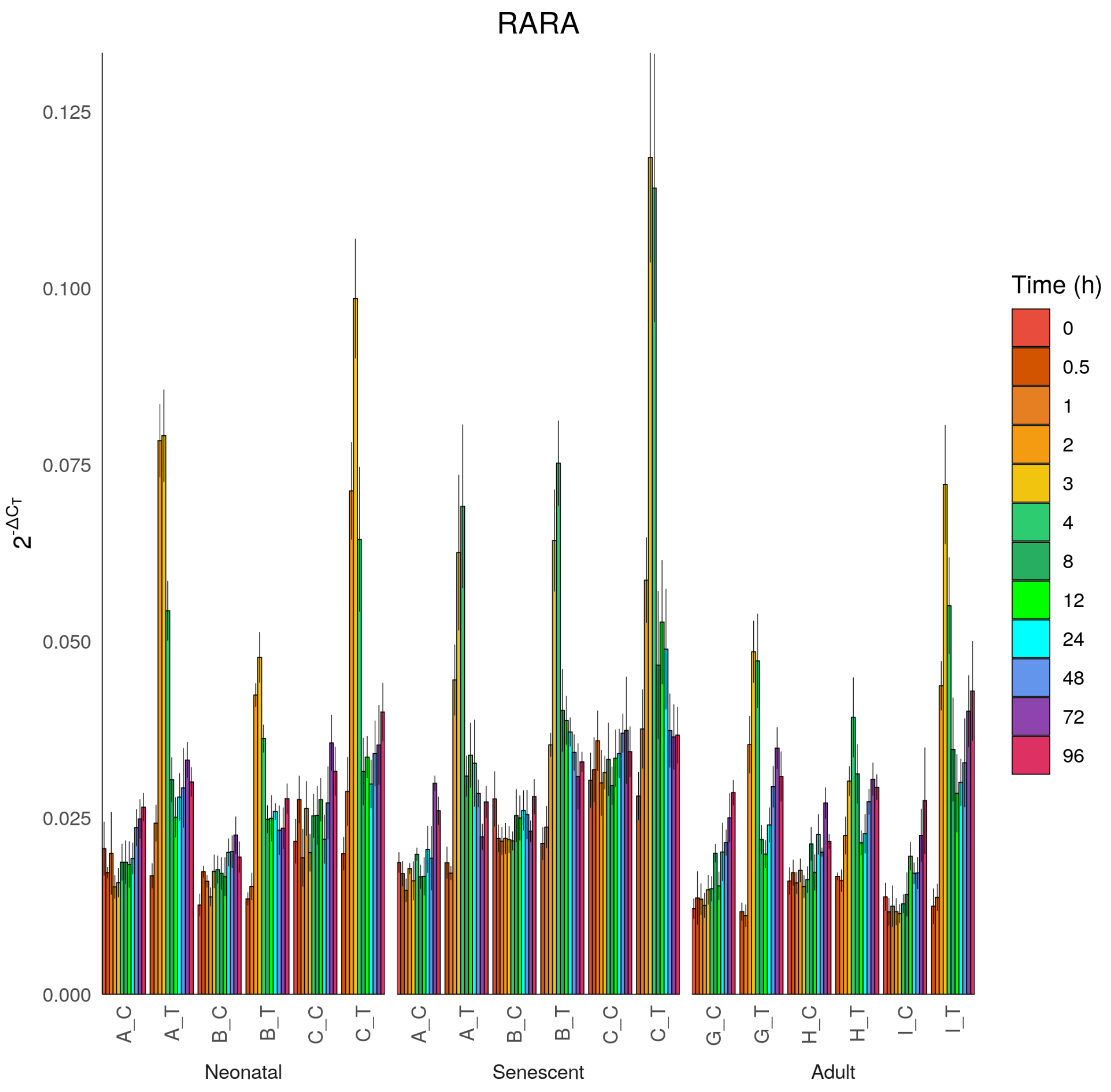
RARA
$2^{-\Delta C_T}$
0.125
0.100
0.075
0.050
0.025
0.000
A_C
A_T
B_C
B_T
C_C
C_T
G_C
G_T
H_C
H_T
I_C
I_T
Neonatal
Senescent
Adult
Time (h)
0
0.5
1
2
3
4
8
12
24
48
72
96

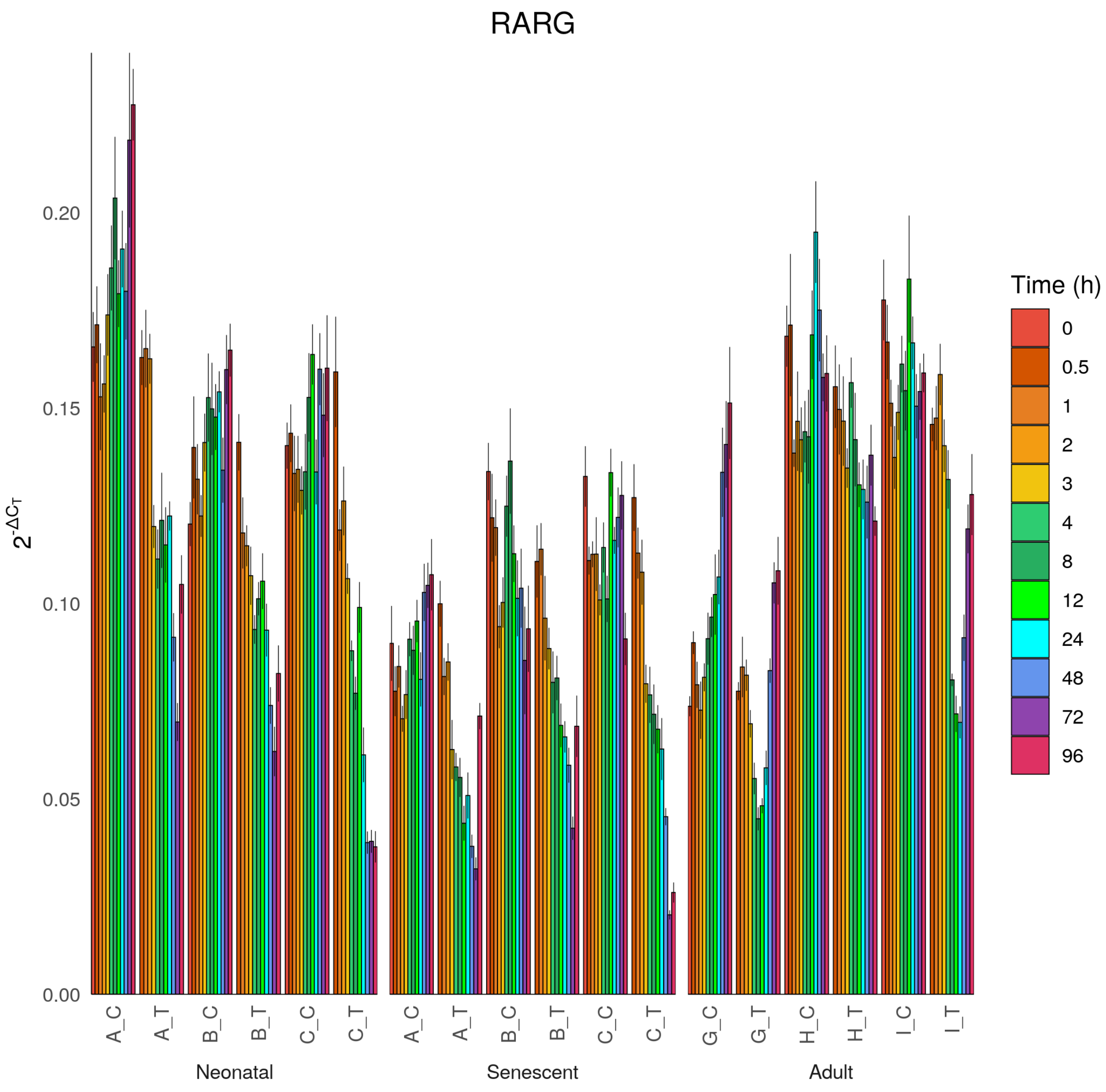

RARG
$2^{-\Delta C_T}$
0.00
0.05
0.10
0.15
0.20
A_C
A_T
B_C
B_T
C_C
C_T
G_C
G_T
H_C
H_T
I_C
I_T
Neonatal
Senescent
Adult
Time (h)
0
0.5
1
2
3
4
8
12
24
48
72
96

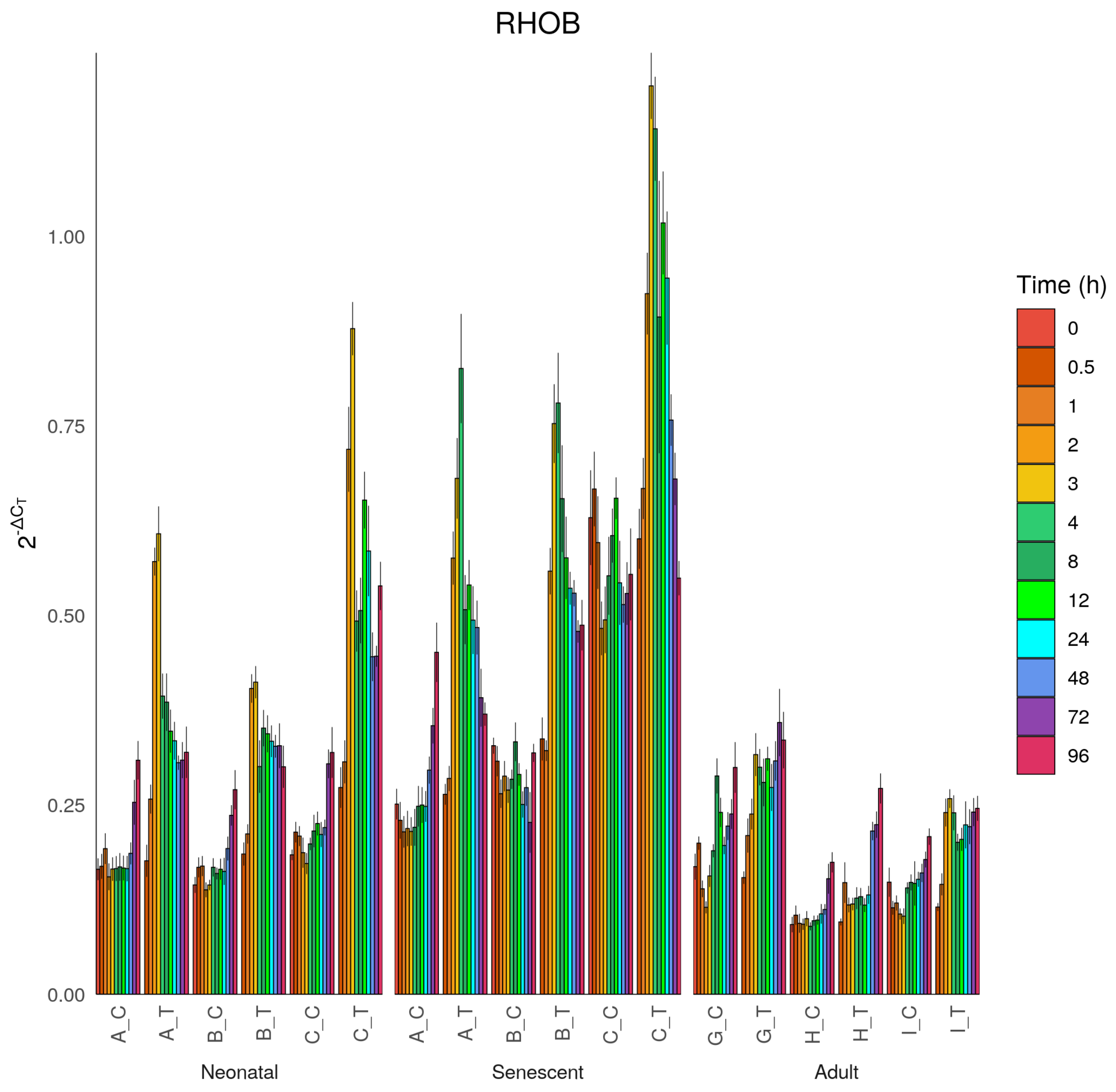

RHOB
$2^{-\Delta C_T}$
1.00
0.75
0.50
0.25
0.00
A_C
A_T
B_C
B_T
C_C
C_T
Neonatal
A_C
A_T
B_C
B_T
C_C
C_T
Senescent
G_C
G_T
H_C
H_T
I_C
I_T
Adult
Time (h)
0
0.5
1
2
3
4
8
12
24
48
72
96

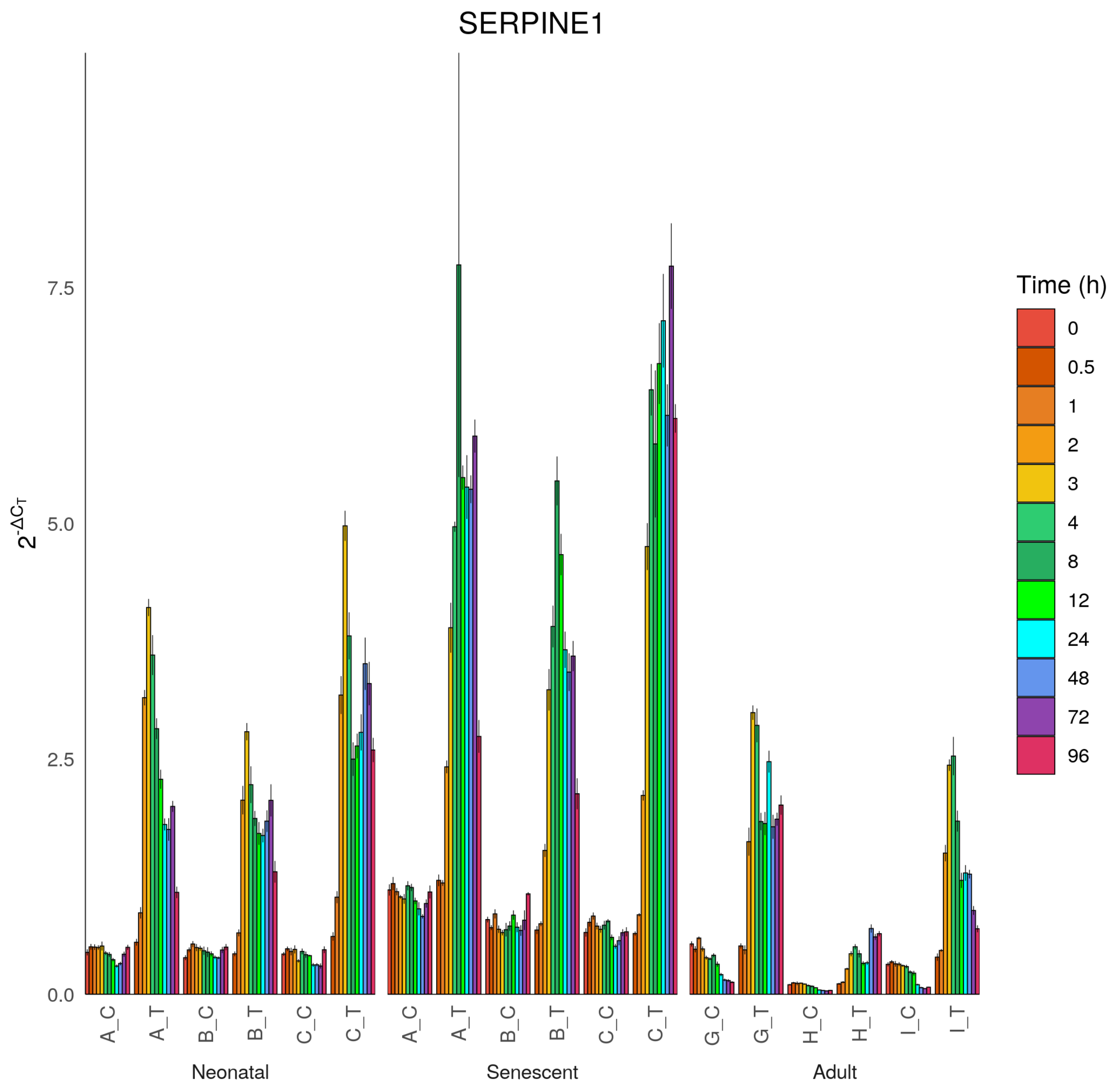
SERPINE1
$2^{-\Delta C_T}$
0.0
2.5
5.0
7.5
A_C
A_T
B_C
B_T
C_C
C_T
A_C
A_T
B_C
B_T
C_C
C_T
G_C
G_T
H_C
H_T
I_C
I_T
Neonatal
Senescent
Adult
Time (h)
0
0.5
1
2
3
4
8
12
24
48
72
96

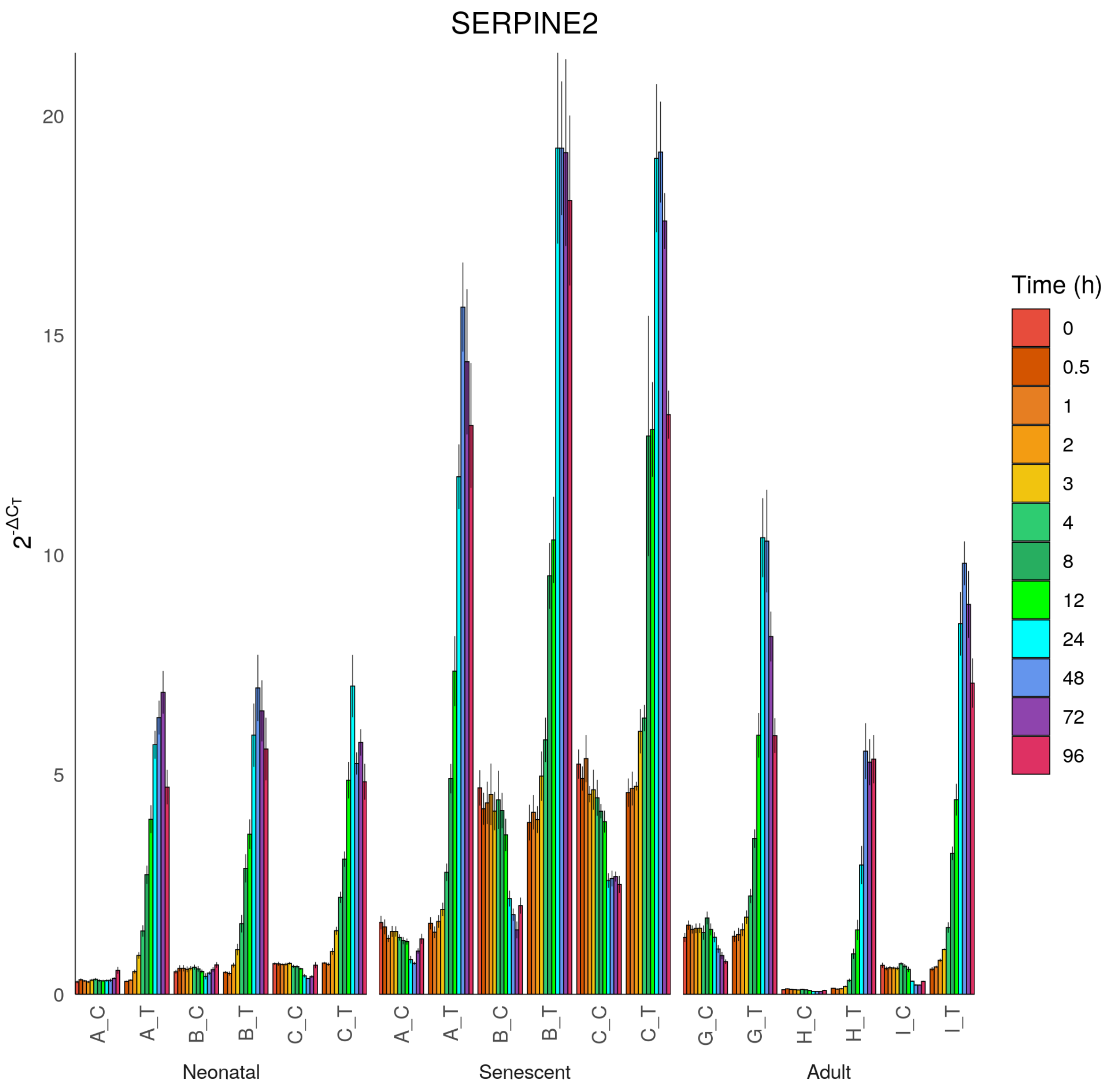

SERPINE2
$2^{-\Delta C_T}$
0
5
10
15
20
A_C
A_T
B_C
B_T
C_C
C_T
A_C
A_T
B_C
B_T
C_C
C_T
G_C
G_T
H_C
H_T
I_C
I_T
Neonatal
Senescent
Adult
Time (h)
0
0.5
1
2
3
4
8
12
24
48
72
96

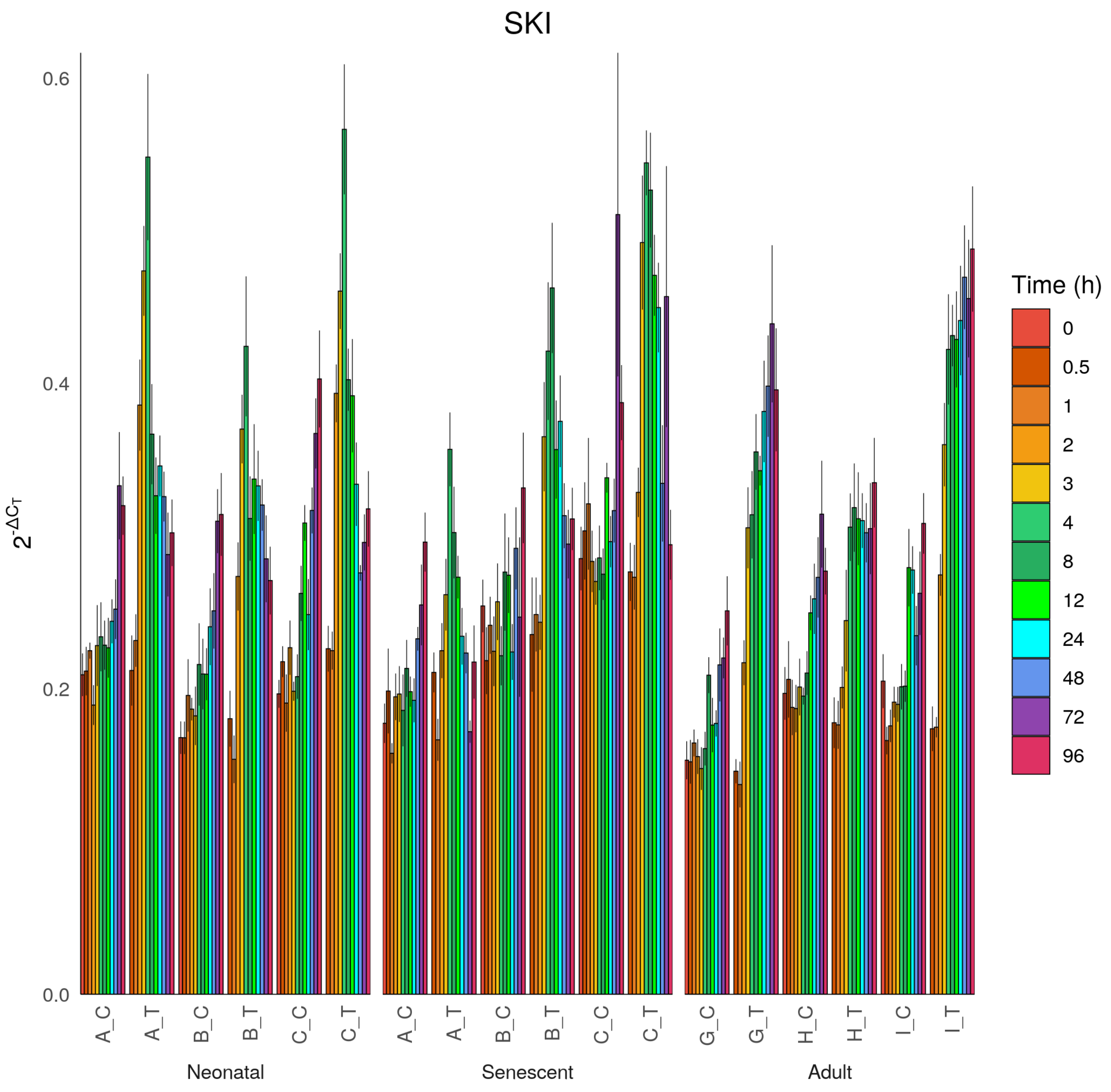

SKI
$2^{-\Delta C_T}$
0.0
0.2
0.4
0.6
A_C
A_T
B_C
B_T
C_C
C_T
G_C
G_T
H_C
H_T
I_C
I_T
Neonatal
Senescent
Adult
Time (h)
0
0.5
1
2
3
4
8
12
24
48
72
96

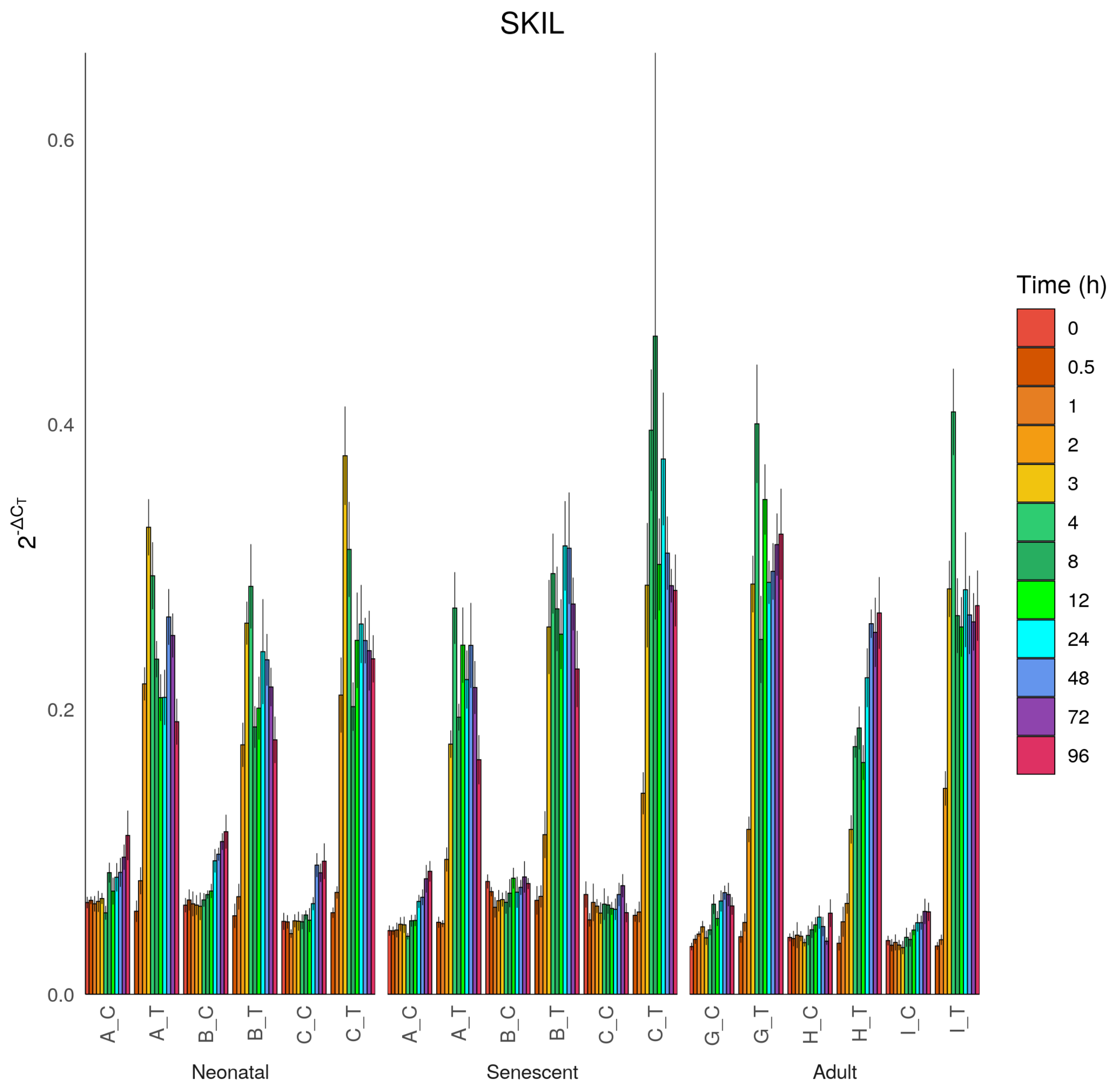
SKIL
$2^{-\Delta C_T}$
0.0
0.2
0.4
0.6
A_C
A_T
B_C
B_T
C_C
C_T
G_C
G_T
H_C
H_T
I_C
I_T
Neonatal
Senescent
Adult
Time (h)
0
0.5
1
2
3
4
8
12
24
48
72
96

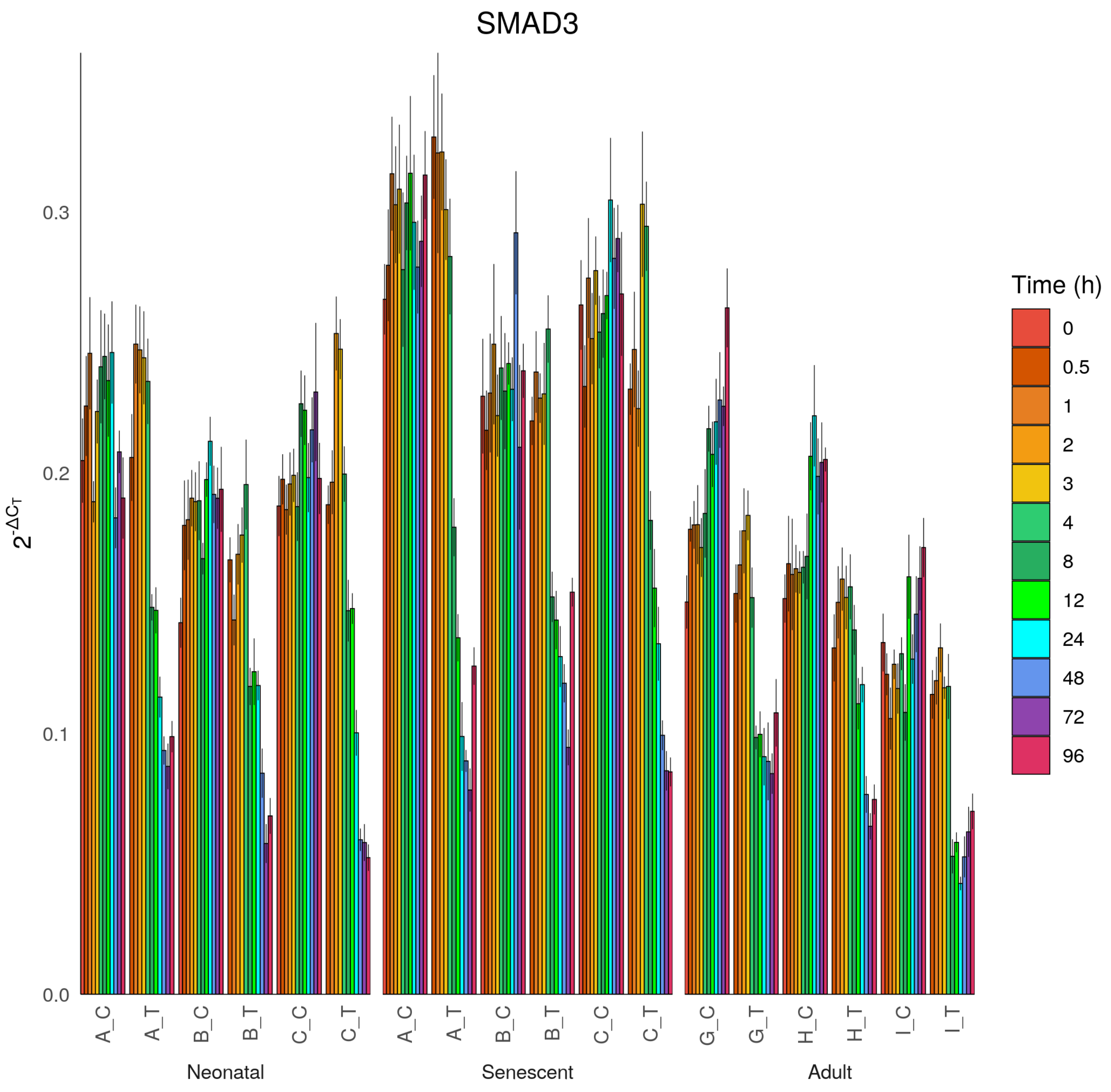
SMAD3
$2^{-\Delta C_T}$
0.0
0.1
0.2
0.3
A_C
A_T
B_C
B_T
C_C
C_T
Neonatal
A_C
A_T
B_C
B_T
C_C
C_T
Senescent
G_C
G_T
H_C
H_T
I_C
I_T
Adult
Time (h)
0
0.5
1
2
3
4
8
12
24
48
72
96

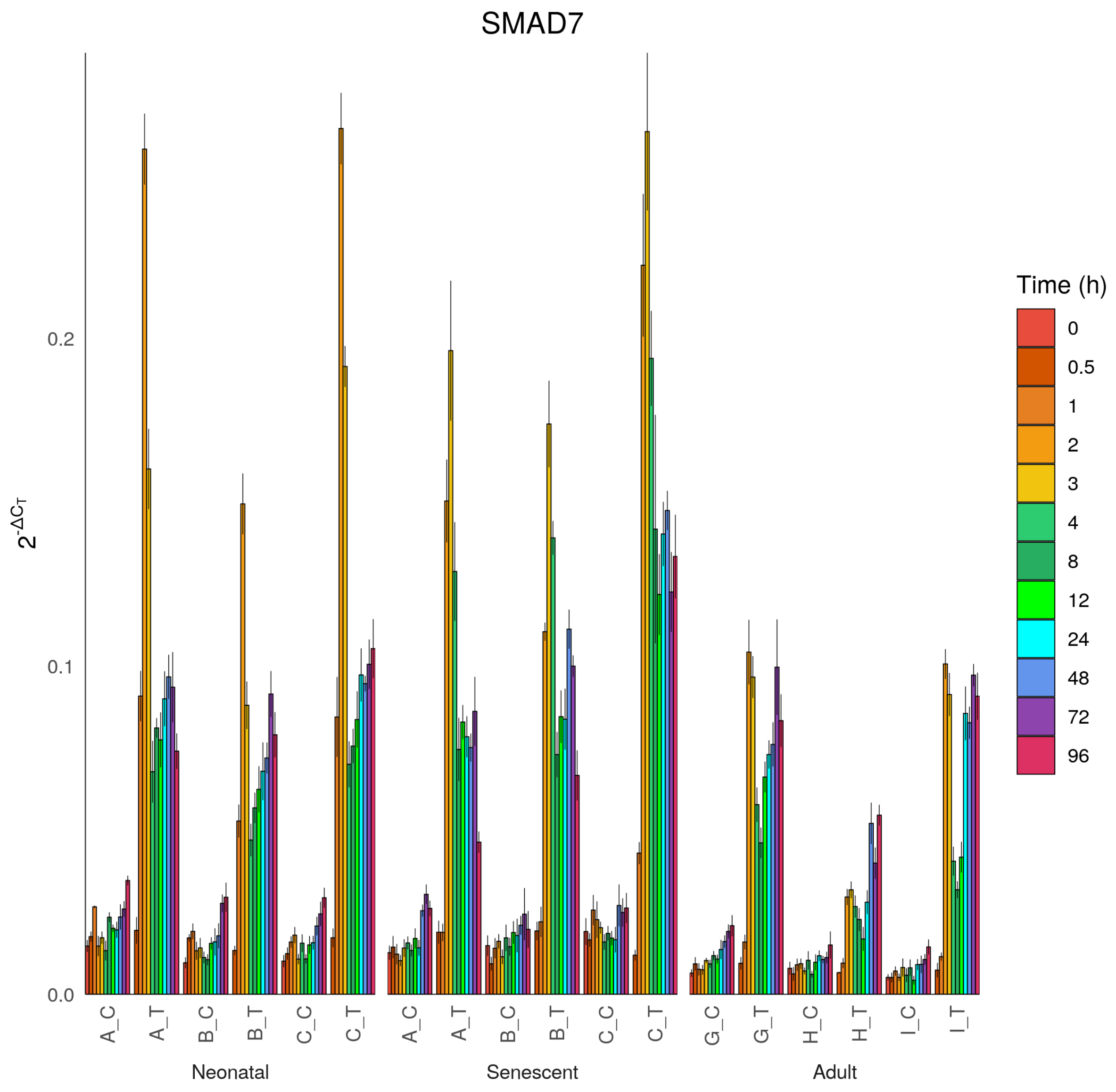

SMAD7
$2^{-\Delta C_T}$
0.0
0.1
0.2
A_C
A_T
B_C
B_T
C_C
C_T
A_C
A_T
B_C
B_T
C_C
C_T
G_C
G_T
H_C
H_T
I_C
I_T
Neonatal
Senescent
Adult
Time (h)
0
0.5
1
2
3
4
8
12
24
48
72
96

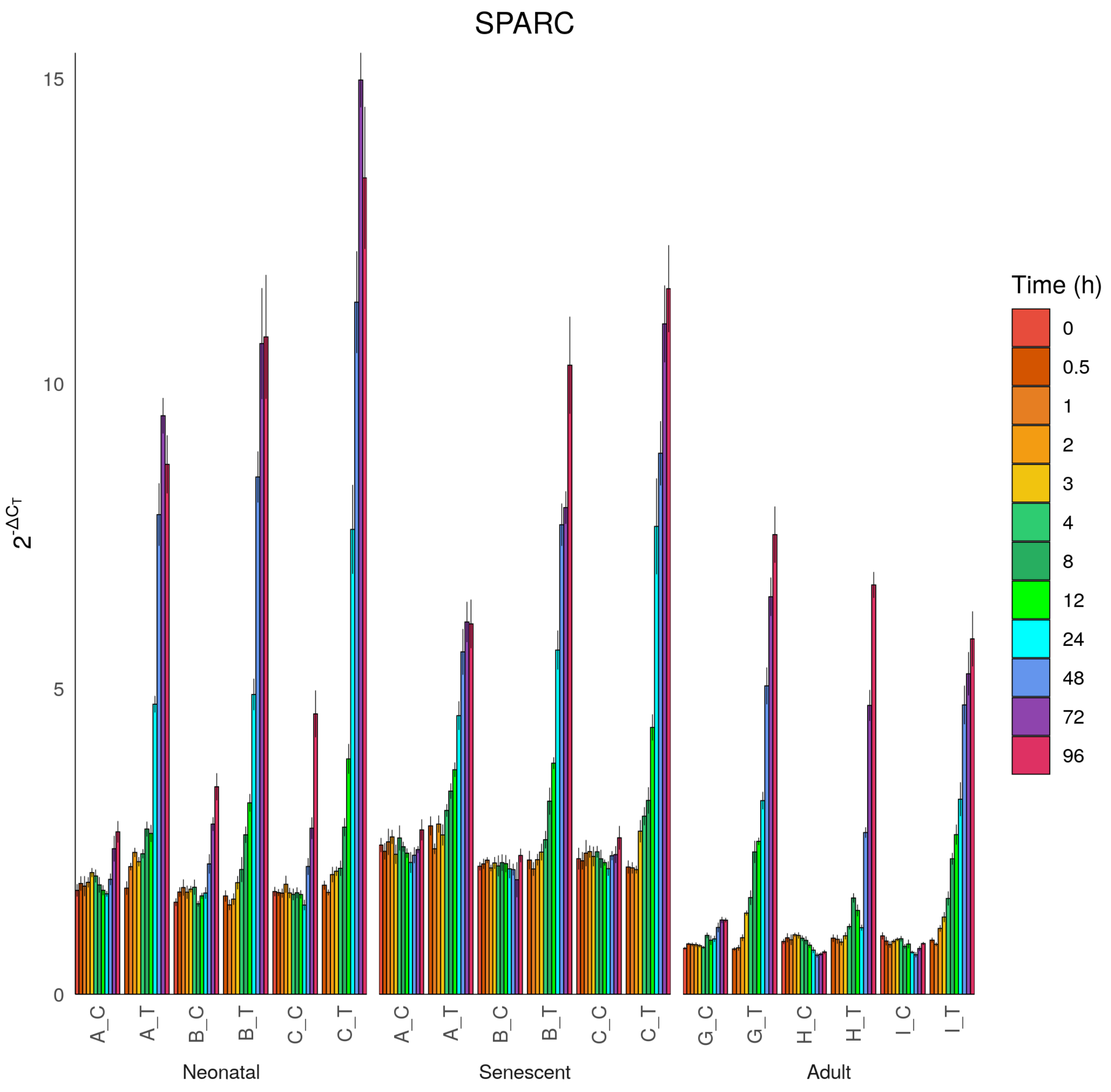

SPARC
$2^{-\Delta C_T}$
0
5
10
15
A_C
A_T
B_C
B_T
C_C
C_T
G_C
G_T
H_C
H_T
I_C
I_T
Neonatal
Senescent
Adult
Time (h)
0
0.5
1
2
3
4
8
12
24
48
72
96

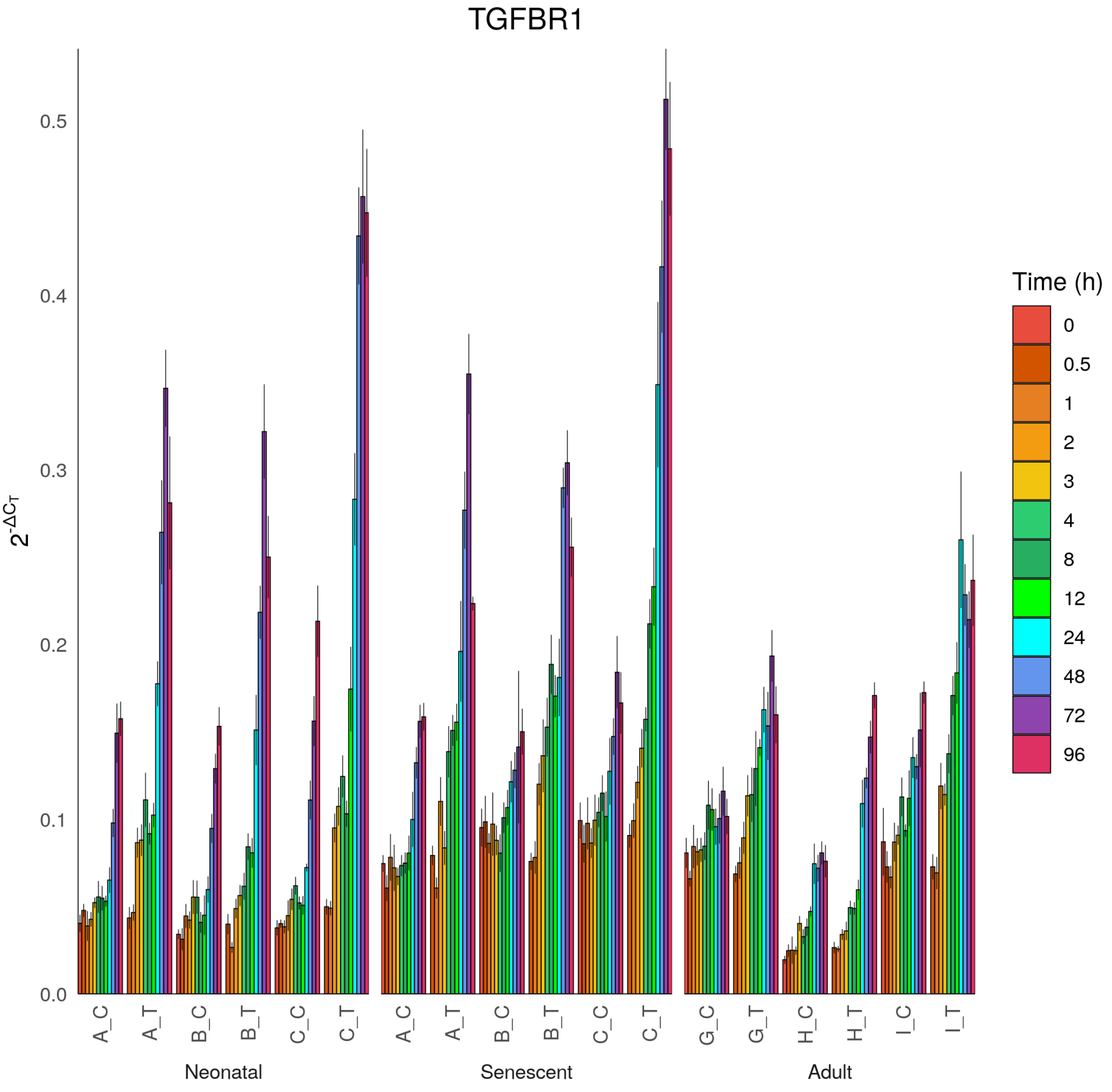

TGFBR1
$2^{-\Delta C_T}$
0.0
0.1
0.2
0.3
0.4
0.5
A_C
A_T
B_C
B_T
C_C
C_T
G_C
G_T
H_C
H_T
I_C
I_T
Neonatal
Senescent
Adult
Time (h)
0
0.5
1
2
3
4
8
12
24
48
72
96

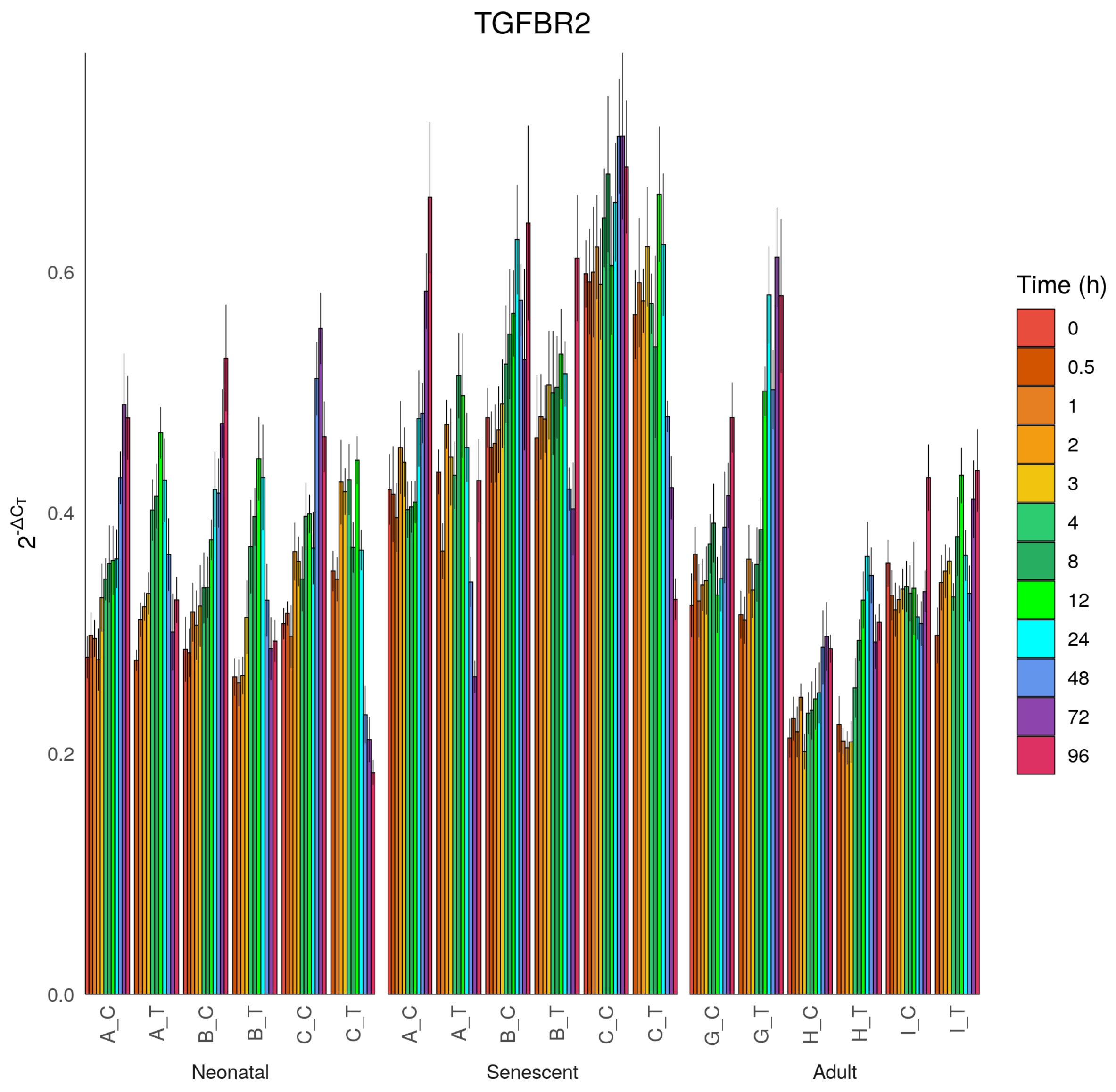

TGFBR2
$2^{-\Delta C_T}$
0.0
0.2
0.4
0.6
A_C
A_T
B_C
B_T
C_C
C_T
A_C
A_T
B_C
B_T
C_C
C_T
G_C
G_T
H_C
H_T
I_C
I_T
Neonatal
Senescent
Adult
Time (h)
0
0.5
1
2
3
4
8
12
24
48
72
96

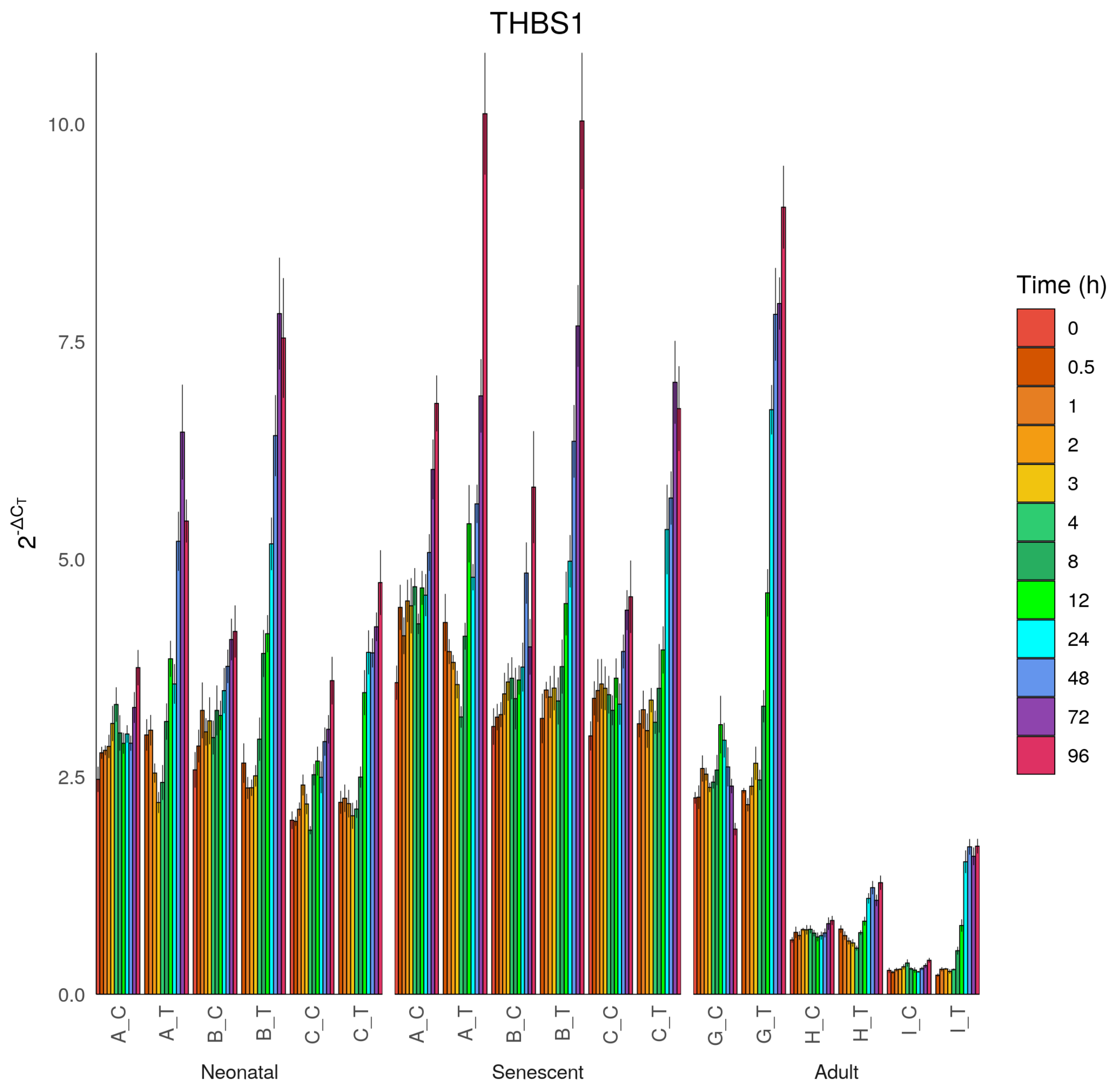
THBS1
$2^{-\Delta C_T}$
0.0
2.5
5.0
7.5
10.0
A_C
A_T
B_C
B_T
C_C
C_T
A_C
A_T
B_C
B_T
C_C
C_T
G_C
G_T
H_C
H_T
I_C
I_T
Neonatal
Senescent
Adult
Time (h)
0
0.5
1
2
3
4
8
12
24
48
72
96

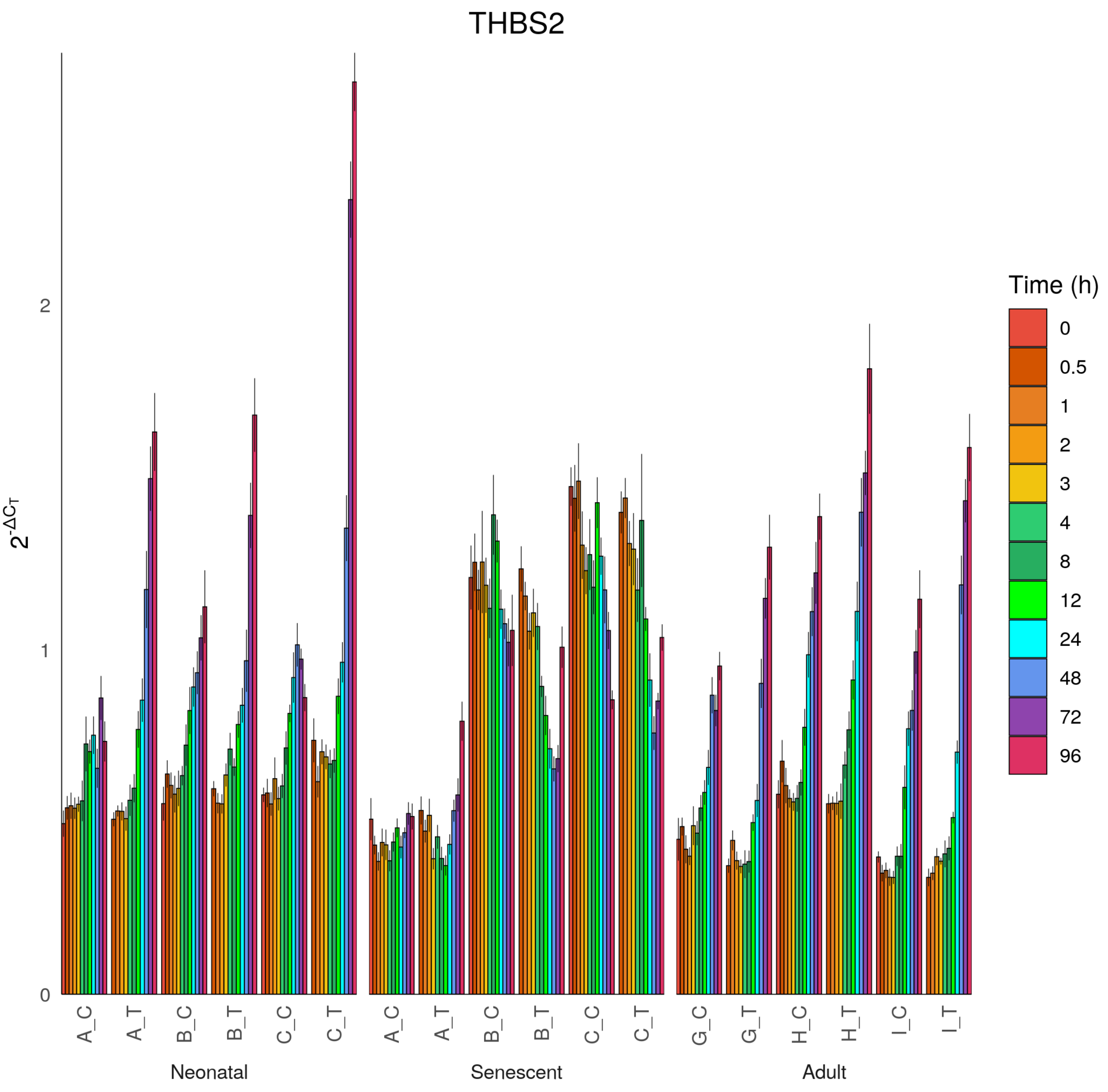
THBS2
$2^{-\Delta C_T}$
0
1
2
A_C
A_T
B_C
B_T
C_C
C_T
Neonatal
A_C
A_T
B_C
B_T
C_C
C_T
Senescent
G_C
G_T
H_C
H_T
I_C
I_T
Adult
Time (h)
0
0.5
1
2
3
4
8
12
24
48
72
96

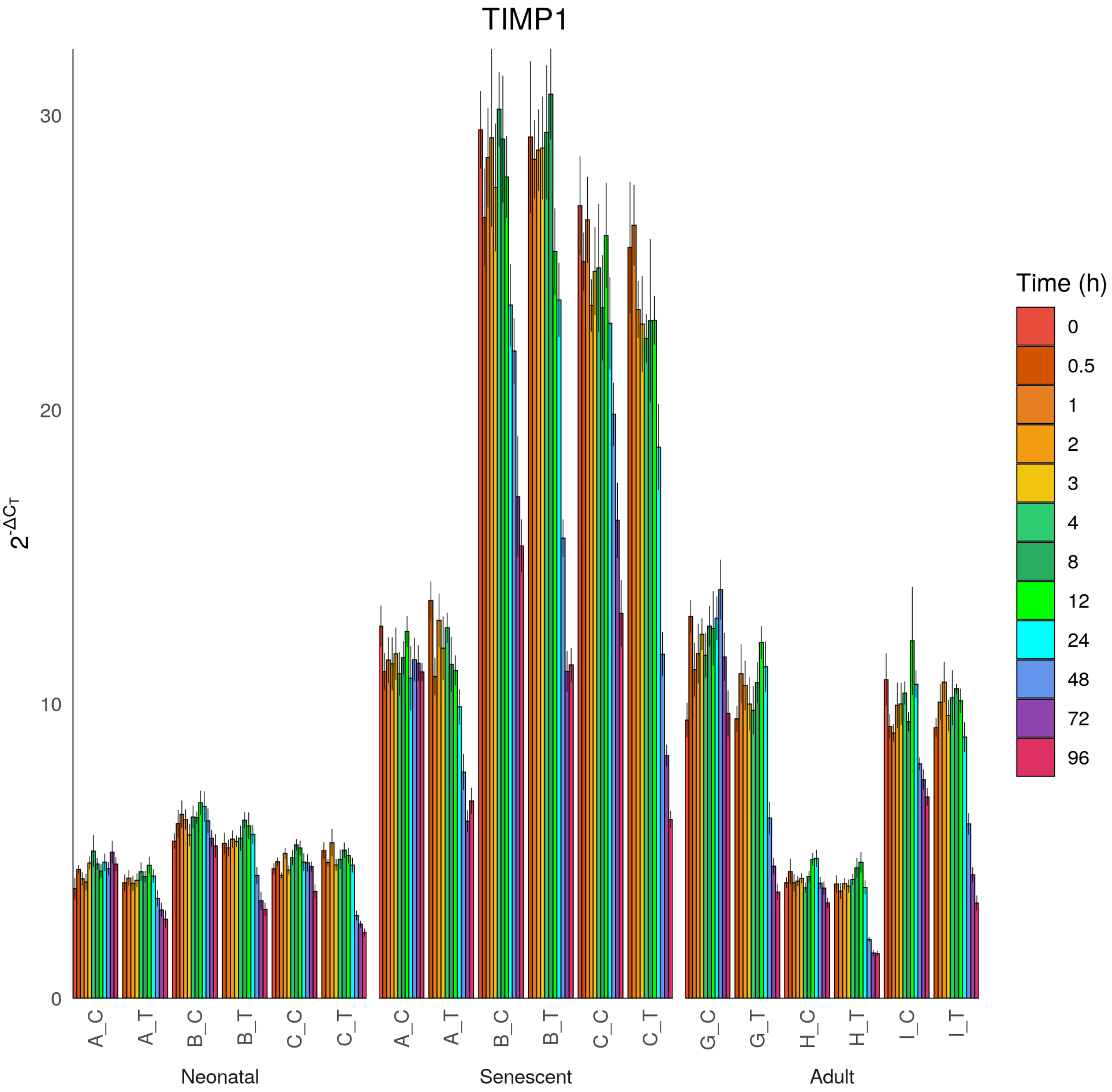
TIMP1
$2^{-\Delta C_T}$
0
10
20
30
A_C
A_T
B_C
B_T
C_C
C_T
Neonatal
A_C
A_T
B_C
B_T
C_C
C_T
Senescent
G_C
G_T
H_C
H_T
I_C
I_T
Adult
Time (h)
0
0.5
1
2
3
4
8
12
24
48
72
96

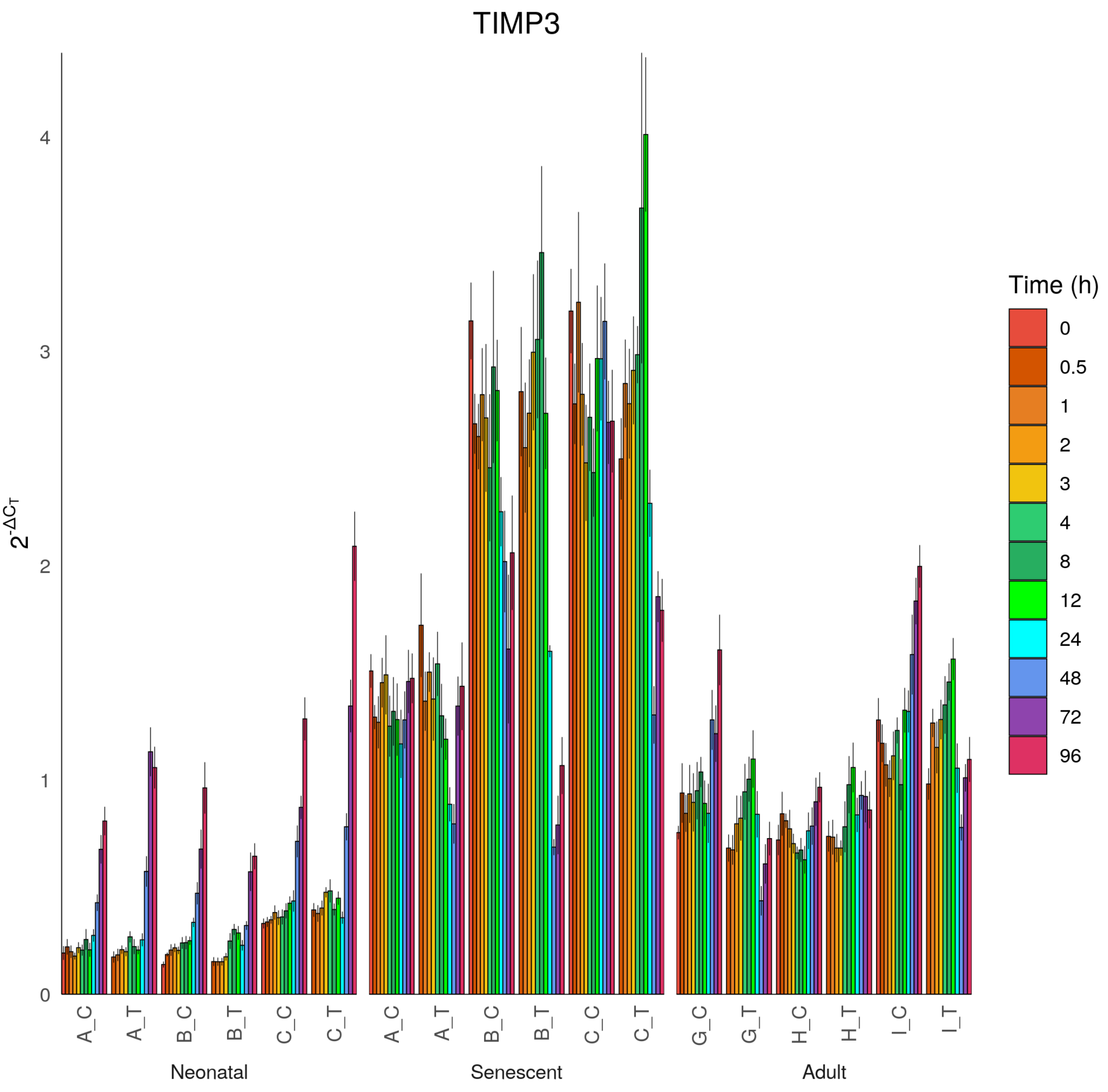

TIMP3
$2^{-\Delta C_T}$
0
1
2
3
4
A_C
A_T
B_C
B_T
C_C
C_T
G_C
G_T
H_C
H_T
I_C
I_T
Neonatal
Senescent
Adult
Time (h)
0
0.5
1
2
3
4
8
12
24
48
72
96

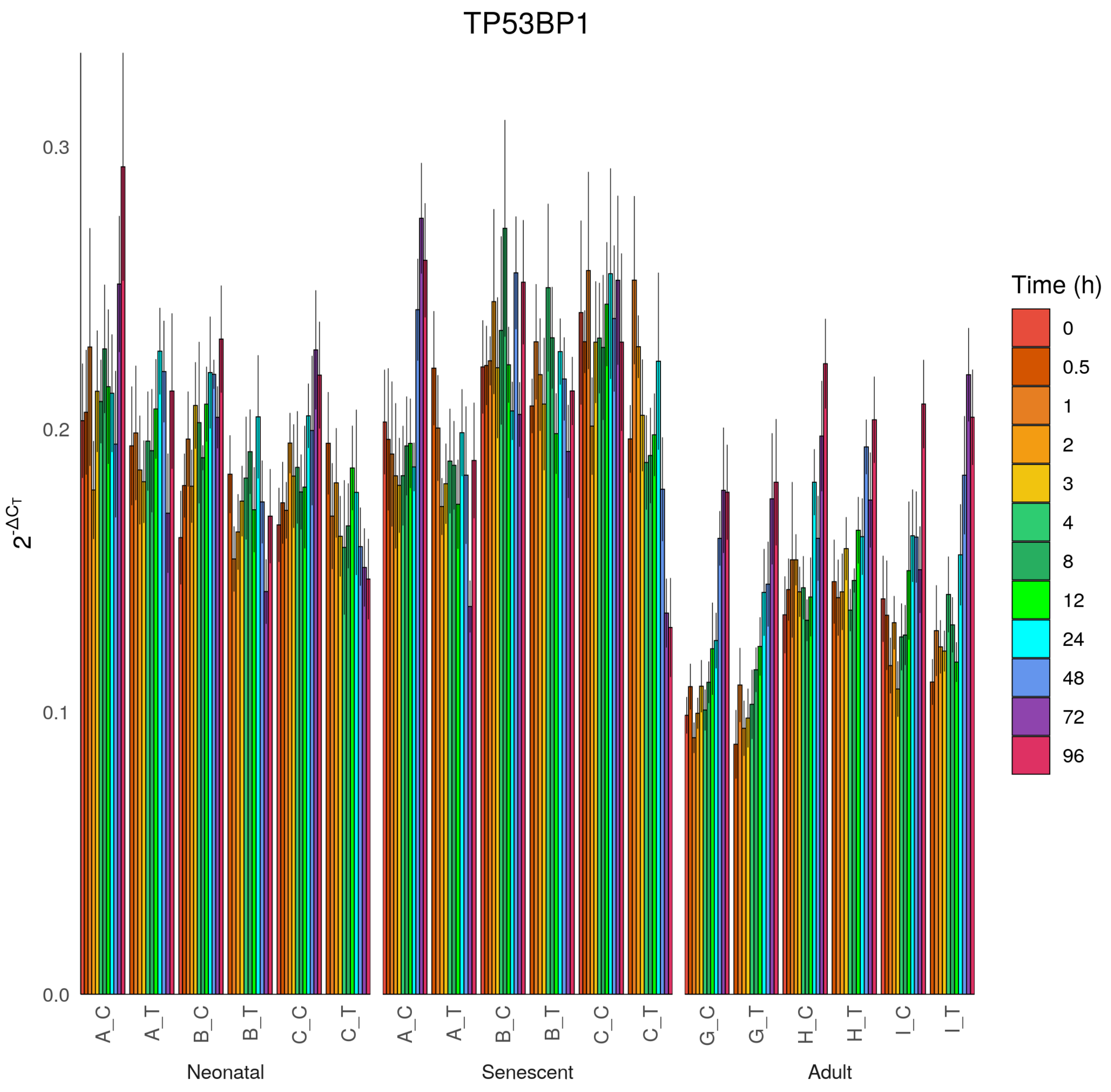
TP53BP1
2^-ΔCT
0.0
0.1
0.2
0.3
A_C
A_T
B_C
B_T
C_C
C_T
Neonatal
A_C
A_T
B_C
B_T
C_C
C_T
Senescent
G_C
G_T
H_C
H_T
I_C
I_T
Adult
Time (h)
0
0.5
1
2
3
4
8
12
24
48
72
96

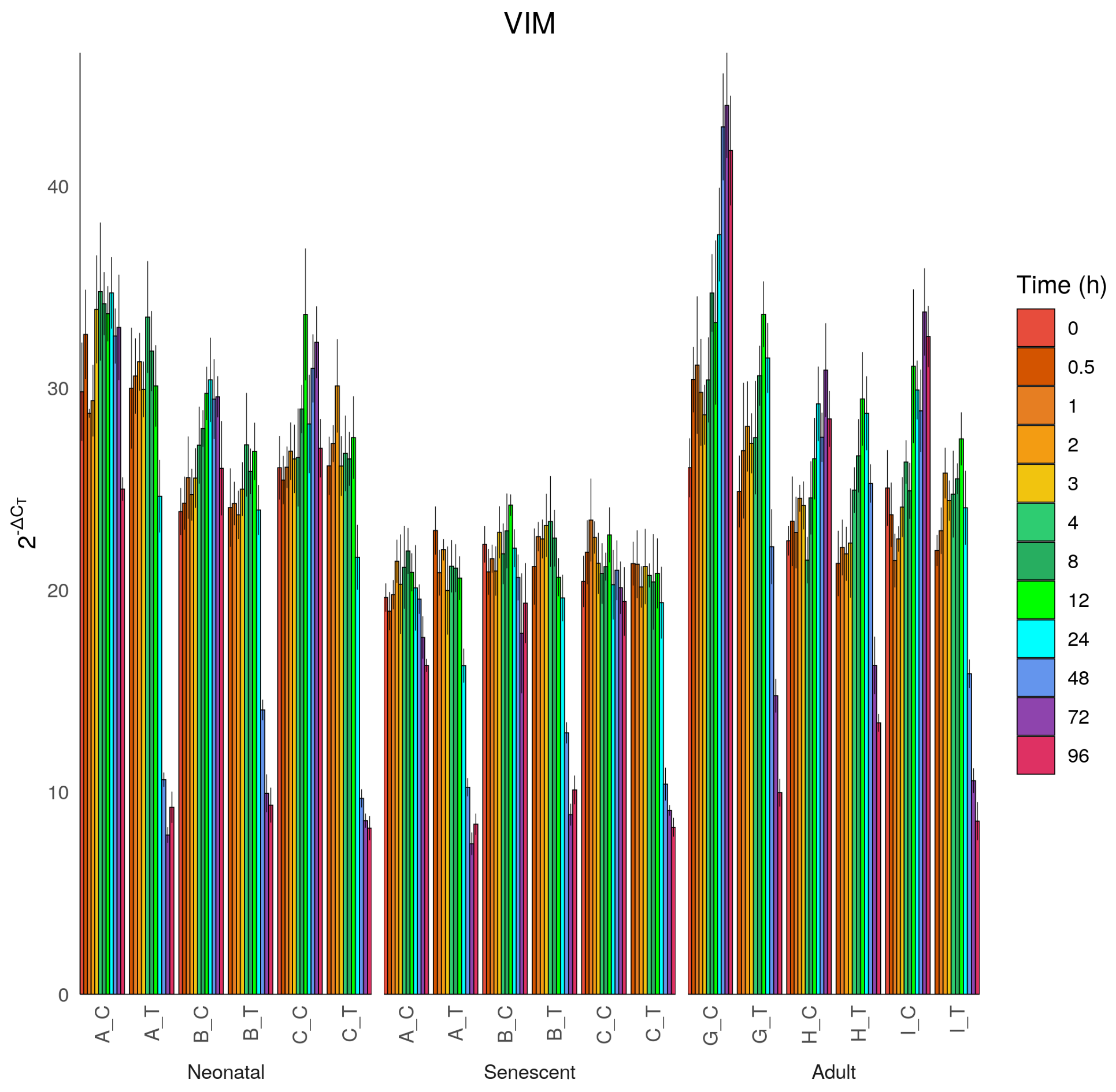

VIM
$2^{-\Delta C_T}$
0
10
20
30
40
A_C
A_T
B_C
B_T
C_C
C_T
A_C
A_T
B_C
B_T
C_C
C_T
G_C
G_T
H_C
H_T
I_C
I_T
Neonatal
Senescent
Adult
Time (h)
0
0.5
1
2
3
4
8
12
24
48
72
96
