## Supplementary file 3 for "Cell Senescence-Independent Ageing of Human Skin"

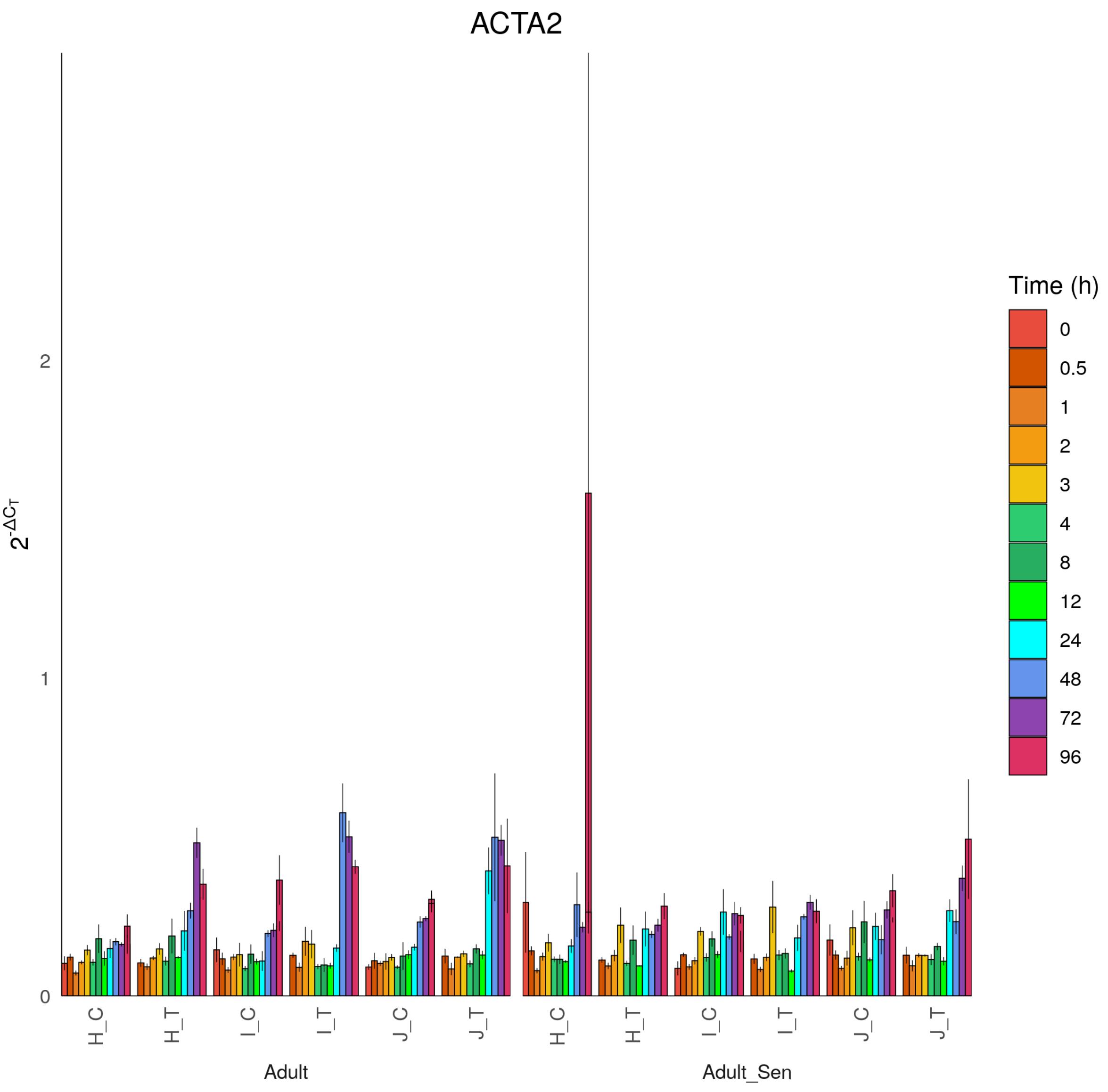

ACTA2
$2^{-\Delta C_T}$
0
1
2
H_C
H_T
I_C
I_T
J_C
J_T
H_C
H_T
I_C
I_T
J_C
J_T
Adult
Adult_Sen
Time (h)
0
0.5
1
2
3
4
8
12
24
48
72
96

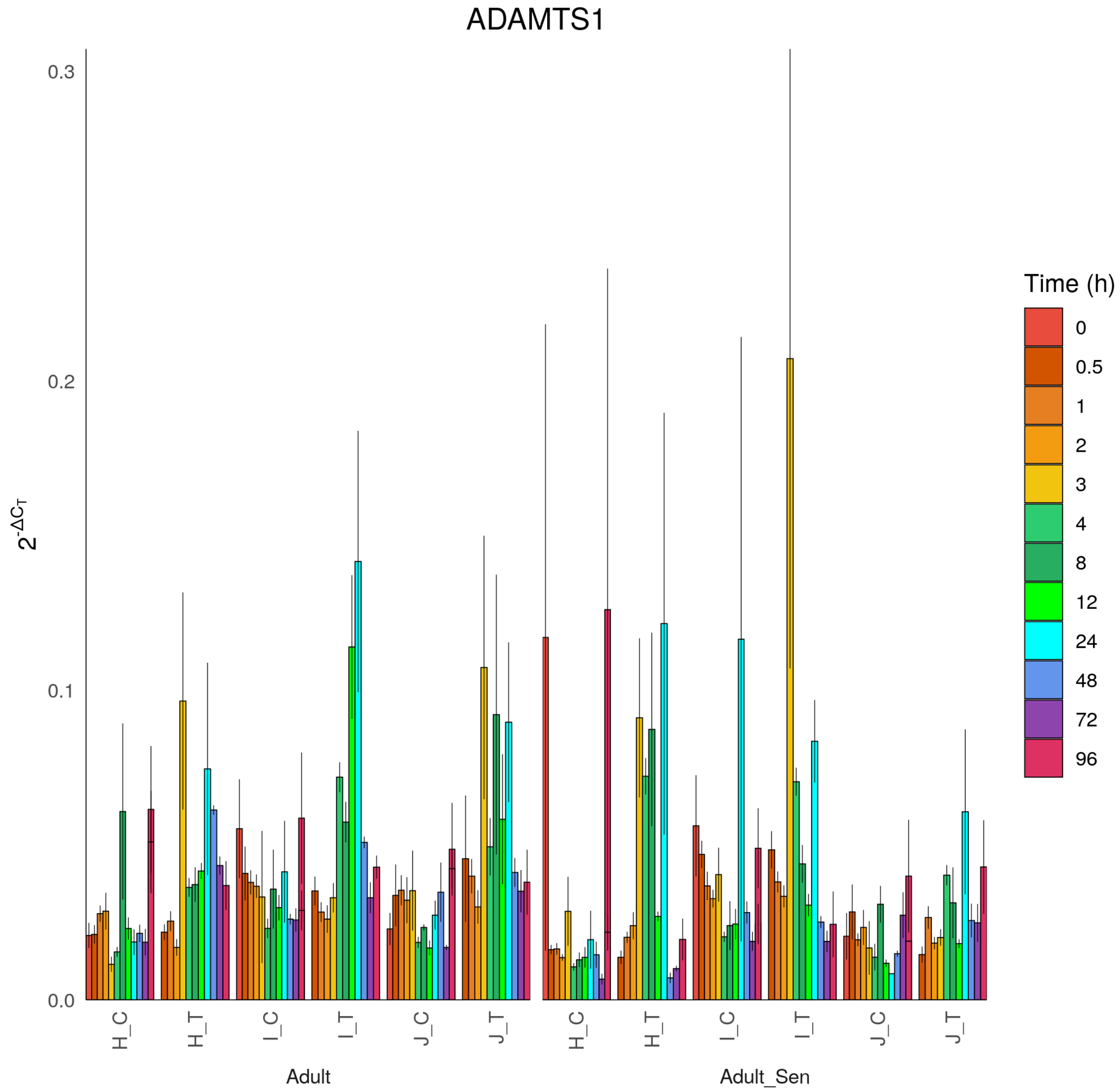
ADAMTS1
$2^{-\Delta C_T}$
0.3
0.2
0.1
0.0
H_C
H_T
I_C
I_T
J_C
J_T
H_C
H_T
I_C
I_T
J_C
J_T
Adult
Adult_Sen
Time (h)
0
0.5
1
2
3
4
8
12
24
48
72
96

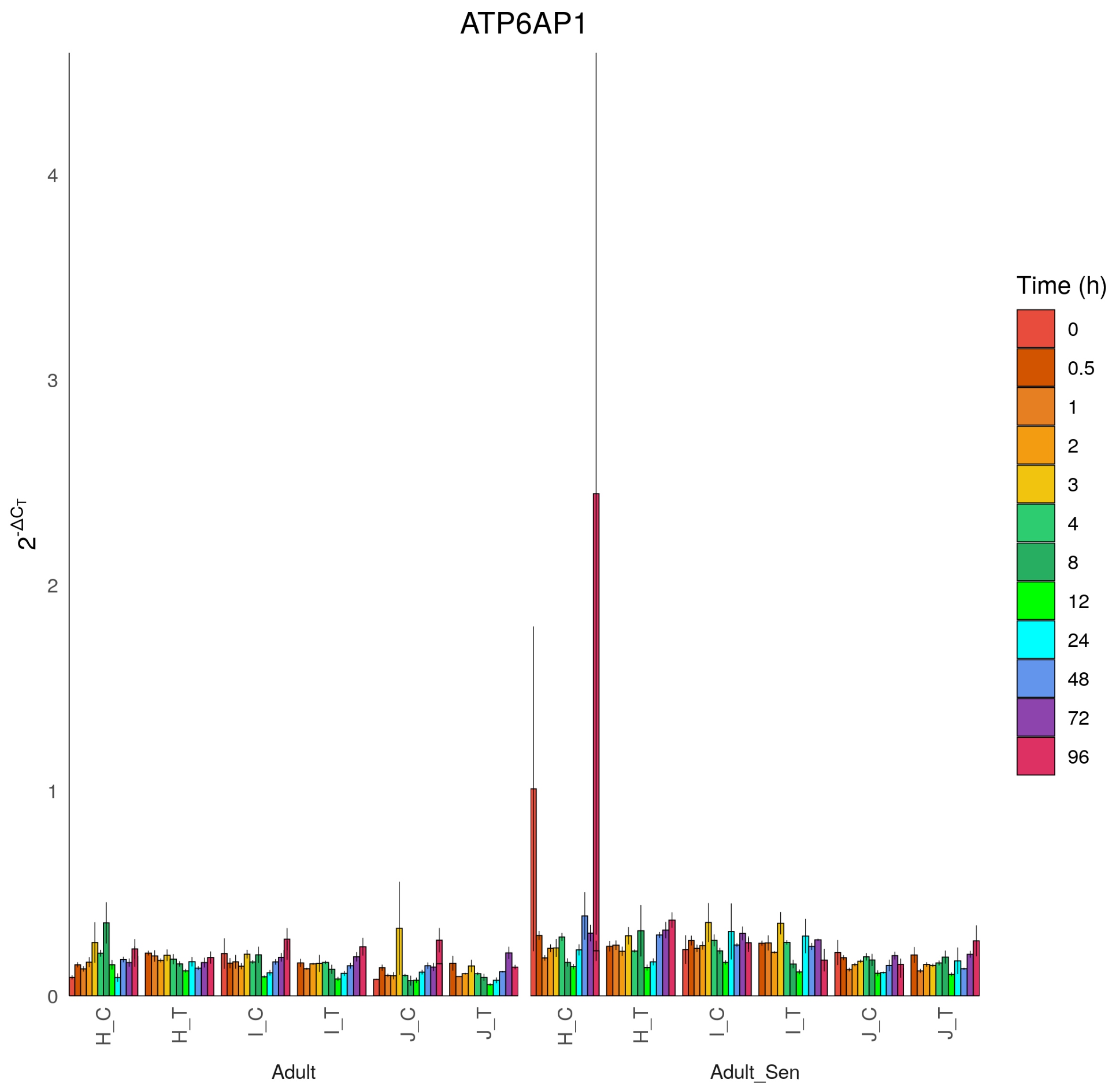
ATP6AP1
$2^{-\Delta C_T}$
0
1
2
3
4
H_C
H_T
I_C
I_T
J_C
J_T
H_C
H_T
I_C
I_T
J_C
J_T
Adult
Adult_Sen
Time (h)
0
0.5
1
2
3
4
8
12
24
48
72
96

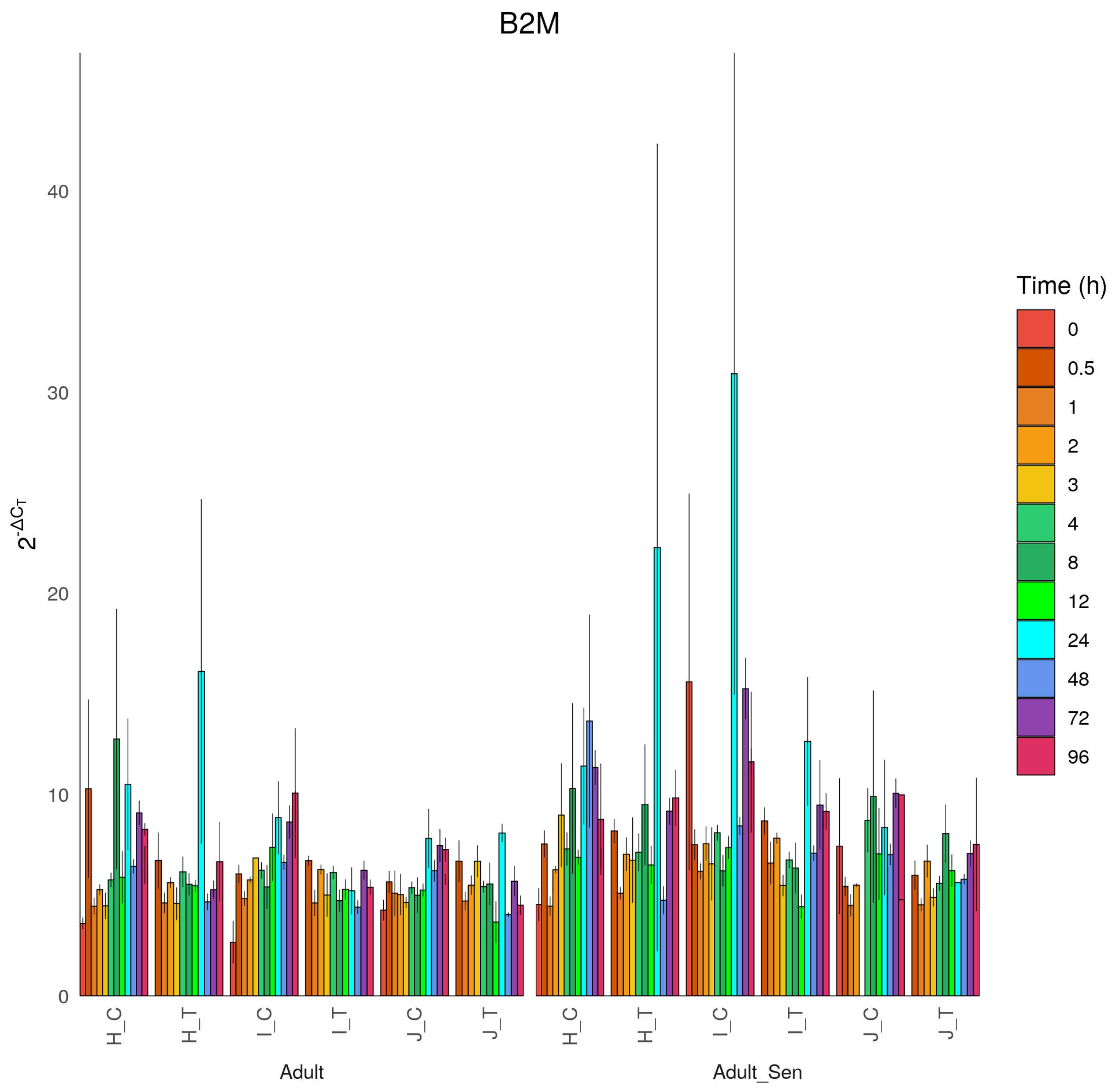

B2M
$2^{-\Delta C_T}$
0
10
20
30
40
H_C
H_T
I_C
I_T
J_C
J_T
Adult
Adult_Sen
Time (h)
0
0.5
1
2
3
4
8
12
24
48
72
96

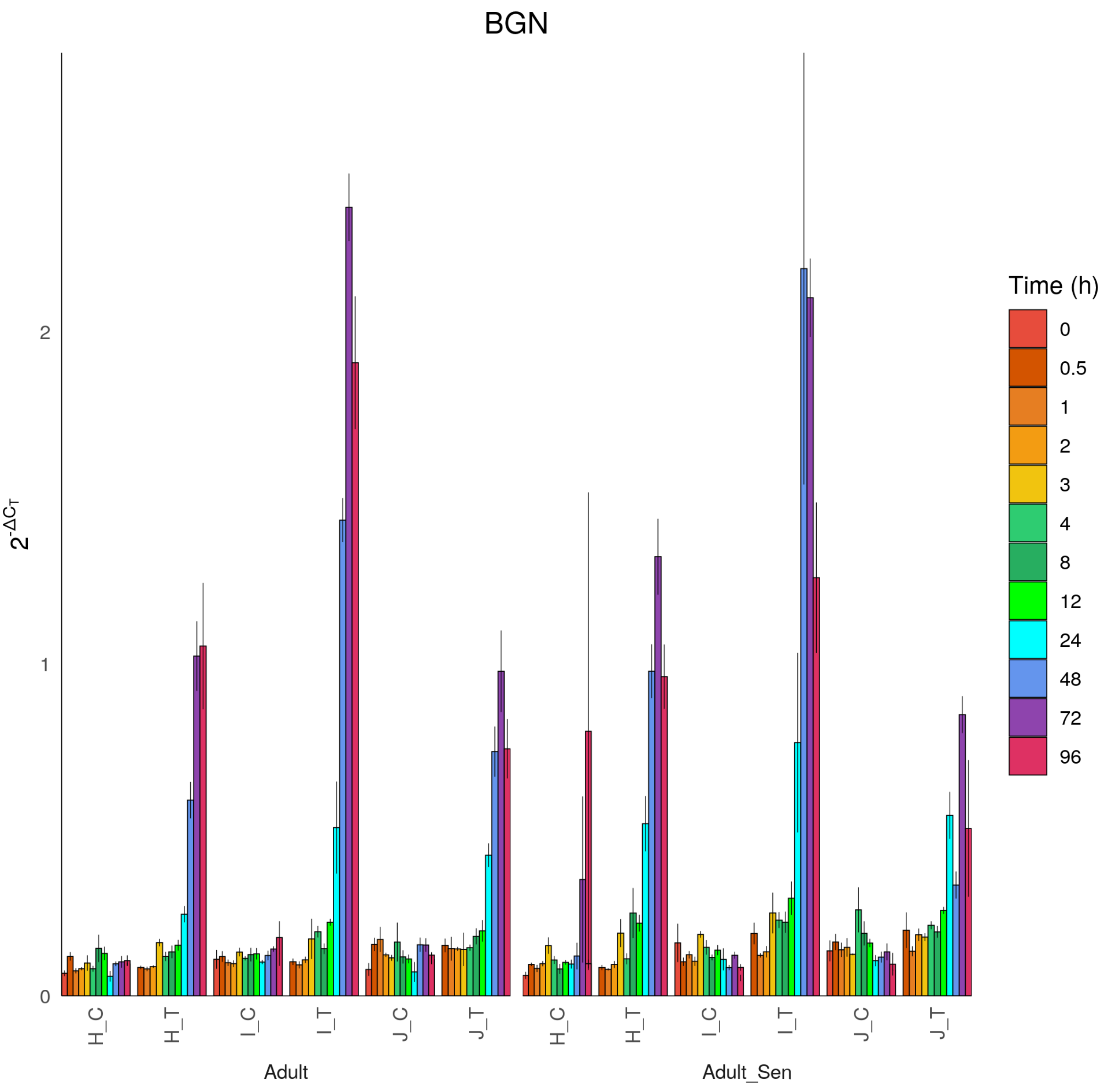

BGN
$2^{-\Delta C_T}$
0
1
2
H_C
H_T
I_C
I_T
J_C
J_T
Adult
Adult_Sen
Time (h)
0
0.5
1
2
3
4
8
12
24
48
72
96

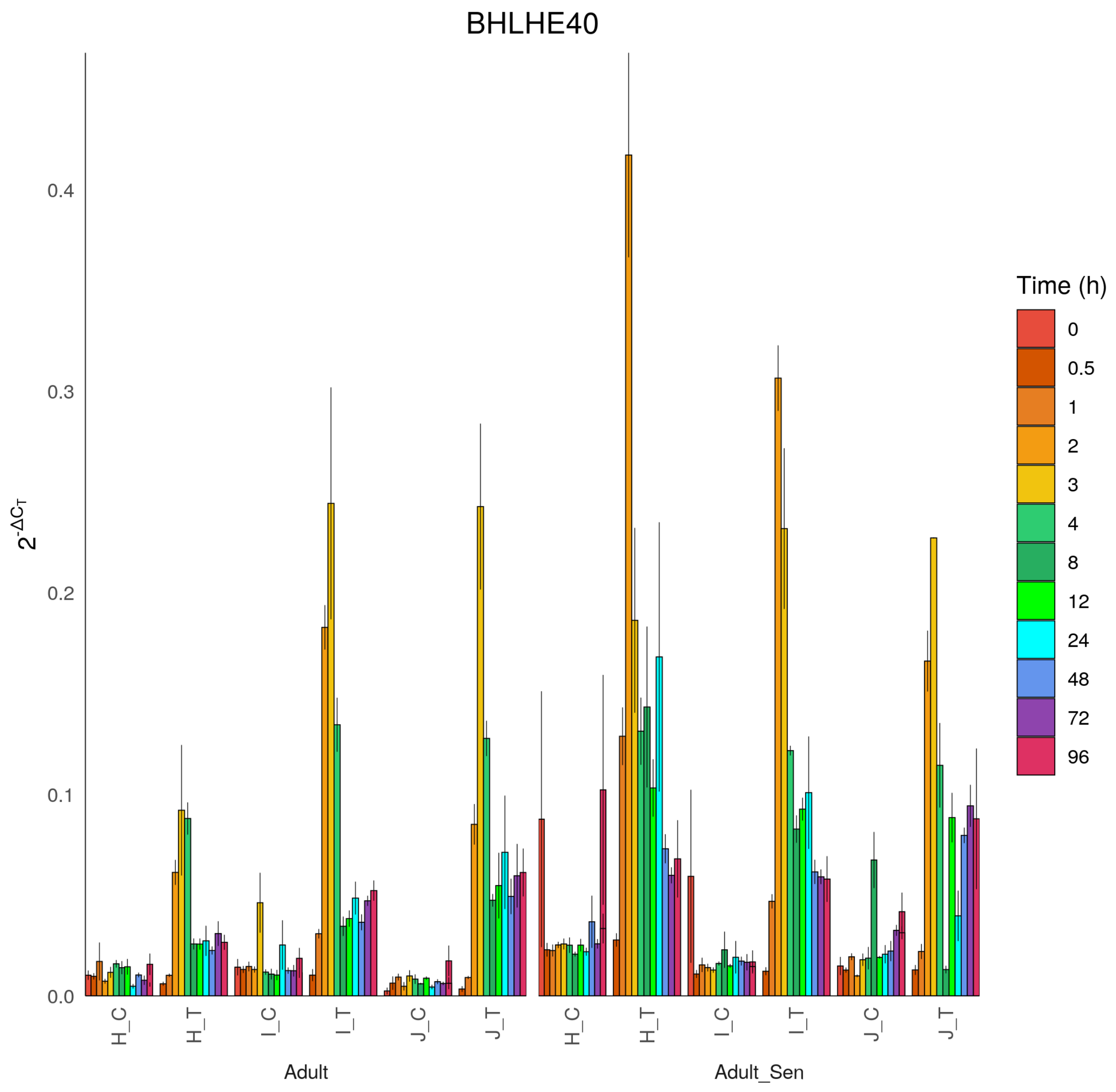

BHLHE40
$2^{-\Delta C_T}$
0.0
0.1
0.2
0.3
0.4
H_C
H_T
I_C
I_T
J_C
J_T
Adult
Adult_Sen
Time (h)
0
0.5
1
2
3
4
8
12
24
48
72
96

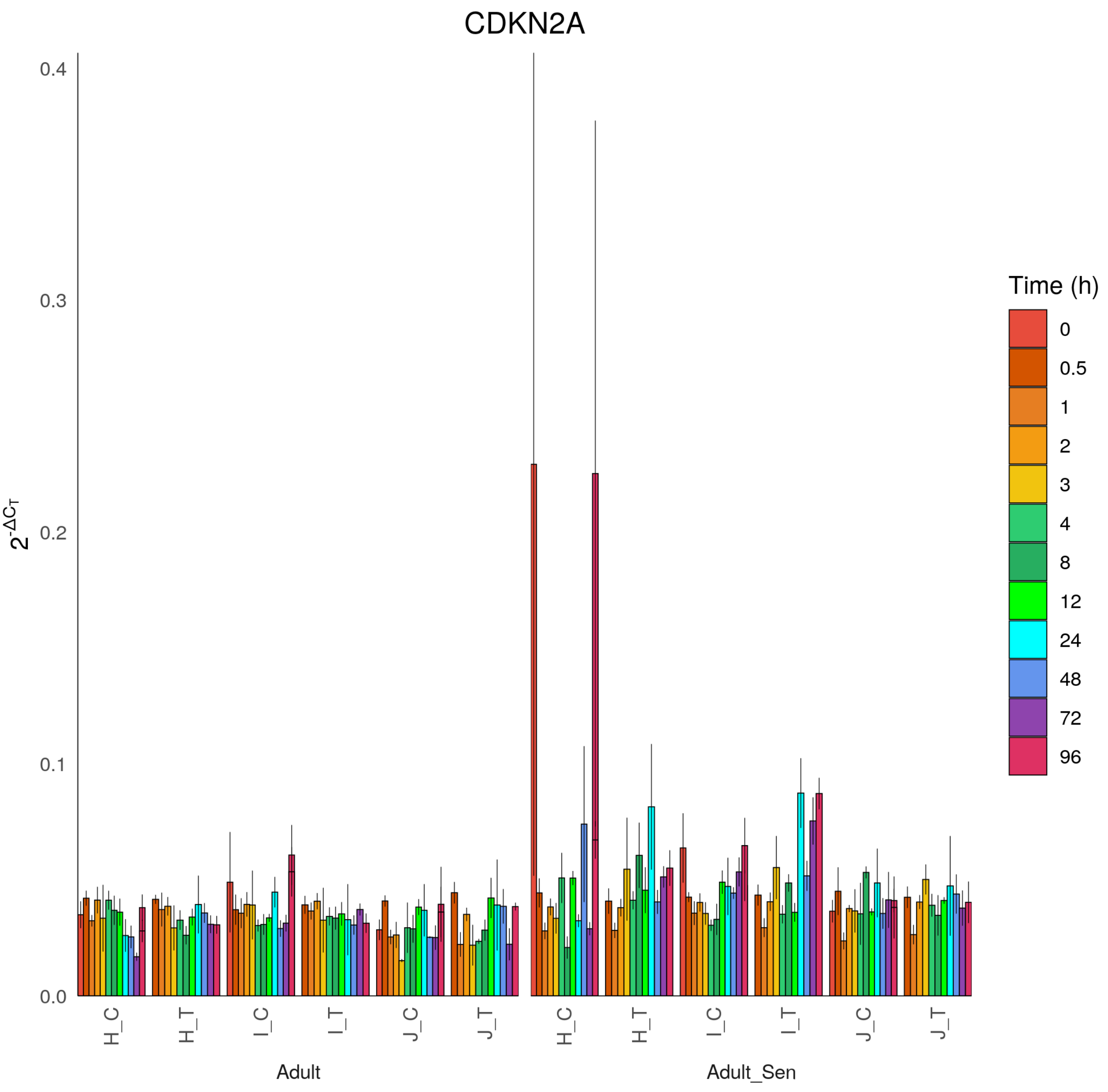

CDKN2A
$2^{-\Delta C_T}$
0.4
0.3
0.2
0.1
0.0
H_C
H_T
I_C
I_T
J_C
J_T
Adult
Adult_Sen
Time (h)
0
0.5
1
2
3
4
8
12
24
48
72
96

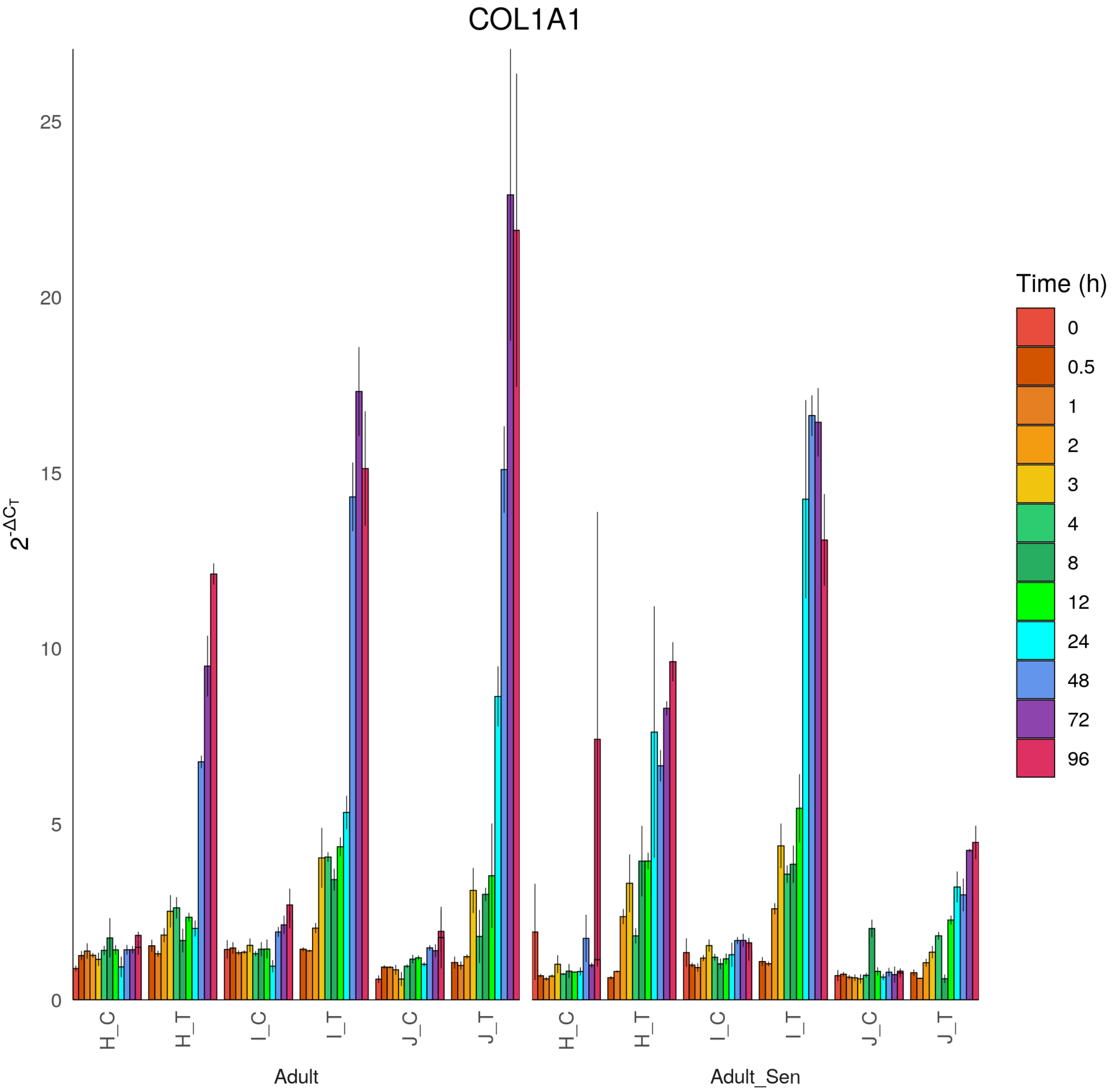
COL1A1
$2^{-\Delta C_T}$
0
5
10
15
20
25
H_C
H_T
I_C
I_T
J_C
J_T
Adult
Adult_Sen
Time (h)
0
0.5
1
2
3
4
8
12
24
48
72
96

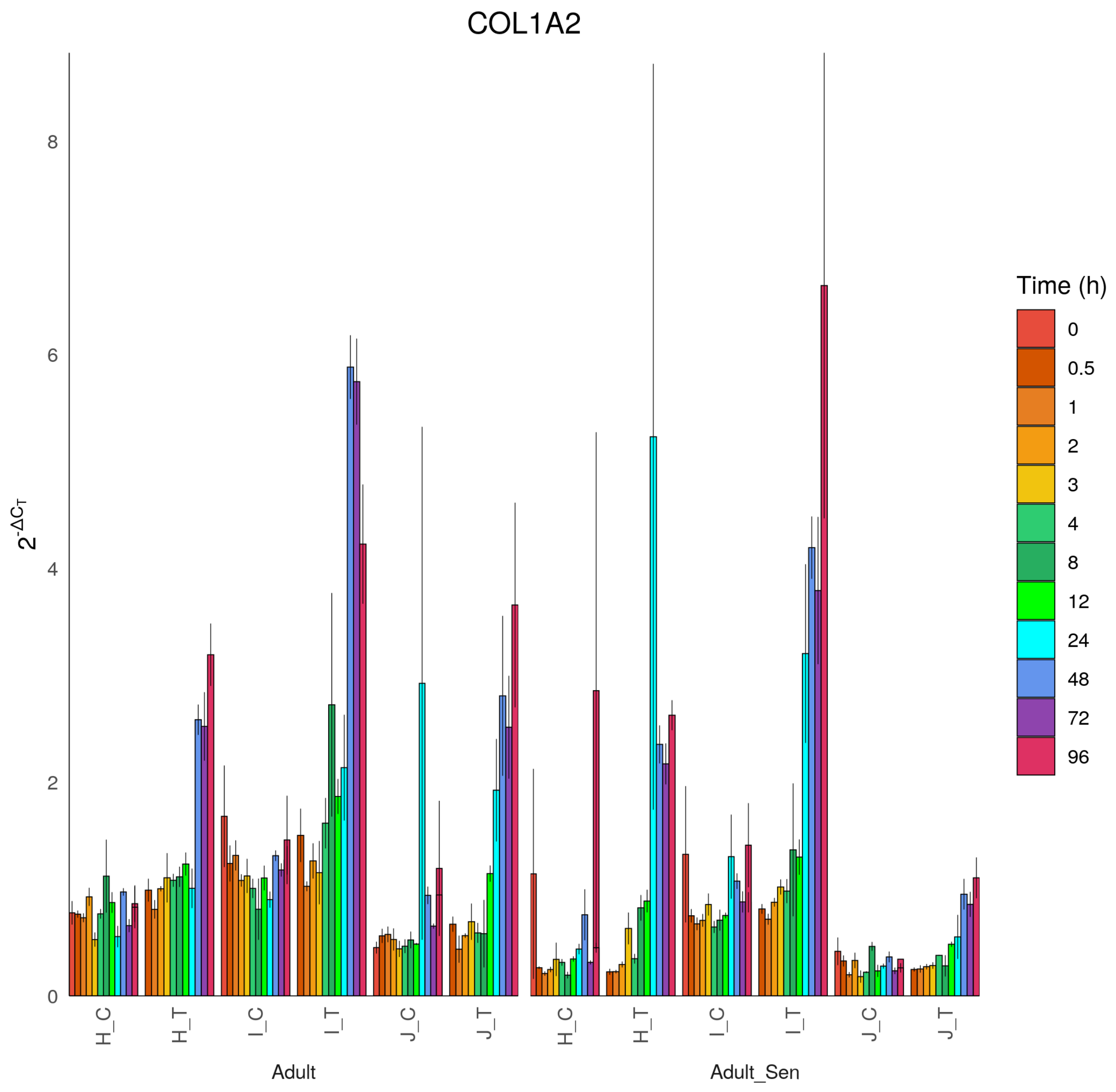

COL1A2
$2^{-\Delta C_T}$
0
2
4
6
8
H_C
H_T
I_C
I_T
J_C
J_T
Adult
Adult_Sen
Time (h)
0
0.5
1
2
3
4
8
12
24
48
72
96

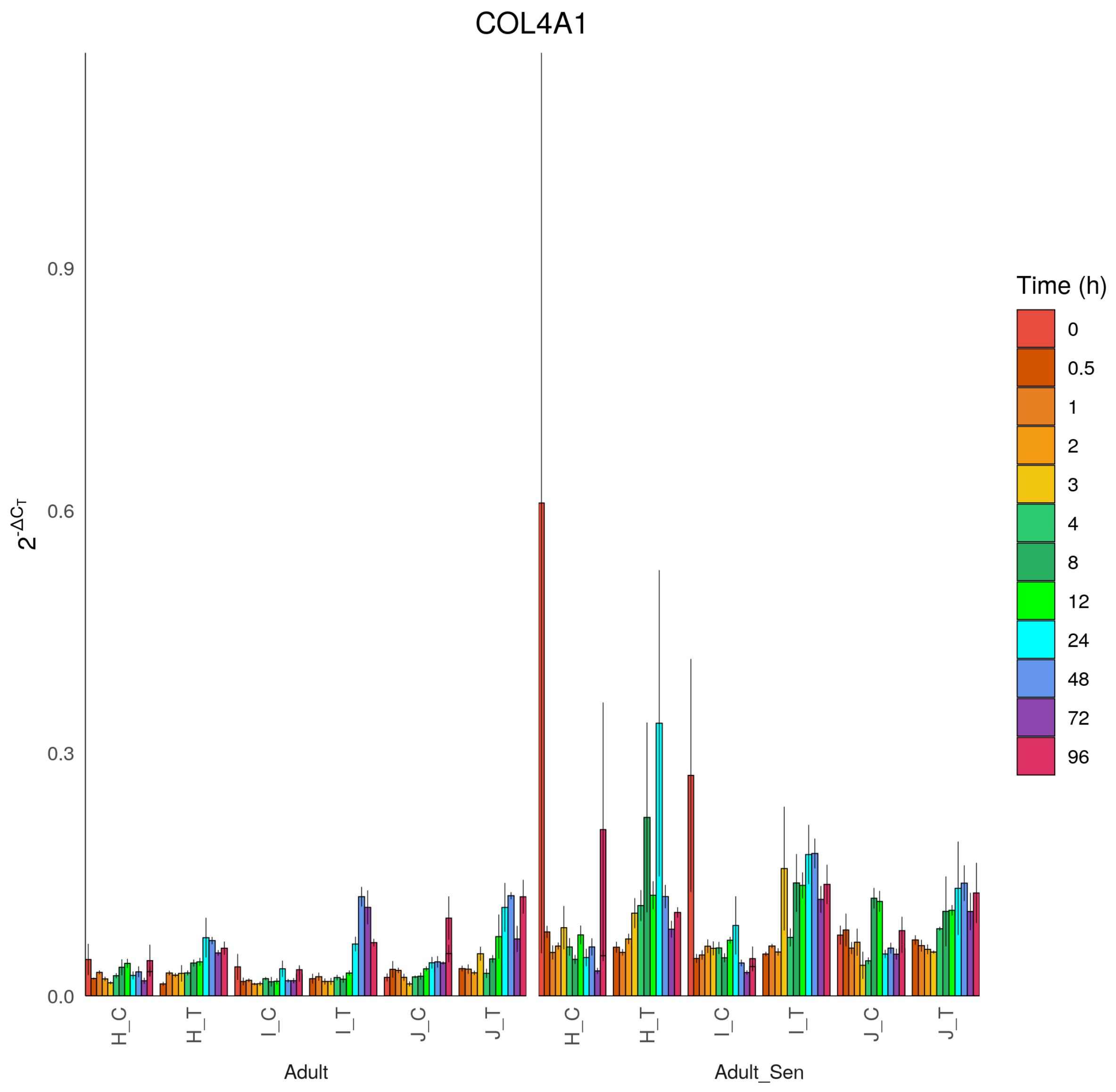

COL4A1
$2^{-\Delta C_T}$
0.0
0.3
0.6
0.9
H_C
H_T
I_C
I_T
J_C
J_T
Adult
Adult_Sen
Time (h)
0
0.5
1
2
3
4
8
12
24
48
72
96

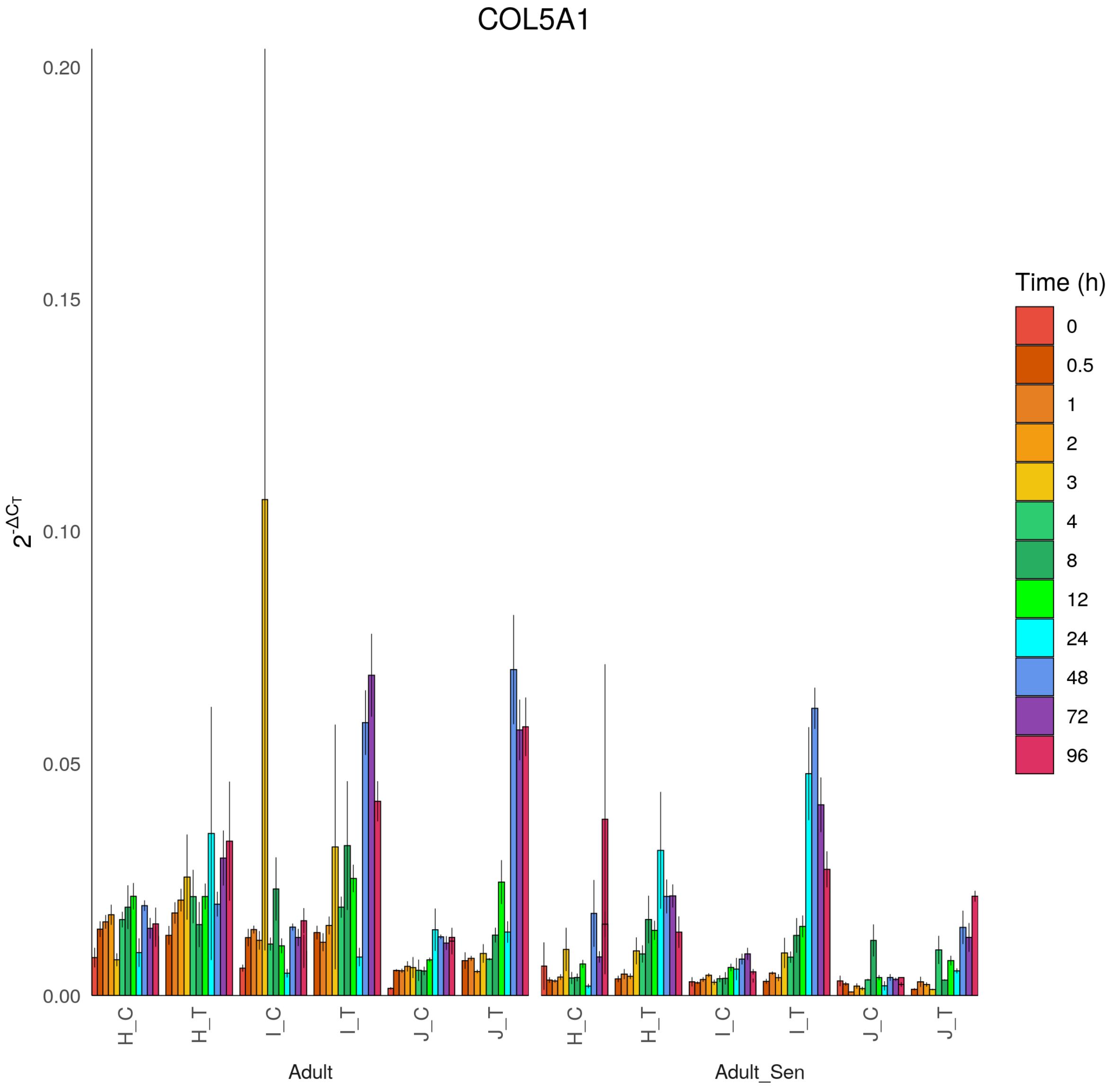

COL5A1
$2^{-\Delta C_T}$
0.20
0.15
0.10
0.05
0.00
H_C
H_T
I_C
I_T
J_C
J_T
Adult
Adult_Sen
Time (h)
0
0.5
1
2
3
4
8
12
24
48
72
96

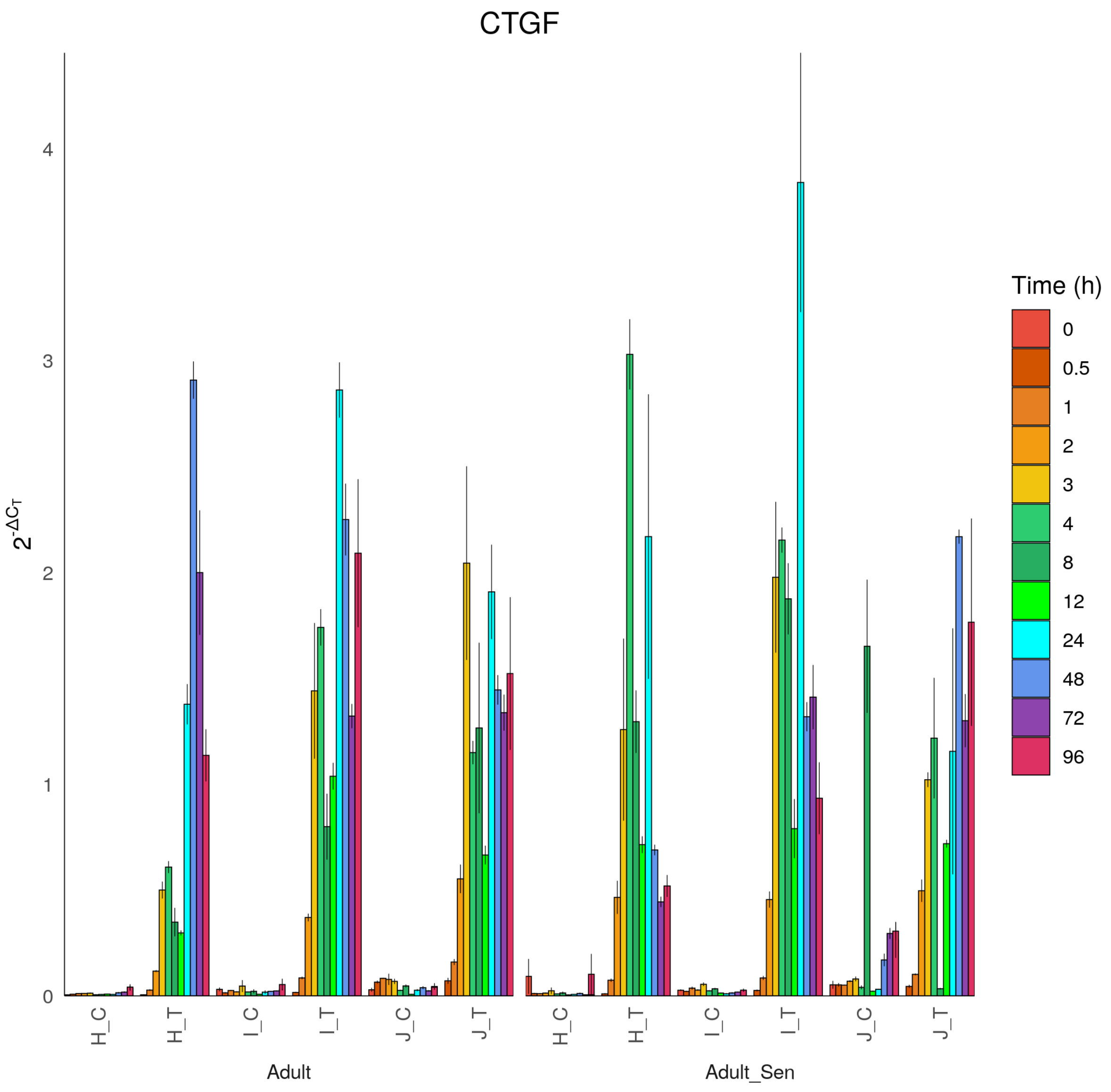

CTGF
$2^{-\Delta C_T}$
0
1
2
3
4
H_C
H_T
I_C
I_T
J_C
J_T
Adult
Adult_Sen
Time (h)
0
0.5
1
2
3
4
8
12
24
48
72
96

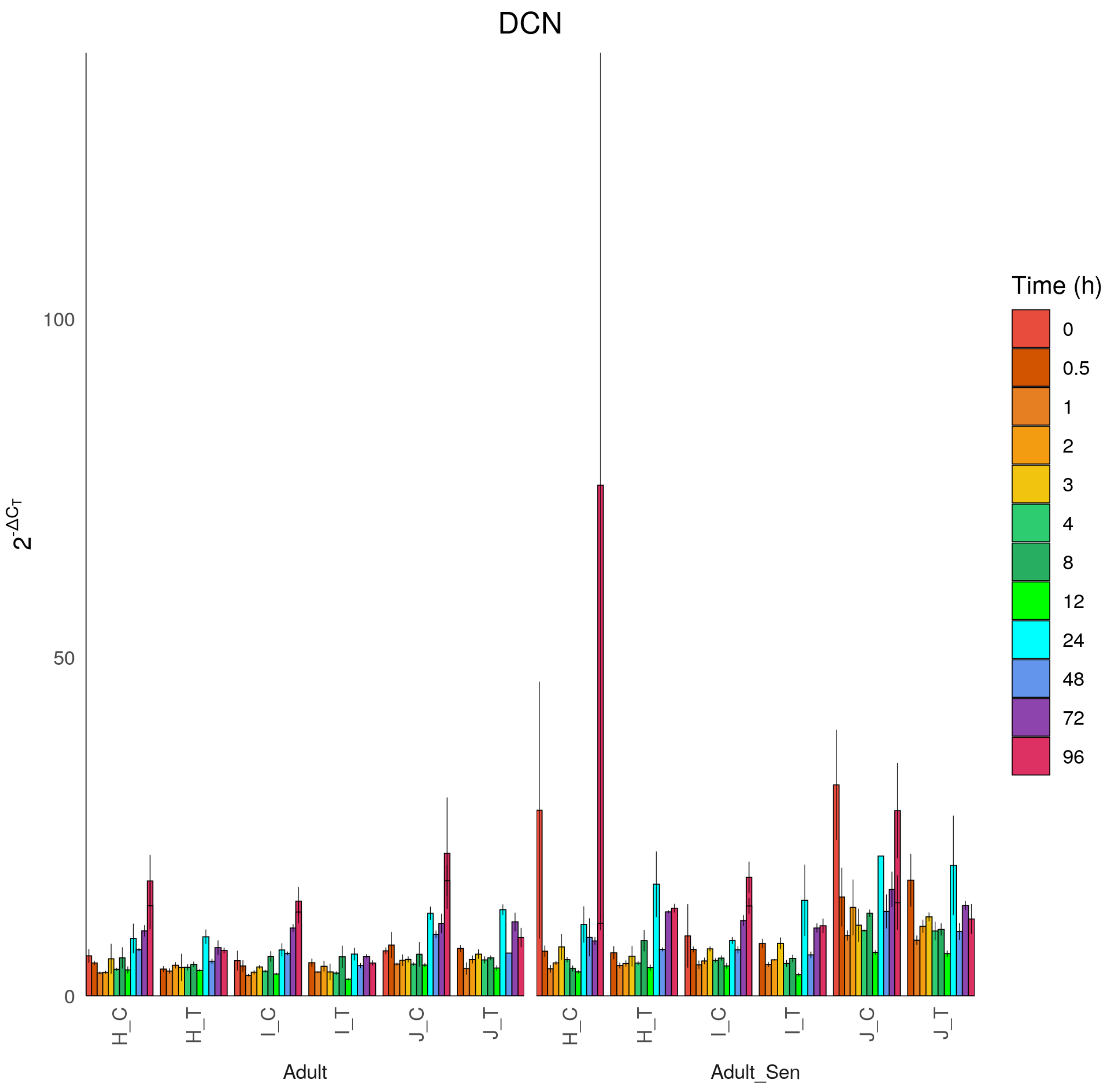
DCN
$2^{-\Delta C_T}$
100
50
0
H_C
H_T
I_C
I_T
J_C
J_T
Adult
Adult_Sen
Time (h)
0
0.5
1
2
3
4
8
12
24
48
72
96

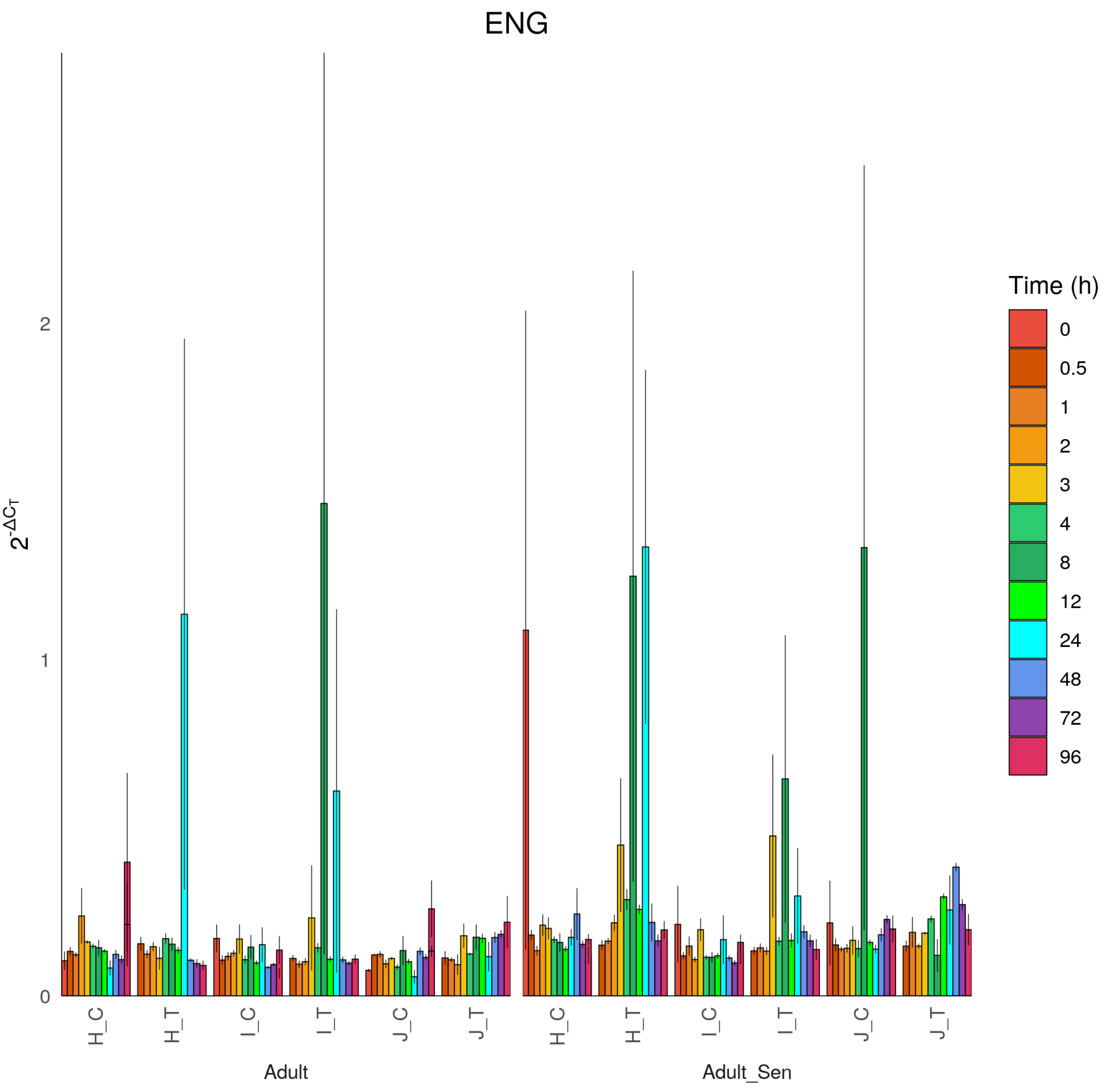
ENG
2
1
0
$2^{-\Delta C_T}$
H_C
H_T
I_C
I_T
J_C
J_T
Adult
Adult_Sen
Time (h)
0
0.5
1
2
3
4
8
12
24
48
72
96

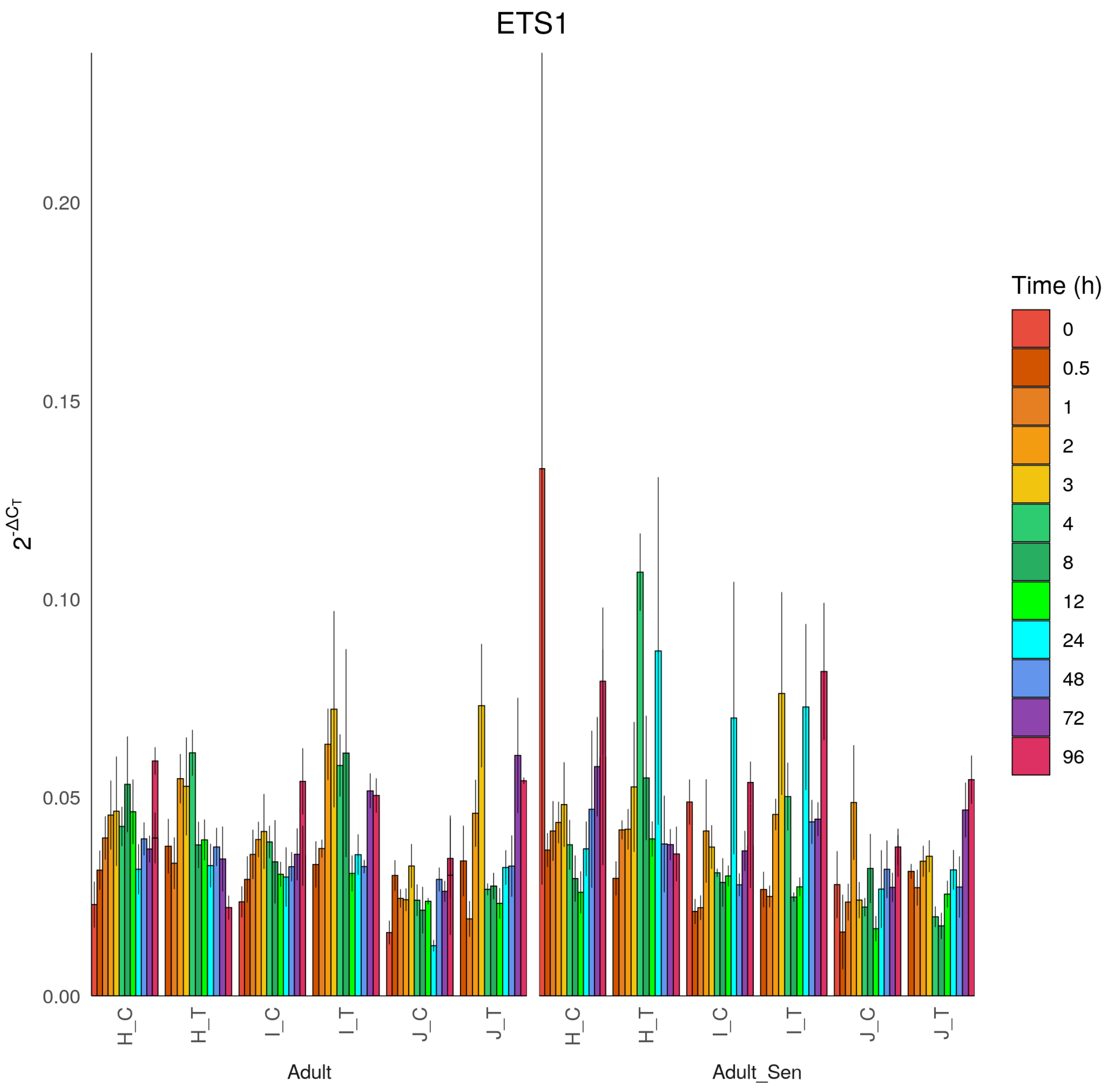
ETS1
2^-ΔCT
0.20
0.15
0.10
0.05
0.00
H_C
H_T
I_C
I_T
J_C
J_T
Adult
Adult_Sen
Time (h)
0
0.5
1
2
3
4
8
12
24
48
72
96

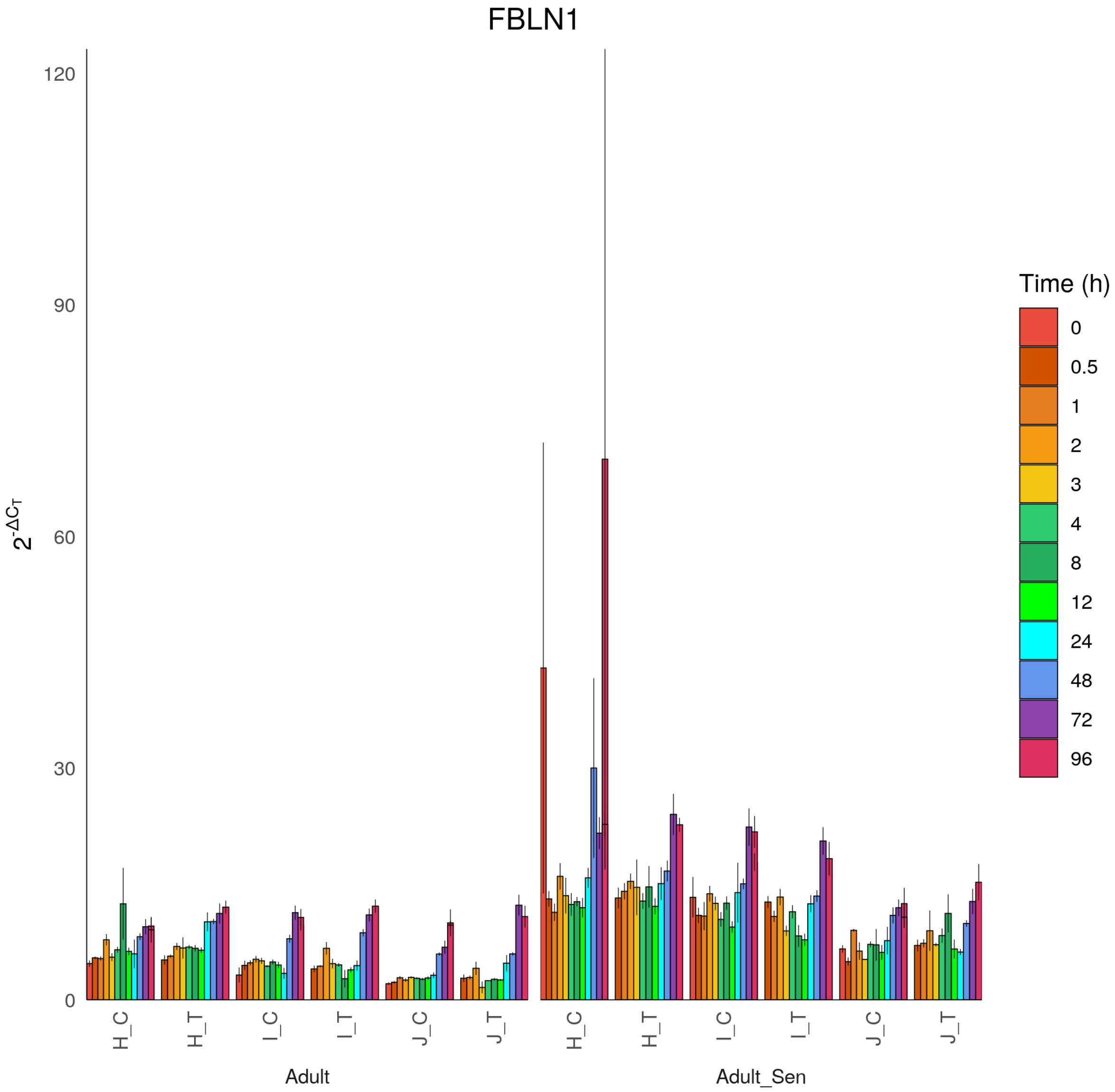

FBLN1
$2^{-\Delta C_T}$
120
90
60
30
0
H_C
H_T
I_C
I_T
J_C
J_T
Adult
Adult_Sen
Time (h)
0
0.5
1
2
3
4
8
12
24
48
72
96

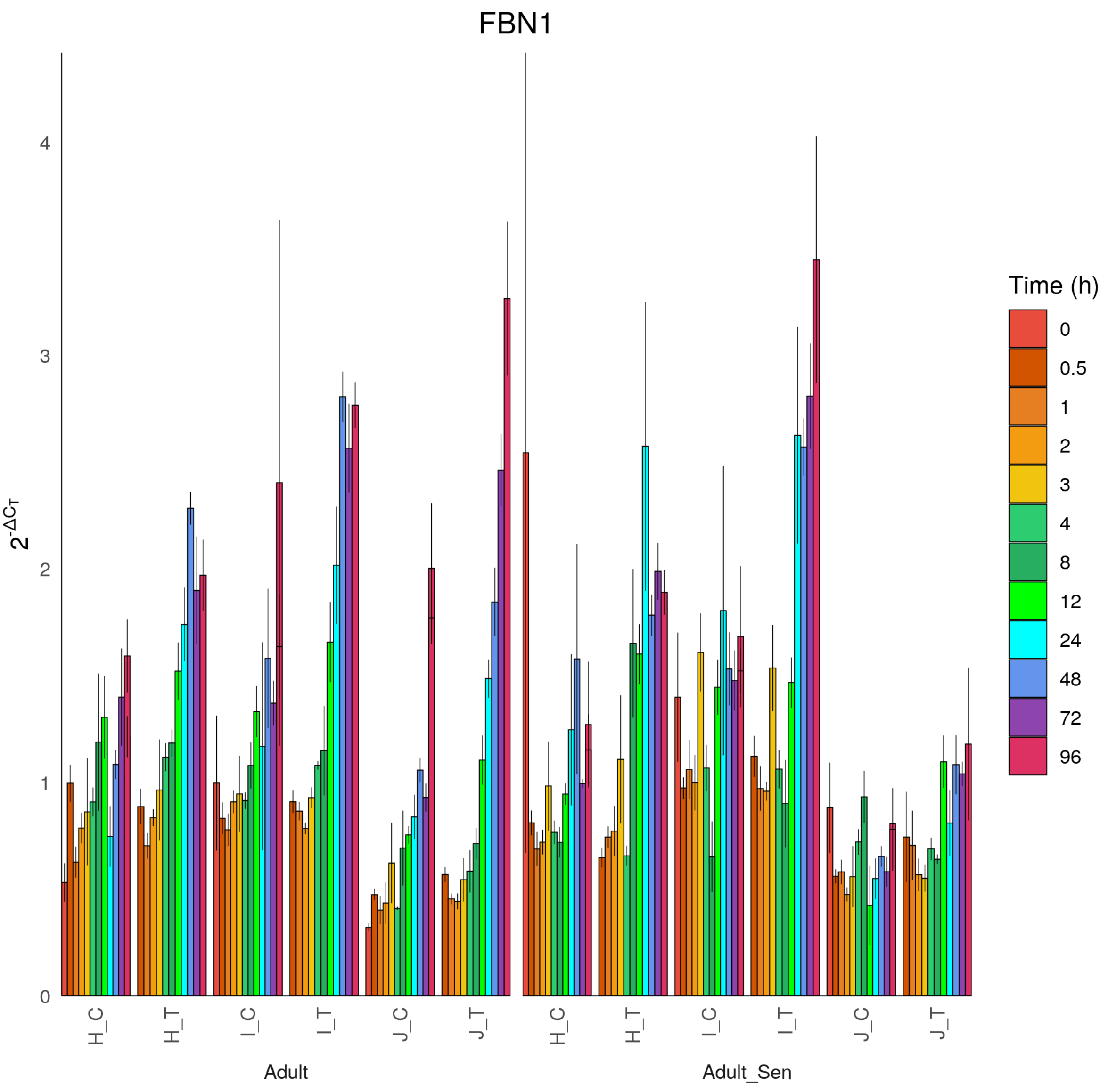
FBN1
$2^{-\Delta C_T}$
0
1
2
3
4
H_C
H_T
I_C
I_T
J_C
J_T
Adult
Adult_Sen
Time (h)
0
0.5
1
2
3
4
8
12
24
48
72
96

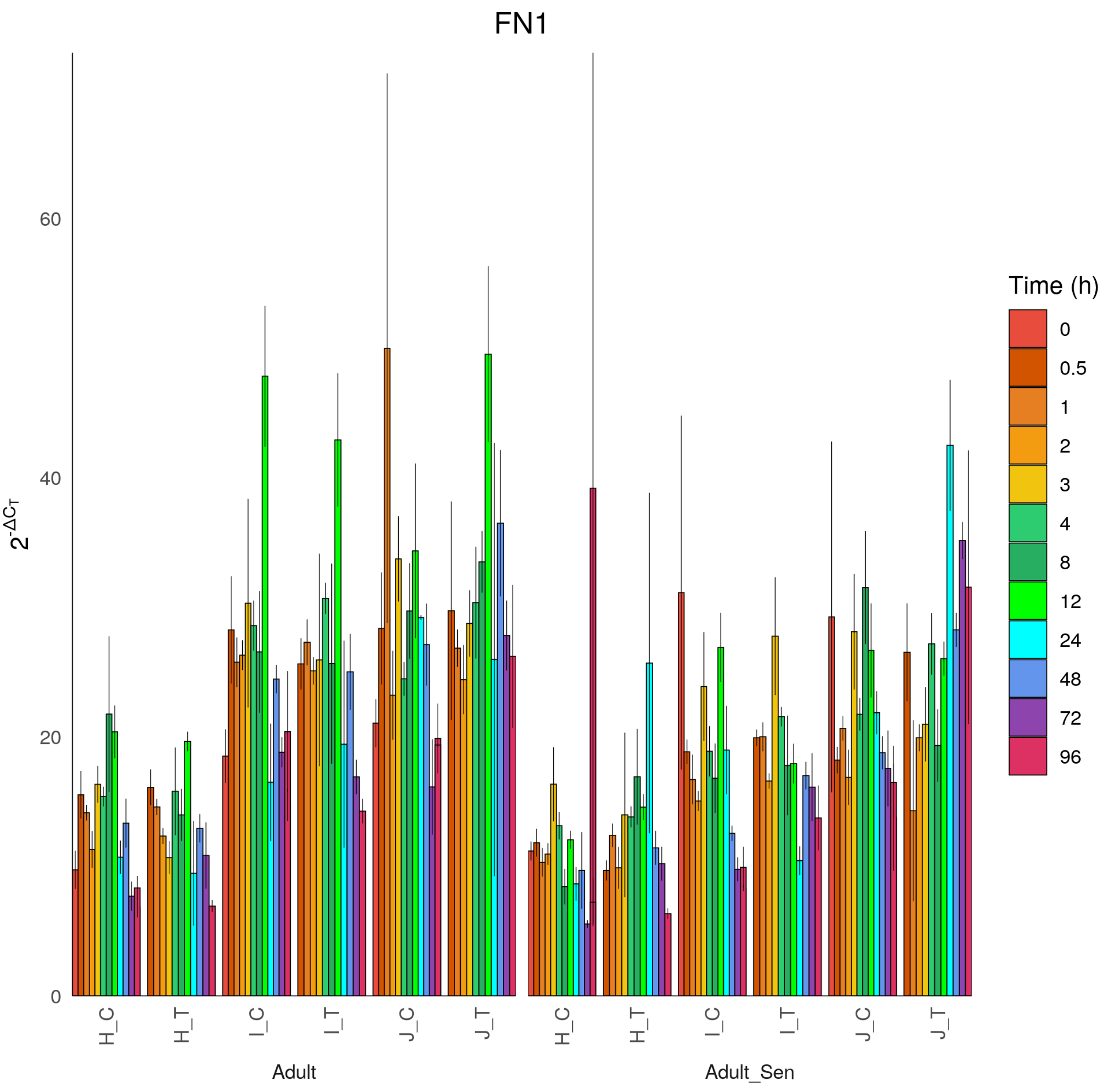
FN1
$2^{-\Delta C_T}$
0
20
40
60
H_C
H_T
I_C
I_T
J_C
J_T
Adult
Adult_Sen
Time (h)
0
0.5
1
2
3
4
8
12
24
48
72
96

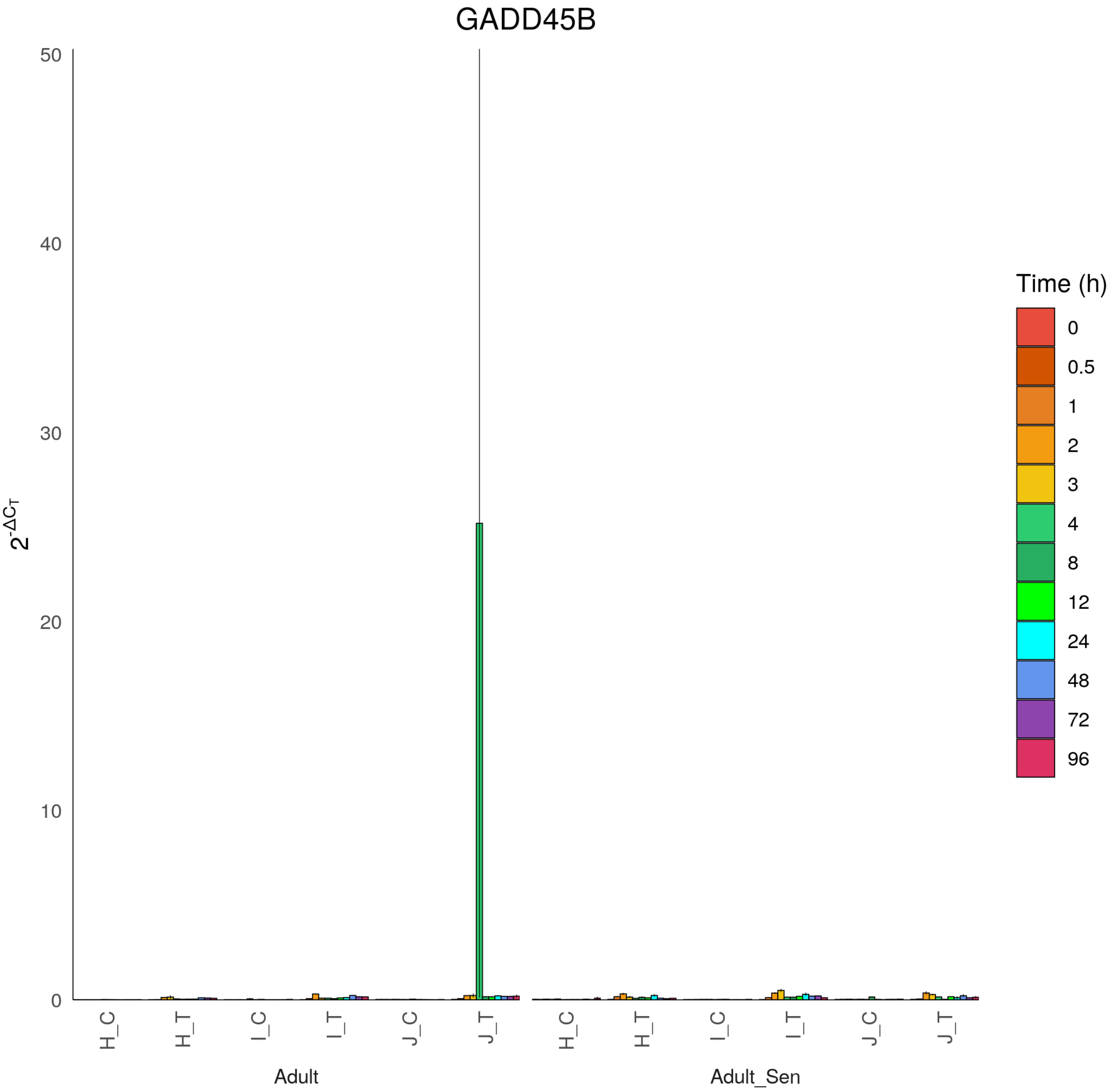

GADD45B
$2^{-\Delta C_T}$
50
40
30
20
10
0
H_C
H_T
I_C
I_T
J_C
J_T
Adult
H_C
H_T
I_C
I_T
J_C
J_T
Adult_Sen
Time (h)
0
0.5
1
2
3
4
8
12
24
48
72
96

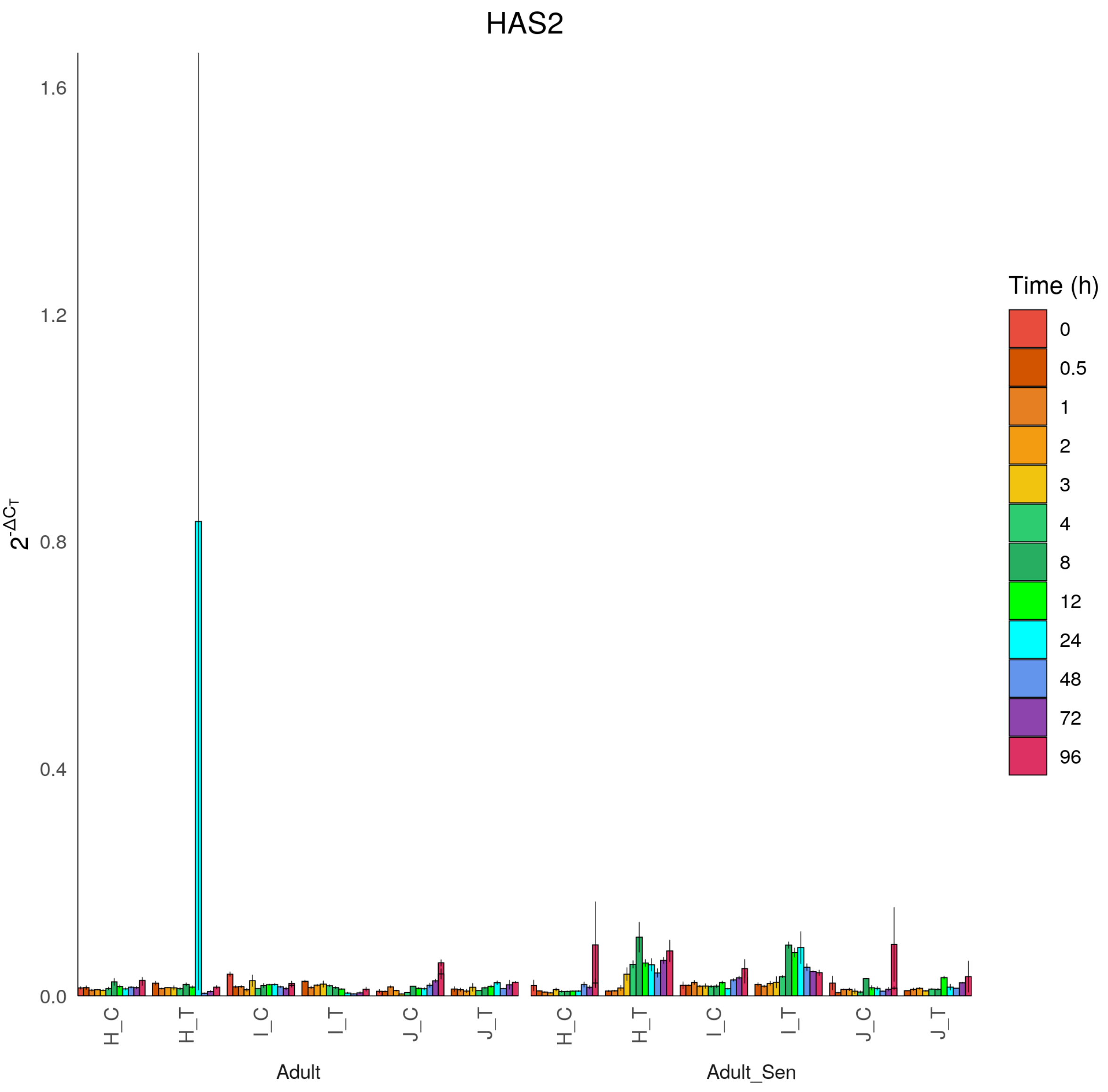

HAS2
2^-ΔCT
1.6
1.2
0.8
0.4
0.0
H_C
H_T
I_C
I_T
J_C
J_T
Adult
Adult_Sen
Time (h)
0
0.5
1
2
3
4
8
12
24
48
72
96

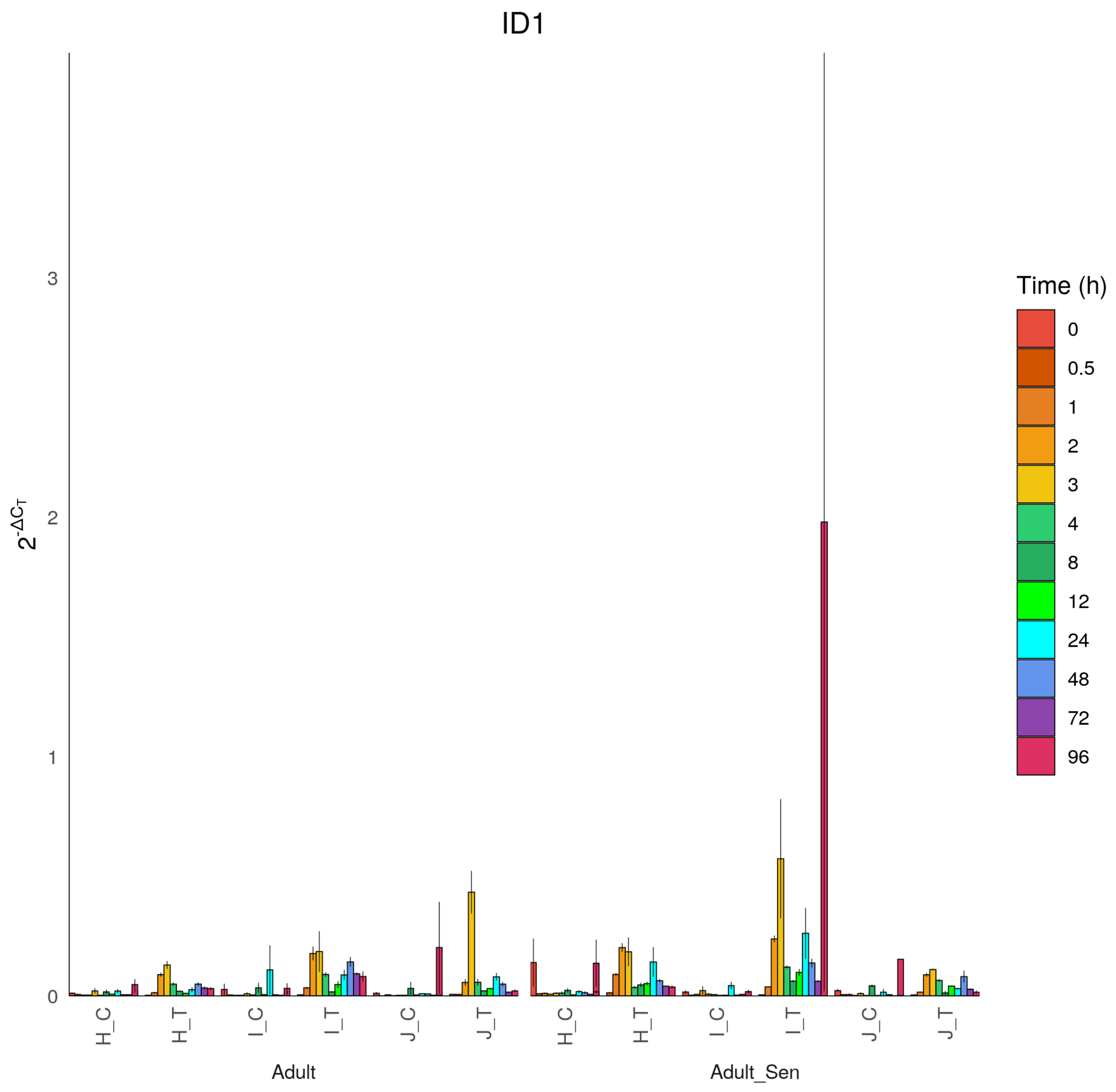

ID1
$2^{-\Delta C_T}$
0
1
2
3
H_C
H_T
I_C
I_T
J_C
J_T
Adult
Adult_Sen
Time (h)
0
0.5
1
2
3
4
8
12
24
48
72
96

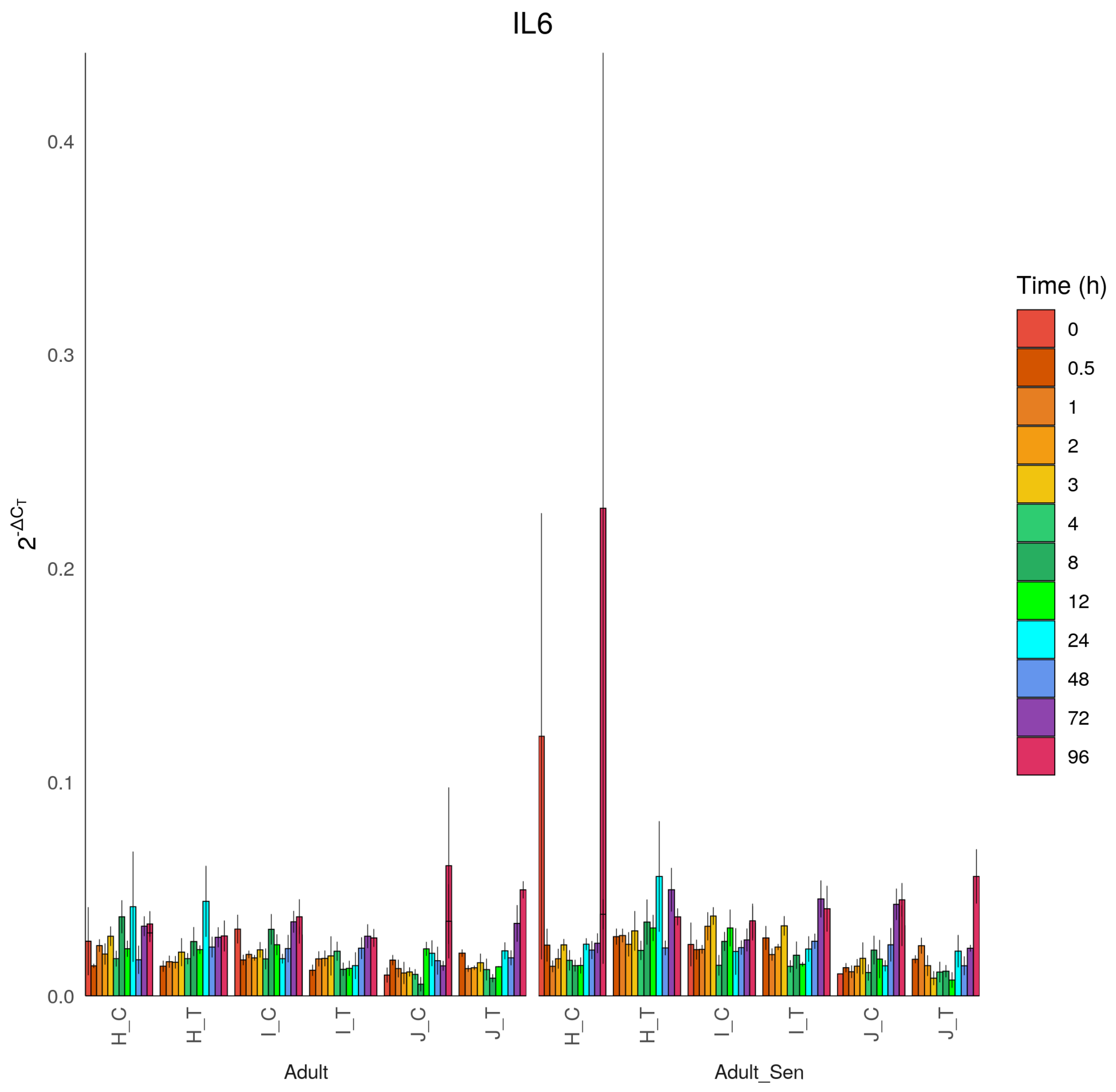

IL6
$2^{-\Delta C_T}$
0.4
0.3
0.2
0.1
0.0
H_C
H_T
I_C
I_T
J_C
J_T
Adult
Adult_Sen
Time (h)
0
0.5
1
2
3
4
8
12
24
48
72
96

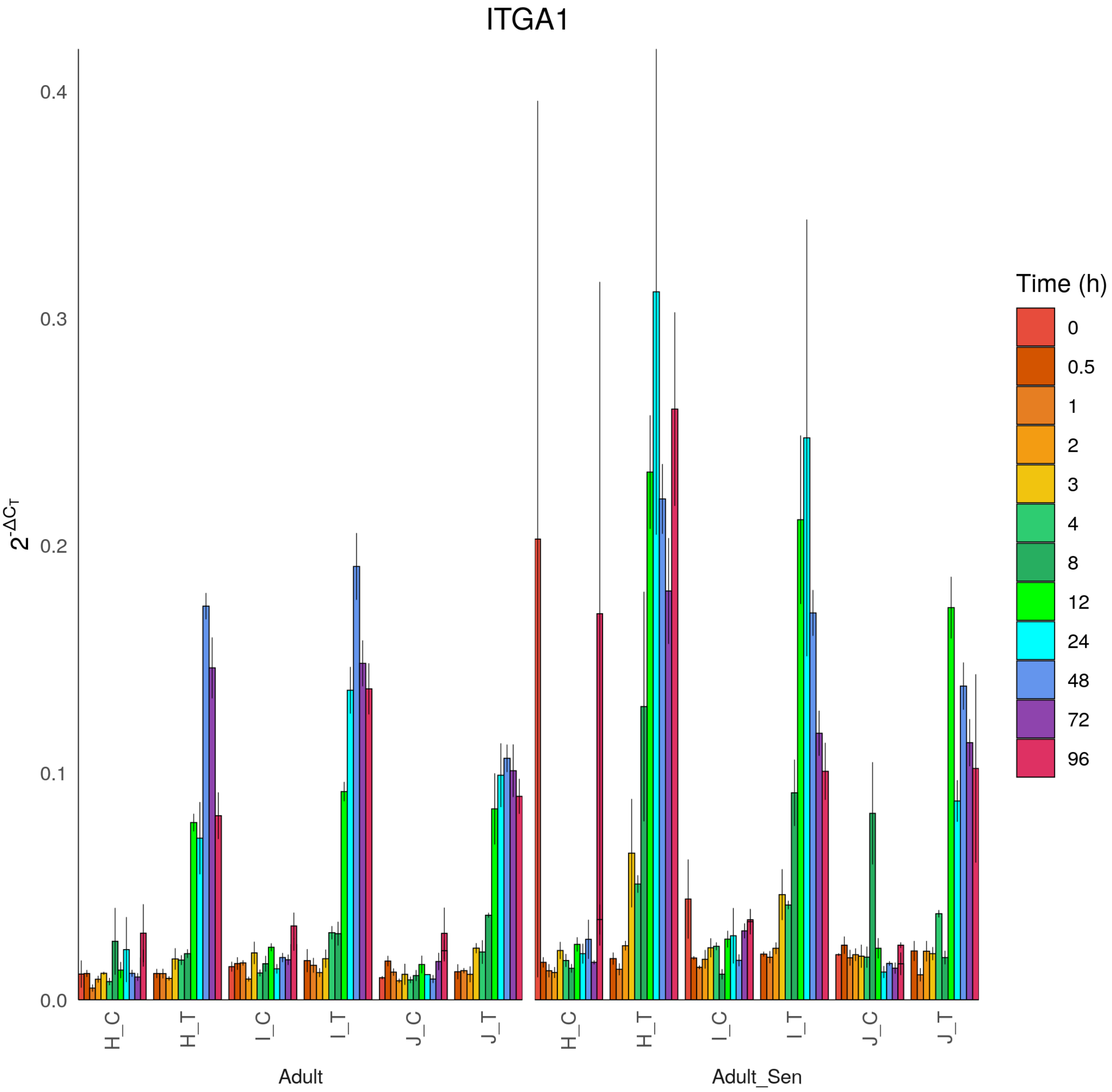

ITGA1
0.4
0.3
0.2
0.1
0.0
$2^{-\Delta C_T}$
H_C
H_T
I_C
I_T
J_C
J_T
Adult
Adult_Sen
Time (h)
0
0.5
1
2
3
4
8
12
24
48
72
96

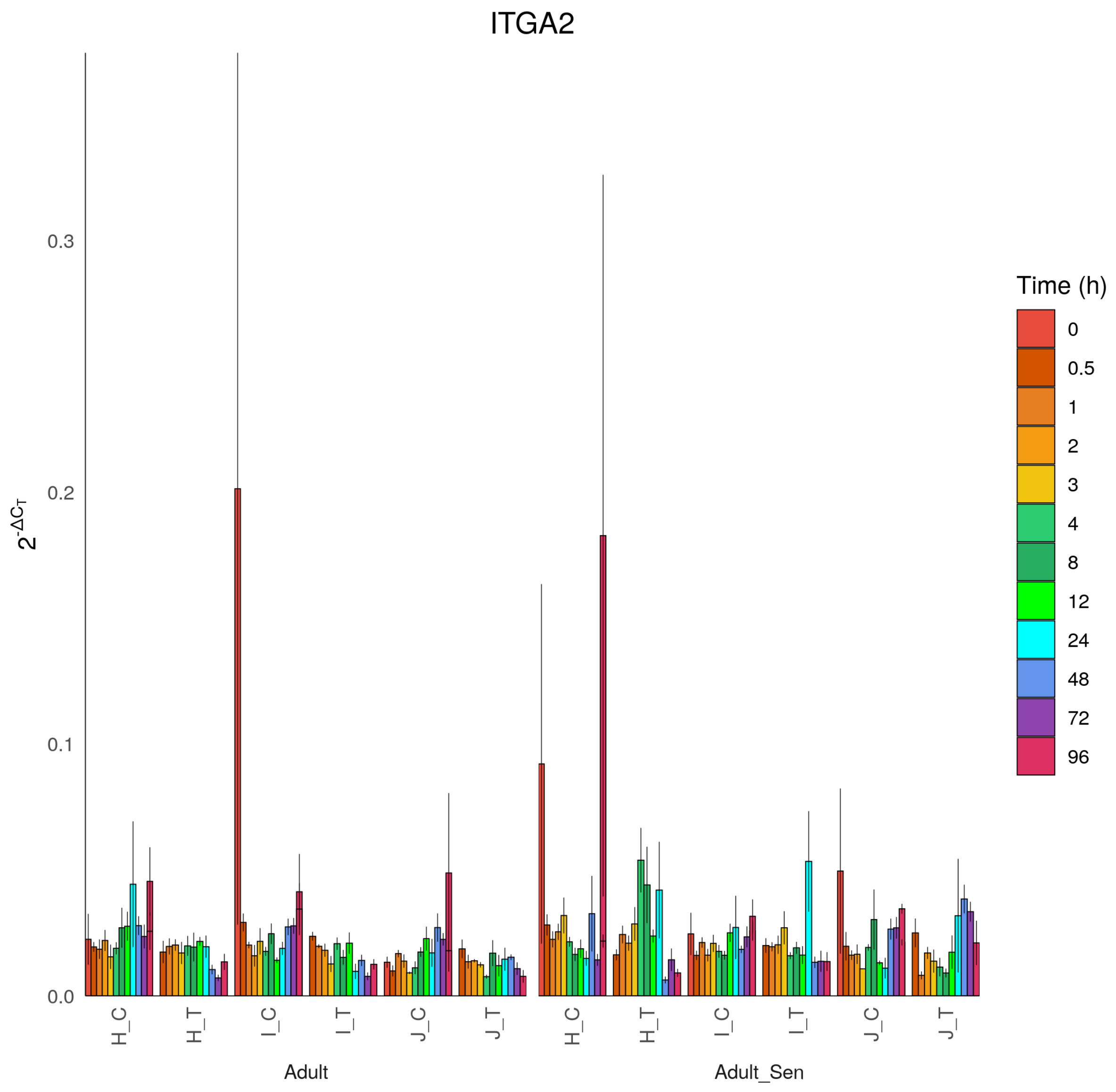
ITGA2
$2^{-\Delta C_T}$
0.0
0.1
0.2
0.3
H_C
H_T
I_C
I_T
J_C
J_T
Adult
Adult_Sen
Time (h)
0
0.5
1
2
3
4
8
12
24
48
72
96

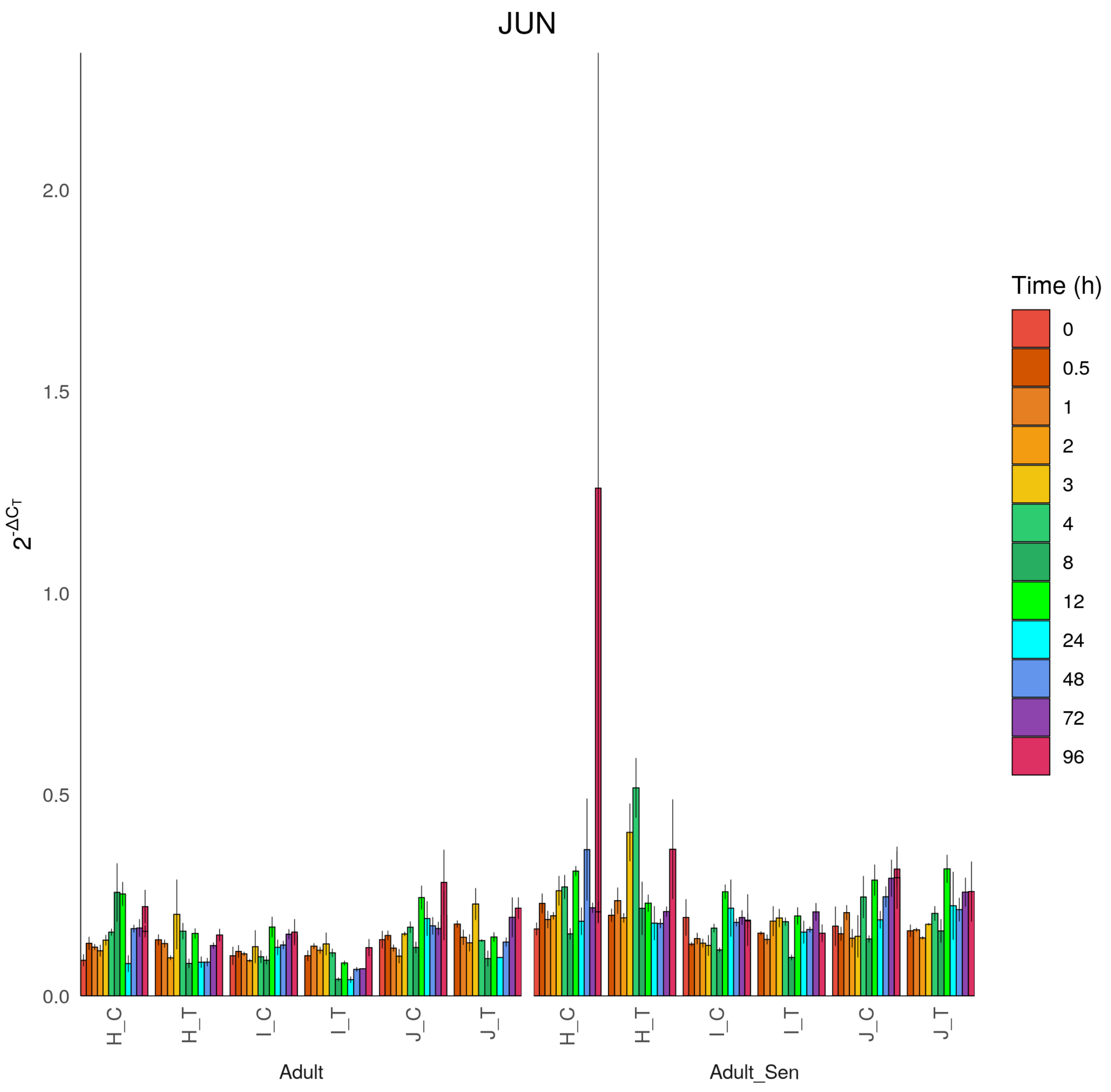
JUN
$2^{-\Delta C_T}$
0.0
0.5
1.0
1.5
2.0
H_C
H_T
I_C
I_T
J_C
J_T
Adult
Adult_Sen
Time (h)
0
0.5
1
2
3
4
8
12
24
48
72
96

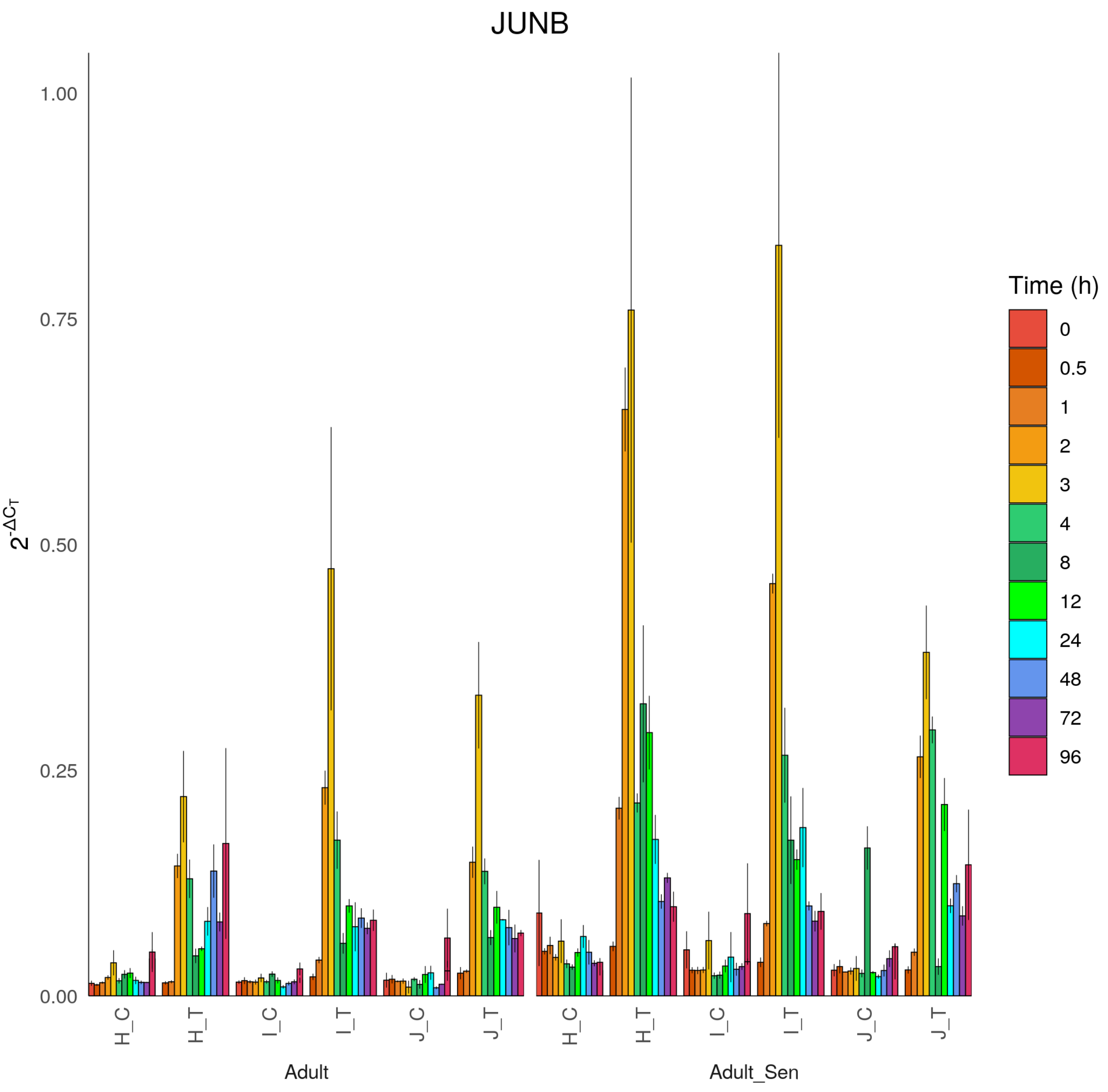

JUNB
1.00
0.75
0.50
0.25
0.00
$2^{-\Delta C_T}$
H_C
H_T
I_C
I_T
J_C
J_T
Adult
Adult_Sen
Time (h)
0
0.5
1
2
3
4
8
12
24
48
72
96

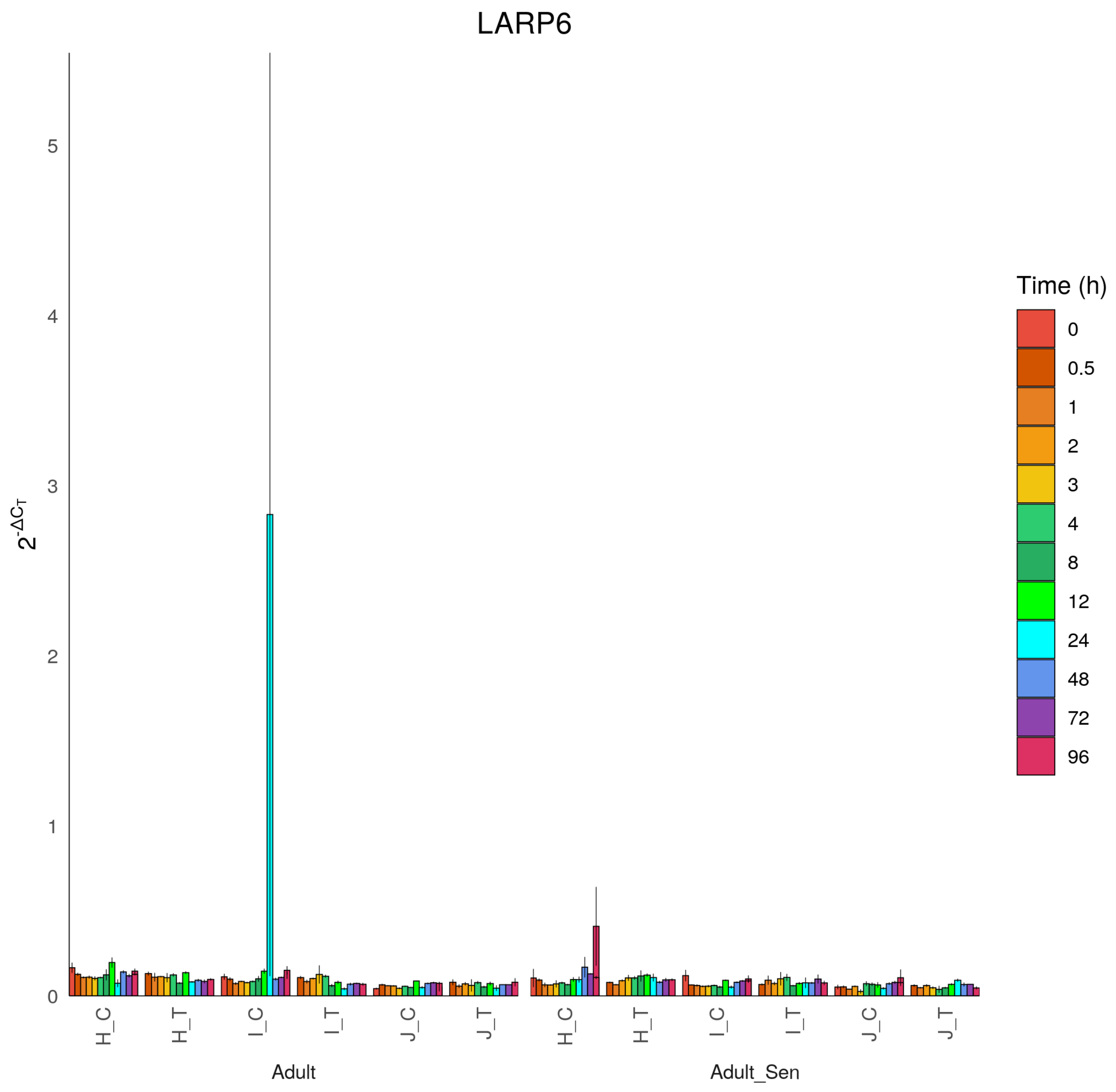

LARP6
$2^{-\Delta C_T}$
0
1
2
3
4
5
H_C
H_T
I_C
I_T
J_C
J_T
Adult
Adult_Sen
Time (h)
0
0.5
1
2
3
4
8
12
24
48
72
96

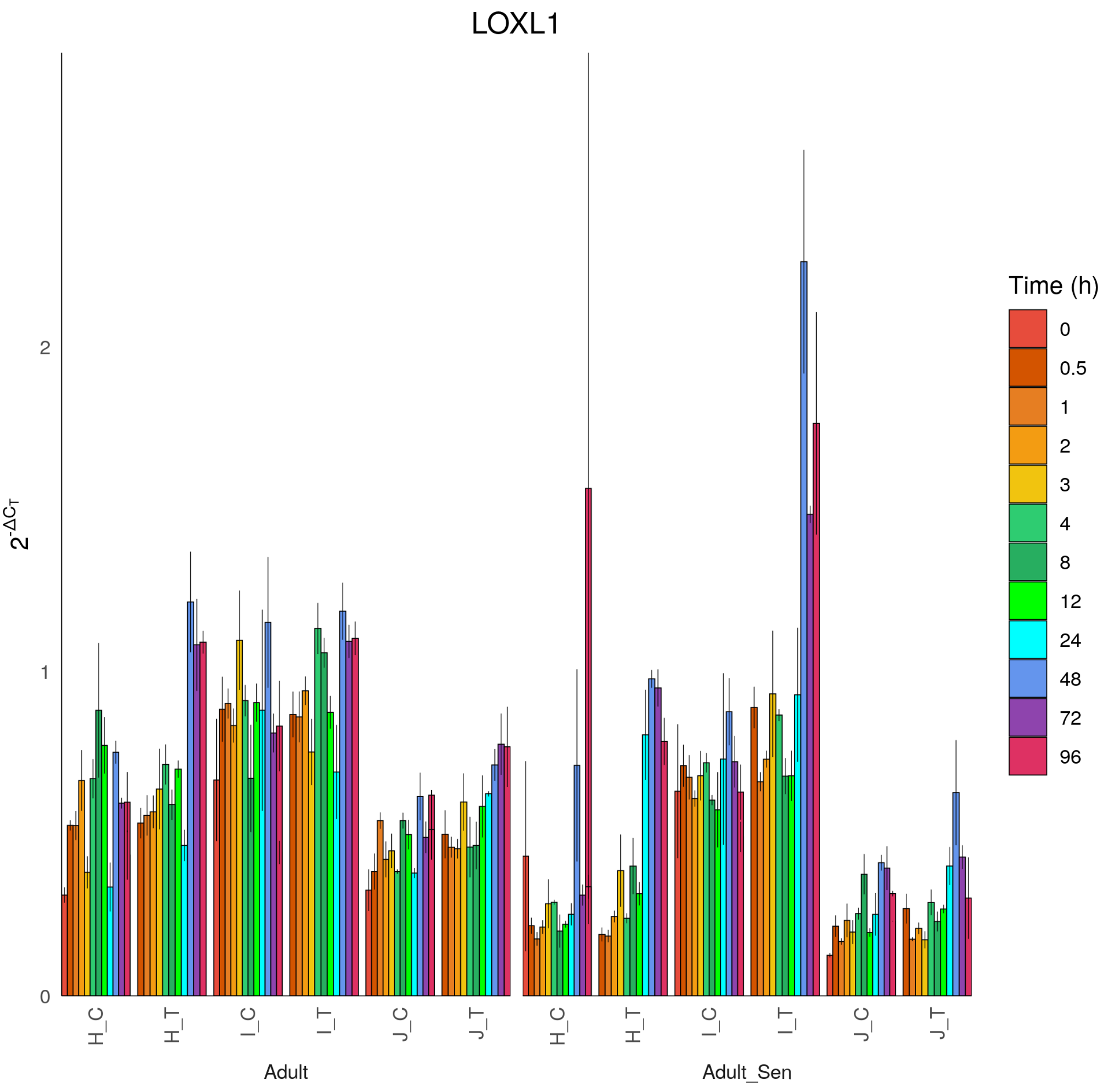
LOXL1
$2^{-\Delta C_T}$
0
1
2
H_C
H_T
I_C
I_T
J_C
J_T
Adult
Adult_Sen
Time (h)
0
0.5
1
2
3
4
8
12
24
48
72
96

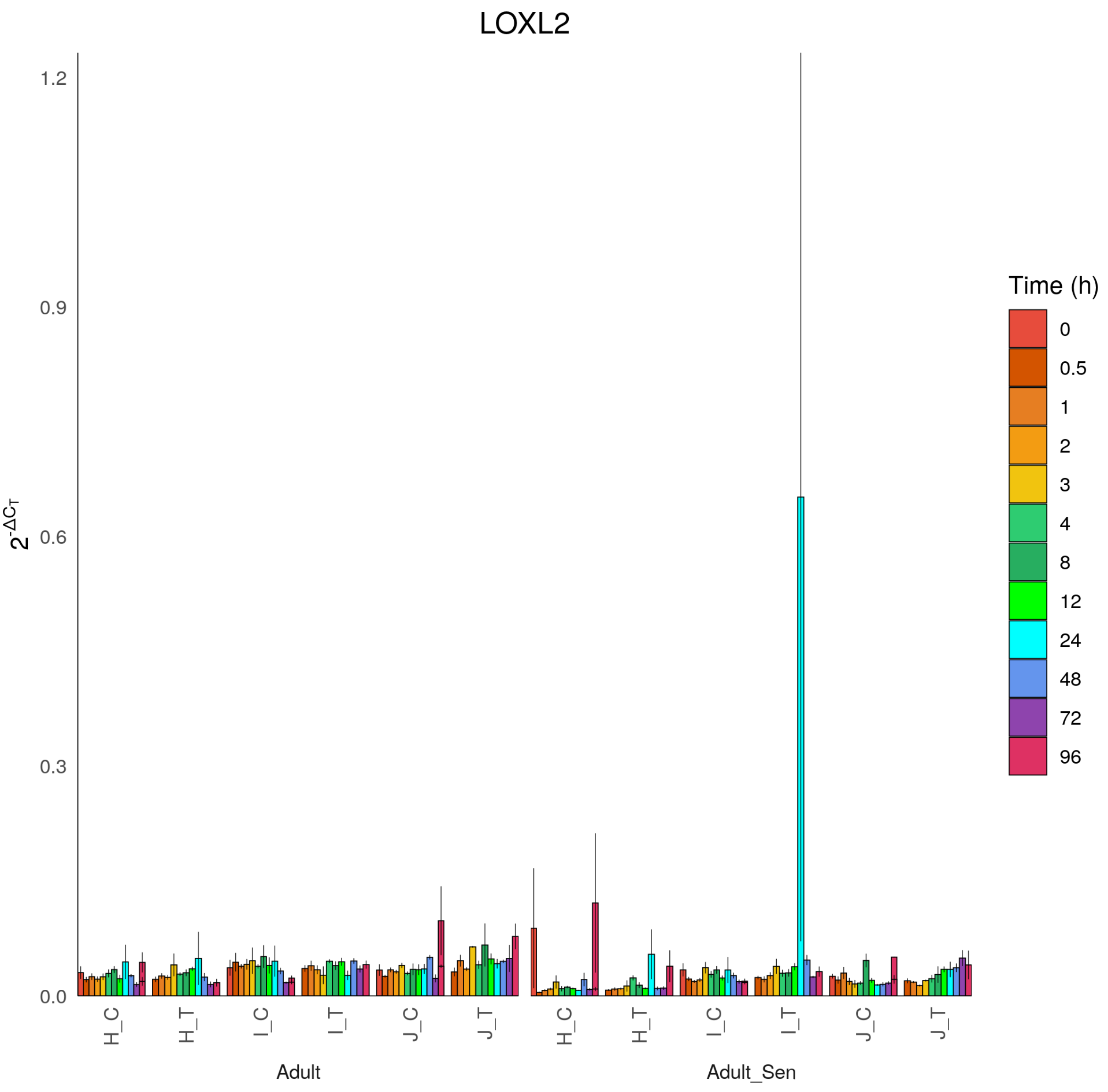

LOXL2
$2^{-\Delta C_T}$
1.2
0.9
0.6
0.3
0.0
H_C
H_T
I_C
I_T
J_C
J_T
Adult
Adult_Sen
Time (h)
0
0.5
1
2
3
4
8
12
24
48
72
96

### LTBP2

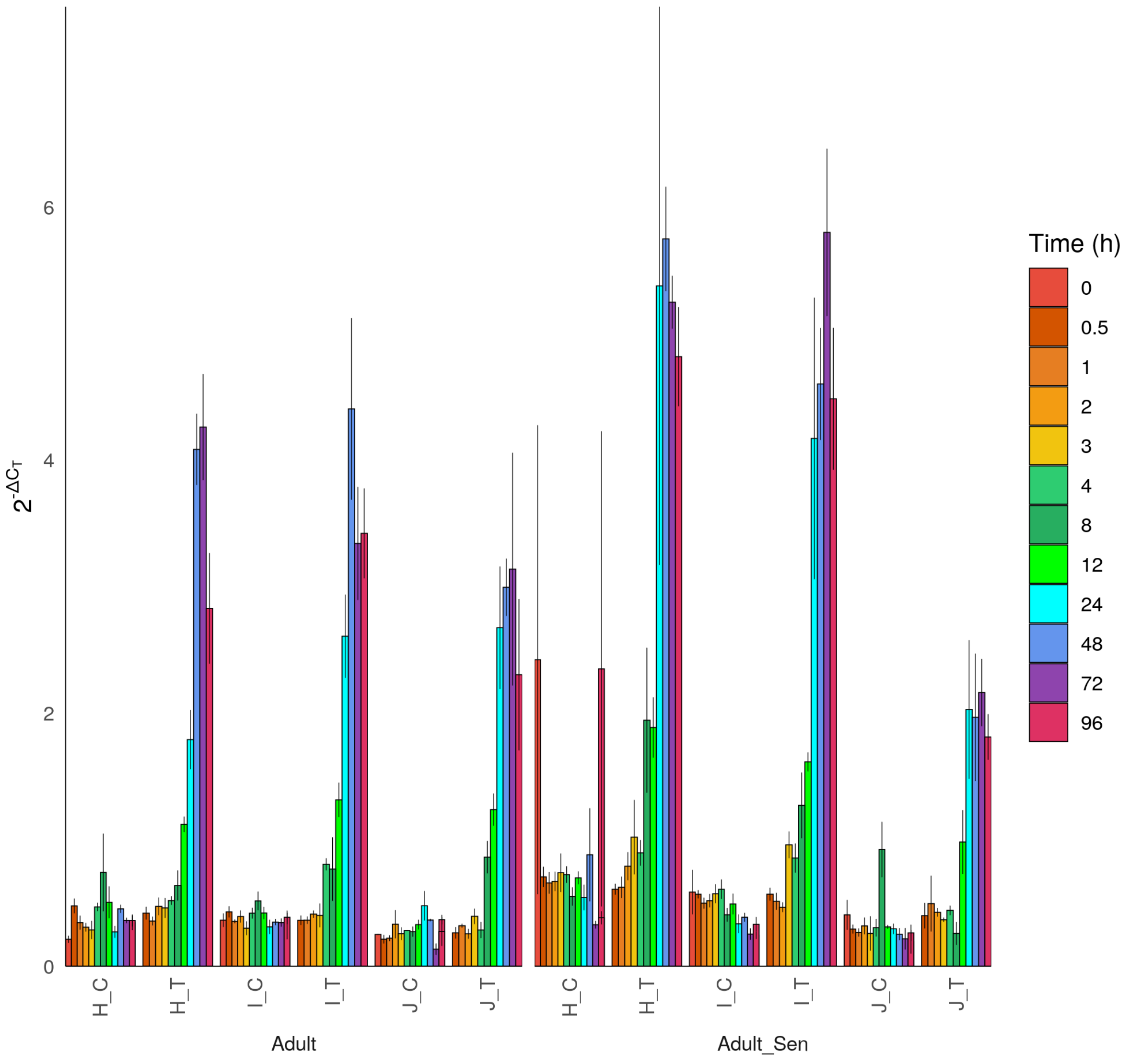

LTBP2
$2^{-\Delta C_T}$
0
2
4
6
H_C
H_T
I_C
I_T
J_C
J_T
Adult
Adult_Sen
Time (h)
0
0.5
1
2
3
4
8
12
24
48
72
96

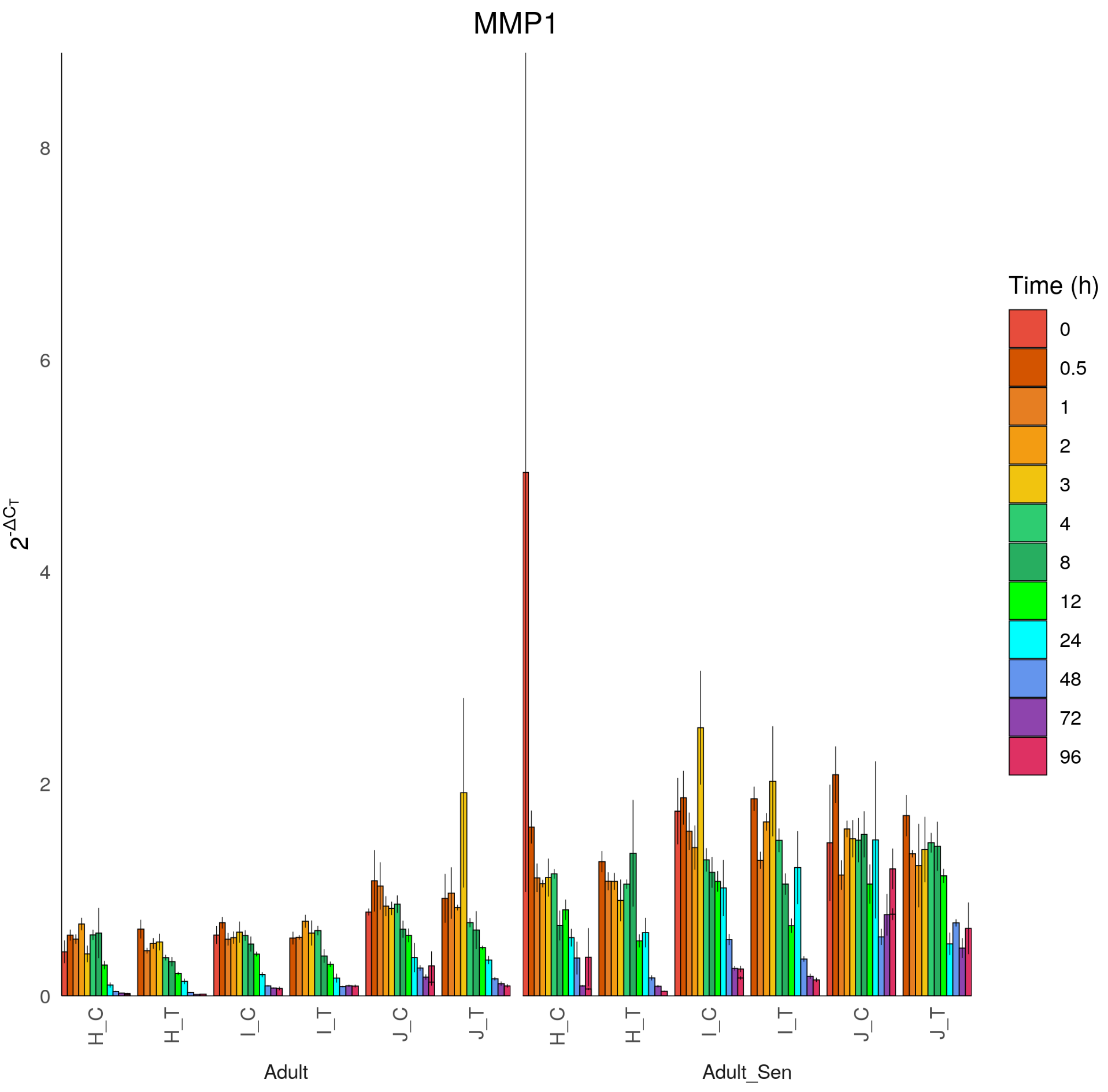

MMP1
2^-ΔCT
0
2
4
6
8
H_C
H_T
I_C
I_T
J_C
J_T
Adult
Adult_Sen
Time (h)
0
0.5
1
2
3
4
8
12
24
48
72
96

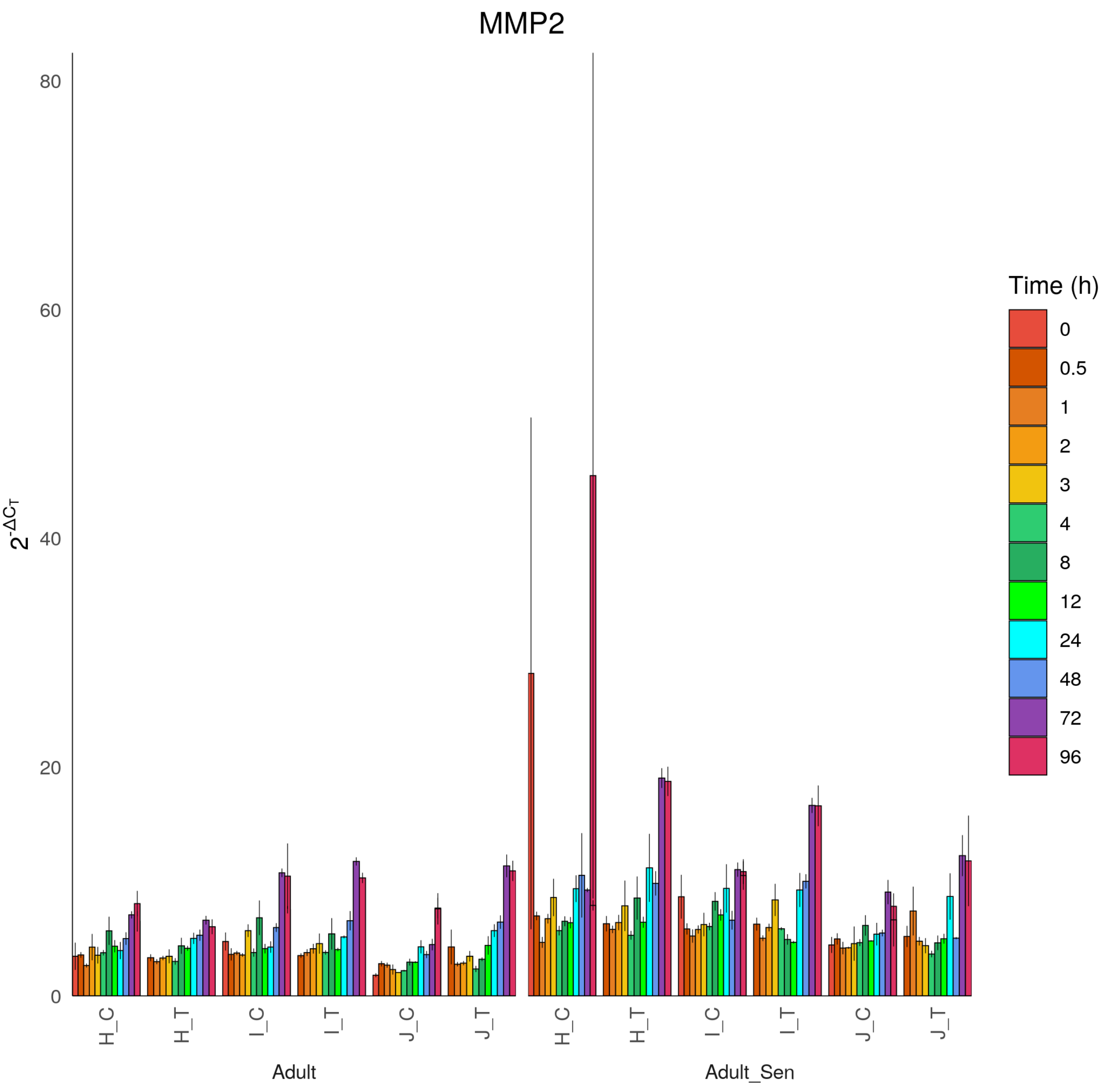
MMP2
$2^{-\Delta C_T}$
0
20
40
60
80
H_C
H_T
I_C
I_T
J_C
J_T
Adult
Adult_Sen
Time (h)
0
0.5
1
2
3
4
8
12
24
48
72
96

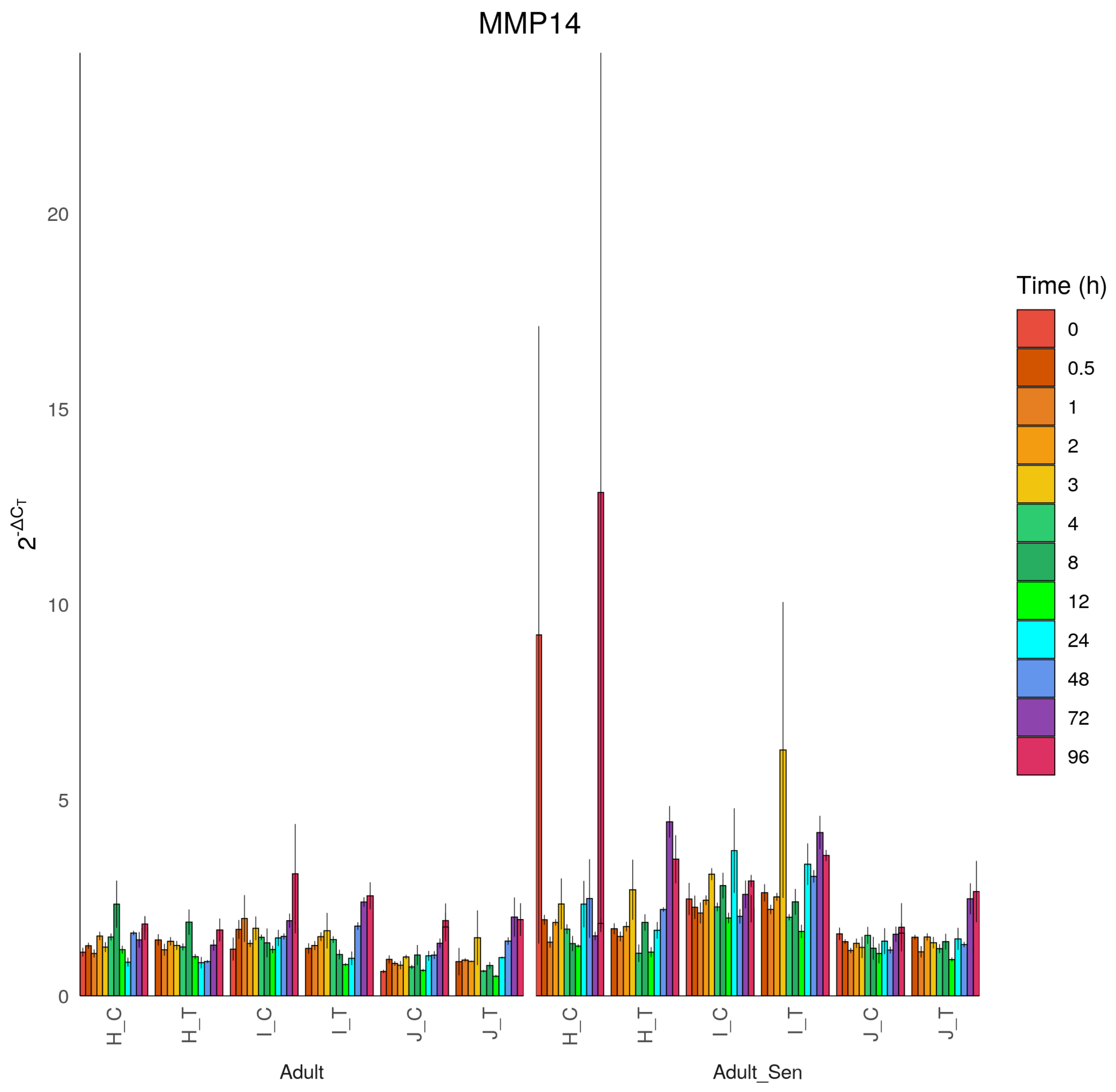
MMP14
2^-ΔCT
20
15
10
5
0
H_C
H_T
I_C
I_T
J_C
J_T
Adult
Adult_Sen
Time (h)
0
0.5
1
2
3
4
8
12
24
48
72
96

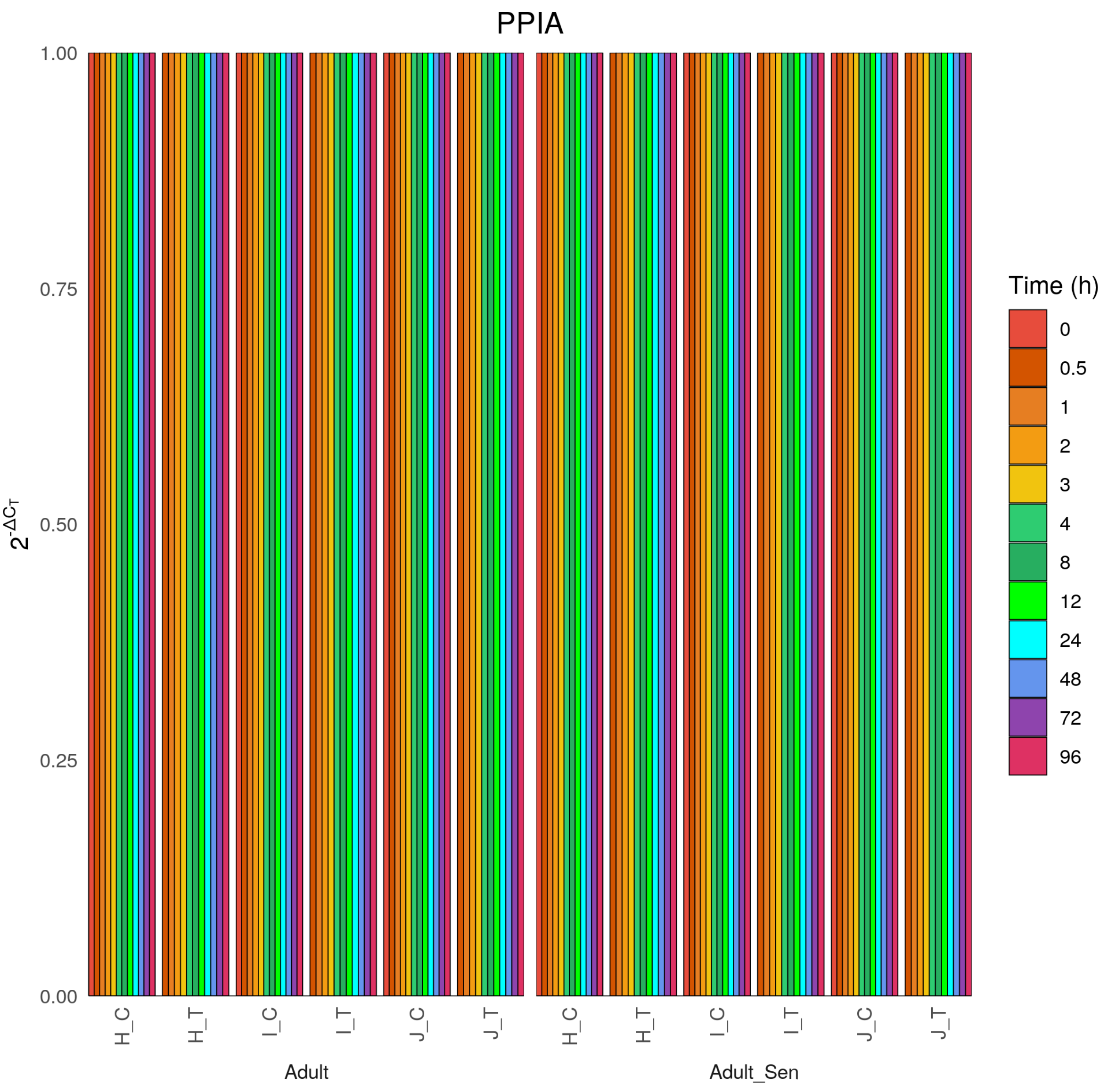
PPIA
1.00
0.75
0.50
0.25
0.00
$2^{-\Delta C_T}$
H_C
H_T
I_C
I_T
J_C
J_T
H_C
H_T
I_C
I_T
J_C
J_T
Adult
Adult_Sen
Time (h)
0
0.5
1
2
3
4
8
12
24
48
72
96

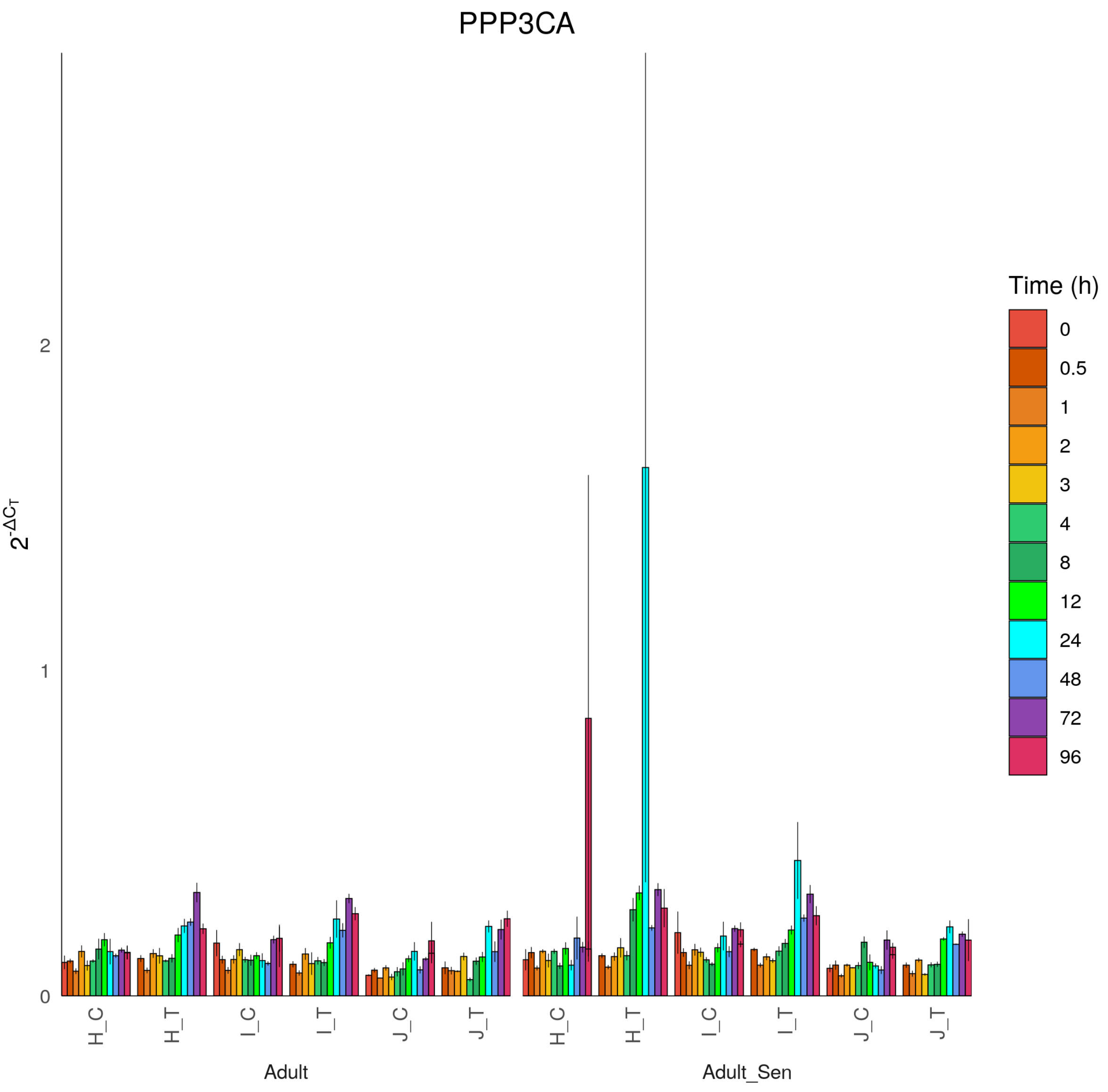

PPP3CA
2^-ΔCT
0
1
2
H_C
H_T
I_C
I_T
J_C
J_T
Adult
Adult_Sen
Time (h)
0
0.5
1
2
3
4
8
12
24
48
72
96

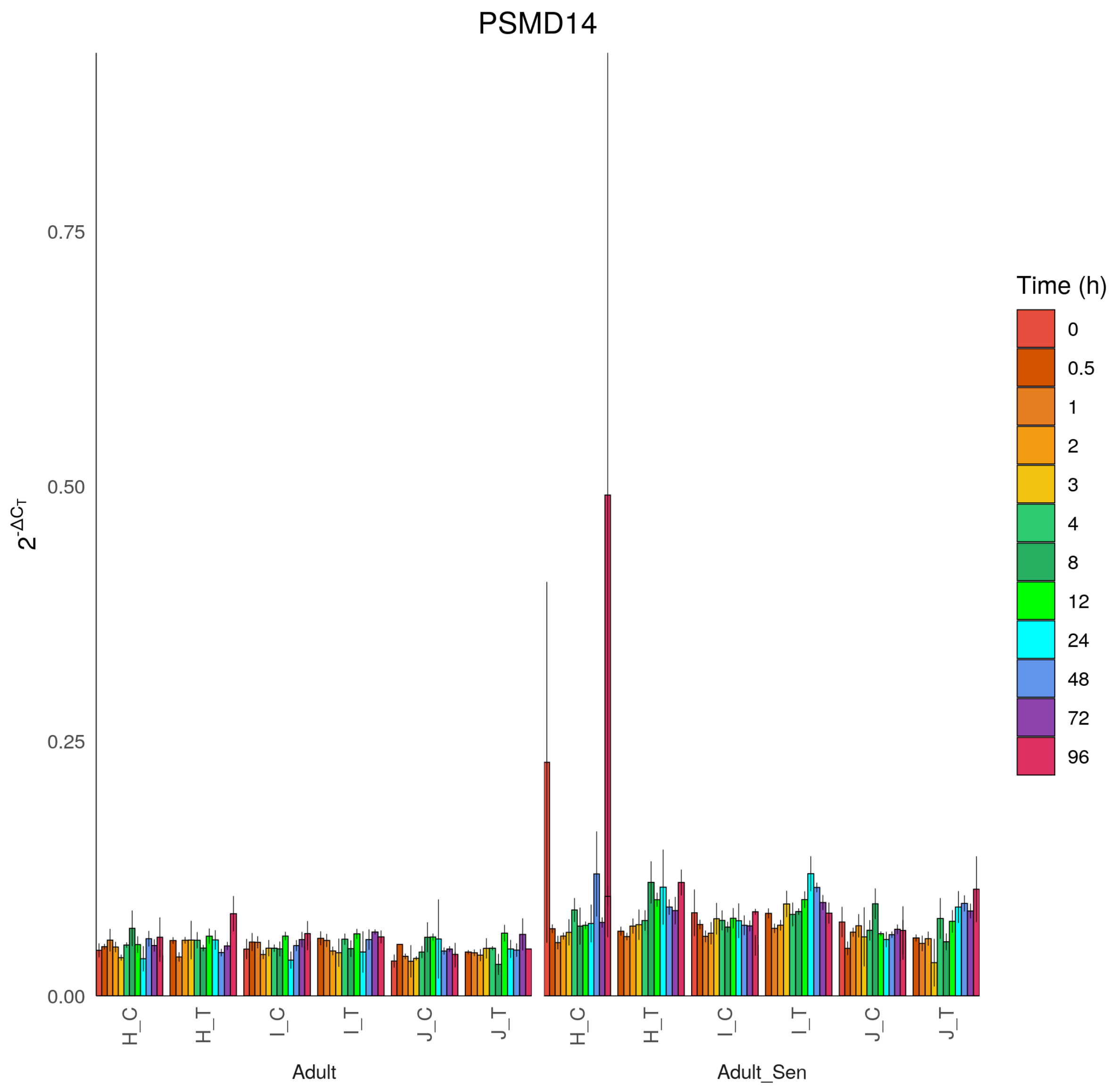
PSMD14
0.75
0.50
0.25
0.00
$2^{-\Delta C_T}$
H_C
H_T
I_C
I_T
J_C
J_T
Adult
Adult_Sen
Time (h)
0
0.5
1
2
3
4
8
12
24
48
72
96

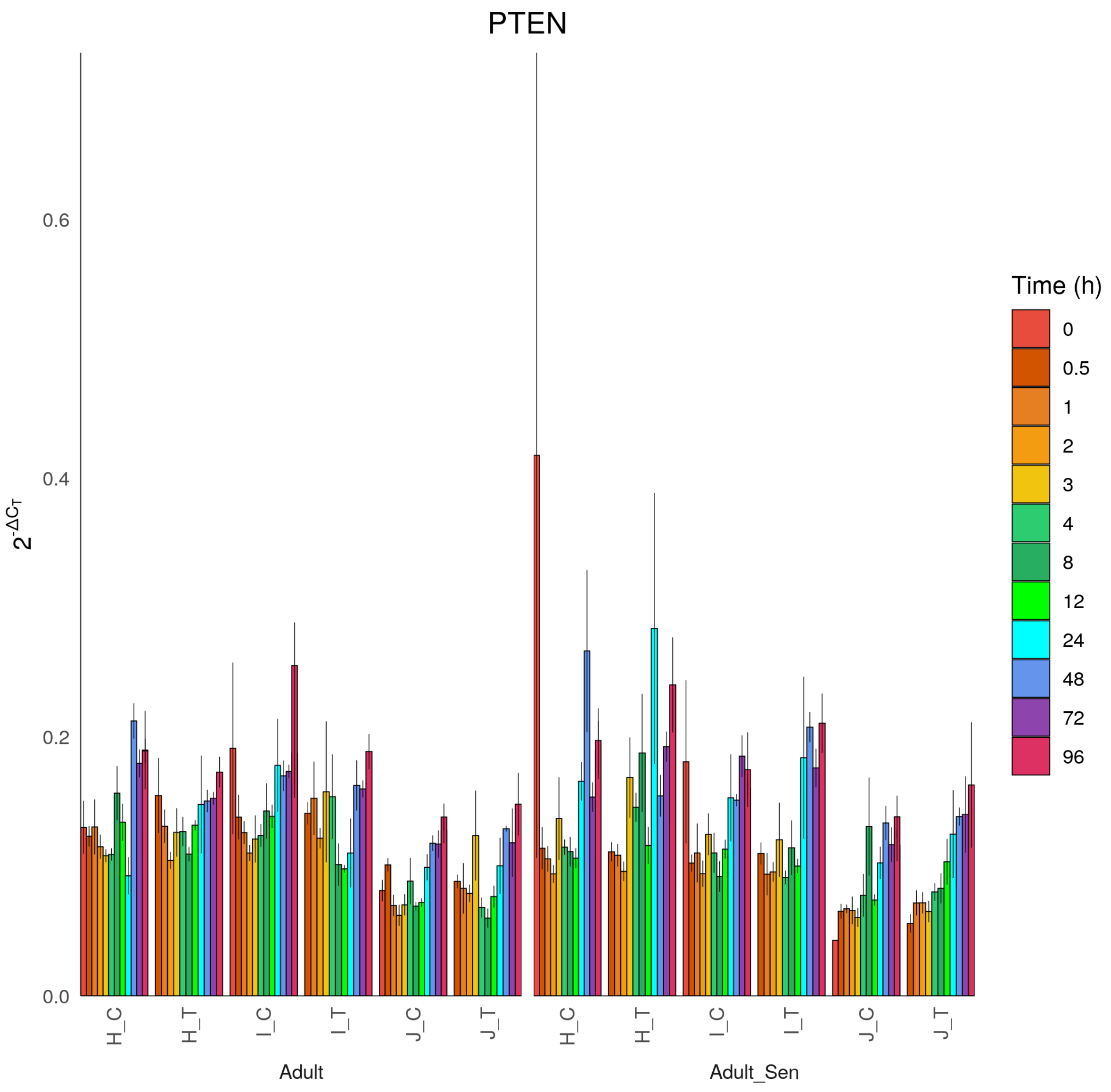
PTEN
$2^{-\Delta C_T}$
0.0
0.2
0.4
0.6
H_C
H_T
I_C
I_T
J_C
J_T
Adult
Adult_Sen
Time (h)
0
0.5
1
2
3
4
8
12
24
48
72
96

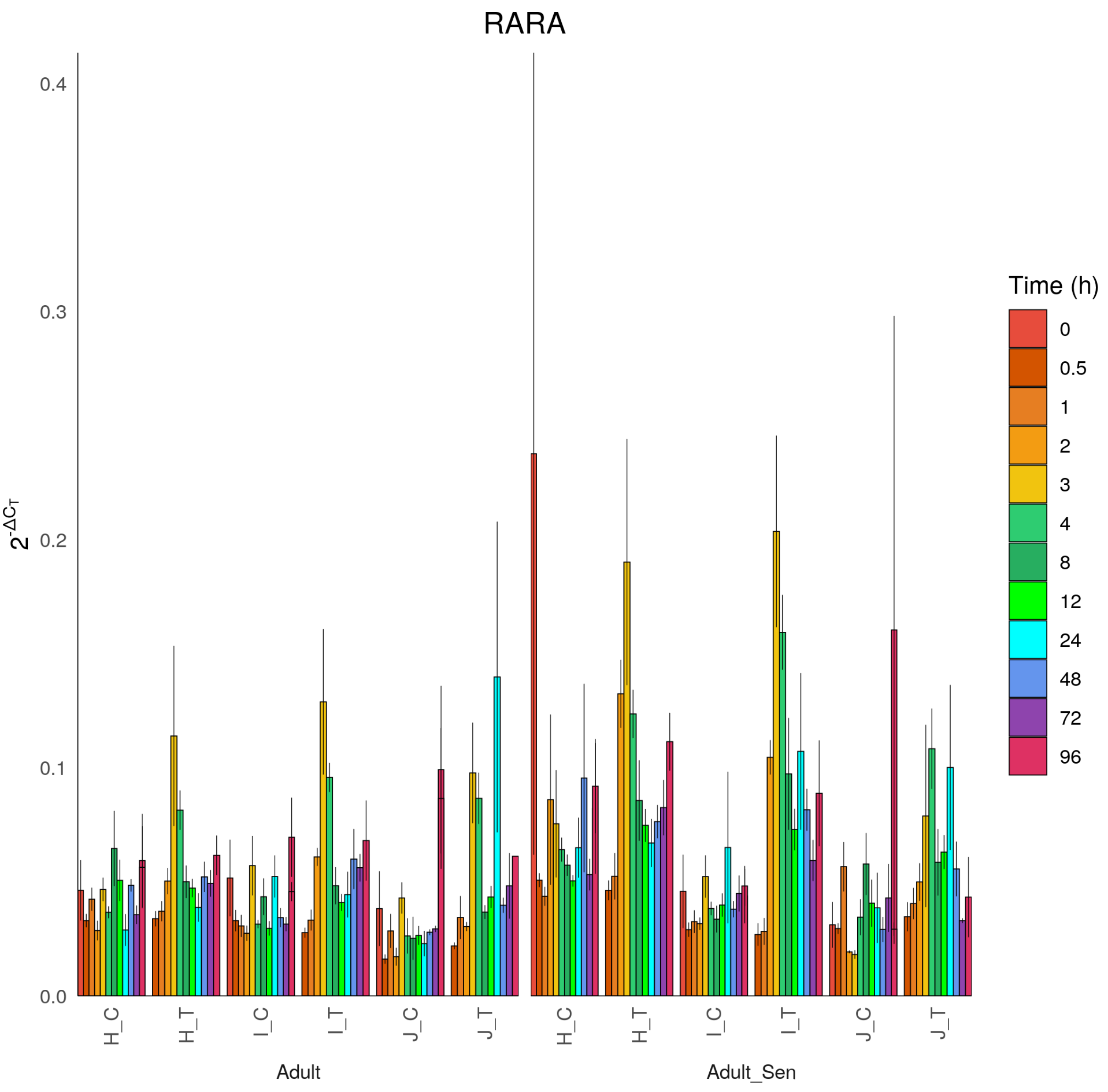

RARA
$2^{-\Delta C_T}$
0.0
0.1
0.2
0.3
0.4
H_C
H_T
I_C
I_T
J_C
J_T
Adult
Adult_Sen
Time (h)
0
0.5
1
2
3
4
8
12
24
48
72
96

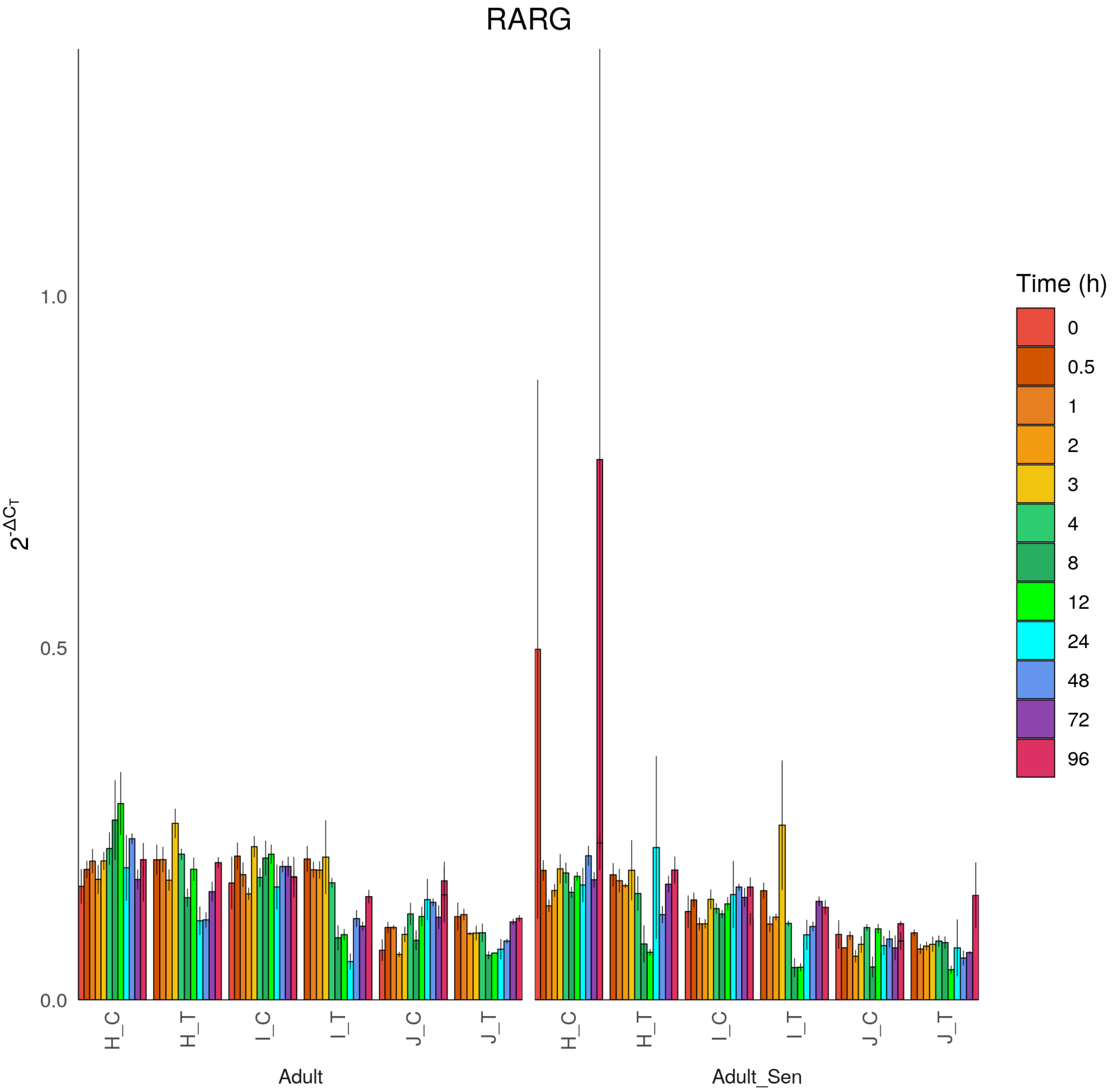
RARG
$2^{-\Delta C_T}$
1.0
0.5
0.0
H_C
H_T
I_C
I_T
J_C
J_T
Adult
Adult_Sen
Time (h)
0
0.5
1
2
3
4
8
12
24
48
72
96

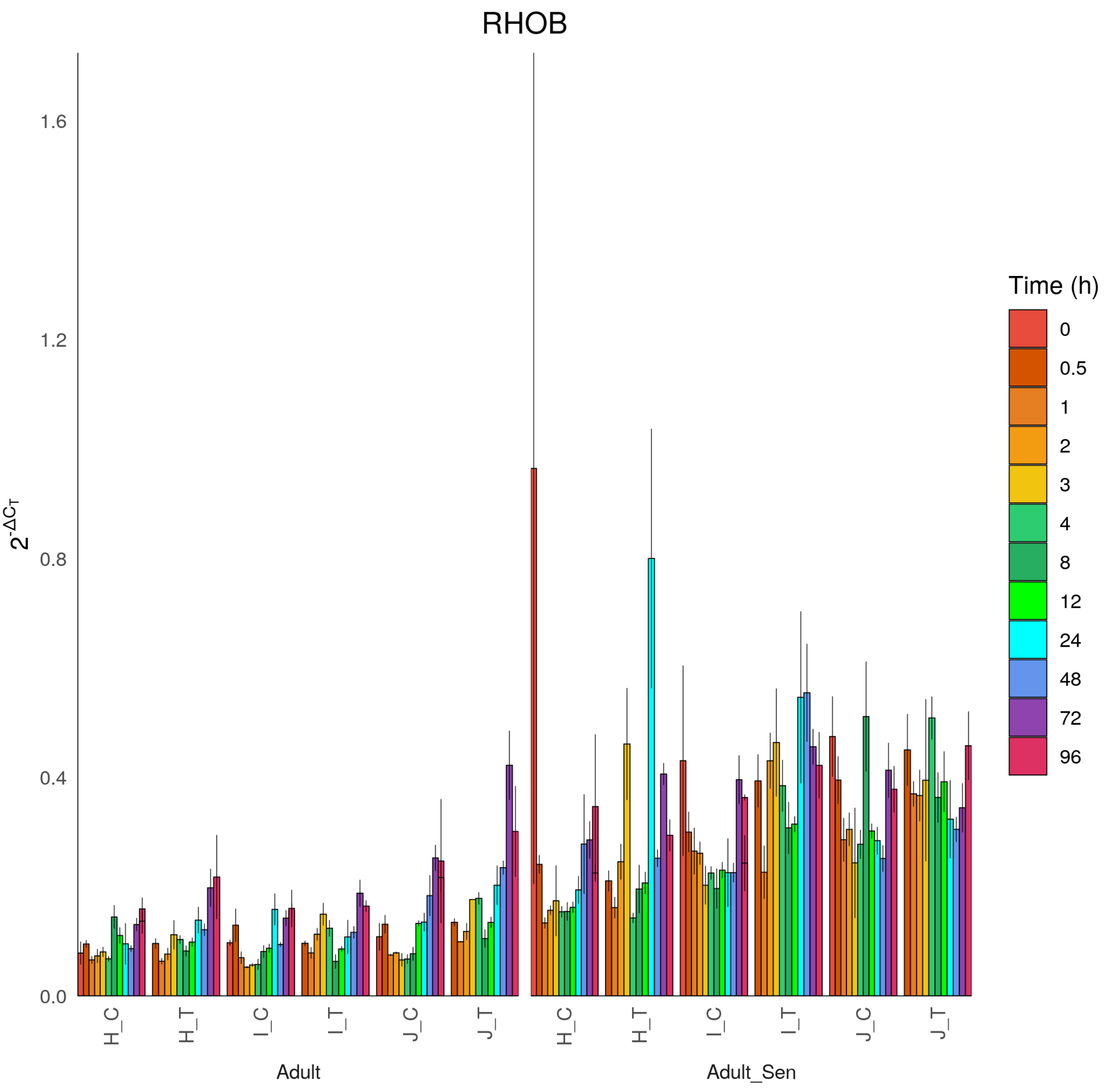
RHOB
$2^{-\Delta C_T}$
1.6
1.2
0.8
0.4
0.0
H_C
H_T
I_C
I_T
J_C
J_T
Adult
Adult_Sen
Time (h)
0
0.5
1
2
3
4
8
12
24
48
72
96

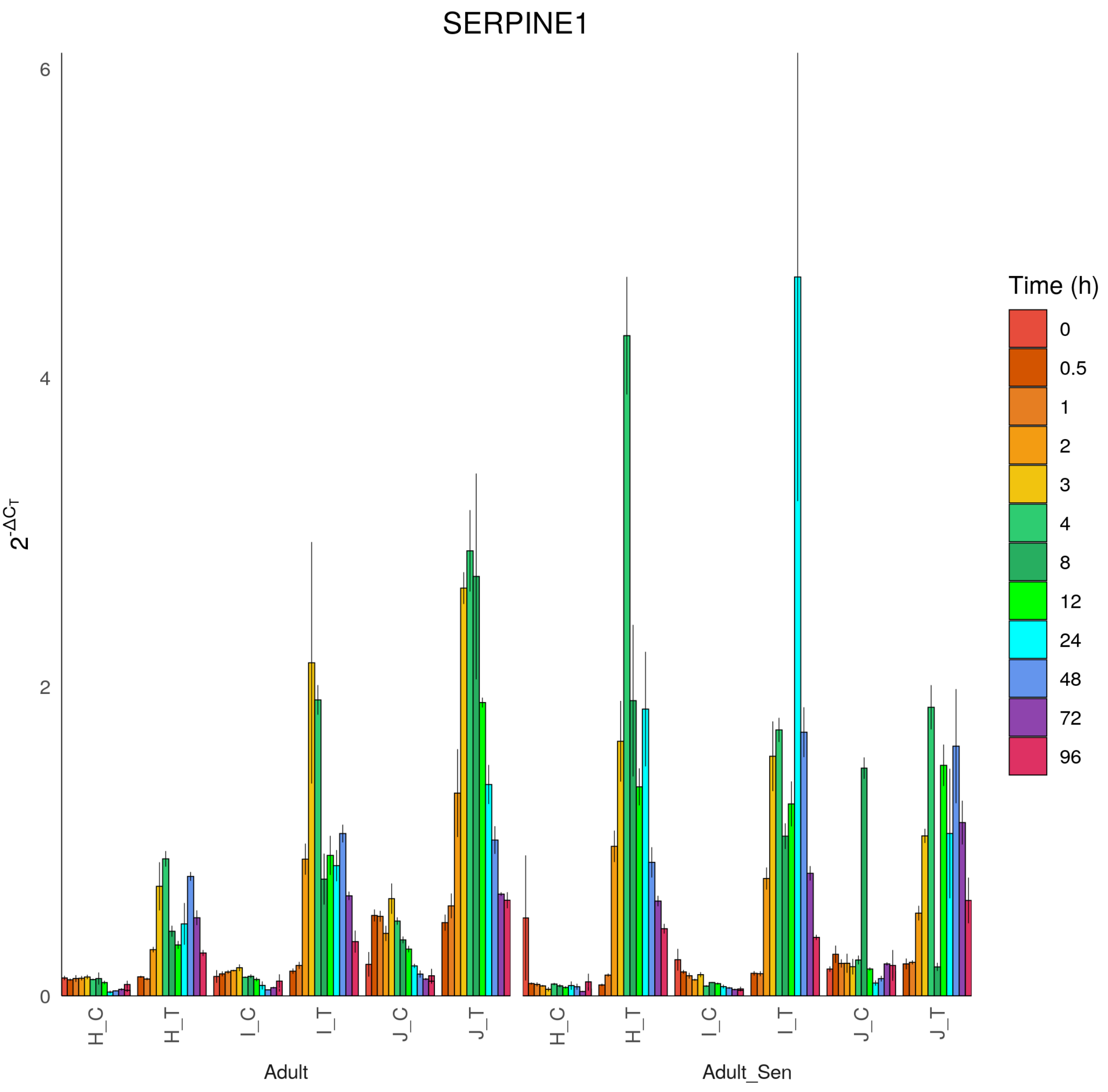

SERPINE1
$2^{-\Delta C_T}$
0
2
4
6
H_C
H_T
I_C
I_T
J_C
J_T
Adult
Adult_Sen
Time (h)
0
0.5
1
2
3
4
8
12
24
48
72
96

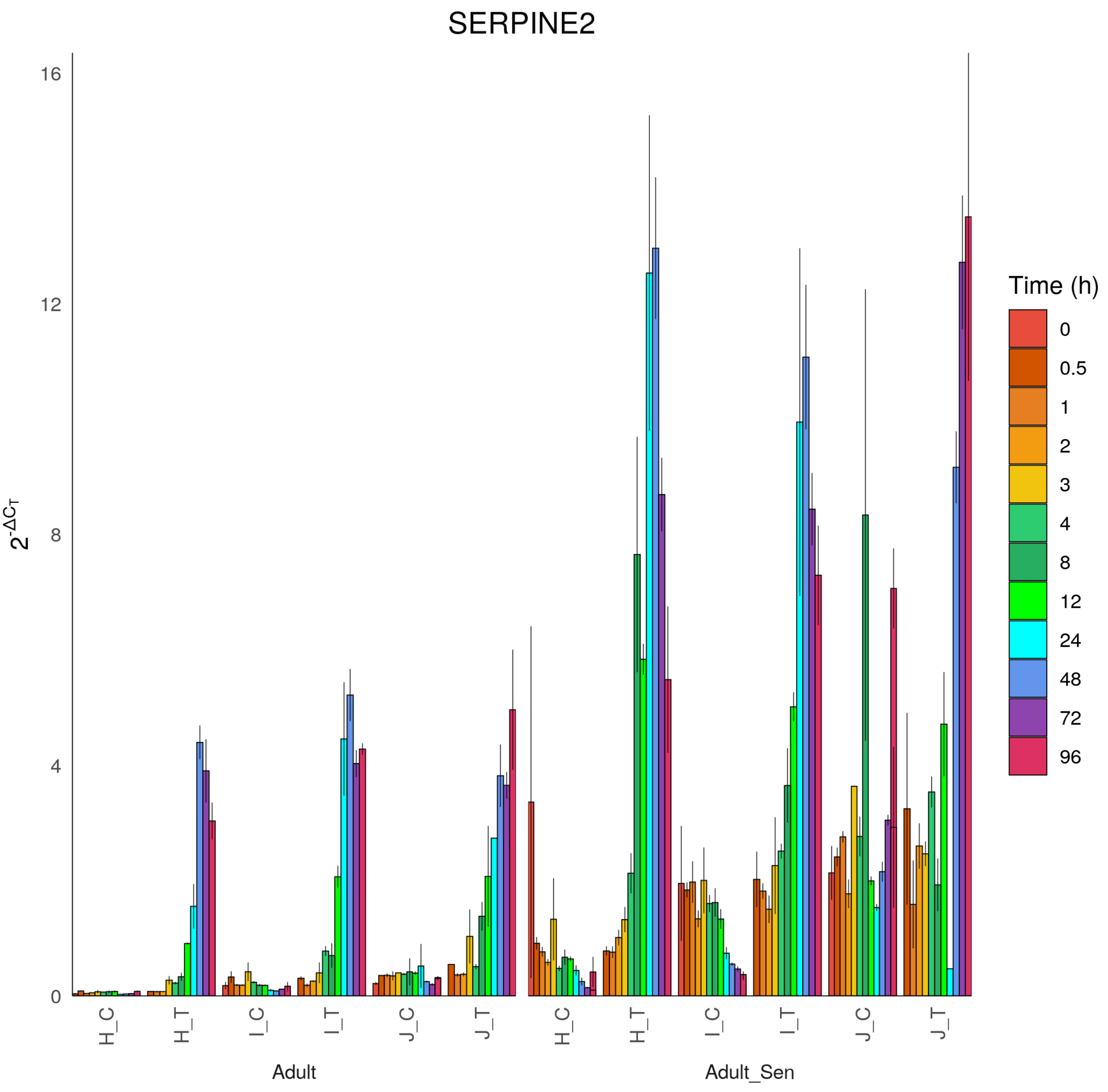

SERPINE2
$2^{-\Delta C_T}$
16
12
8
4
0
H_C
H_T
I_C
I_T
J_C
J_T
Adult
Adult_Sen
Time (h)
0
0.5
1
2
3
4
8
12
24
48
72
96

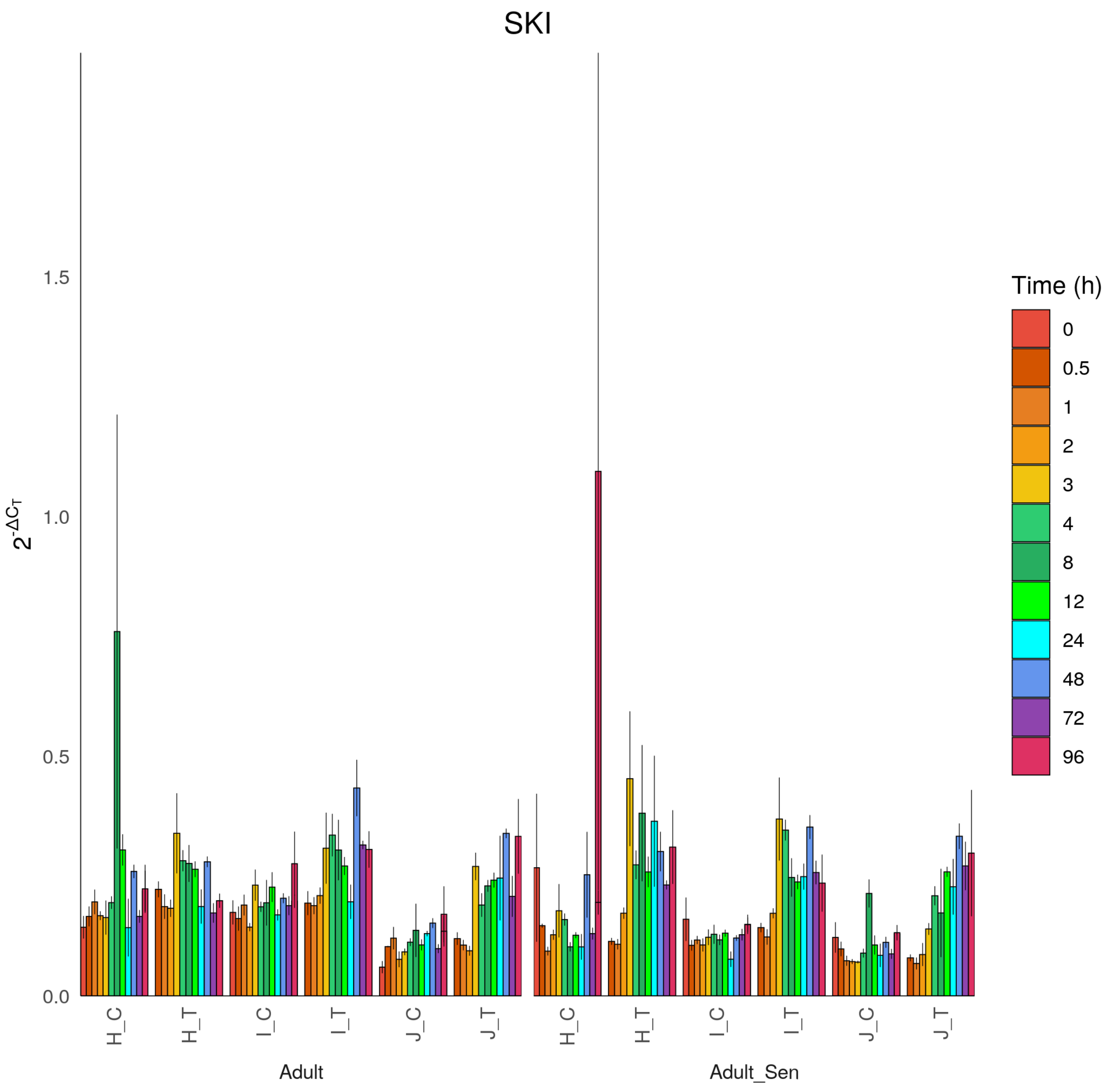
SKI
$2^{-\Delta C_T}$
0.0
0.5
1.0
1.5
H_C
H_T
I_C
I_T
J_C
J_T
Adult
Adult_Sen
Time (h)
0
0.5
1
2
3
4
8
12
24
48
72
96

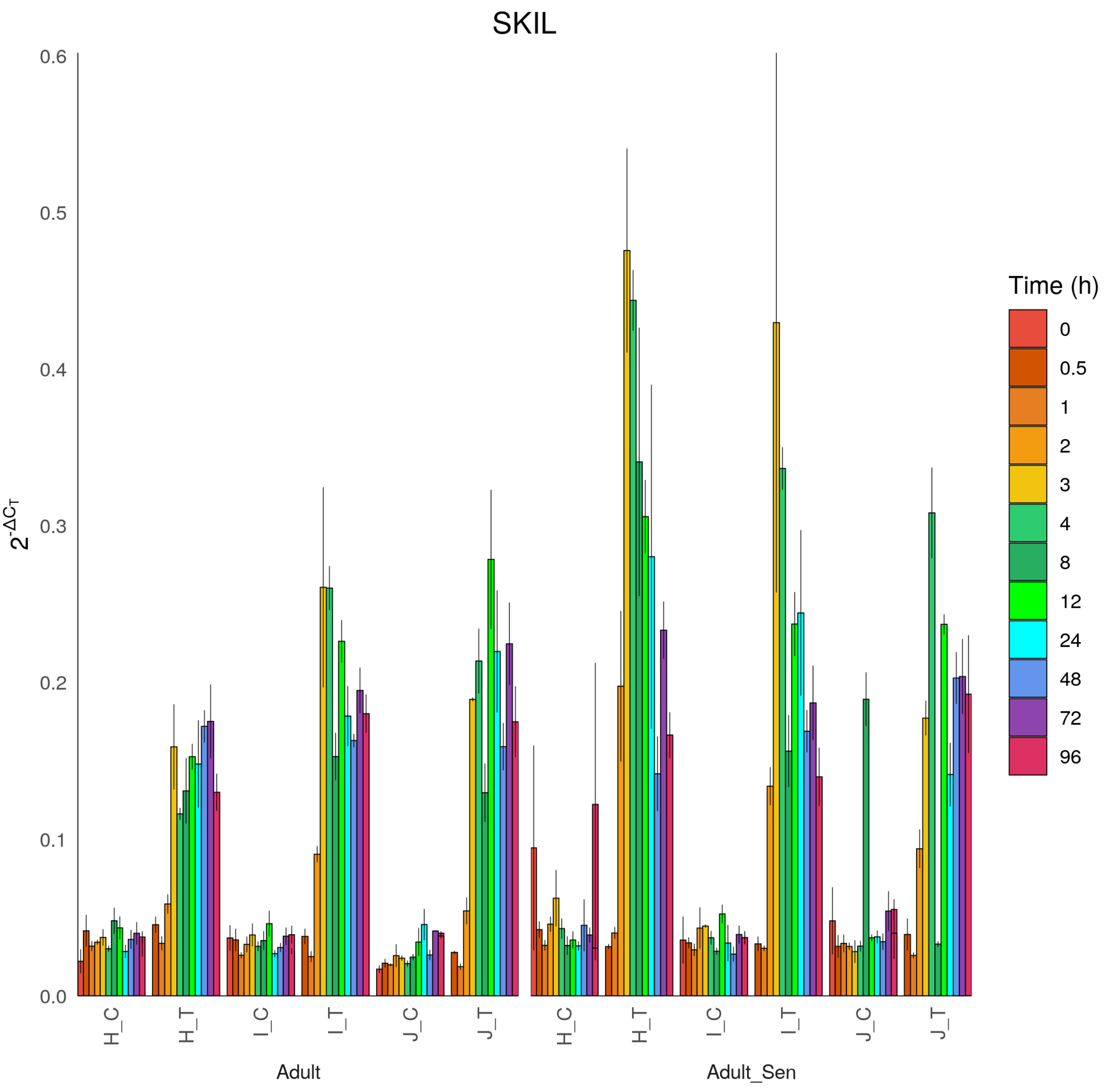
SKIL
$2^{-\Delta C_T}$
0.0
0.1
0.2
0.3
0.4
0.5
0.6
H_C
H_T
I_C
I_T
J_C
J_T
Adult
Adult_Sen
Time (h)
0
0.5
1
2
3
4
8
12
24
48
72
96

SMAD3

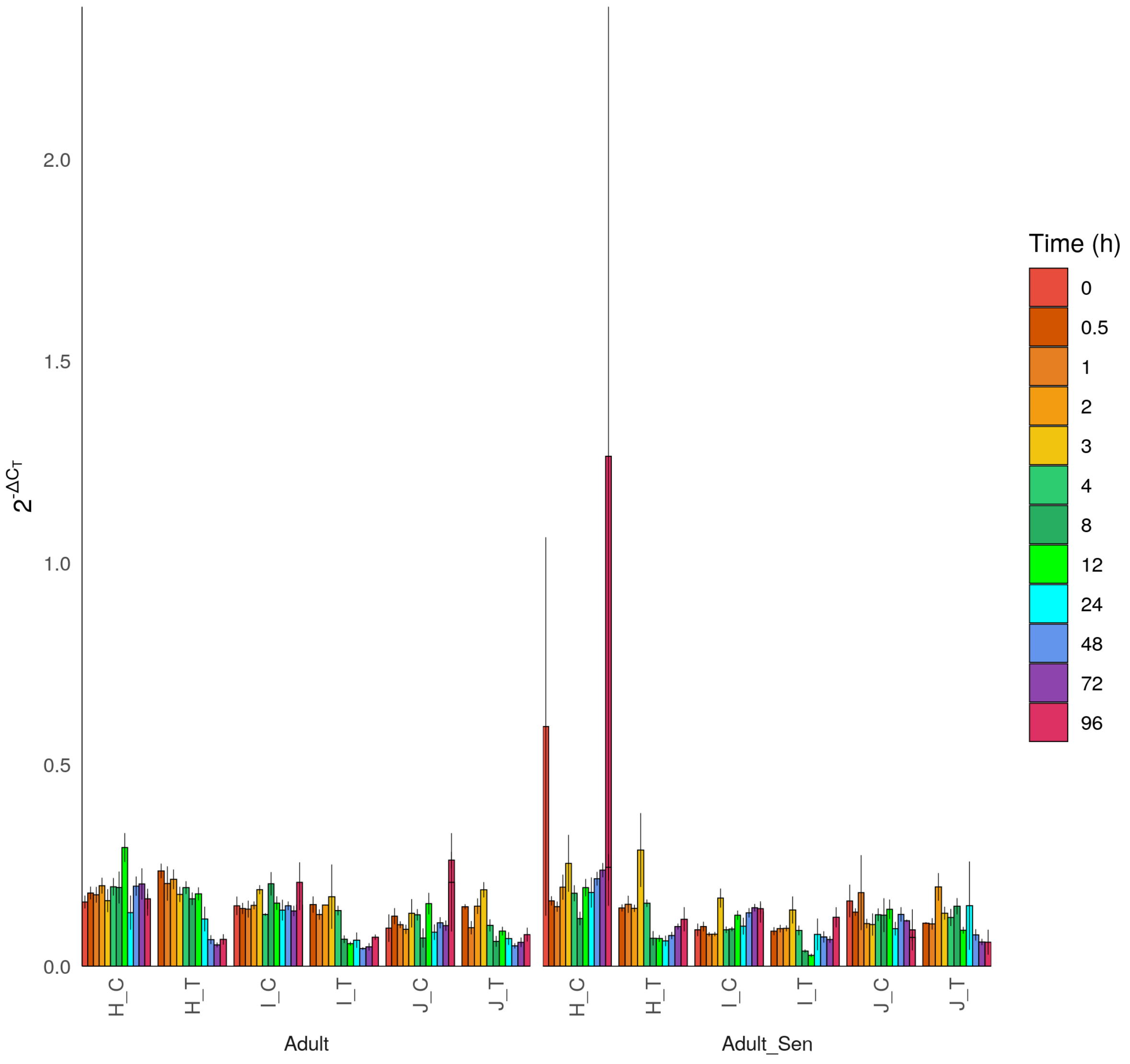

0.0
0.5
1.0
1.5
2.0
$2^{-\Delta C_T}$
H_C
H_T
I_C
I_T
J_C
J_T
Adult
Adult_Sen
Time (h)
0
0.5
1
2
3
4
8
12
24
48
72
96

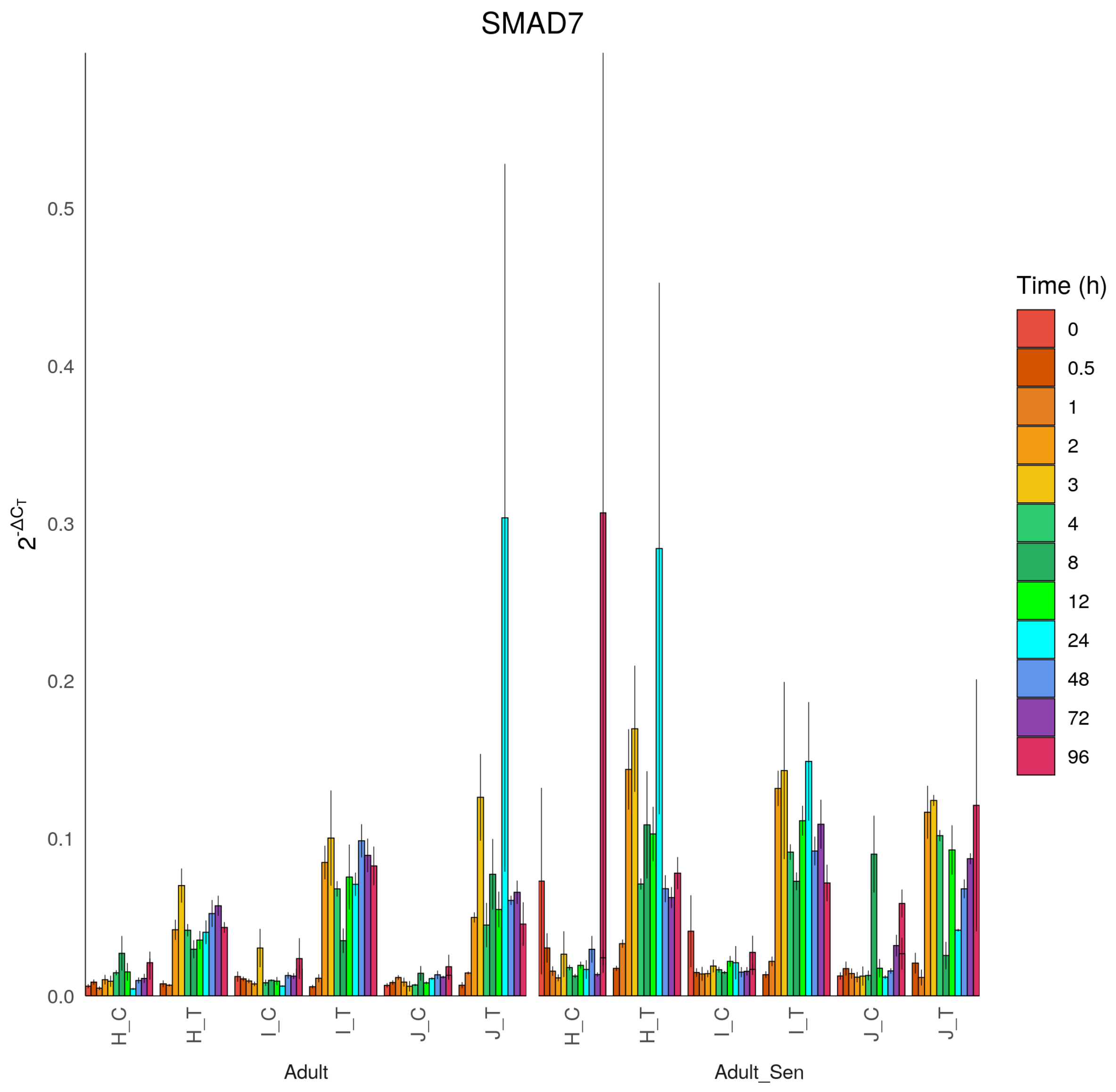
SMAD7
$2^{-\Delta C_T}$
0.0
0.1
0.2
0.3
0.4
0.5
H_C
H_T
I_C
I_T
J_C
J_T
Adult
Adult_Sen
Time (h)
0
0.5
1
2
3
4
8
12
24
48
72
96

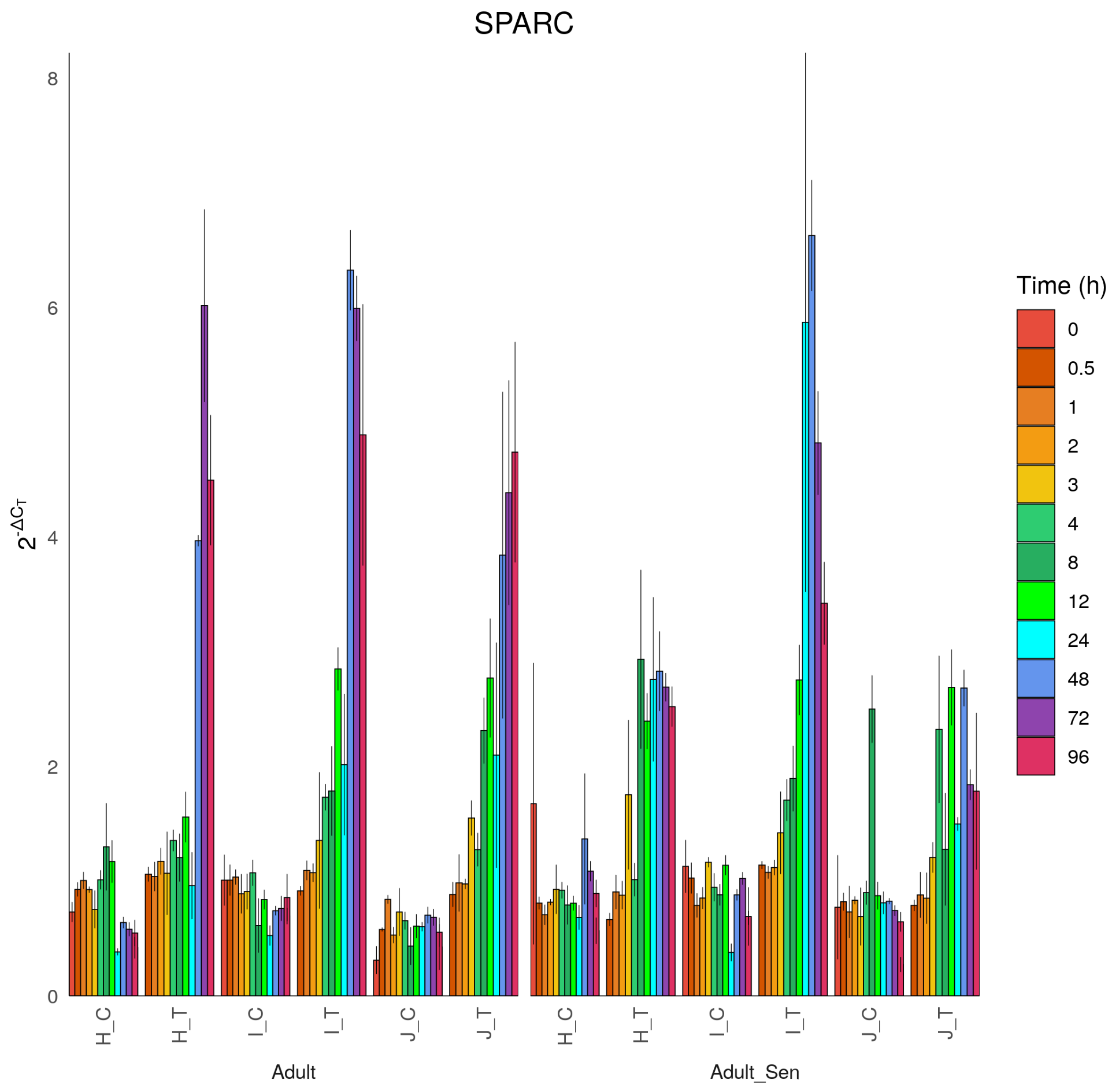
SPARC
$2^{-\Delta C_T}$
0
2
4
6
8
H_C
H_T
I_C
I_T
J_C
J_T
Adult
Adult_Sen
Time (h)
0
0.5
1
2
3
4
8
12
24
48
72
96

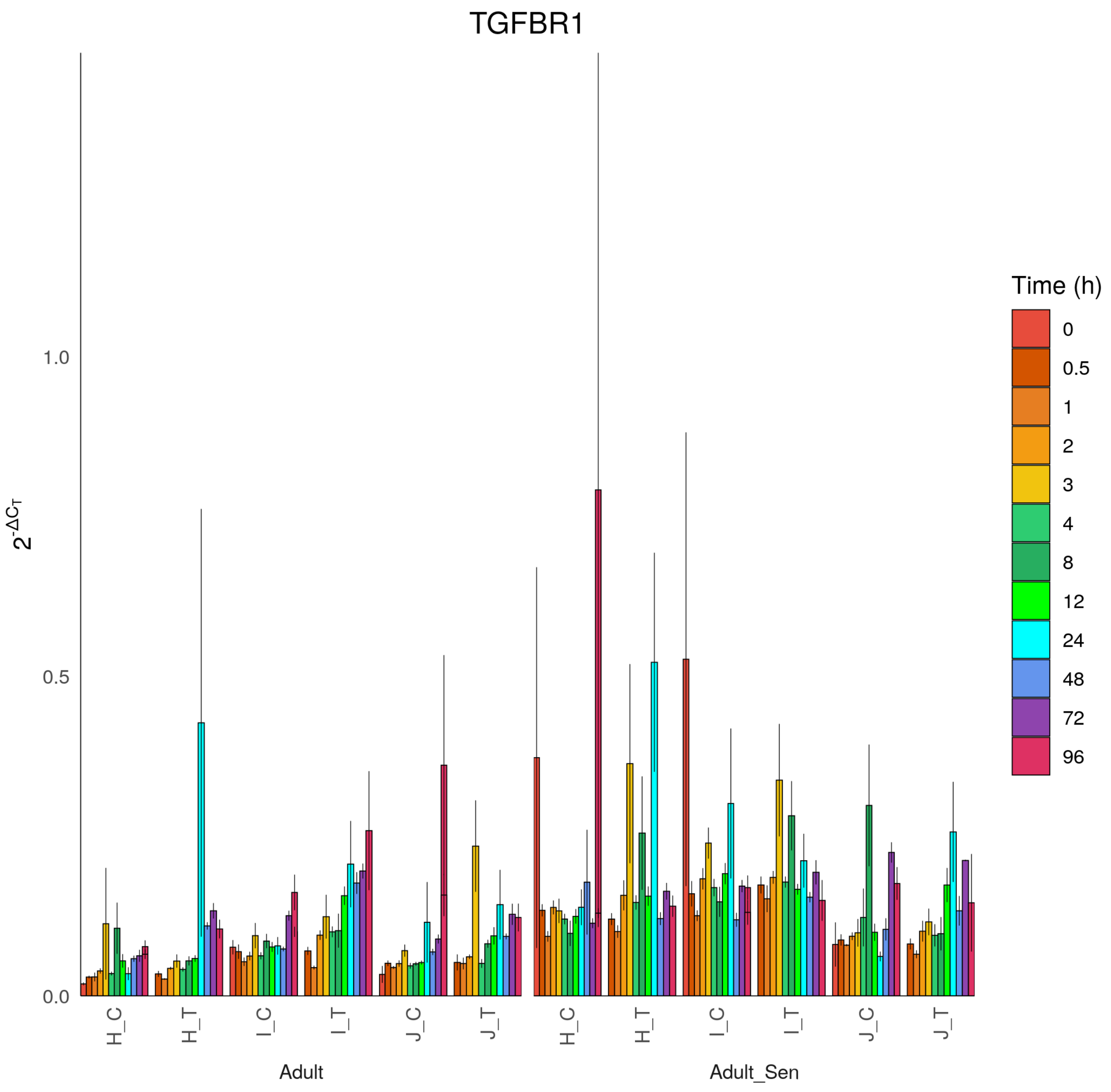
TGFBR1
$2^{-\Delta C_T}$
0.0
0.5
1.0
H_C
H_T
I_C
I_T
J_C
J_T
Adult
Adult_Sen
Time (h)
0
0.5
1
2
3
4
8
12
24
48
72
96

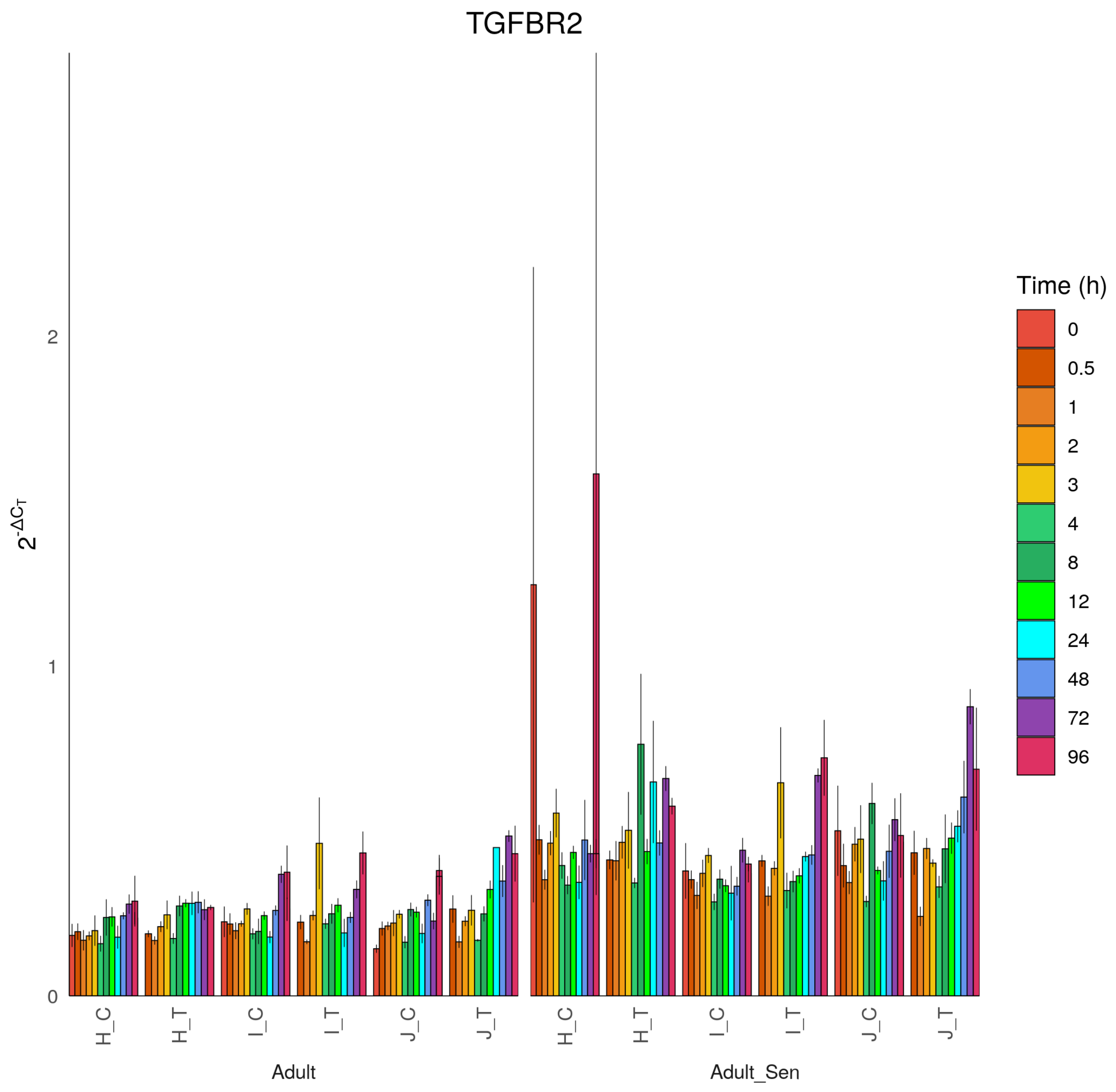

TGFBR2
$2^{-\Delta C_T}$
0
1
2
H_C
H_T
I_C
I_T
J_C
J_T
Adult
Adult_Sen
Time (h)
0
0.5
1
2
3
4
8
12
24
48
72
96

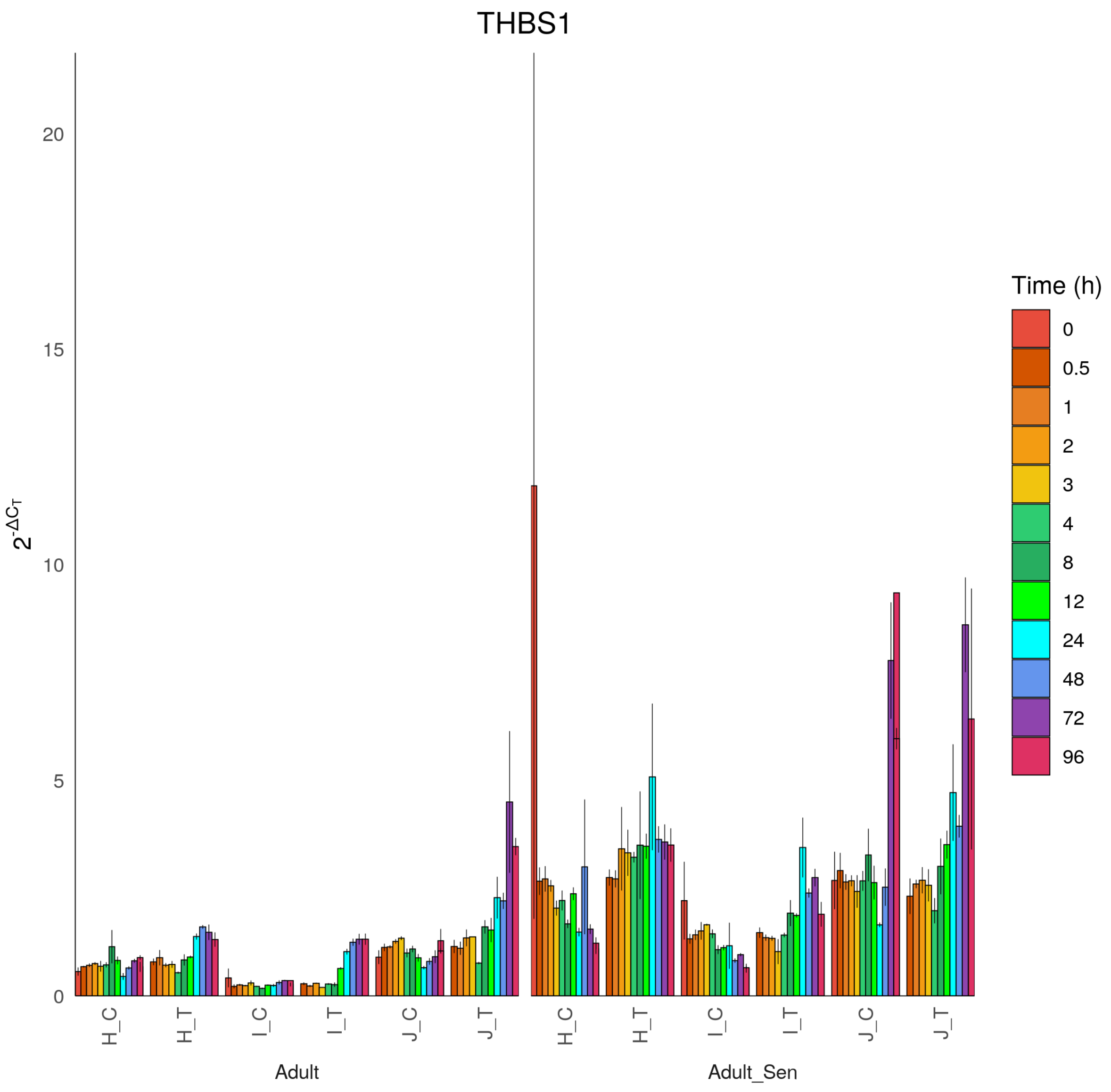
THBS1
$2^{-\Delta C_T}$
0
5
10
15
20
H_C
H_T
I_C
I_T
J_C
J_T
Adult
Adult_Sen
Time (h)
0
0.5
1
2
3
4
8
12
24
48
72
96

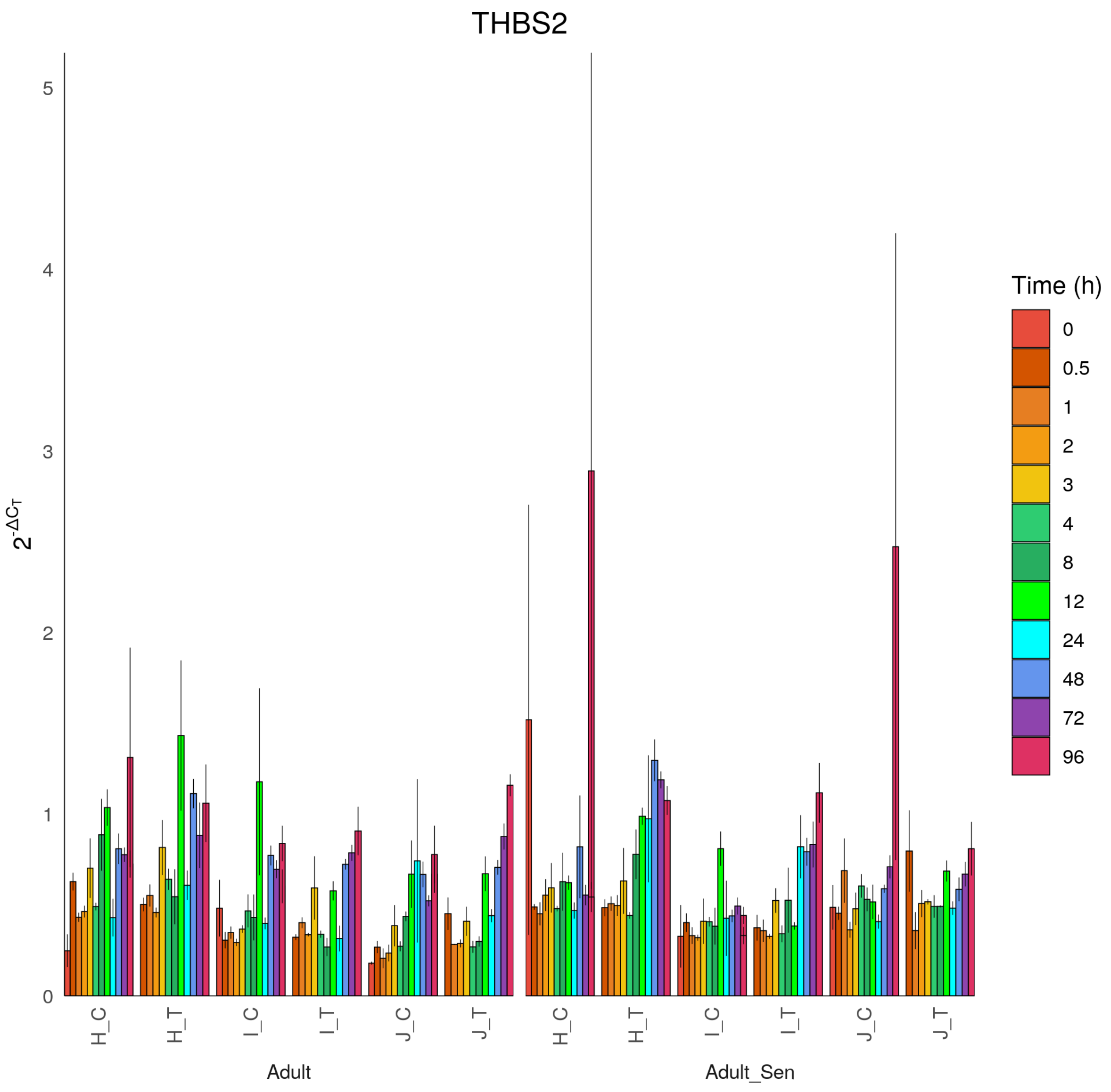

THBS2
$2^{-\Delta C_T}$
0
1
2
3
4
5
H_C
H_T
I_C
I_T
J_C
J_T
H_C
H_T
I_C
I_T
J_C
J_T
Adult
Adult_Sen
Time (h)
0
0.5
1
2
3
4
8
12
24
48
72
96

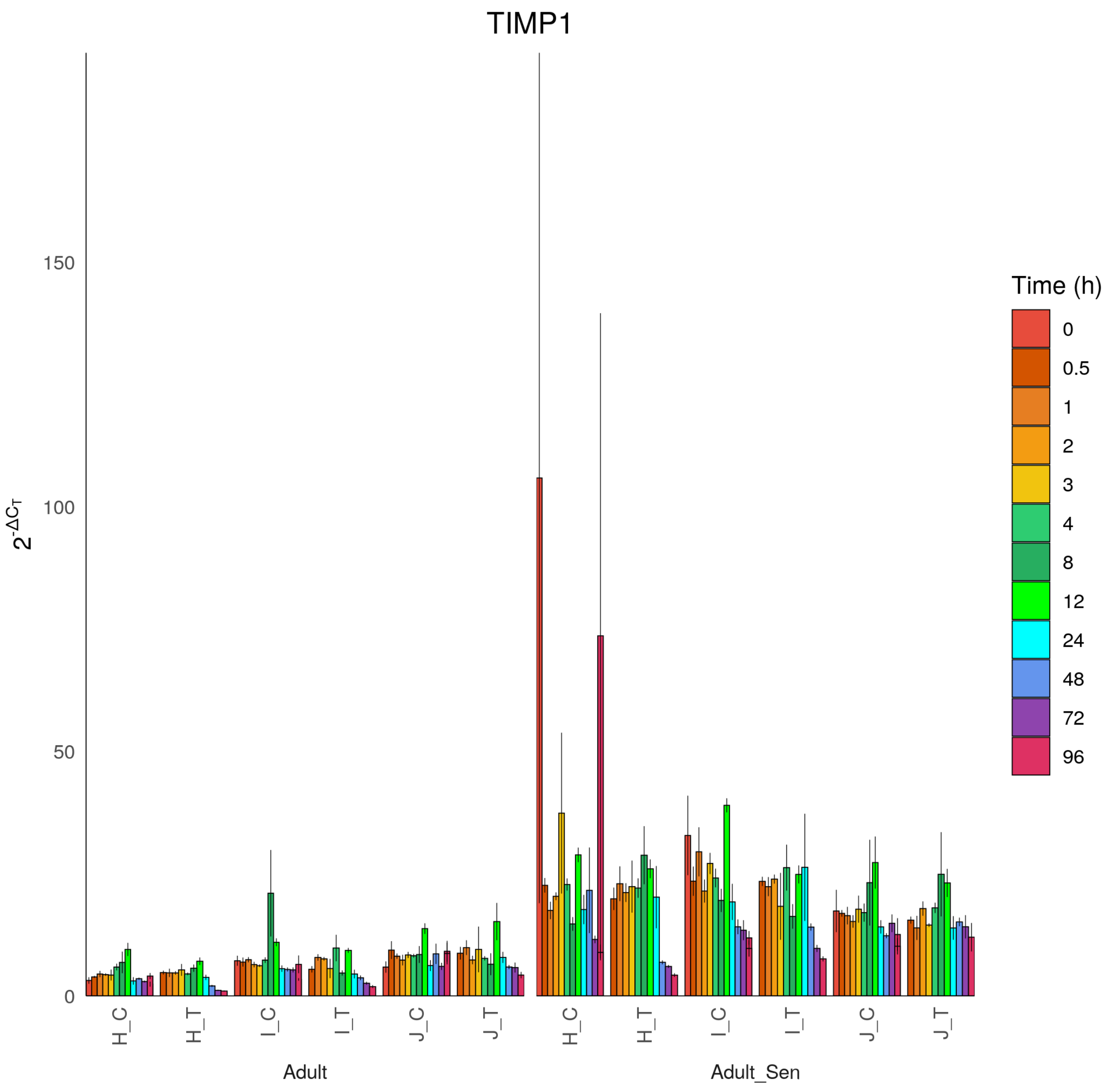

TIMP1
$2^{-\Delta C_T}$
0
50
100
150
H_C
H_T
I_C
I_T
J_C
J_T
Adult
H_C
H_T
I_C
I_T
J_C
J_T
Adult_Sen
Time (h)
0
0.5
1
2
3
4
8
12
24
48
72
96

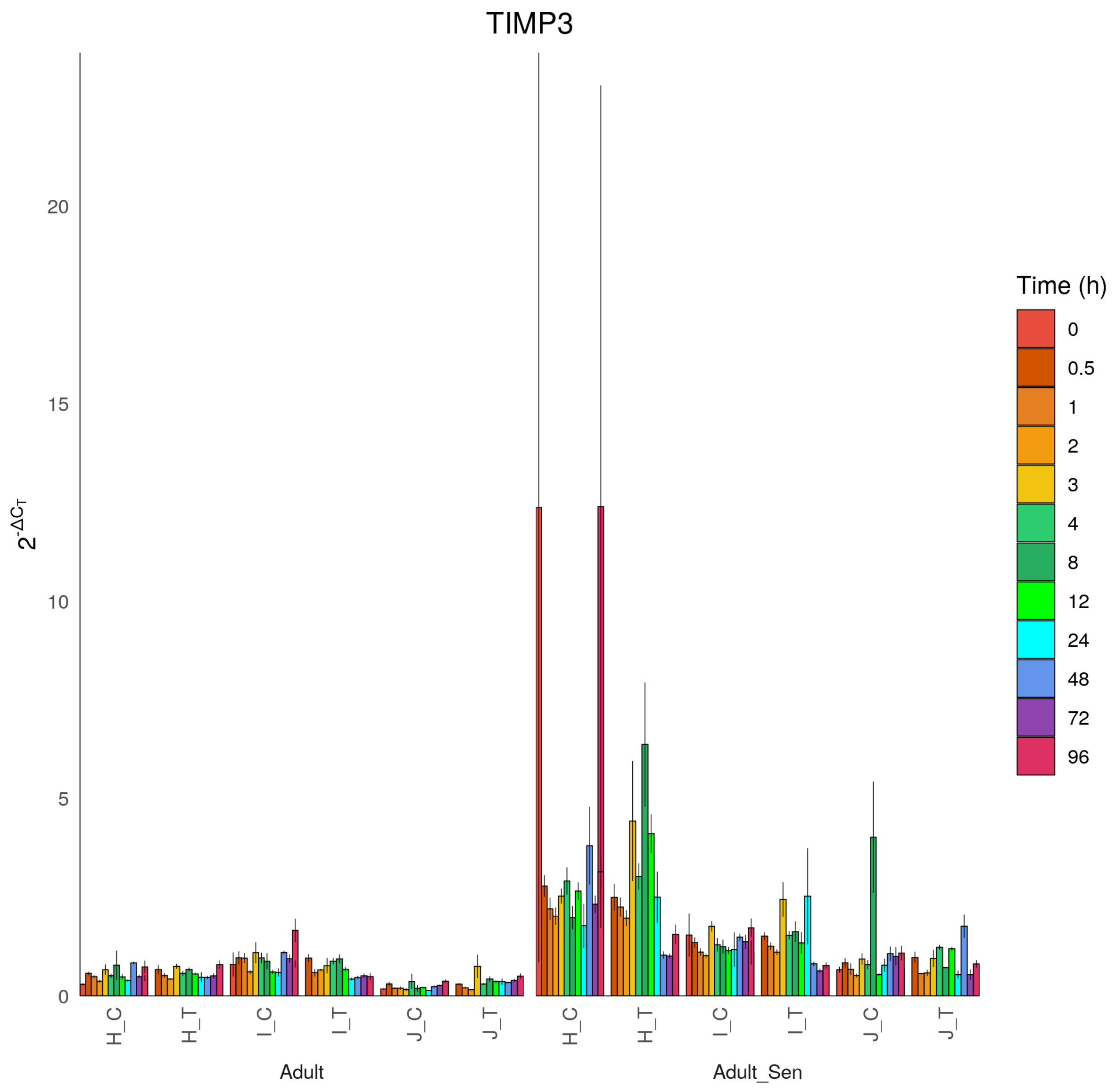

TIMP3
$2^{-\Delta C_T}$
0
5
10
15
20
H_C
H_T
I_C
I_T
J_C
J_T
Adult
Adult_Sen
Time (h)
0
0.5
1
2
3
4
8
12
24
48
72
96

TP53BP1

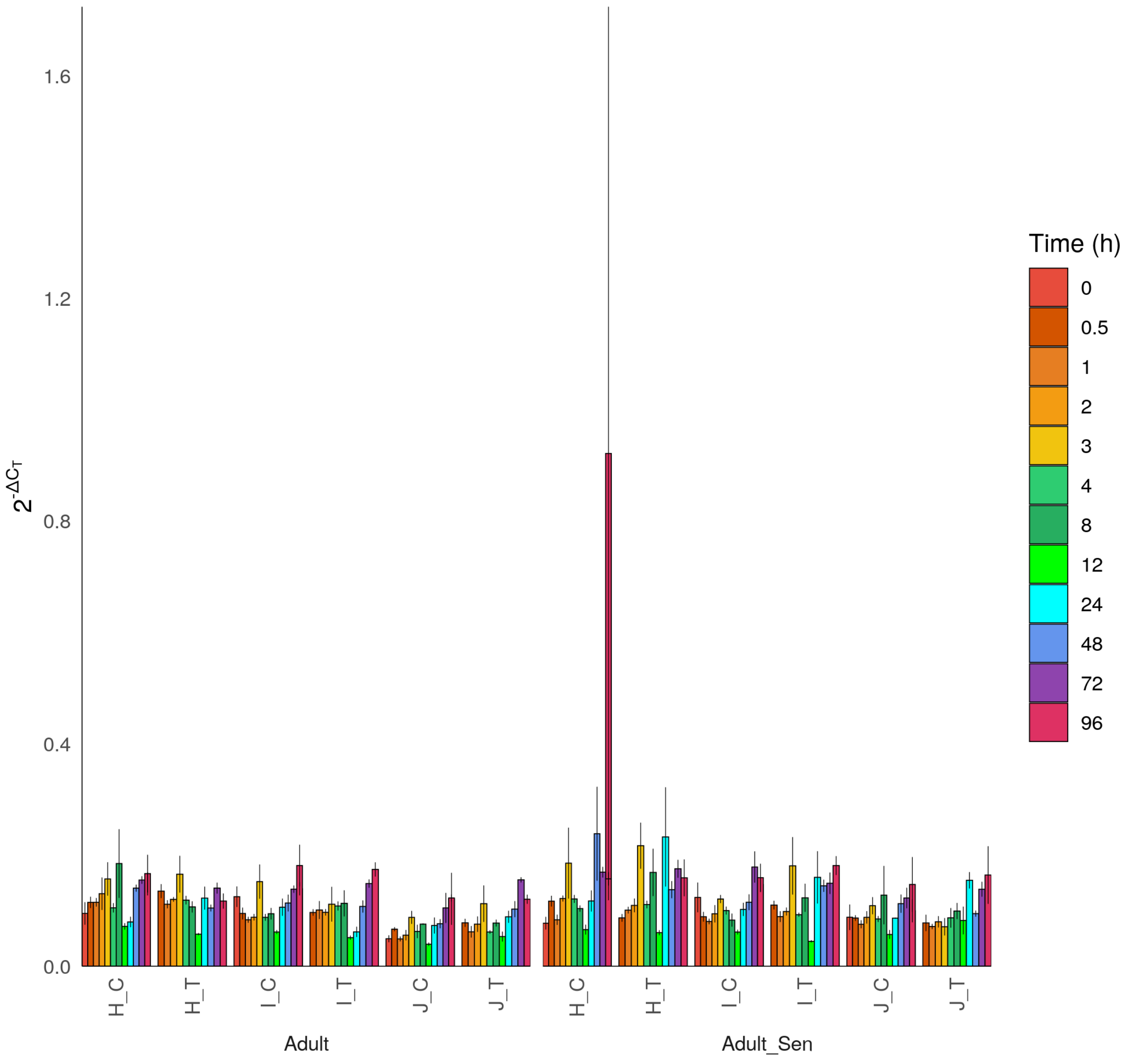

1.6
1.2
0.8
0.4
0.0
$2^{-\Delta C_T}$
H_C
H_T
I_C
I_T
J_C
J_T
Adult
Adult_Sen
Time (h)
0
0.5
1
2
3
4
8
12
24
48
72
96

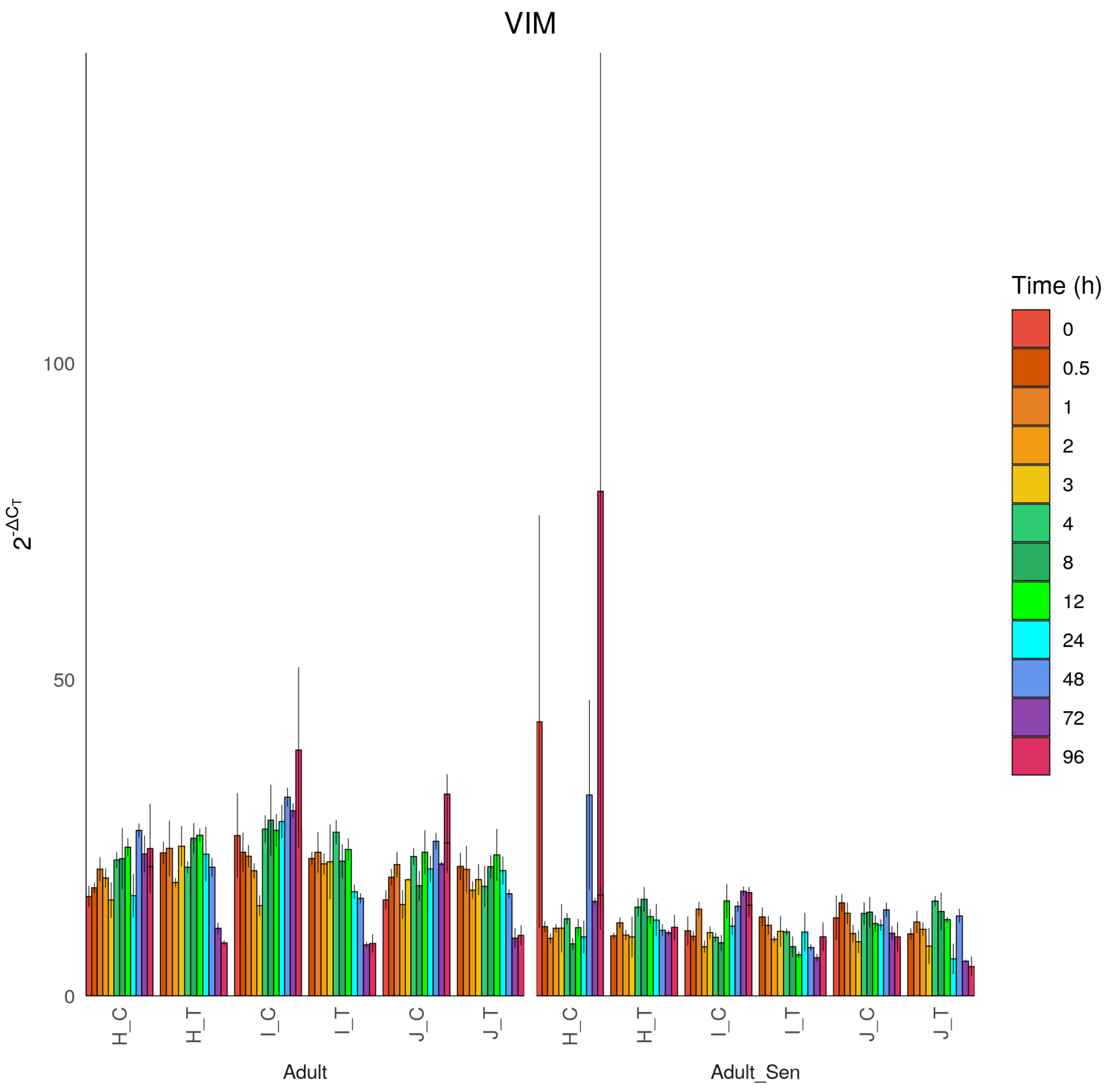
VIM
$2^{-\Delta C_T}$
0
50
100
H_C
H_T
I_C
I_T
J_C
J_T
Adult
Adult_Sen
Time (h)
0
0.5
1
2
3
4
8
12
24
48
72
96
