## Supplementary file 4 for "Cell Senescence-Independent Ageing of Human Skin"

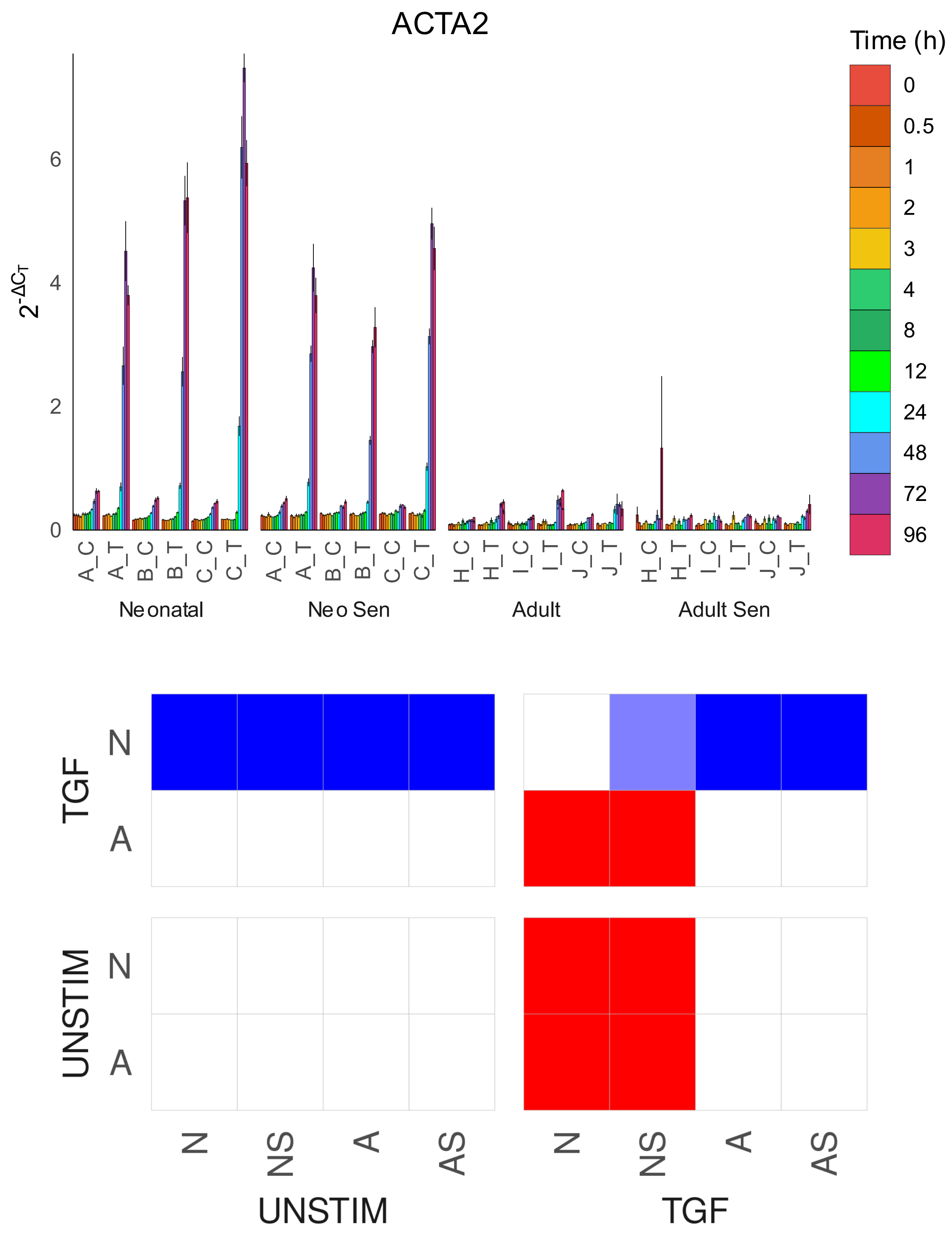

ACTA2
Time (h)
0
0.5
1
2
3
4
8
12
24
48
72
96
$2^{-\Delta C_T}$
0
2
4
6
A_C
A_T
B_C
B_T
C_C
C_T
H_C
H_T
I_C
I_T
J_C
J_T
Neonatal
Neo Sen
Adult
Adult Sen
TGF
UNSTIM
N
A
N
NS
A
AS
UNSTIM
TGF

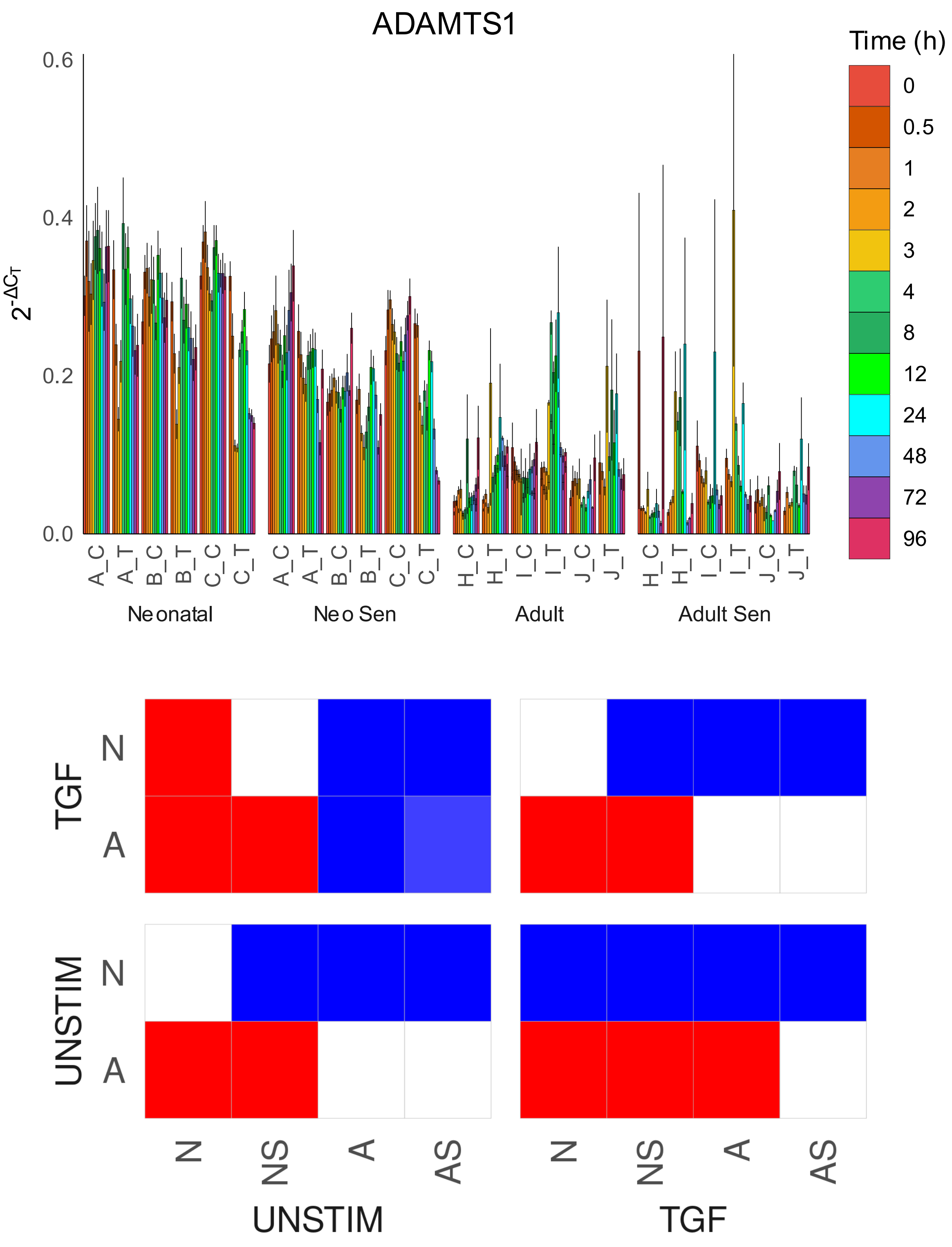

ADAMTS1
Time (h)
0
0.5
1
2
3
4
8
12
24
48
72
96
0.6
0.4
0.2
0.0
$2^{-\Delta C_T}$
A_C
A_T
B_C
B_T
C_C
C_T
H_C
H_T
I_C
I_T
J_C
J_T
Neonatal
Neo Sen
Adult
Adult Sen
TGF
UNSTIM
N
A
N
NS
A
AS
UNSTIM
TGF

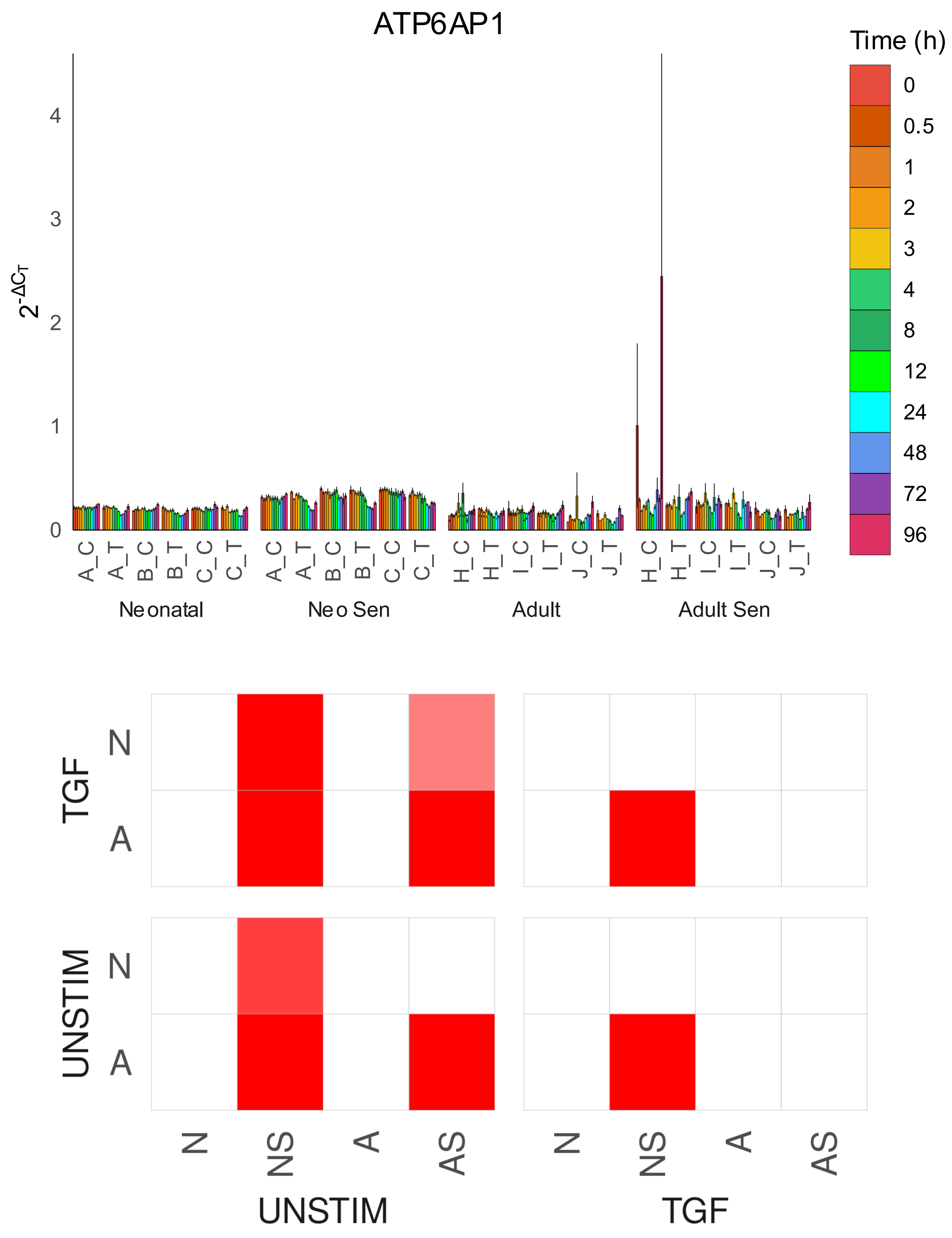

ATP6AP1
Time (h)
0
0.5
1
2
3
4
8
12
24
48
72
96
$2^{-\Delta C_T}$
Neonatal
Neo Sen
Adult
Adult Sen
A_C
A_T
B_C
B_T
C_C
C_T
H_C
H_T
I_C
I_T
J_C
J_T
TGF
UNSTIM
N
A
NS
AS

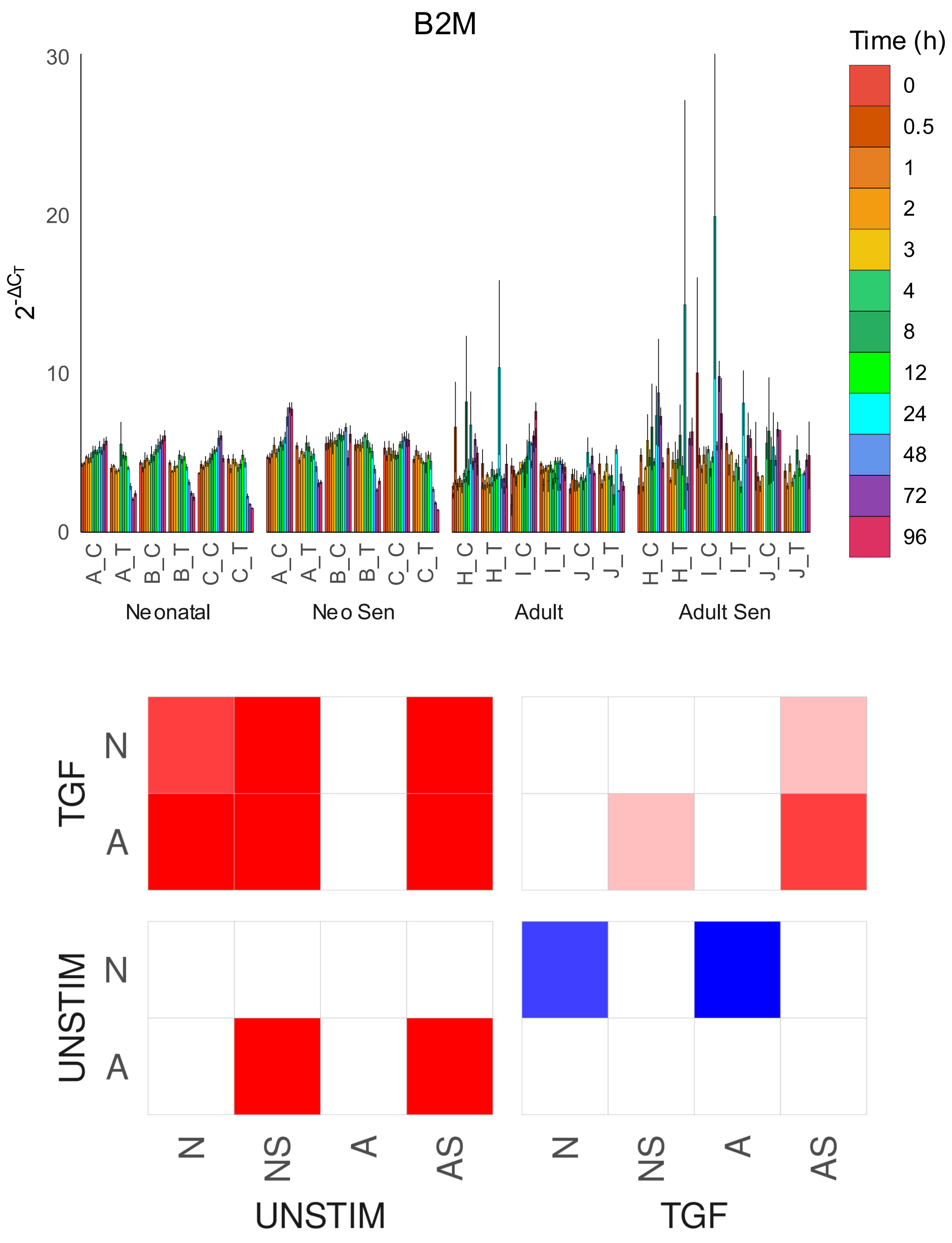

B2M
Time (h)
0
0.5
1
2
3
4
8
12
24
48
72
96
30
20
10
0
A_C
A_T
B_C
B_T
C_C
C_T
H_C
H_T
I_C
I_T
J_C
J_T
Neonatal
Neo Sen
Adult
Adult Sen
TGF
UNSTIM
N
A
N
NS
A
AS
UNSTIM
TGF

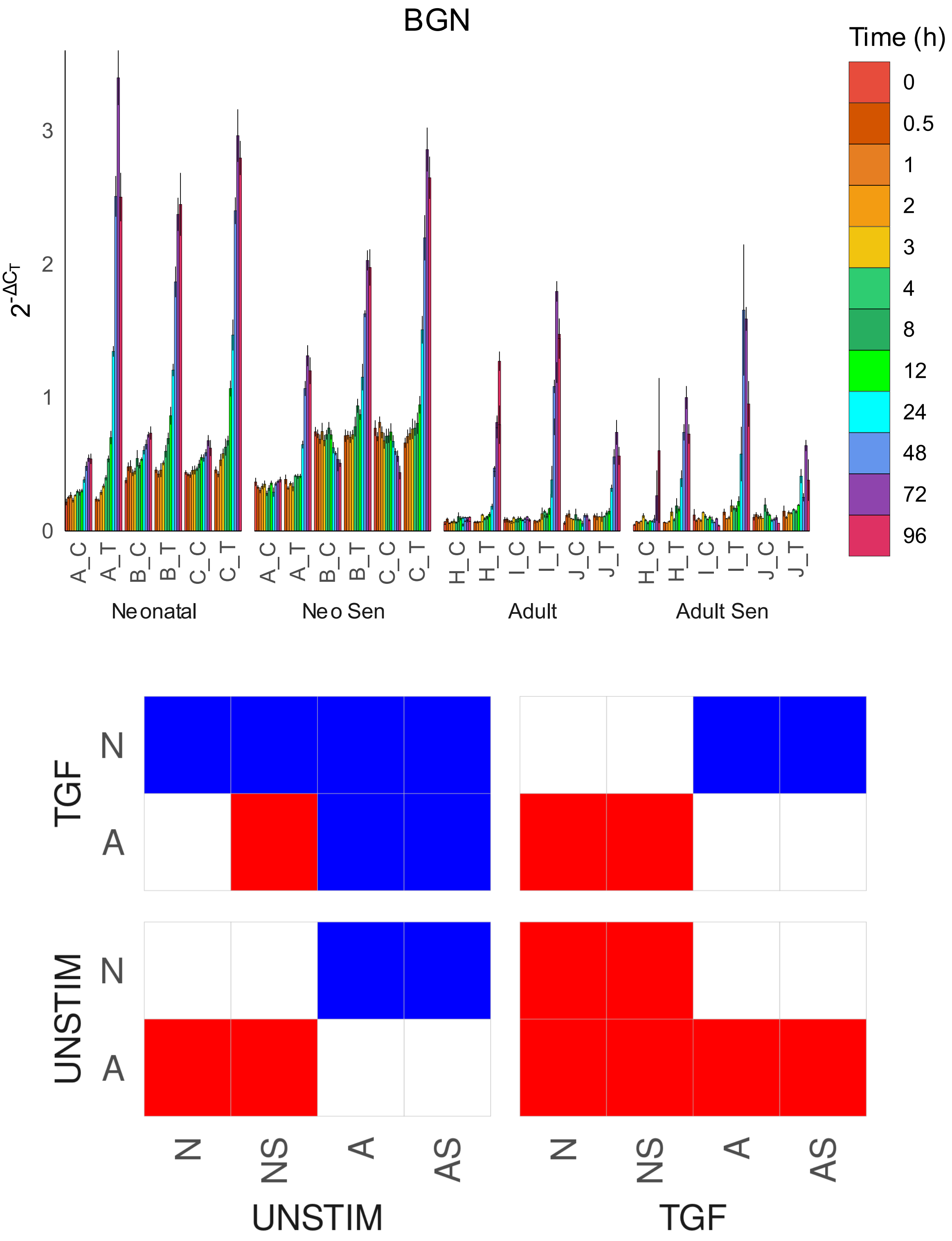

BGN
Time (h)
0
0.5
1
2
3
4
8
12
24
48
72
96
$2^{-\Delta C_T}$
0
1
2
3
A_C
A_T
B_C
B_T
C_C
C_T
H_C
H_T
I_C
I_T
J_C
J_T
Neonatal
Neo Sen
Adult
Adult Sen
TGF
UNSTIM
N
A
N
NS
A
AS
UNSTIM
TGF

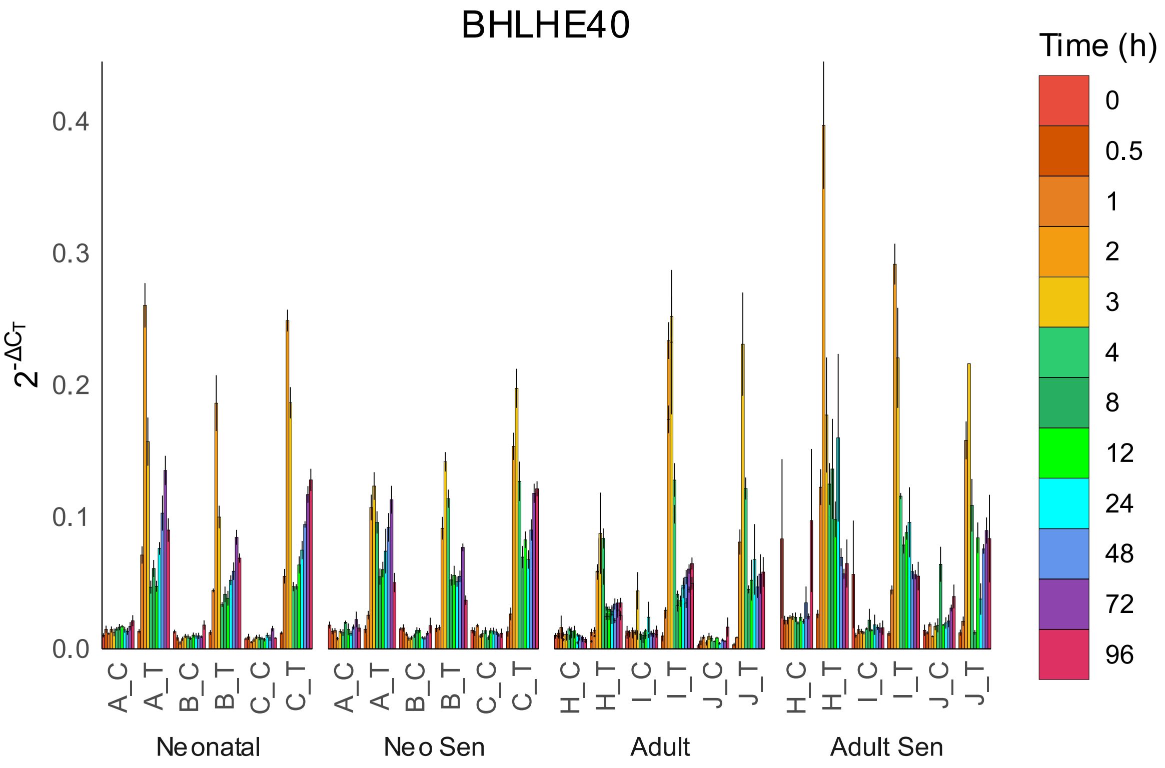

BHLHE40
Time (h)
0
0.5
1
2
3
4
8
12
24
48
72
96
$2^{-\Delta C_T}$
0.4
0.3
0.2
0.1
0.0
A_C
A_T
B_C
B_T
C_C
C_T
H_C
H_T
I_C
I_T
J_C
J_T
Neonatal
Neo Sen
Adult
Adult Sen


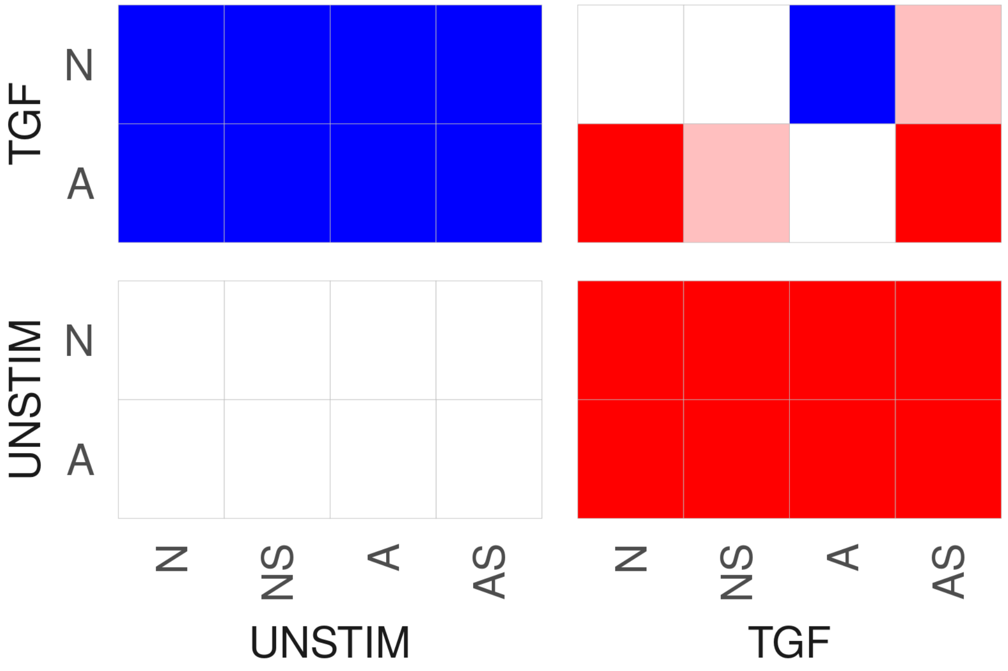

TGF
N
A
UNSTIM
N
A
N
NS
A
AS
N
NS
A
AS
UNSTIM
TGF

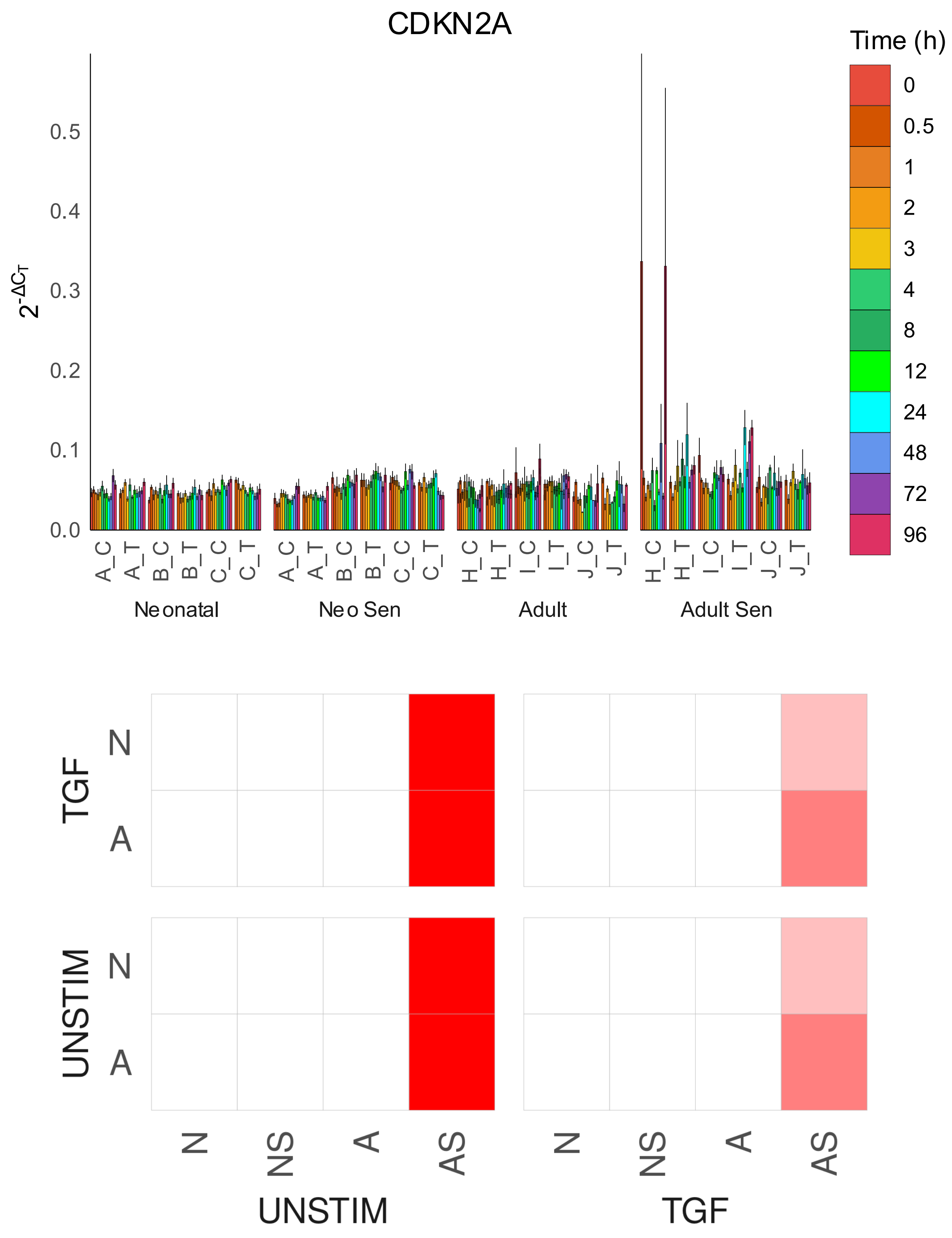

CDKN2A
Time (h)
0
0.5
1
2
3
4
8
12
24
48
72
96
$2^{-\Delta C_T}$
0.0
0.1
0.2
0.3
0.4
0.5
A_C
A_T
B_C
B_T
C_C
C_T
H_C
H_T
I_C
I_T
J_C
J_T
Neonatal
Neo Sen
Adult
Adult Sen
TGF
UNSTIM
N
A
N
NS
A
AS
UNSTIM
TGF

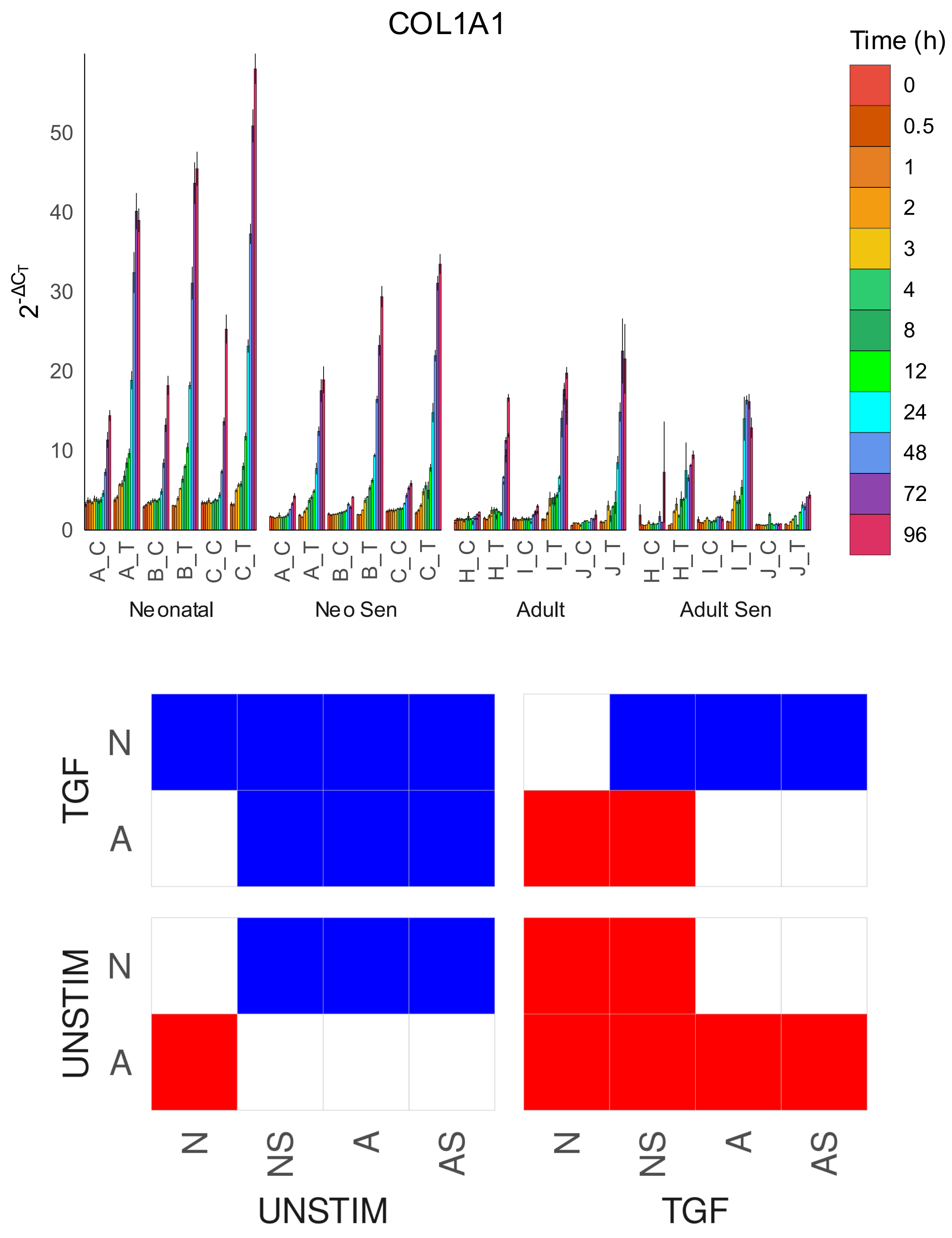

COL1A1
Time (h)
0
0.5
1
2
3
4
8
12
24
48
72
96
50
40
30
20
10
0
$2^{-\Delta C_T}$
A_C
A_T
B_C
B_T
C_C
C_T
H_C
H_T
I_C
I_T
J_C
J_T
Neonatal
Neo Sen
Adult
Adult Sen
TGF
UNSTIM
N
A
N
NS
A
AS
UNSTIM
TGF

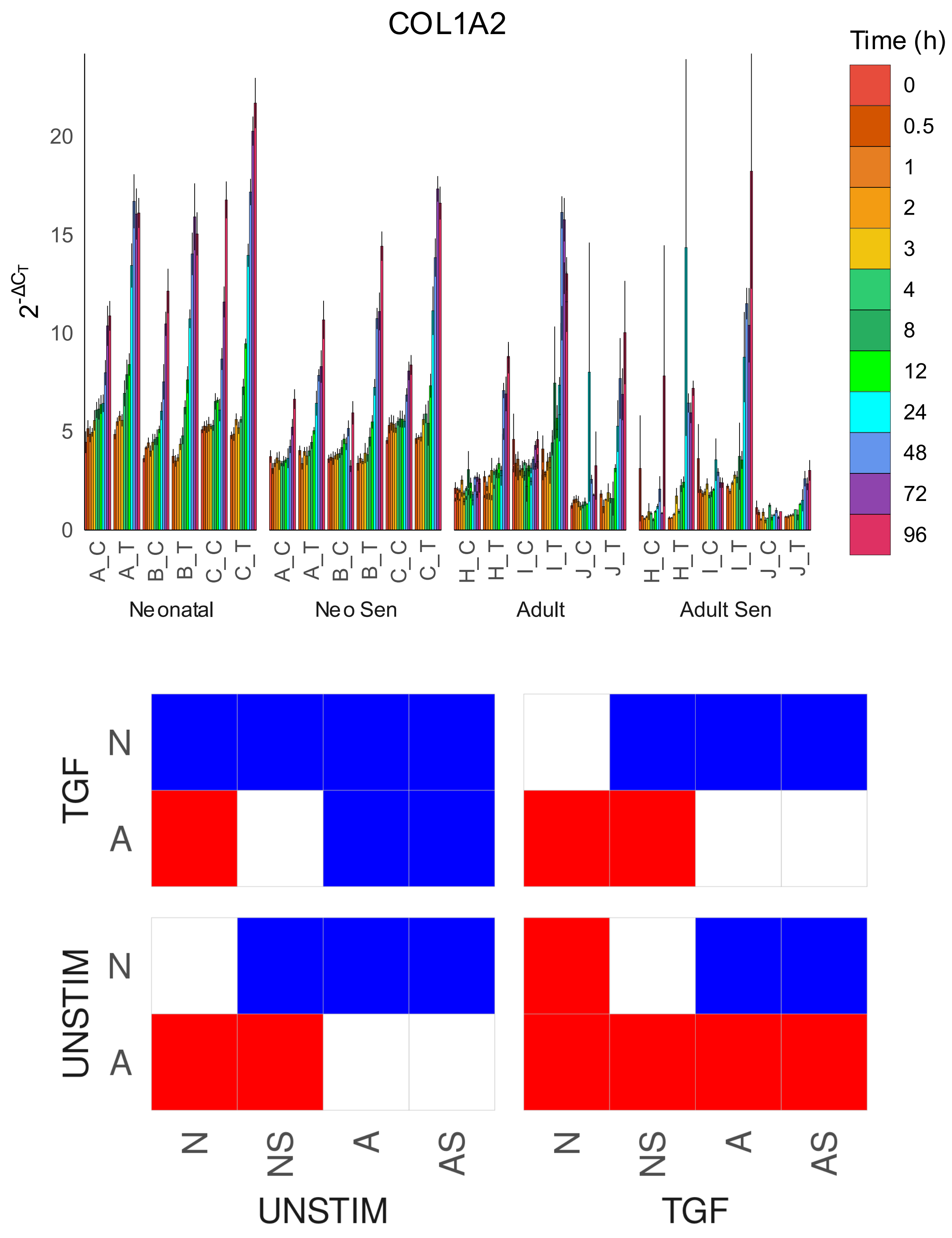

COL1A2
Time (h)
0
0.5
1
2
3
4
8
12
24
48
72
96
2-ΔCT
0
5
10
15
20
A_C
A_T
B_C
B_T
C_C
C_T
H_C
H_T
I_C
I_T
J_C
J_T
Neonatal
Neo Sen
Adult
Adult Sen
TGF
UNSTIM
N
A
N
NS
A
AS
UNSTIM
TGF

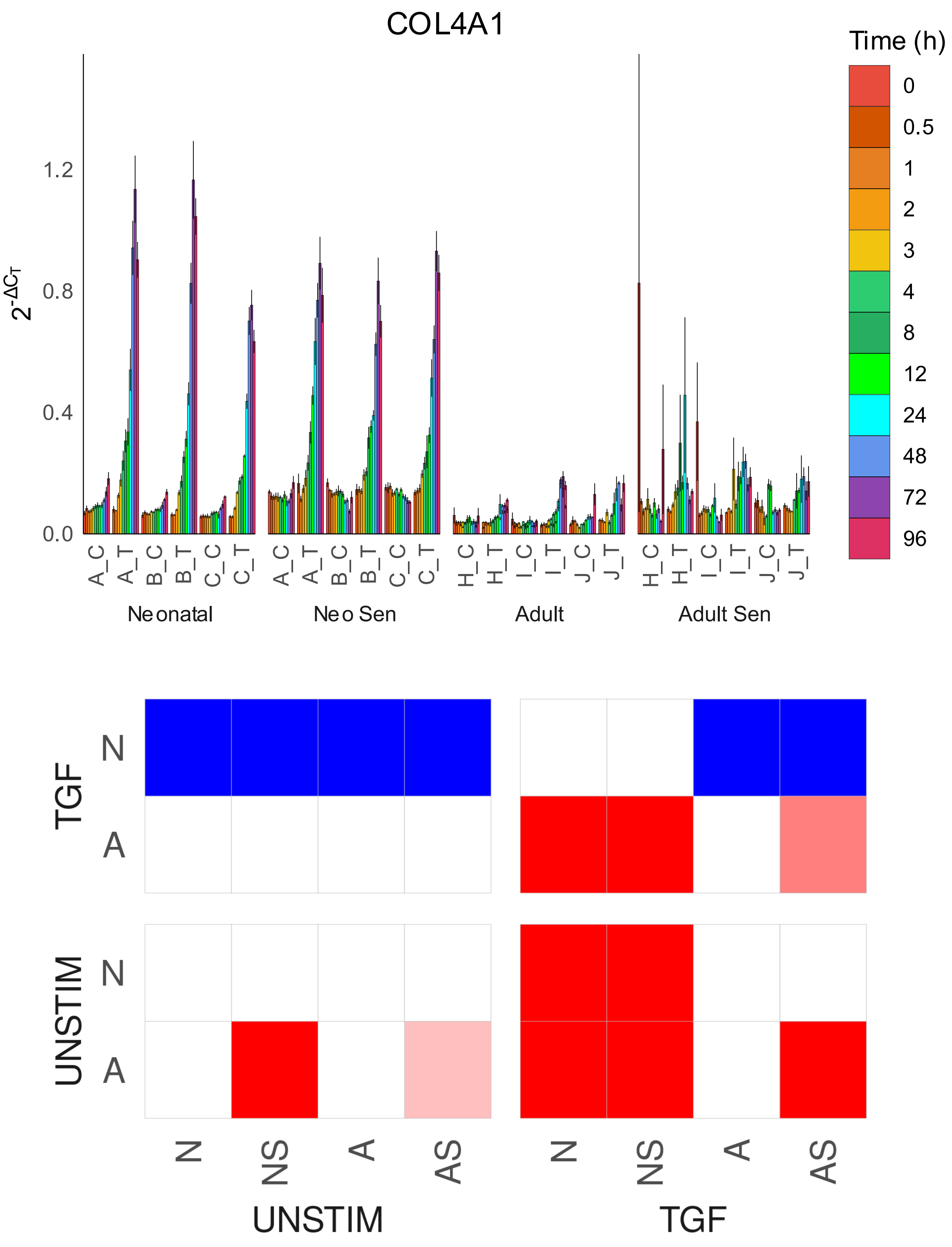
COL4A1
Time (h)
0
0.5
1
2
3
4
8
12
24
48
72
96
1.2
0.8
0.4
0.0
A_C
A_T
B_C
B_T
C_C
C_T
H_C
H_T
I_C
I_T
J_C
J_T
Neonatal
Neo Sen
Adult
Adult Sen
TGF
UNSTIM
N
A
N
NS
A
AS
UNSTIM
TGF

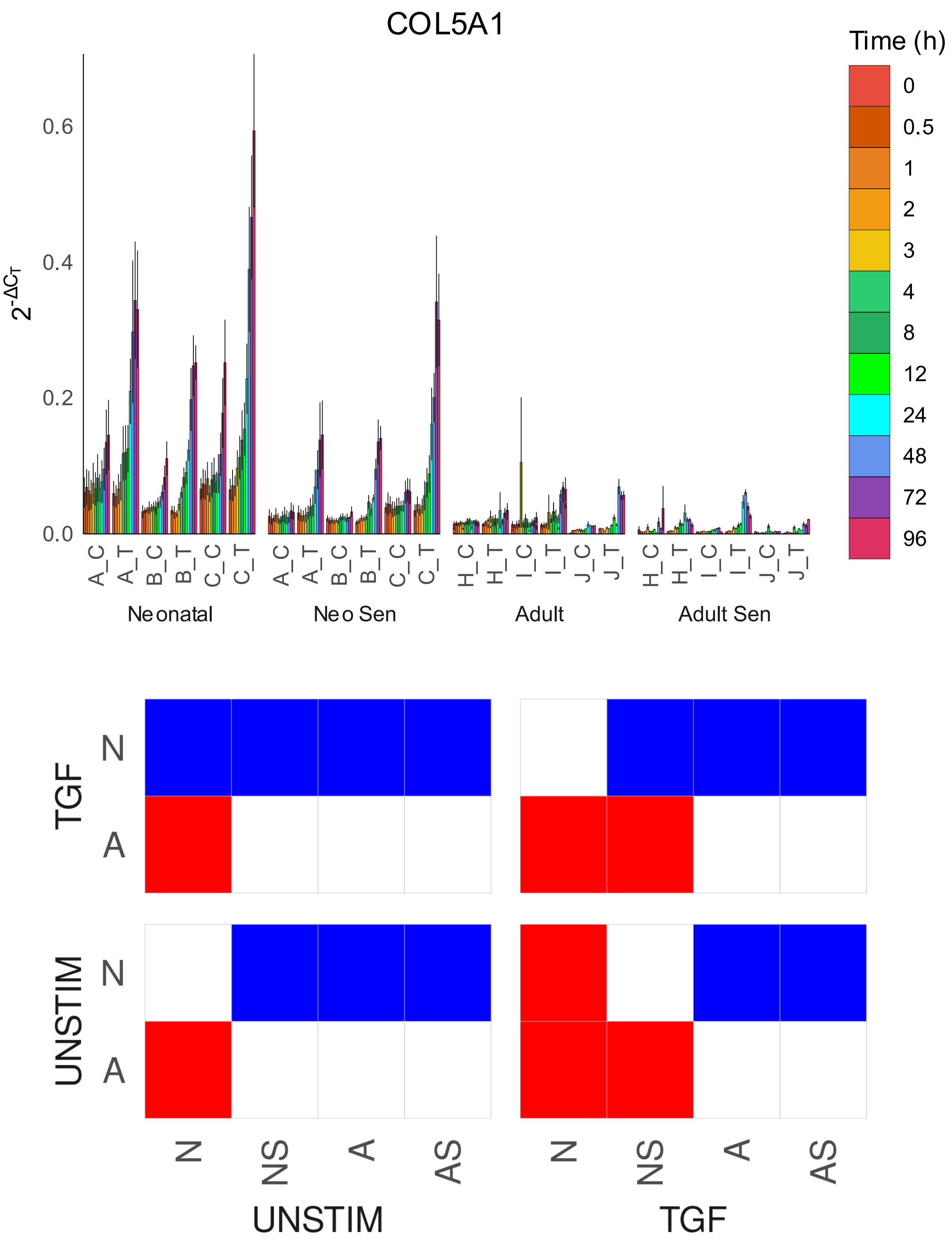
COL5A1
Time (h)
0
0.5
1
2
3
4
8
12
24
48
72
96
$2^{-\Delta C_T}$
0.0
0.2
0.4
0.6
A_C
A_T
B_C
B_T
C_C
C_T
H_C
H_T
I_C
I_T
J_C
J_T
Neonatal
Neo Sen
Adult
Adult Sen
TGF
UNSTIM
N
A
N
NS
A
AS
UNSTIM
TGF

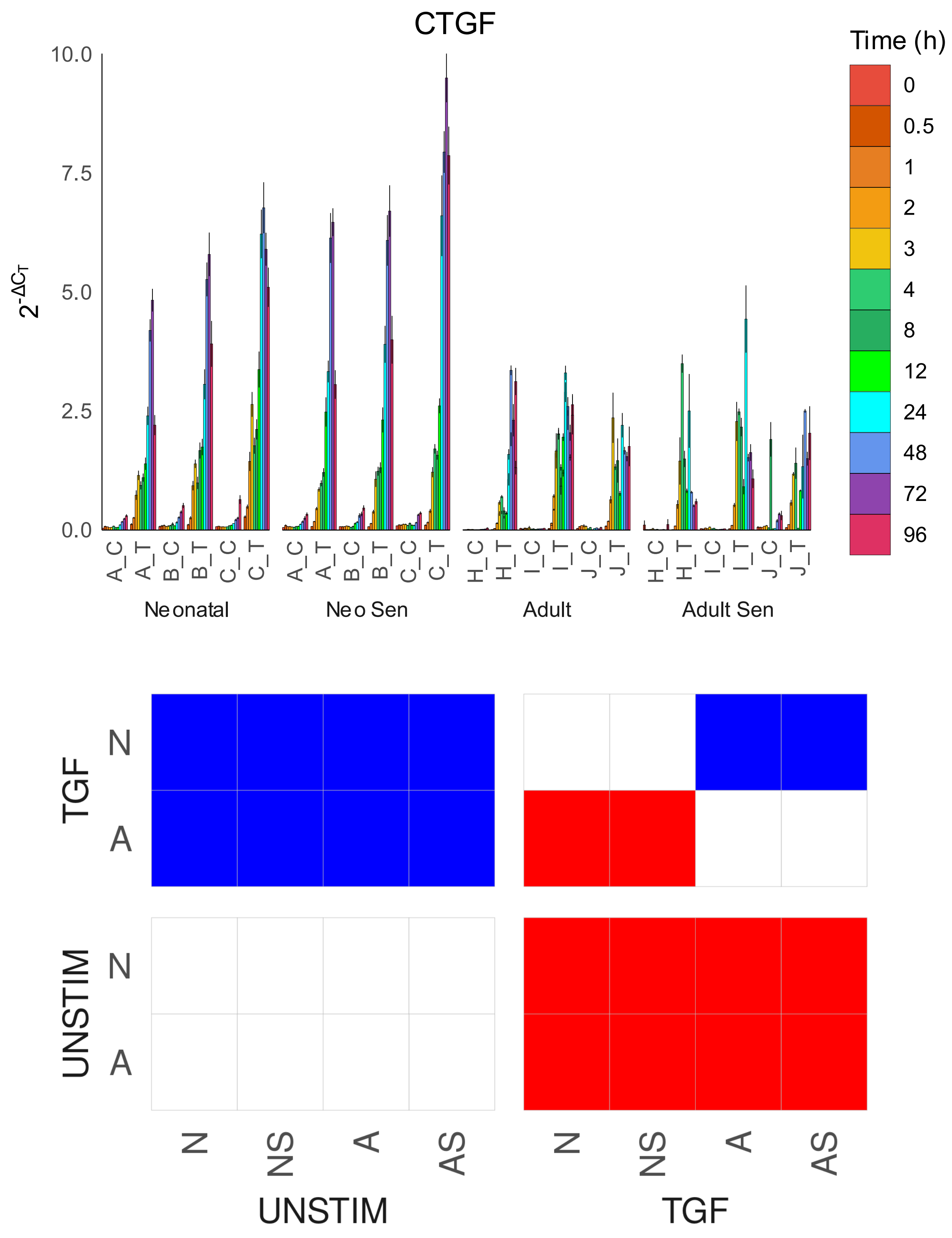

CTGF
Time (h)
0
0.5
1
2
3
4
8
12
24
48
72
96
$2^{-\Delta C_T}$
10.0
7.5
5.0
2.5
0.0
A_C
A_T
B_C
B_T
C_C
C_T
H_C
H_T
I_C
I_T
J_C
J_T
Neonatal
Neo Sen
Adult
Adult Sen
TGF
UNSTIM
N
A
N
NS
A
AS
UNSTIM
TGF

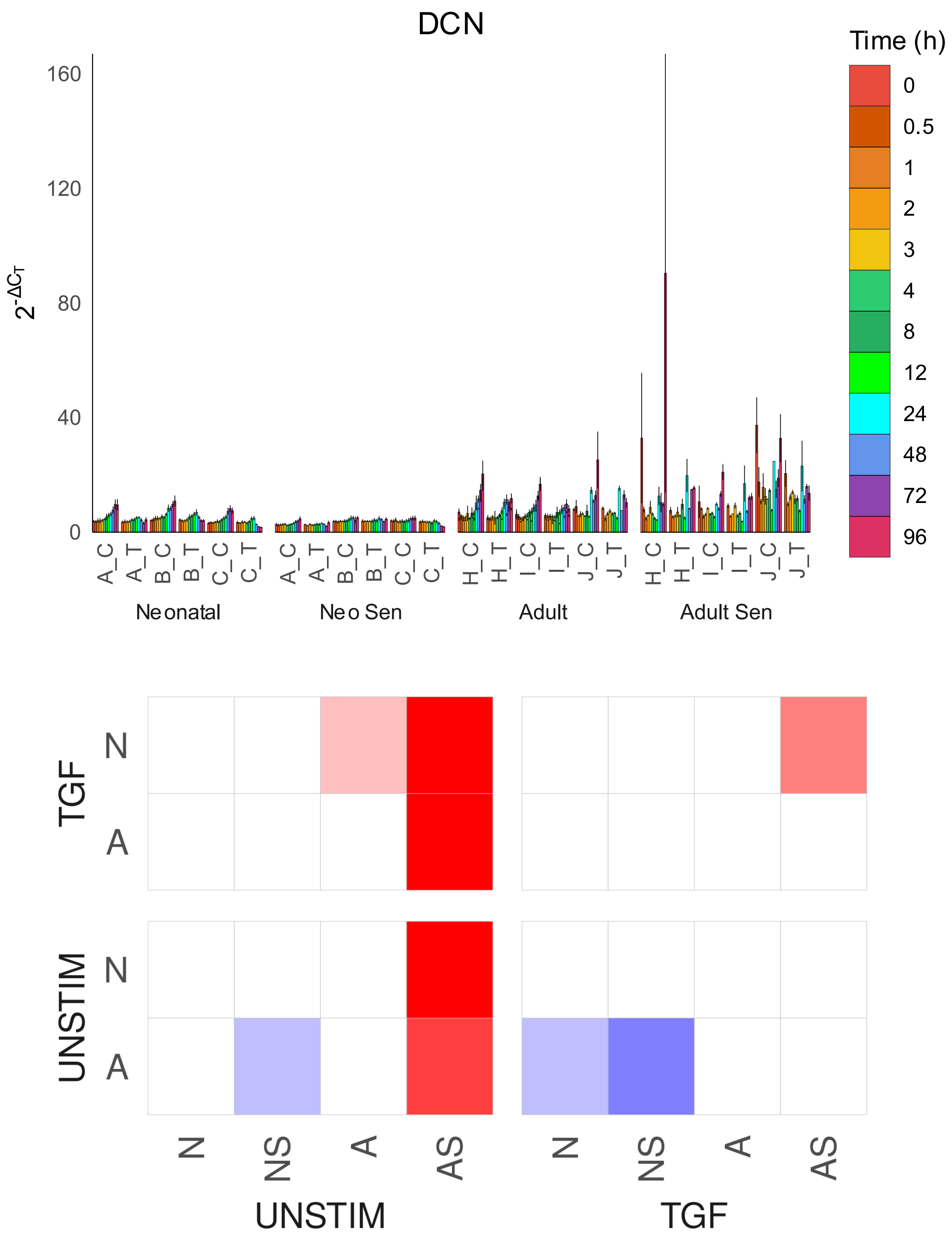
DCN
Time (h)
0
0.5
1
2
3
4
8
12
24
48
72
96
160
120
80
40
0
$2^{-\Delta C_T}$
A_C
A_T
B_C
B_T
C_C
C_T
H_C
H_T
I_C
I_T
J_C
J_T
Neonatal
Neo Sen
Adult
Adult Sen
TGF
UNSTIM
N
A
N
NS
A
AS
UNSTIM
TGF

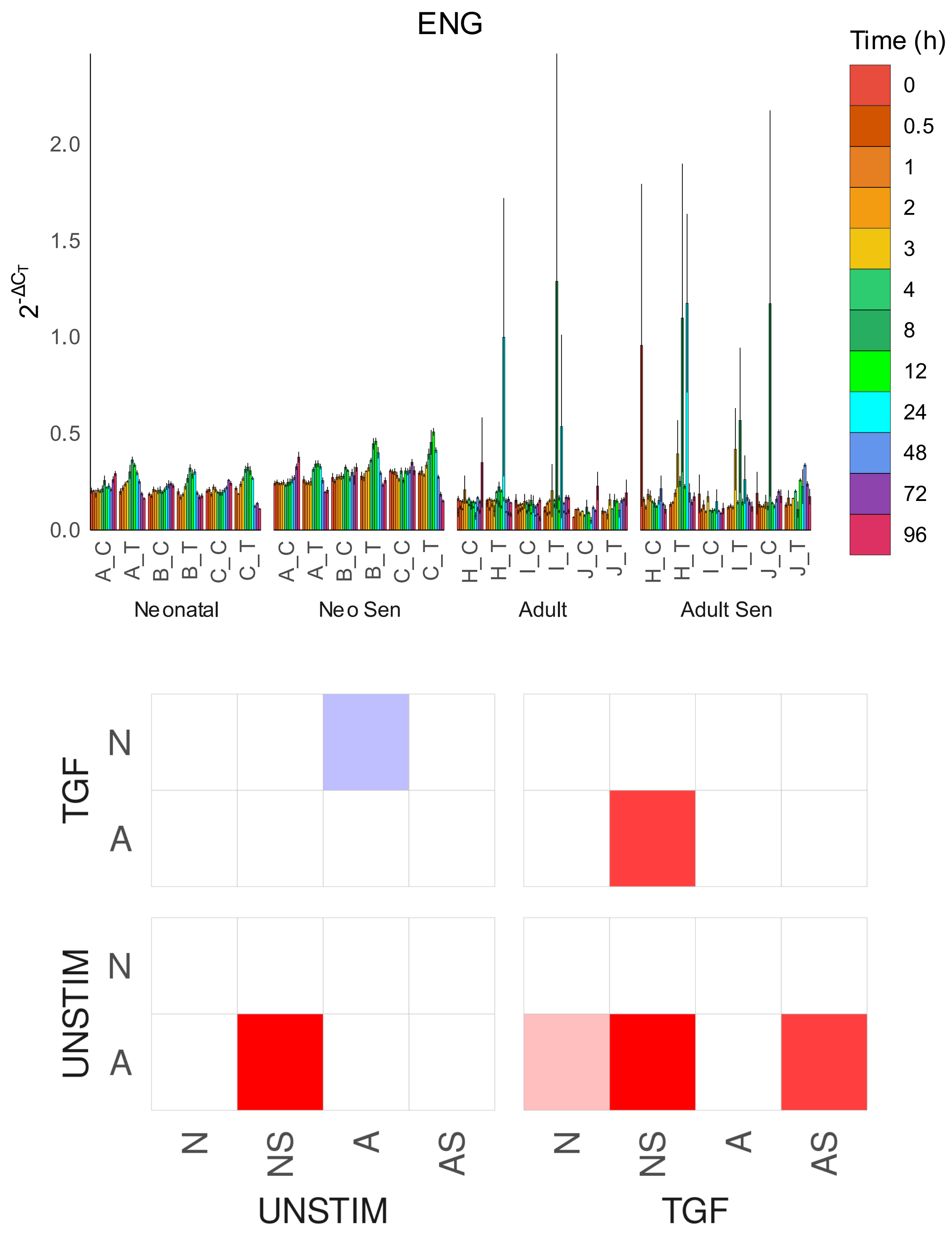

ENG
Time (h)
0
0.5
1
2
3
4
8
12
24
48
72
96
2.0
1.5
1.0
0.5
0.0
2^-ΔCT
A_C
A_T
B_C
B_T
C_C
C_T
H_C
H_T
I_C
I_T
J_C
J_T
Neonatal
Neo Sen
Adult
Adult Sen
TGF
UNSTIM
N
A
N
NS
A
AS
UNSTIM
TGF

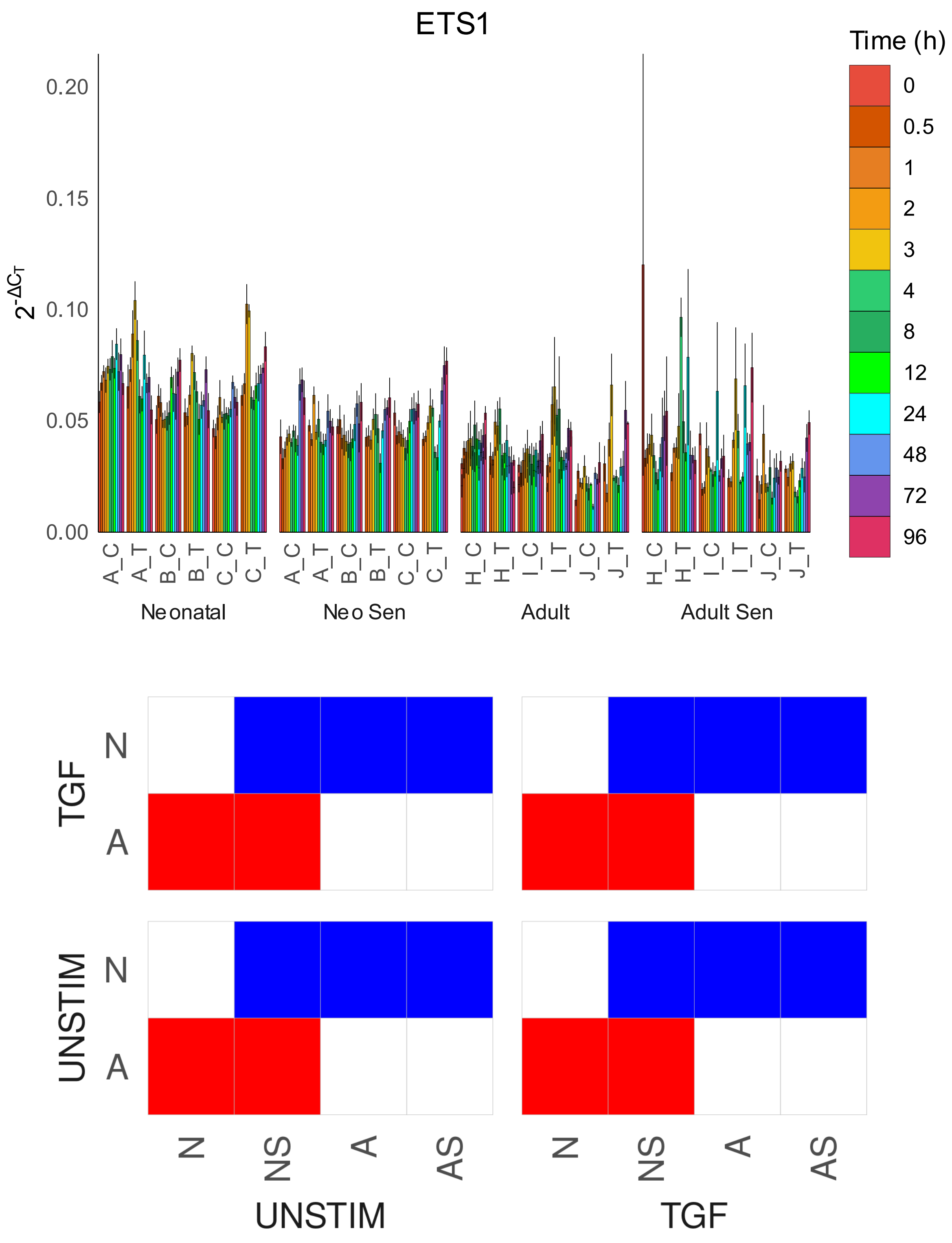
ETS1
Time (h)
0
0.5
1
2
3
4
8
12
24
48
72
96
0.20
0.15
0.10
0.05
0.00
$2^{-\Delta C_T}$
A_C
A_T
B_C
B_T
C_C
C_T
H_C
H_T
I_C
I_T
J_C
J_T
Neonatal
Neo Sen
Adult
Adult Sen
TGF
UNSTIM
N
A
N
NS
A
AS
UNSTIM
TGF

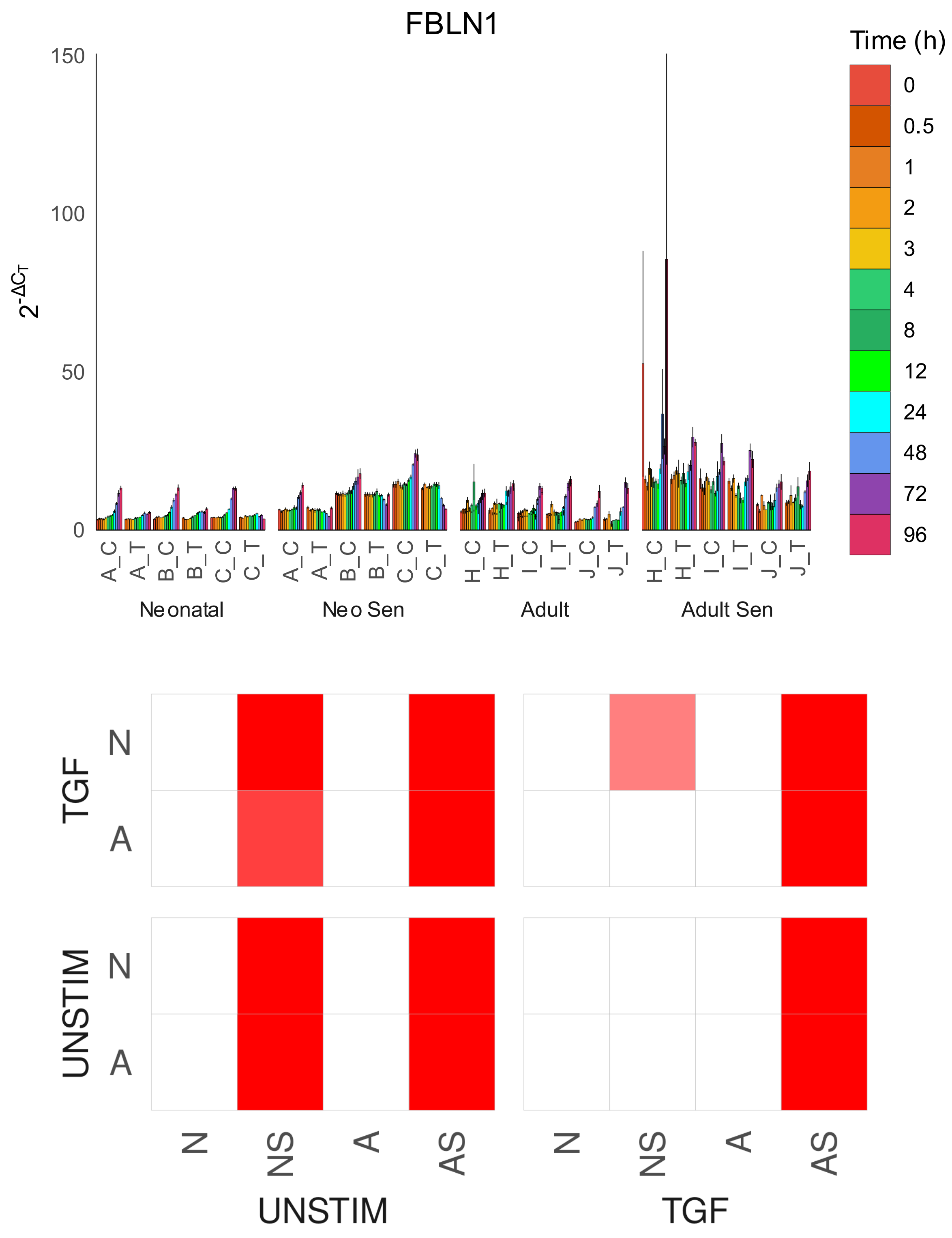

FBLN1
Time (h)
0
0.5
1
2
3
4
8
12
24
48
72
96
150
100
50
0
$2^{-\Delta C_T}$
A_C
A_T
B_C
B_T
C_C
C_T
A_C
A_T
B_C
B_T
C_C
C_T
H_C
H_T
I_C
I_T
J_C
J_T
H_C
H_T
I_C
I_T
J_C
J_T
Neonatal
Neo Sen
Adult
Adult Sen
TGF
N
A
UNSTIM
N
A
N
NS
A
AS
UNSTIM
N
NS
A
AS
TGF

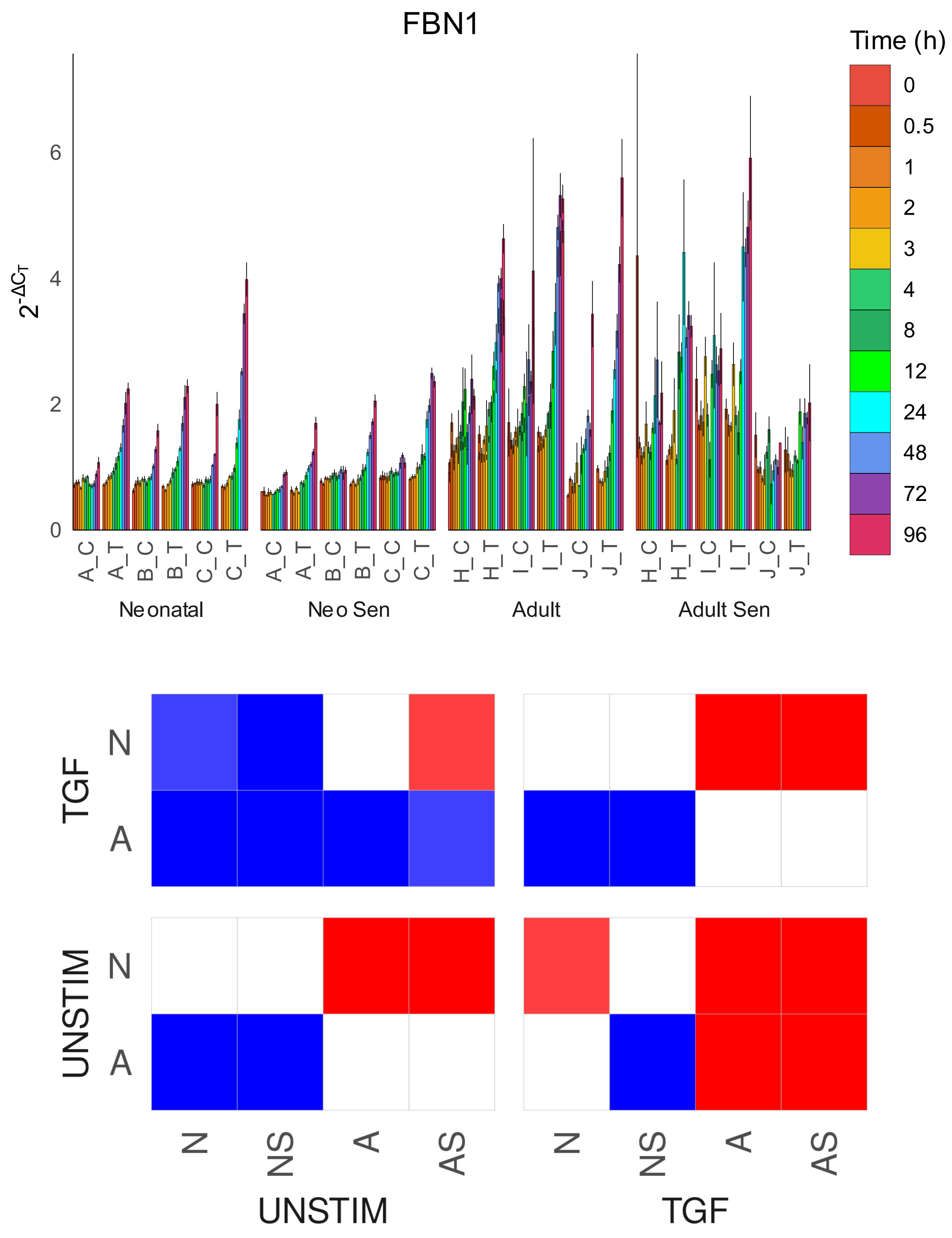

FBN1
Time (h)
0
0.5
1
2
3
4
8
12
24
48
72
96
$2^{-\Delta C_T}$
0
2
4
6
A_C A_T B_C B_T C_C C_T
Neonatal
A_C A_T B_C B_T C_C C_T
Neo Sen
H_C H_T I_C I_T J_C J_T
Adult
H_C H_T I_C I_T J_C J_T
Adult Sen
TGF
UNSTIM
N
A
N NS A AS
UNSTIM
N NS A AS
TGF

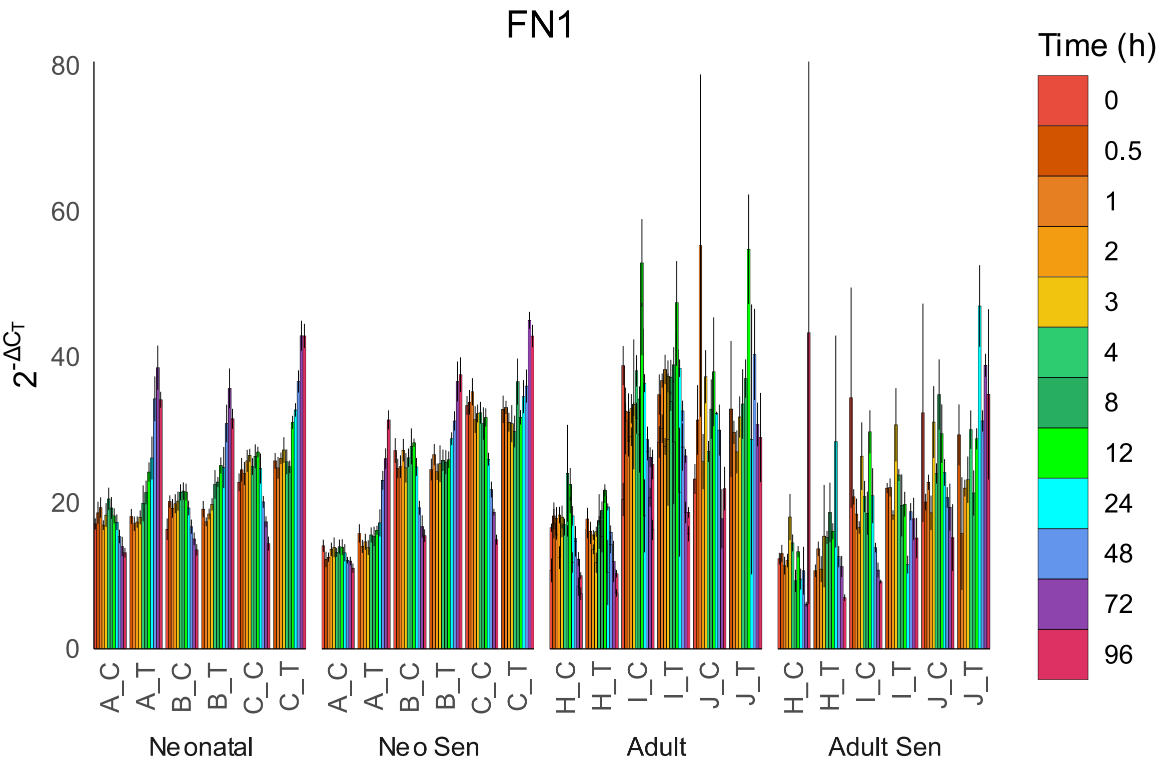

FN1
Time (h)
0
0.5
1
2
3
4
8
12
24
48
72
96
0
20
40
60
80
A_C
A_T
B_C
B_T
C_C
C_T
H_C
H_T
I_C
I_T
J_C
J_T
Neonatal
Neo Sen
Adult
Adult Sen


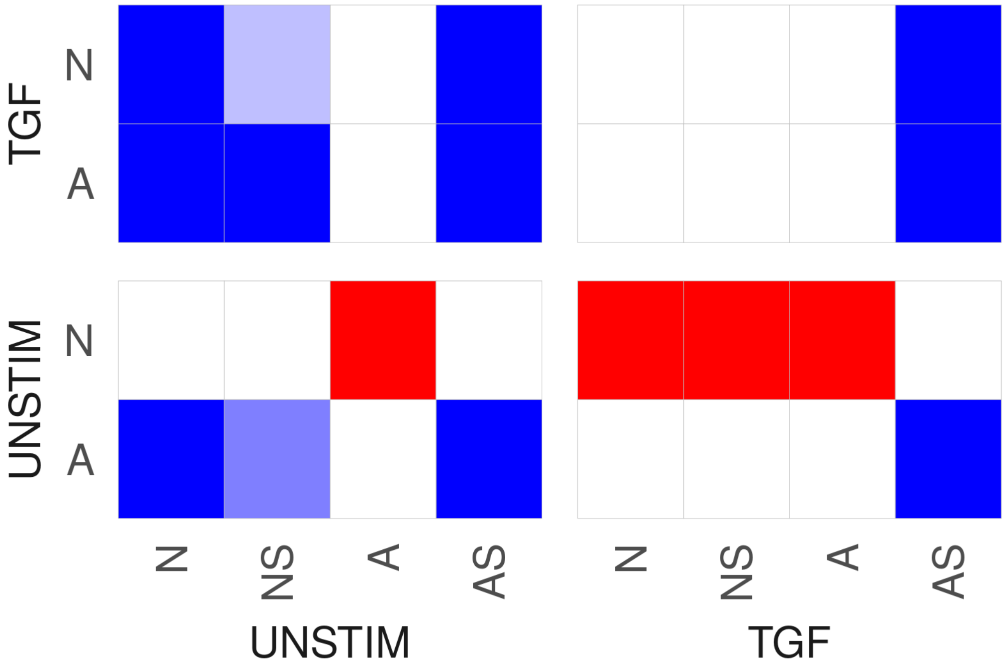

TGF
UNSTIM
N
A
N
NS
A
AS
UNSTIM
TGF

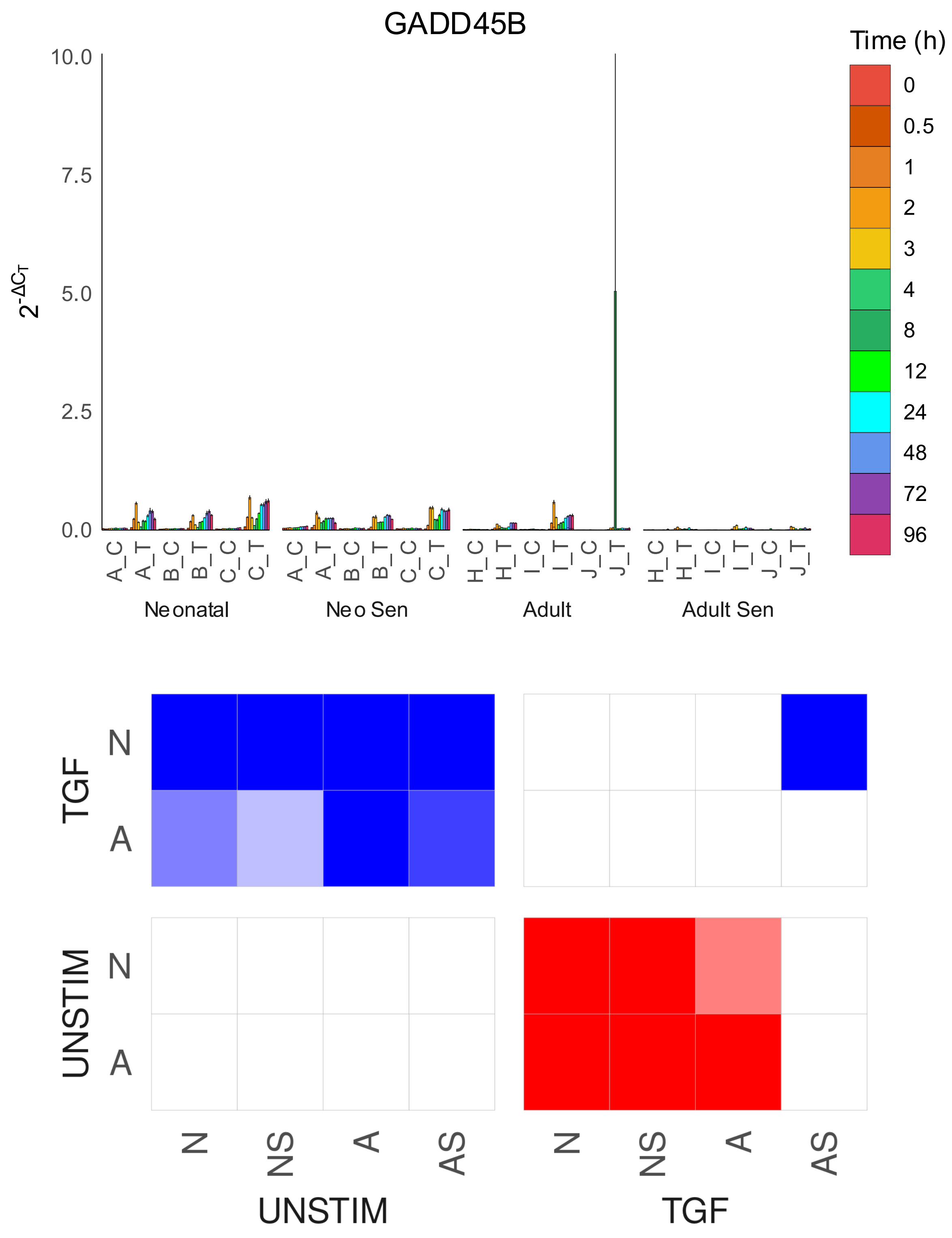
GADD45B
Time (h)
0
0.5
1
2
3
4
8
12
24
48
72
96
10.0
7.5
5.0
2.5
0.0
2-ΔCT
A_C
A_T
B_C
B_T
C_C
C_T
H_C
H_T
I_C
I_T
J_C
J_T
Neonatal
Neo Sen
Adult
Adult Sen
TGF
UNSTIM
N
A
N
NS
A
AS
UNSTIM
TGF

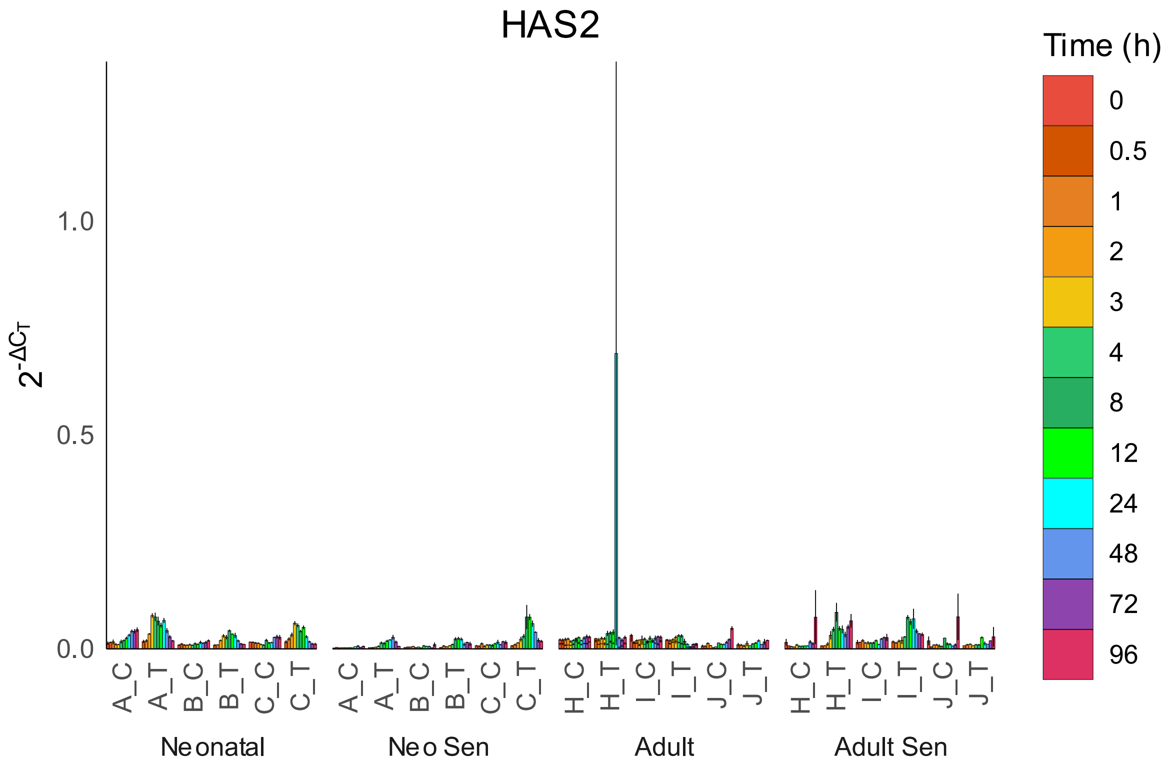

HAS2
Time (h)
0
0.5
1
2
3
4
8
12
24
48
72
96
1.0
0.5
0.0
A_C
A_T
B_C
B_T
C_C
C_T
H_C
H_T
I_C
I_T
J_C
J_T
Neonatal
Neo Sen
Adult
Adult Sen


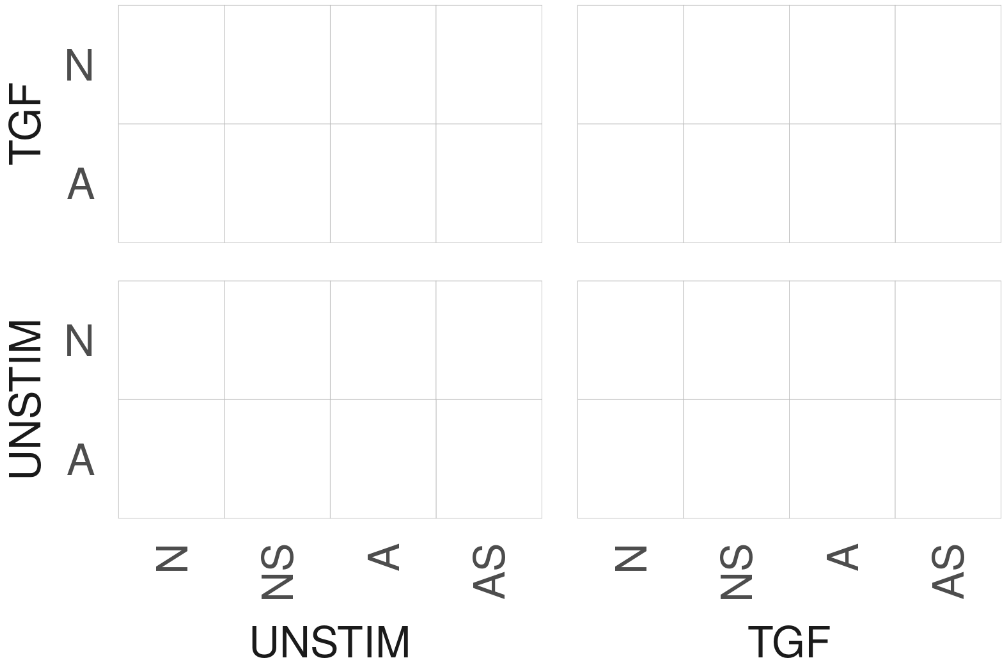

TGF
UNSTIM
N
A
N
NS
A
AS
UNSTIM
TGF

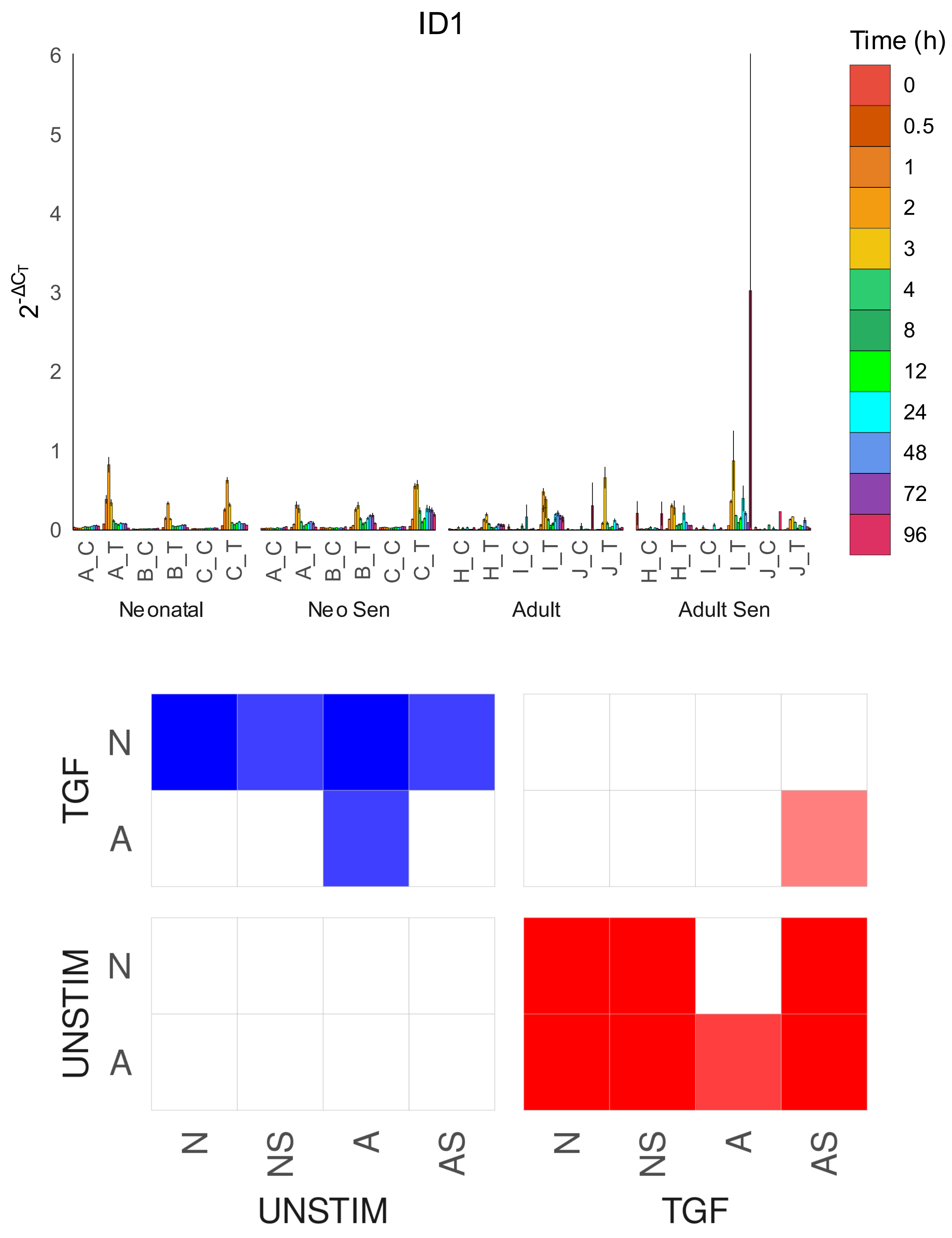
ID1
Time (h)
0
0.5
1
2
3
4
8
12
24
48
72
96
$2^{-\Delta C_T}$
6
5
4
3
2
1
0
A_C
A_T
B_C
B_T
C_C
C_T
H_C
H_T
I_C
I_T
J_C
J_T
Neonatal
Neo Sen
Adult
Adult Sen
TGF
UNSTIM
N
A
N
NS
A
AS
UNSTIM
TGF

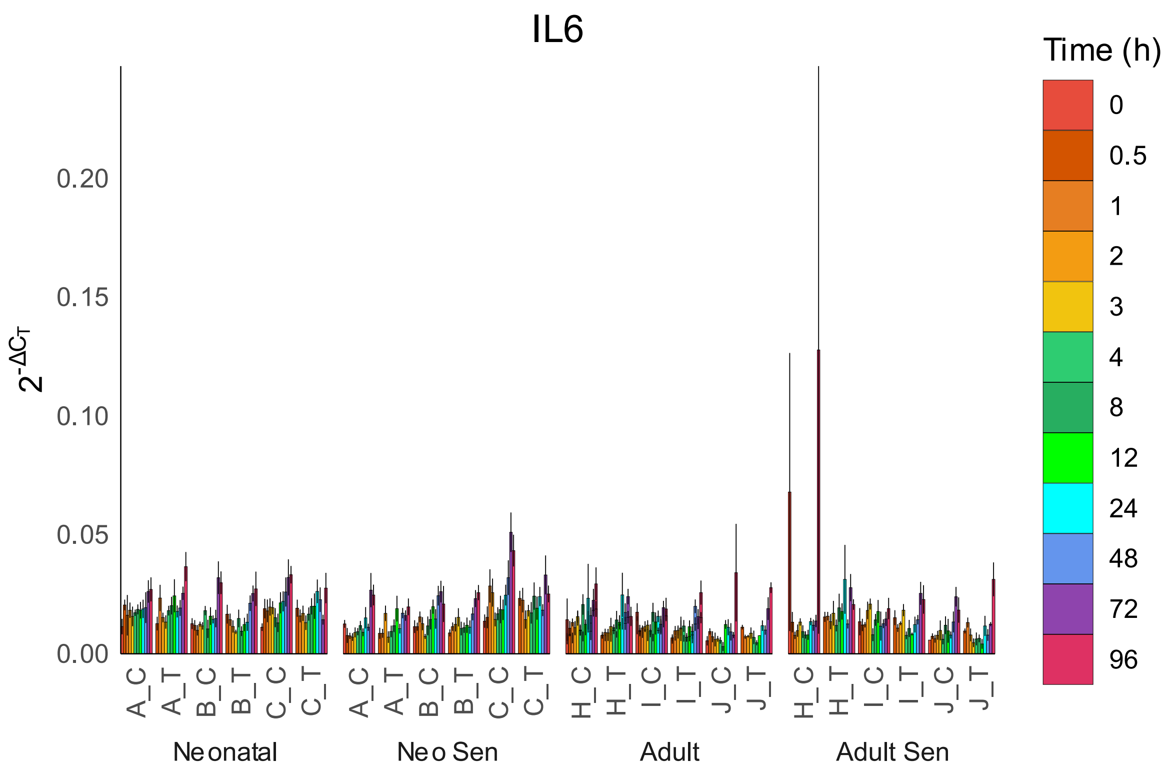
IL6
Time (h)
0
0.5
1
2
3
4
8
12
24
48
72
96
0.20
0.15
0.10
0.05
0.00
A_C
A_T
B_C
B_T
C_C
C_T
H_C
H_T
I_C
I_T
J_C
J_T
Neonatal
Neo Sen
Adult
Adult Sen

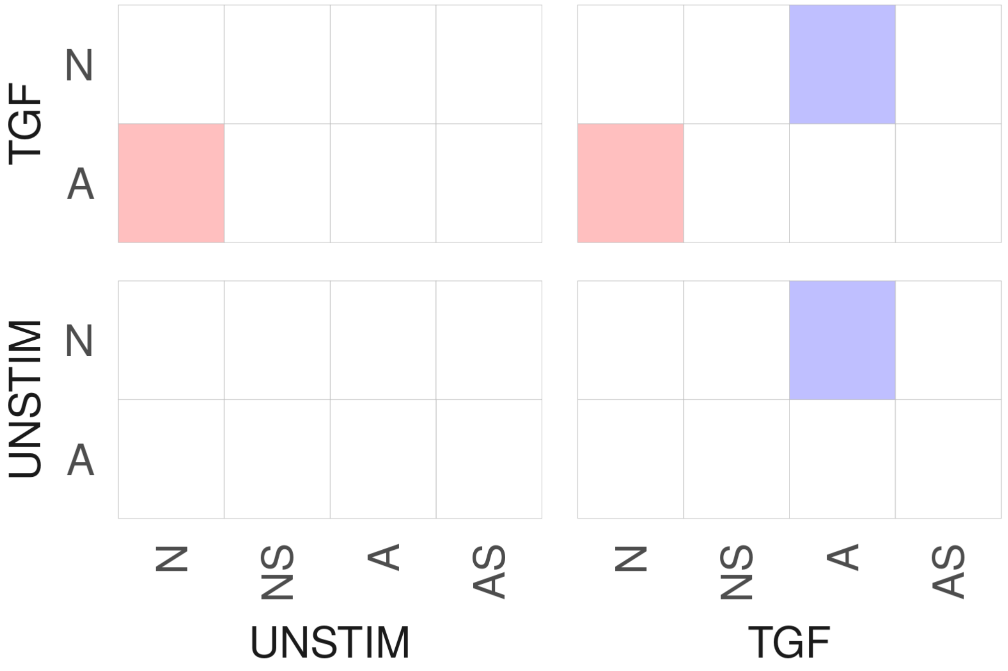
TGF
UNSTIM
N
A
N
NS
A
AS
UNSTIM
TGF

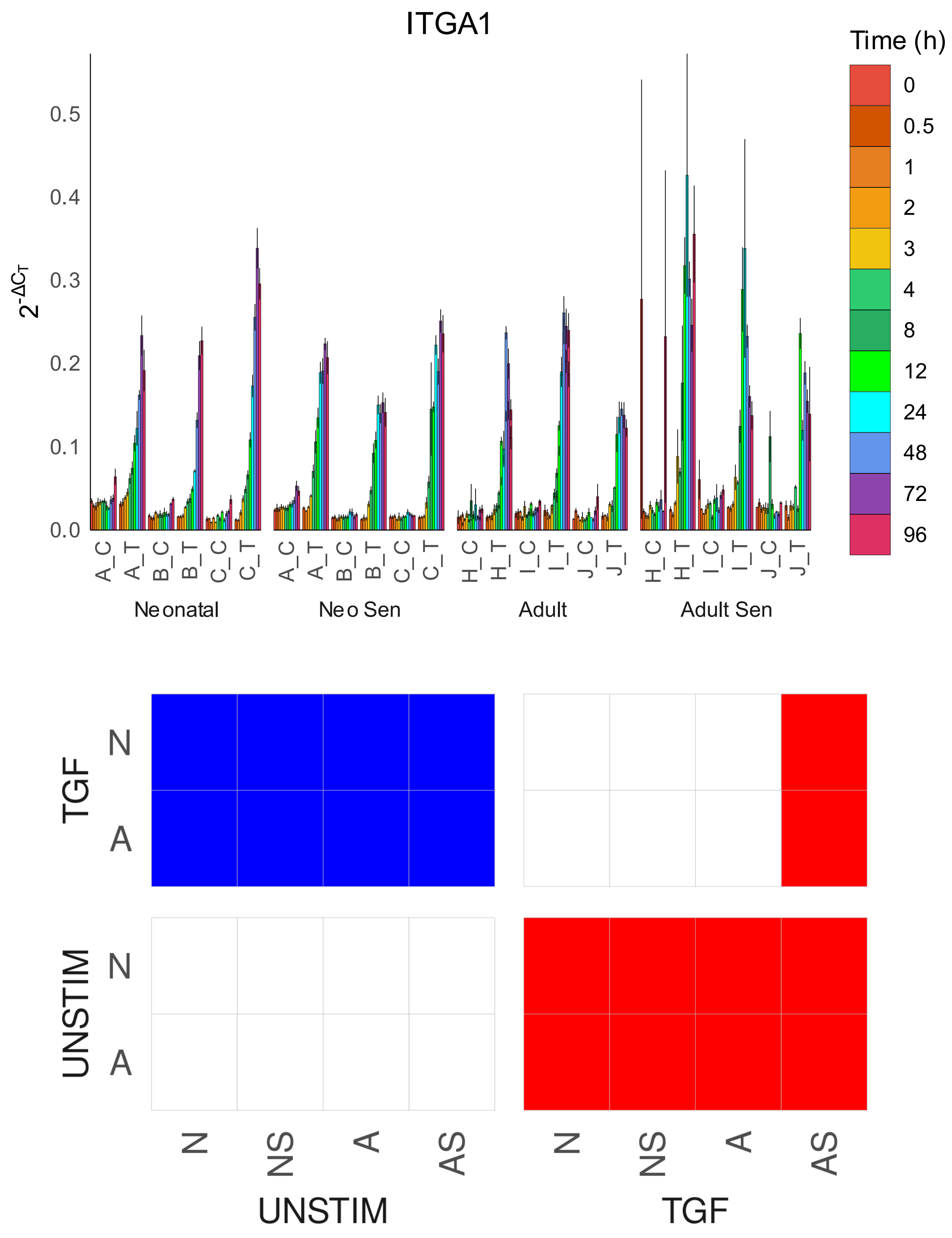

ITGA1
Time (h)
0
0.5
1
2
3
4
8
12
24
48
72
96
$2^{-\Delta C_T}$
0.0
0.1
0.2
0.3
0.4
0.5
A_C
A_T
B_C
B_T
C_C
C_T
H_C
H_T
I_C
I_T
J_C
J_T
Neonatal
Neo Sen
Adult
Adult Sen
TGF
UNSTIM
N
A
N
NS
A
AS
UNSTIM
TGF

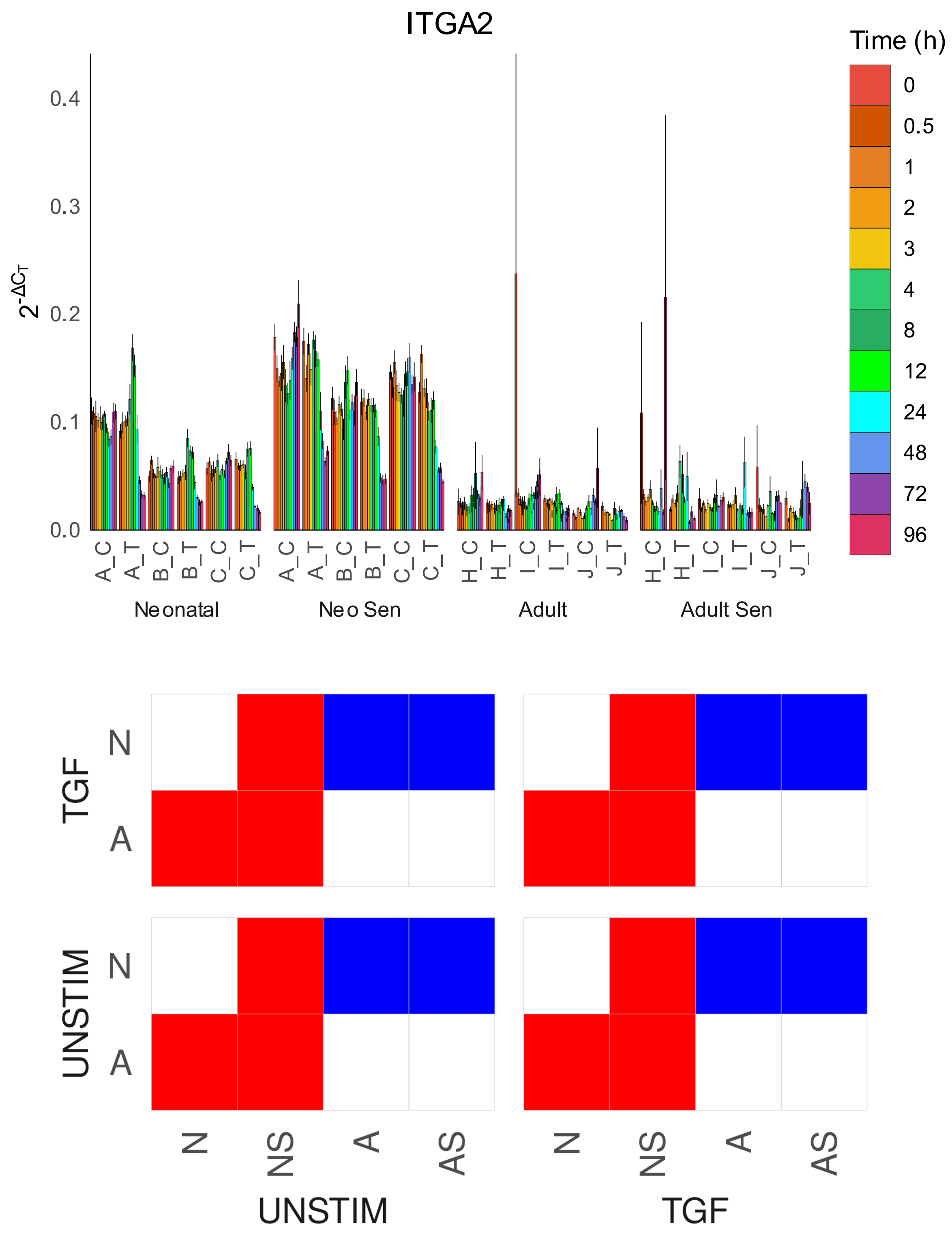

ITGA2
Time (h)
0
0.5
1
2
3
4
8
12
24
48
72
96
0.4
0.3
0.2
0.1
0.0
A_C
A_T
B_C
B_T
C_C
C_T
H_C
H_T
I_C
I_T
J_C
J_T
Neonatal
Neo Sen
Adult
Adult Sen
TGF
UNSTIM
N
A
N
NS
A
AS
UNSTIM
TGF

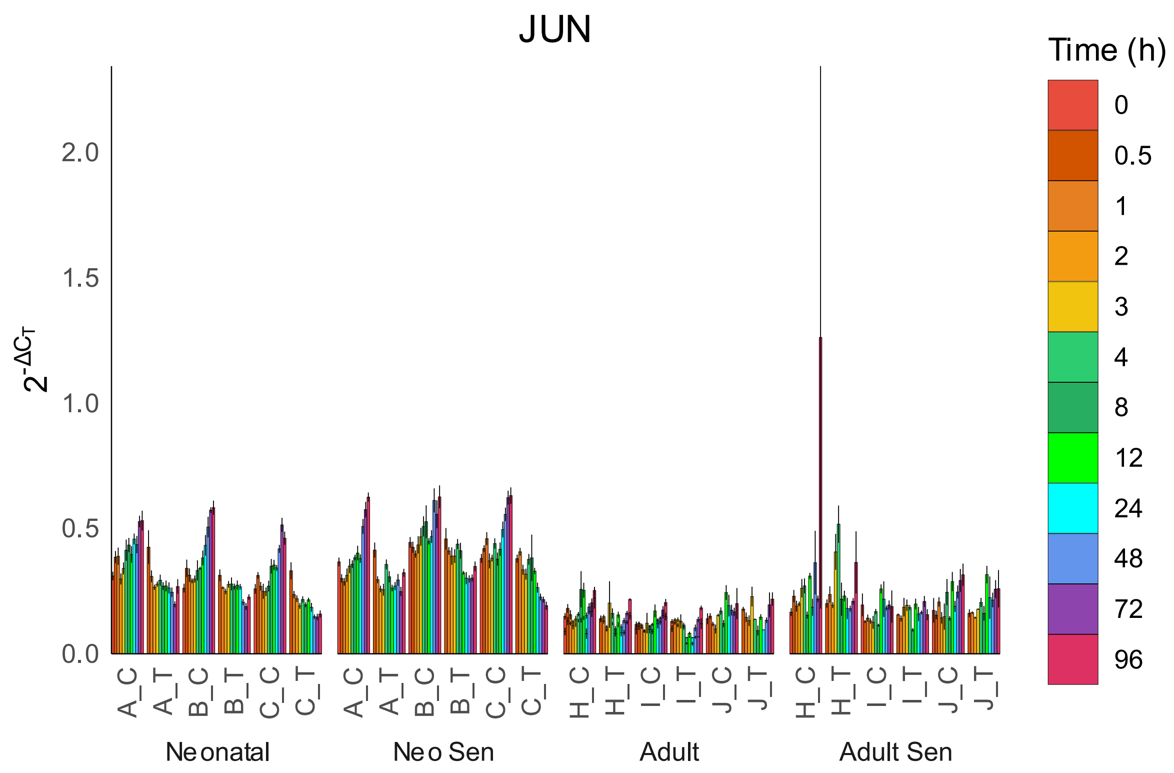
JUN
Time (h)
0
0.5
1
2
3
4
8
12
24
48
72
96
2.0
1.5
1.0
0.5
0.0
$2^{-\Delta C_T}$
A_C
A_T
B_C
B_T
C_C
C_T
H_C
H_T
I_C
I_T
J_C
J_T
Neonatal
Neo Sen
Adult
Adult Sen

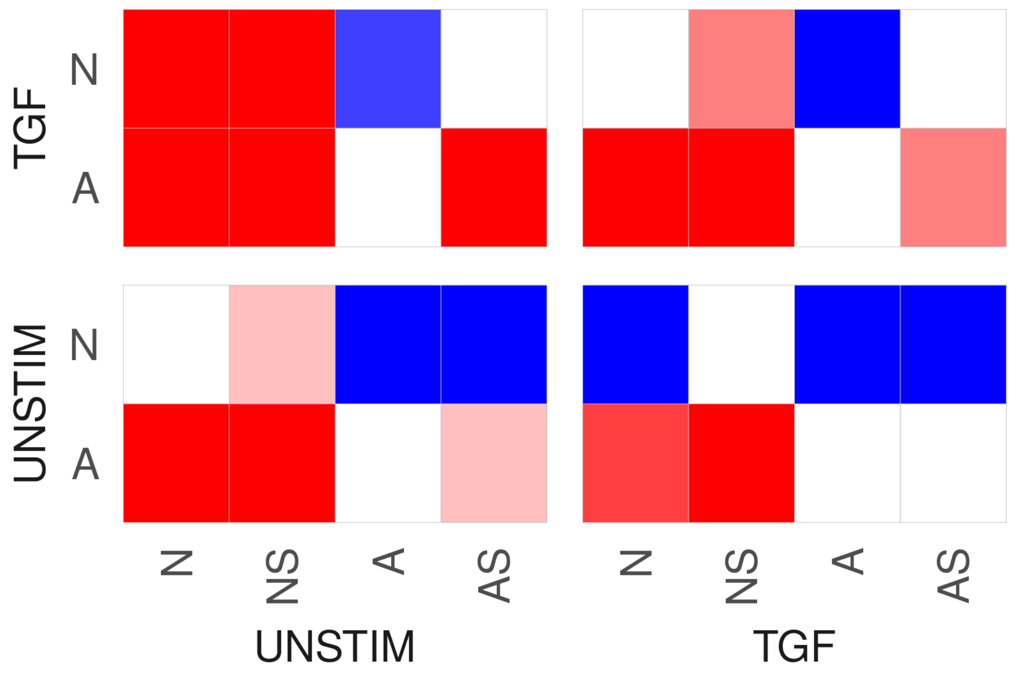
TGF
UNSTIM
N
A
N
NS
A
AS
UNSTIM
TGF

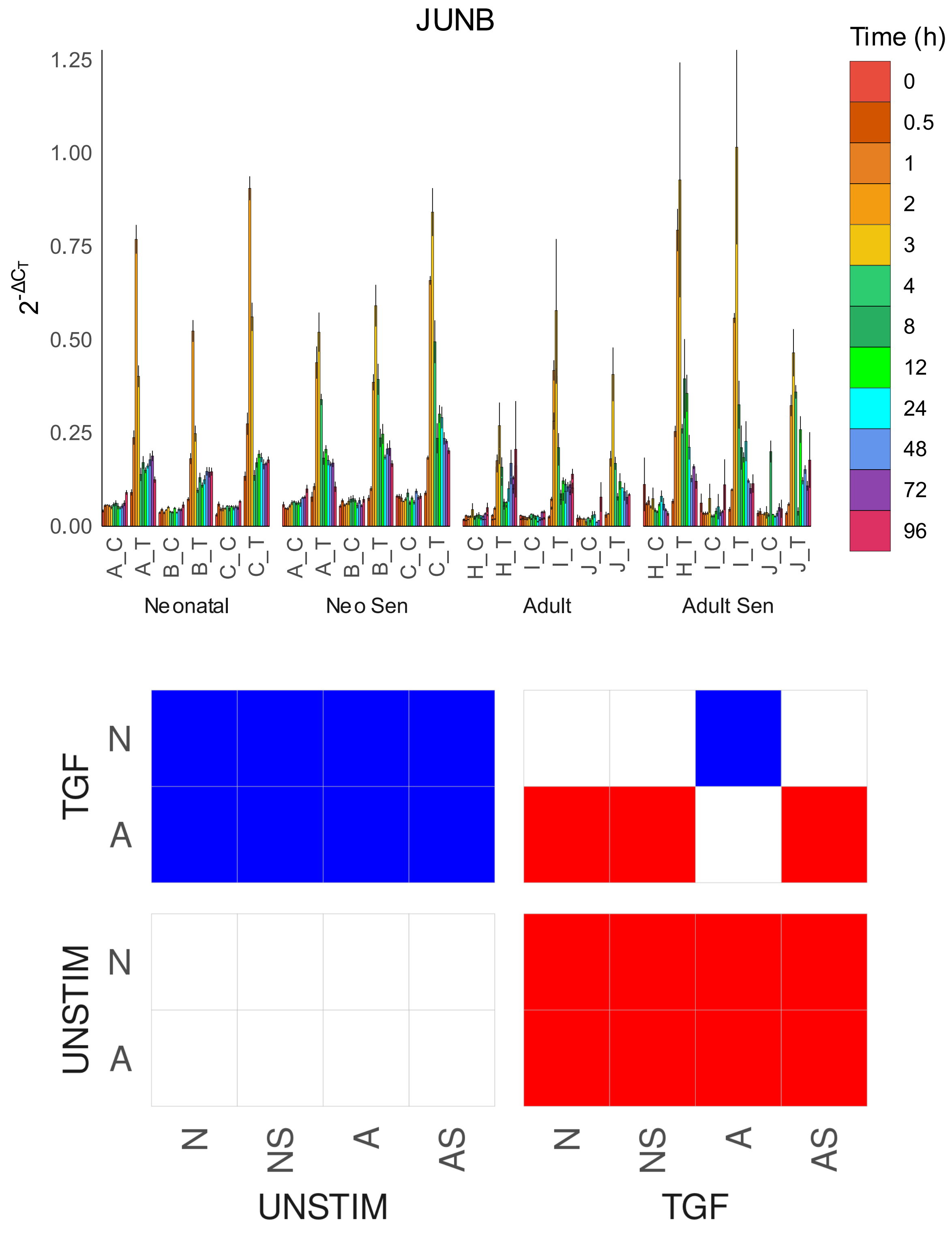

JUNB
Time (h)
0
0.5
1
2
3
4
8
12
24
48
72
96
1.25
1.00
0.75
0.50
0.25
0.00
$2^{-\Delta C_T}$
A_C
A_T
B_C
B_T
C_C
C_T
H_C
H_T
I_C
I_T
J_C
J_T
Neonatal
Neo Sen
Adult
Adult Sen
TGF
UNSTIM
N
A
N
NS
A
AS
UNSTIM
TGF

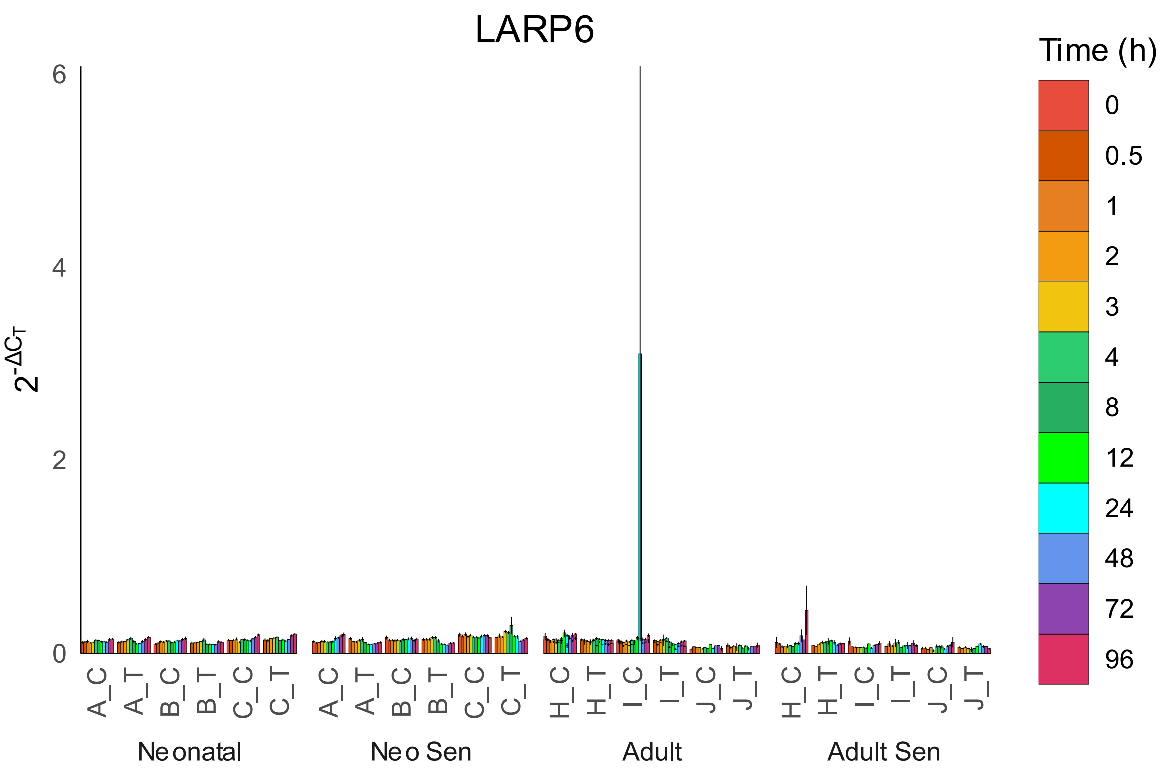

LARP6
Time (h)
0
0.5
1
2
3
4
8
12
24
48
72
96
6
4
2
0
2-ΔCT
A_C
A_T
B_C
B_T
C_C
C_T
H_C
H_T
I_C
I_T
J_C
J_T
Neonatal
Neo Sen
Adult
Adult Sen


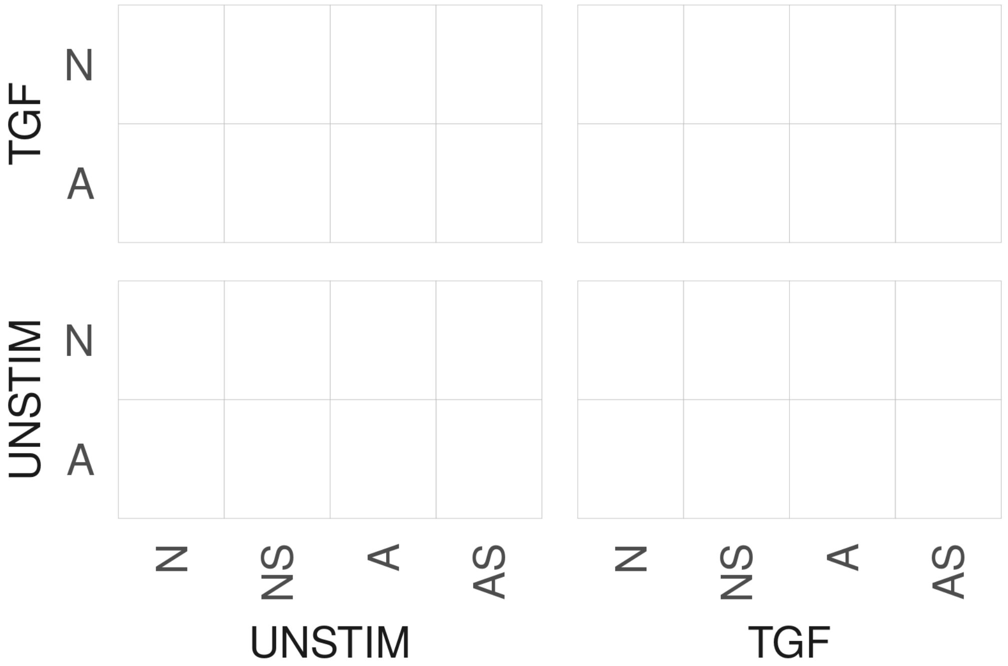

TGF
N
A
UNSTIM
N
A
N
NS
A
AS
UNSTIM
TGF

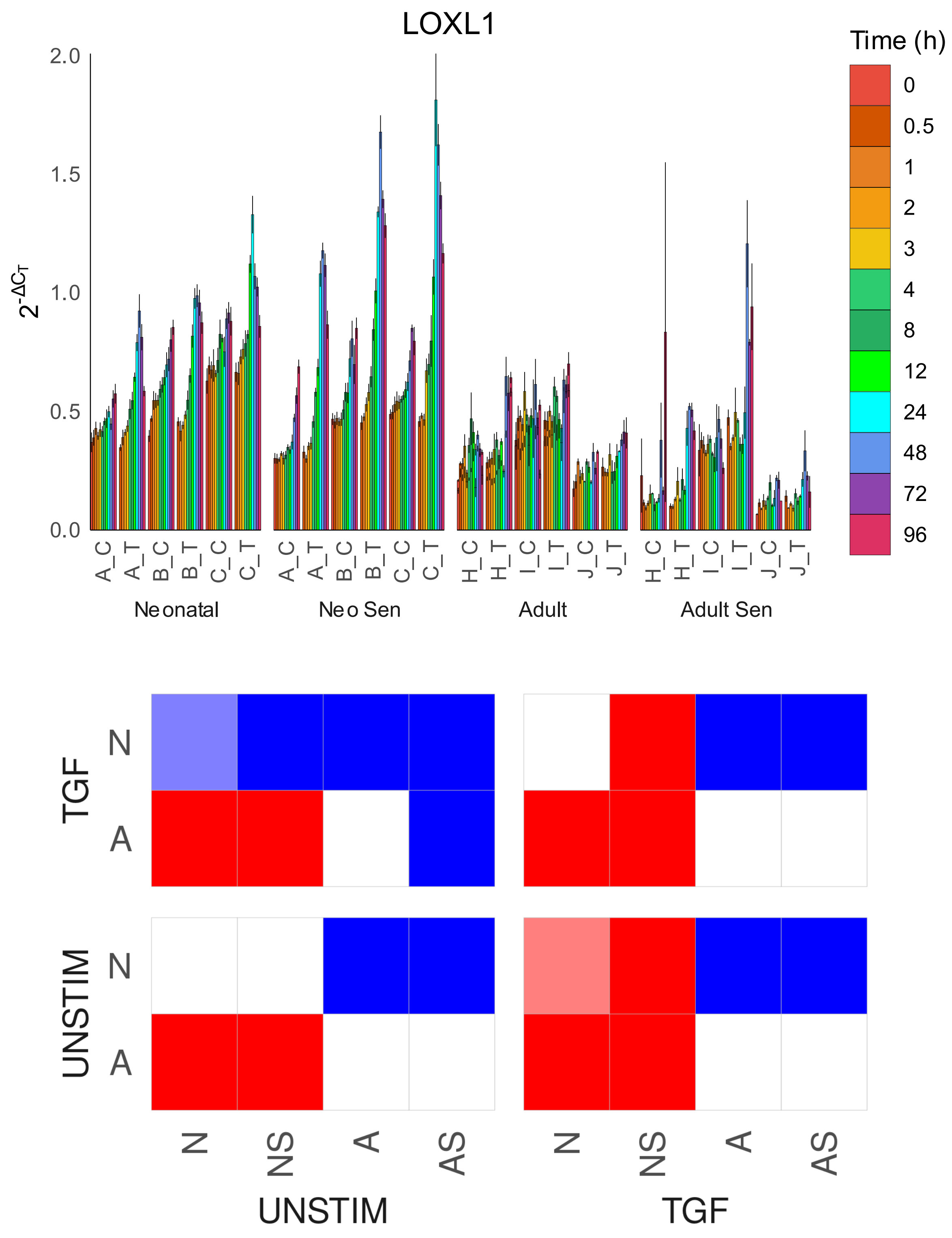

LOXL1
Time (h)
0
0.5
1
2
3
4
8
12
24
48
72
96
2.0
1.5
1.0
0.5
0.0
$2^{-\Delta C_T}$
A_C
A_T
B_C
B_T
C_C
C_T
H_C
H_T
I_C
I_T
J_C
J_T
Neonatal
Neo Sen
Adult
Adult Sen
TGF
UNSTIM
N
A
N
NS
A
AS
UNSTIM
TGF

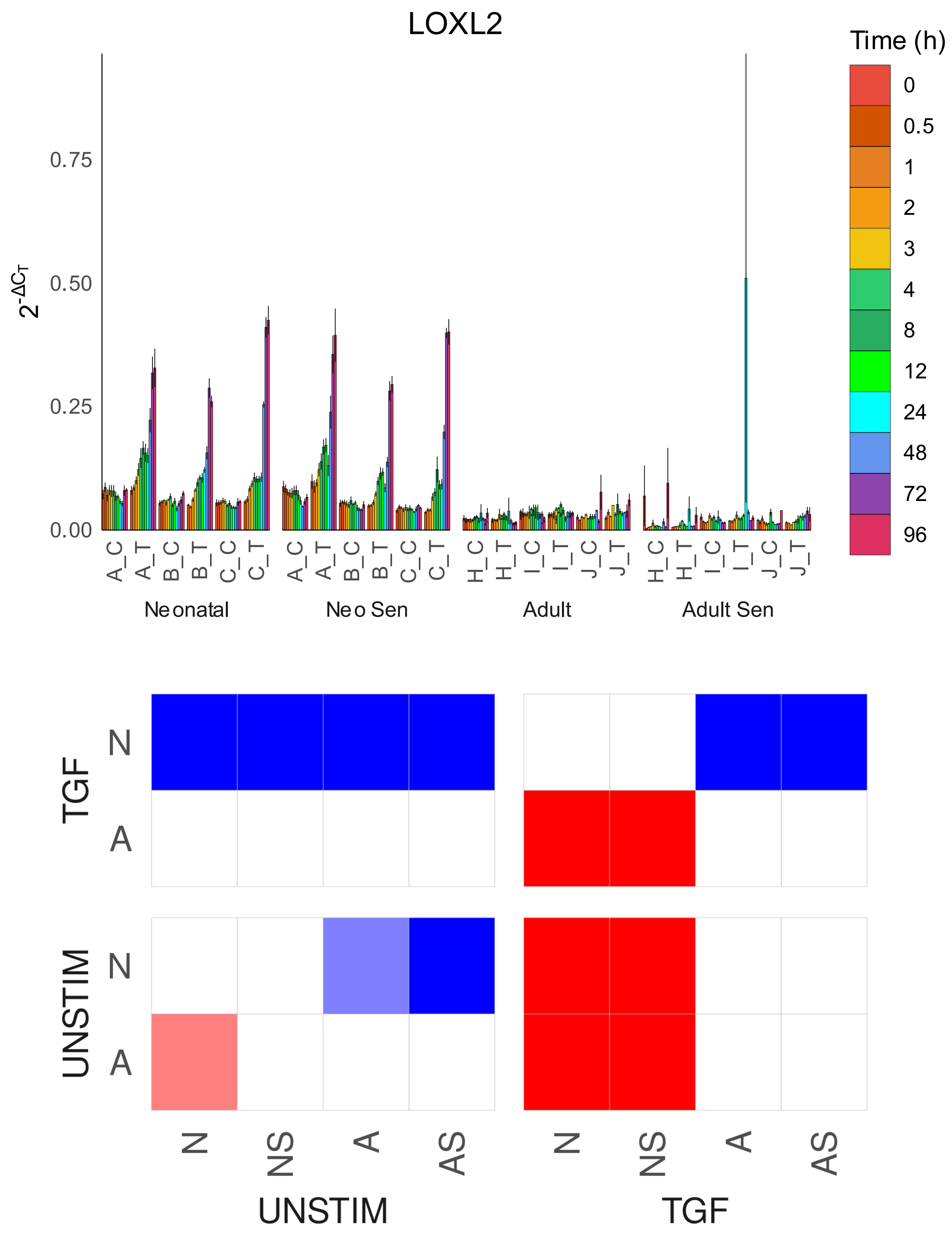
LOXL2
Time (h)
0
0.5
1
2
3
4
8
12
24
48
72
96
0.75
0.50
0.25
0.00
A_C
A_T
B_C
B_T
C_C
C_T
H_C
H_T
I_C
I_T
J_C
J_T
Neonatal
Neo Sen
Adult
Adult Sen
TGF
UNSTIM
N
A
N
NS
A
AS
UNSTIM
TGF

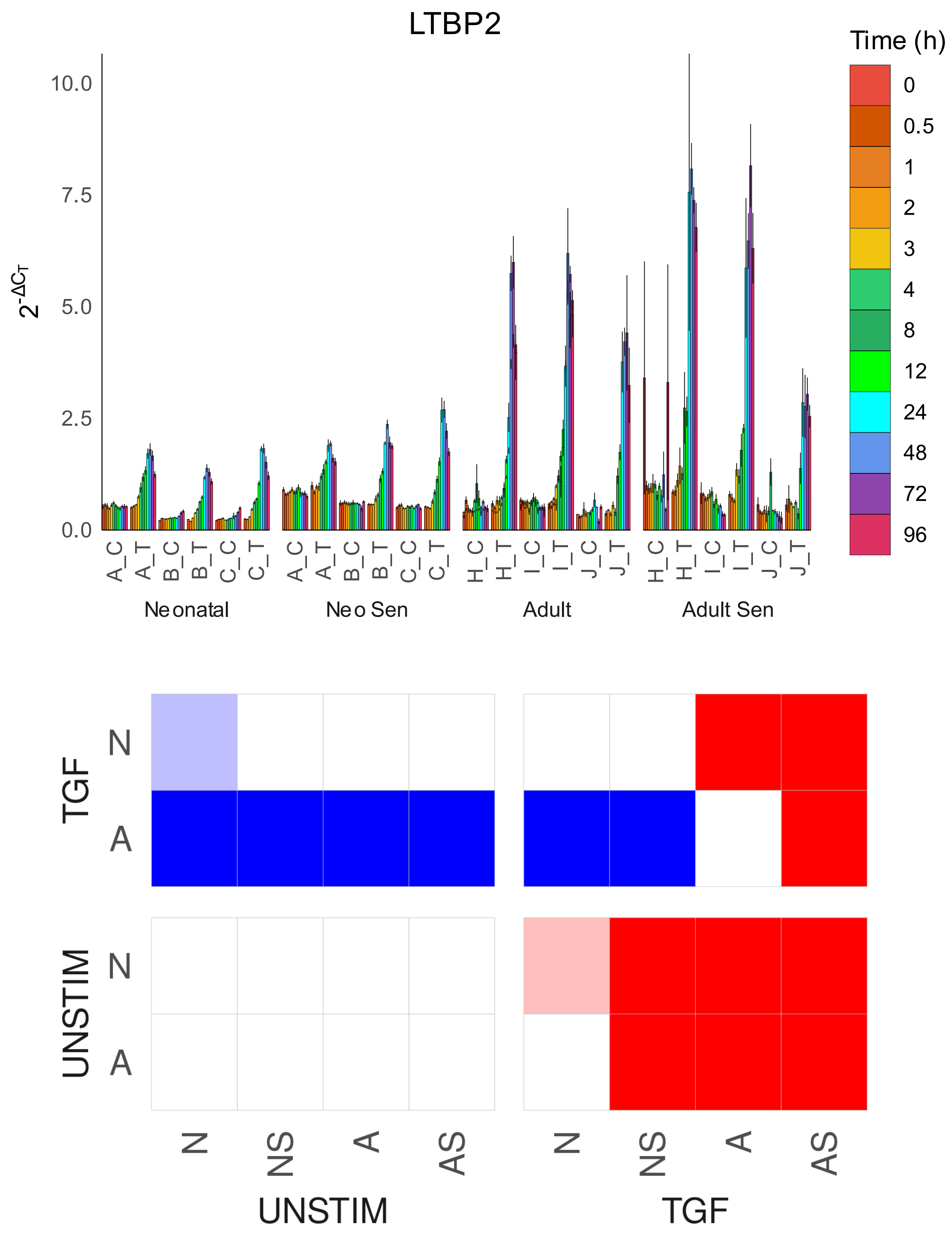

LTBP2
Time (h)
0
0.5
1
2
3
4
8
12
24
48
72
96
10.0
7.5
5.0
2.5
0.0
$2^{-\Delta C_T}$
A_C
A_T
B_C
B_T
C_C
C_T
H_C
H_T
I_C
I_T
J_C
J_T
Neonatal
Neo Sen
Adult
Adult Sen
TGF
UNSTIM
N
A
N
NS
A
AS
UNSTIM
TGF

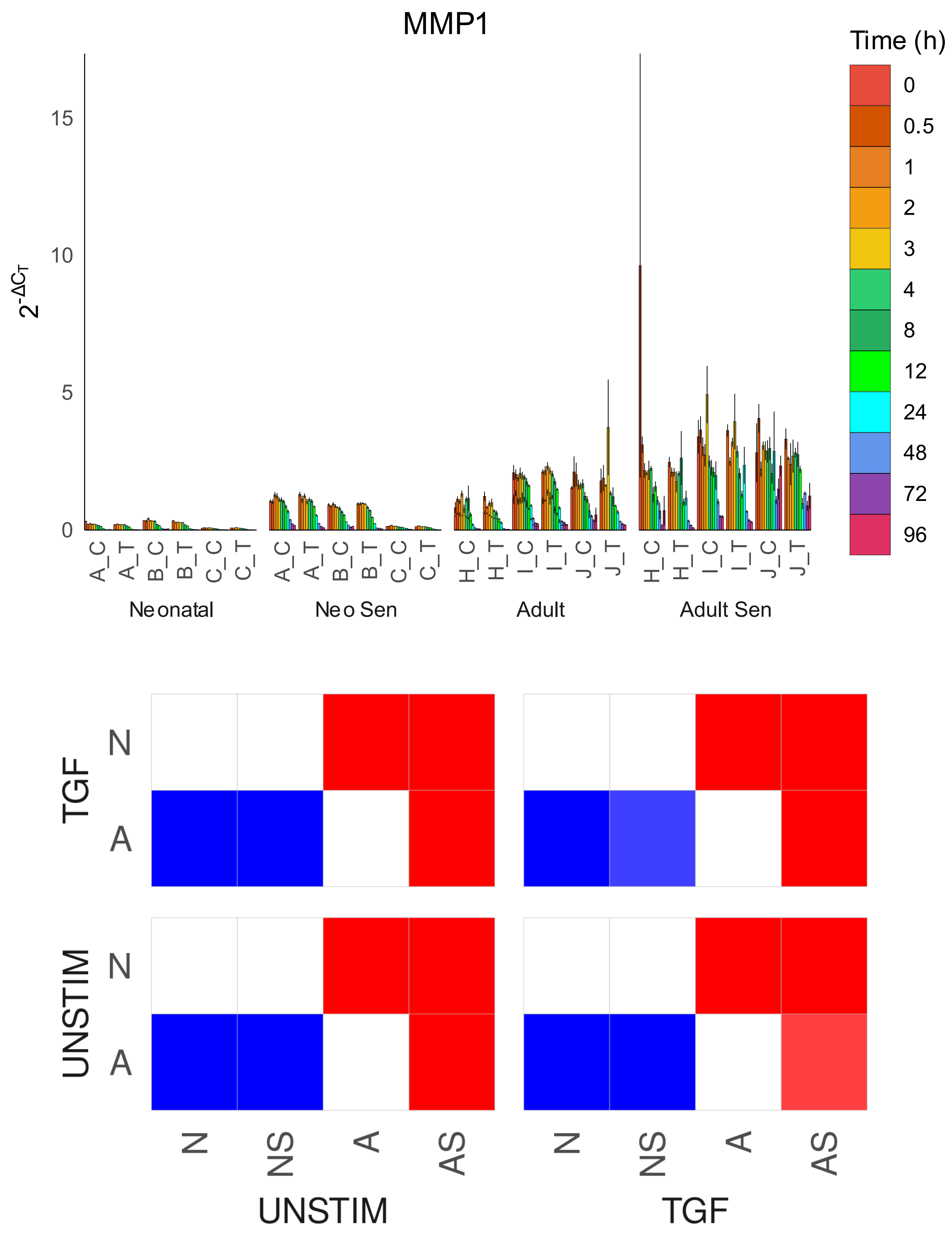

MMP1
Time (h)
0
0.5
1
2
3
4
8
12
24
48
72
96
15
10
5
0
$2^{-\Delta C_T}$
A_C
A_T
B_C
B_T
C_C
C_T
H_C
H_T
I_C
I_T
J_C
J_T
Neonatal
Neo Sen
Adult
Adult Sen
TGF
UNSTIM
N
A
N
NS
A
AS
UNSTIM
TGF

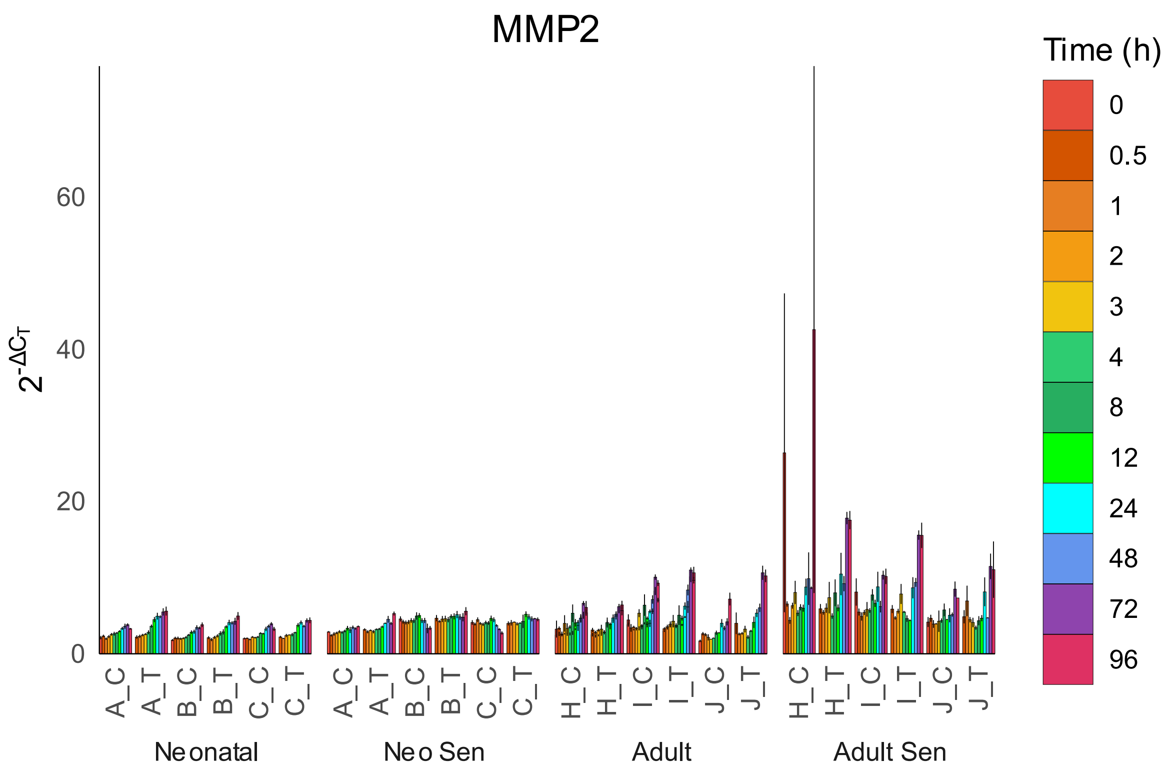
MMP2
Time (h)
0
0.5
1
2
3
4
8
12
24
48
72
96
$2^{-\Delta C_T}$
0
20
40
60
A_C
A_T
B_C
B_T
C_C
C_T
H_C
H_T
I_C
I_T
J_C
J_T
Neonatal
Neo Sen
Adult
Adult Sen

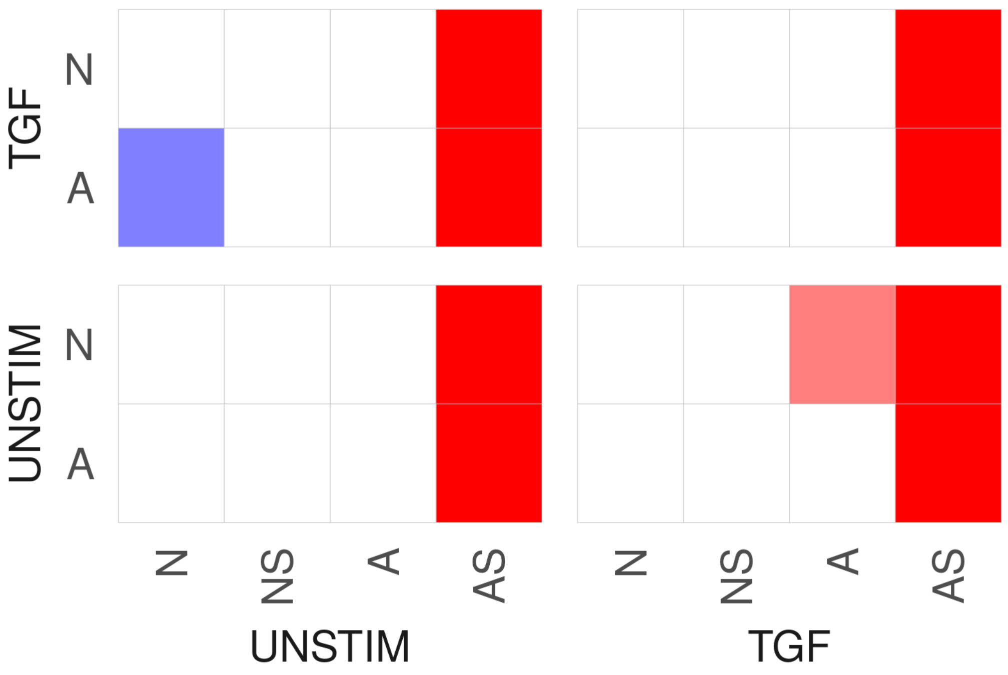
TGF
N
A
UNSTIM
N
A
N
NS
A
AS
UNSTIM
N
NS
A
AS
TGF

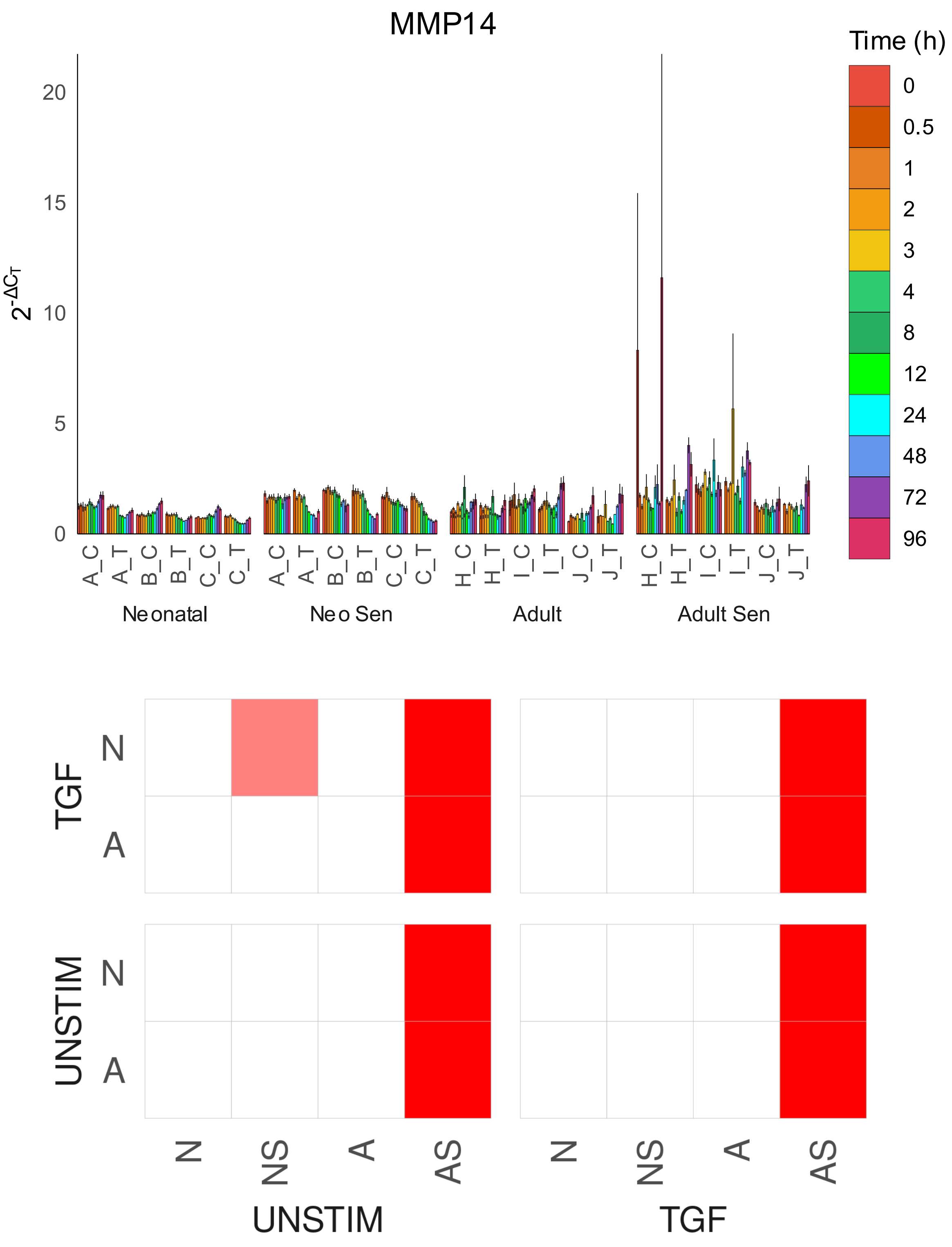
MMP14
Time (h)
0
0.5
1
2
3
4
8
12
24
48
72
96
20
15
10
5
0
A_C
A_T
B_C
B_T
C_C
C_T
H_C
H_T
I_C
I_T
J_C
J_T
Neonatal
Neo Sen
Adult
Adult Sen
TGF
UNSTIM
N
A
N
NS
A
AS
UNSTIM
TGF

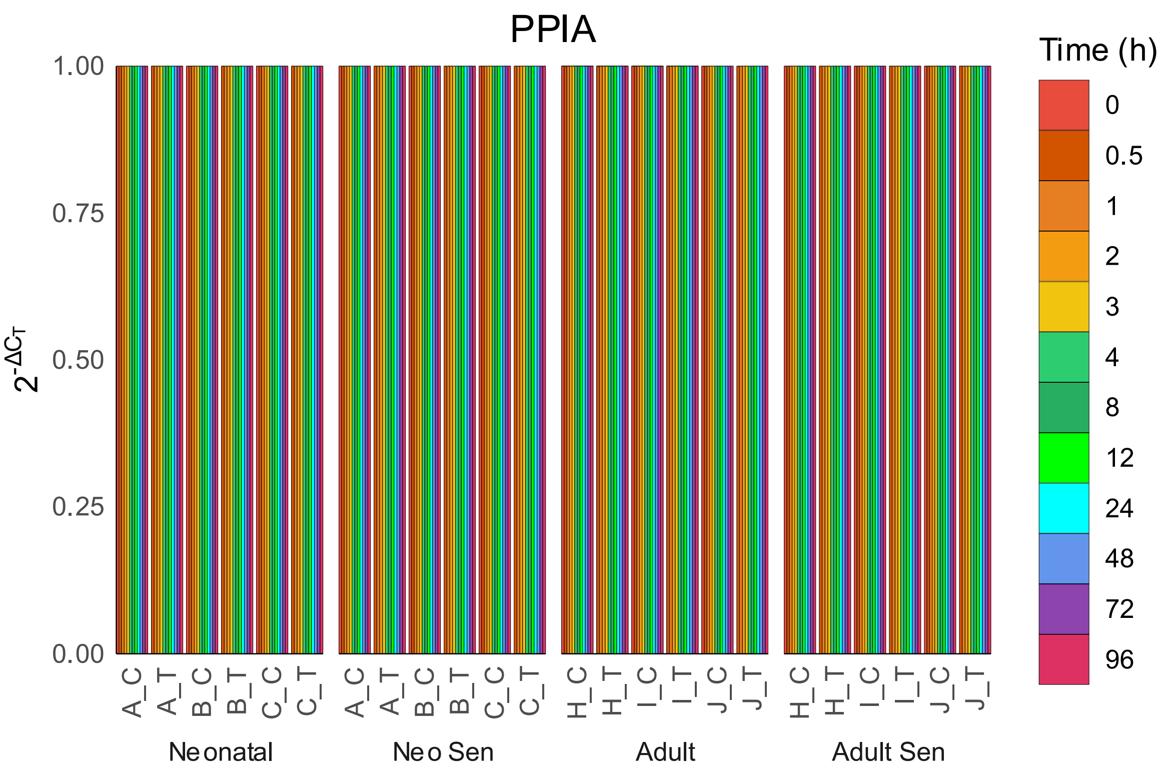
PPIA
Time (h)
0
0.5
1
2
3
4
8
12
24
48
72
96
1.00
0.75
0.50
0.25
0.00
$2^{-\Delta C_T}$
A_C
A_T
B_C
B_T
C_C
C_T
H_C
H_T
I_C
I_T
J_C
J_T
Neonatal
Neo Sen
Adult
Adult Sen

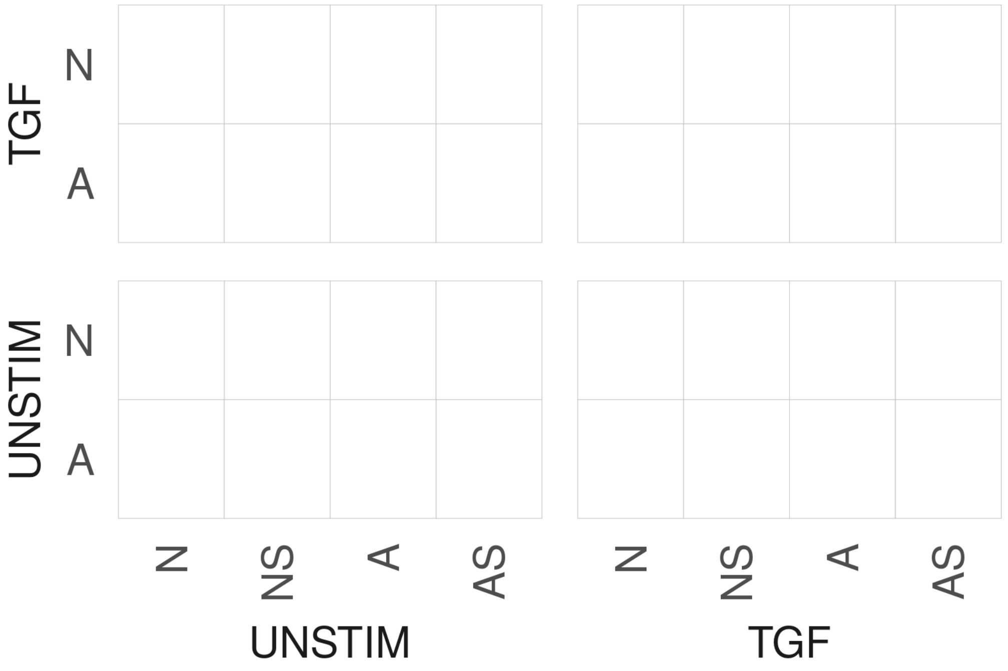
TGF
UNSTIM
N
A
N
NS
A
AS
UNSTIM
TGF

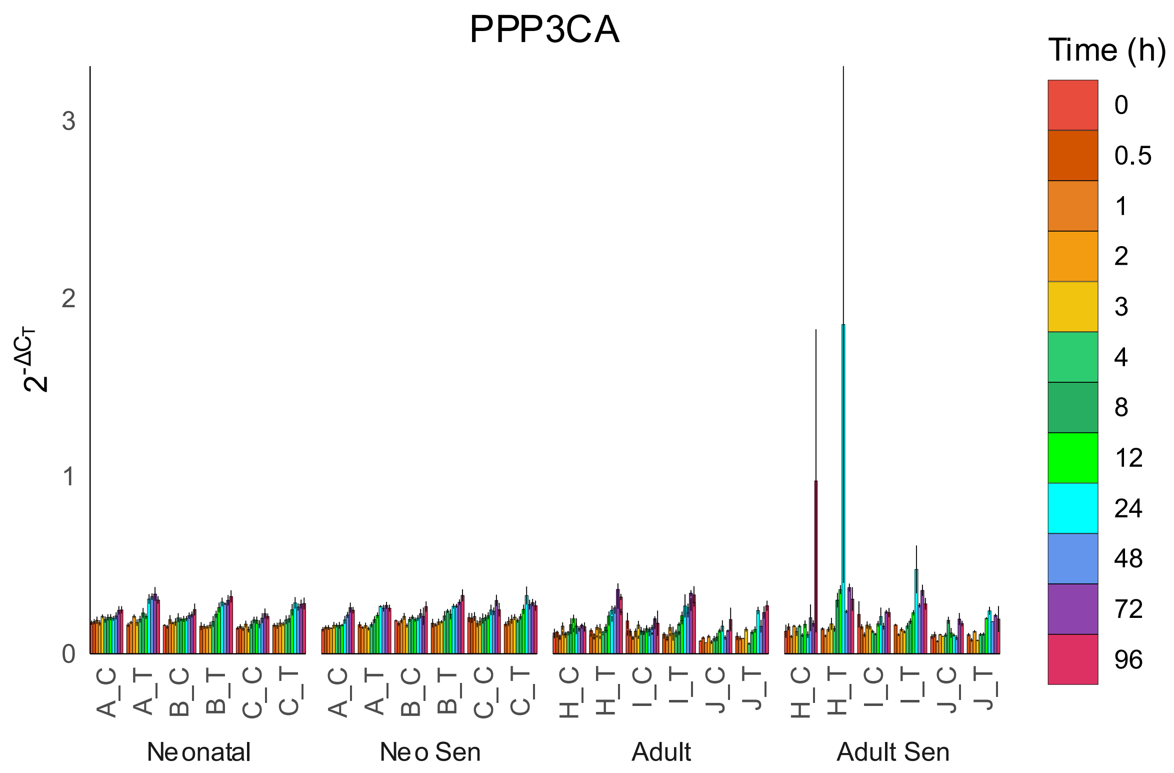
PPP3CA
Time (h)
0
0.5
1
2
3
4
8
12
24
48
72
96
$2^{-\Delta C_T}$
0
1
2
3
A_C
A_T
B_C
B_T
C_C
C_T
H_C
H_T
I_C
I_T
J_C
J_T
Neonatal
Neo Sen
Adult
Adult Sen

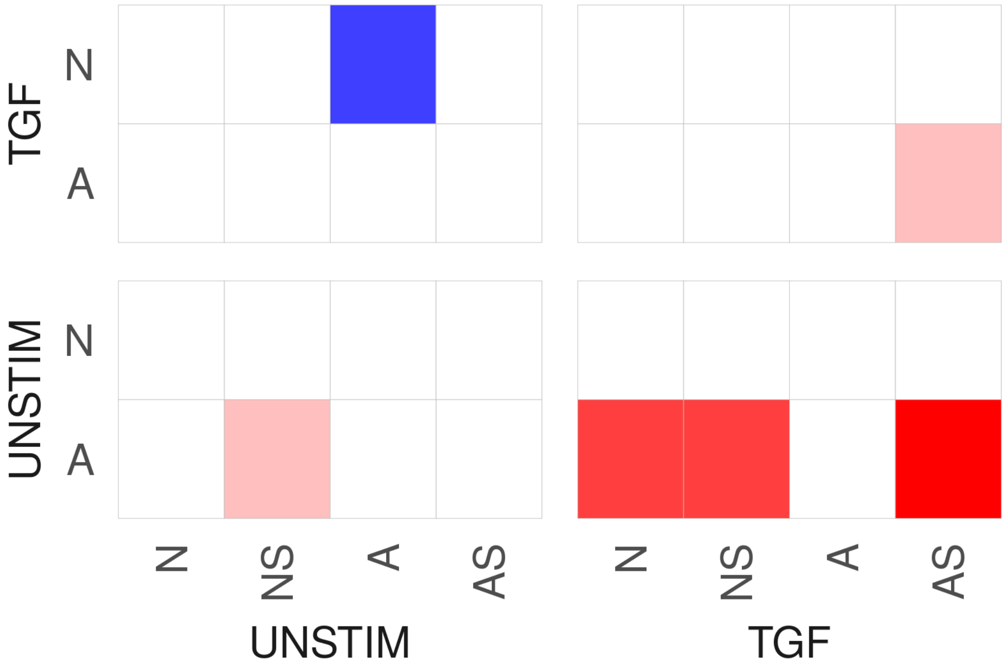
TGF
UNSTIM
N
A
N
NS
A
AS
UNSTIM
TGF

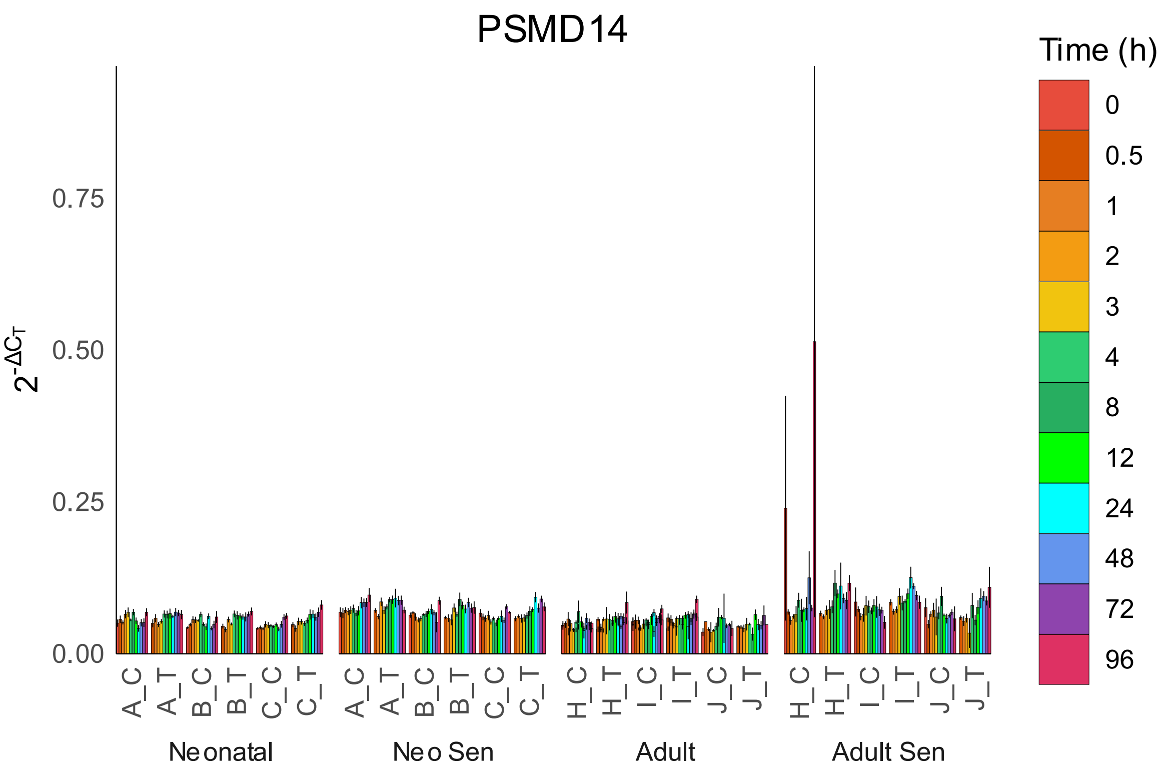
PSMD14
Time (h)
0
0.5
1
2
3
4
8
12
24
48
72
96
0.75
0.50
0.25
0.00
A_C
A_T
B_C
B_T
C_C
C_T
H_C
H_T
I_C
I_T
J_C
J_T
Neonatal
Neo Sen
Adult
Adult Sen

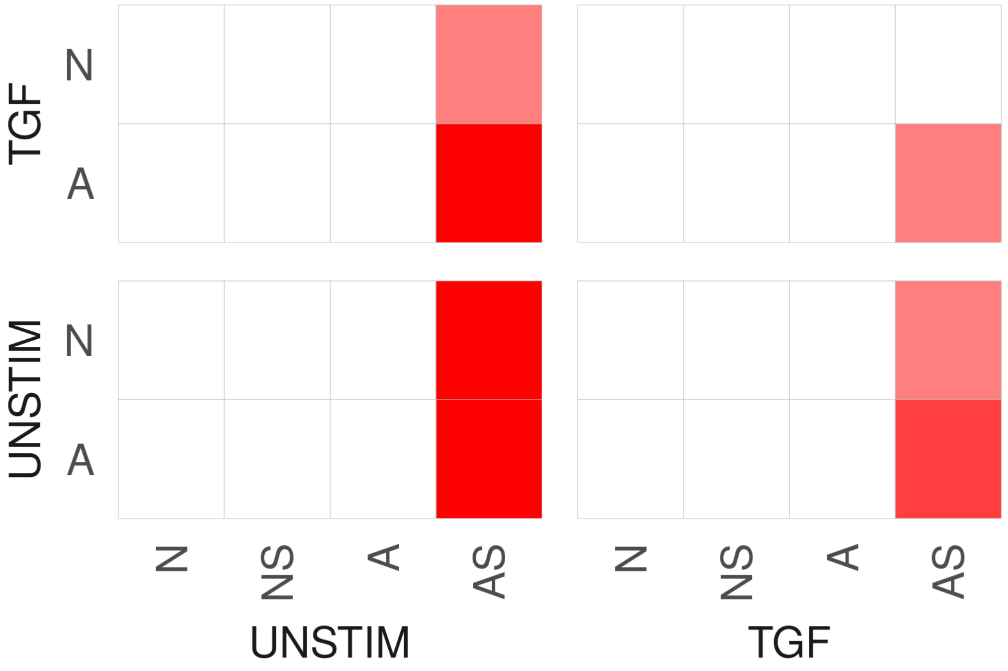
TGF
UNSTIM
N
A
N
NS
A
AS
UNSTIM
TGF

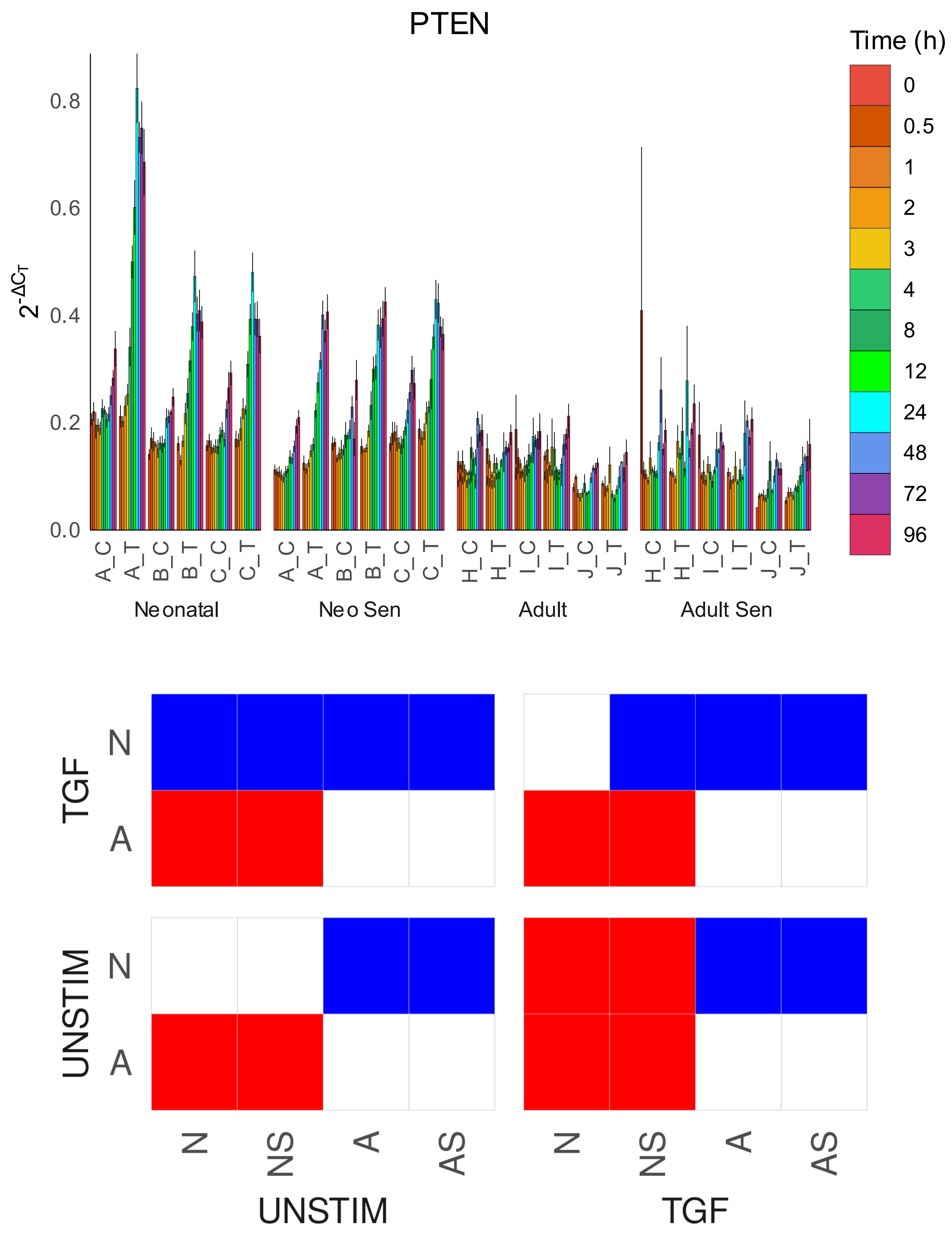
PTEN
Time (h)
0
0.5
1
2
3
4
8
12
24
48
72
96
$2^{-\Delta C_T}$
0.0
0.2
0.4
0.6
0.8
A_C
A_T
B_C
B_T
C_C
C_T
H_C
H_T
I_C
I_T
J_C
J_T
Neonatal
Neo Sen
Adult
Adult Sen
TGF
UNSTIM
N
A
N
NS
A
AS
UNSTIM
TGF

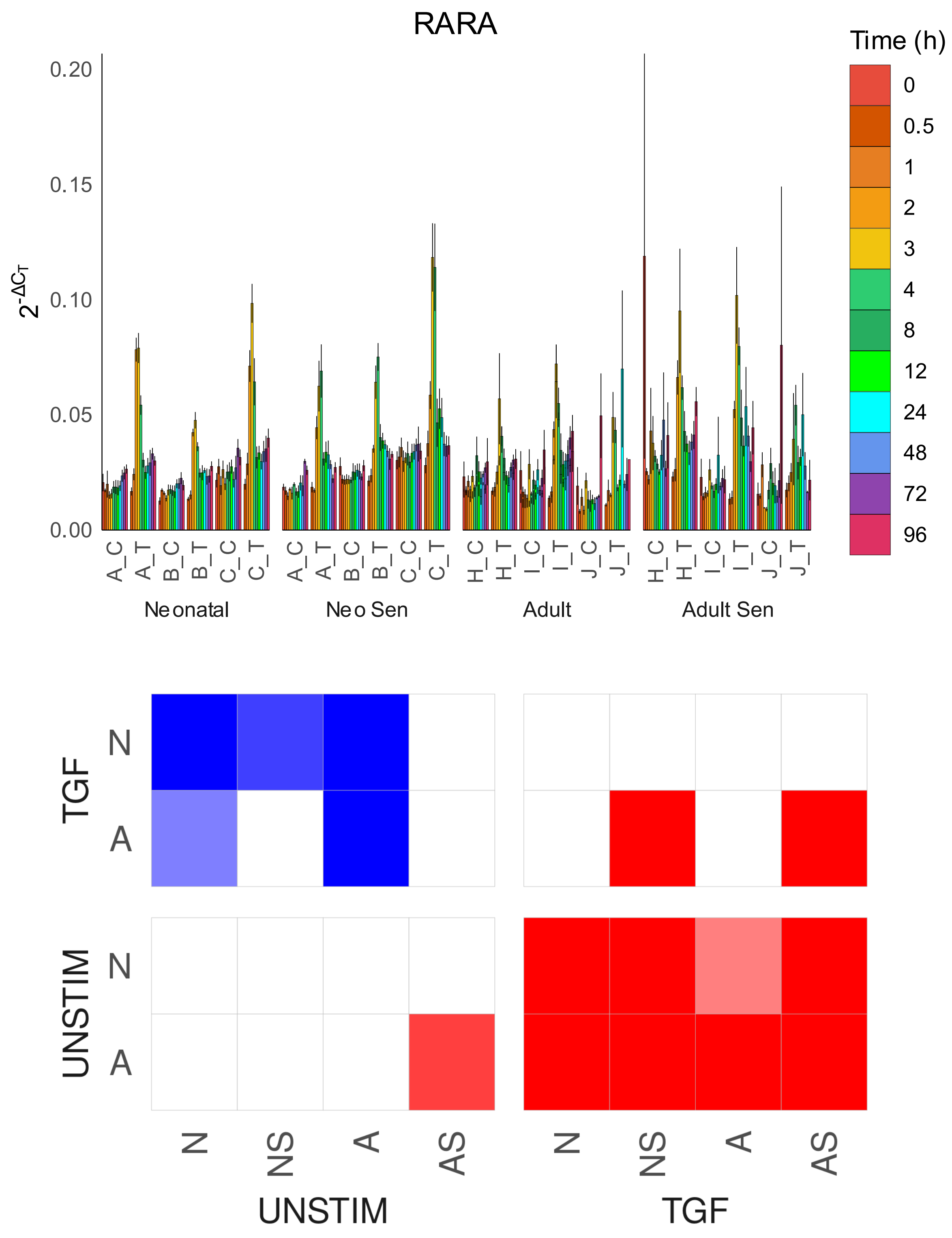

RARA
Time (h)
0
0.5
1
2
3
4
8
12
24
48
72
96
0.20
0.15
0.10
0.05
0.00
$2^{-\Delta C_T}$
A_C
A_T
B_C
B_T
C_C
C_T
H_C
H_T
I_C
I_T
J_C
J_T
Neonatal
Neo Sen
Adult
Adult Sen
TGF
UNSTIM
N
A
NS
AS

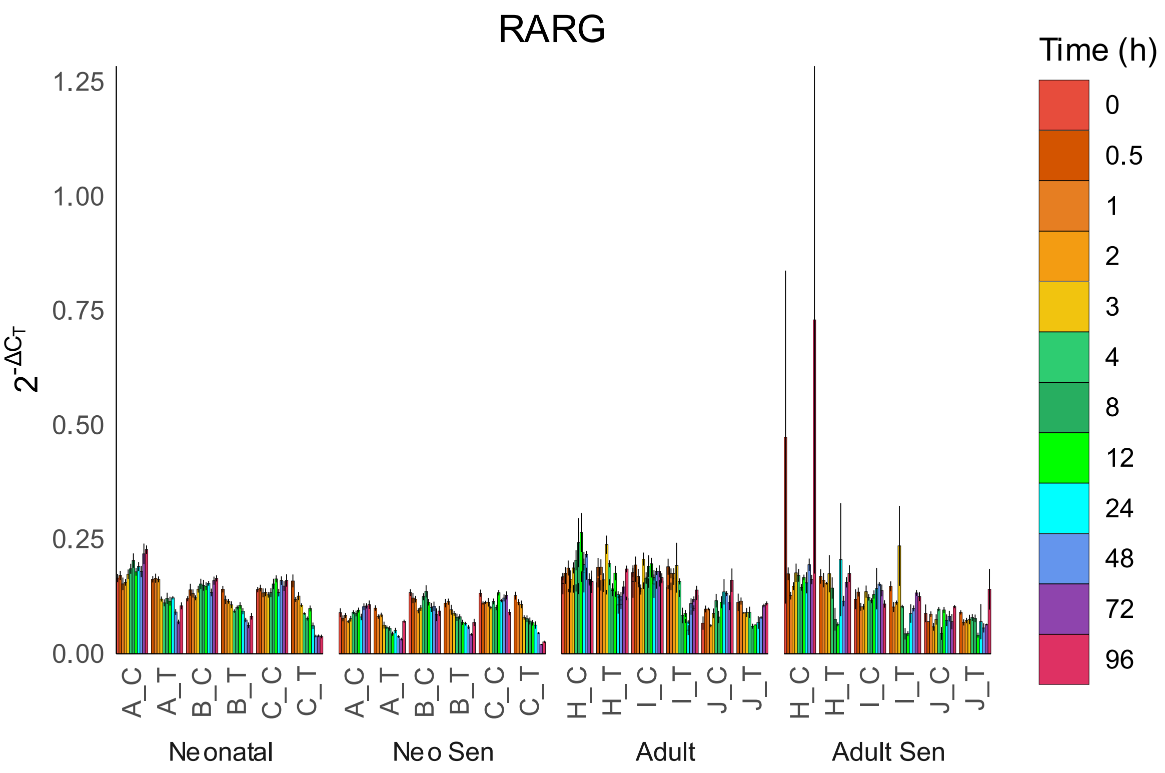
RARG
Time (h)
0
0.5
1
2
3
4
8
12
24
48
72
96
1.25
1.00
0.75
0.50
0.25
0.00
A_C
A_T
B_C
B_T
C_C
C_T
H_C
H_T
I_C
I_T
J_C
J_T
Neonatal
Neo Sen
Adult
Adult Sen

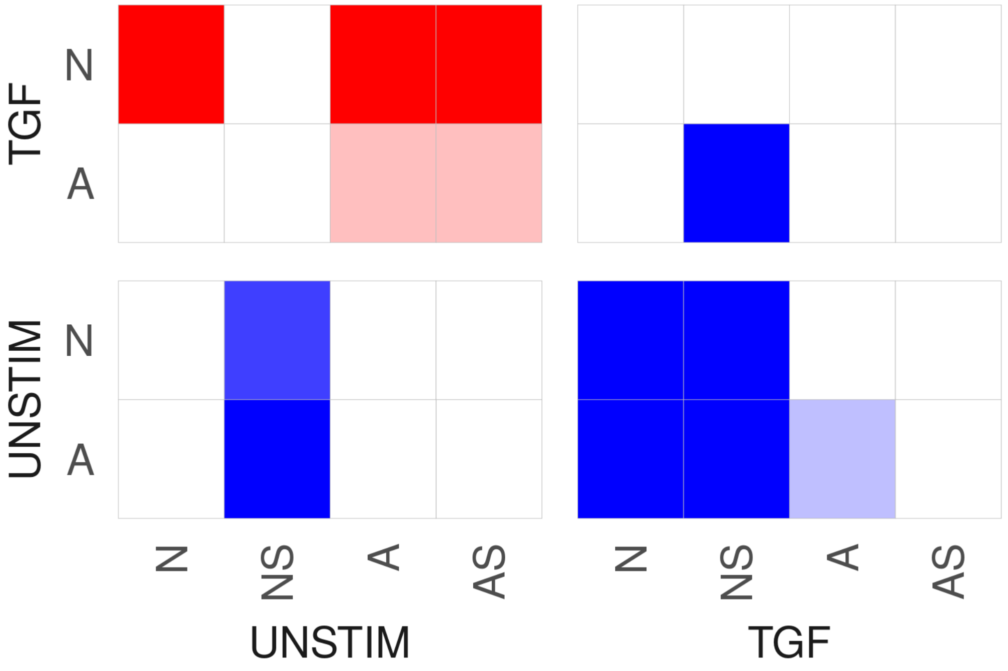
TGF
N
A
UNSTIM
N
A
N
NS
A
AS
UNSTIM
TGF

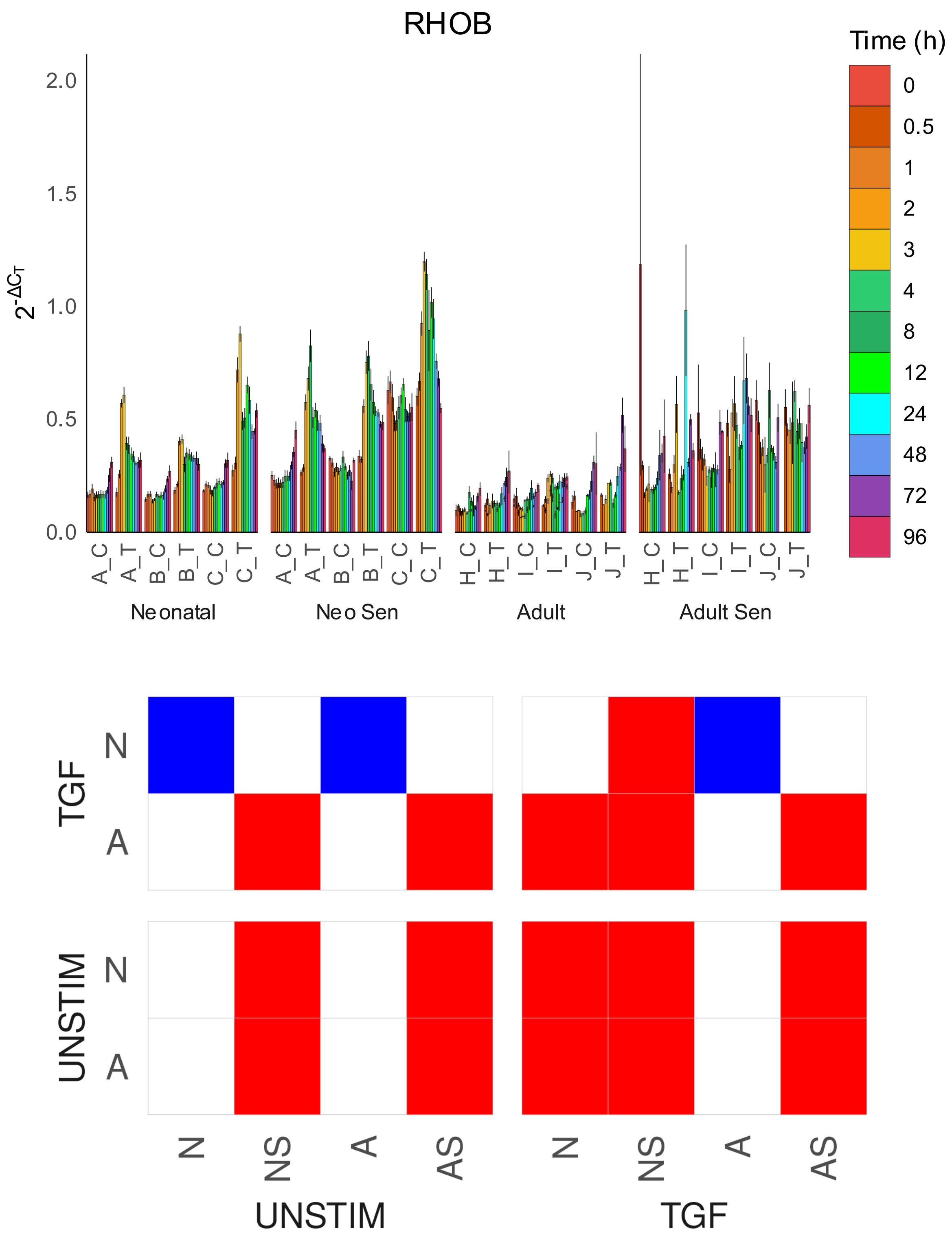
RHOB
Time (h)
0
0.5
1
2
3
4
8
12
24
48
72
96
2.0
1.5
1.0
0.5
0.0
$2^{-\Delta C_T}$
A_C
A_T
B_C
B_T
C_C
C_T
H_C
H_T
I_C
I_T
J_C
J_T
Neonatal
Neo Sen
Adult
Adult Sen
TGF
UNSTIM
N
A
N
NS
A
AS
UNSTIM
TGF

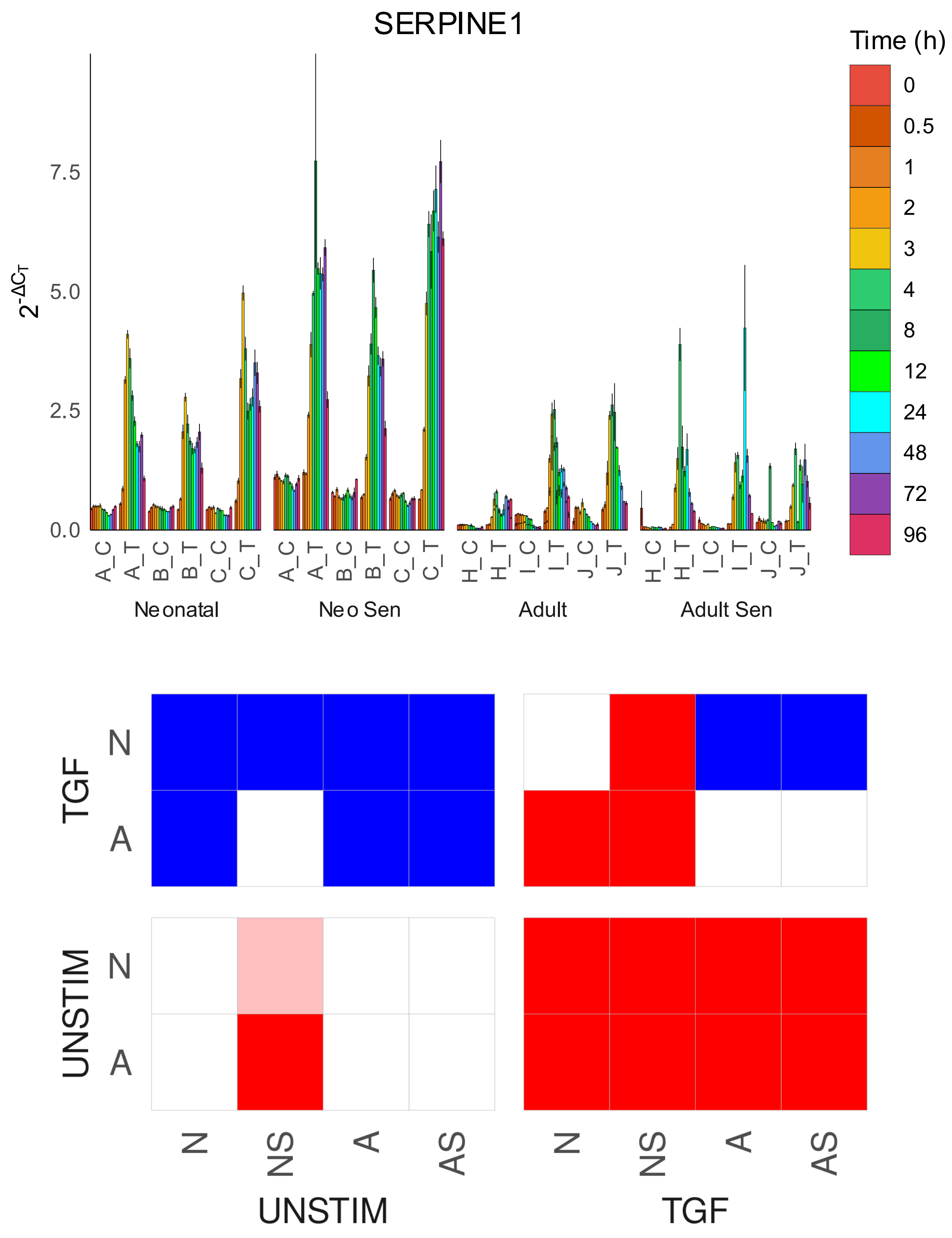
SERPINE1
Time (h)
0
0.5
1
2
3
4
8
12
24
48
72
96
2-ΔCT
7.5
5.0
2.5
0.0
A_C
A_T
B_C
B_T
C_C
C_T
H_C
H_T
I_C
I_T
J_C
J_T
Neonatal
Neo Sen
Adult
Adult Sen
TGF
UNSTIM
N
A
N
NS
A
AS
UNSTIM
TGF

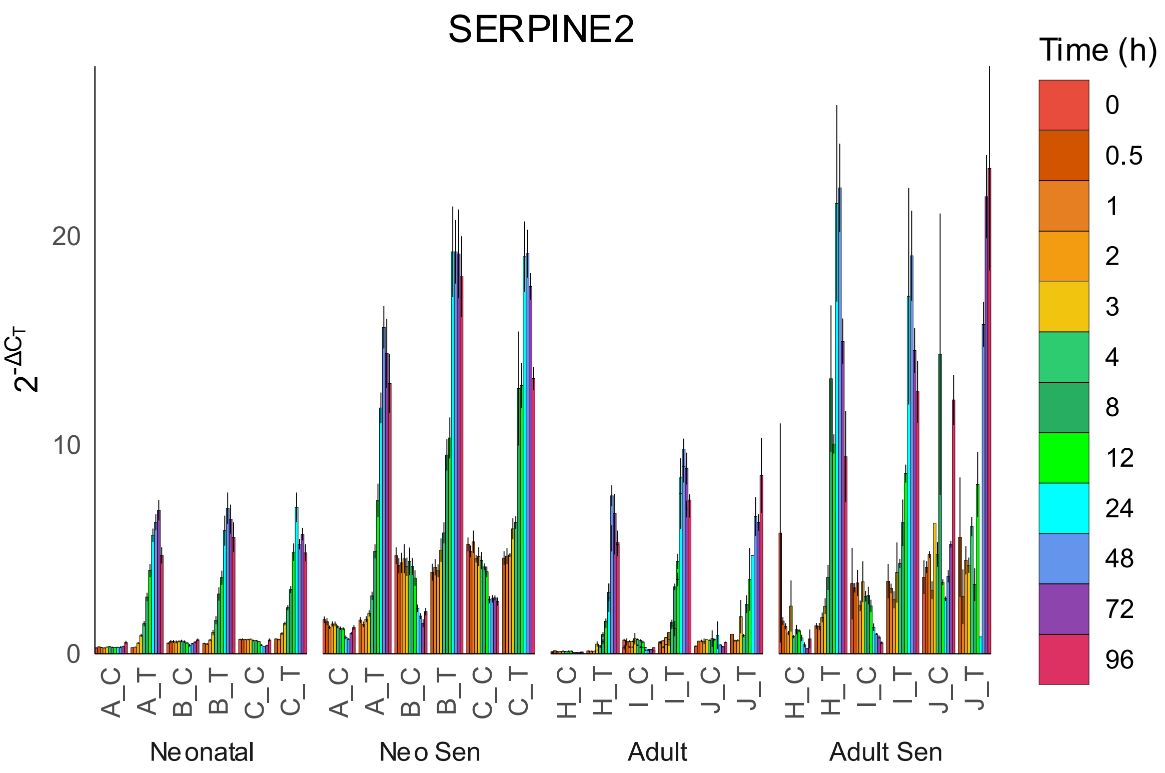

SERPINE2
Time (h)
0
0.5
1
2
3
4
8
12
24
48
72
96
2-ΔCT
20
10
0
A_C
A_T
B_C
B_T
C_C
C_T
A_C
A_T
B_C
B_T
C_C
C_T
H_C
H_T
I_C
I_T
J_C
J_T
H_C
H_T
I_C
I_T
J_C
J_T
Neonatal
Neo Sen
Adult
Adult Sen


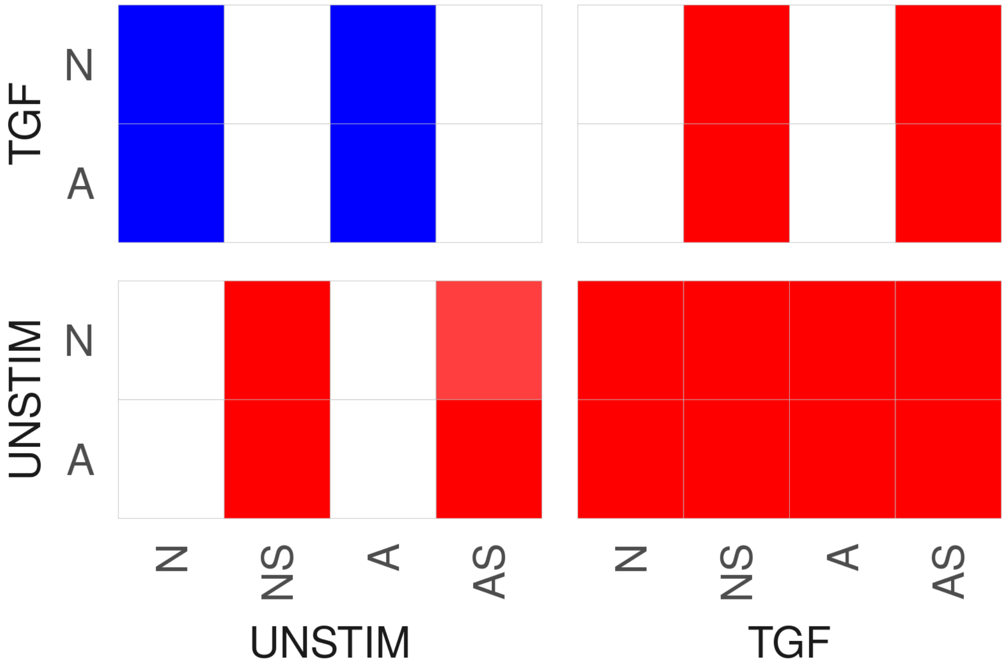

TGF
N
A
UNSTIM
N
A
N
NS
A
AS
N
NS
A
AS
UNSTIM
TGF

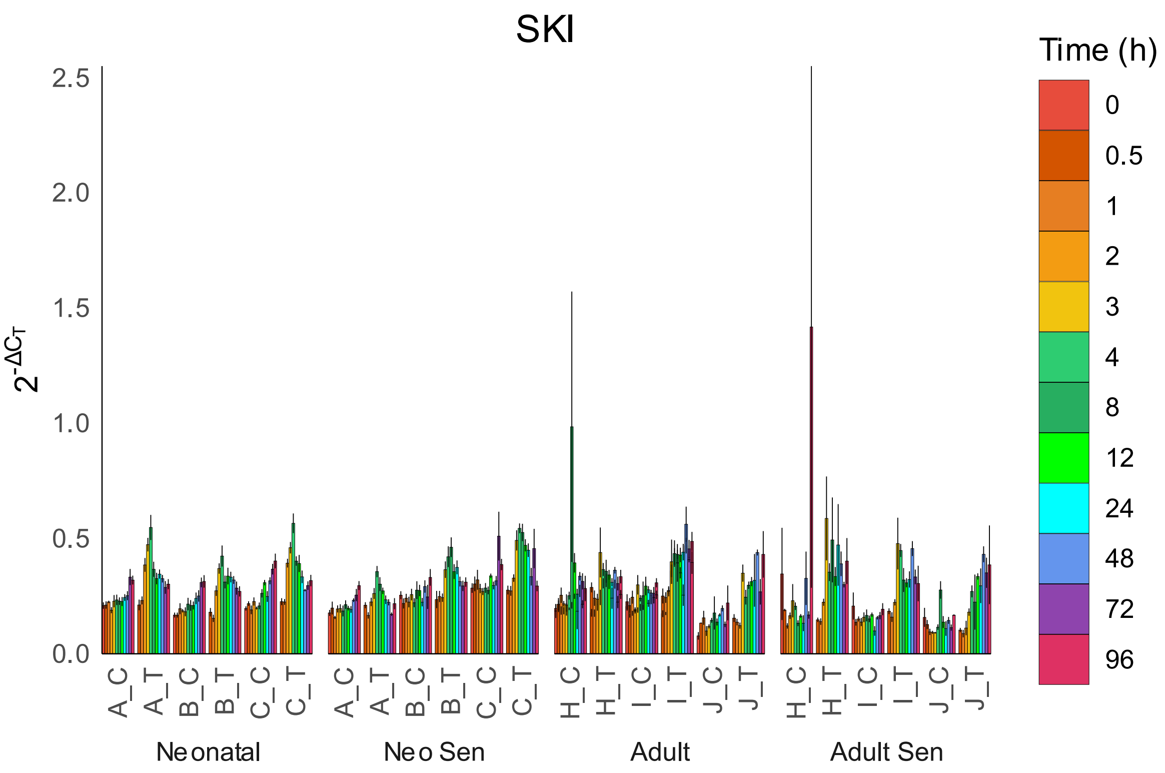
SKI
Time (h)
0
0.5
1
2
3
4
8
12
24
48
72
96
2.5
2.0
1.5
1.0
0.5
0.0
$2^{-\Delta C_T}$
A_C
A_T
B_C
B_T
C_C
C_T
H_C
H_T
I_C
I_T
J_C
J_T
Neonatal
Neo Sen
Adult
Adult Sen

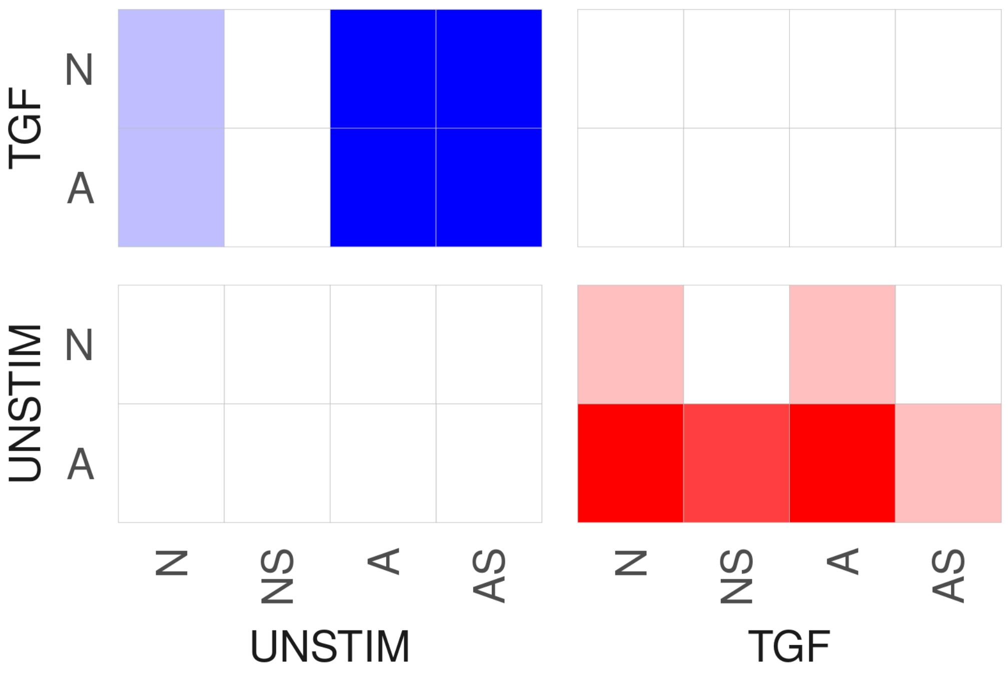
TGF
N
A
UNSTIM
N
A
N
NS
A
AS
UNSTIM
N
NS
A
AS
TGF

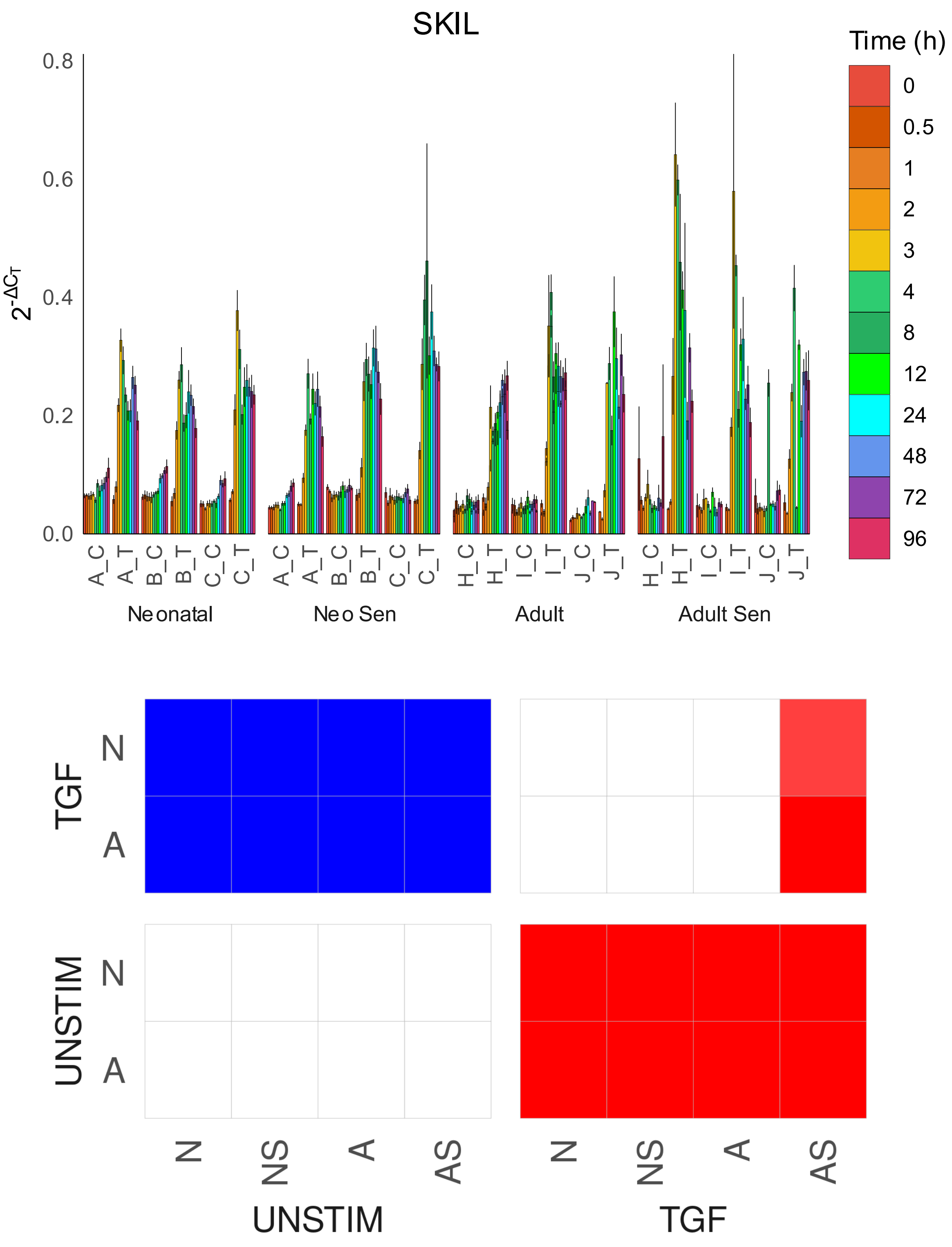
SKIL
Time (h)
0
0.5
1
2
3
4
8
12
24
48
72
96
0.0
0.2
0.4
0.6
0.8
$2^{-\Delta C_T}$
A_C A_T B_C B_T C_C C_T
A_C A_T B_C B_T C_C C_T
H_C H_T I_C I_T J_C J_T
H_C H_T I_C I_T J_C J_T
Neonatal
Neo Sen
Adult
Adult Sen
TGF
UNSTIM
N
A
N
A
N NS A AS
N NS A AS
UNSTIM
TGF

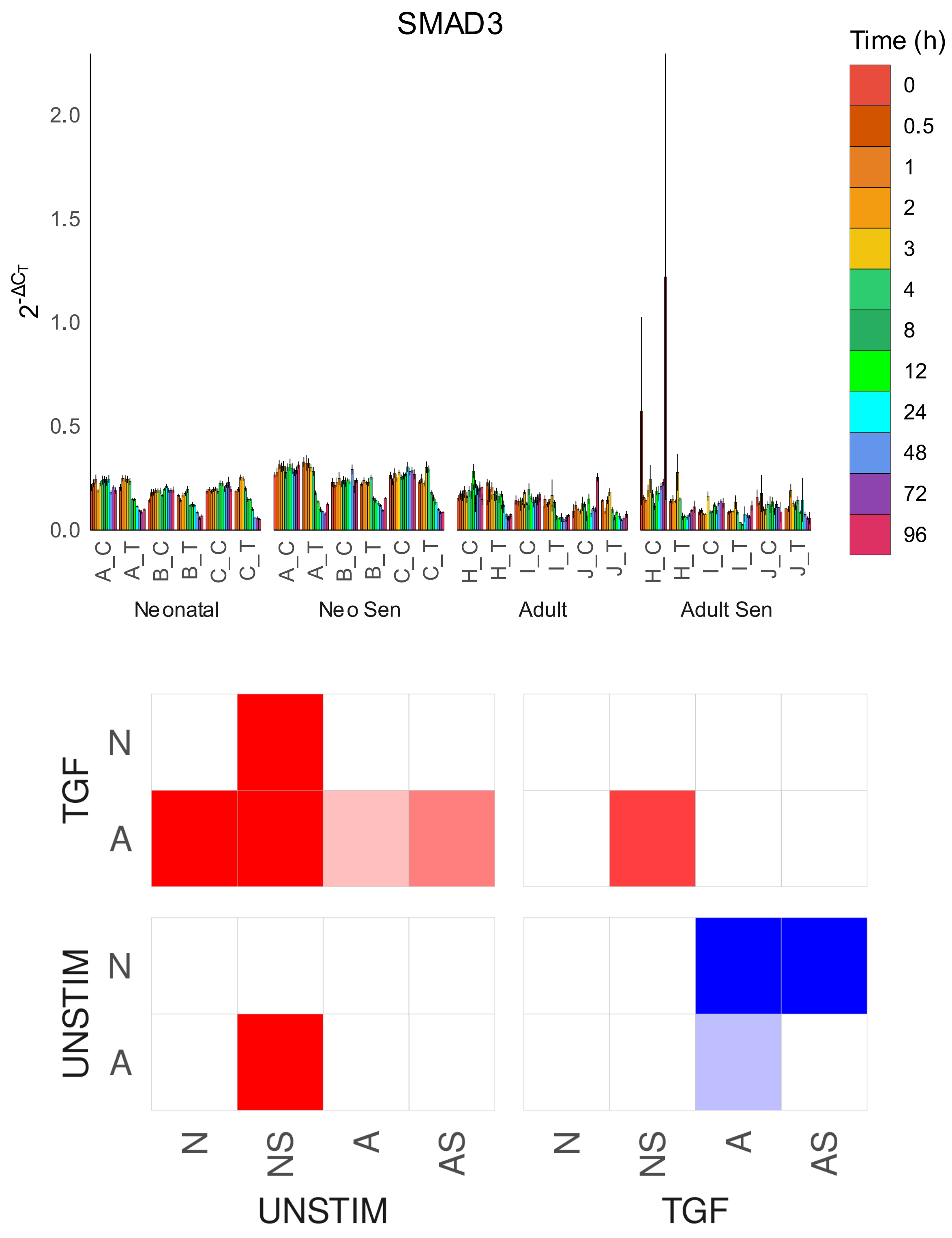

SMAD3
Time (h)
0
0.5
1
2
3
4
8
12
24
48
72
96
$2^{-\Delta C_T}$
2.0
1.5
1.0
0.5
0.0
A_C
A_T
B_C
B_T
C_C
C_T
H_C
H_T
I_C
I_T
J_C
J_T
Neonatal
Neo Sen
Adult
Adult Sen
TGF
UNSTIM
N
A
N
NS
A
AS
UNSTIM
TGF

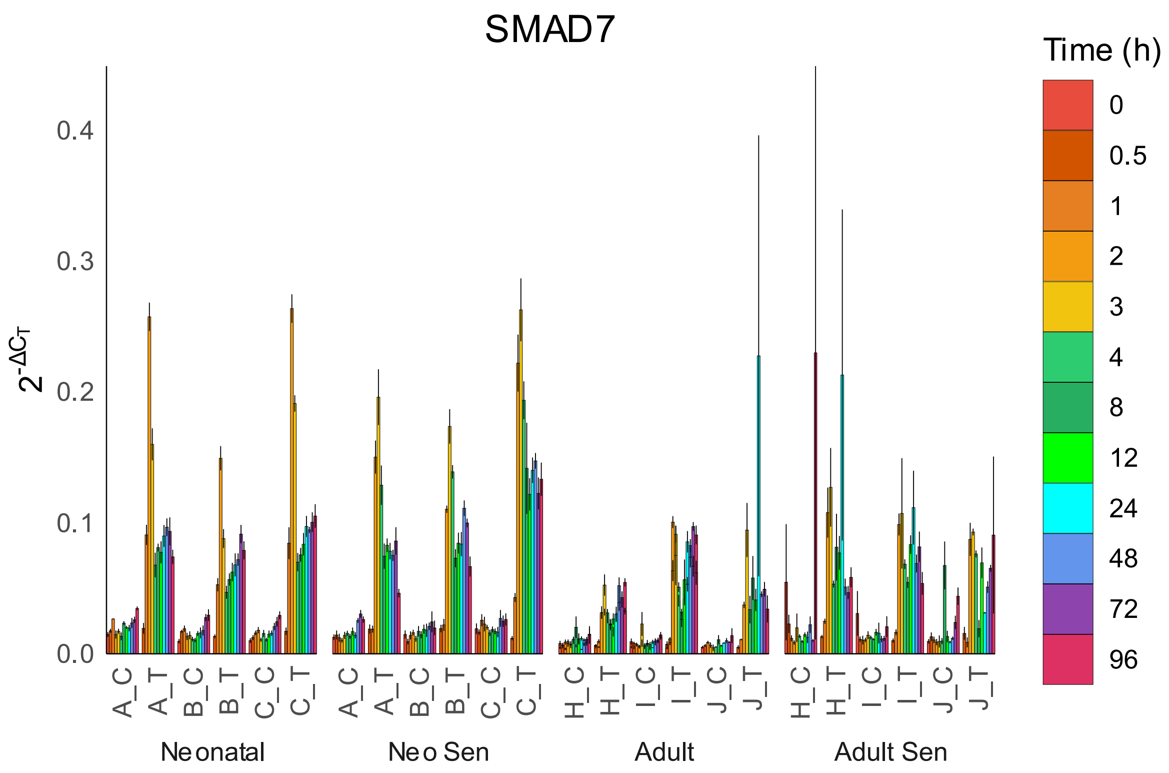

SMAD7
Time (h)
0
0.5
1
2
3
4
8
12
24
48
72
96
$2^{-\Delta C_T}$
0.0
0.1
0.2
0.3
0.4
A_C
A_T
B_C
B_T
C_C
C_T
H_C
H_T
I_C
I_T
J_C
J_T
Neonatal
Neo Sen
Adult
Adult Sen


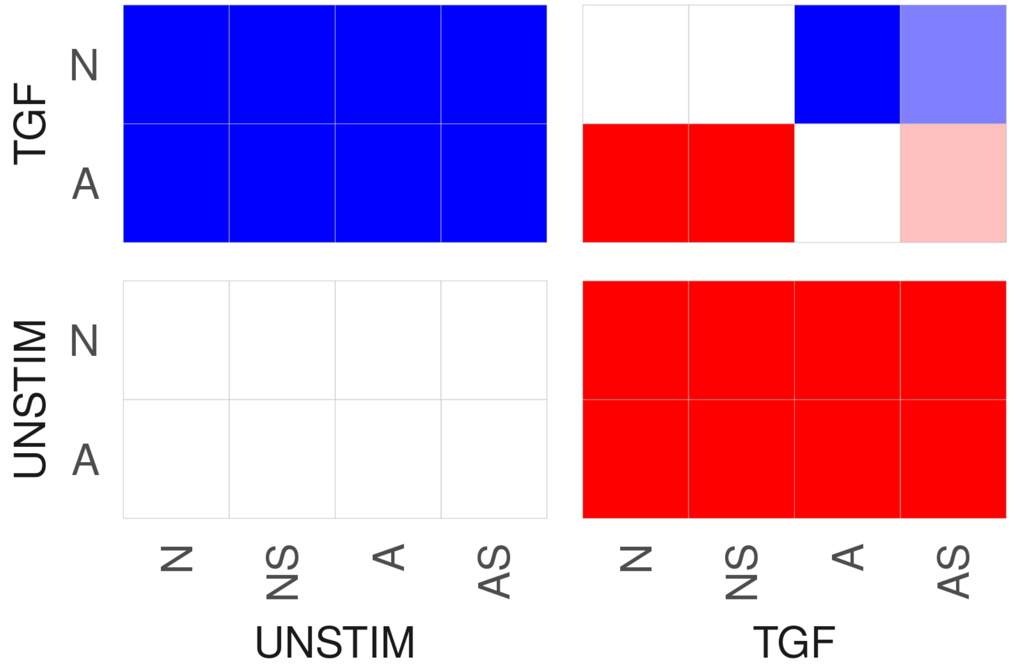

TGF
N
A
UNSTIM
N
A
N
NS
A
AS
UNSTIM
TGF

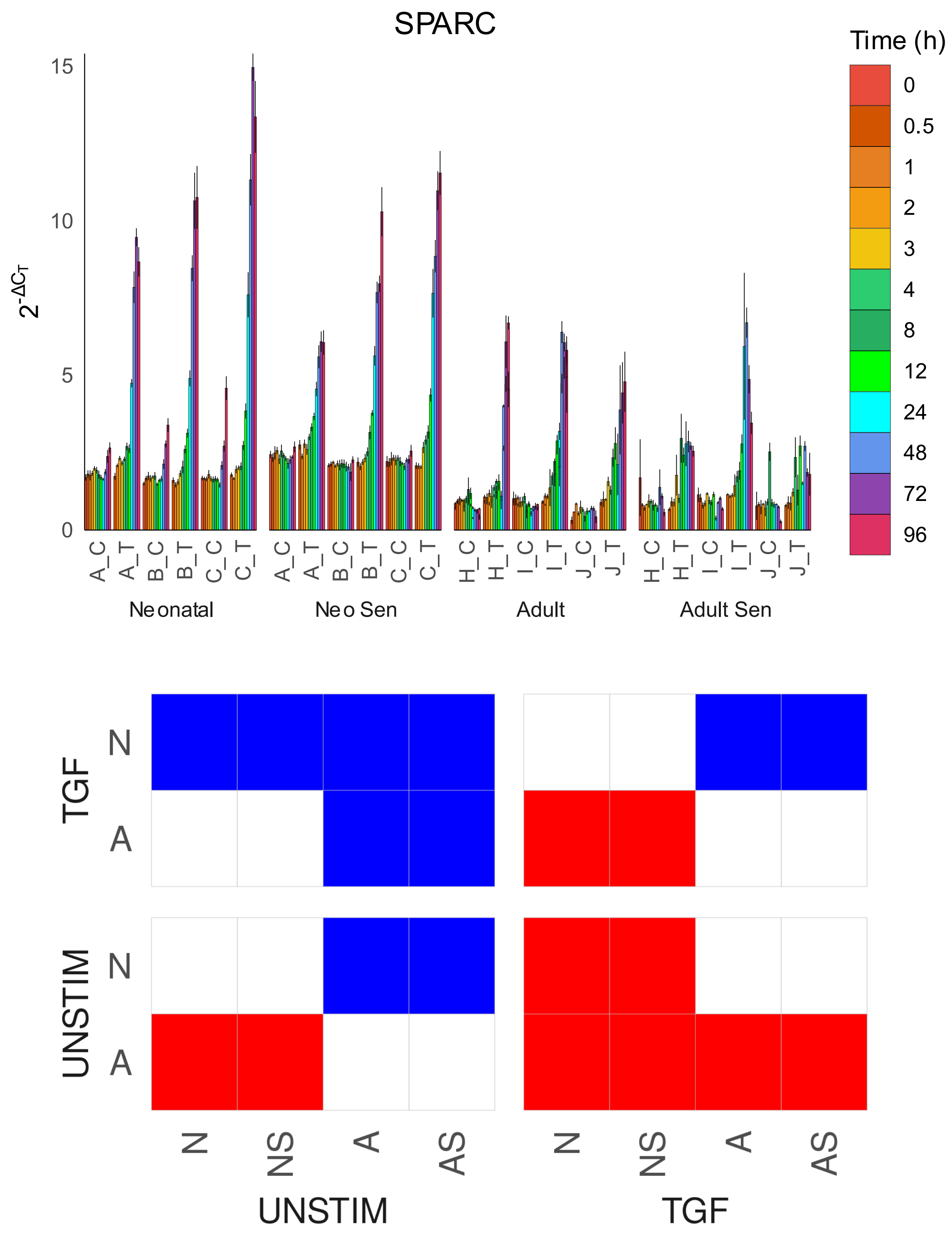

SPARC
Time (h)
0
0.5
1
2
3
4
8
12
24
48
72
96
15
10
5
0
$2^{-\Delta C_T}$
A_C
A_T
B_C
B_T
C_C
C_T
H_C
H_T
I_C
I_T
J_C
J_T
Neonatal
Neo Sen
Adult
Adult Sen
TGF
N
A
UNSTIM
N
A
N
NS
A
AS
UNSTIM
N
NS
A
AS
TGF

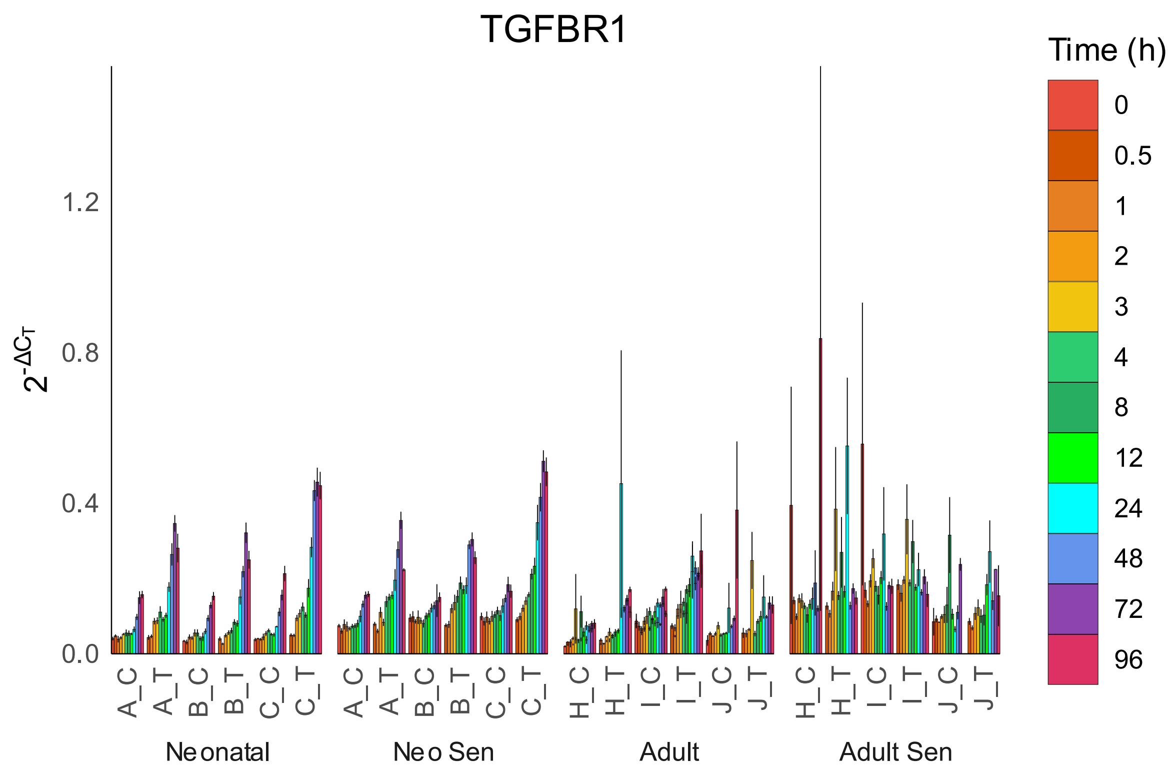
TGFBR1
Time (h)
0
0.5
1
2
3
4
8
12
24
48
72
96
$2^{-\Delta C_T}$
1.2
0.8
0.4
0.0
A_C
A_T
B_C
B_T
C_C
C_T
H_C
H_T
I_C
I_T
J_C
J_T
Neonatal
Neo Sen
Adult
Adult Sen

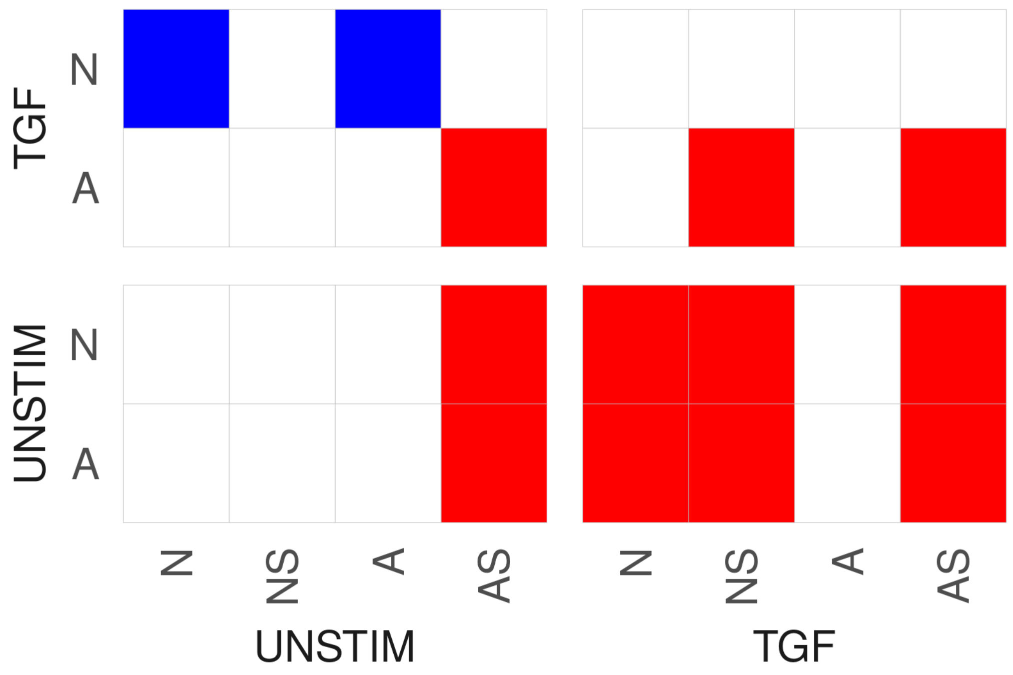
TGF
UNSTIM
N
A
N
NS
A
AS
UNSTIM
TGF

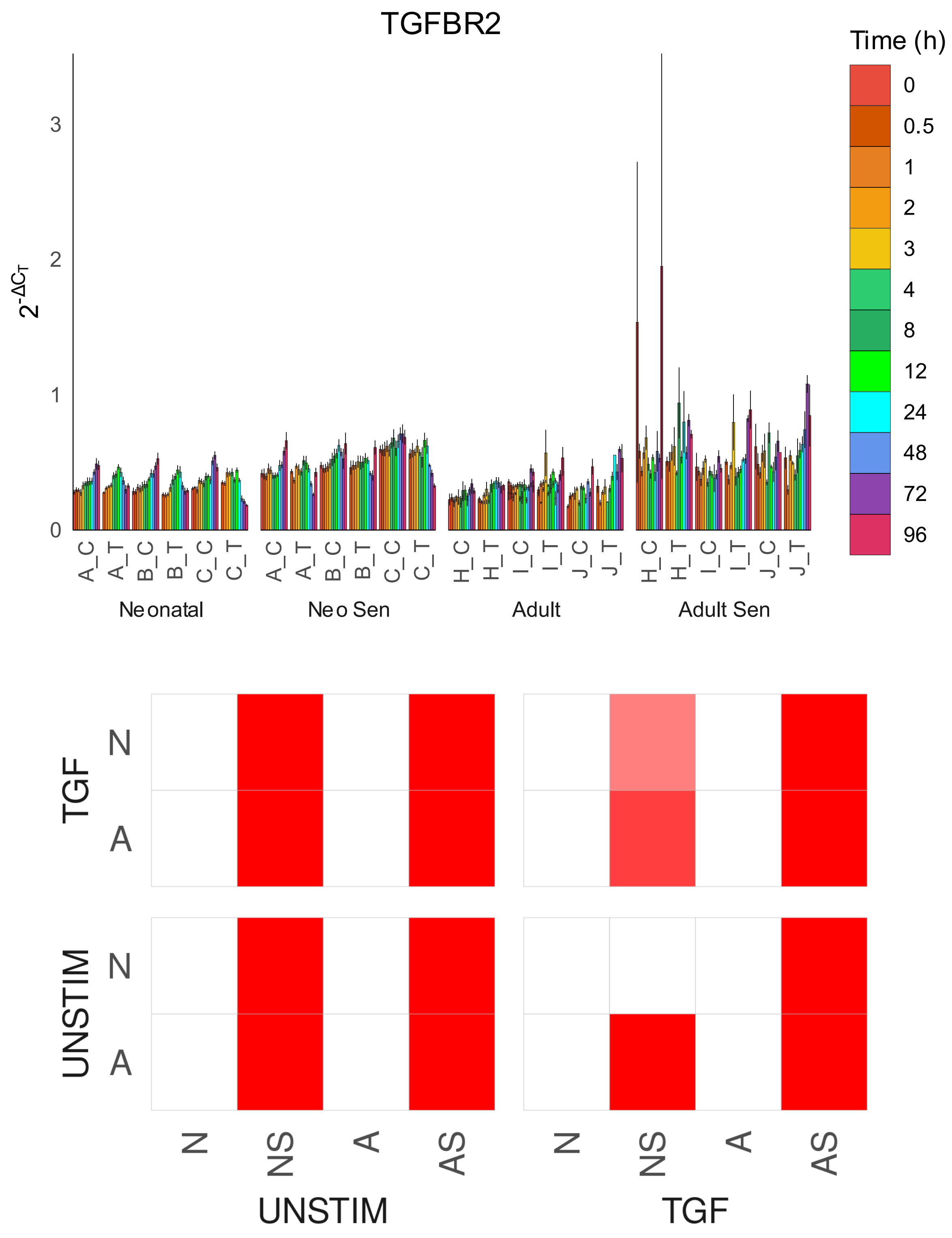
TGFBR2
Time (h)
0
0.5
1
2
3
4
8
12
24
48
72
96
$2^{-\Delta C_T}$
0
1
2
3
A_C
A_T
B_C
B_T
C_C
C_T
H_C
H_T
I_C
I_T
J_C
J_T
Neonatal
Neo Sen
Adult
Adult Sen
TGF
UNSTIM
N
A
N
NS
A
AS
UNSTIM
TGF

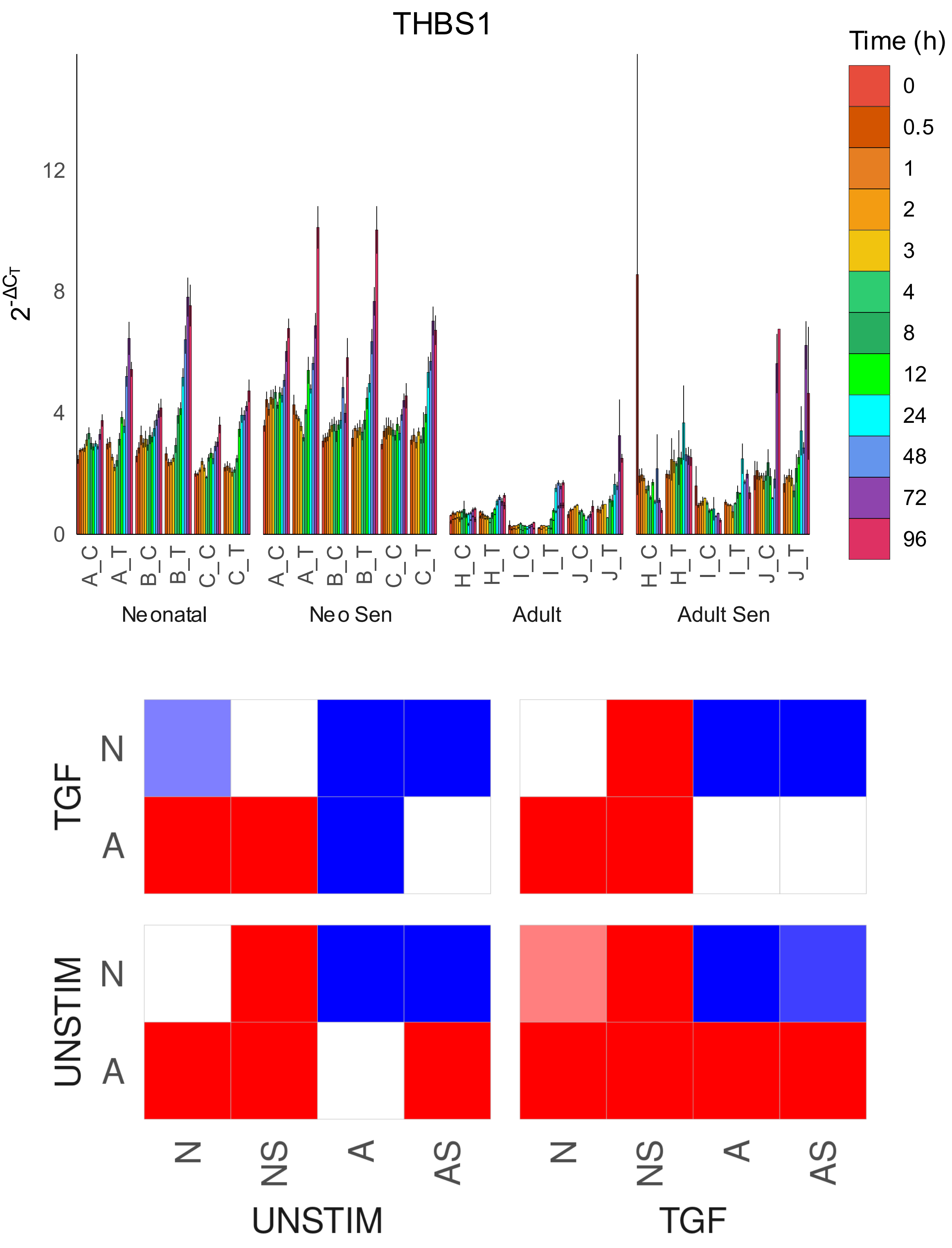

THBS1
Time (h)
0
0.5
1
2
3
4
8
12
24
48
72
96
$2^{-\Delta C_T}$
0
4
8
12
A_C
A_T
B_C
B_T
C_C
C_T
H_C
H_T
I_C
I_T
J_C
J_T
Neonatal
Neo Sen
Adult
Adult Sen
TGF
UNSTIM
N
A
N
A
N
NS
A
AS
N
NS
A
AS
UNSTIM
TGF

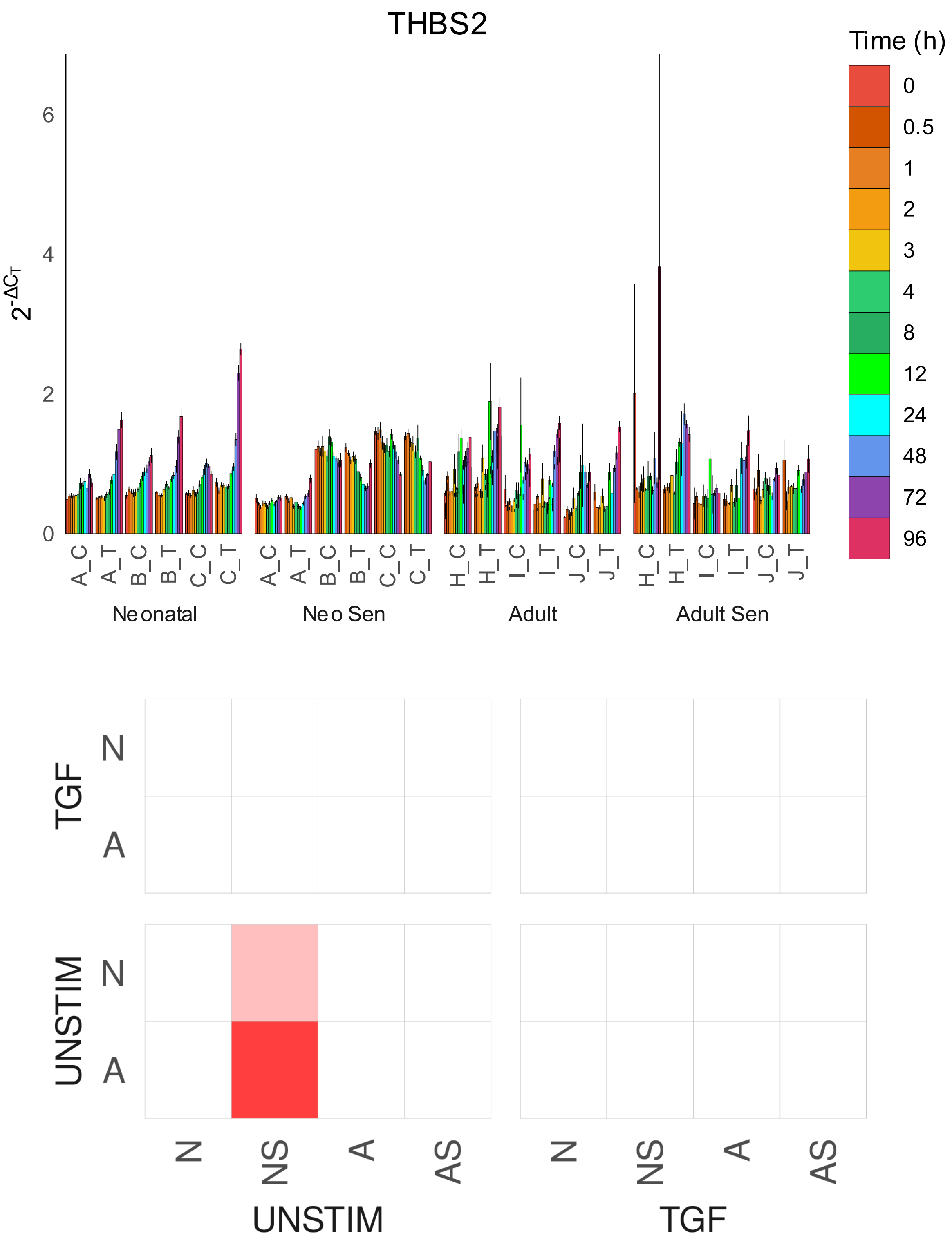
THBS2
Time (h)
0
0.5
1
2
3
4
8
12
24
48
72
96
$2^{-\Delta C_T}$
0
2
4
6
A_C
A_T
B_C
B_T
C_C
C_T
H_C
H_T
I_C
I_T
J_C
J_T
Neonatal
Neo Sen
Adult
Adult Sen
TGF
UNSTIM
N
A
N
NS
A
AS
UNSTIM
TGF

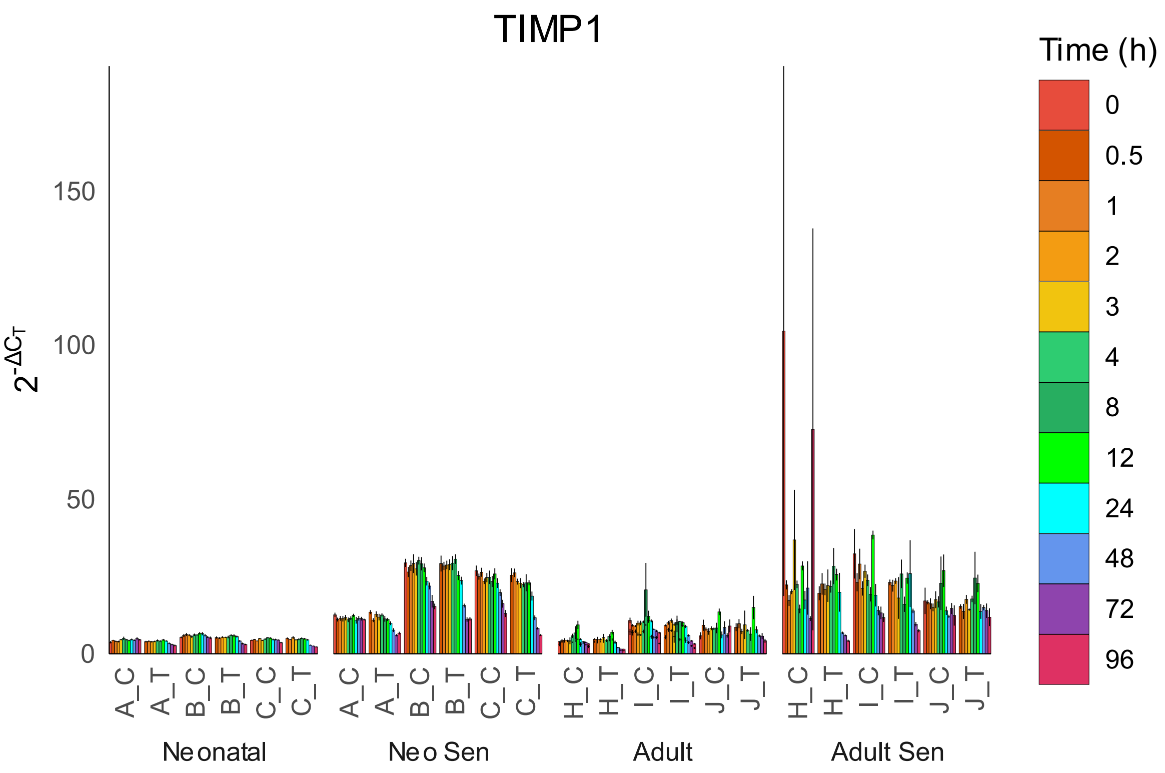
TIMP1
Time (h)
0
0.5
1
2
3
4
8
12
24
48
72
96
$2^{-\Delta C_T}$
150
100
50
0
A_C
A_T
B_C
B_T
C_C
C_T
H_C
H_T
I_C
I_T
J_C
J_T
Neonatal
Neo Sen
Adult
Adult Sen

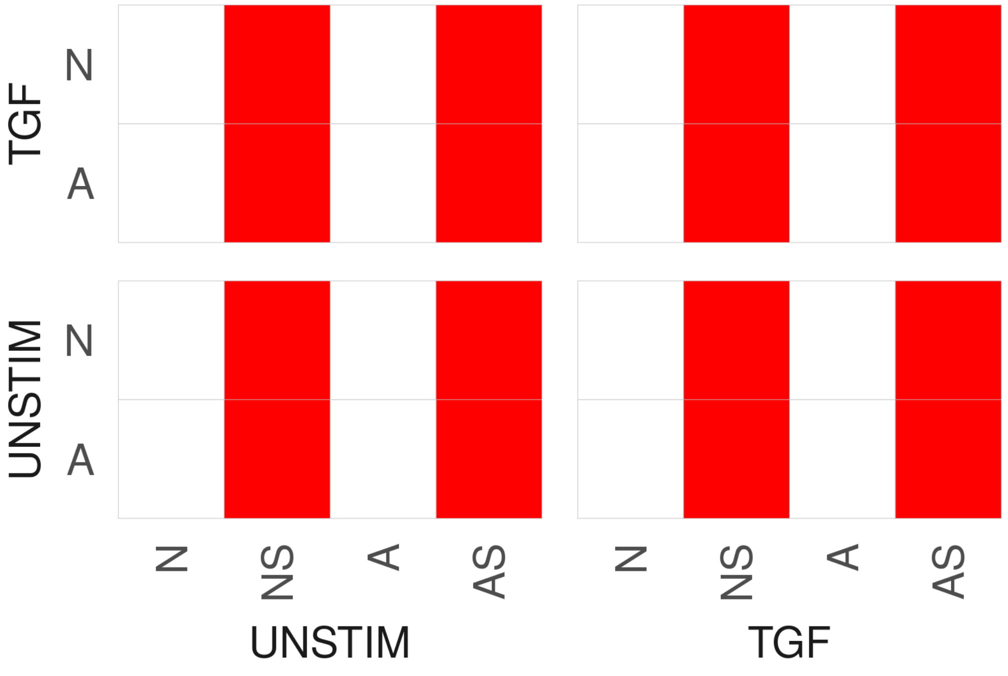
TGF
N
A
UNSTIM
N
A
N
NS
A
AS
UNSTIM
N
NS
A
AS
TGF

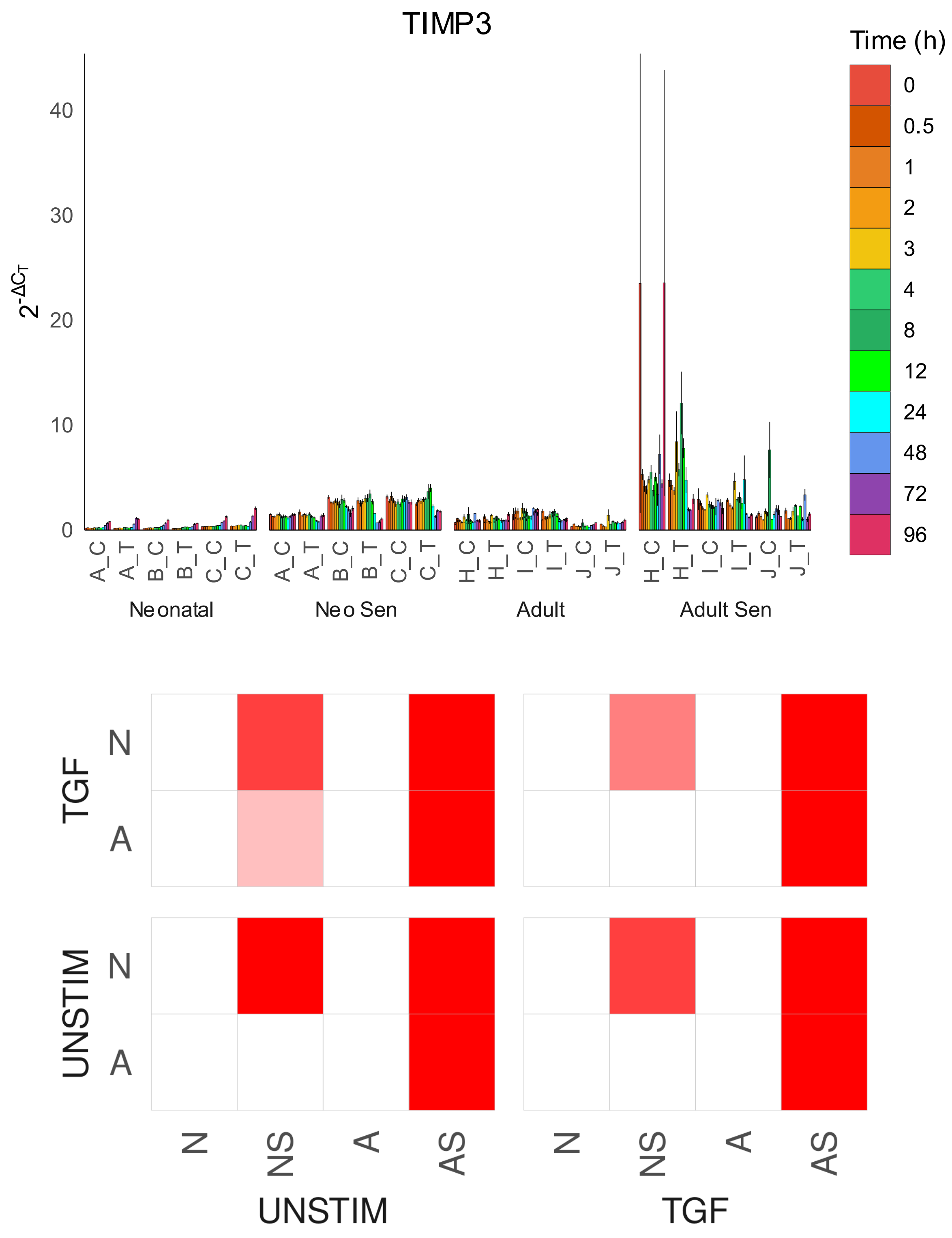
TIMP3
Time (h)
0
0.5
1
2
3
4
8
12
24
48
72
96
$2^{-\Delta C_T}$
40
30
20
10
0
A_C
A_T
B_C
B_T
C_C
C_T
H_C
H_T
I_C
I_T
J_C
J_T
Neonatal
Neo Sen
Adult
Adult Sen
TGF
UNSTIM
N
A
N
NS
A
AS
UNSTIM
TGF

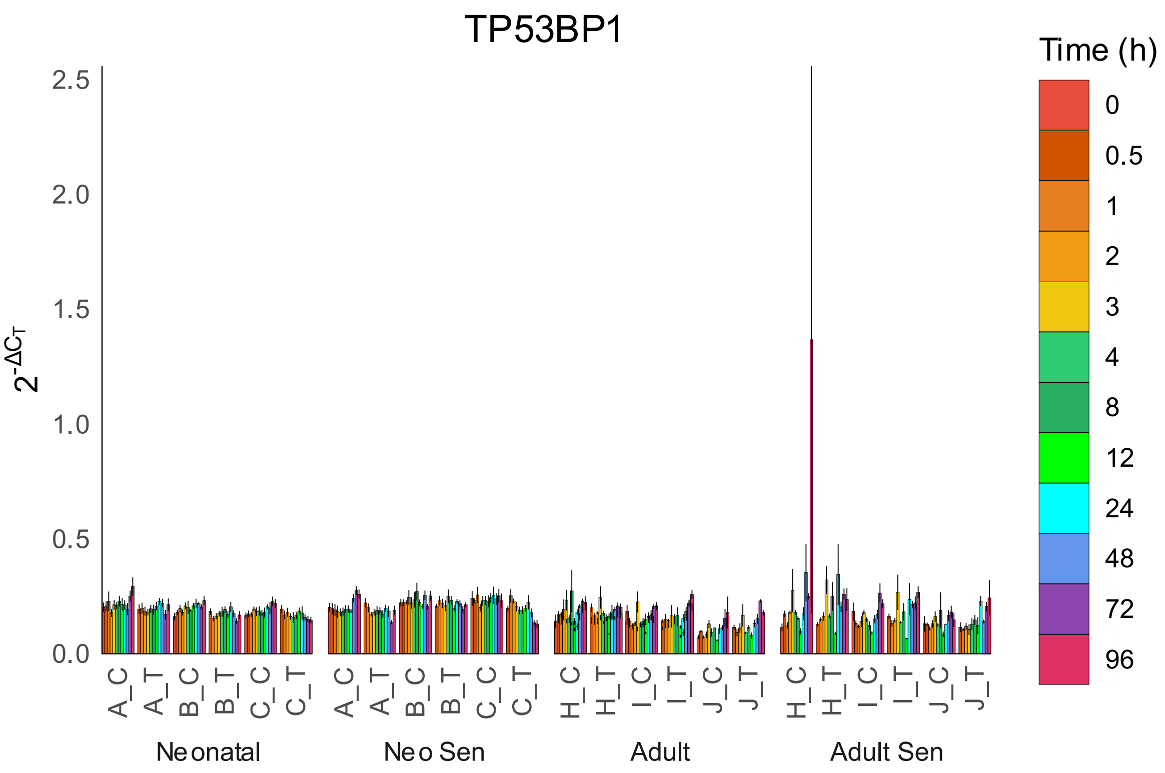

TP53BP1
Time (h)
0
0.5
1
2
3
4
8
12
24
48
72
96
2.5
2.0
1.5
1.0
0.5
0.0
$2^{-\Delta C_T}$
A_C
A_T
B_C
B_T
C_C
C_T
H_C
H_T
I_C
I_T
J_C
J_T
Neonatal
Neo Sen
Adult
Adult Sen


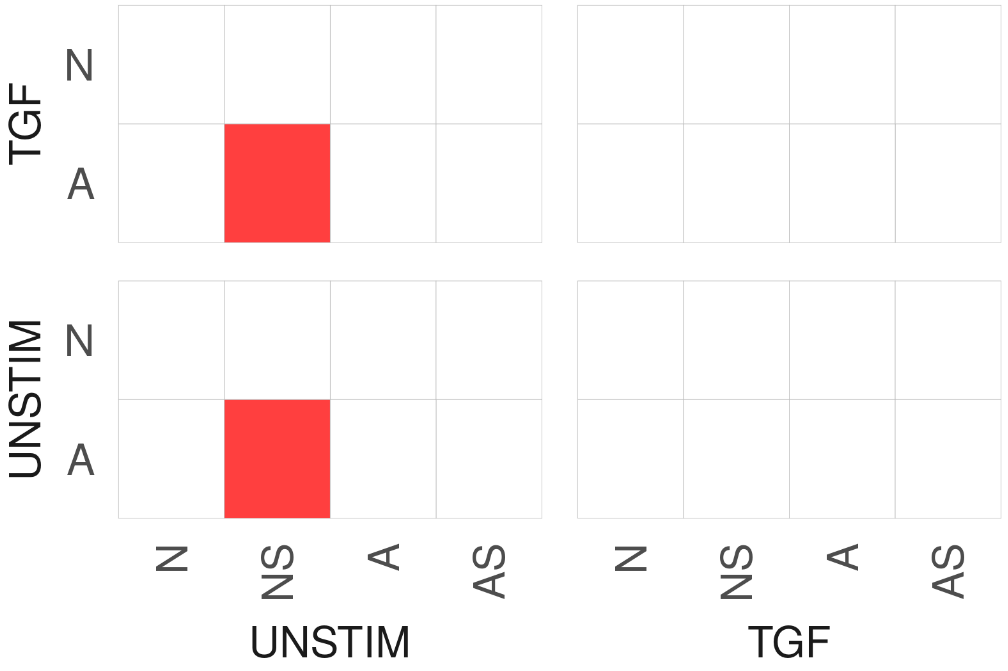

TGF
N
A
UNSTIM
N
A
N
NS
A
AS
UNSTIM
N
NS
A
AS
TGF

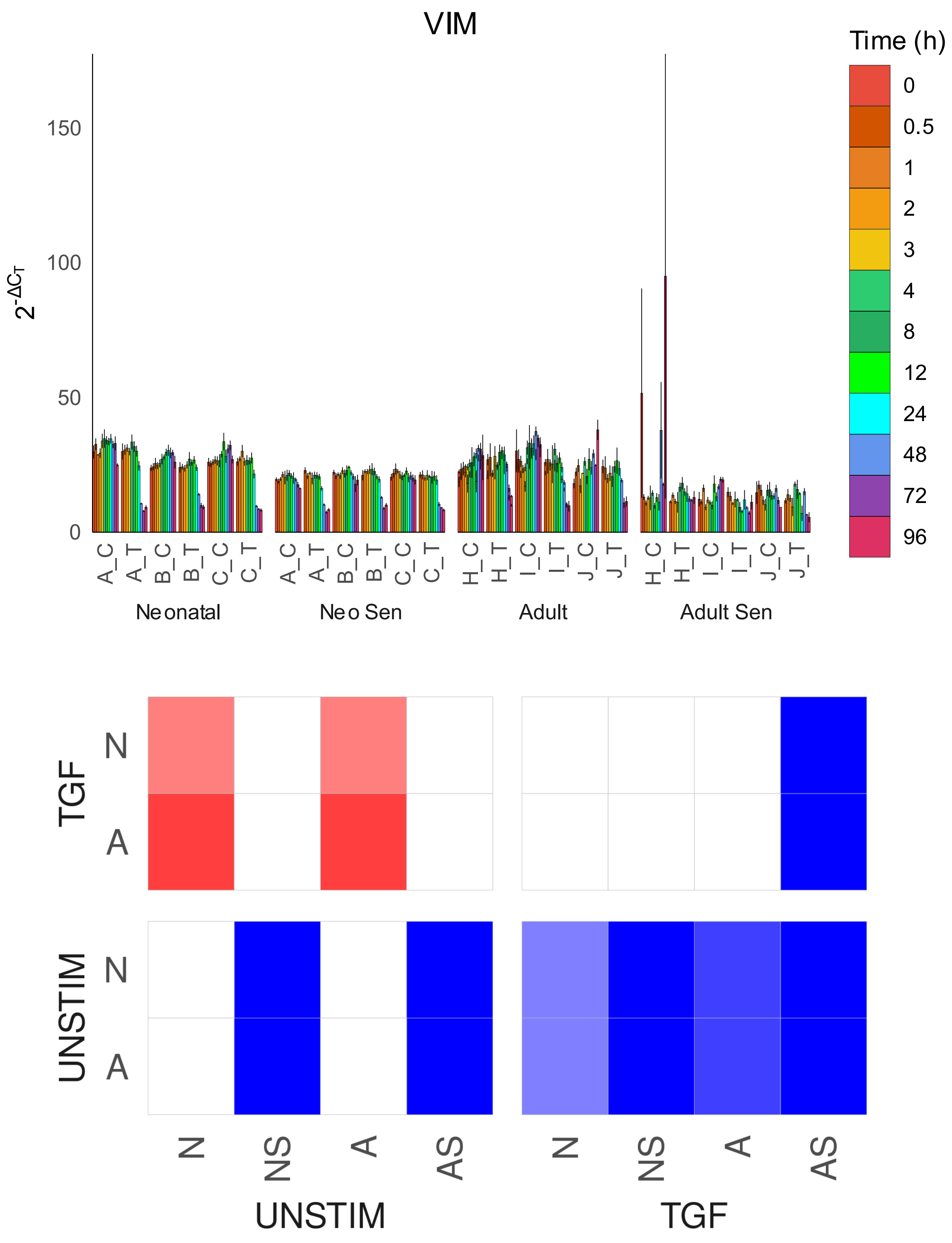
VIM
Time (h)
0
0.5
1
2
3
4
8
12
24
48
72
96
$2^{-\Delta C_T}$
150
100
50
0
A_C
A_T
B_C
B_T
C_C
C_T
A_C
A_T
B_C
B_T
C_C
C_T
H_C
H_T
I_C
I_T
J_C
J_T
H_C
H_T
I_C
I_T
J_C
J_T
Neonatal
Neo Sen
Adult
Adult Sen
TGF
N
A
UNSTIM
N
A
N
NS
A
AS
UNSTIM
N
NS
A
AS
TGF
