## Supplementary file 5 for "Cell Senescence-Independent Ageing of Human Skin"

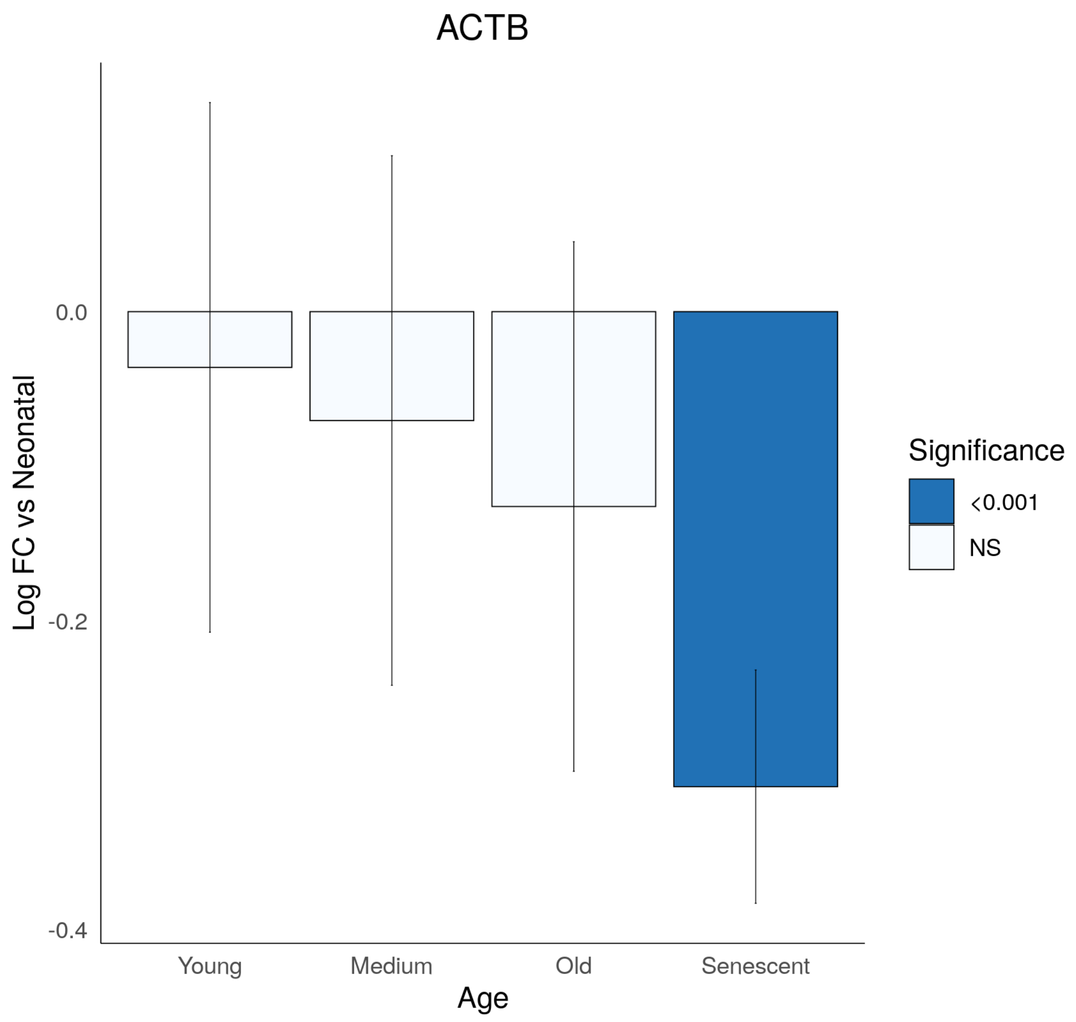
ACTB
Log FC vs Neonatal
0.0
-0.2
-0.4
Young
Medium
Old
Senescent
Age
Significance
<0.001
NS

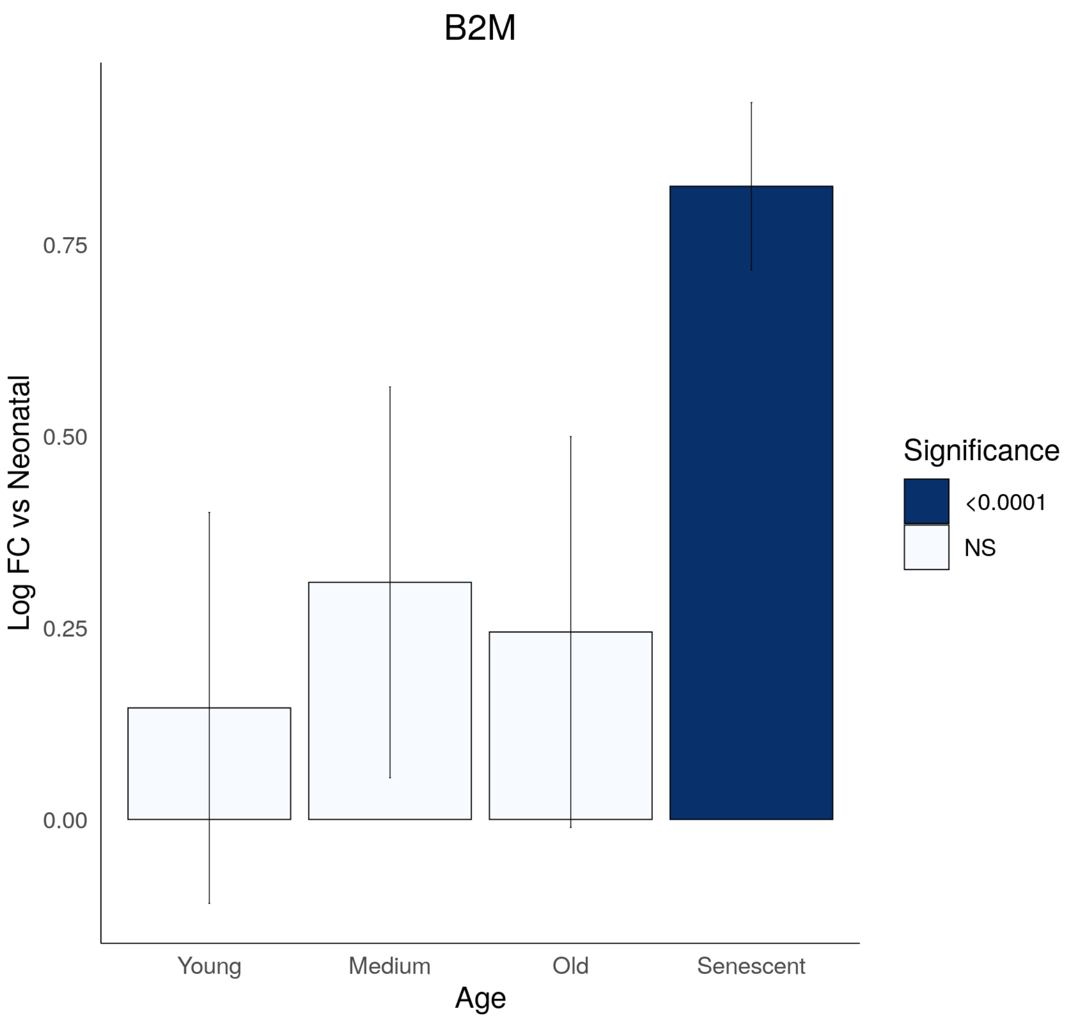
B2M
Log FC vs Neonatal
0.75
0.50
0.25
0.00
Young
Medium
Old
Senescent
Age
Significance
<0.0001
NS

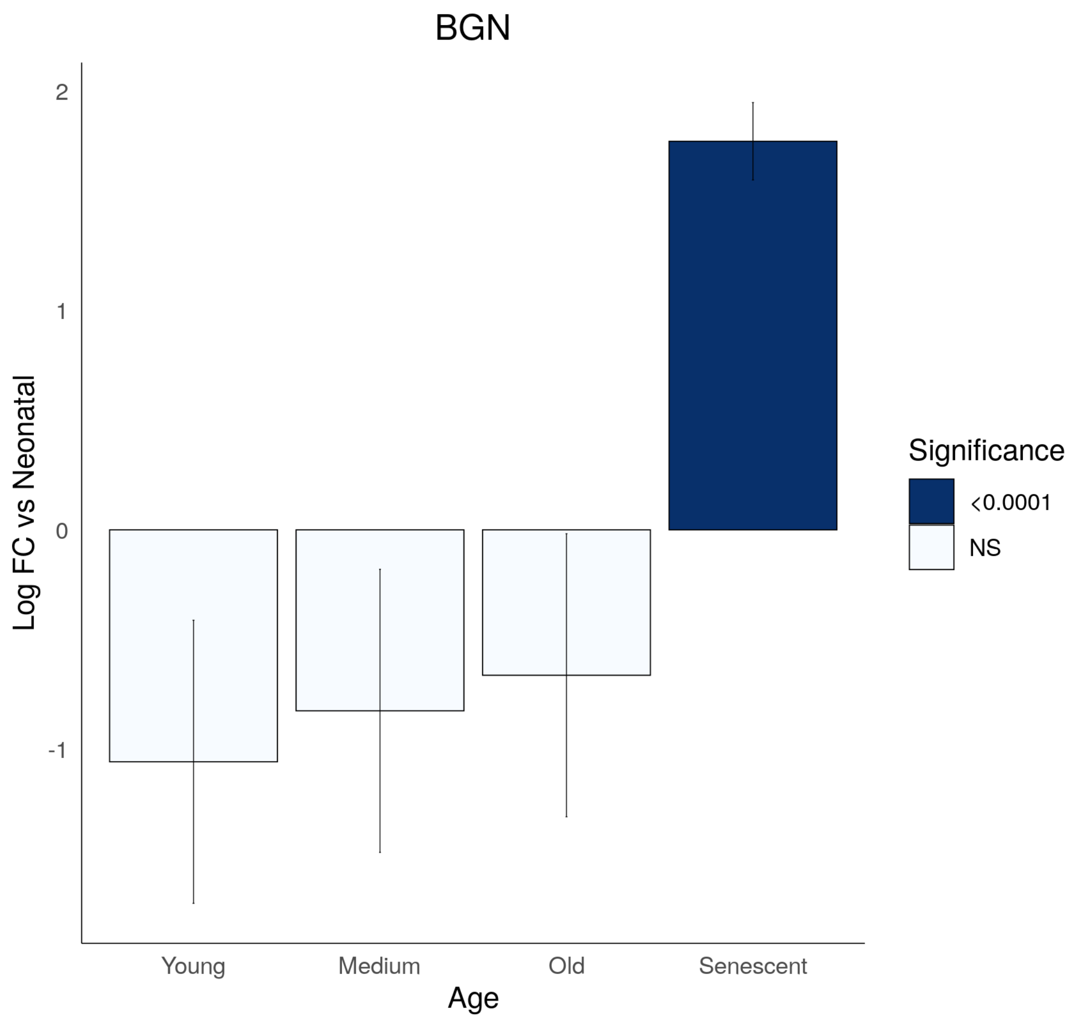

BGN
2
1
0
-1
Log FC vs Neonatal
Young
Medium
Old
Senescent
Age
Significance
<0.0001
NS

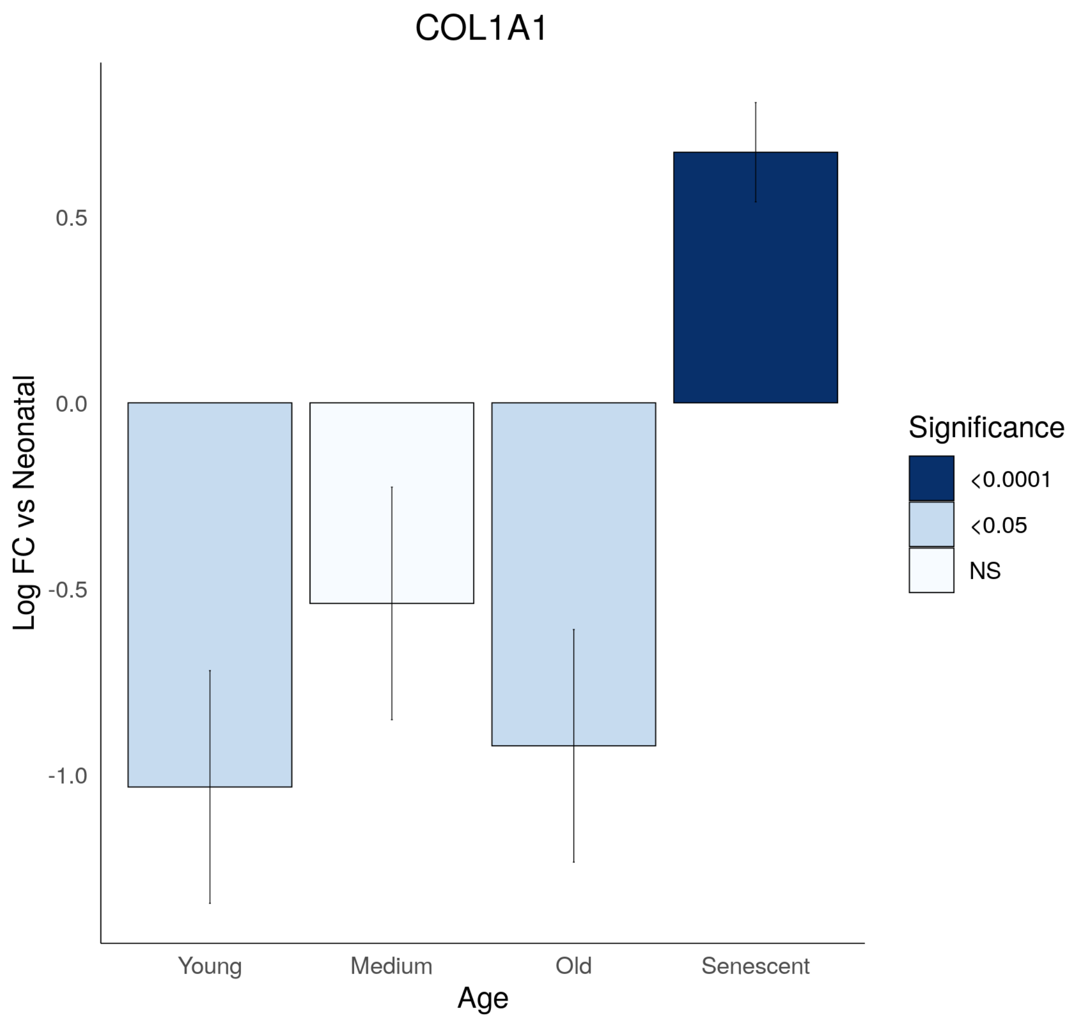
COL1A1
Log FC vs Neonatal
0.5
0.0
-0.5
-1.0
Young
Medium
Old
Senescent
Age
Significance
<0.0001
<0.05
NS

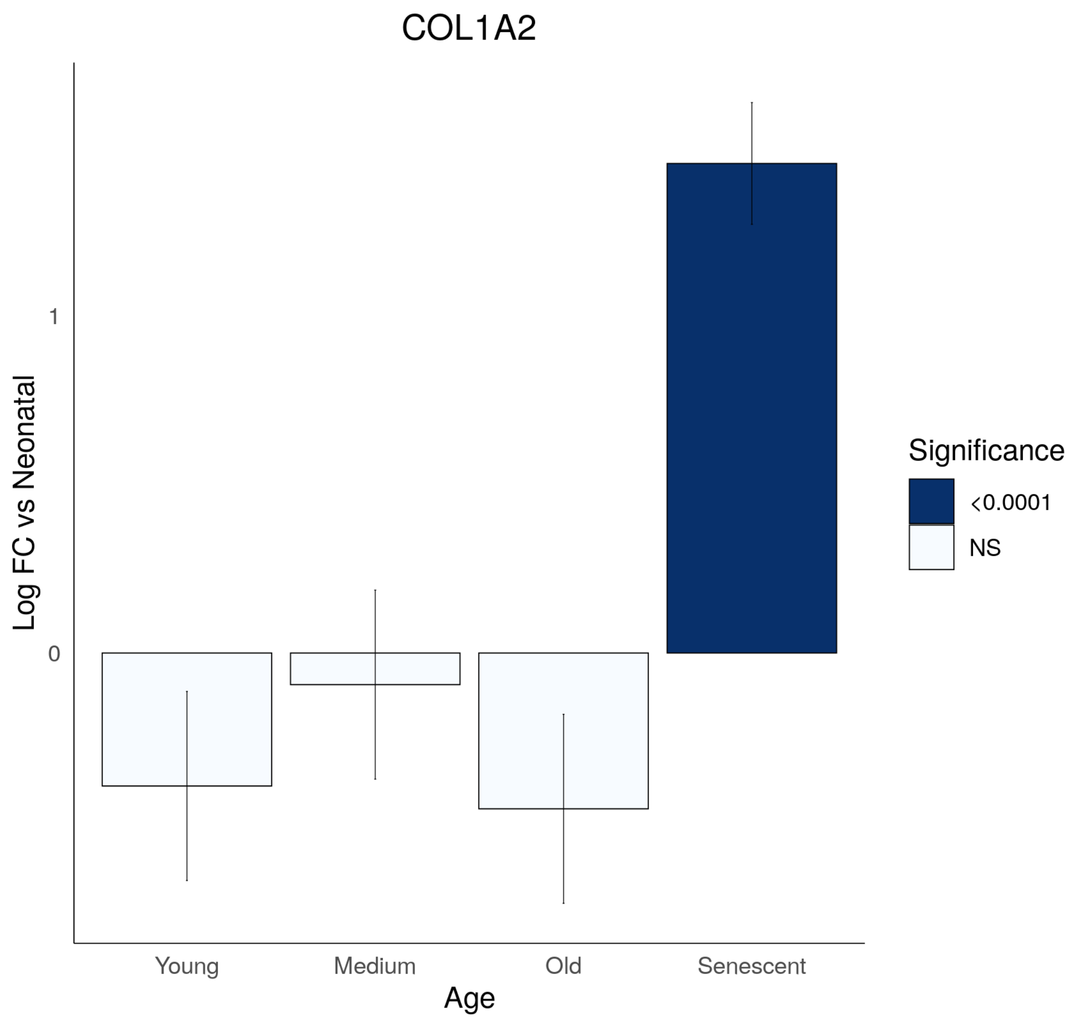
COL1A2
Log FC vs Neonatal
1
0
Young
Medium
Old
Senescent
Age
Significance
<0.0001
NS

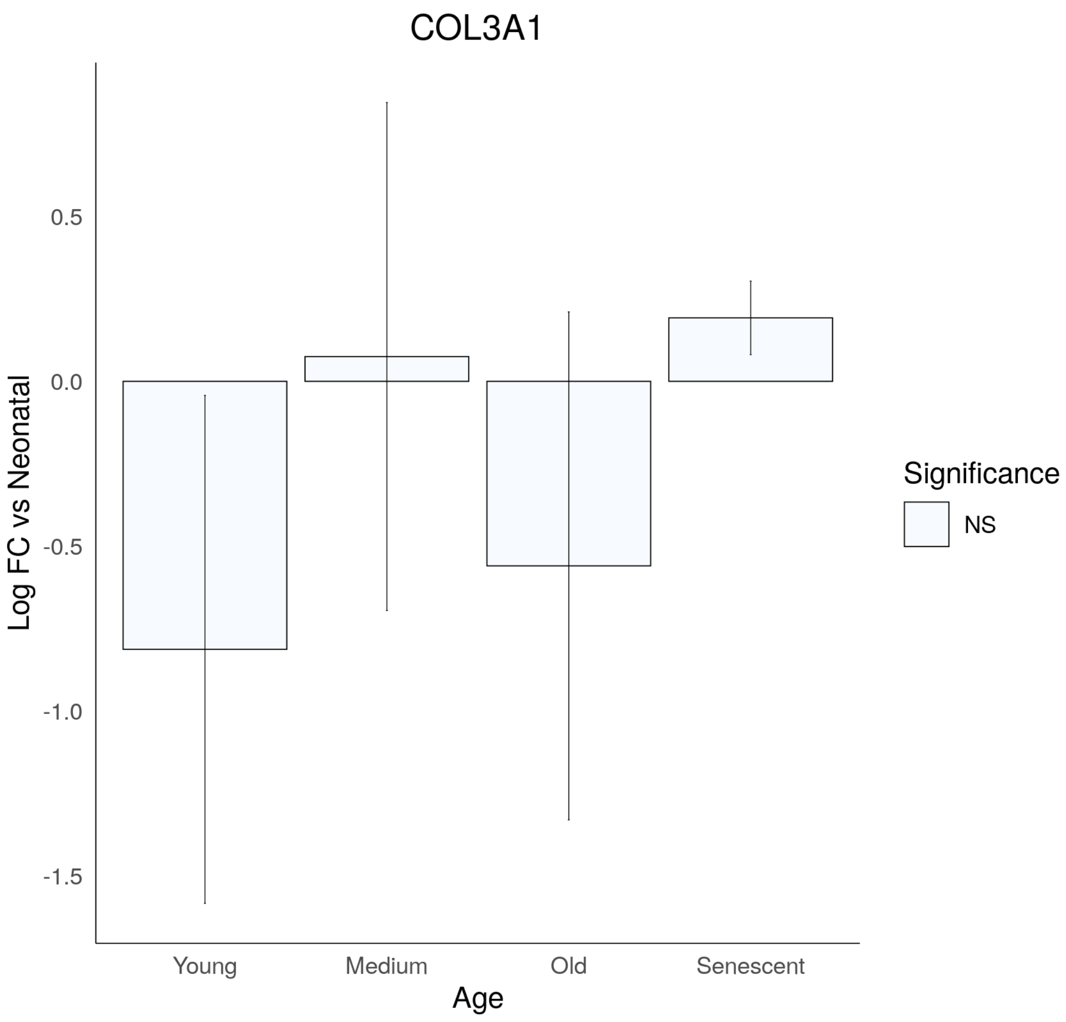
COL3A1
Log FC vs Neonatal
0.5
0.0
-0.5
-1.0
-1.5
Young
Medium
Old
Senescent
Age
Significance
NS

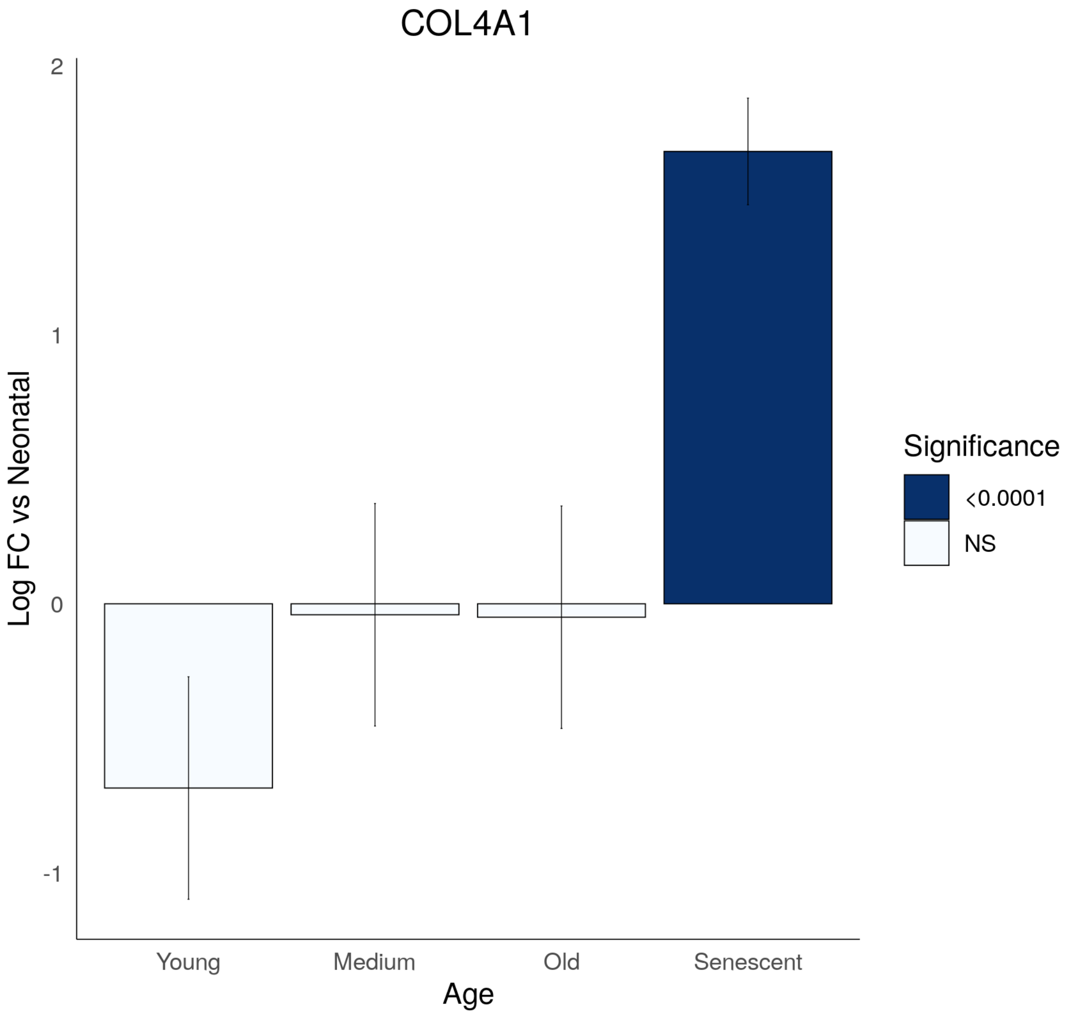

COL4A1
2
1
0
-1
Log FC vs Neonatal
Young
Medium
Old
Senescent
Age
Significance
<0.0001
NS

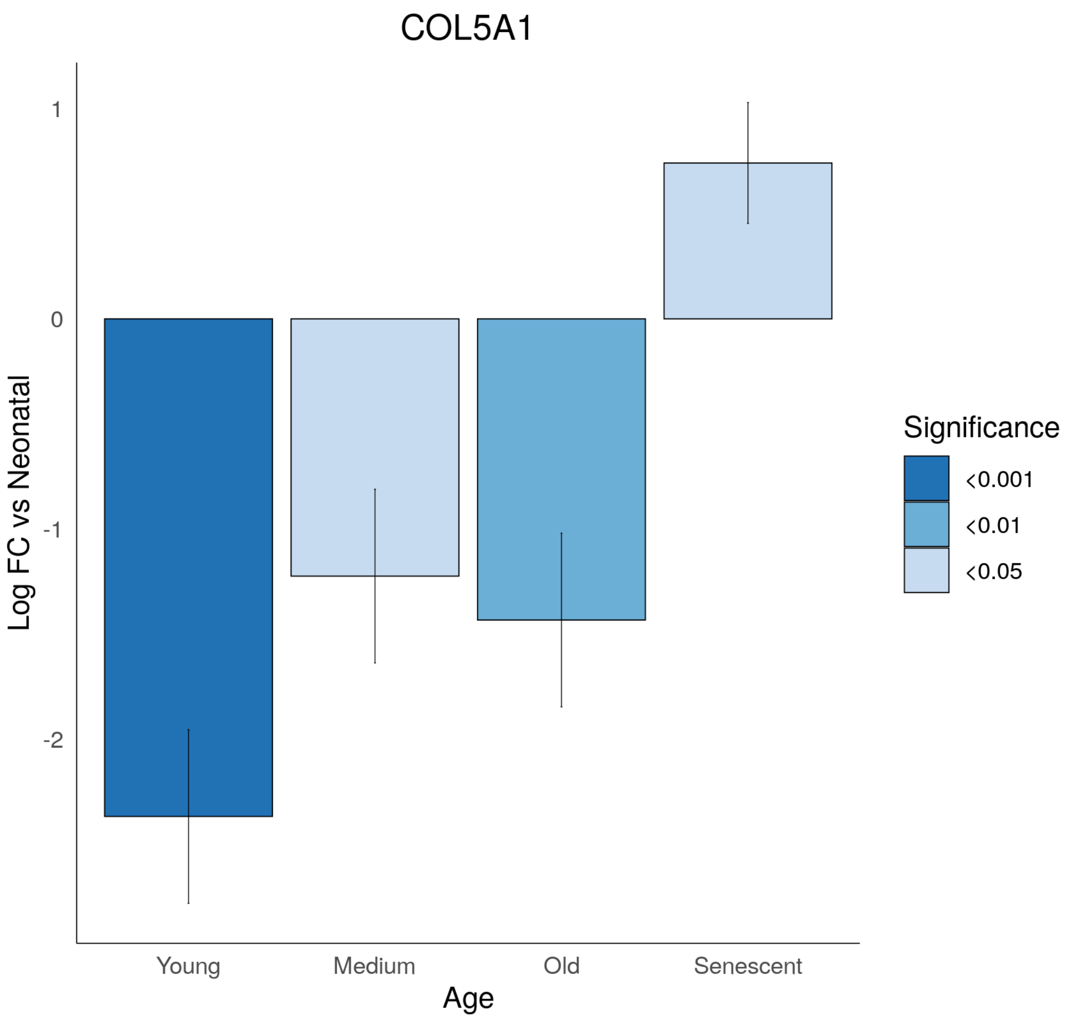
COL5A1
1
0
-1
-2
Log FC vs Neonatal
Young
Medium
Old
Senescent
Age
Significance
<0.001
<0.01
<0.05

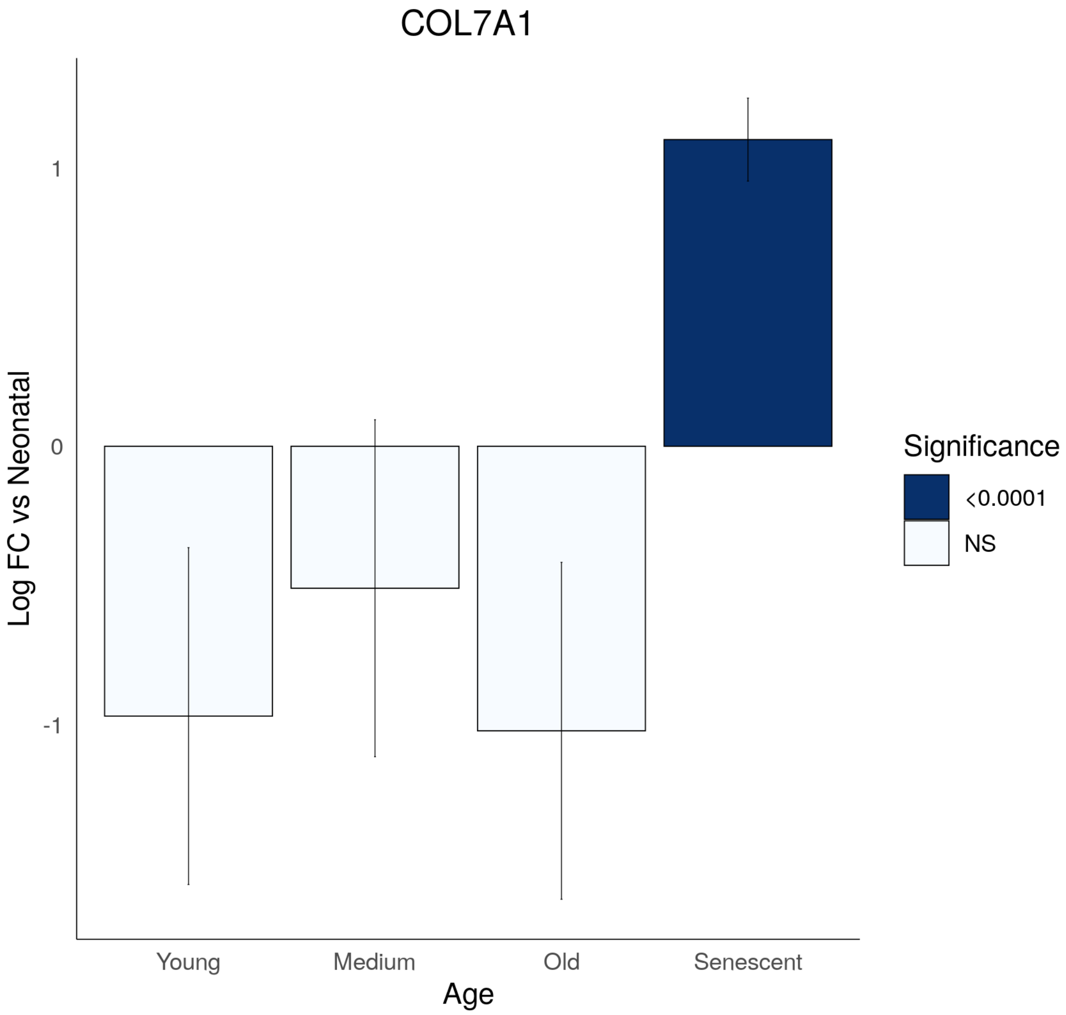
COL7A1
Log FC vs Neonatal
1
0
-1
Young
Medium
Old
Senescent
Age
Significance
<0.0001
NS

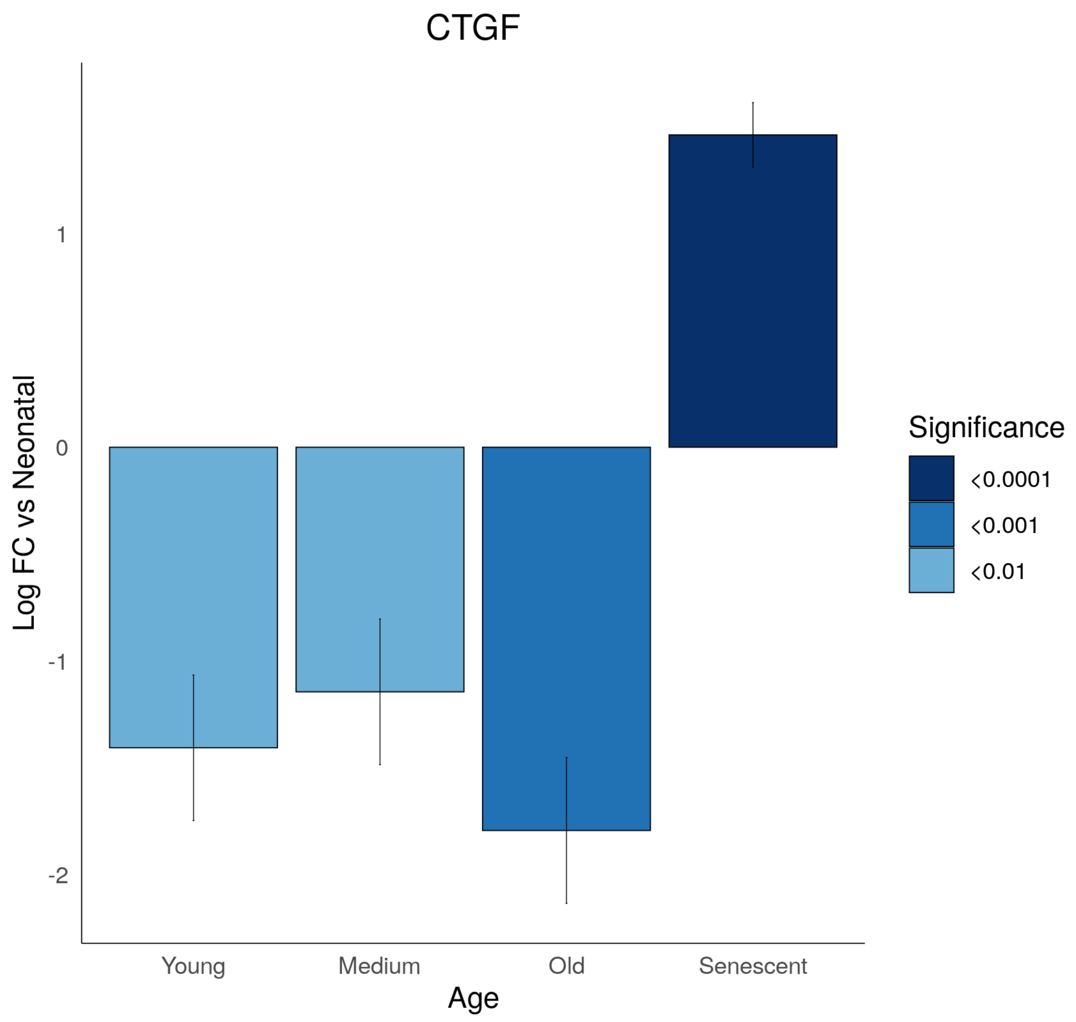

CTGF
1
0
-1
-2
Log FC vs Neonatal
Young
Medium
Old
Senescent
Age
Significance
<0.0001
<0.001
<0.01

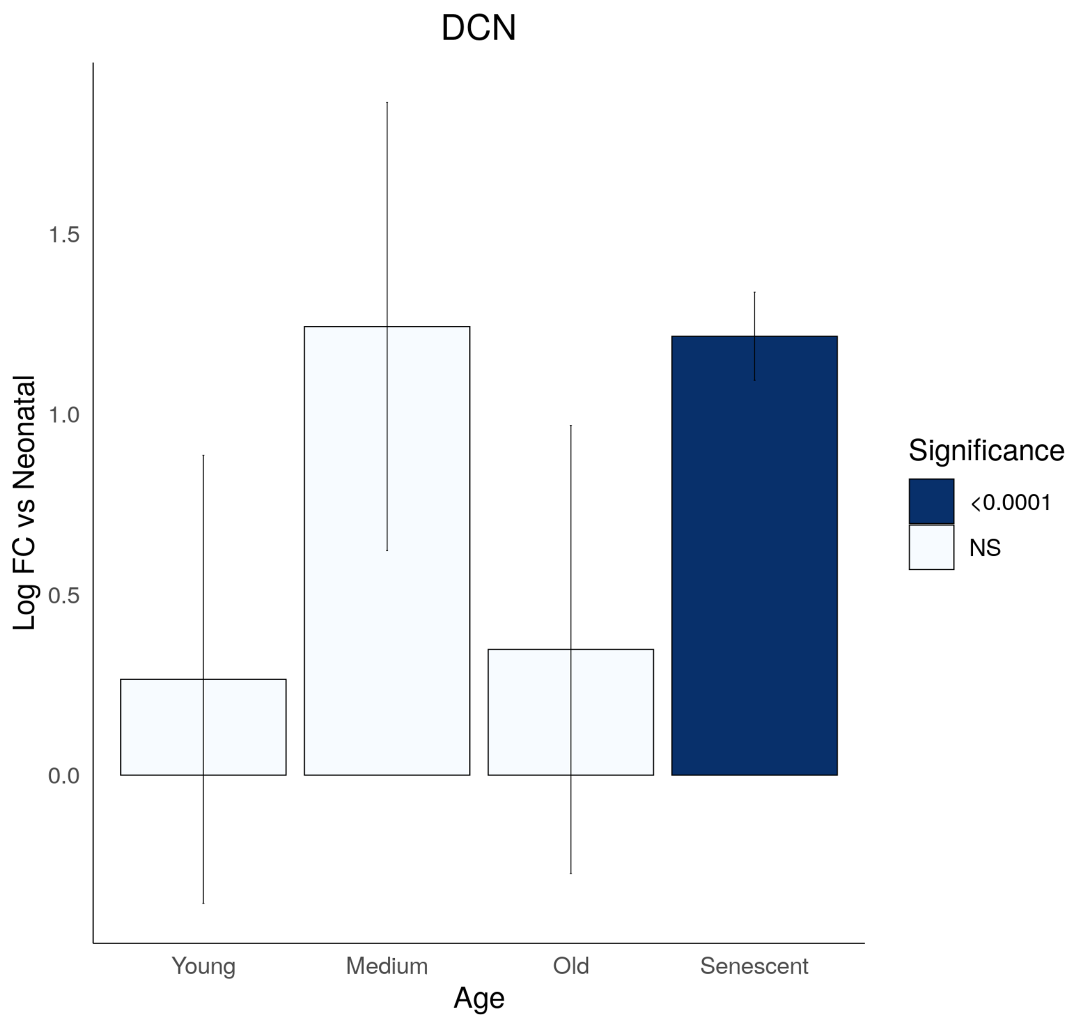

DCN
Log FC vs Neonatal
1.5
1.0
0.5
0.0
Young
Medium
Old
Senescent
Age
Significance
<0.0001
NS

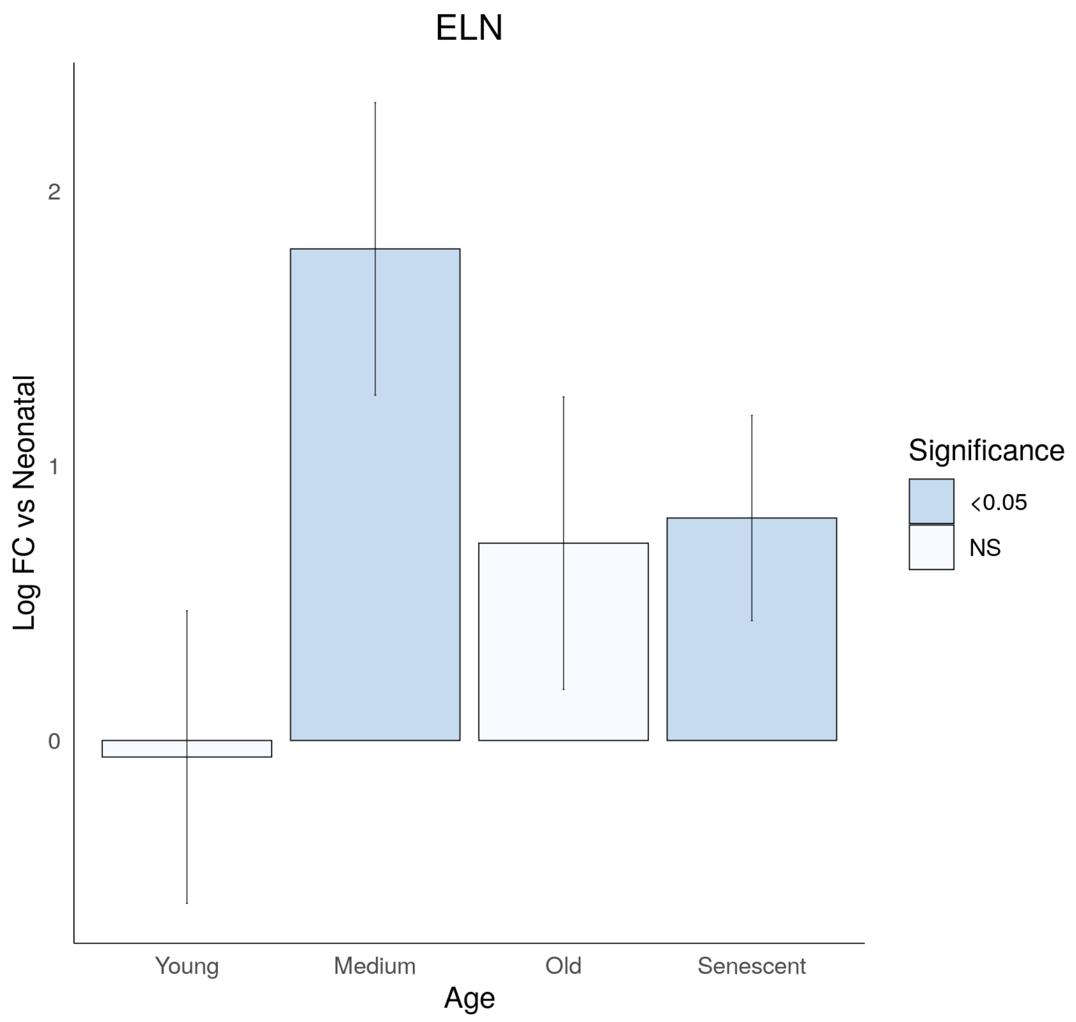
ELN
Log FC vs Neonatal
2
1
0
Young
Medium
Old
Senescent
Age
Significance
<0.05
NS

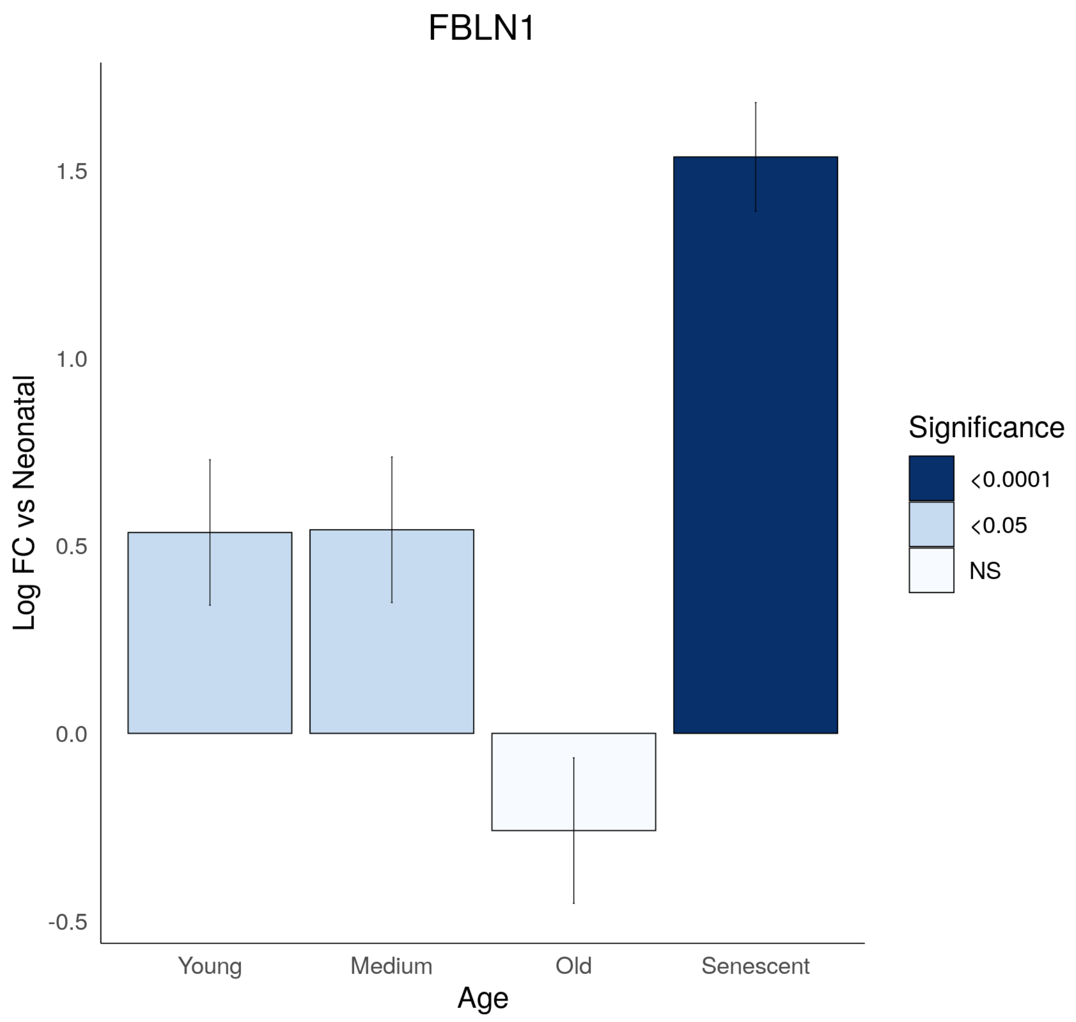

FBLN1
Log FC vs Neonatal
1.5
1.0
0.5
0.0
-0.5
Young
Medium
Old
Senescent
Age
Significance
<0.0001
<0.05
NS

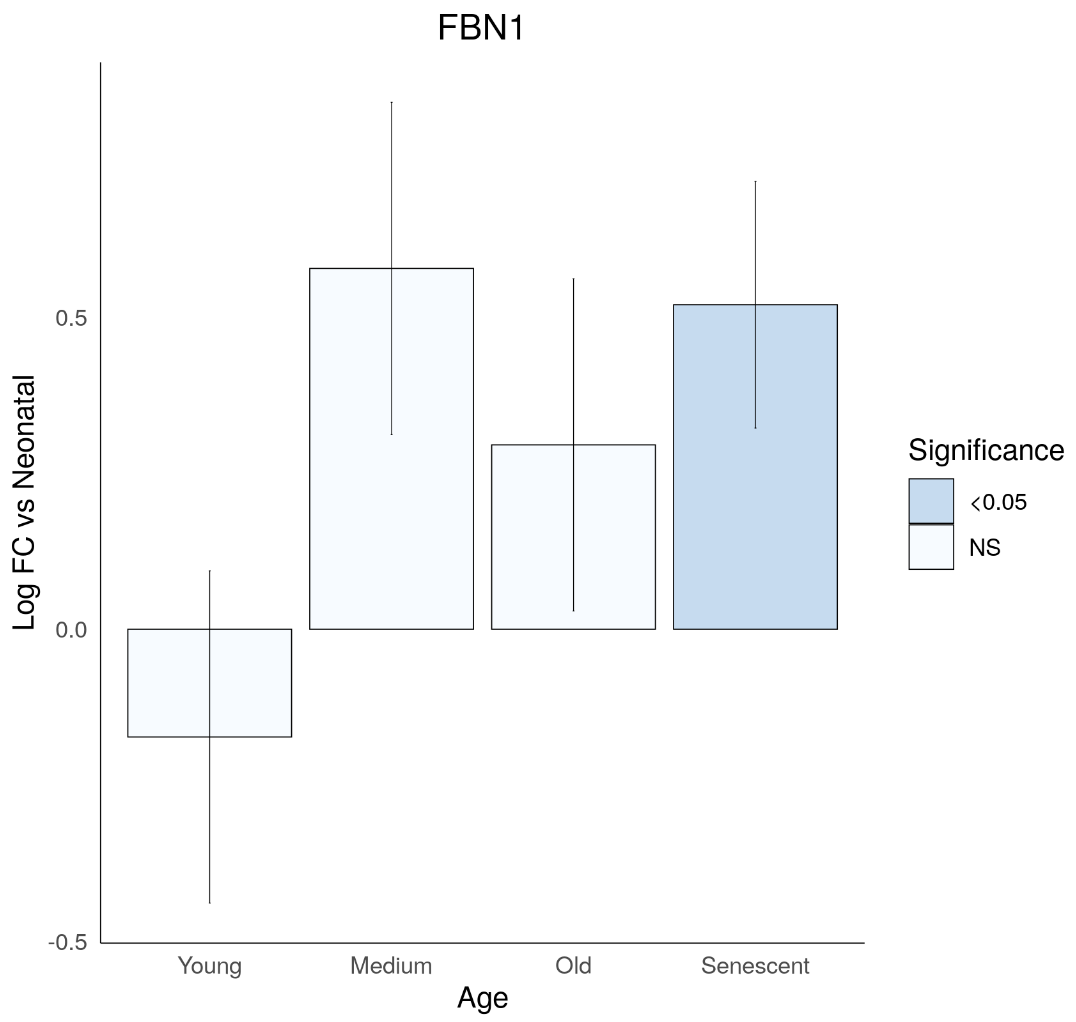

FBN1
Log FC vs Neonatal
0.5
0.0
-0.5
Young
Medium
Old
Senescent
Age
Significance
<0.05
NS

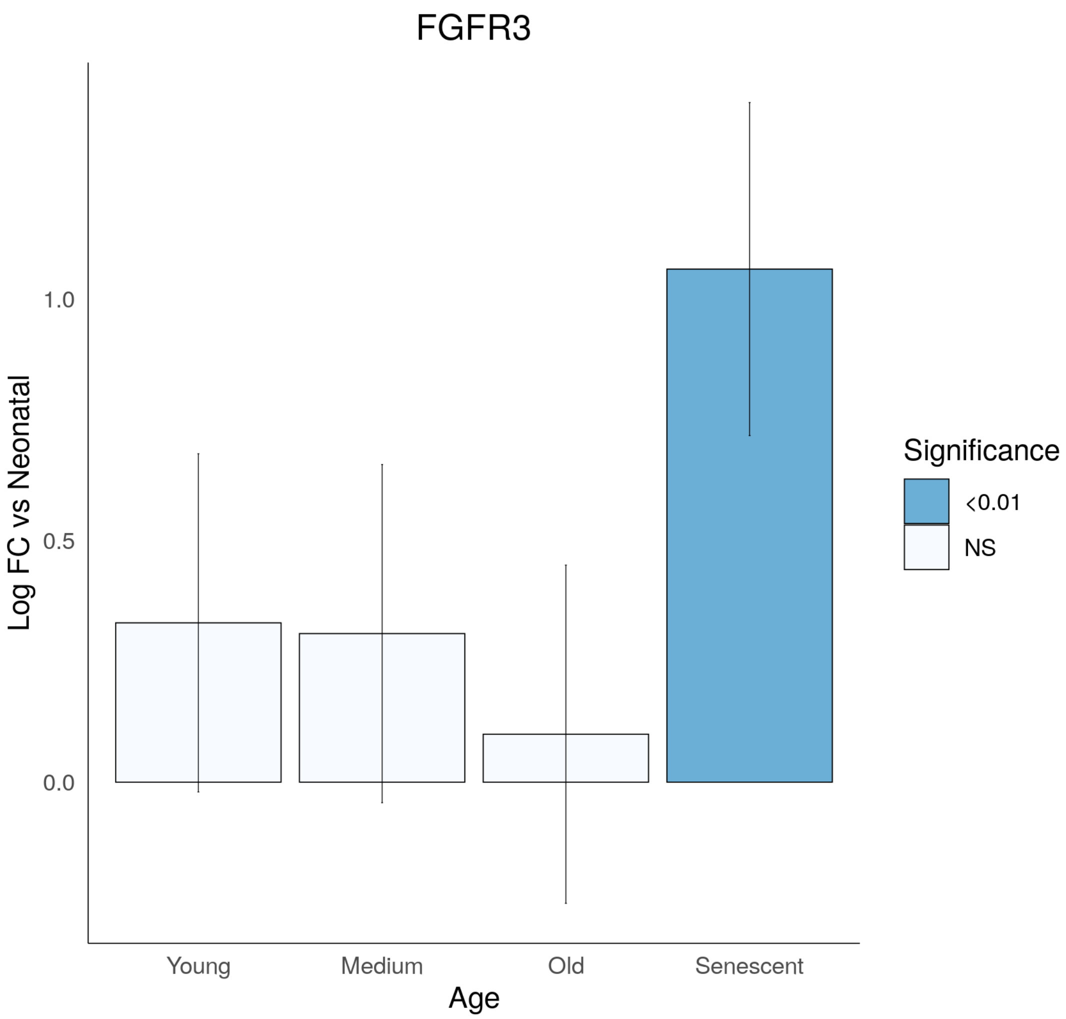

FGFR3
Log FC vs Neonatal
1.0
0.5
0.0
Young
Medium
Old
Senescent
Age
Significance
<0.01
NS

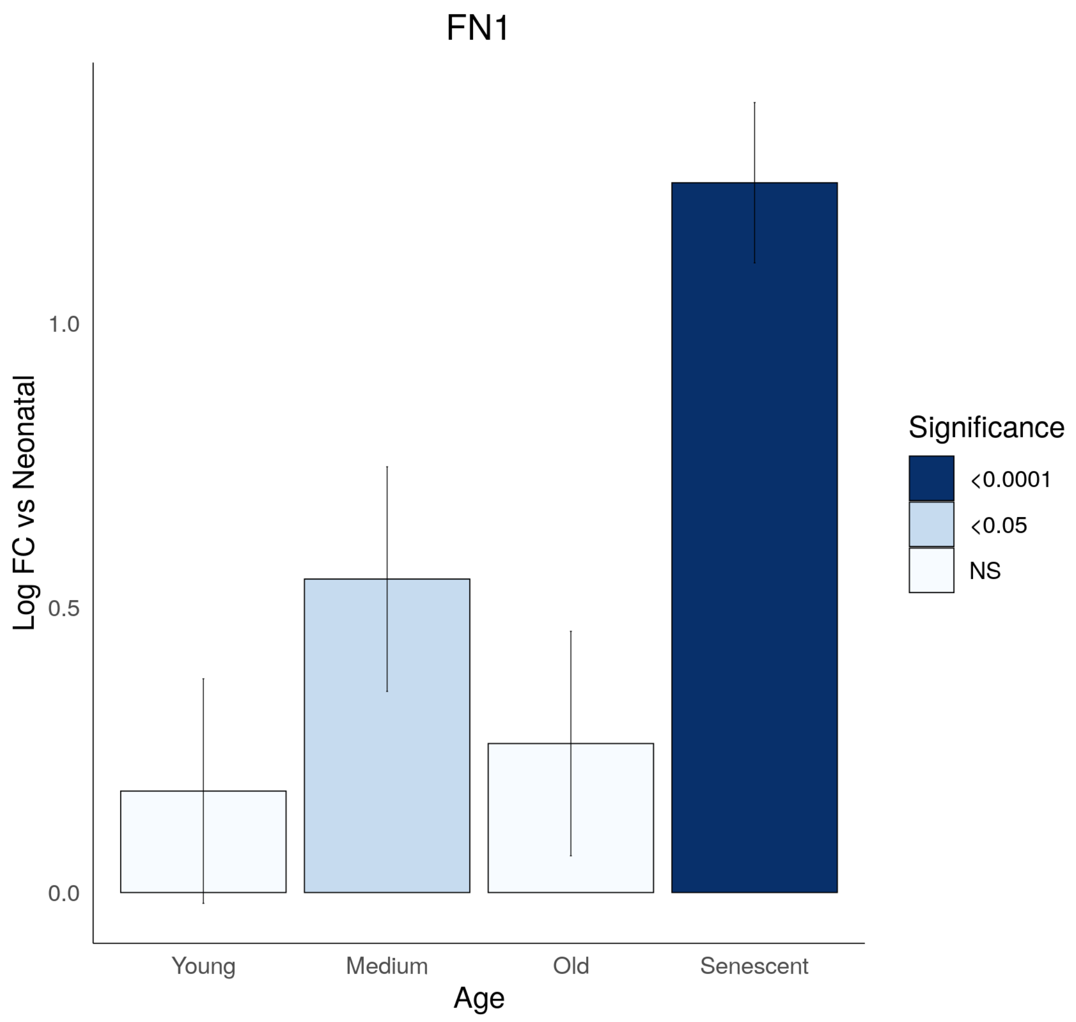

FN1
Log FC vs Neonatal
1.0
0.5
0.0
Young
Medium
Old
Senescent
Age
Significance
<0.0001
<0.05
NS

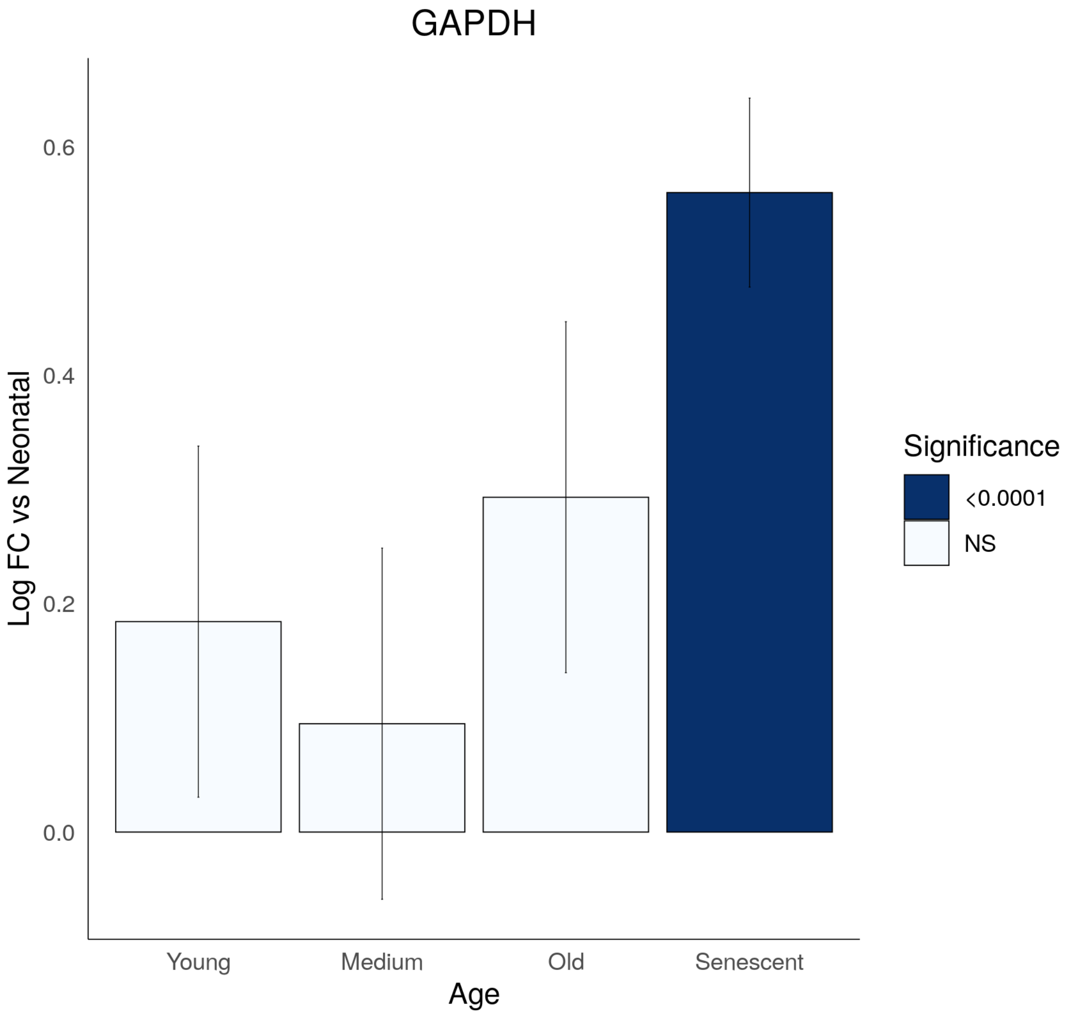

GAPDH
Log FC vs Neonatal
0.6
0.4
0.2
0.0
Young
Medium
Old
Senescent
Age
Significance
<0.0001
NS

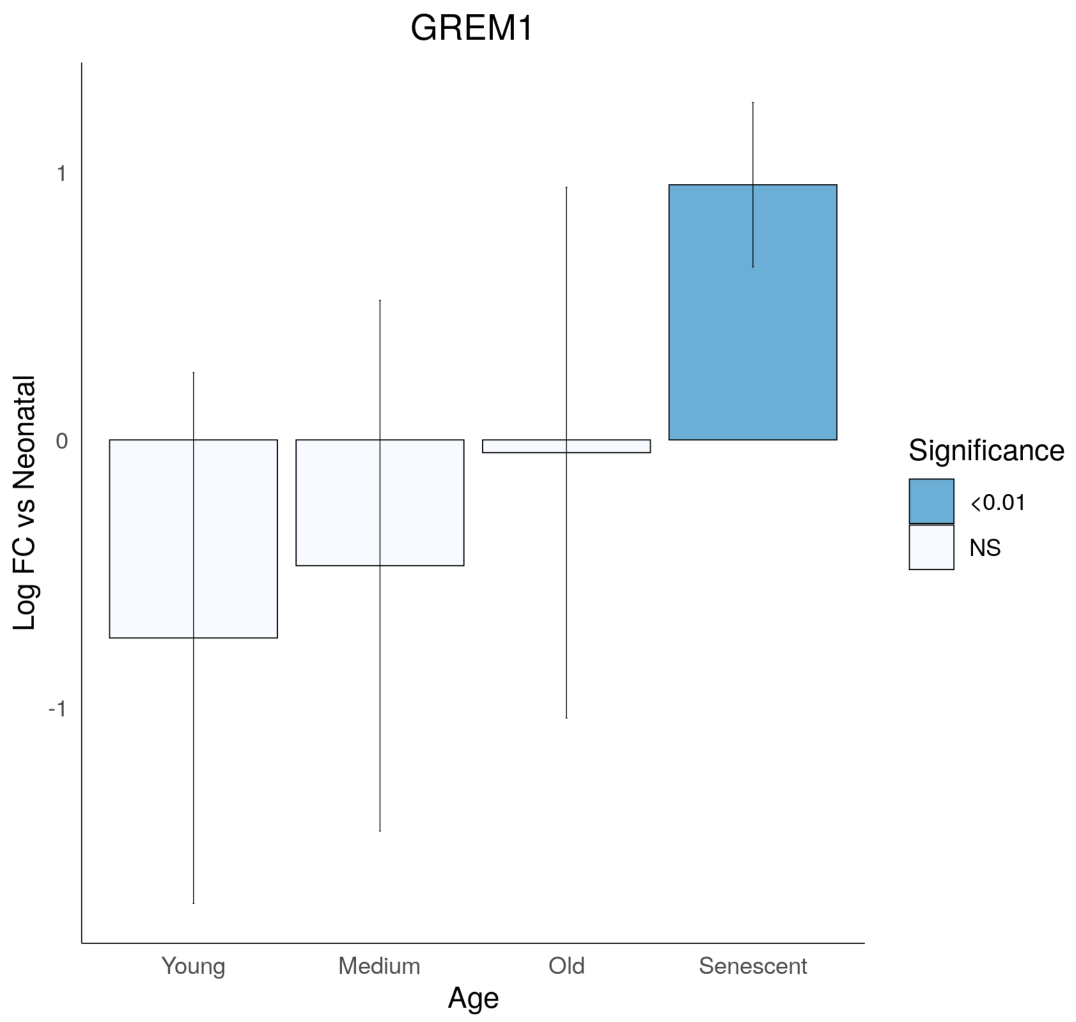
GREM1
Log FC vs Neonatal
1
0
-1
Young
Medium
Old
Senescent
Age
Significance
<0.01
NS

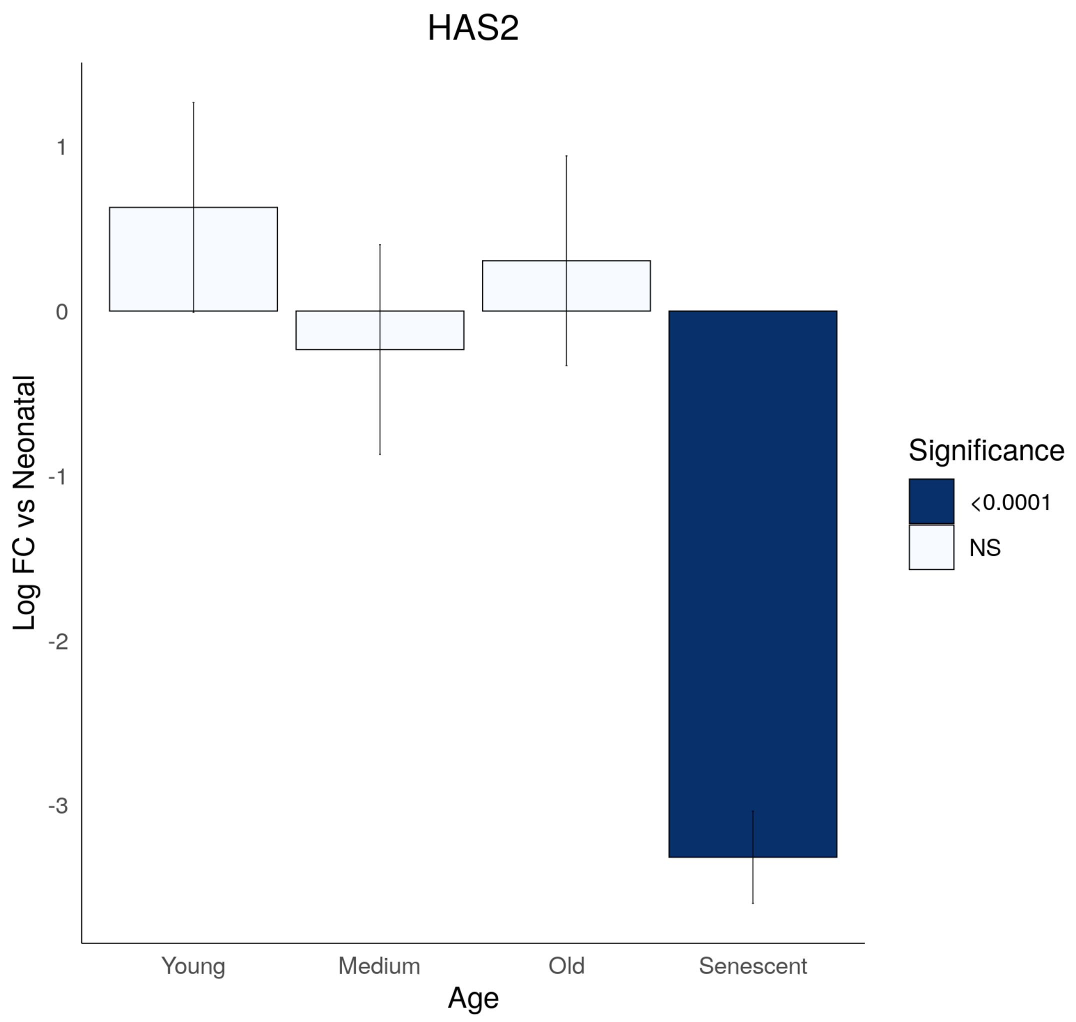
HAS2
1
0
-1
-2
-3
Log FC vs Neonatal
Young
Medium
Old
Senescent
Age
Significance
<0.0001
NS

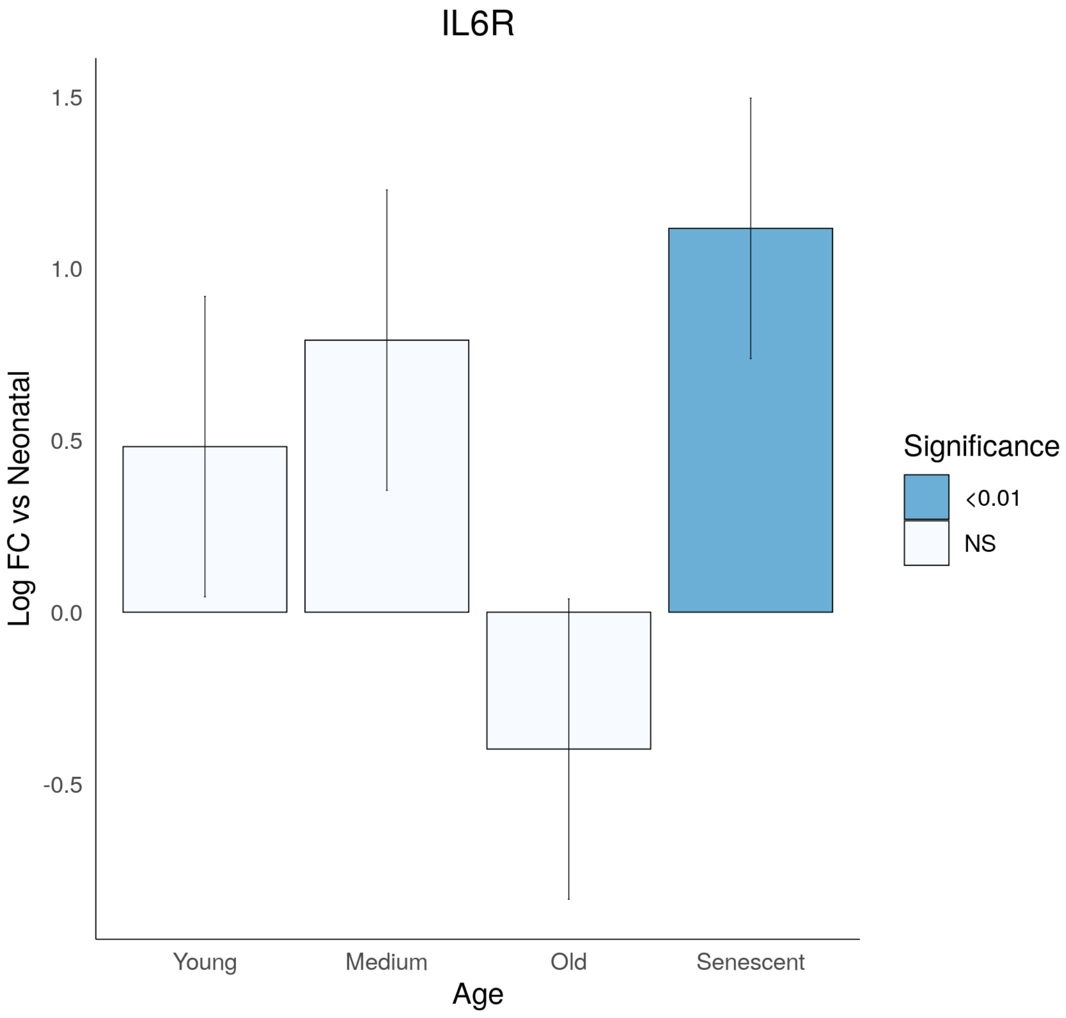

IL6R
Log FC vs Neonatal
1.5
1.0
0.5
0.0
-0.5
Young
Medium
Old
Senescent
Age
Significance
<0.01
NS

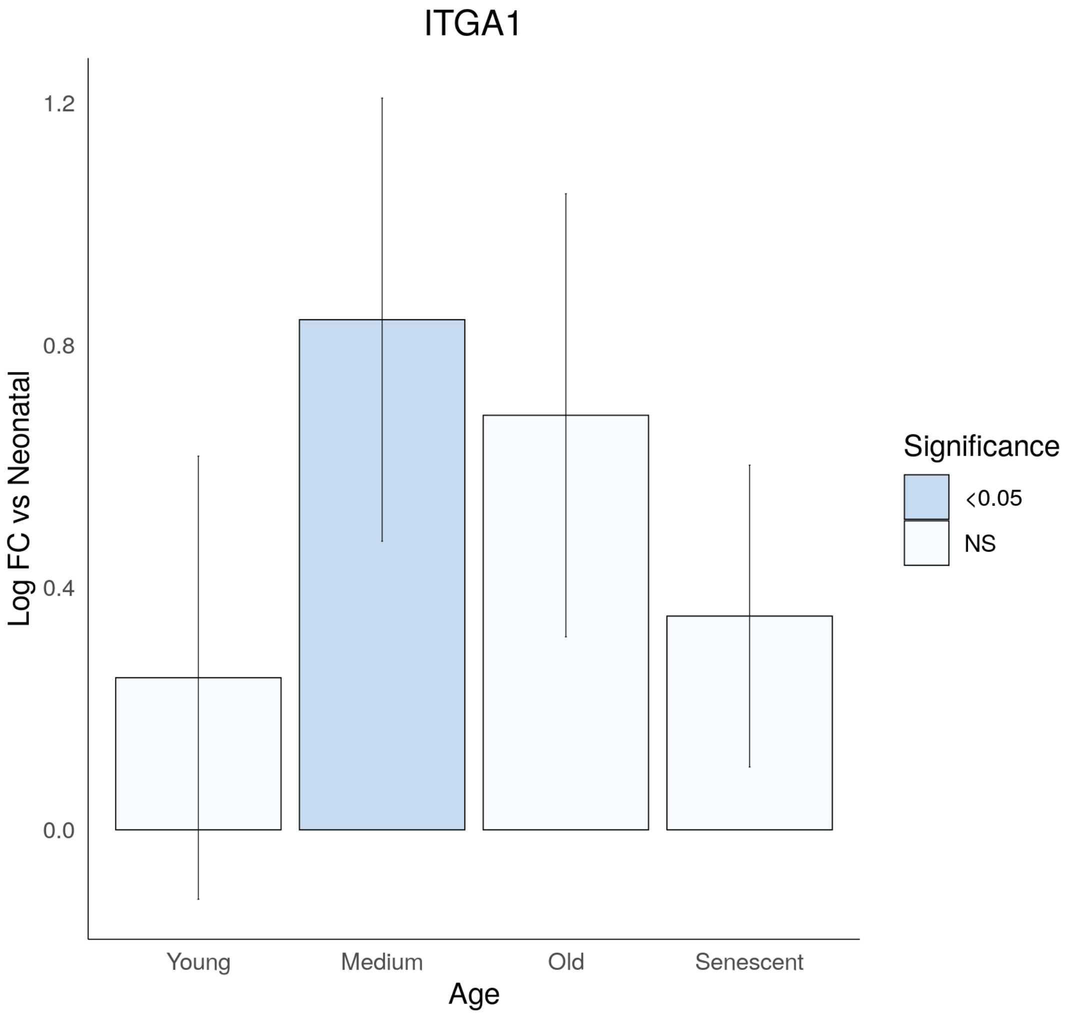
ITGA1
1.2
0.8
0.4
0.0
Log FC vs Neonatal
Young
Medium
Old
Senescent
Age
Significance
<0.05
NS

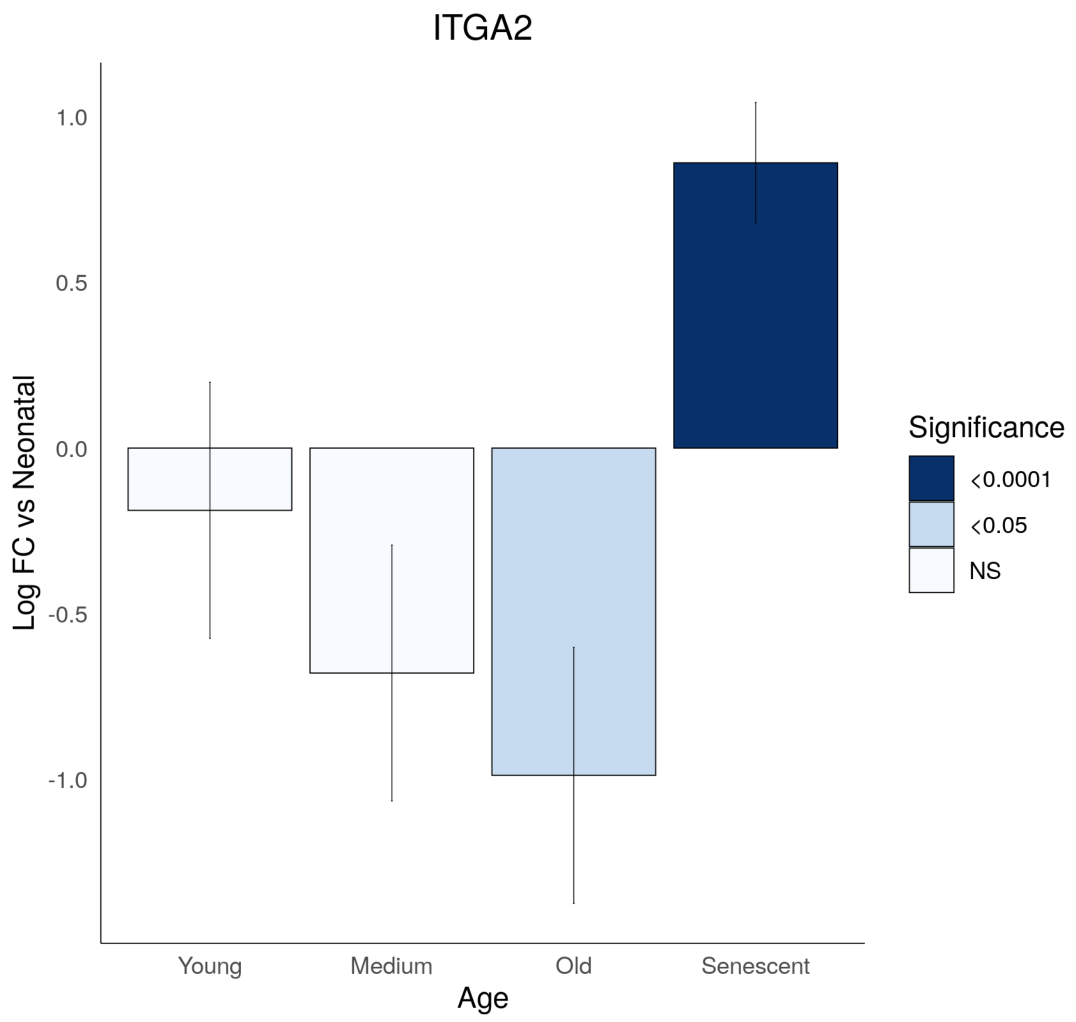

ITGA2
Log FC vs Neonatal
1.0
0.5
0.0
-0.5
-1.0
Young
Medium
Old
Senescent
Age
Significance
<0.0001
<0.05
NS

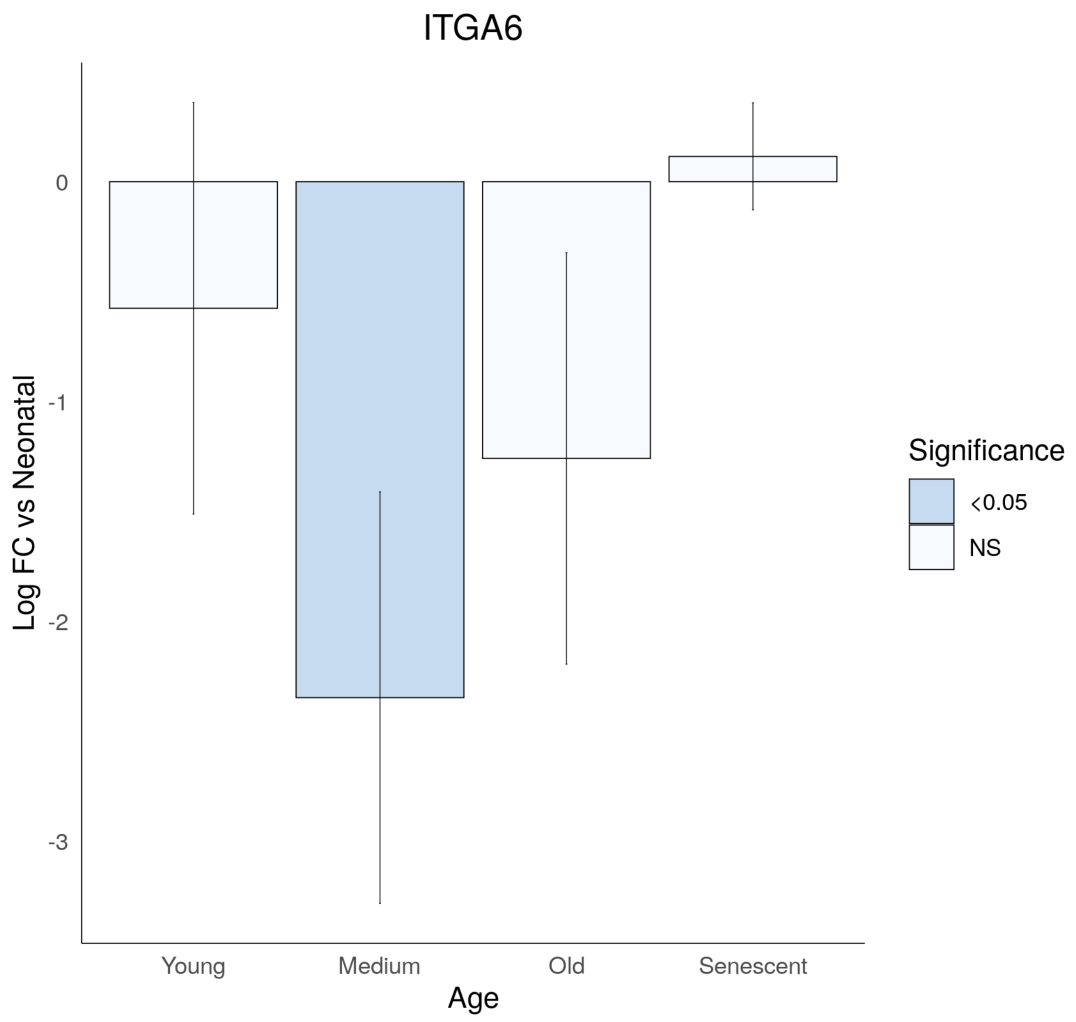
ITGA6
Log FC vs Neonatal
0
-1
-2
-3
Young
Medium
Old
Senescent
Age
Significance
<0.05
NS

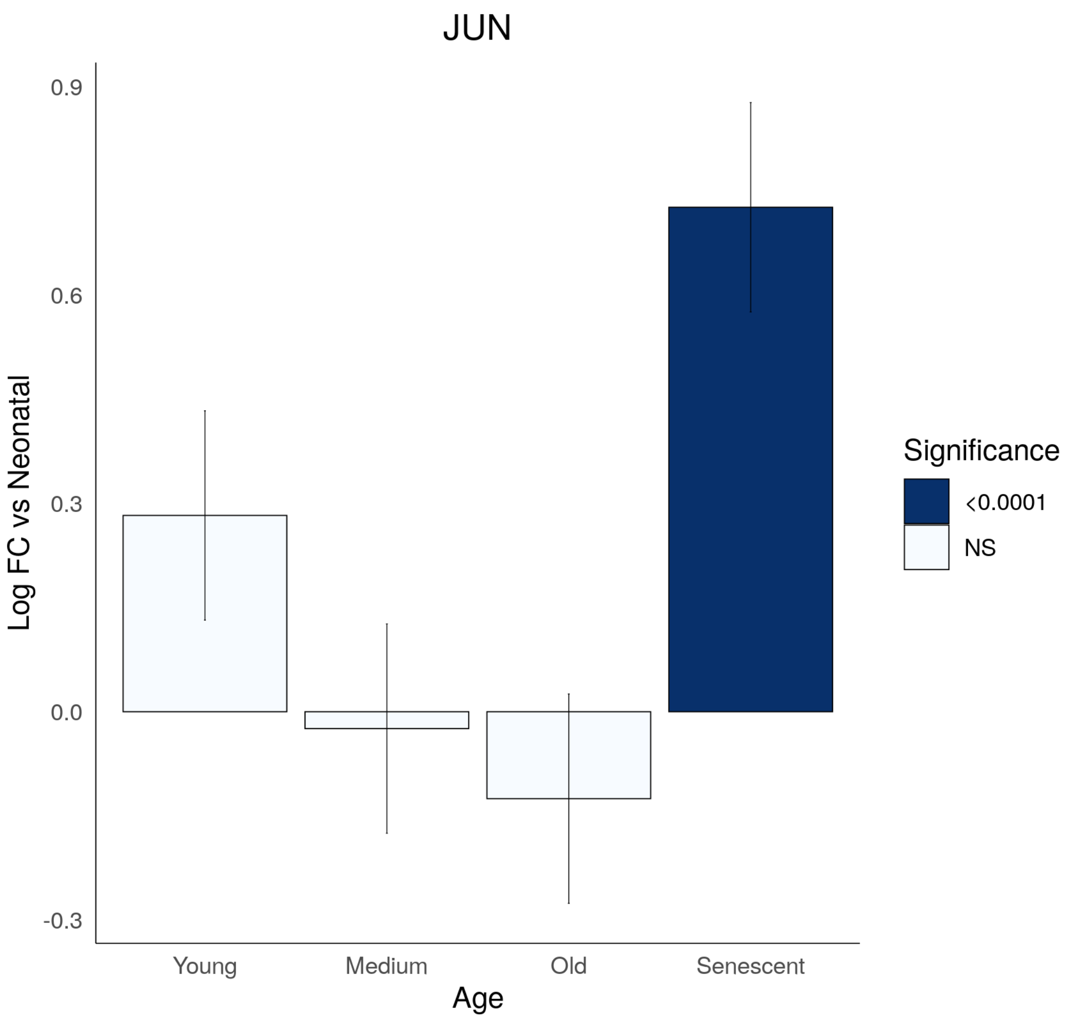
JUN
0.9
0.6
0.3
0.0
-0.3
Log FC vs Neonatal
Young
Medium
Old
Senescent
Age
Significance
<0.0001
NS

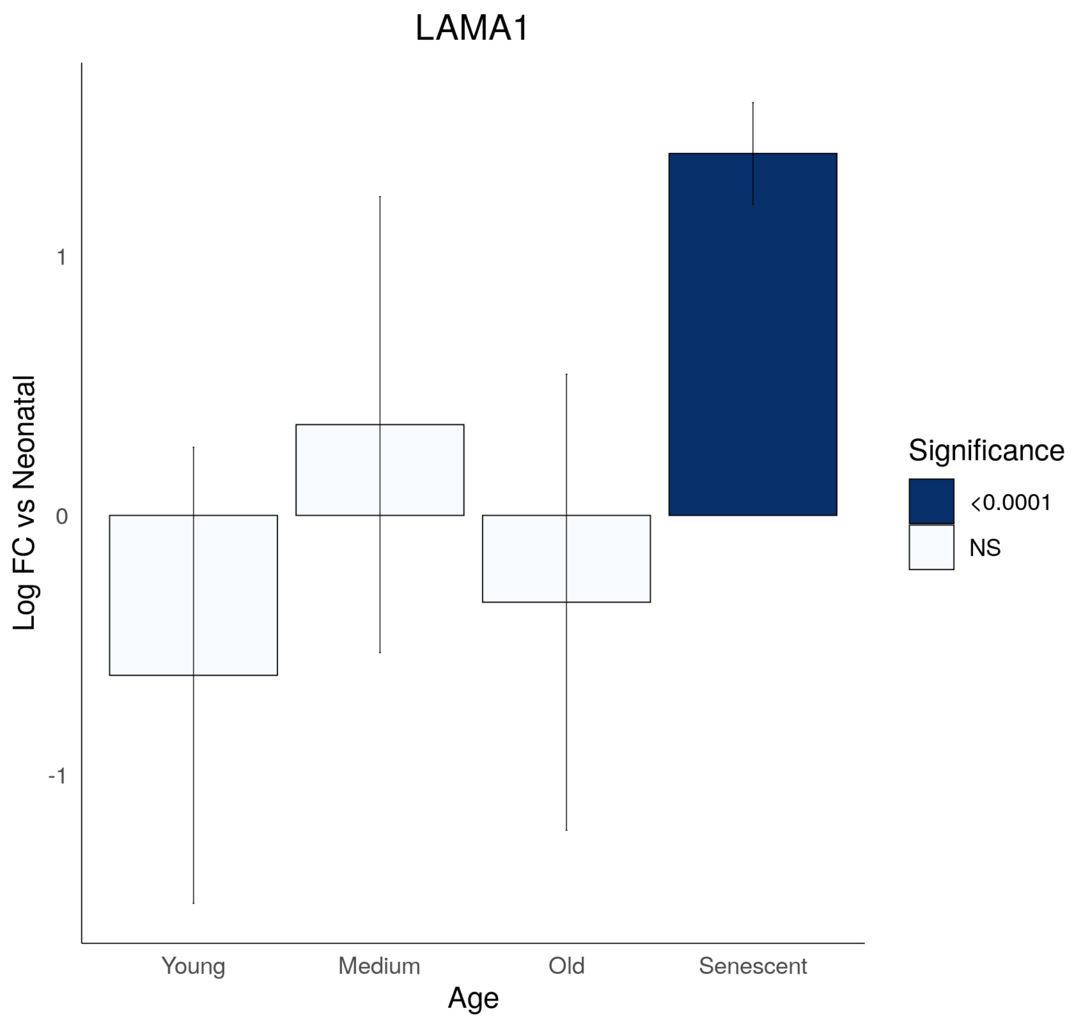
LAMA1
Log FC vs Neonatal
1
0
-1
Young
Medium
Old
Senescent
Age
Significance
<0.0001
NS

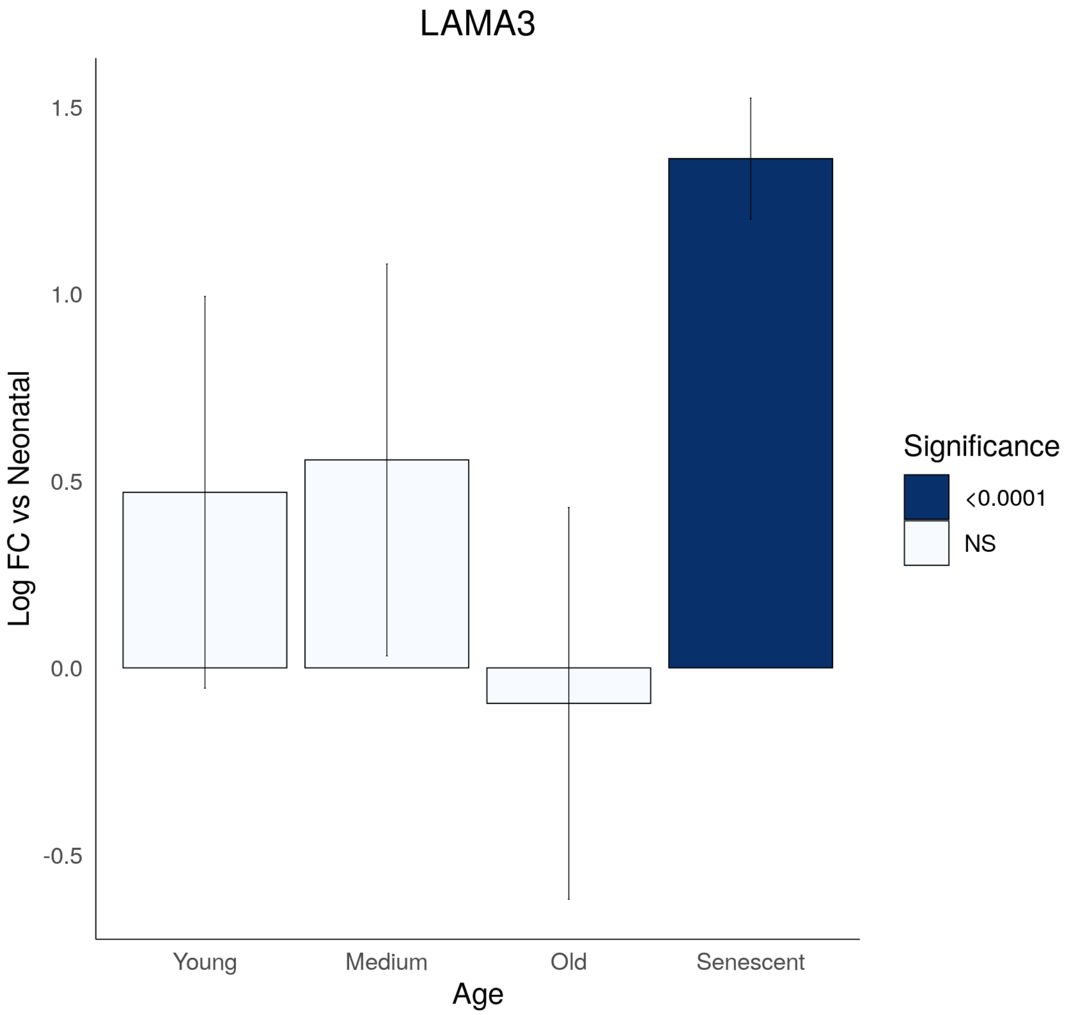

LAMA3
1.5
1.0
0.5
0.0
-0.5
Log FC vs Neonatal
Young
Medium
Old
Senescent
Age
Significance
<0.0001
NS

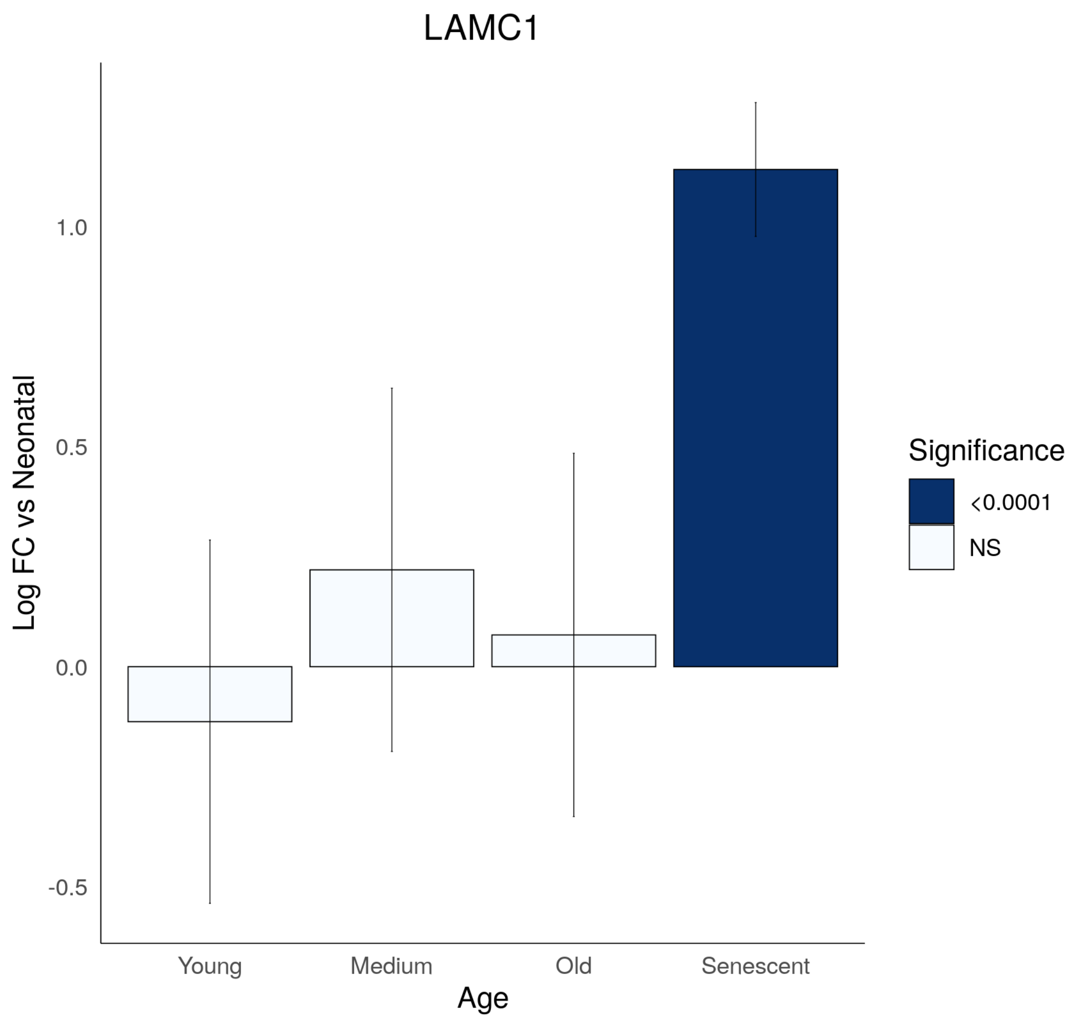

LAMC1
Log FC vs Neonatal
1.0
0.5
0.0
-0.5
Young
Medium
Old
Senescent
Age
Significance
<0.0001
NS

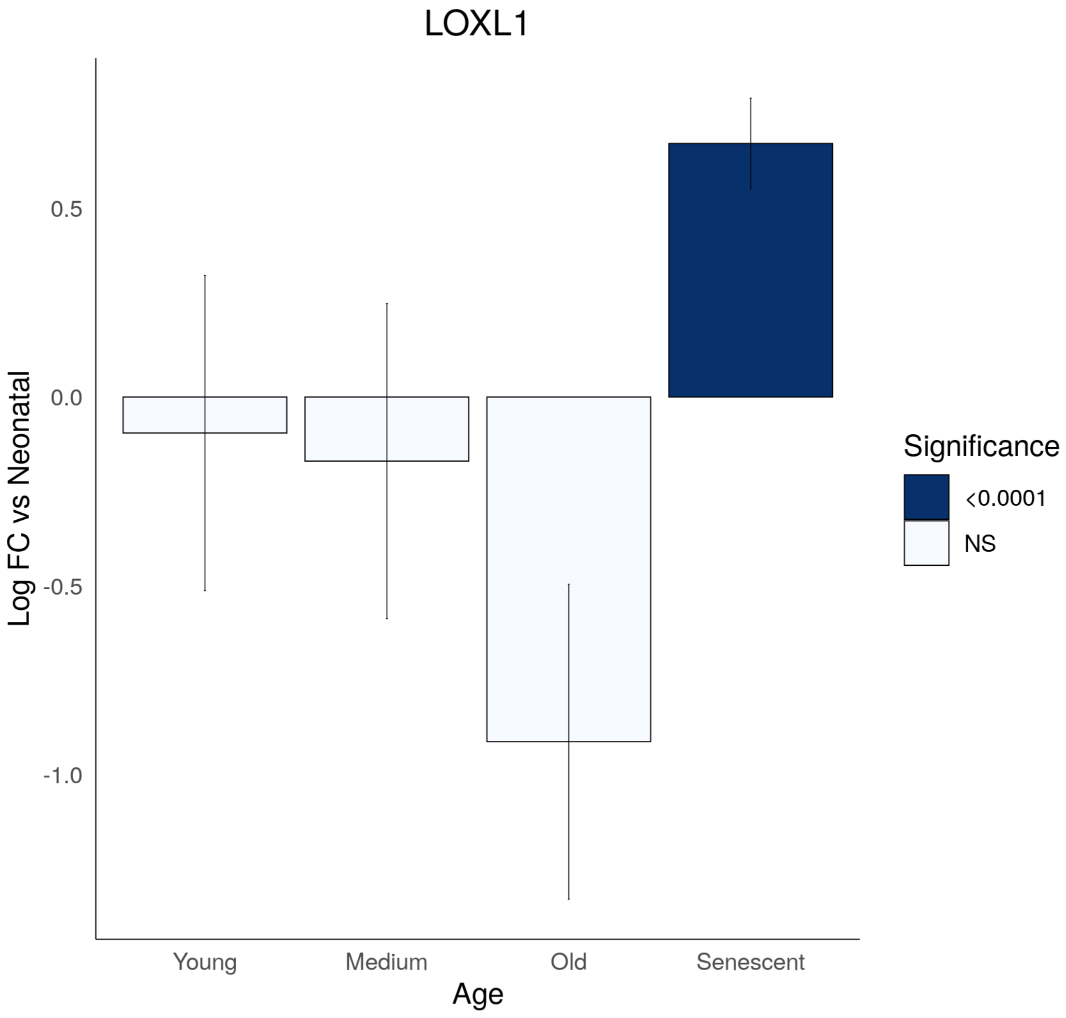

LOXL1
0.5
0.0
-0.5
-1.0
Log FC vs Neonatal
Young
Medium
Old
Senescent
Age
Significance
<0.0001
NS

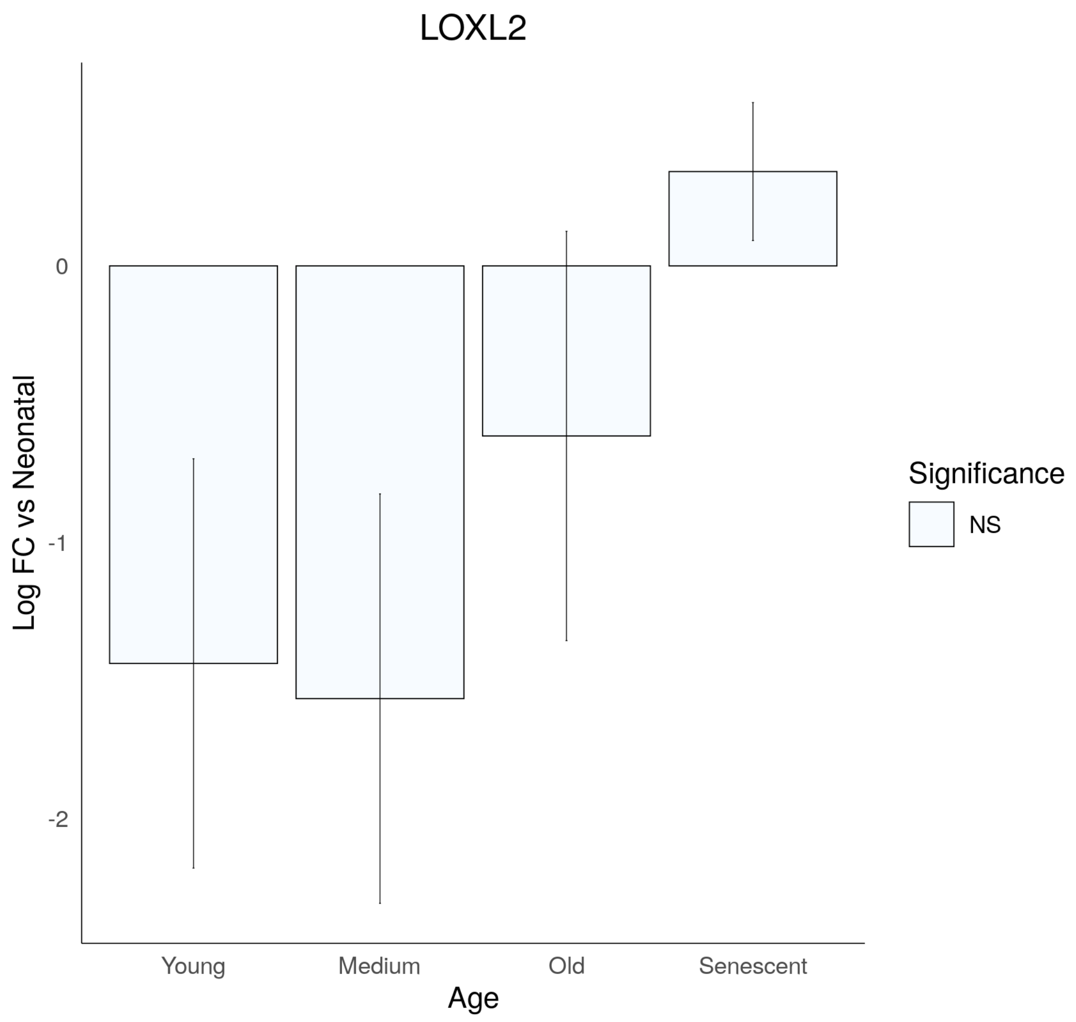

LOXL2
Log FC vs Neonatal
0
-1
-2
Young
Medium
Old
Senescent
Age
Significance
NS

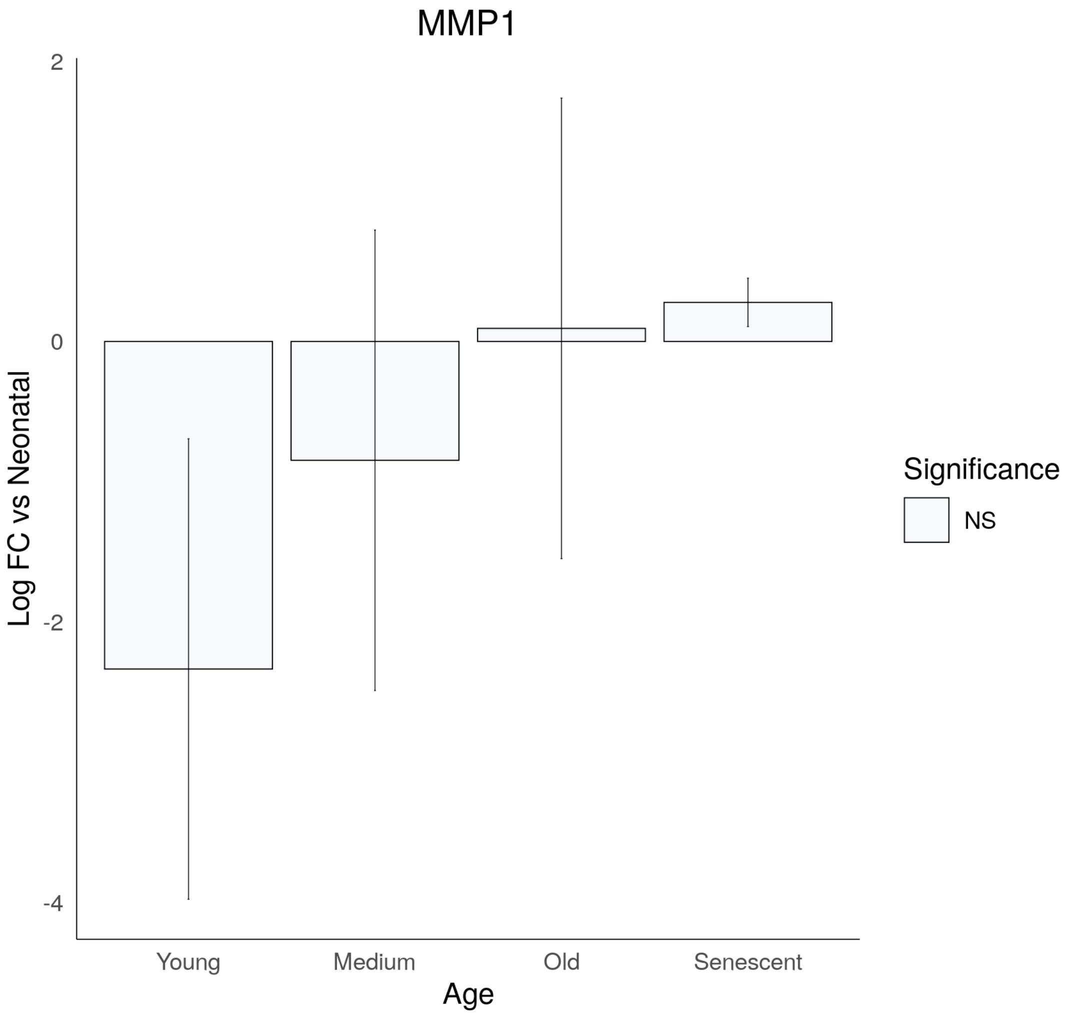
MMP1
Log FC vs Neonatal
2
0
-2
-4
Young
Medium
Old
Senescent
Age
Significance
NS

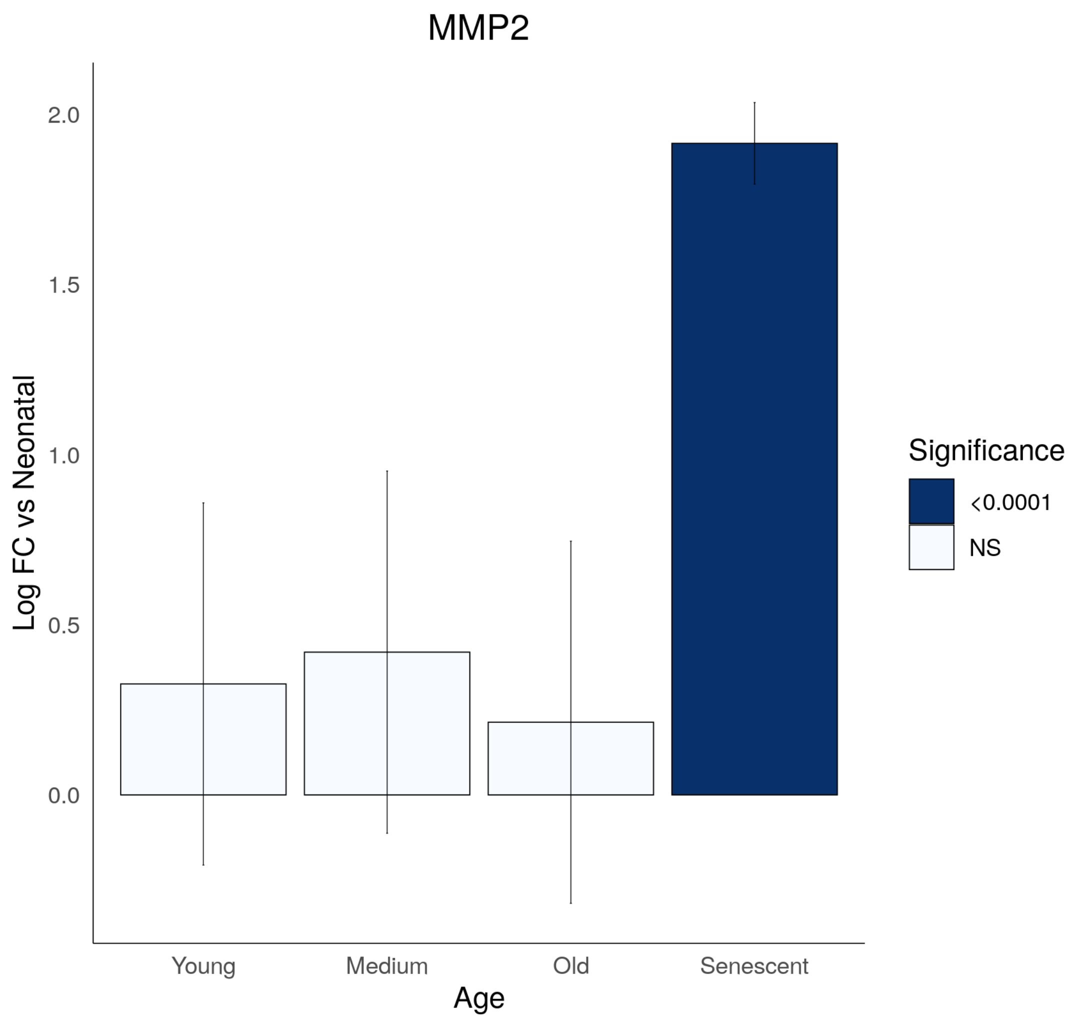
MMP2
2.0
1.5
1.0
0.5
0.0
Log FC vs Neonatal
Young
Medium
Old
Senescent
Age
Significance
<0.0001
NS

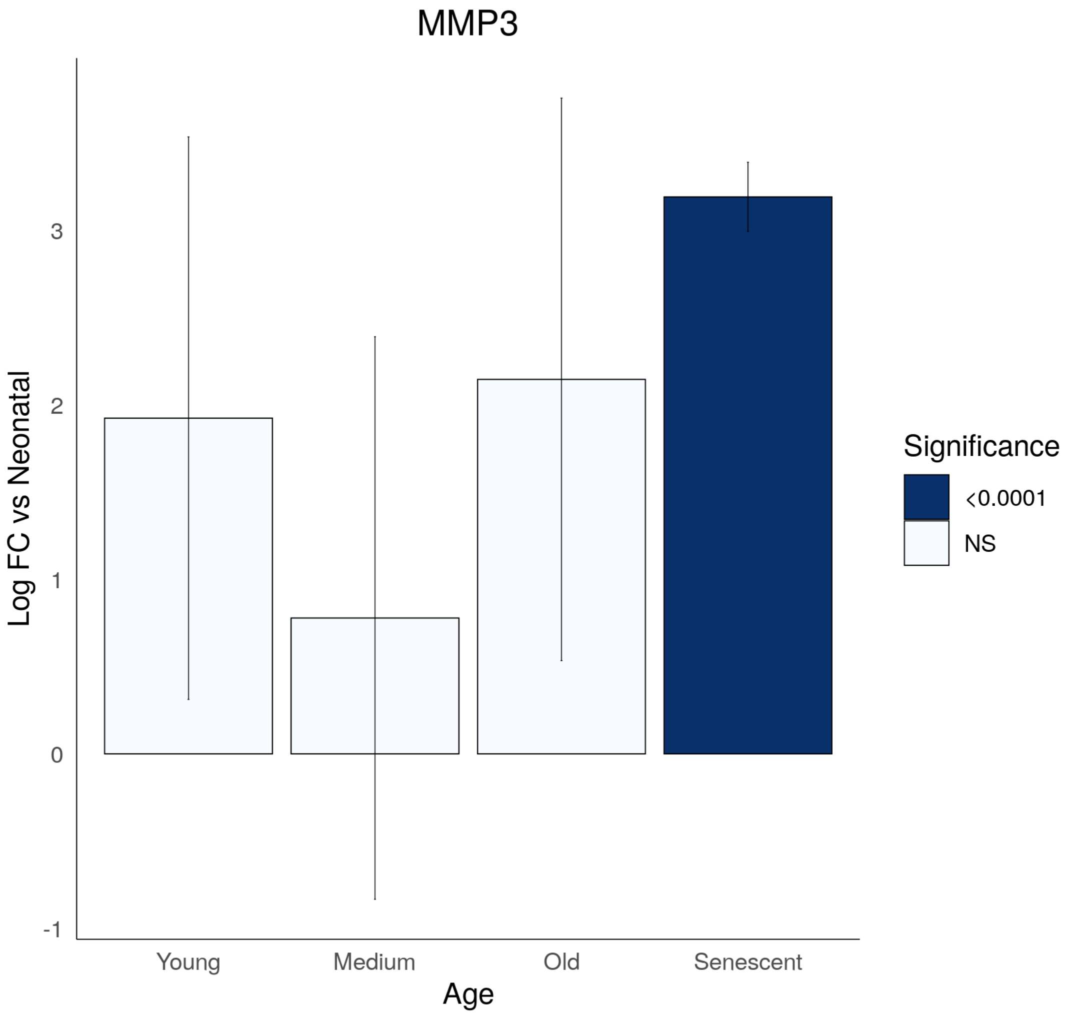
MMP3
Log FC vs Neonatal
3
2
1
0
-1
Young
Medium
Old
Senescent
Age
Significance
<0.0001
NS

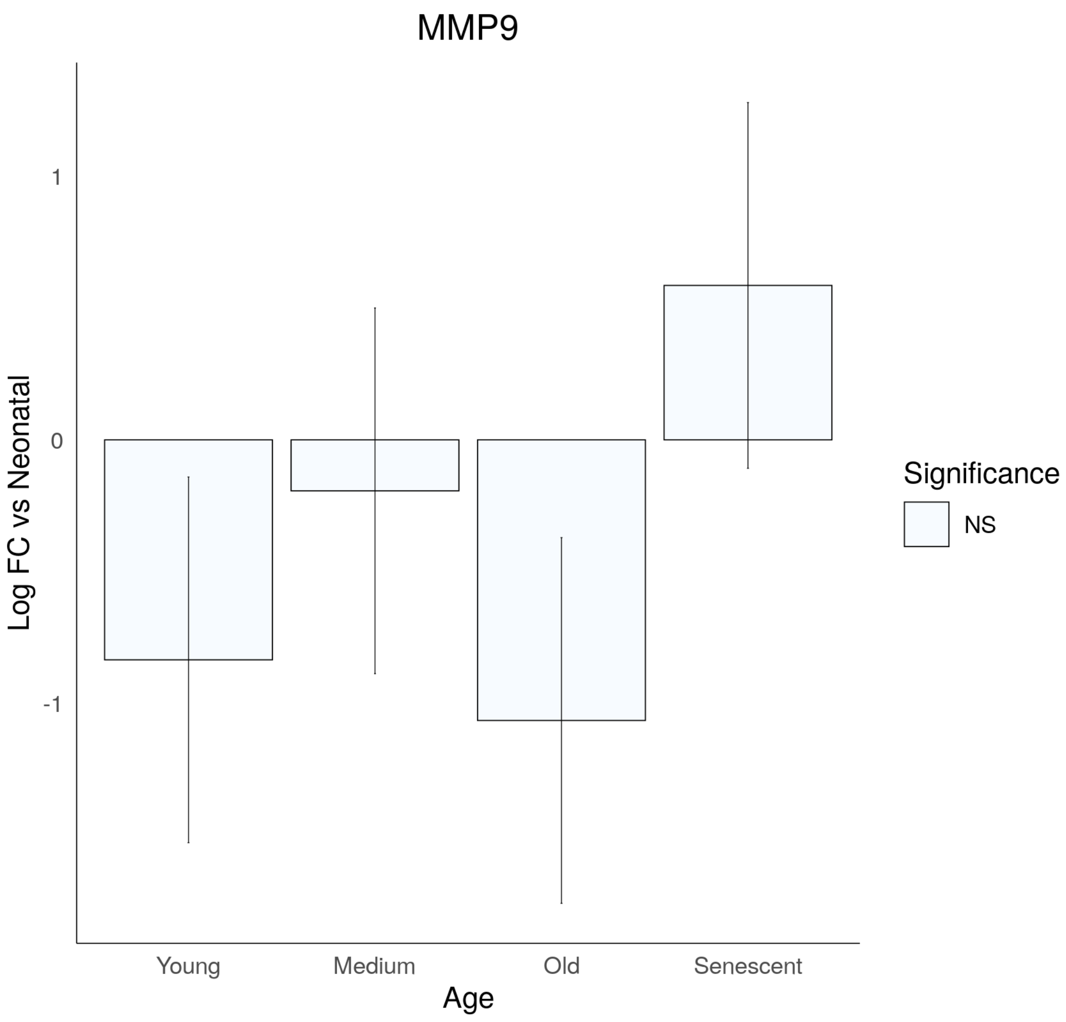

MMP9
Log FC vs Neonatal
1
0
-1
Young
Medium
Old
Senescent
Age
Significance
NS

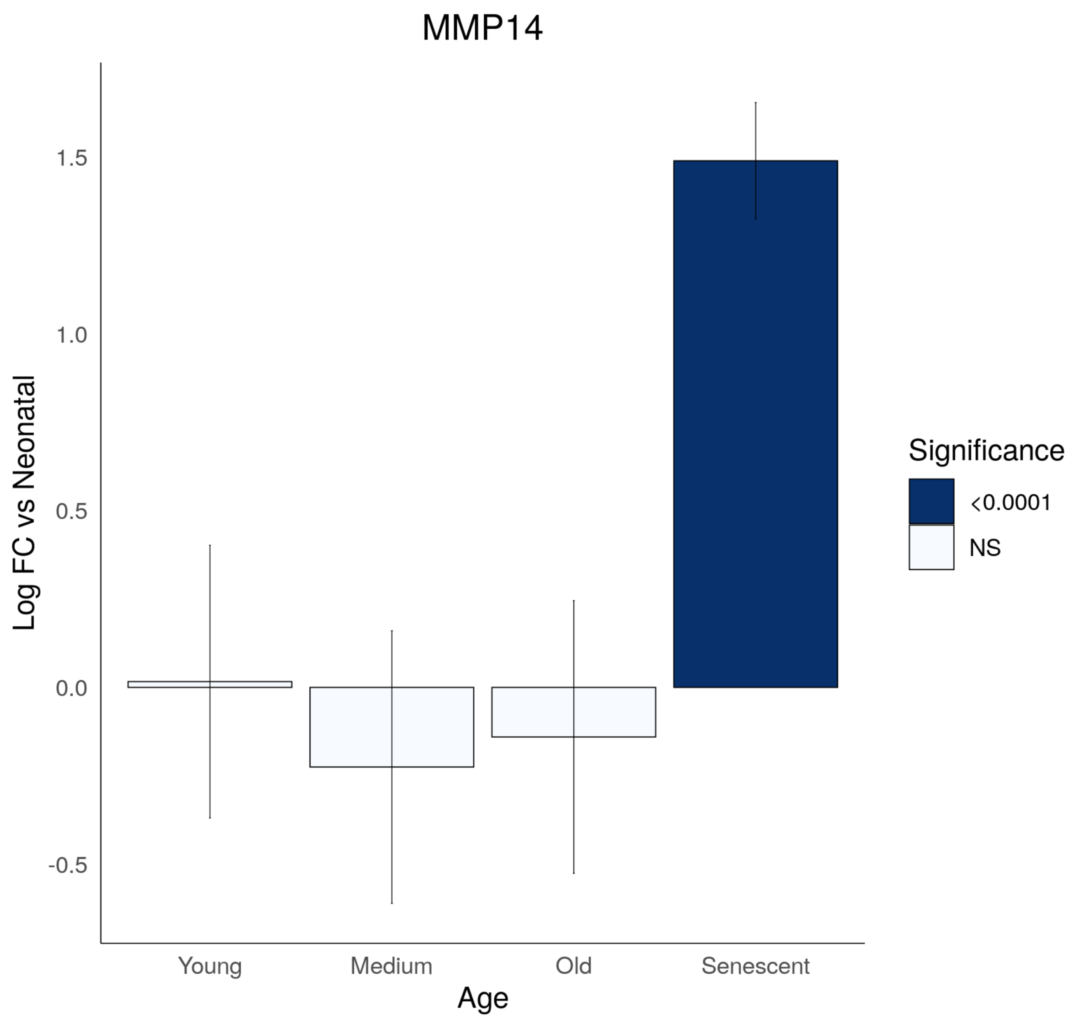

MMP14
Log FC vs Neonatal
1.5
1.0
0.5
0.0
-0.5
Young
Medium
Old
Senescent
Age
Significance
<0.0001
NS

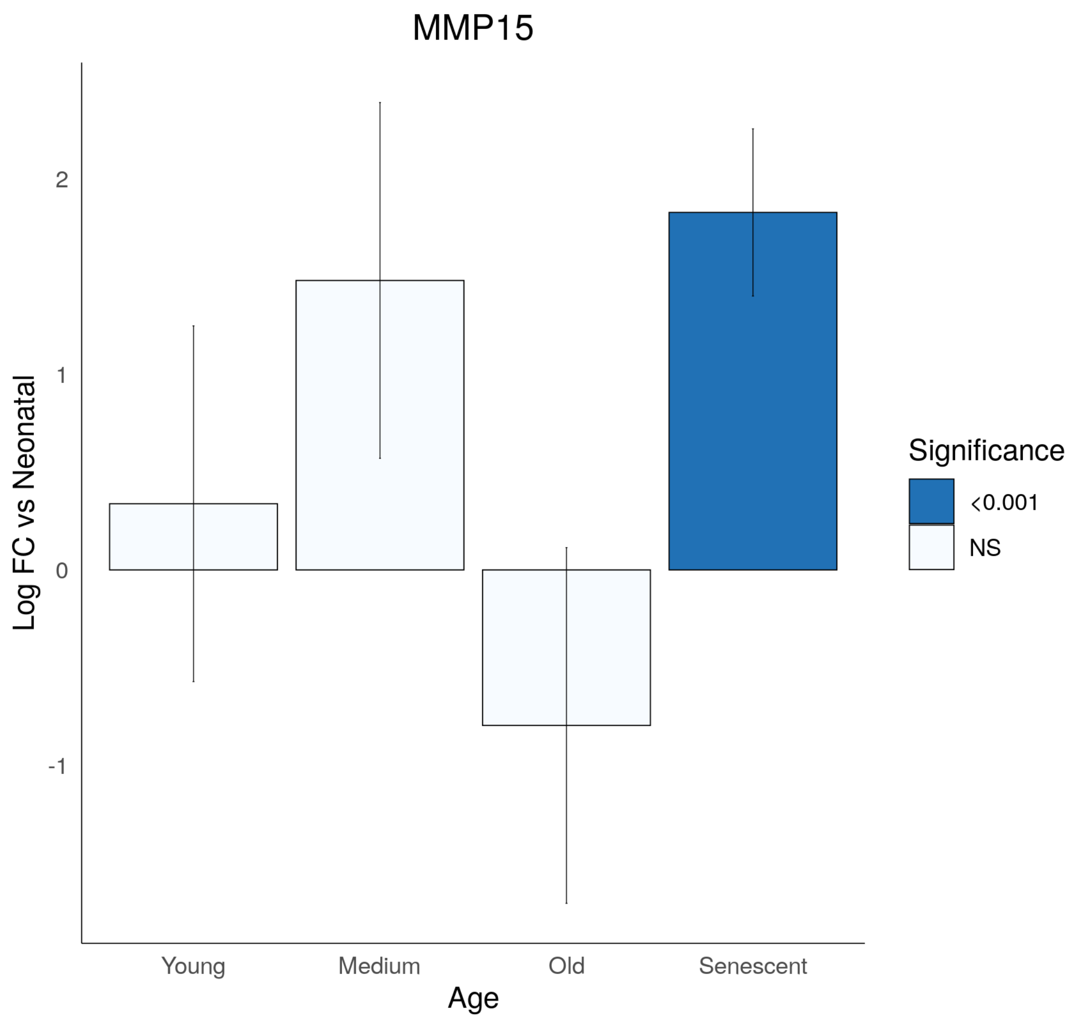

MMP15
Log FC vs Neonatal
2
1
0
-1
Young
Medium
Old
Senescent
Age
Significance
<0.001
NS

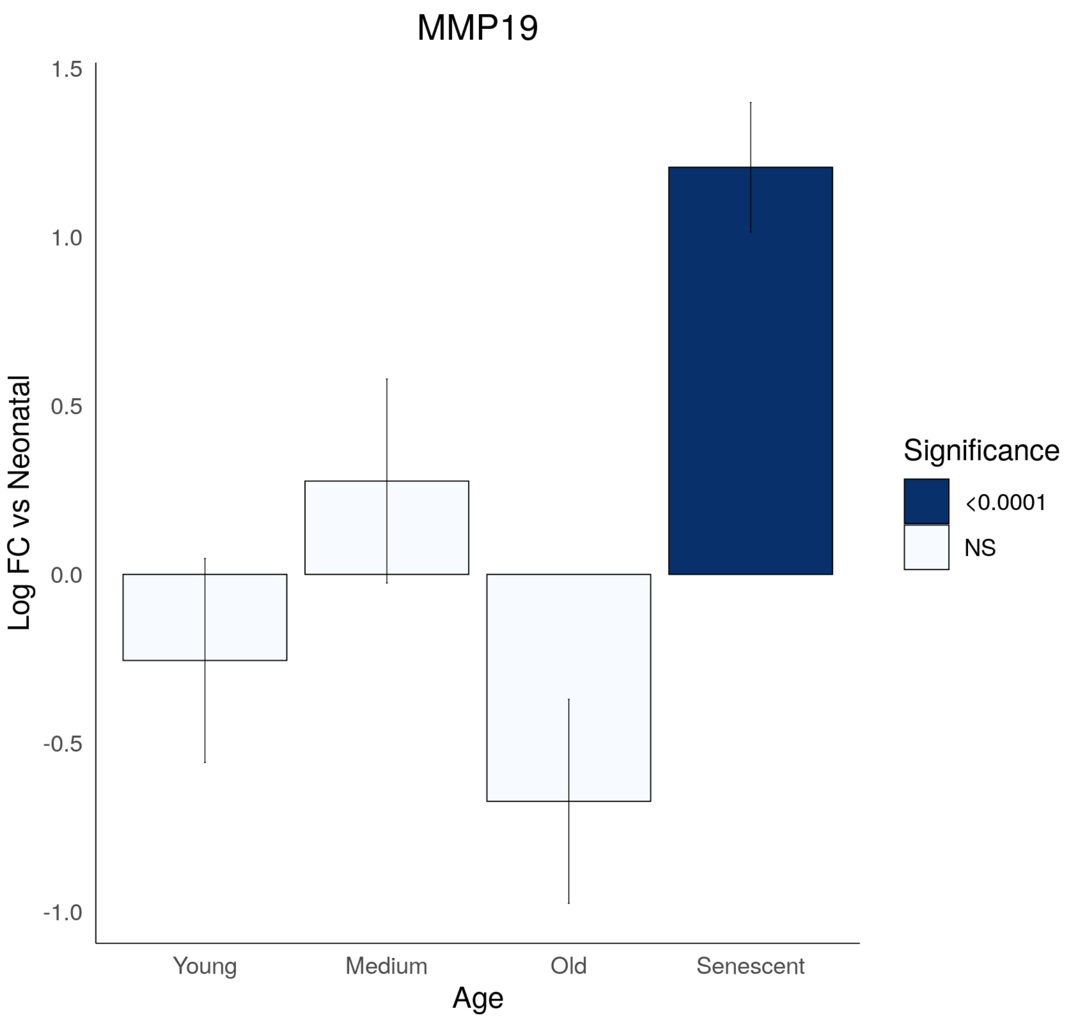

MMP19
Log FC vs Neonatal
1.5
1.0
0.5
0.0
-0.5
-1.0
Young
Medium
Old
Senescent
Age
Significance
<0.0001
NS

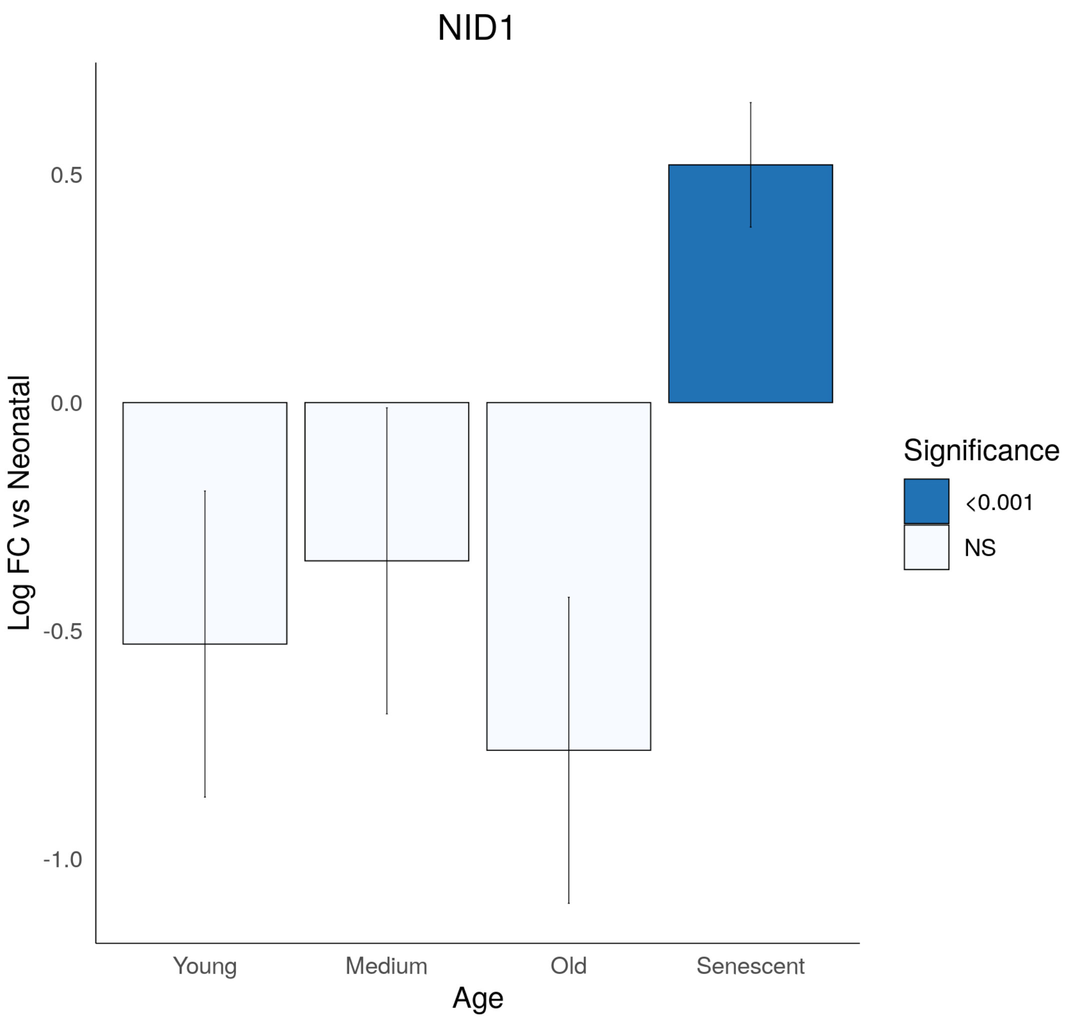

NID1
Log FC vs Neonatal
0.5
0.0
-0.5
-1.0
Young
Medium
Old
Senescent
Age
Significance
<0.001
NS

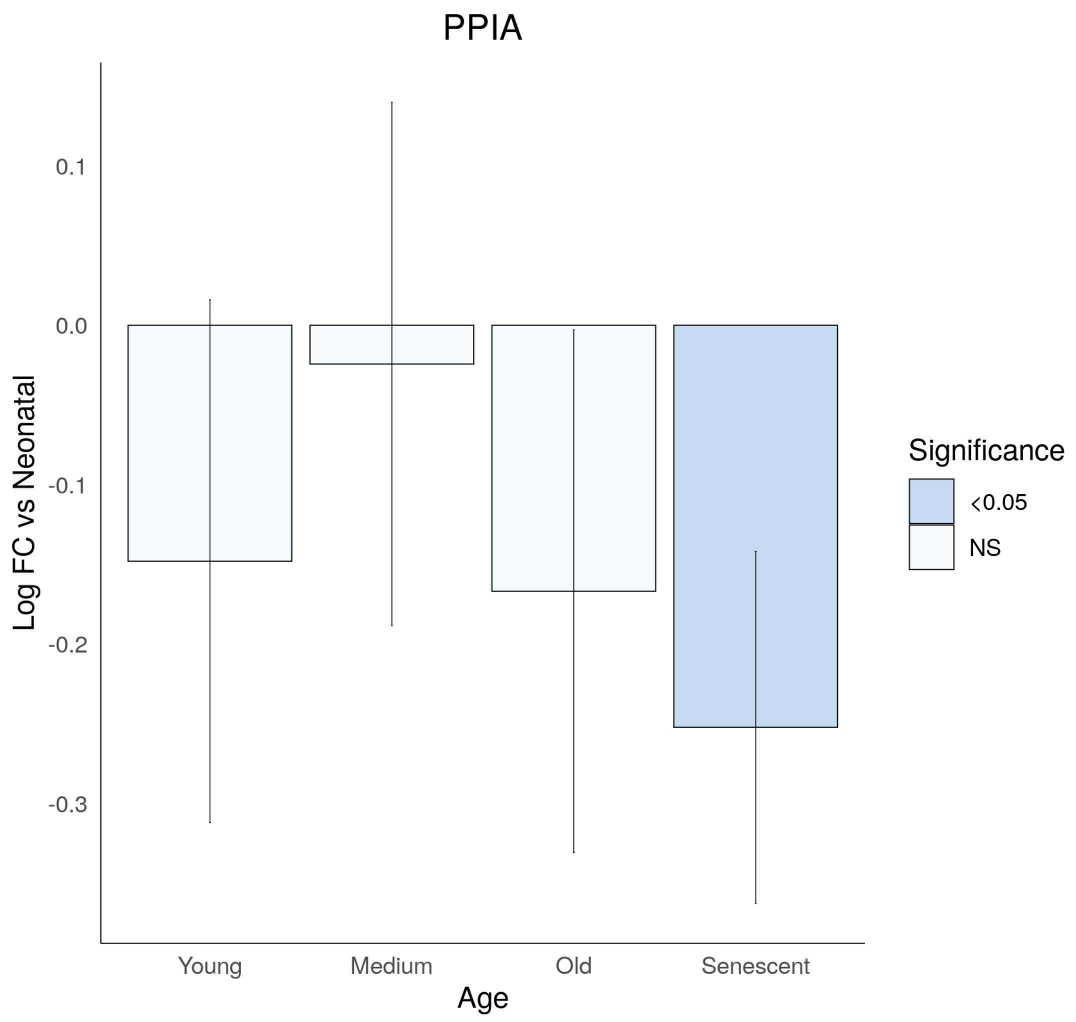
PPIA
Log FC vs Neonatal
0.1
0.0
-0.1
-0.2
-0.3
Young
Medium
Old
Senescent
Age
Significance
<0.05
NS

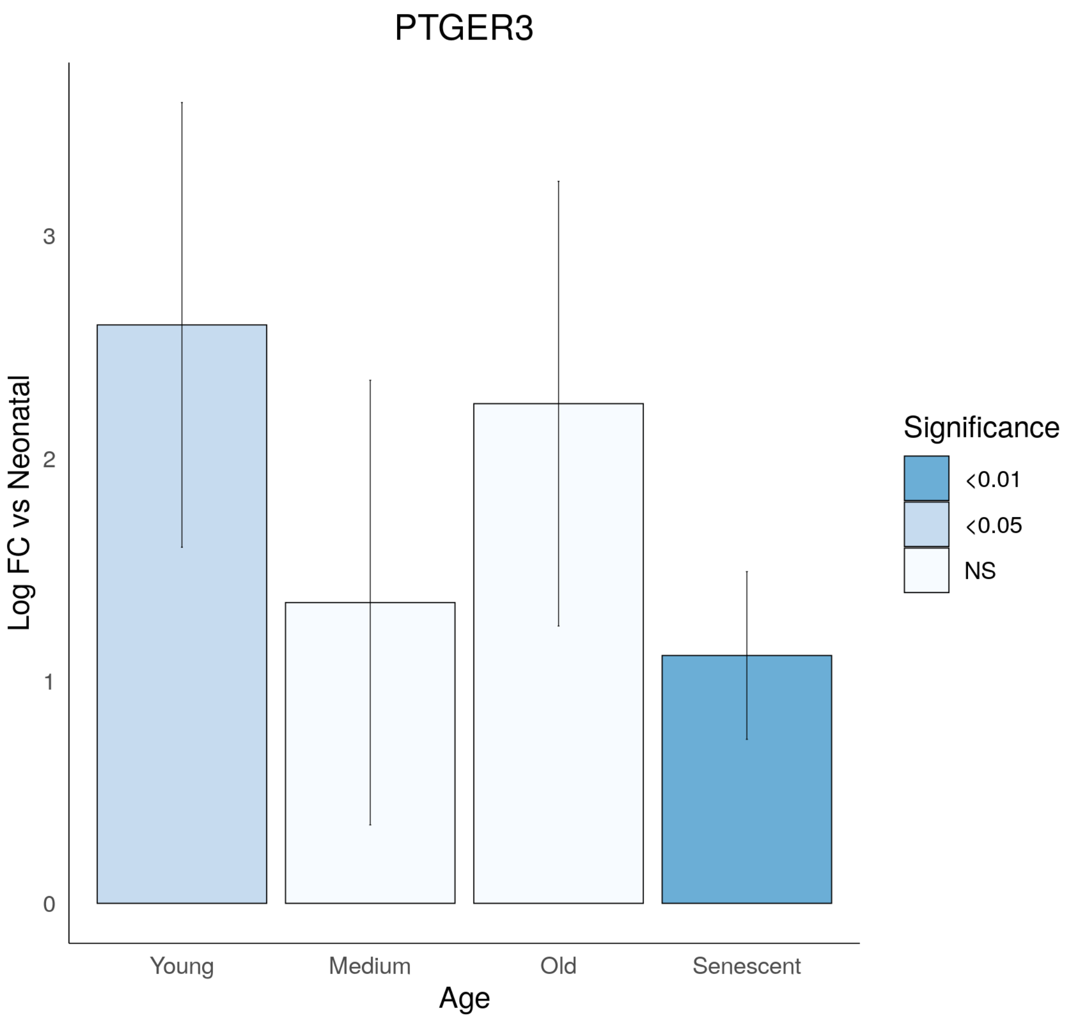

PTGER3
Log FC vs Neonatal
0
1
2
3
Young
Medium
Old
Senescent
Age
Significance
<0.01
<0.05
NS

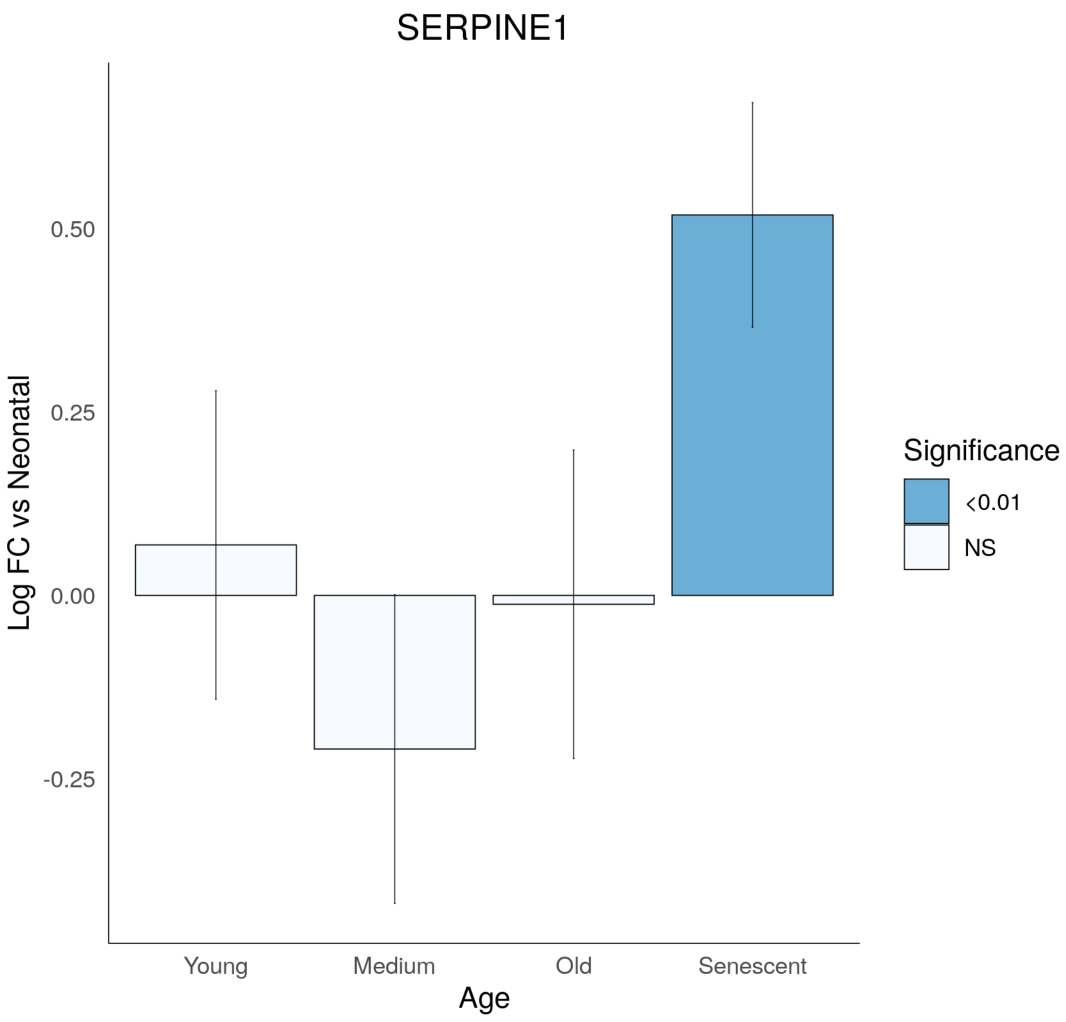
SERPINE1
Log FC vs Neonatal
0.50
0.25
0.00
-0.25
Young
Medium
Old
Senescent
Age
Significance
<0.01
NS

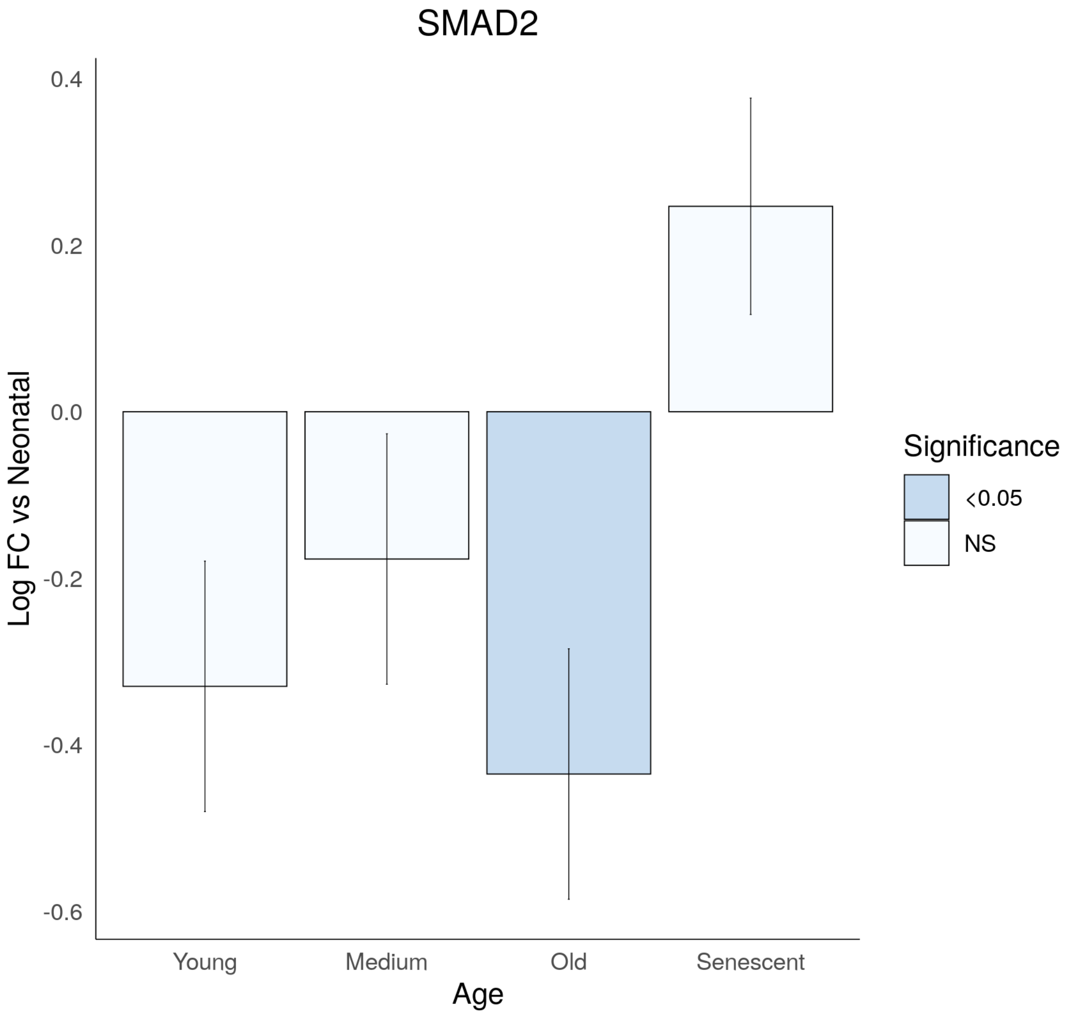
SMAD2
Log FC vs Neonatal
0.4
0.2
0.0
-0.2
-0.4
-0.6
Young
Medium
Old
Senescent
Age
Significance
<0.05
NS

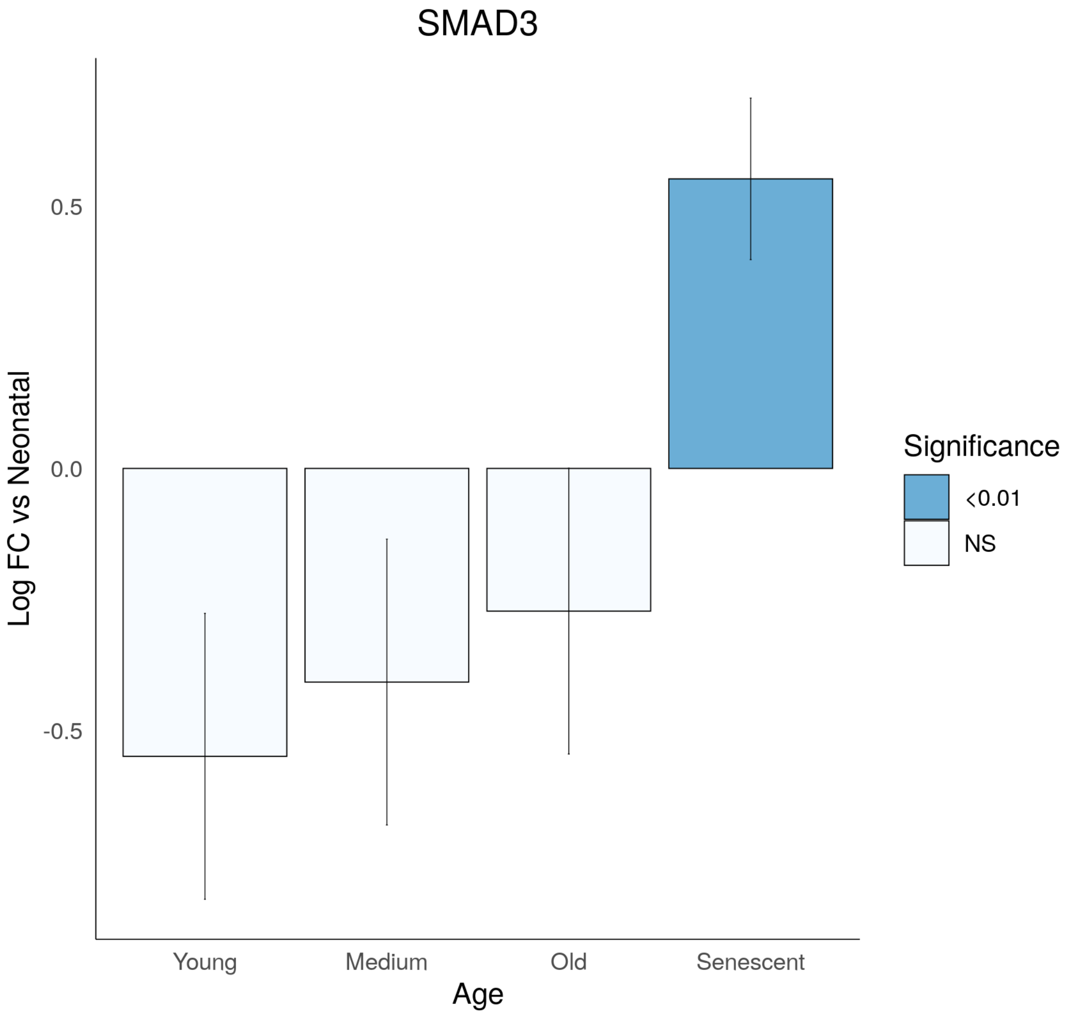
SMAD3
Log FC vs Neonatal
0.5
0.0
-0.5
Young
Medium
Old
Senescent
Age
Significance
<0.01
NS

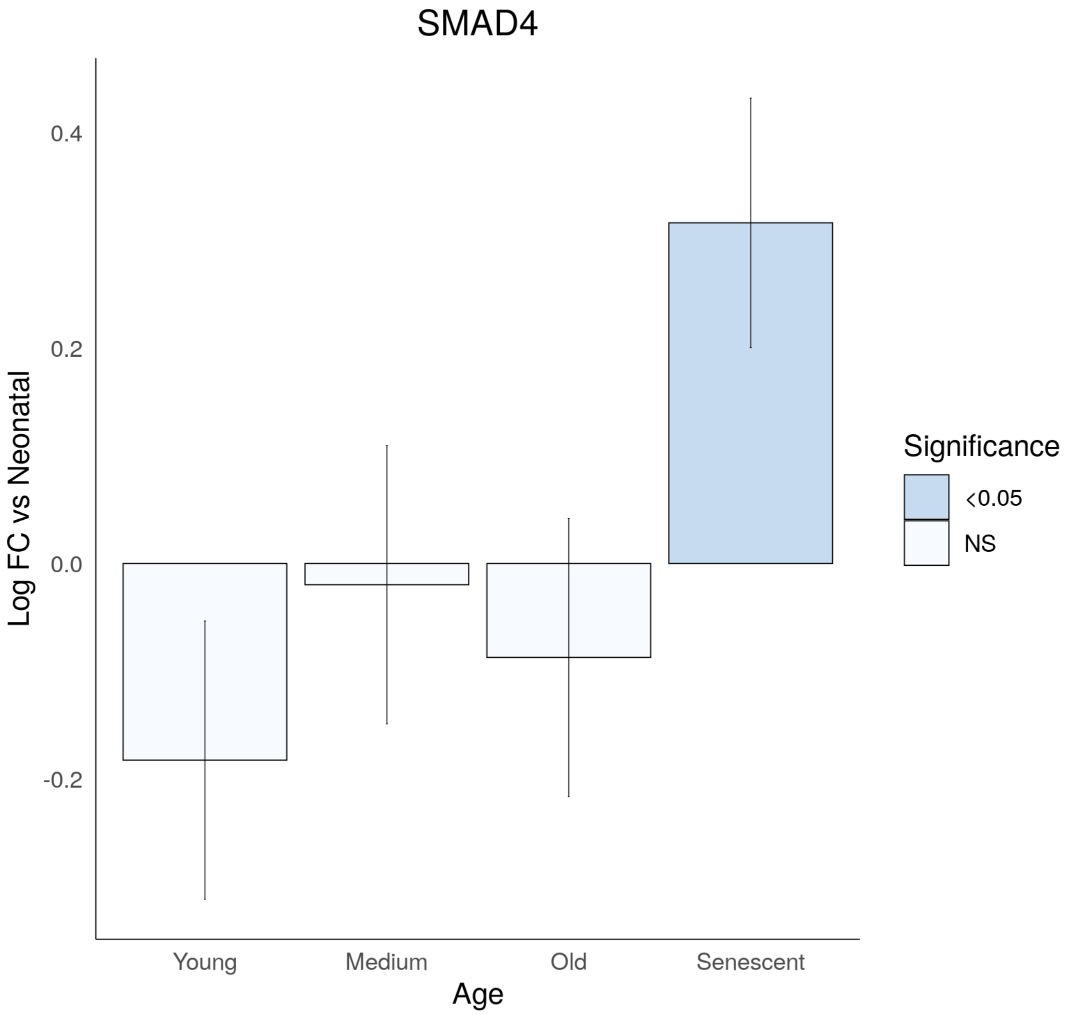
SMAD4
Log FC vs Neonatal
0.4
0.2
0.0
-0.2
Young
Medium
Old
Senescent
Age
Significance
<0.05
NS

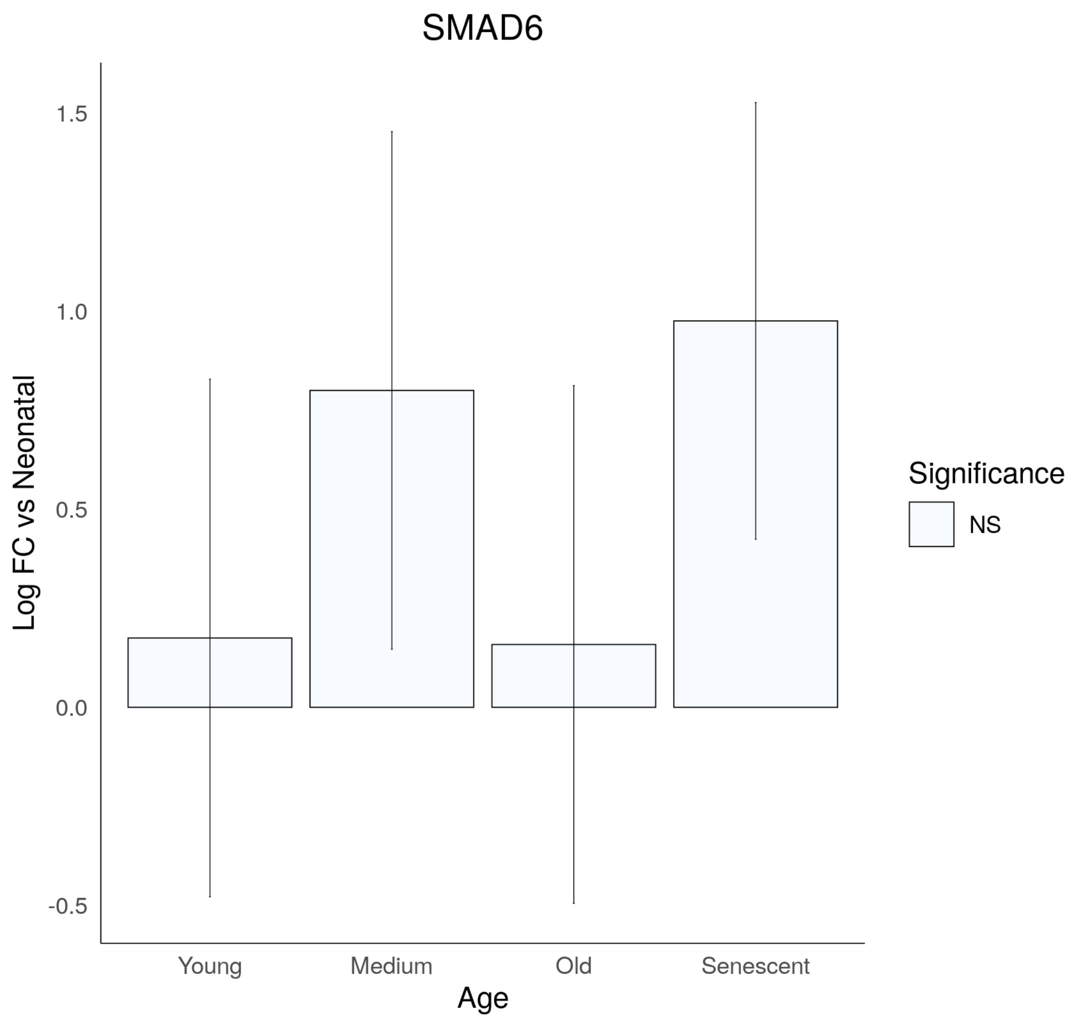
SMAD6
Log FC vs Neonatal
1.5
1.0
0.5
0.0
-0.5
Young
Medium
Old
Senescent
Age
Significance
NS

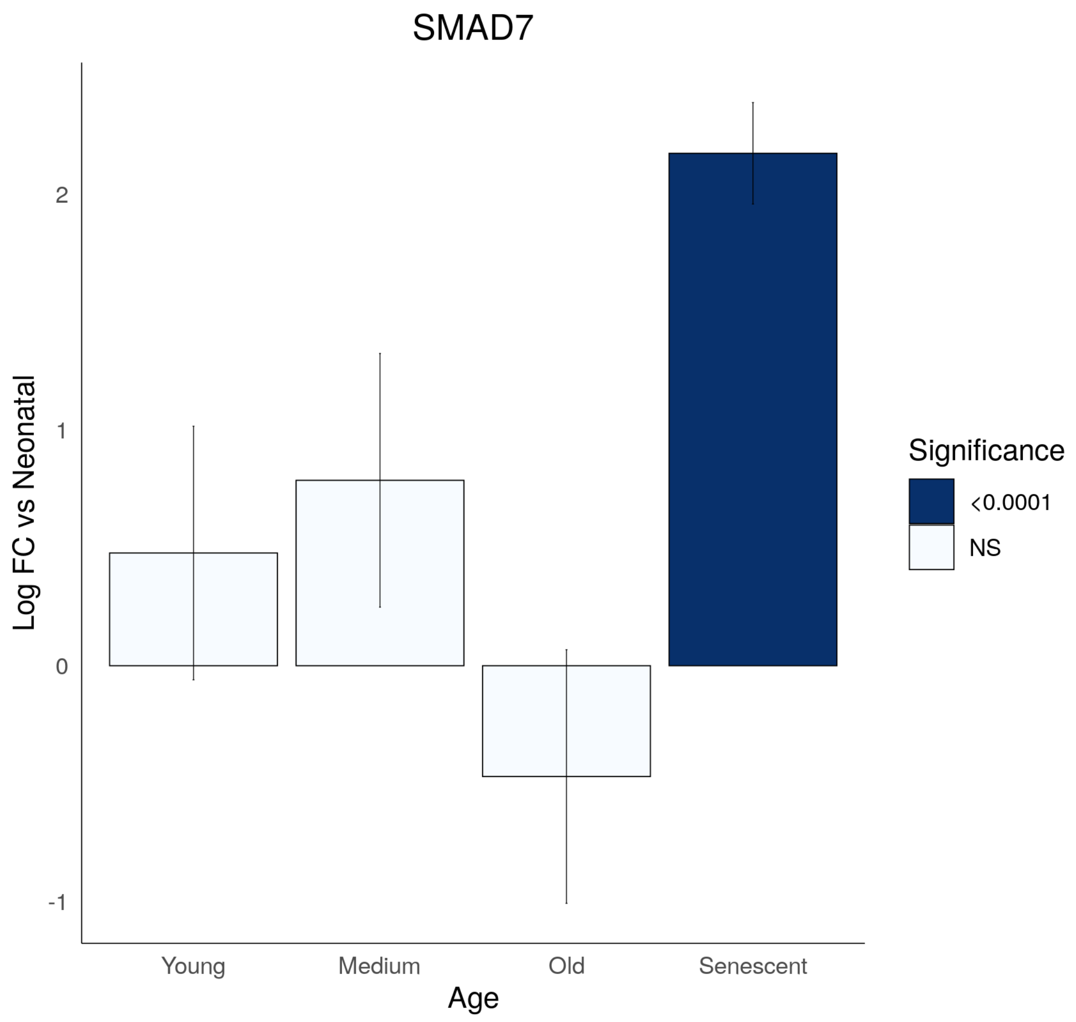

SMAD7
Log FC vs Neonatal
2
1
0
-1
Young
Medium
Old
Senescent
Age
Significance
<0.0001
NS

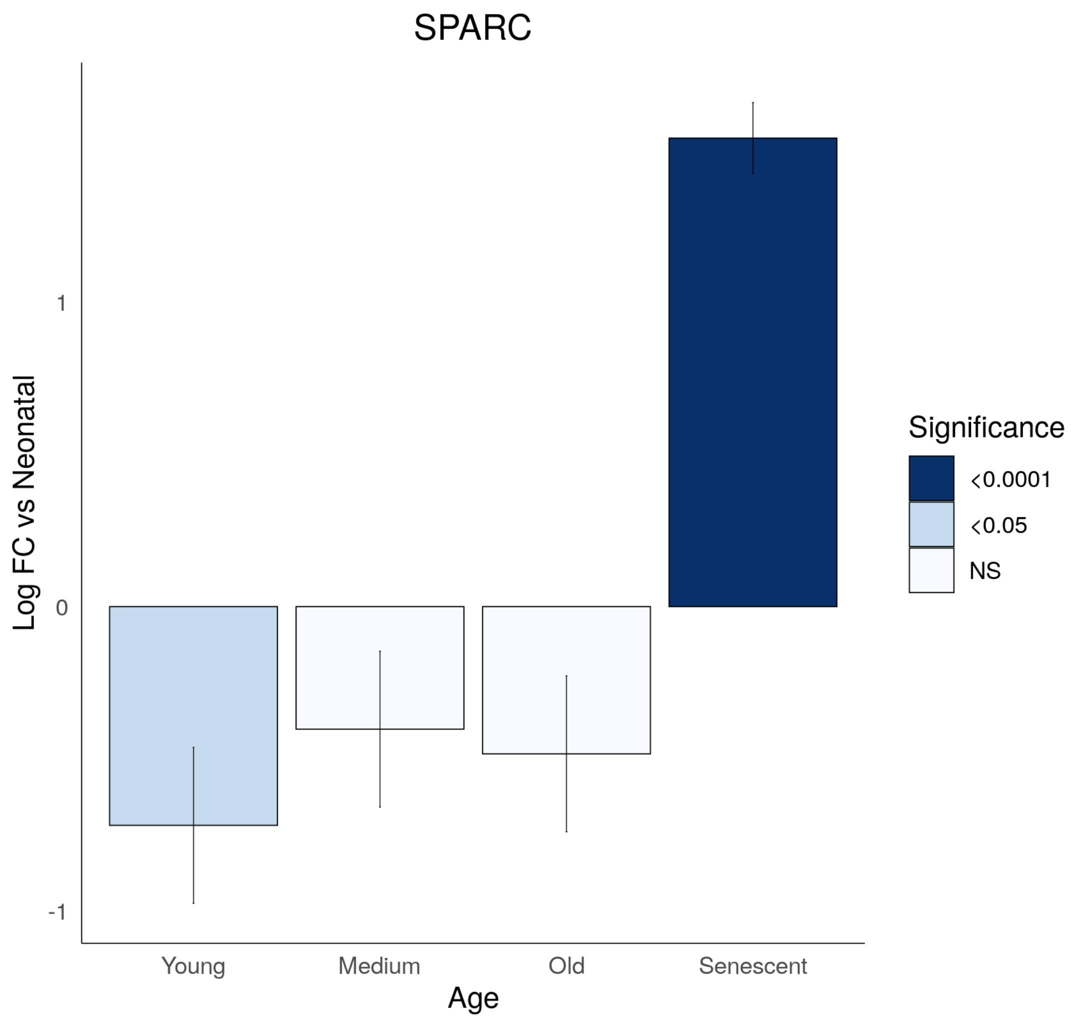

SPARC
Log FC vs Neonatal
1
0
-1
Young
Medium
Old
Senescent
Age
Significance
<0.0001
<0.05
NS

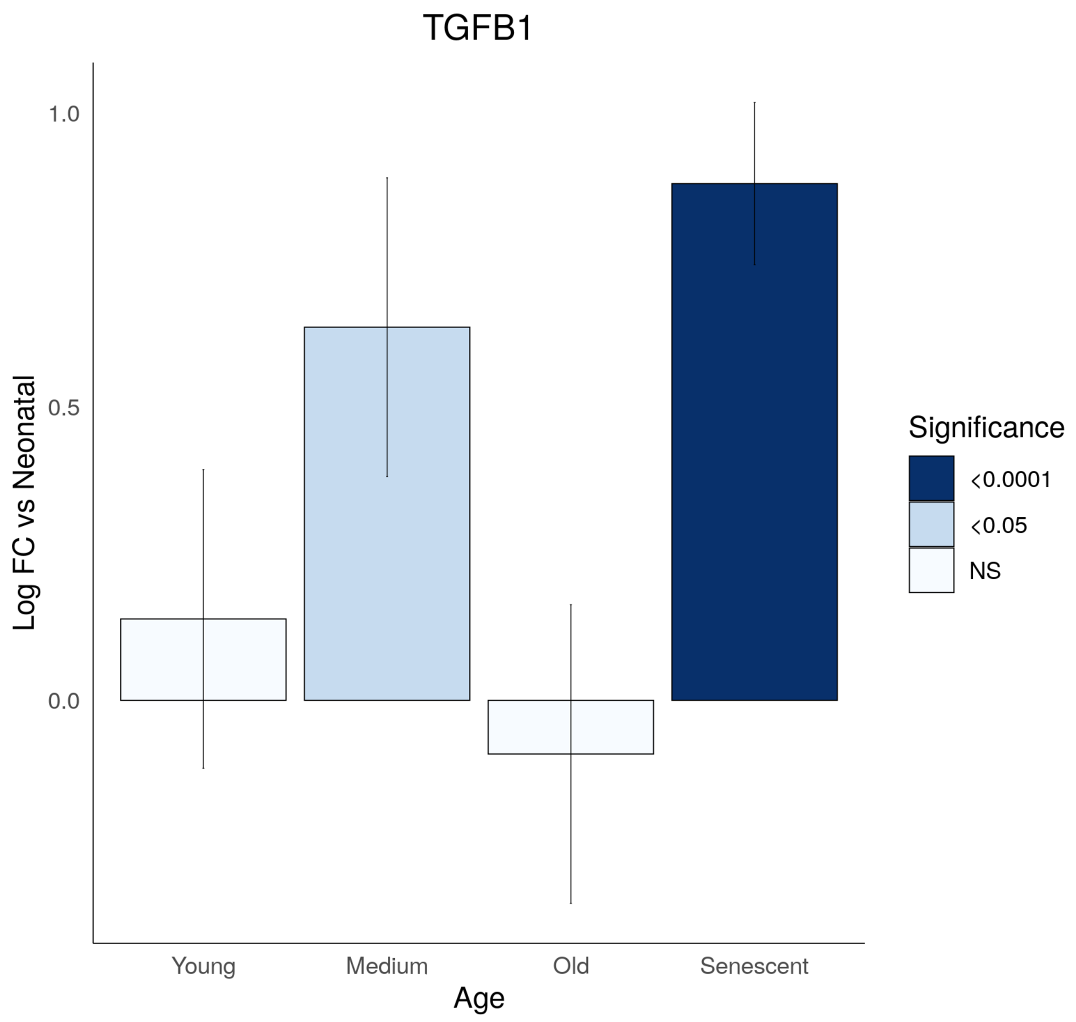
TGFB1
1.0
0.5
0.0
Log FC vs Neonatal
Young
Medium
Old
Senescent
Age
Significance
<0.0001
<0.05
NS

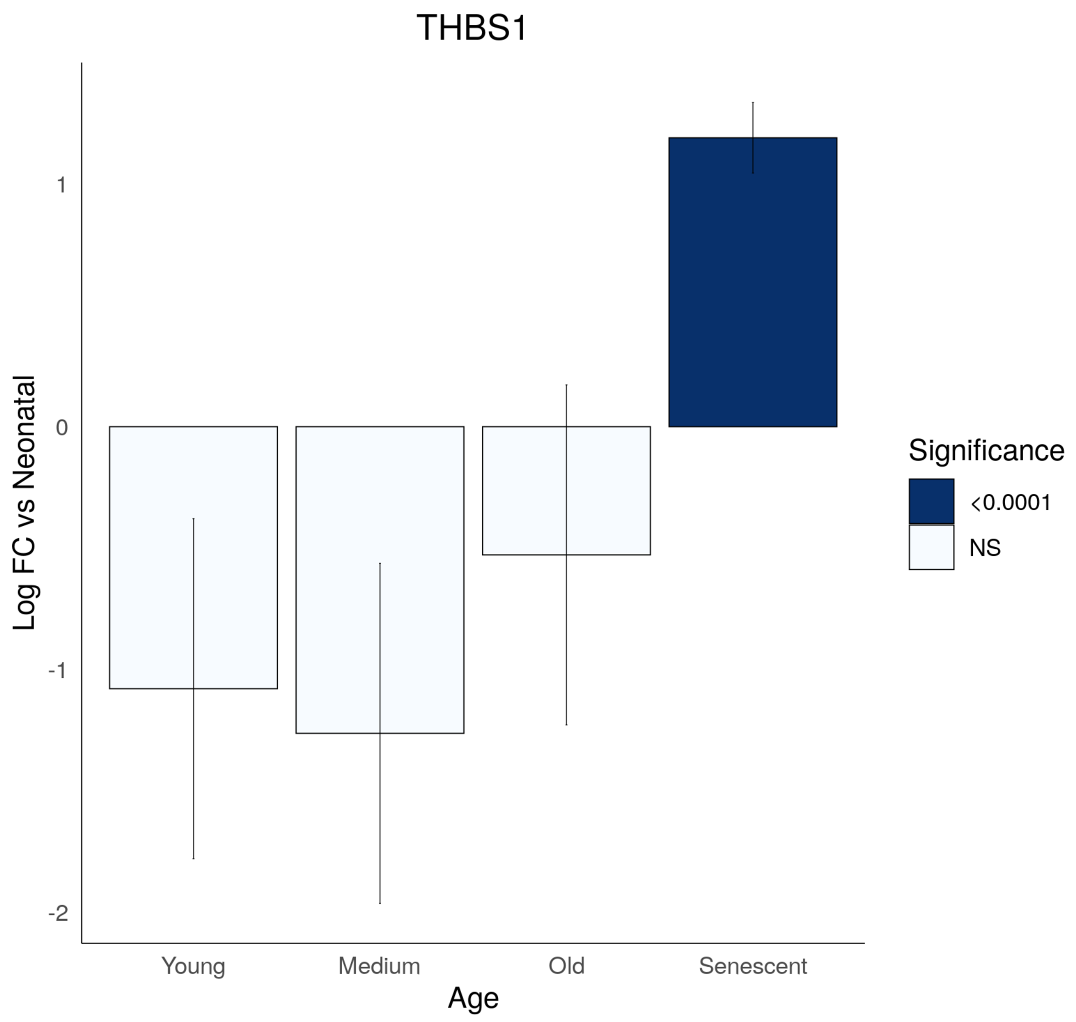
THBS1
Log FC vs Neonatal
1
0
-1
-2
Young
Medium
Old
Senescent
Age
Significance
<0.0001
NS

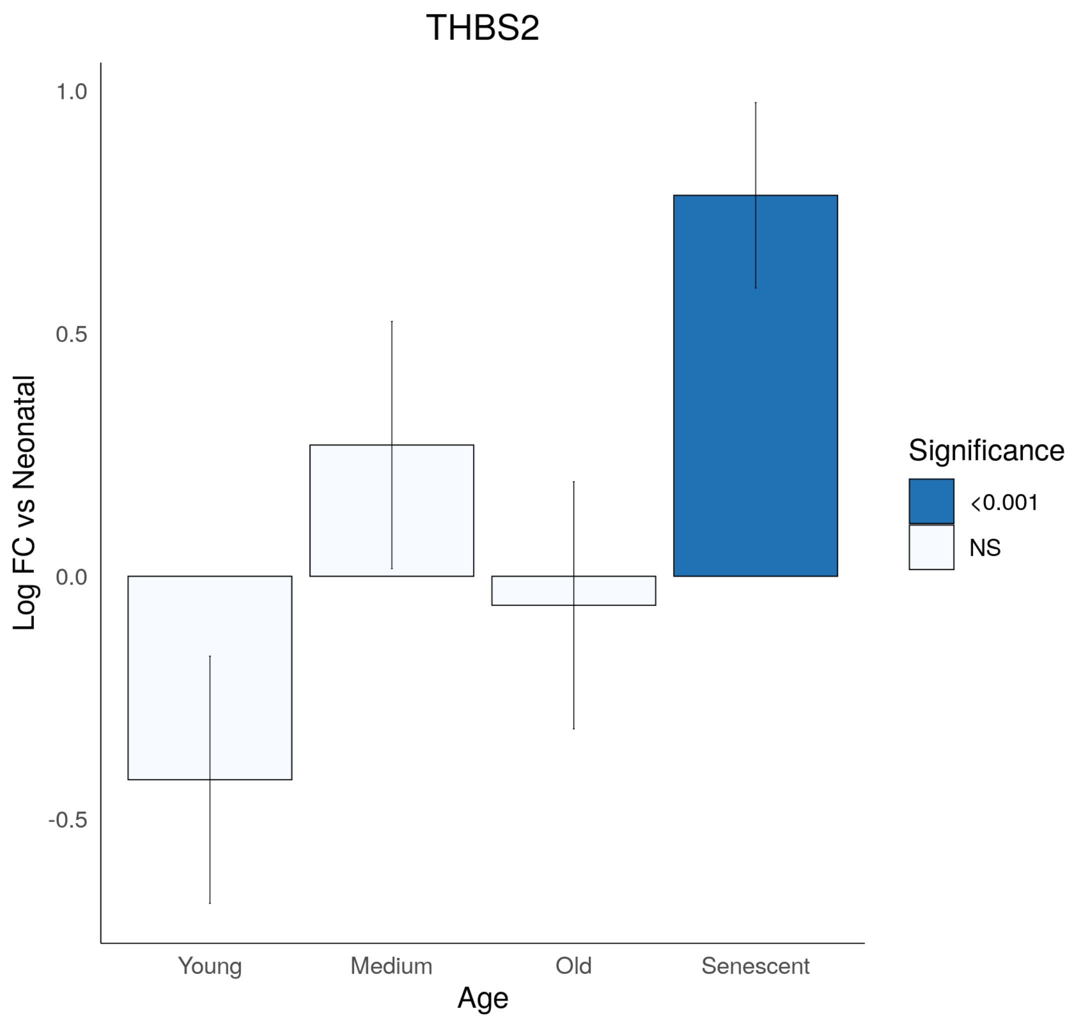
THBS2
1.0
0.5
0.0
-0.5
Log FC vs Neonatal
Young
Medium
Old
Senescent
Age
Significance
<0.001
NS

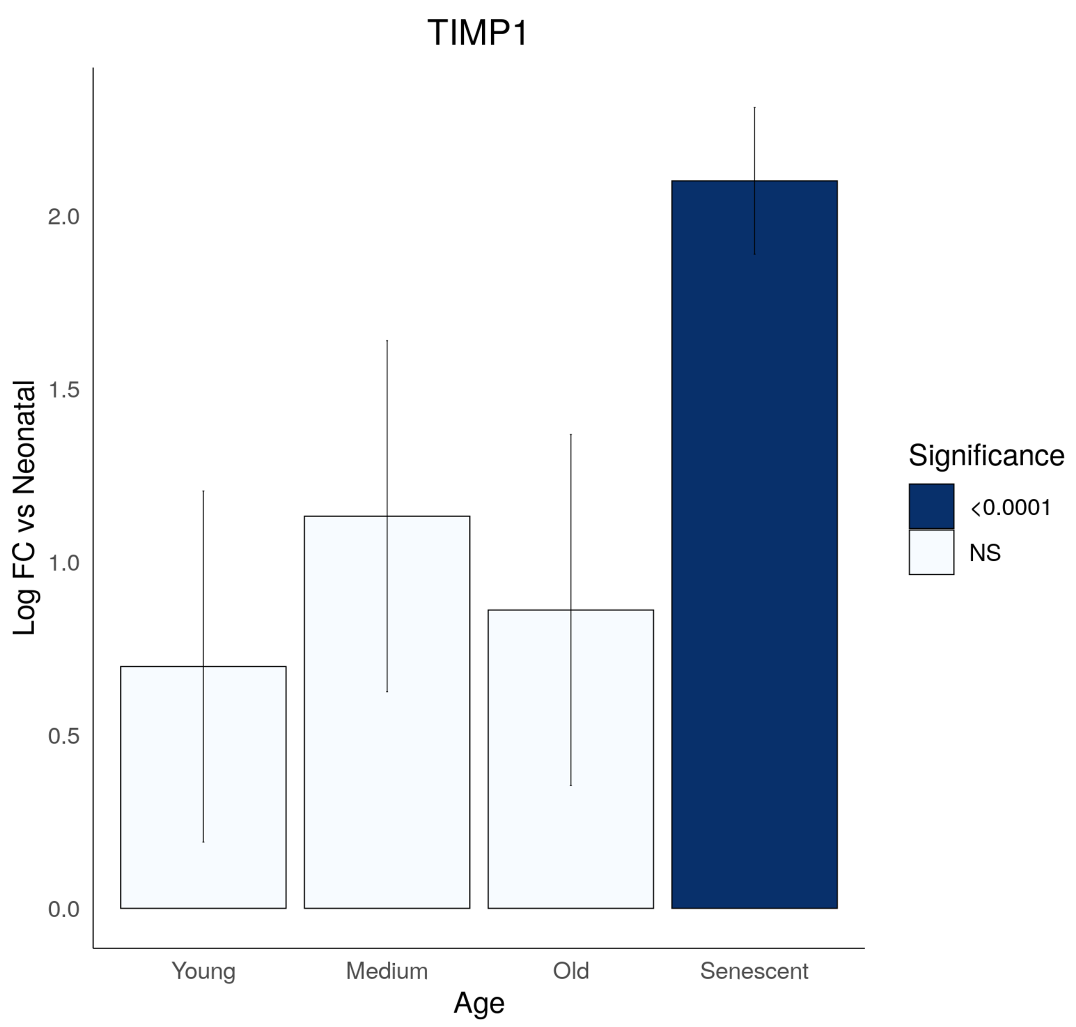
TIMP1
Log FC vs Neonatal
2.0
1.5
1.0
0.5
0.0
Young
Medium
Old
Senescent
Age
Significance
<0.0001
NS

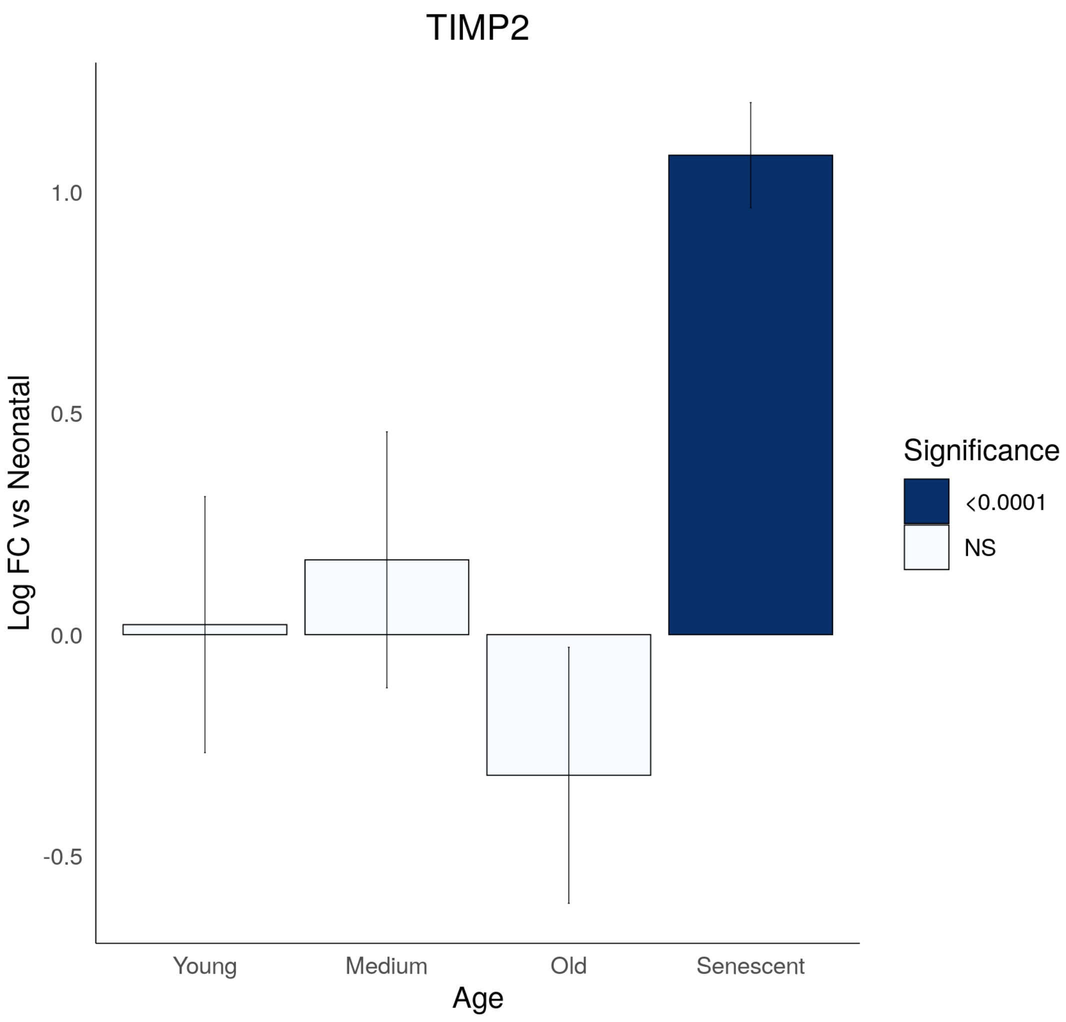
TIMP2
Log FC vs Neonatal
1.0
0.5
0.0
-0.5
Young
Medium
Old
Senescent
Age
Significance
<0.0001
NS

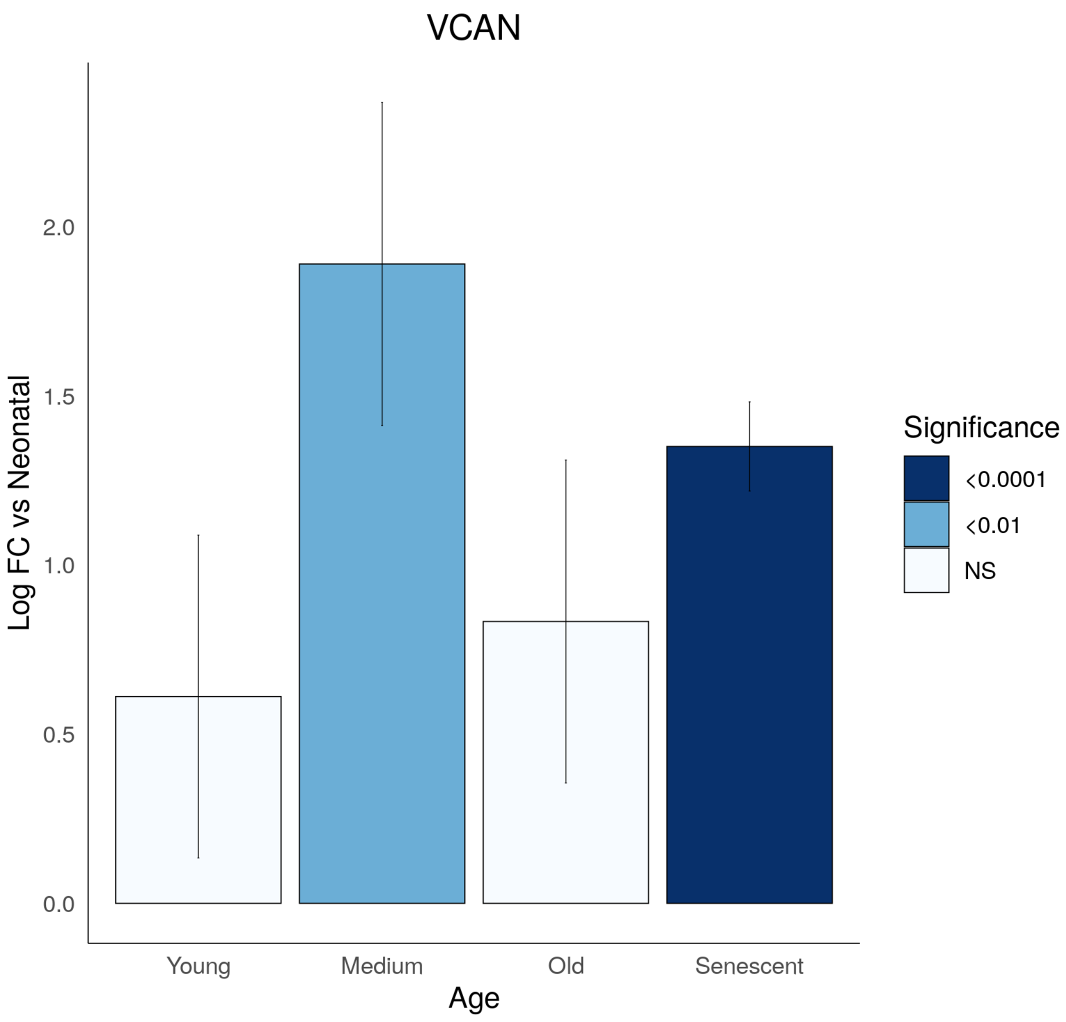
VCAN
Log FC vs Neonatal
2.0
1.5
1.0
0.5
0.0
Young
Medium
Old
Senescent
Age
Significance
<0.0001
<0.01
NS

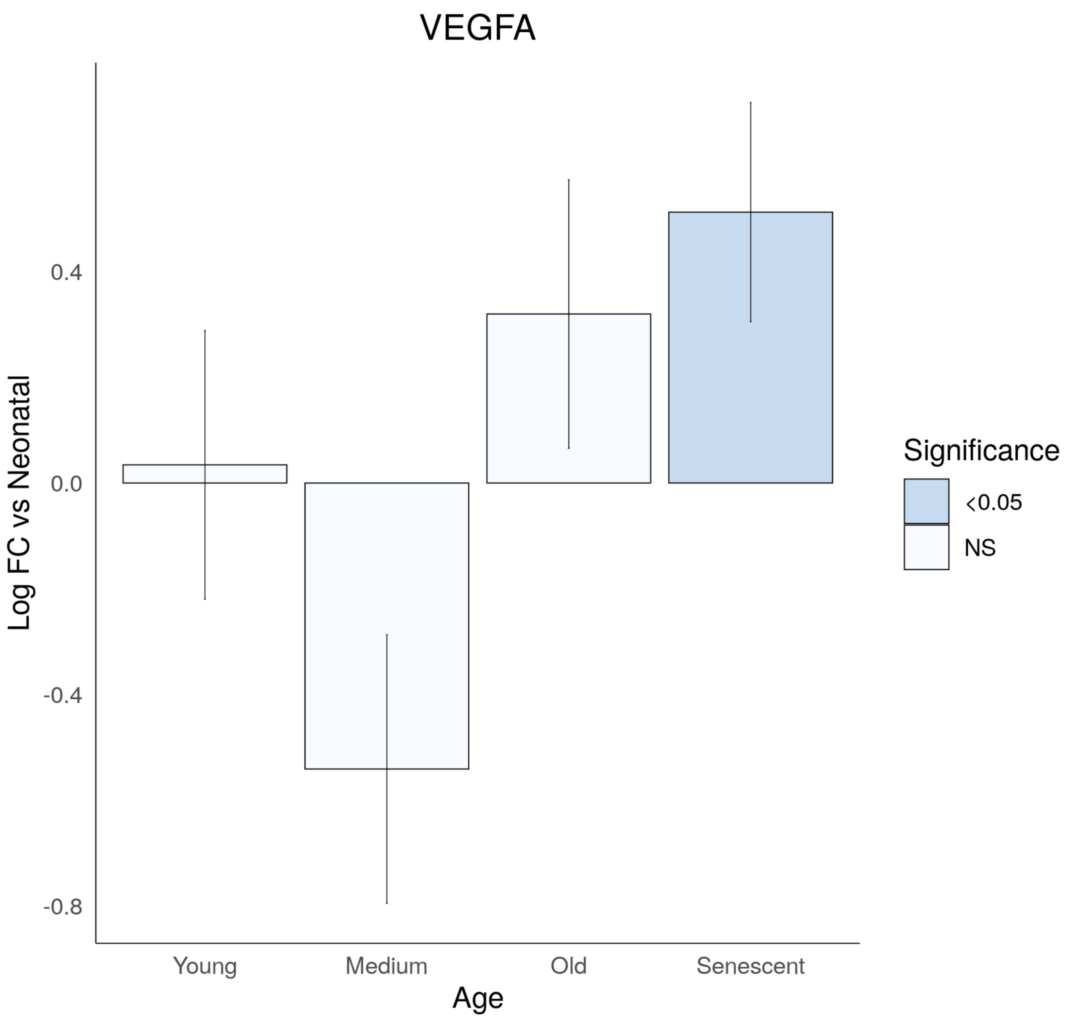
VEGFA
Log FC vs Neonatal
0.4
0.0
-0.4
-0.8
Young
Medium
Old
Senescent
Age
Significance
<0.05
NS

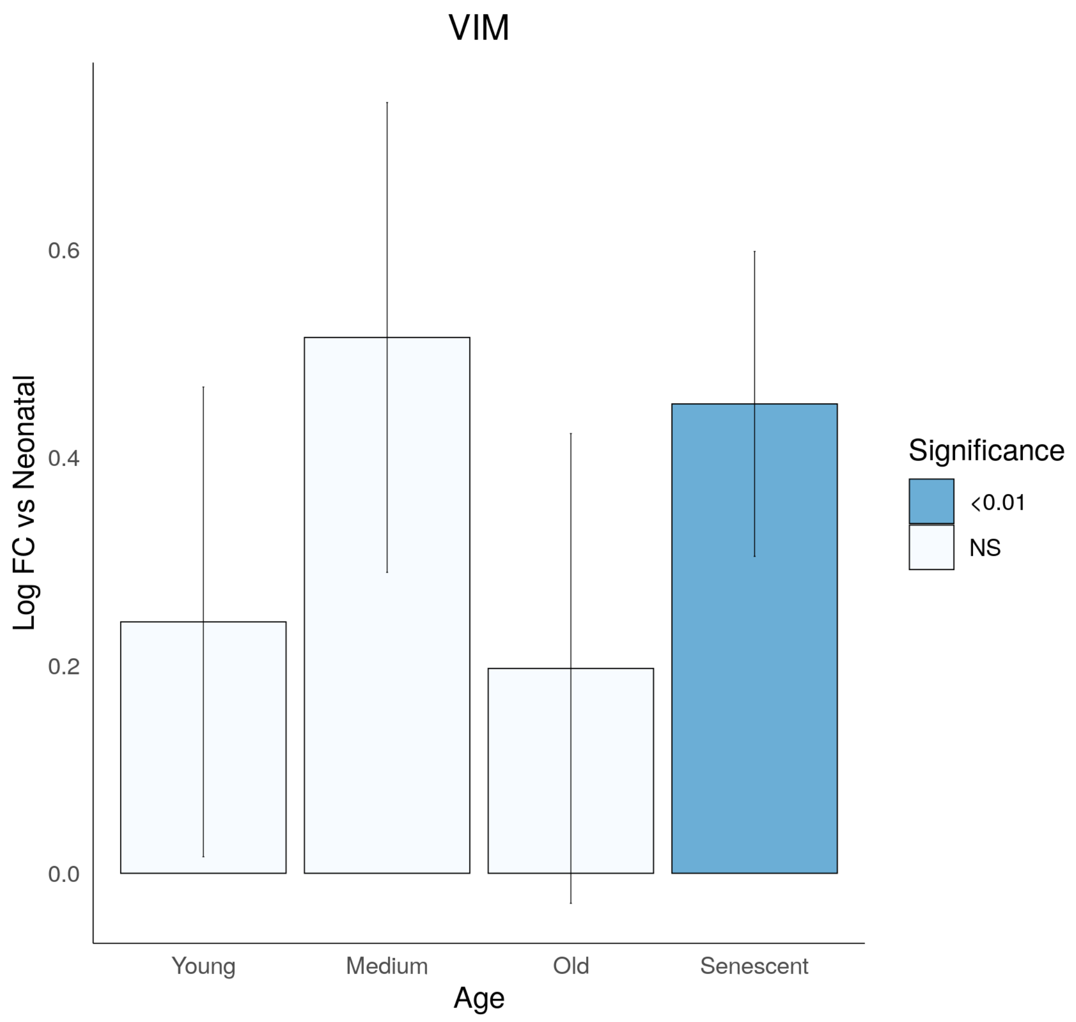

VIM
0.6
0.4
0.2
0.0
Log FC vs Neonatal
Young
Medium
Old
Senescent
Age
Significance
<0.01
NS
