## Supplementary file 6 for "Cell Senescence-Independent Ageing of Human Skin"

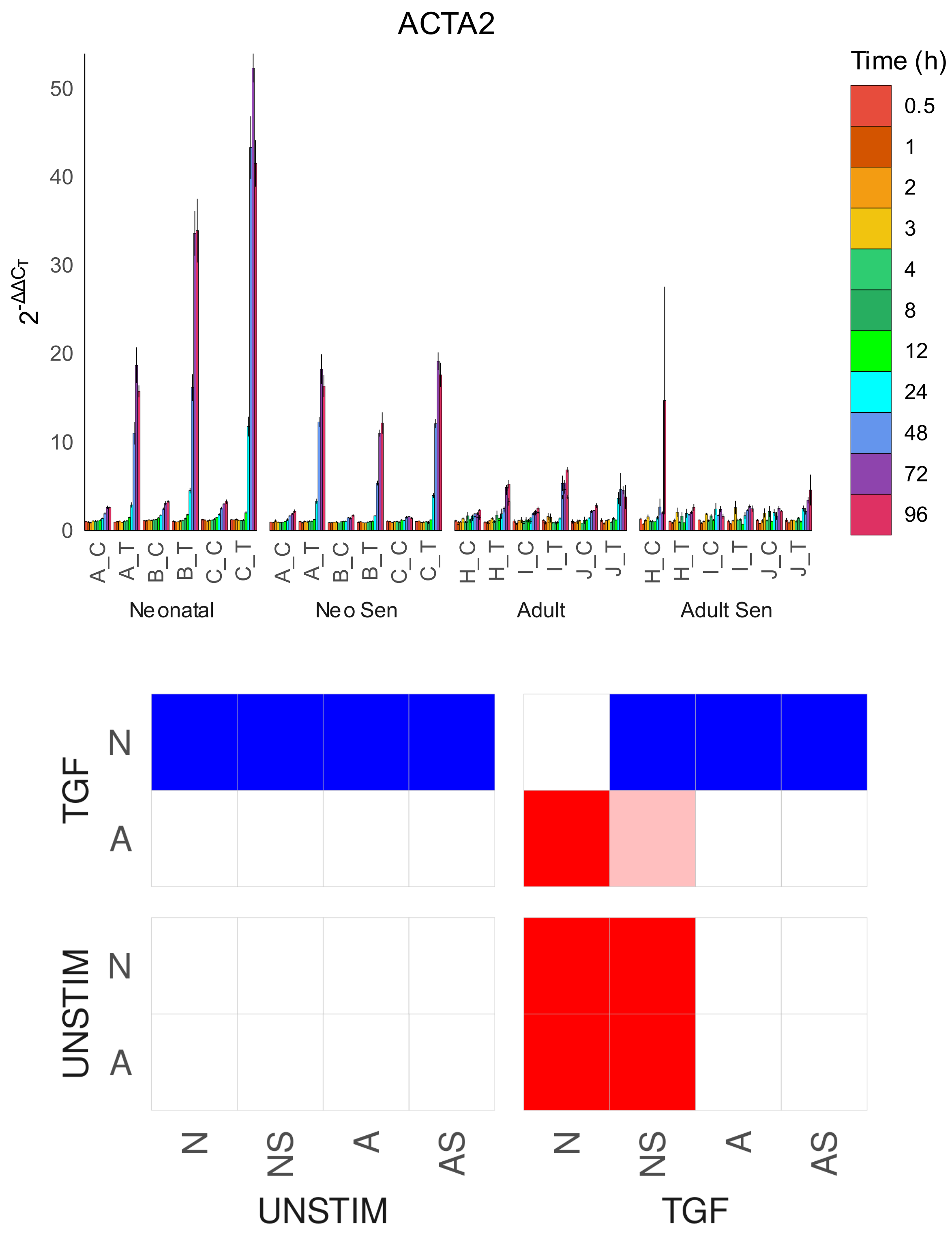
ACTA2
Time (h)
0.5
1
2
3
4
8
12
24
48
72
96
$2^{-\Delta\Delta C_T}$
0
10
20
30
40
50
A_C
A_T
B_C
B_T
C_C
C_T
H_C
H_T
I_C
I_T
J_C
J_T
Neonatal
Neo Sen
Adult
Adult Sen
TGF
UNSTIM
N
A
N
NS
A
AS
UNSTIM
TGF

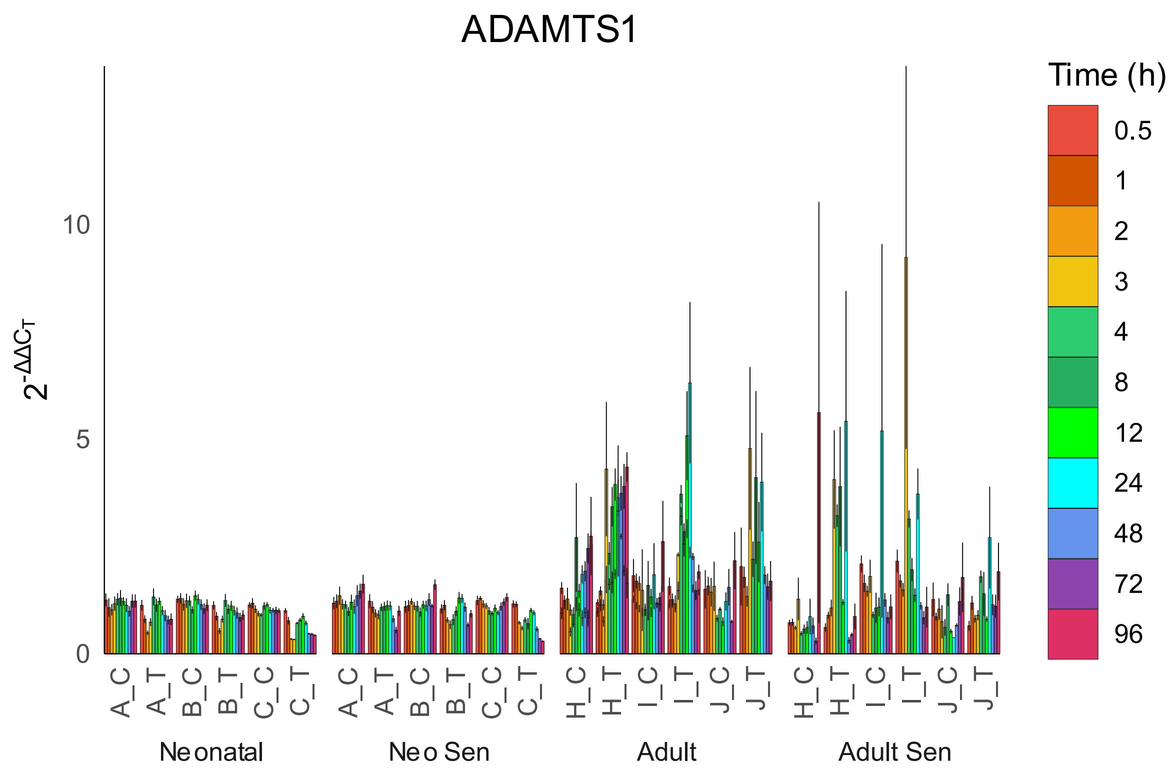
ADAMTS1
Time (h)
0.5
1
2
3
4
8
12
24
48
72
96
$2^{-\Delta\Delta C_T}$
10
5
0
A_C
A_T
B_C
B_T
C_C
C_T
Neonatal
Neo Sen
H_C
H_T
I_C
I_T
J_C
J_T
Adult
Adult Sen

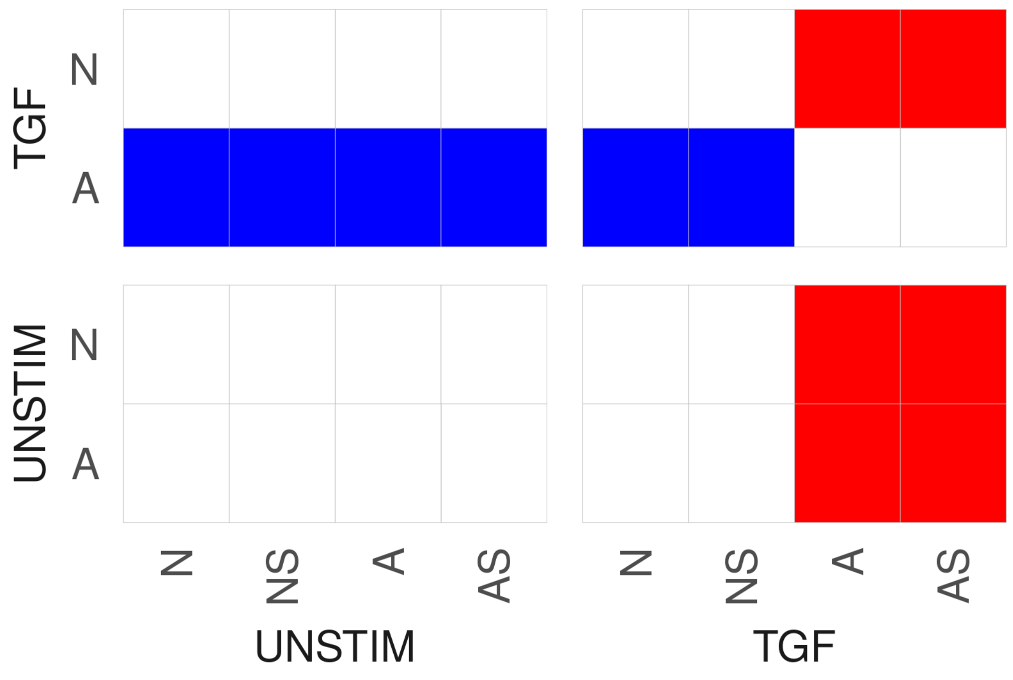
TGF
UNSTIM
N
A
N
NS
A
AS
UNSTIM
TGF

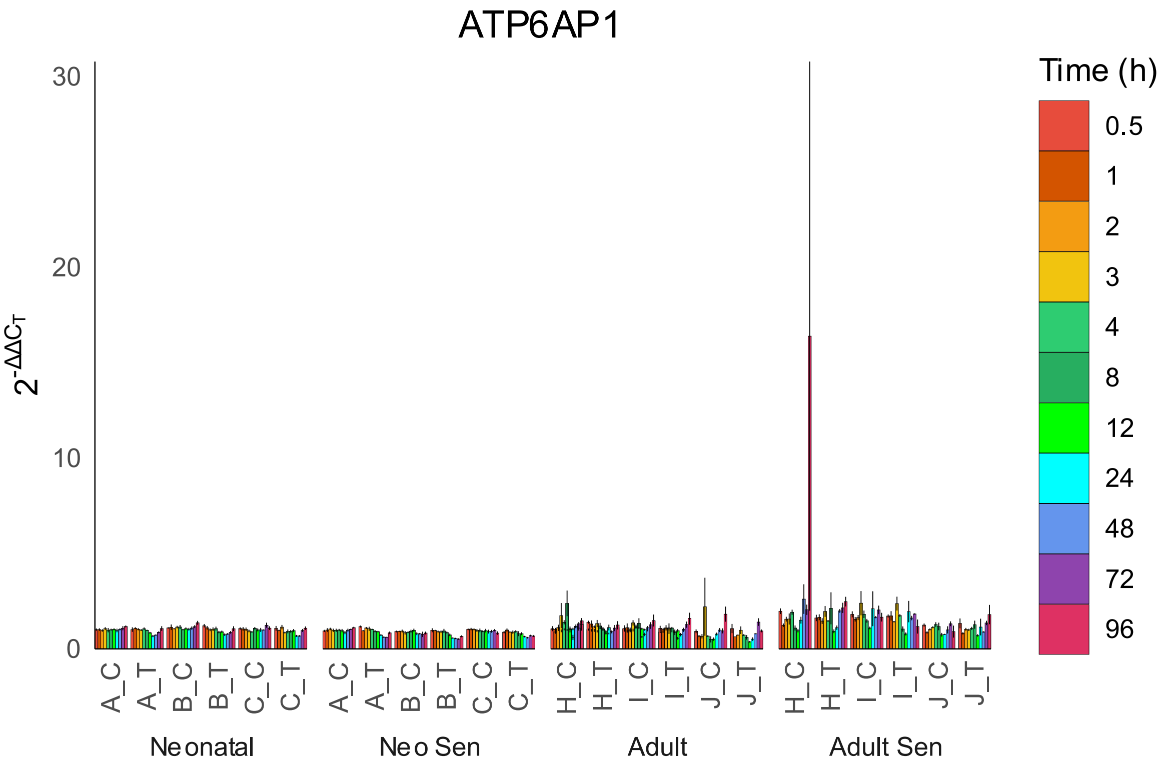

ATP6AP1
$2^{-\Delta\Delta C_T}$
30
20
10
0
Time (h)
0.5
1
2
3
4
8
12
24
48
72
96
A_C
A_T
B_C
B_T
C_C
C_T
H_C
H_T
I_C
I_T
J_C
J_T
Neonatal
Neo Sen
Adult
Adult Sen


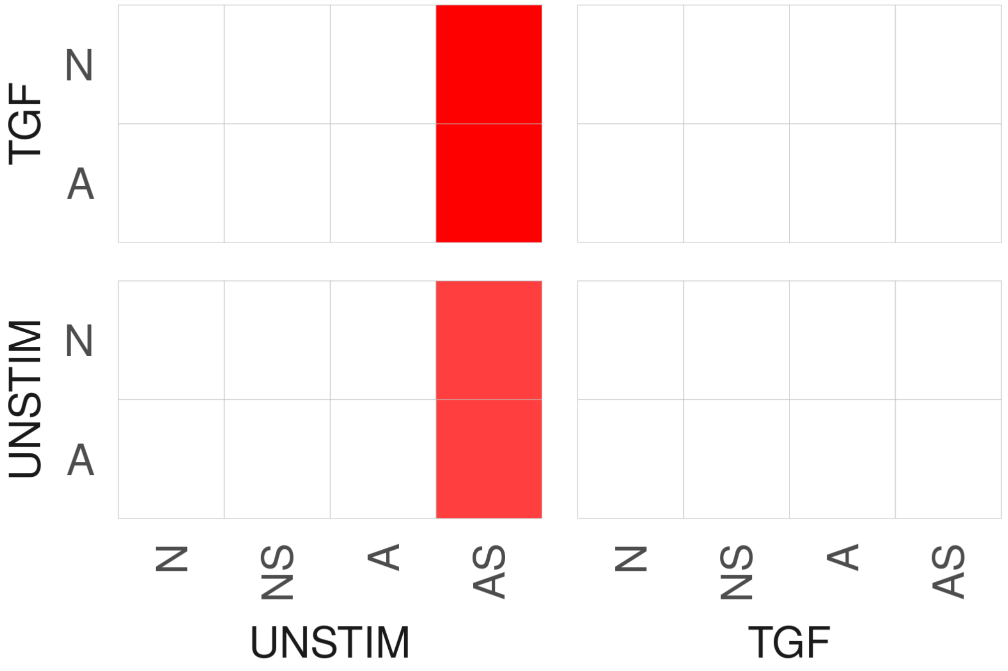

TGF
UNSTIM
N
A
N
NS
A
AS
UNSTIM
TGF

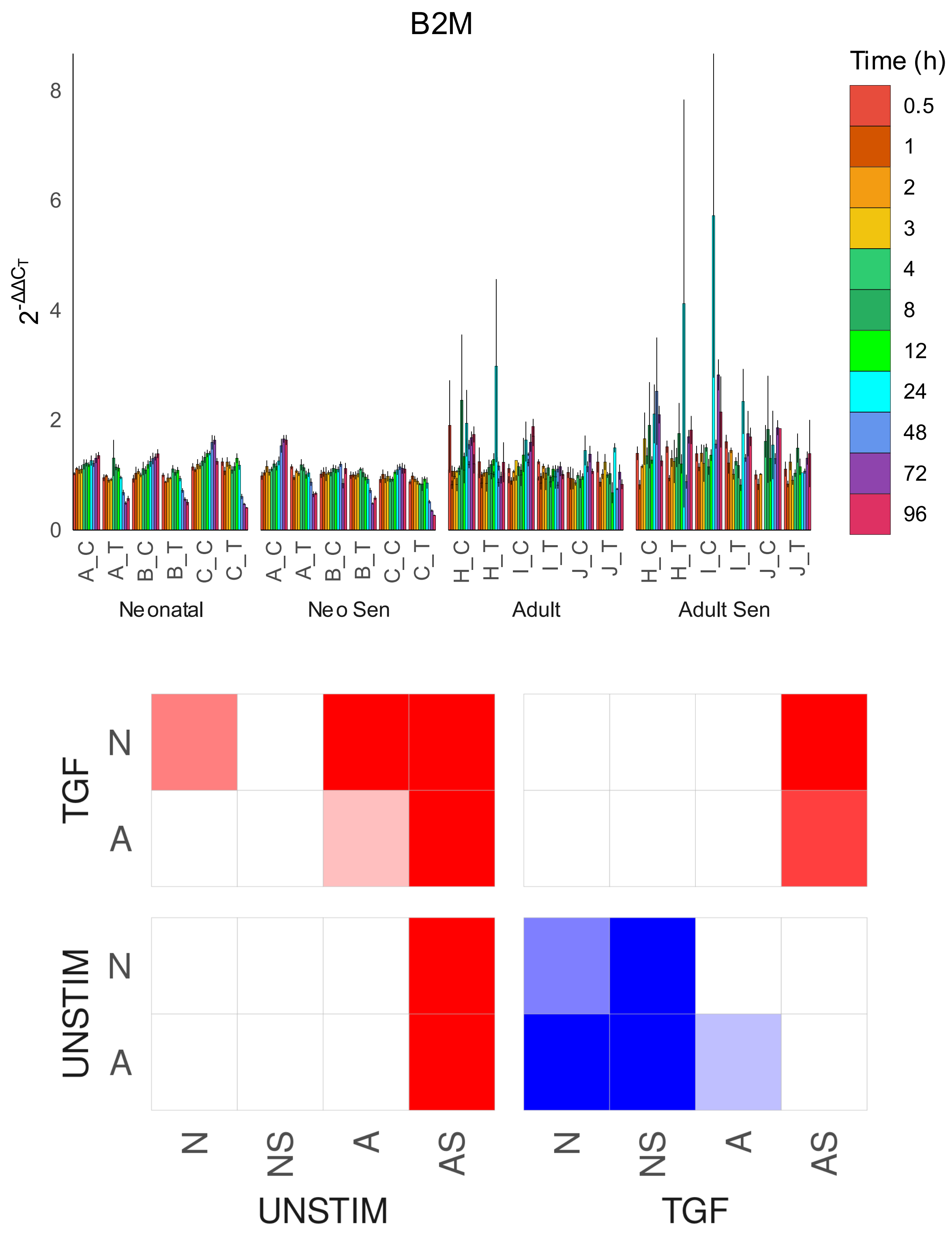

B2M
Time (h)
0.5
1
2
3
4
8
12
24
48
72
96
$2^{-\Delta\Delta C_T}$
0
2
4
6
8
A_C
A_T
B_C
B_T
C_C
C_T
H_C
H_T
I_C
I_T
J_C
J_T
Neonatal
Neo Sen
Adult
Adult Sen
TGF
UNSTIM
N
A
N
NS
A
AS
UNSTIM
TGF

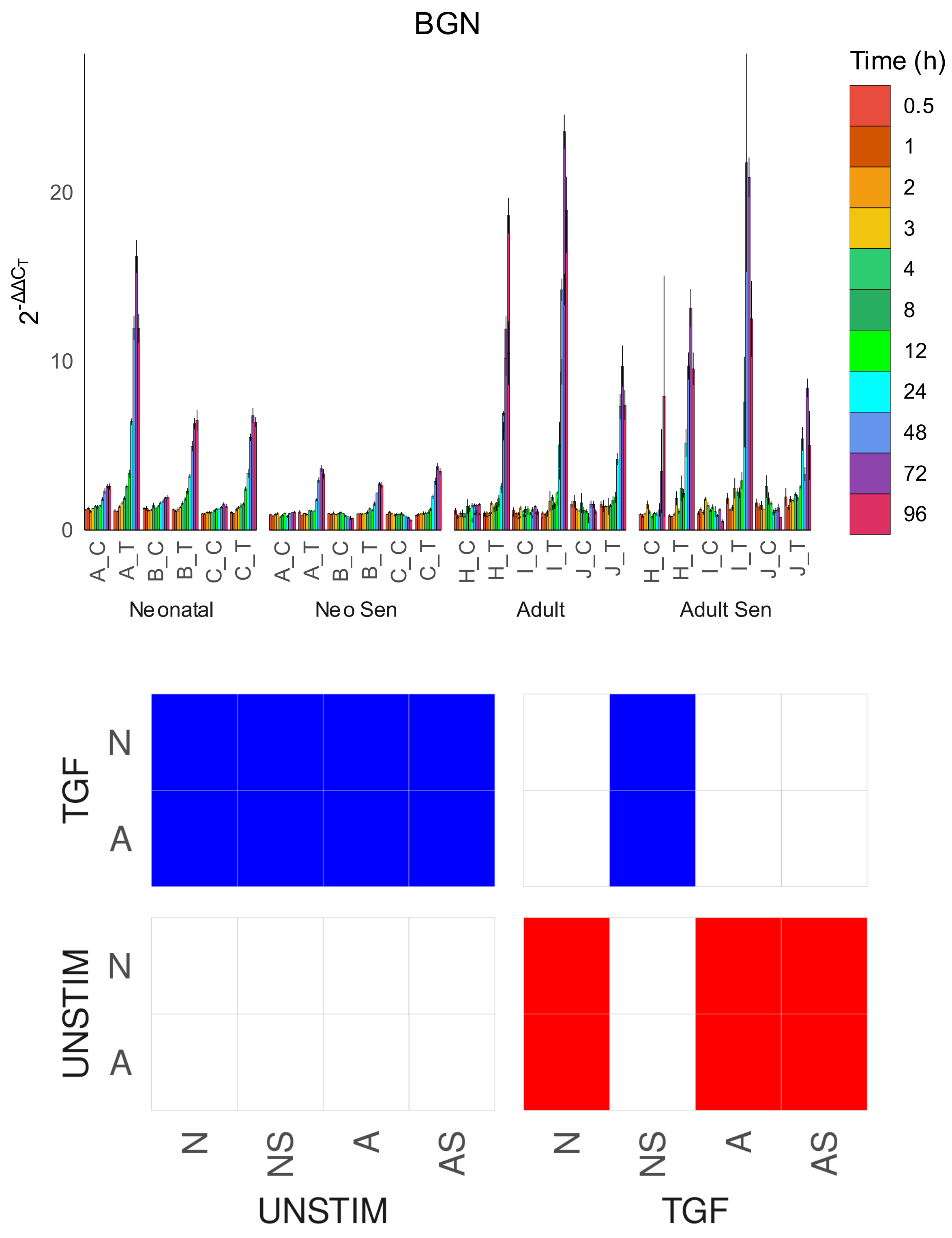

BGN
Time (h)
0.5
1
2
3
4
8
12
24
48
72
96
$2^{-\Delta\Delta C_T}$
20
10
0
A_C
A_T
B_C
B_T
C_C
C_T
H_C
H_T
I_C
I_T
J_C
J_T
Neonatal
Neo Sen
Adult
Adult Sen
TGF
UNSTIM
N
A
N
NS
A
AS
UNSTIM
TGF

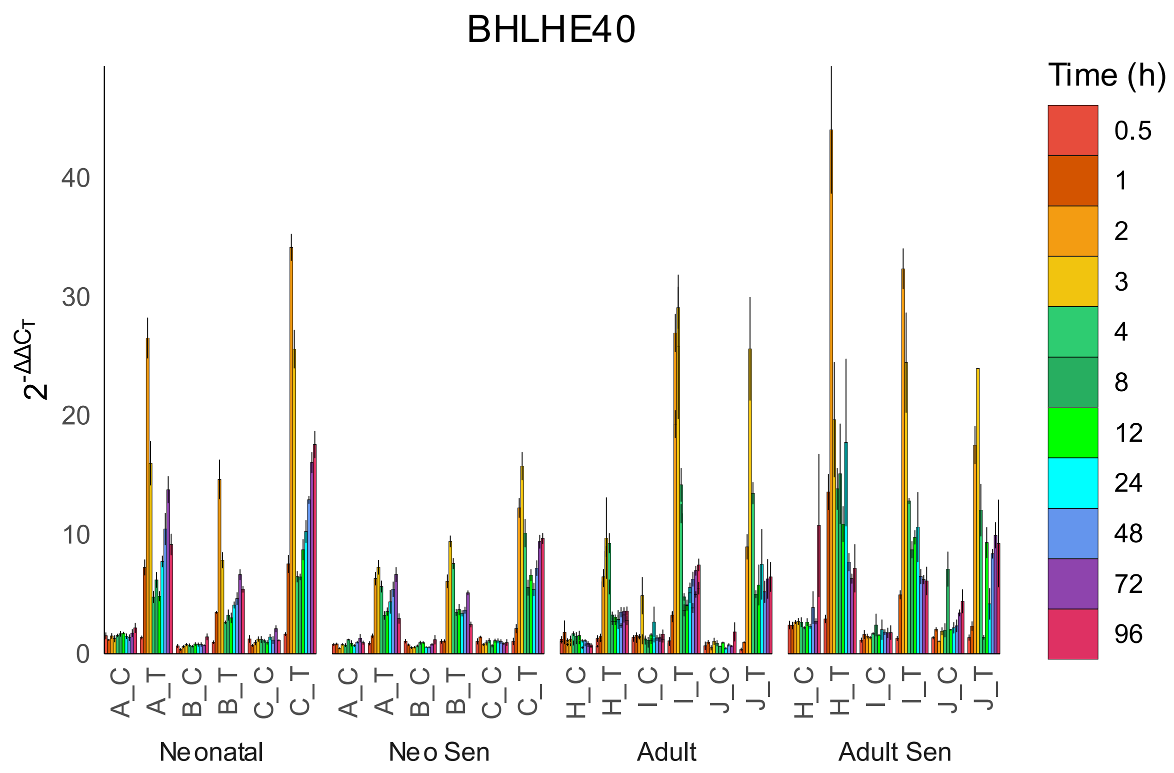

BHLHE40
Time (h)
0.5
1
2
3
4
8
12
24
48
72
96
$2^{-\Delta\Delta C_T}$
0
10
20
30
40
A_C
A_T
B_C
B_T
C_C
C_T
H_C
H_T
I_C
I_T
J_C
J_T
Neonatal
Neo Sen
Adult
Adult Sen


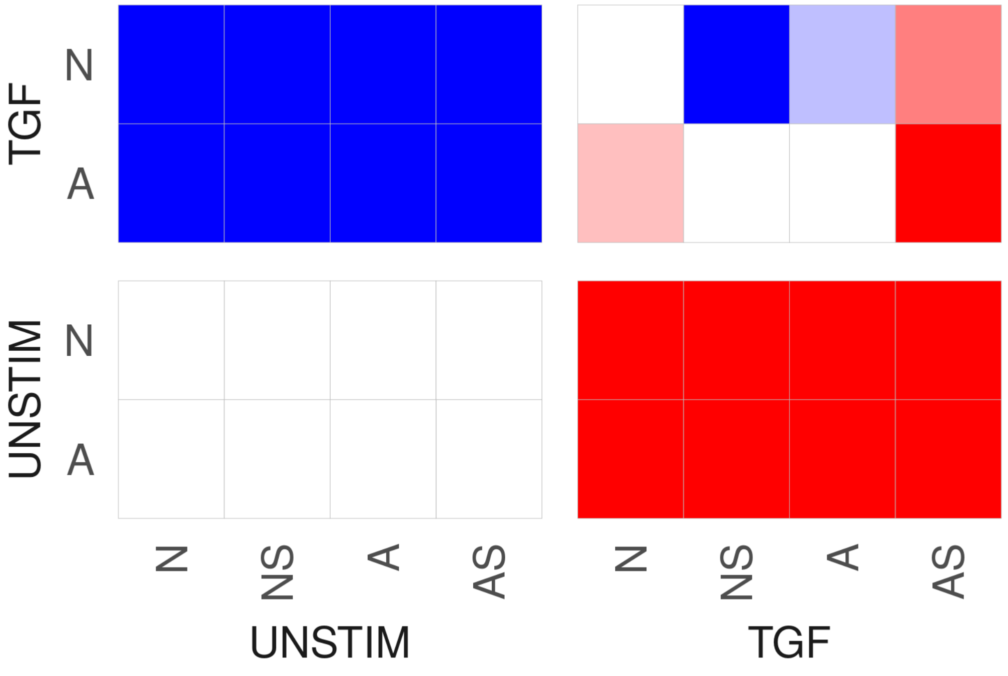

TGF
UNSTIM
N
A
N
NS
A
AS
UNSTIM
TGF

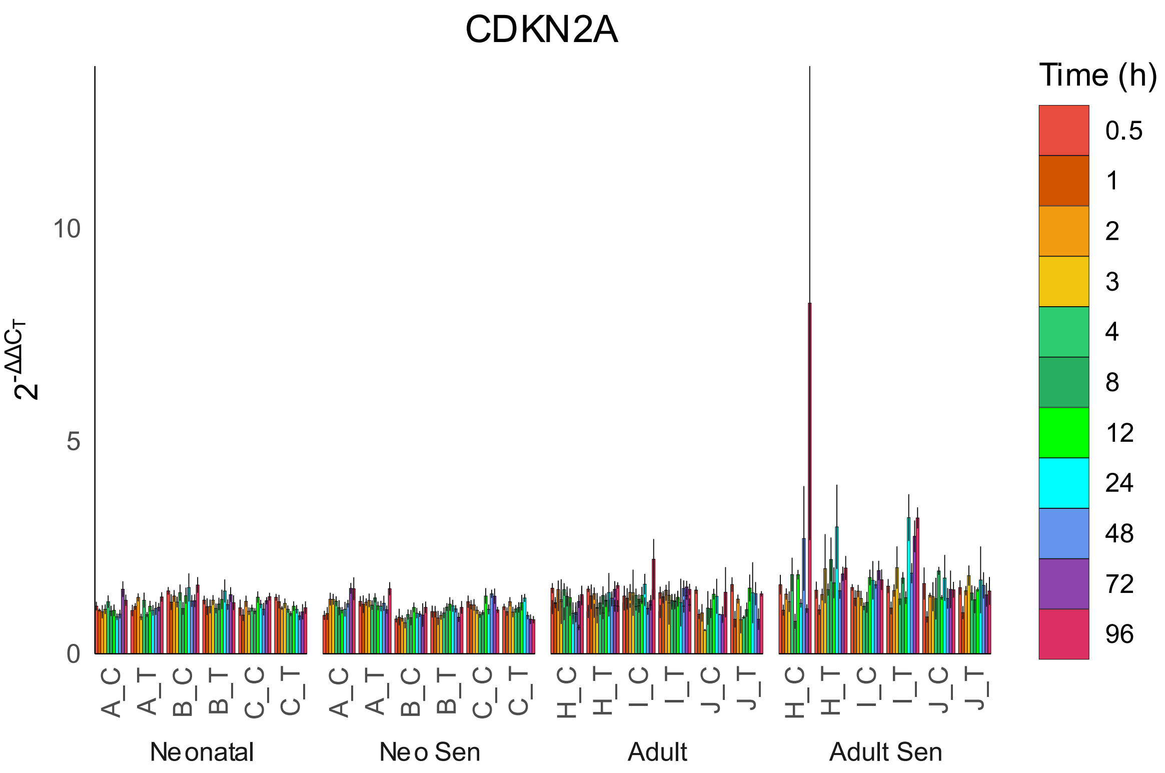

CDKN2A
Time (h)
0.5
1
2
3
4
8
12
24
48
72
96
$2^{-\Delta\Delta C_T}$
10
5
0
A_C
A_T
B_C
B_T
C_C
C_T
H_C
H_T
I_C
I_T
J_C
J_T
Neonatal
Neo Sen
Adult
Adult Sen


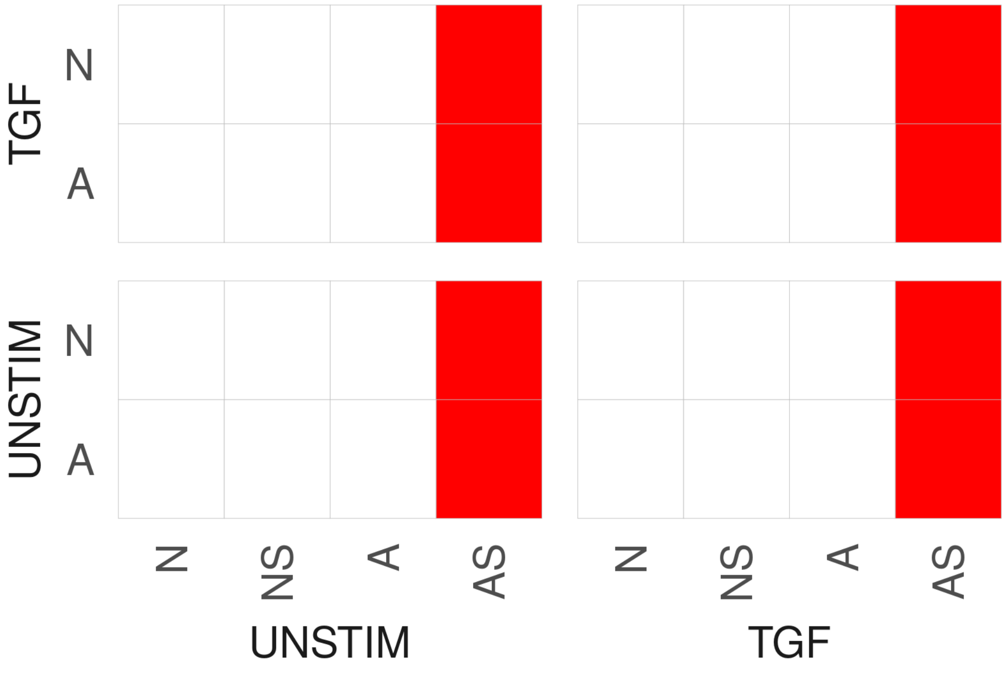

TGF
UNSTIM
N
A
N
NS
A
AS
UNSTIM
TGF

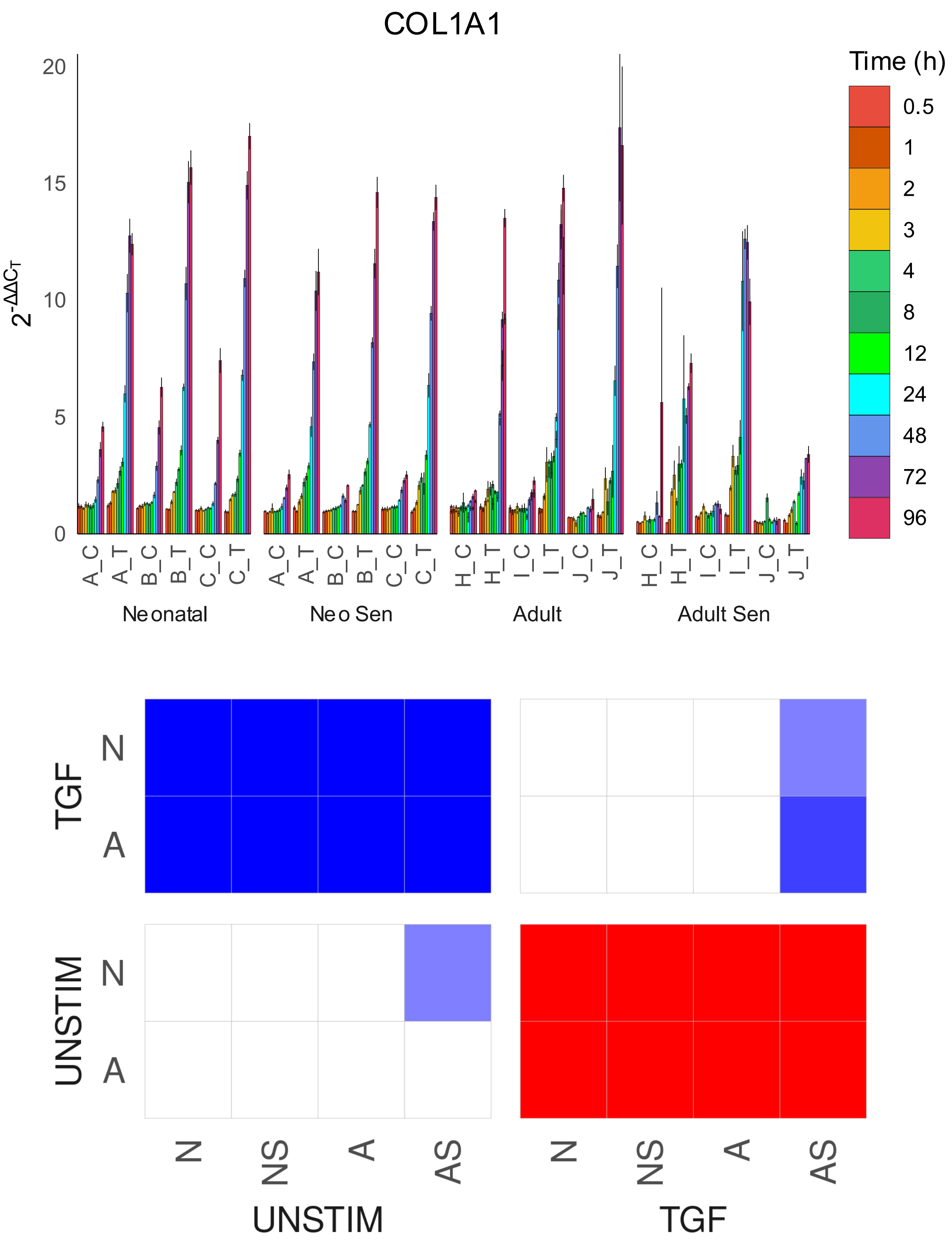

COL1A1
Time (h)
0.5
1
2
3
4
8
12
24
48
72
96
$2^{-\Delta\Delta C_T}$
0
5
10
15
20
A_C
A_T
B_C
B_T
C_C
C_T
H_C
H_T
I_C
I_T
J_C
J_T
Neonatal
Neo Sen
Adult
Adult Sen
TGF
UNSTIM
N
A
N
NS
A
AS
UNSTIM
TGF

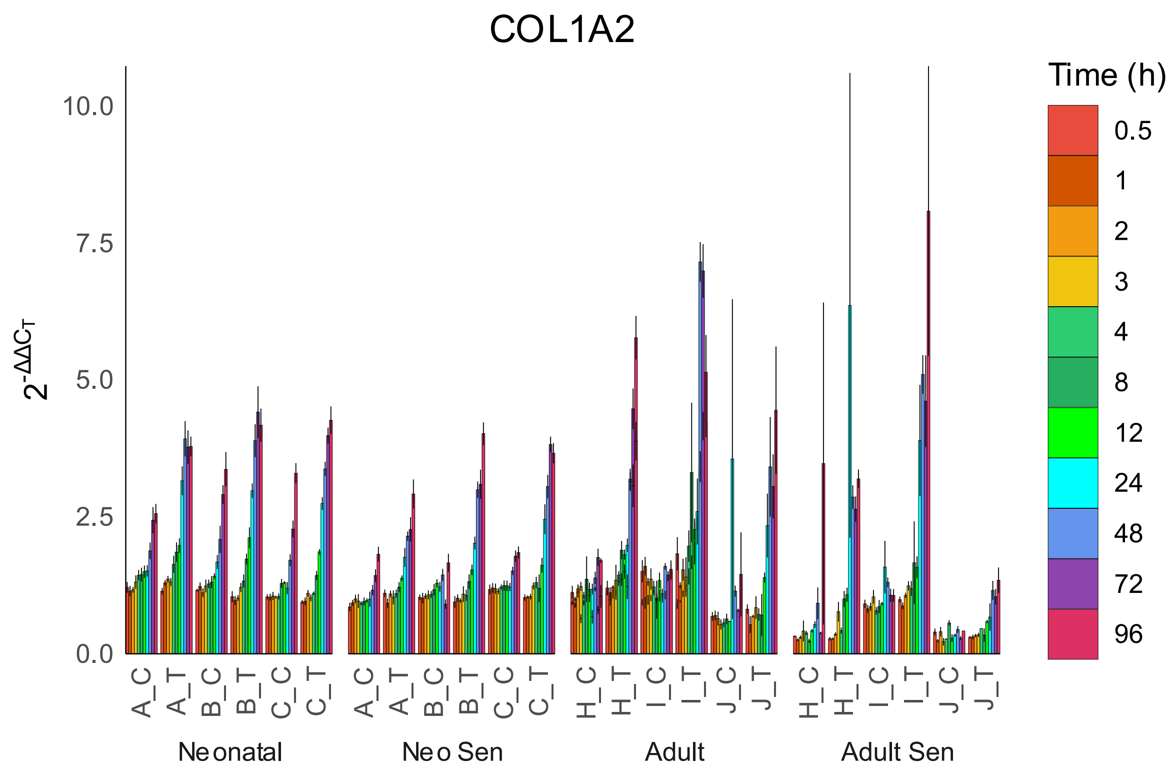
COL1A2
Time (h)
0.5
1
2
3
4
8
12
24
48
72
96
10.0
7.5
5.0
2.5
0.0
$2^{-\Delta\Delta C_T}$
A_C
A_T
B_C
B_T
C_C
C_T
H_C
H_T
I_C
I_T
J_C
J_T
Neonatal
Neo Sen
Adult
Adult Sen

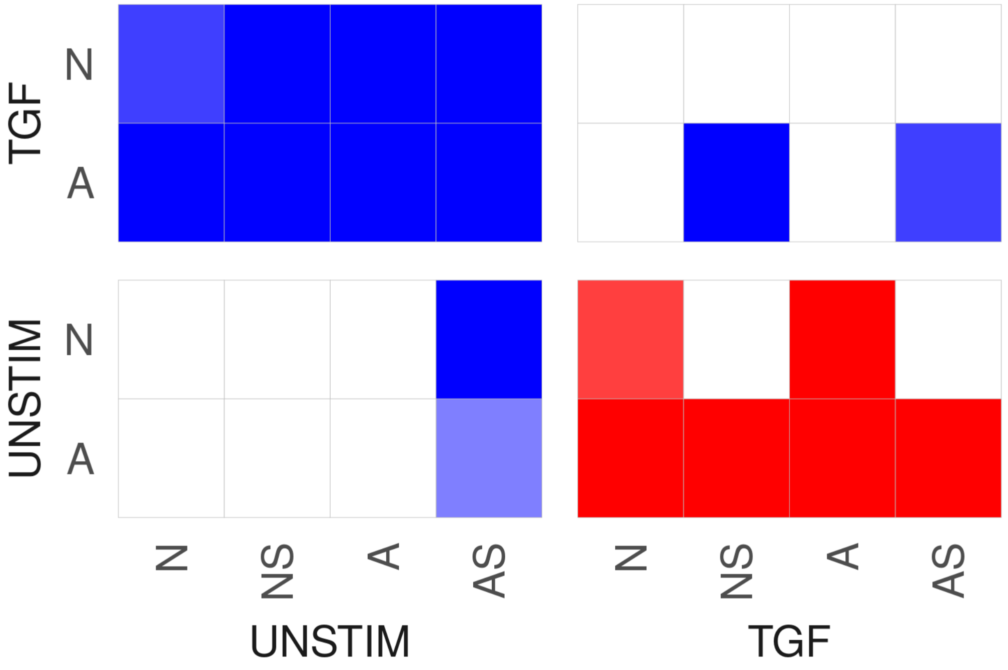
TGF
UNSTIM
N
A
N
NS
A
AS
UNSTIM
TGF

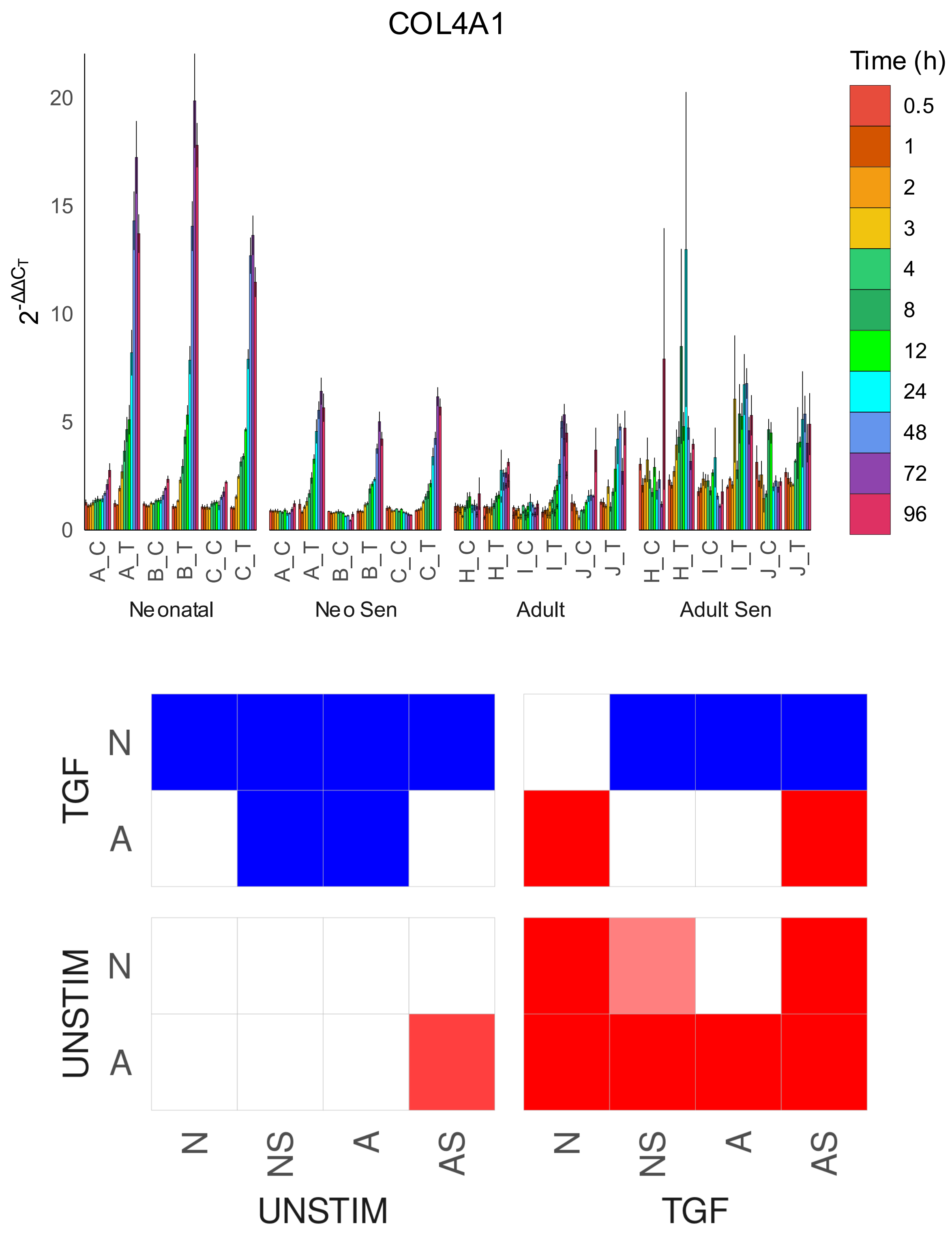

COL4A1
Time (h)
0.5
1
2
3
4
8
12
24
48
72
96
20
15
10
5
0
A_C
A_T
B_C
B_T
C_C
C_T
H_C
H_T
I_C
I_T
J_C
J_T
Neonatal
Neo Sen
Adult
Adult Sen
TGF
UNSTIM
N
A
N
NS
A
AS
UNSTIM
TGF

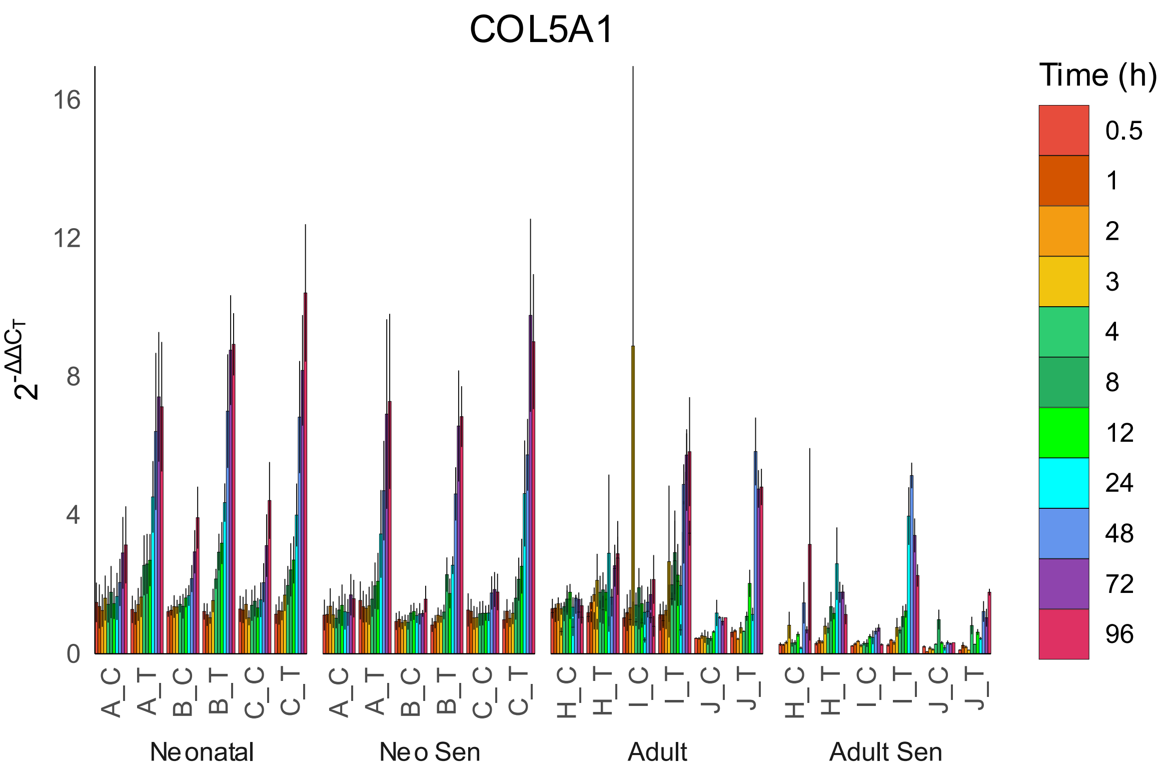

COL5A1
Time (h)
0.5
1
2
3
4
8
12
24
48
72
96
$2^{-\Delta\Delta C_T}$
16
12
8
4
0
A_C
A_T
B_C
B_T
C_C
C_T
H_C
H_T
I_C
I_T
J_C
J_T
Neonatal
Neo Sen
Adult
Adult Sen


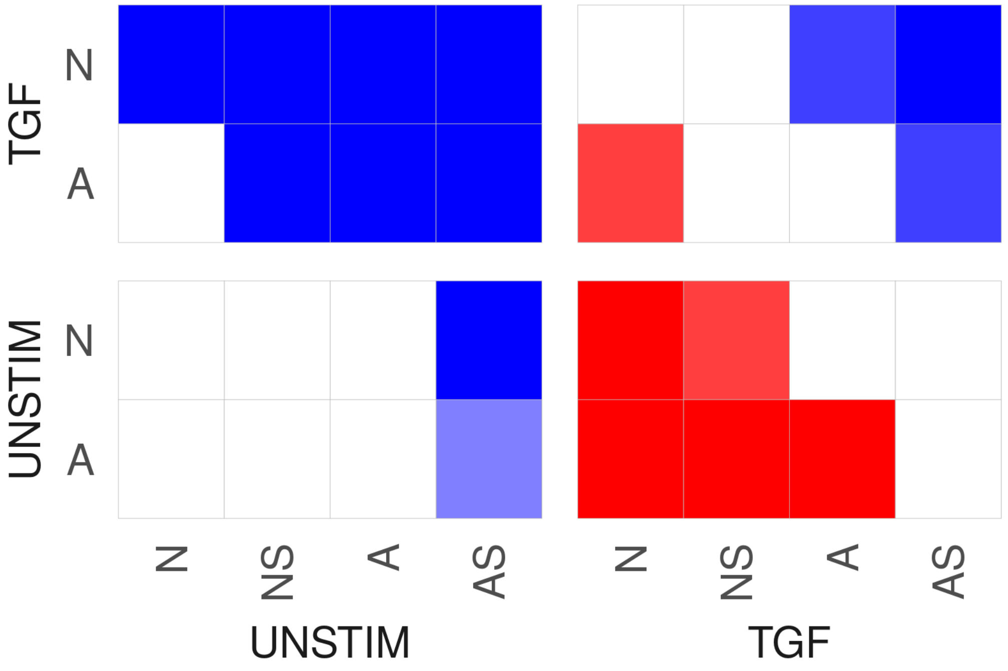

TGF
N
A
UNSTIM
N
A
N
NS
A
AS
UNSTIM
N
NS
A
AS
TGF

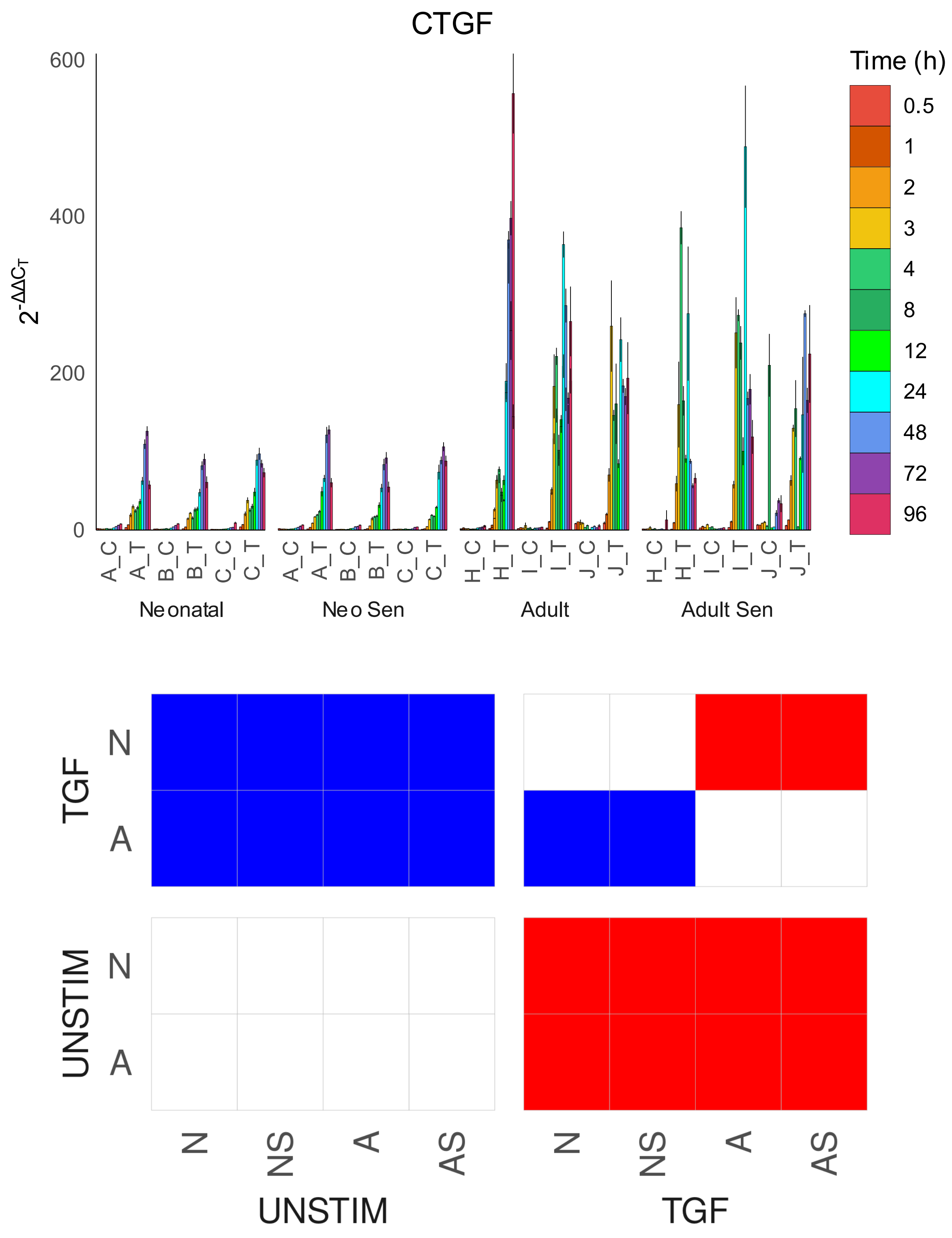

CTGF
600
400
200
0
2^-ΔΔCT
Time (h)
0.5
1
2
3
4
8
12
24
48
72
96
A_C A_T B_C B_T C_C C_T
Neonatal
A_C A_T B_C B_T C_C C_T
Neo Sen
H_C H_T I_C I_T J_C J_T
Adult
H_C H_T I_C I_T J_C J_T
Adult Sen
TGF
N
A
UNSTIM
N
A
N NS A AS
UNSTIM
N NS A AS
TGF

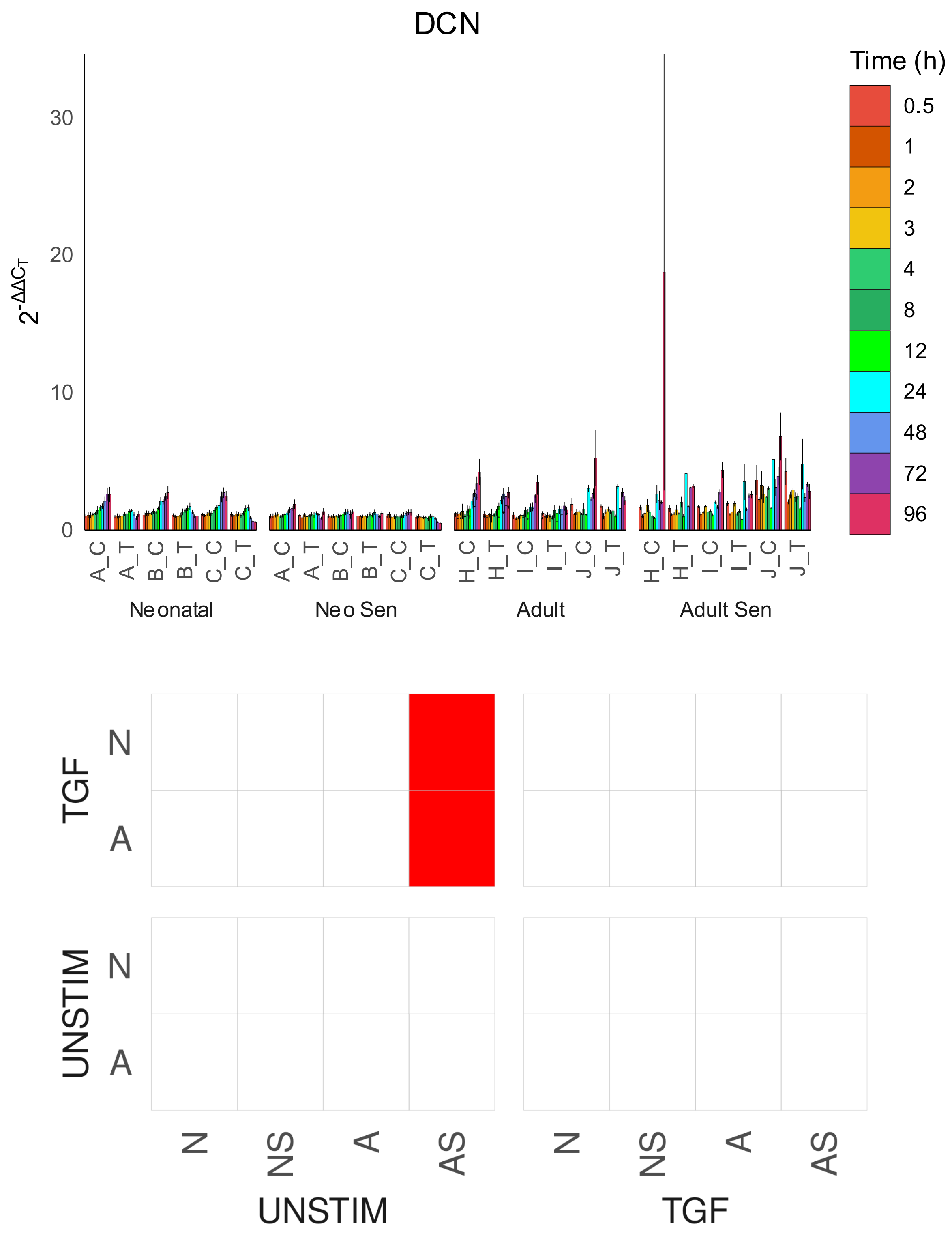
DCN
Time (h)
0.5
1
2
3
4
8
12
24
48
72
96
$2^{-\Delta\Delta C_T}$
0
10
20
30
A_C
A_T
B_C
B_T
C_C
C_T
H_C
H_T
I_C
I_T
J_C
J_T
Neonatal
Neo Sen
Adult
Adult Sen
TGF
UNSTIM
N
A
N
NS
A
AS
UNSTIM
TGF

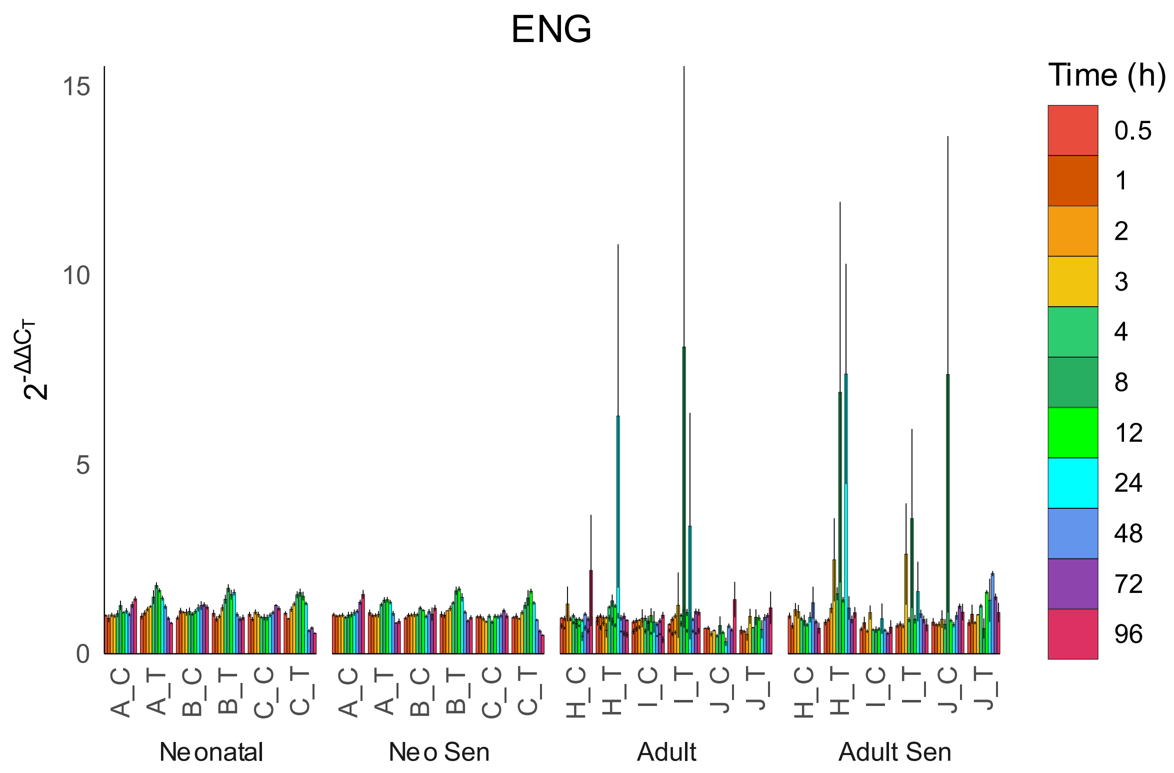

ENG
$2^{-\Delta\Delta C_T}$
15
10
5
0
A_C
A_T
B_C
B_T
C_C
C_T
A_C
A_T
B_C
B_T
C_C
C_T
H_C
H_T
I_C
I_T
J_C
J_T
H_C
H_T
I_C
I_T
J_C
J_T
Neonatal
Neo Sen
Adult
Adult Sen
Time (h)
0.5
1
2
3
4
8
12
24
48
72
96


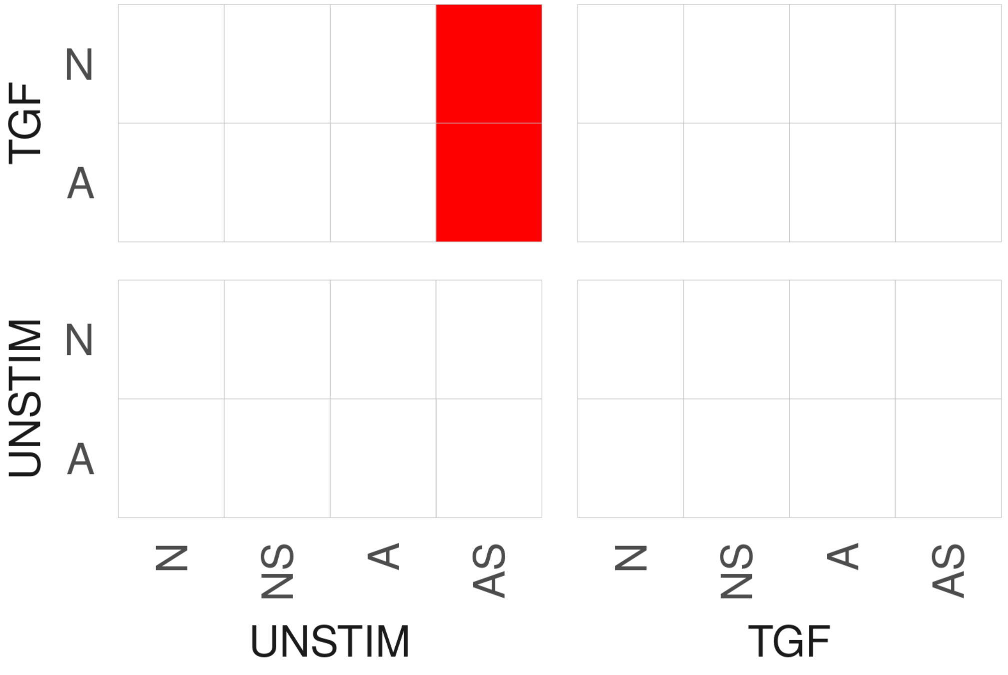

TGF
N
A
UNSTIM
N
A
N
NS
A
AS
N
NS
A
AS
UNSTIM
TGF

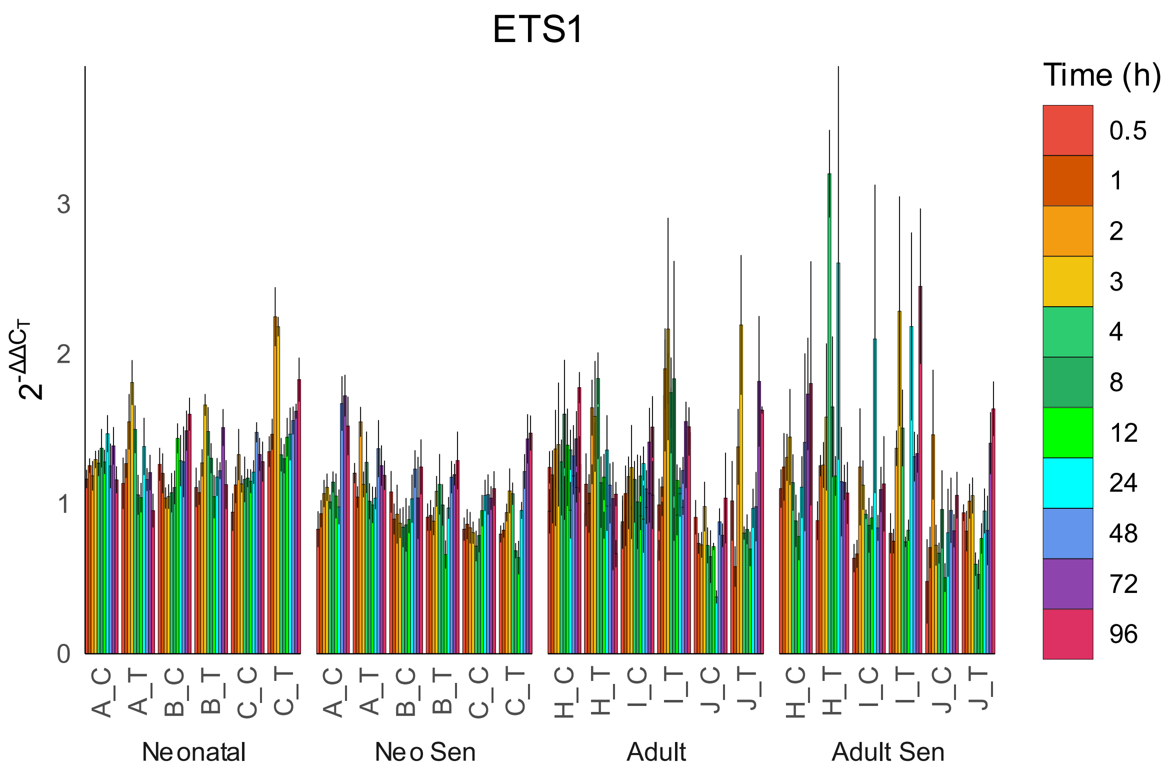
ETS1
Time (h)
0.5
1
2
3
4
8
12
24
48
72
96
$2^{-\Delta\Delta C_T}$
0
1
2
3
A_C
A_T
B_C
B_T
C_C
C_T
H_C
H_T
I_C
I_T
J_C
J_T
Neonatal
Neo Sen
Adult
Adult Sen

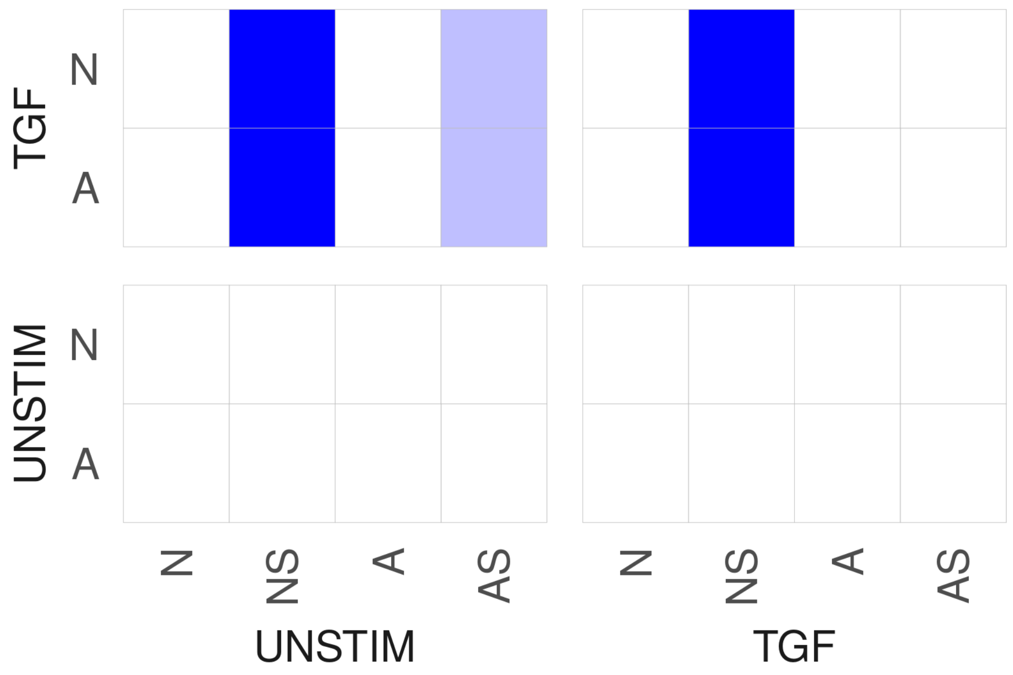
TGF
N
A
UNSTIM
N
A
N
NS
A
AS
UNSTIM
N
NS
A
AS
TGF

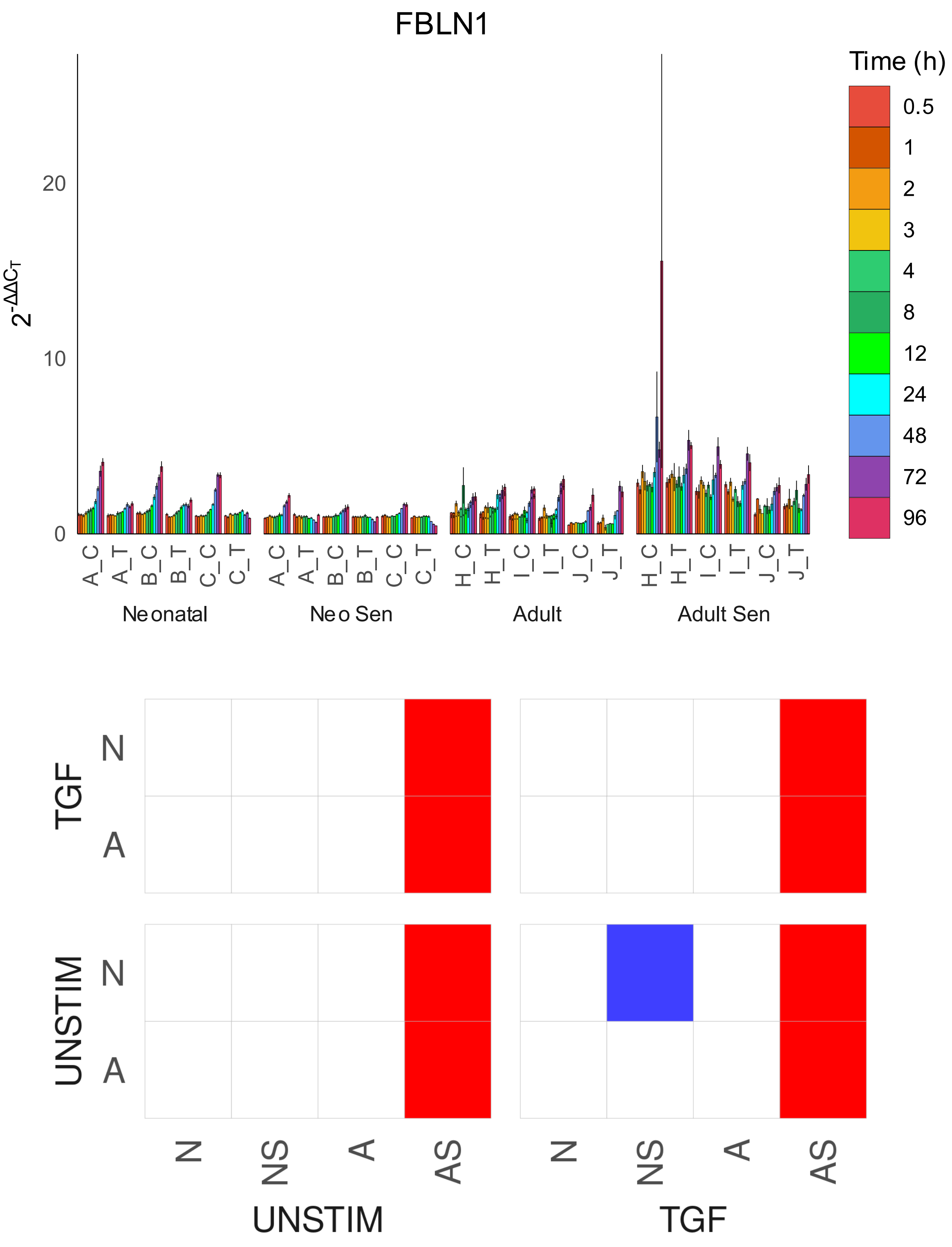

FBLN1
Time (h)
0.5
1
2
3
4
8
12
24
48
72
96
$2^{-\Delta\Delta C_T}$
20
10
0
A_C
A_T
B_C
B_T
C_C
C_T
H_C
H_T
I_C
I_T
J_C
J_T
Neonatal
Neo Sen
Adult
Adult Sen
TGF
UNSTIM
N
A
N
NS
A
AS
UNSTIM
TGF

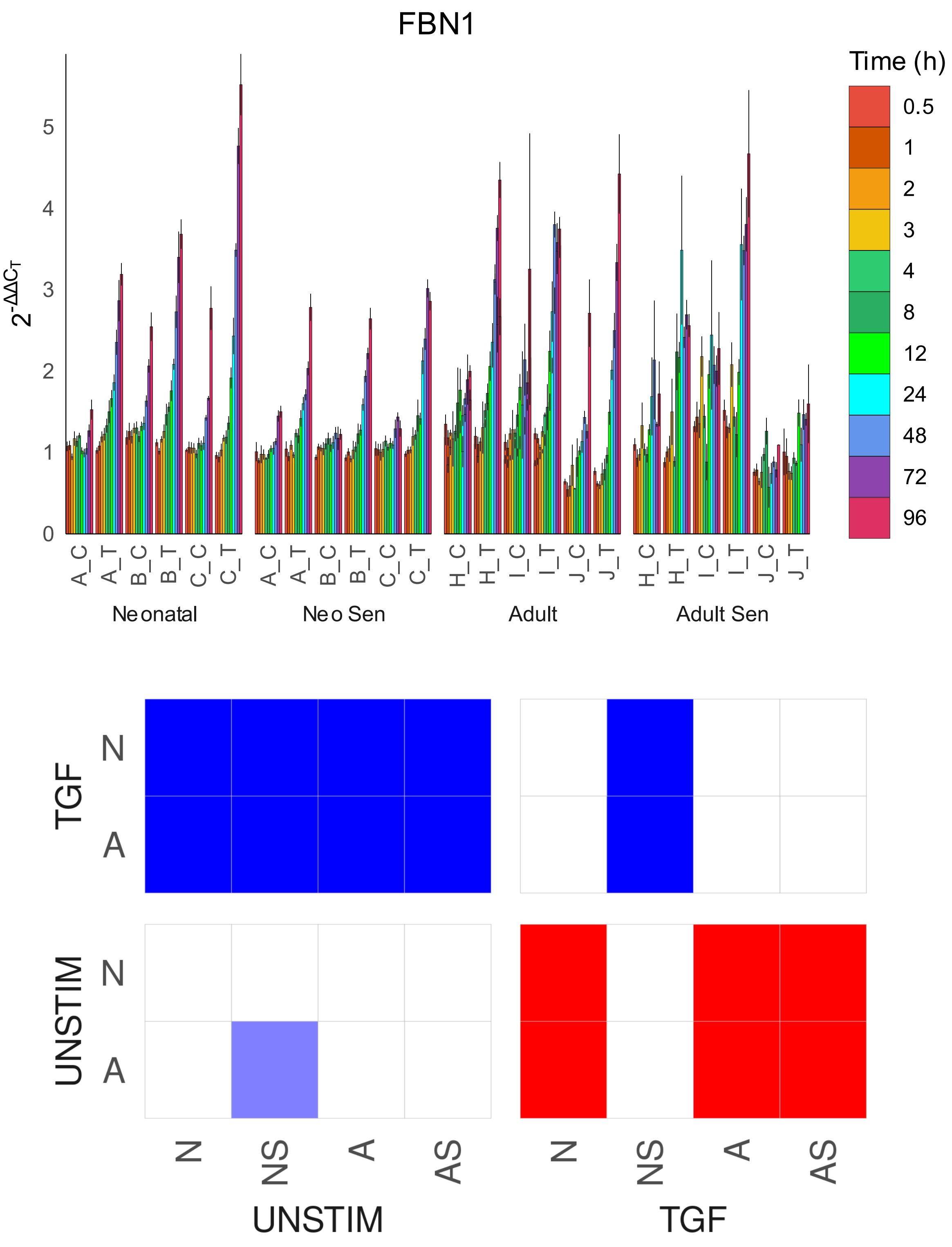
FBN1
Time (h)
0.5
1
2
3
4
8
12
24
48
72
96
$2^{-\Delta\Delta C_T}$
0
1
2
3
4
5
A_C
A_T
B_C
B_T
C_C
C_T
H_C
H_T
I_C
I_T
J_C
J_T
Neonatal
Neo Sen
Adult
Adult Sen
TGF
UNSTIM
N
A
N
NS
A
AS
UNSTIM
TGF

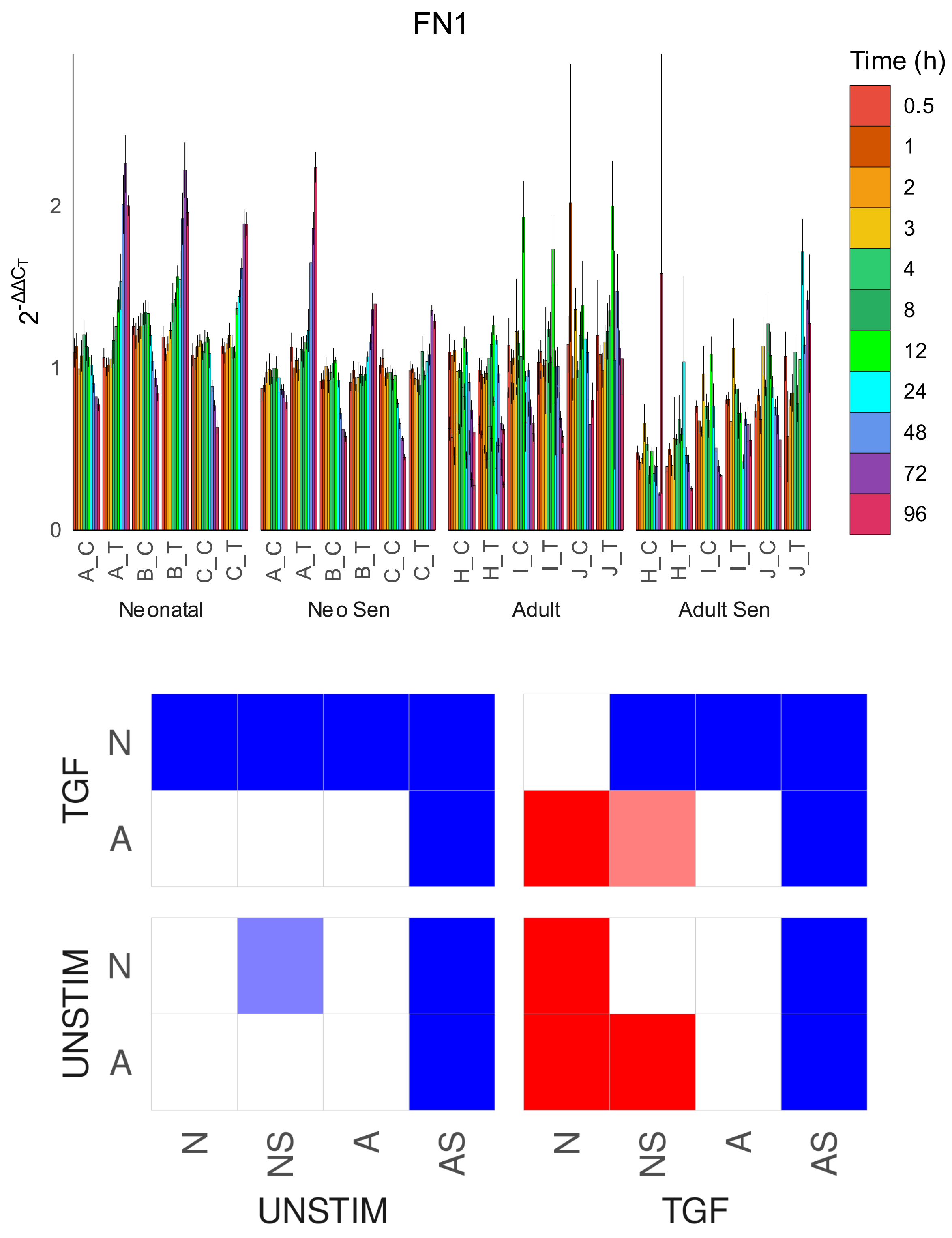
FN1
Time (h)
0.5
1
2
3
4
8
12
24
48
72
96
Neonatal
Neo Sen
Adult
Adult Sen
A_C
A_T
B_C
B_T
C_C
C_T
H_C
H_T
I_C
I_T
J_C
J_T
TGF
UNSTIM
N
A
NS
AS

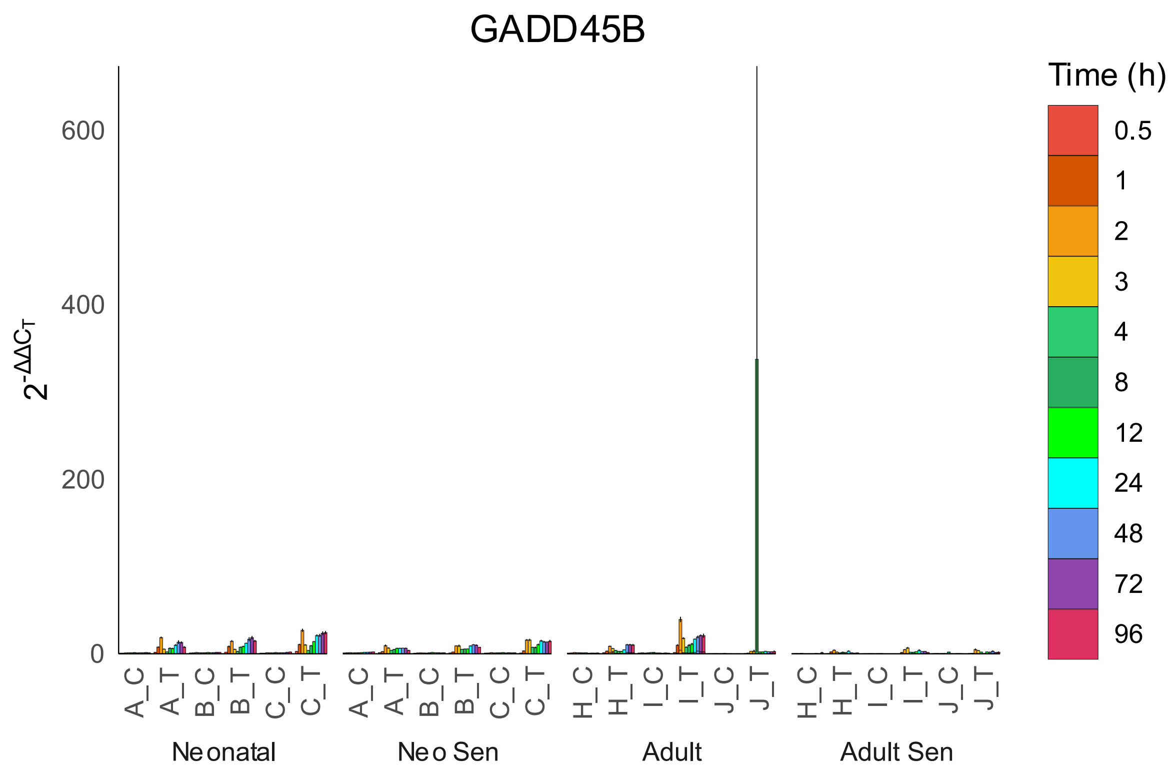
GADD45B
Time (h)
0.5
1
2
3
4
8
12
24
48
72
96
600
400
200
0
A_C
A_T
B_C
B_T
C_C
C_T
H_C
H_T
I_C
I_T
J_C
J_T
Neonatal
Neo Sen
Adult
Adult Sen

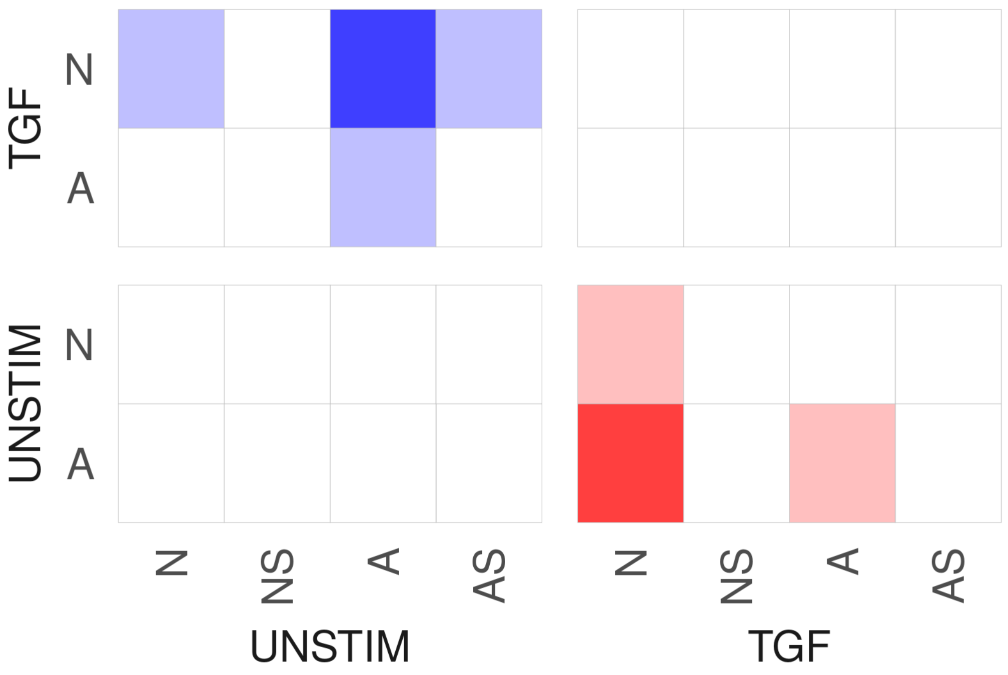
TGF
UNSTIM
N
A
N
NS
A
AS
UNSTIM
TGF

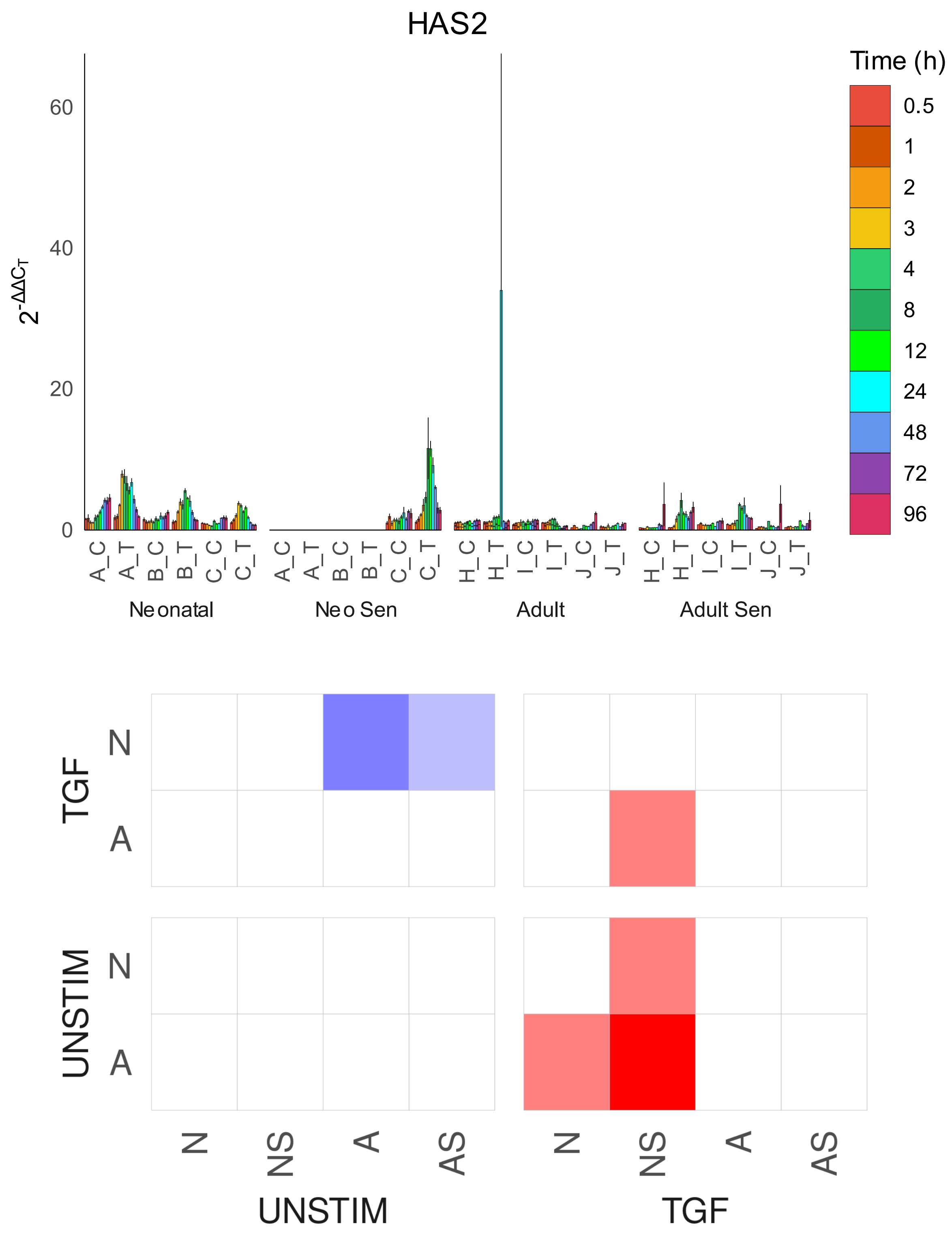

HAS2
Time (h)
0.5
1
2
3
4
8
12
24
48
72
96
$2^{-\Delta\Delta C_T}$
0
20
40
60
A_C A_T B_C B_T C_C C_T
Neonatal
A_C A_T B_C B_T C_C C_T
Neo Sen
H_C H_T I_C I_T J_C J_T
Adult
H_C H_T I_C I_T J_C J_T
Adult Sen
TGF
N
A
UNSTIM
N
A
N NS A AS
UNSTIM
N NS A AS
TGF

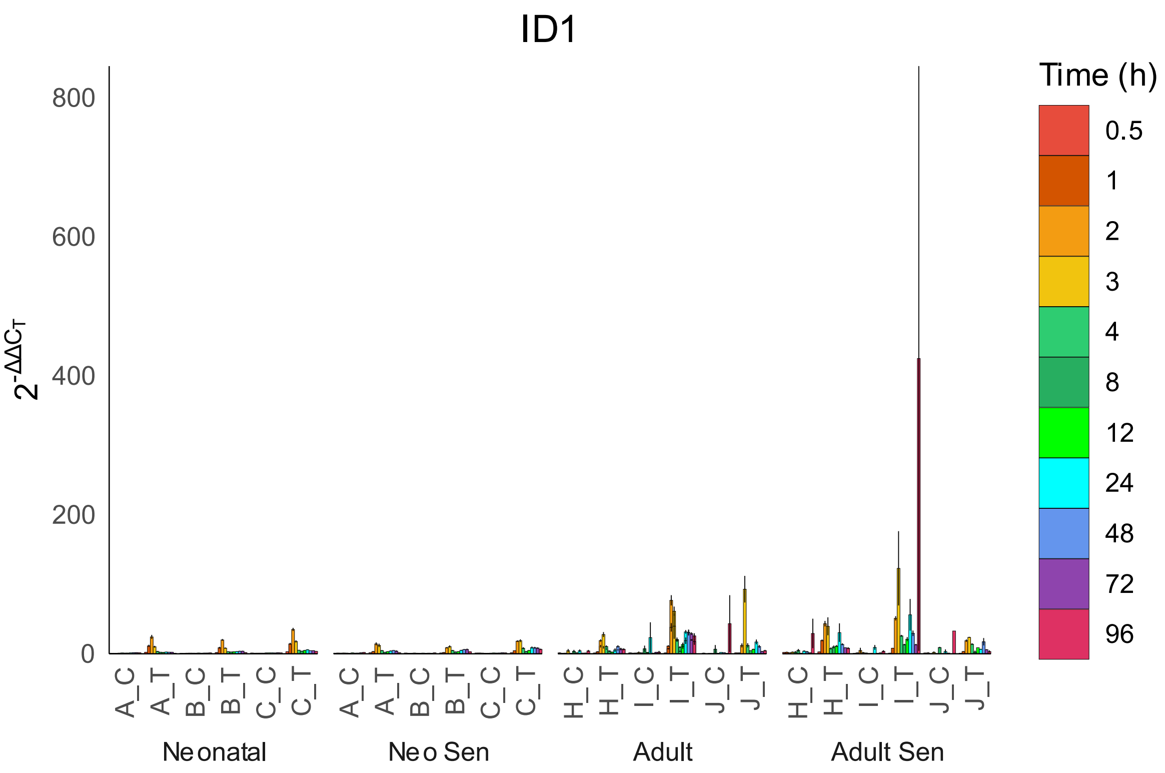

ID1
Time (h)
0.5
1
2
3
4
8
12
24
48
72
96
800
600
400
200
0
$2^{-\Delta\Delta C_T}$
A_C
A_T
B_C
B_T
C_C
C_T
H_C
H_T
I_C
I_T
J_C
J_T
Neonatal
Neo Sen
Adult
Adult Sen


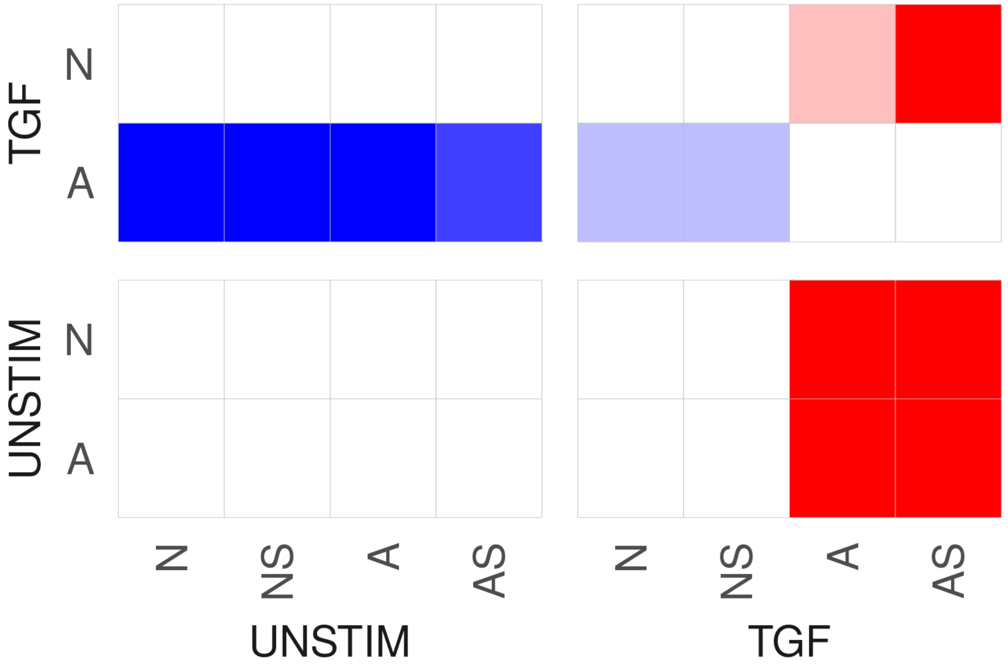

TGF
UNSTIM
N
A
N
NS
A
AS
UNSTIM
TGF

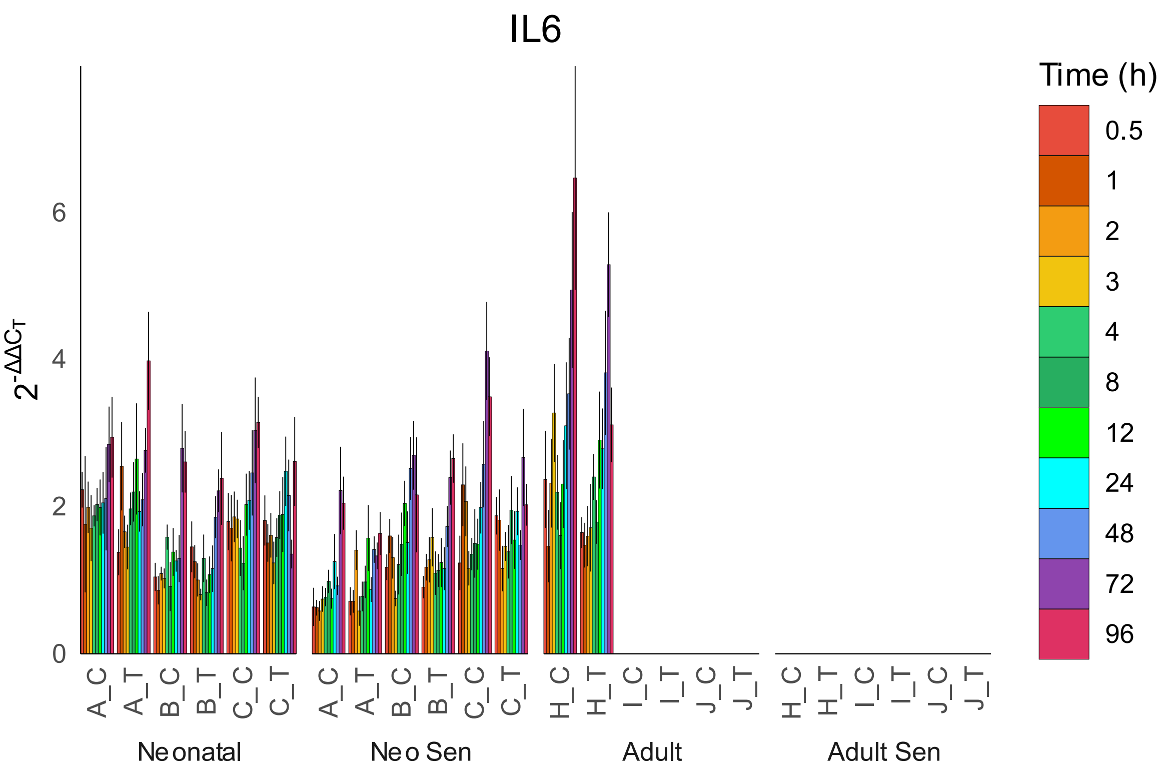

IL6
Time (h)
0.5
1
2
3
4
8
12
24
48
72
96
0
2
4
6
A_C
A_T
B_C
B_T
C_C
C_T
H_C
H_T
I_C
I_T
J_C
J_T
Neonatal
Neo Sen
Adult
Adult Sen


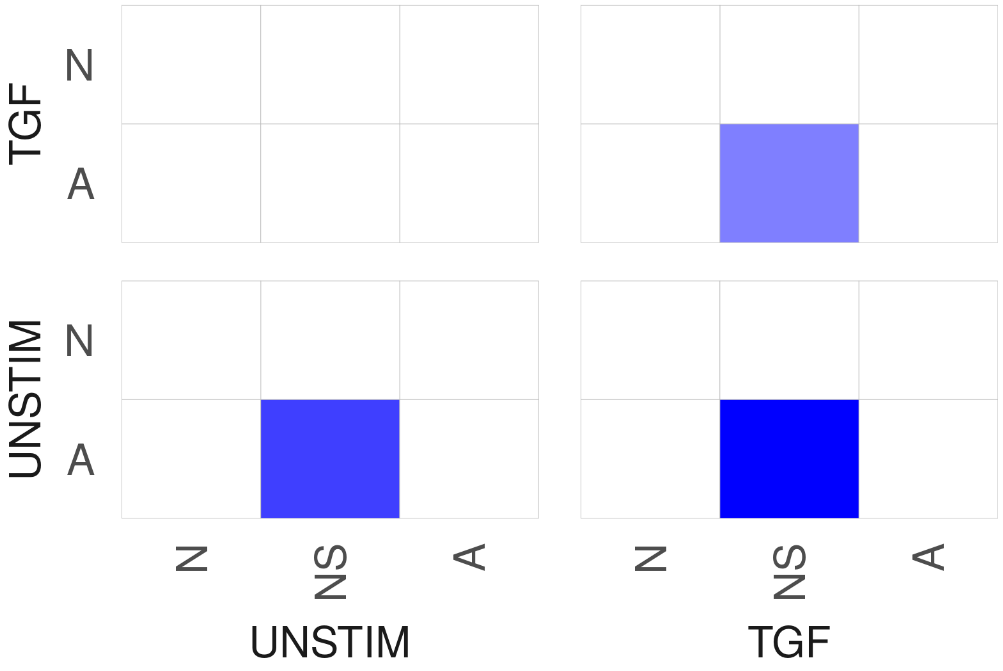

TGF
UNSTIM
N
A
N
NS
A
UNSTIM
TGF

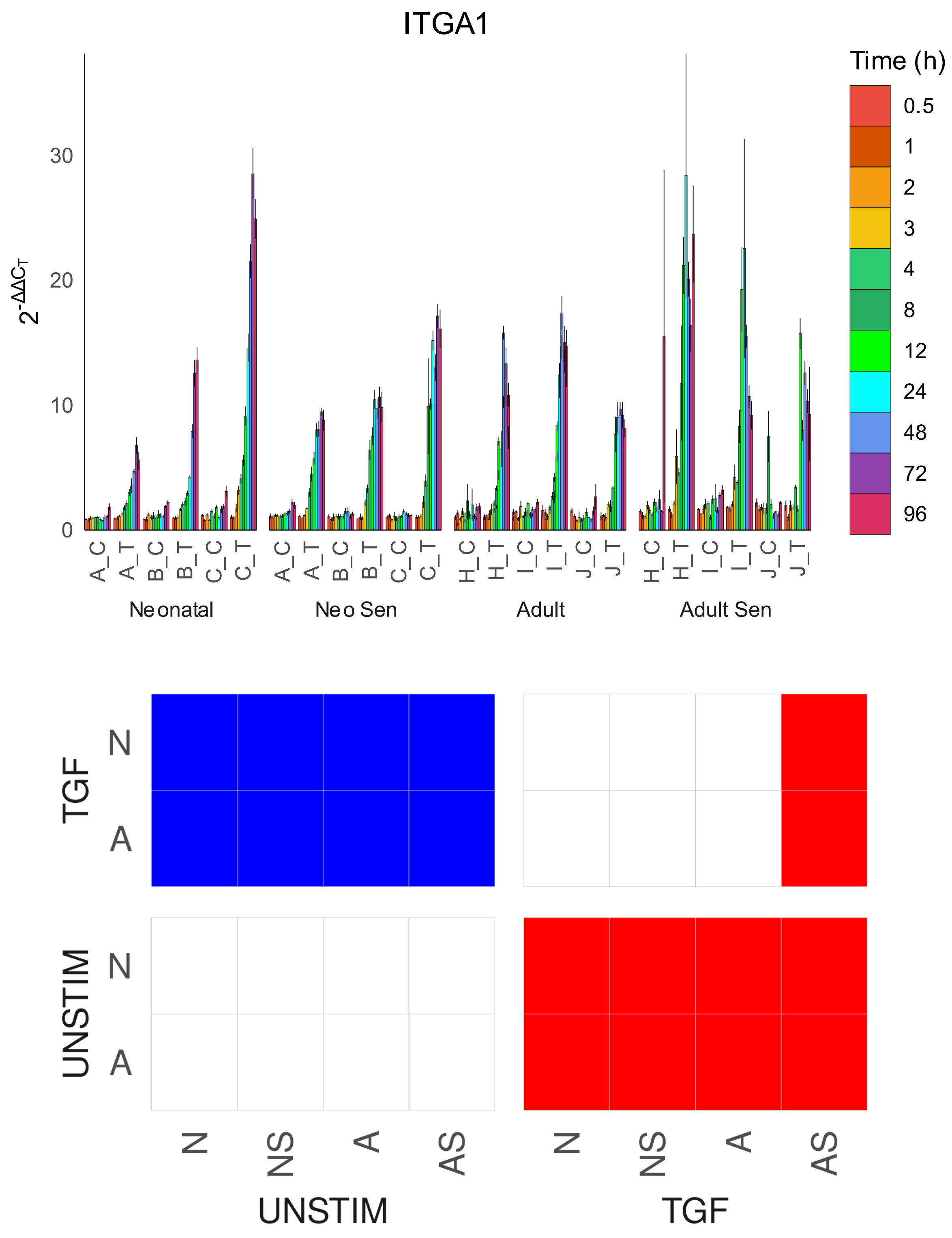
ITGA1
Time (h)
0.5
1
2
3
4
8
12
24
48
72
96
30
20
10
0
A_C
A_T
B_C
B_T
C_C
C_T
H_C
H_T
I_C
I_T
J_C
J_T
Neonatal
Neo Sen
Adult
Adult Sen
TGF
UNSTIM
N
A
N
NS
A
AS
UNSTIM
TGF

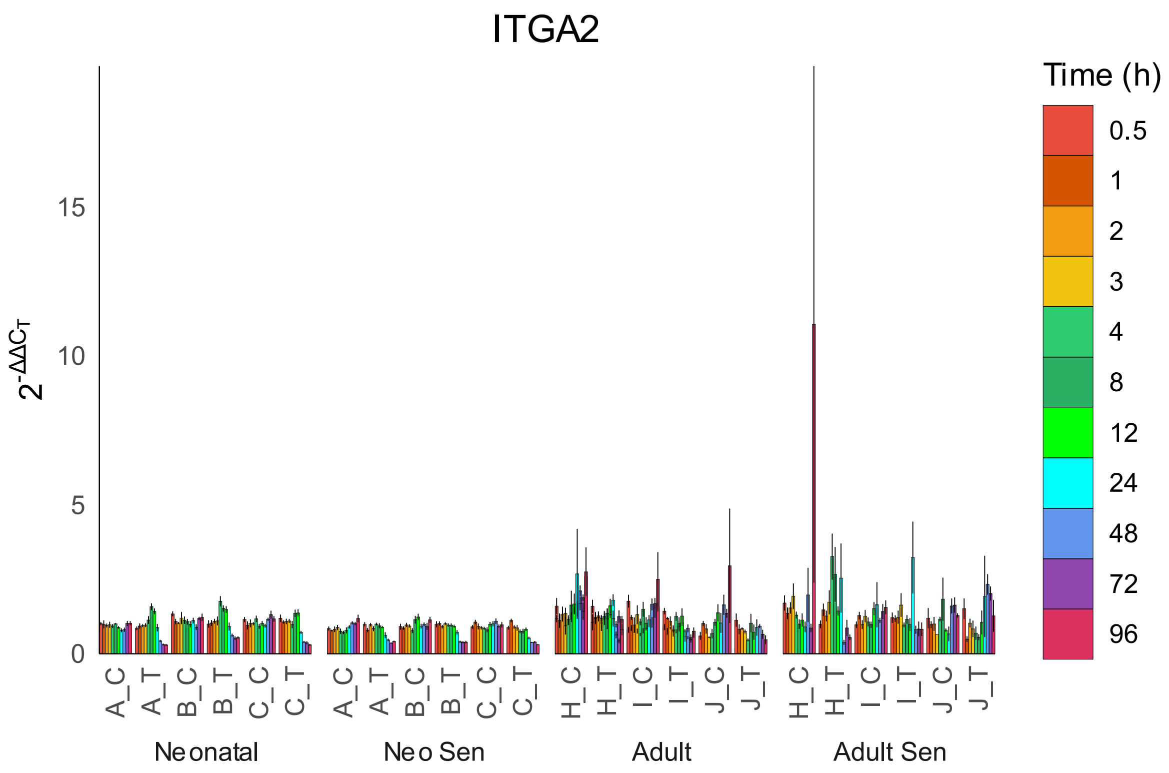
ITGA2
$2^{-\Delta\Delta C_T}$
0
5
10
15
A_C
A_T
B_C
B_T
C_C
C_T
H_C
H_T
I_C
I_T
J_C
J_T
Neonatal
Neo Sen
Adult
Adult Sen
Time (h)
0.5
1
2
3
4
8
12
24
48
72
96

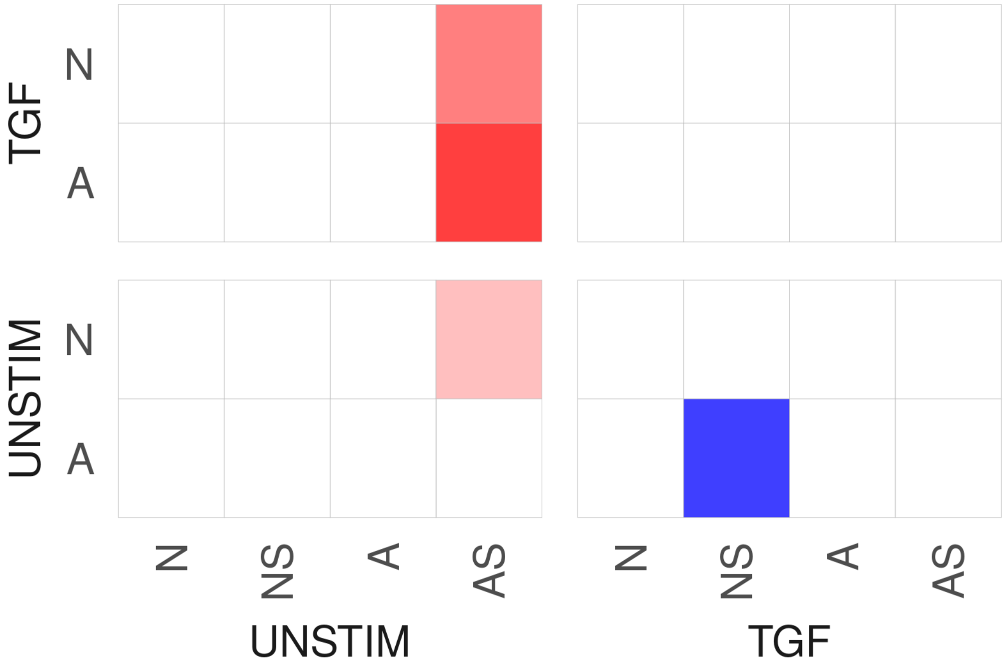
TGF
UNSTIM
N
A
N
NS
A
AS
UNSTIM
TGF

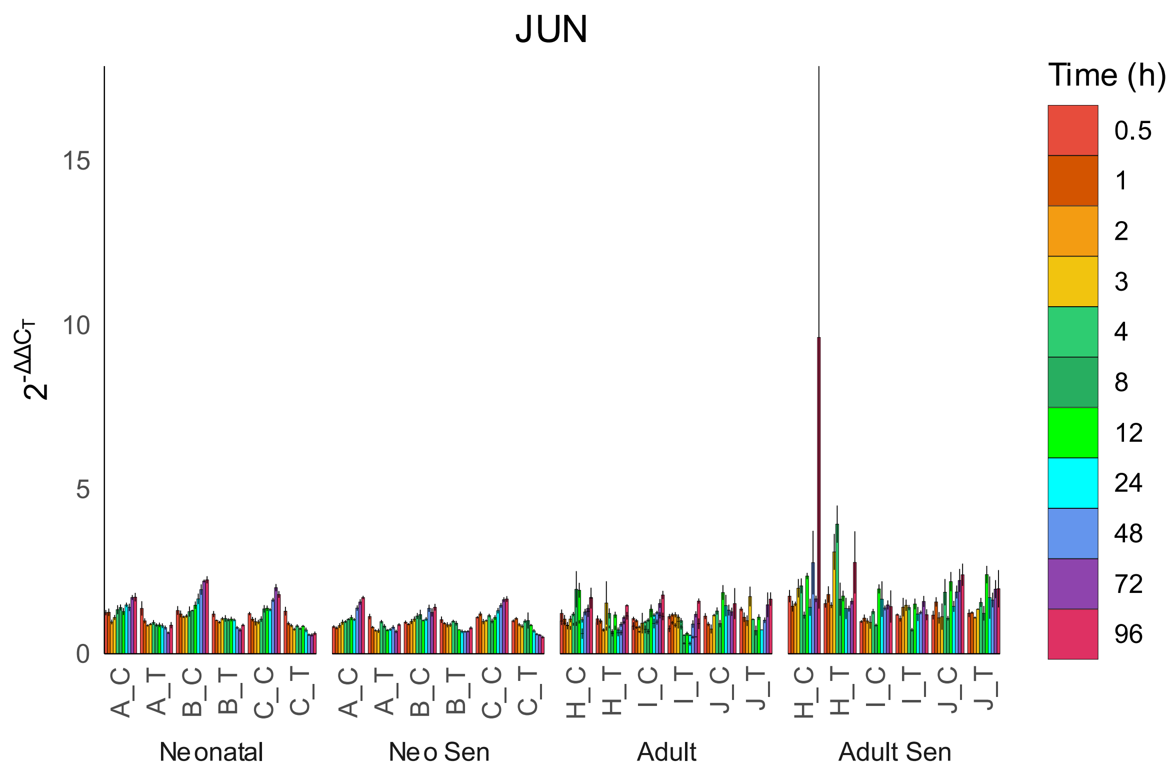
JUN
Time (h)
0.5
1
2
3
4
8
12
24
48
72
96
15
10
5
0
A_C
A_T
B_C
B_T
C_C
C_T
H_C
H_T
I_C
I_T
J_C
J_T
Neonatal
Neo Sen
Adult
Adult Sen

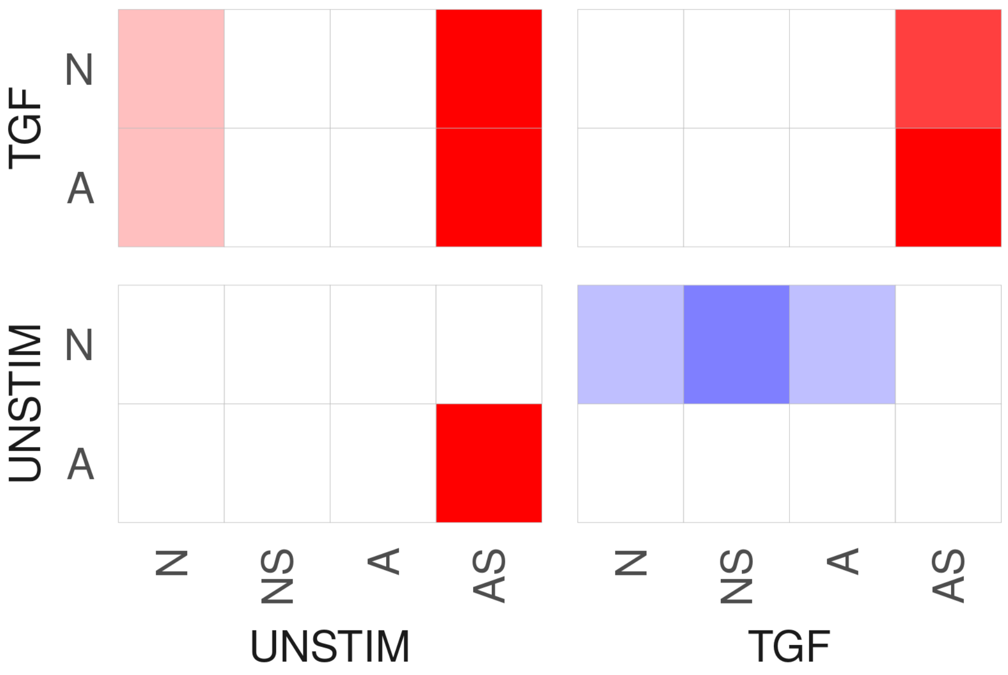
TGF
UNSTIM
N
A
N
NS
A
AS
UNSTIM
TGF

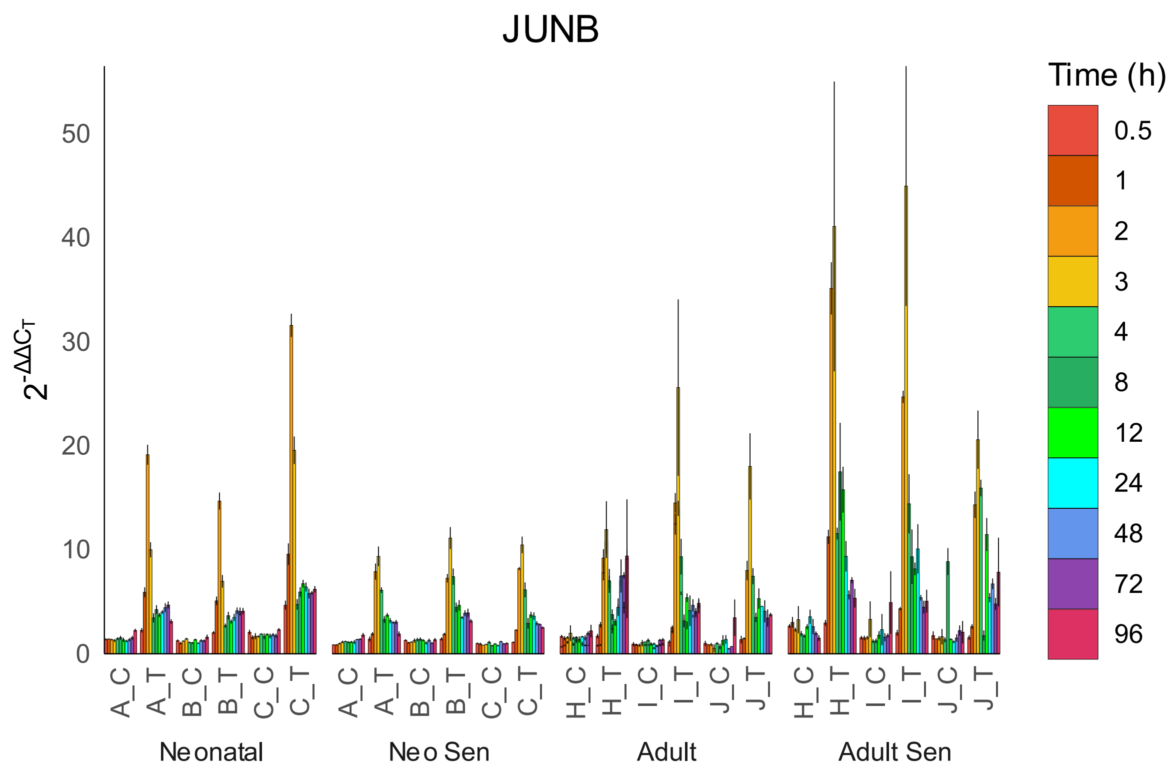
JUNB
Time (h)
0.5
1
2
3
4
8
12
24
48
72
96
50
40
30
20
10
0
$2^{-\Delta\Delta C_T}$
A_C
A_T
B_C
B_T
C_C
C_T
H_C
H_T
I_C
I_T
J_C
J_T
Neonatal
Neo Sen
Adult
Adult Sen

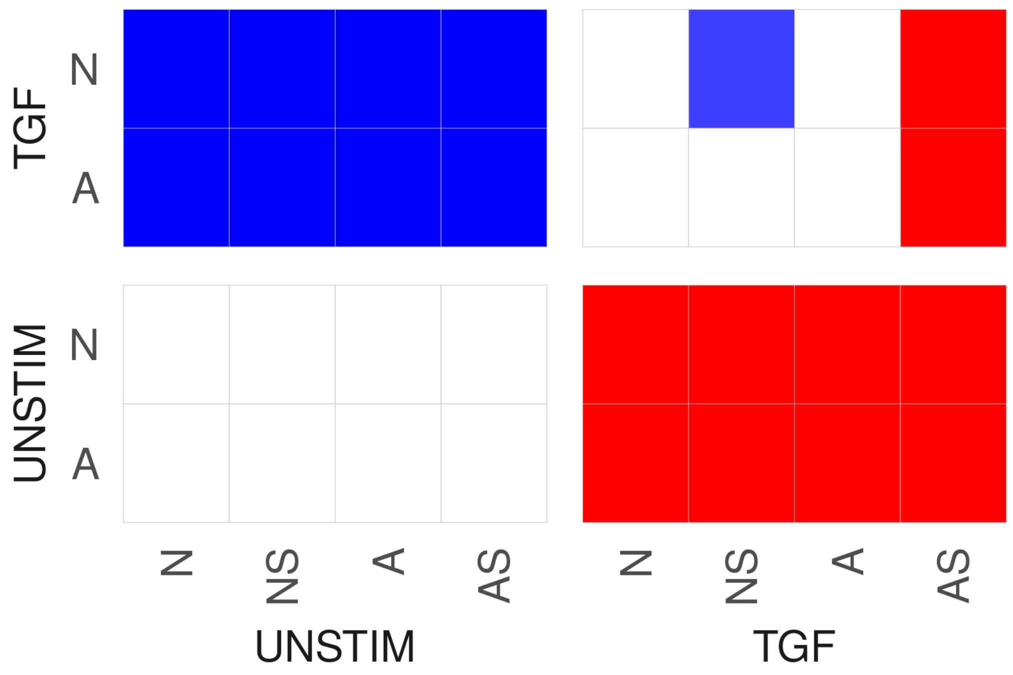
TGF
N
A
UNSTIM
N
A
N
NS
A
AS
UNSTIM
N
NS
A
AS
TGF

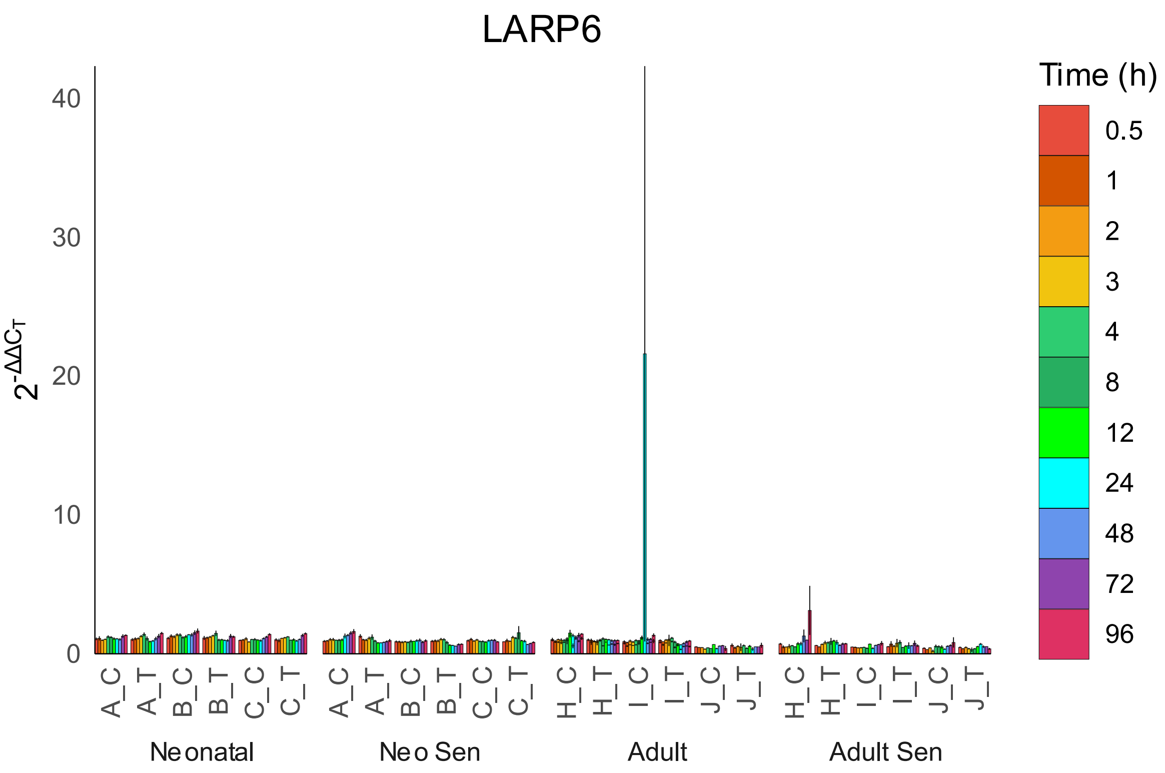
LARP6
Time (h)
0.5
1
2
3
4
8
12
24
48
72
96
40
30
20
10
0
$2^{-\Delta\Delta C_T}$
A_C
A_T
B_C
B_T
C_C
C_T
H_C
H_T
I_C
I_T
J_C
J_T
Neonatal
Neo Sen
Adult
Adult Sen

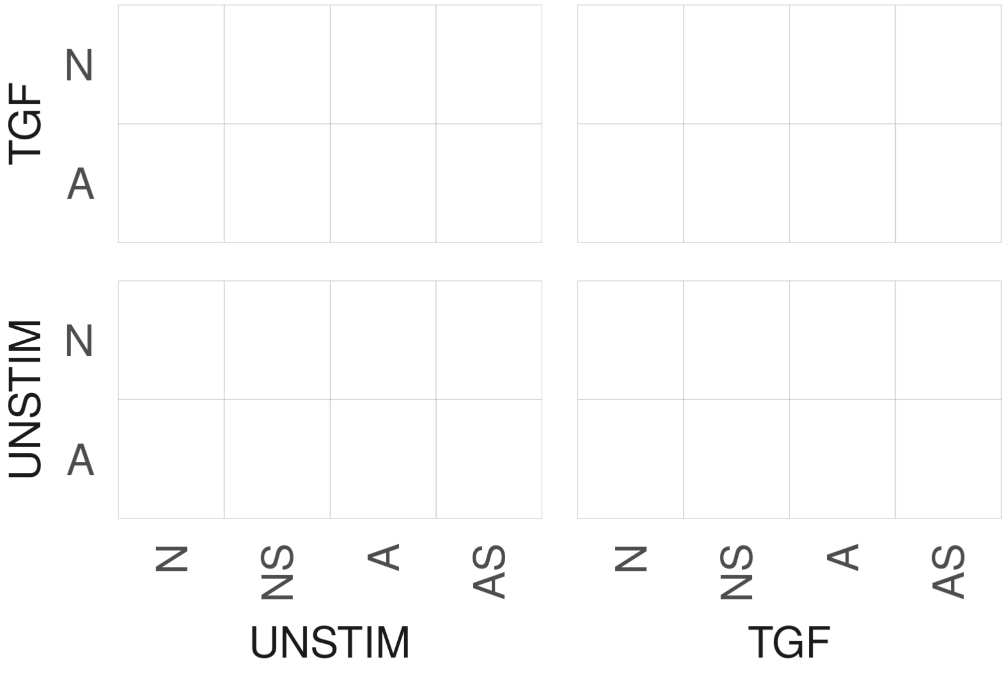
TGF
N
A
UNSTIM
N
A
N
NS
A
AS
UNSTIM
TGF

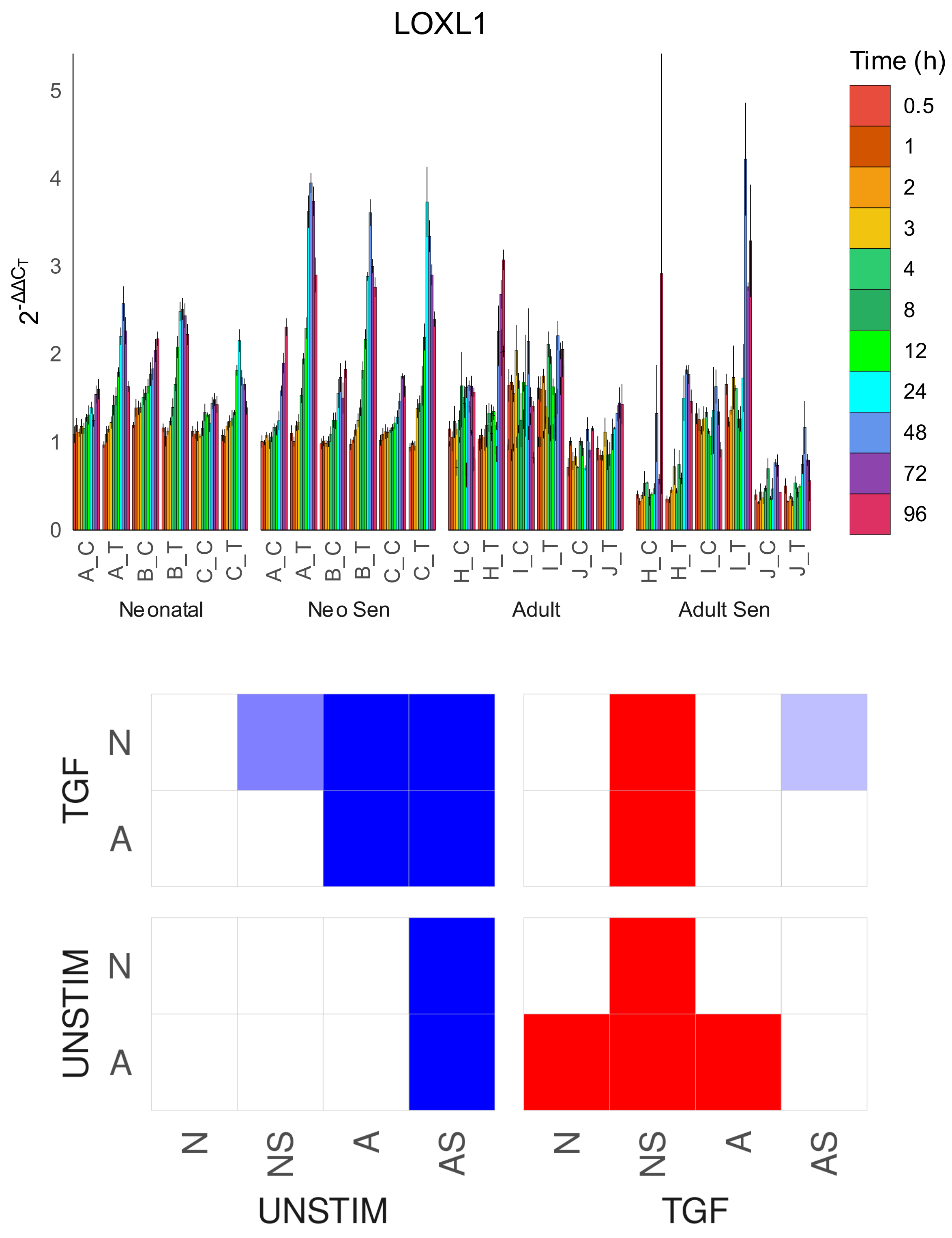

LOXL1
Time (h)
0.5
1
2
3
4
8
12
24
48
72
96
$2^{-\Delta\Delta C_T}$
0
1
2
3
4
5
A_C
A_T
B_C
B_T
C_C
C_T
H_C
H_T
I_C
I_T
J_C
J_T
Neonatal
Neo Sen
Adult
Adult Sen
TGF
UNSTIM
N
A
N
NS
A
AS
UNSTIM
TGF

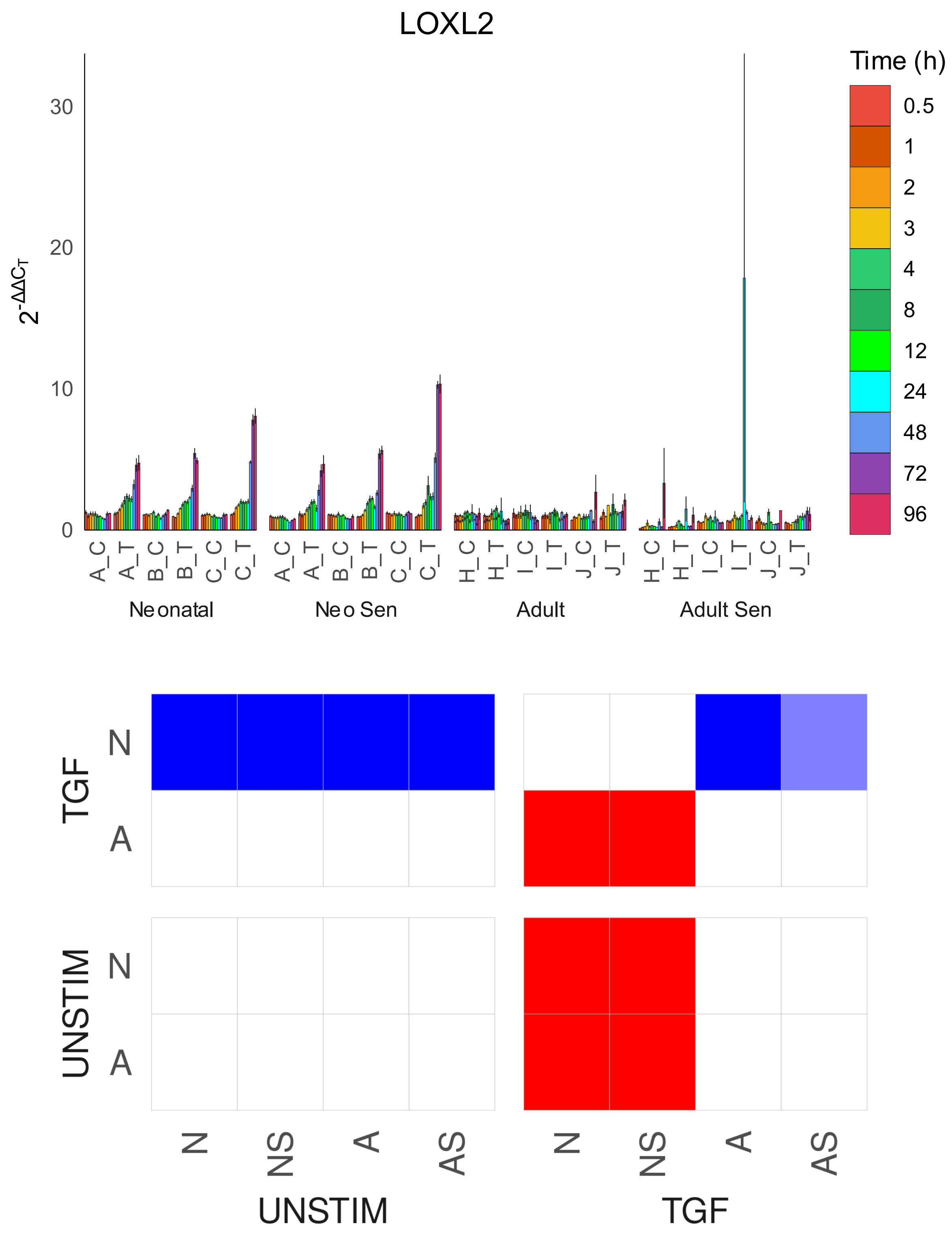

LOXL2
Time (h)
0.5
1
2
3
4
8
12
24
48
72
96
30
20
10
0
A_C
A_T
B_C
B_T
C_C
C_T
H_C
H_T
I_C
I_T
J_C
J_T
Neonatal
Neo Sen
Adult
Adult Sen
TGF
UNSTIM
N
A
N
NS
A
AS
UNSTIM
TGF

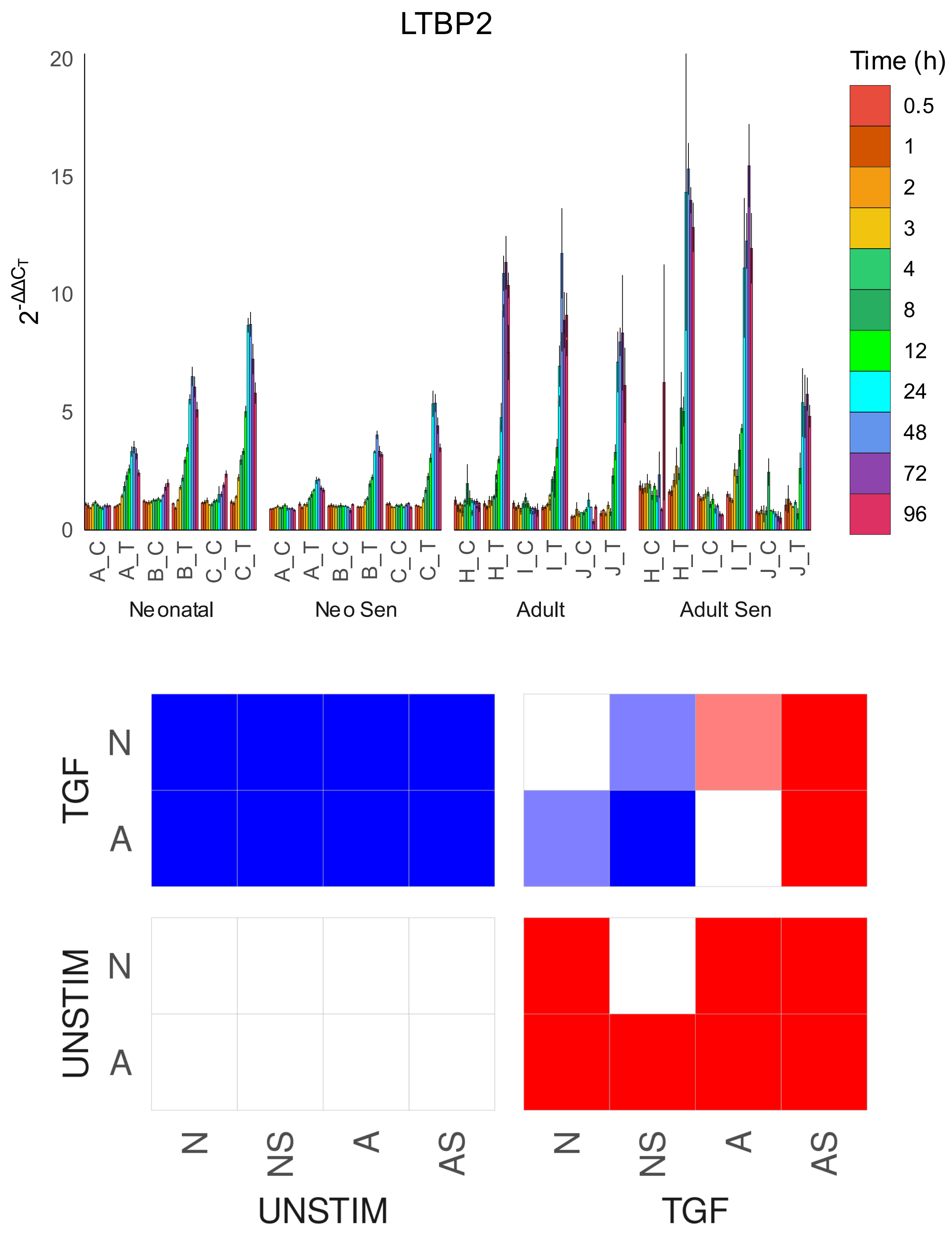

LTBP2
Time (h)
0.5
1
2
3
4
8
12
24
48
72
96
$2^{-\Delta\Delta C_T}$
20
15
10
5
0
A_C
A_T
B_C
B_T
C_C
C_T
H_C
H_T
I_C
I_T
J_C
J_T
Neonatal
Neo Sen
Adult
Adult Sen
TGF
UNSTIM
N
A
N
NS
A
AS
UNSTIM
TGF

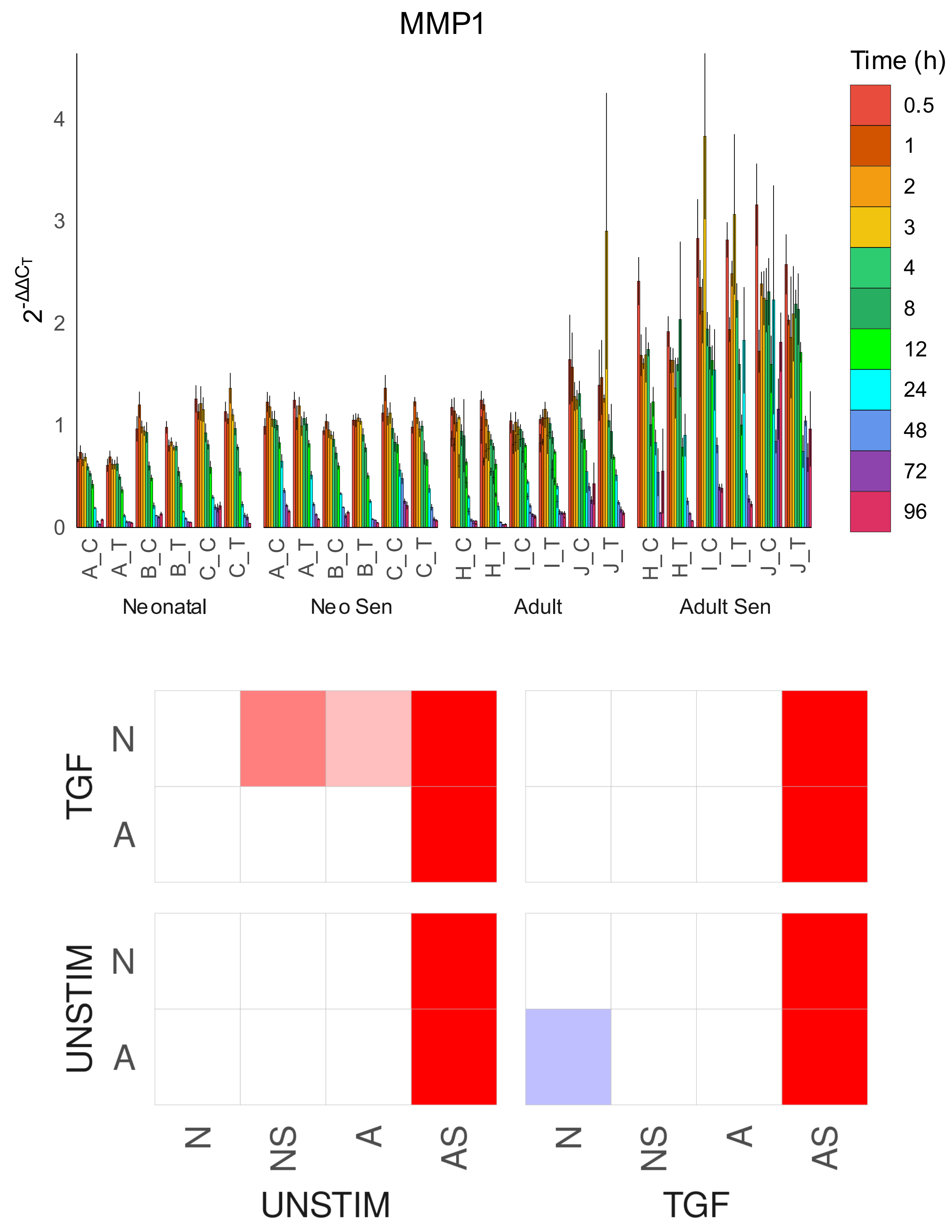

MMP1
Time (h)
0.5
1
2
3
4
8
12
24
48
72
96
$2^{-\Delta\Delta C_T}$
0
1
2
3
4
A_C
A_T
B_C
B_T
C_C
C_T
H_C
H_T
I_C
I_T
J_C
J_T
Neonatal
Neo Sen
Adult
Adult Sen
TGF
UNSTIM
N
A
N
NS
A
AS
UNSTIM
TGF

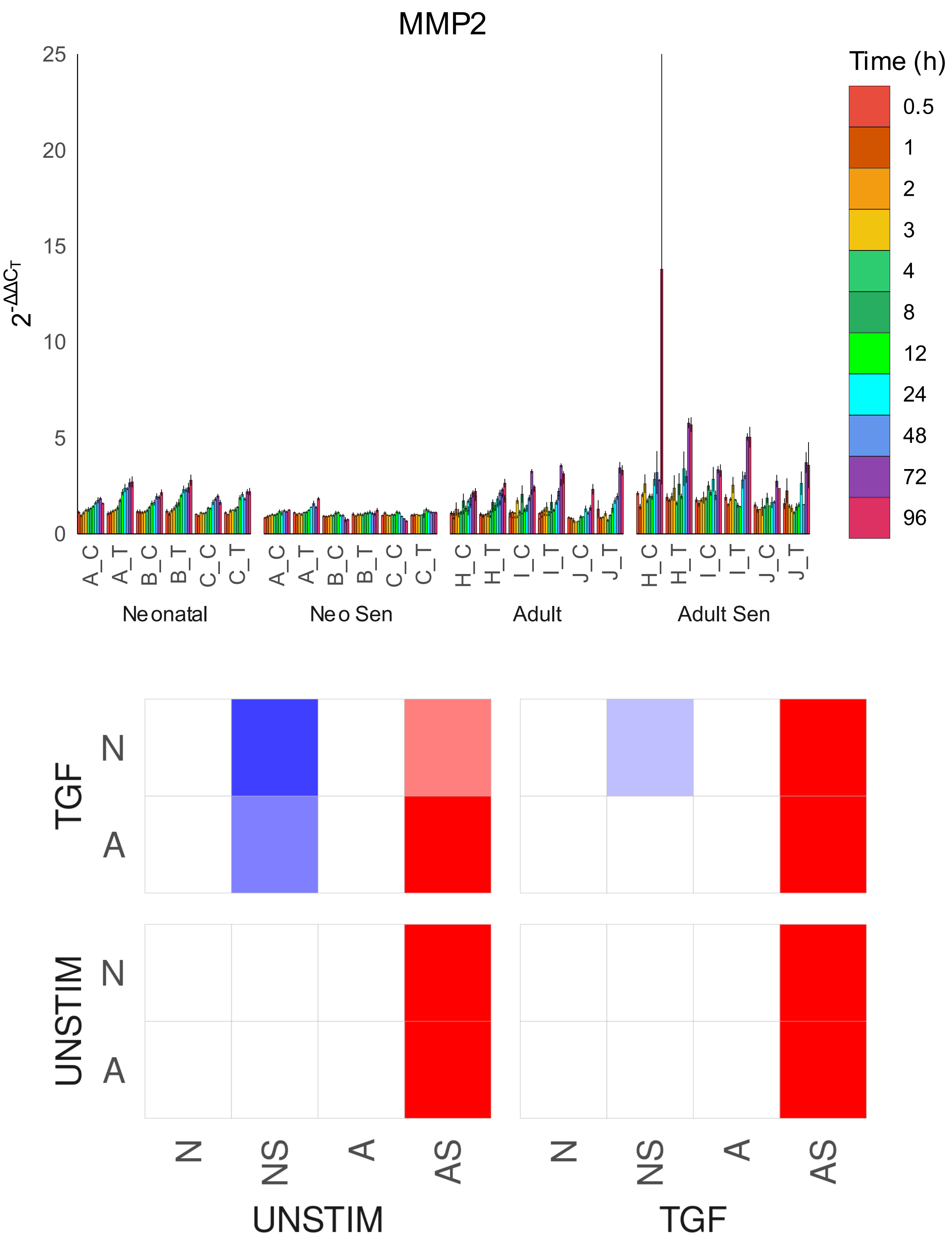
MMP2
25
20
15
10
5
0
$2^{-\Delta\Delta C_T}$
Time (h)
0.5
1
2
3
4
8
12
24
48
72
96
A_C
A_T
B_C
B_T
C_C
C_T
H_C
H_T
I_C
I_T
J_C
J_T
Neonatal
Neo Sen
Adult
Adult Sen
TGF
UNSTIM
N
A
N
NS
A
AS
UNSTIM
TGF

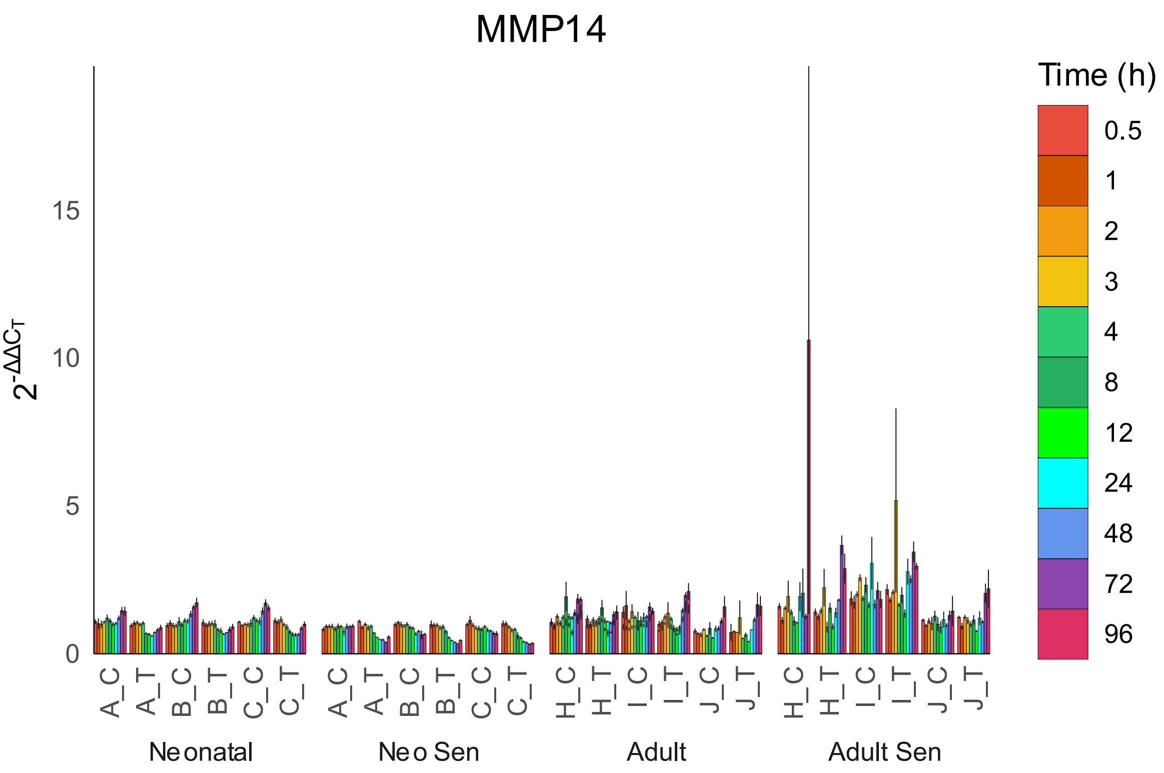
MMP14
Time (h)
0.5
1
2
3
4
8
12
24
48
72
96
$2^{-\Delta\Delta C_T}$
0
5
10
15
A_C
A_T
B_C
B_T
C_C
C_T
H_C
H_T
I_C
I_T
J_C
J_T
Neonatal
Neo Sen
Adult
Adult Sen

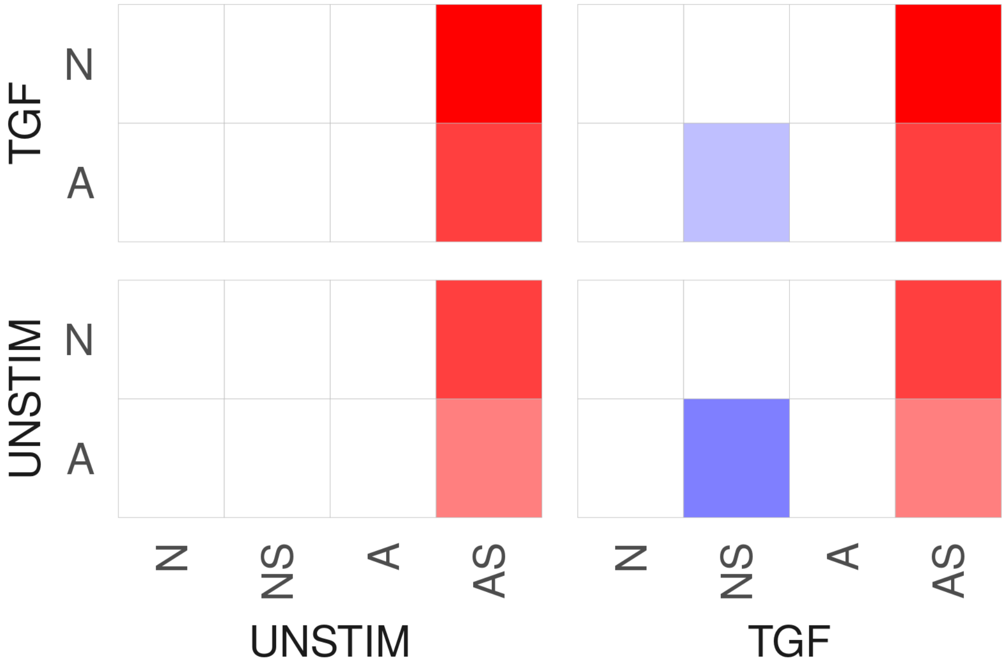
TGF
UNSTIM
N
A
N
NS
A
AS
UNSTIM
TGF

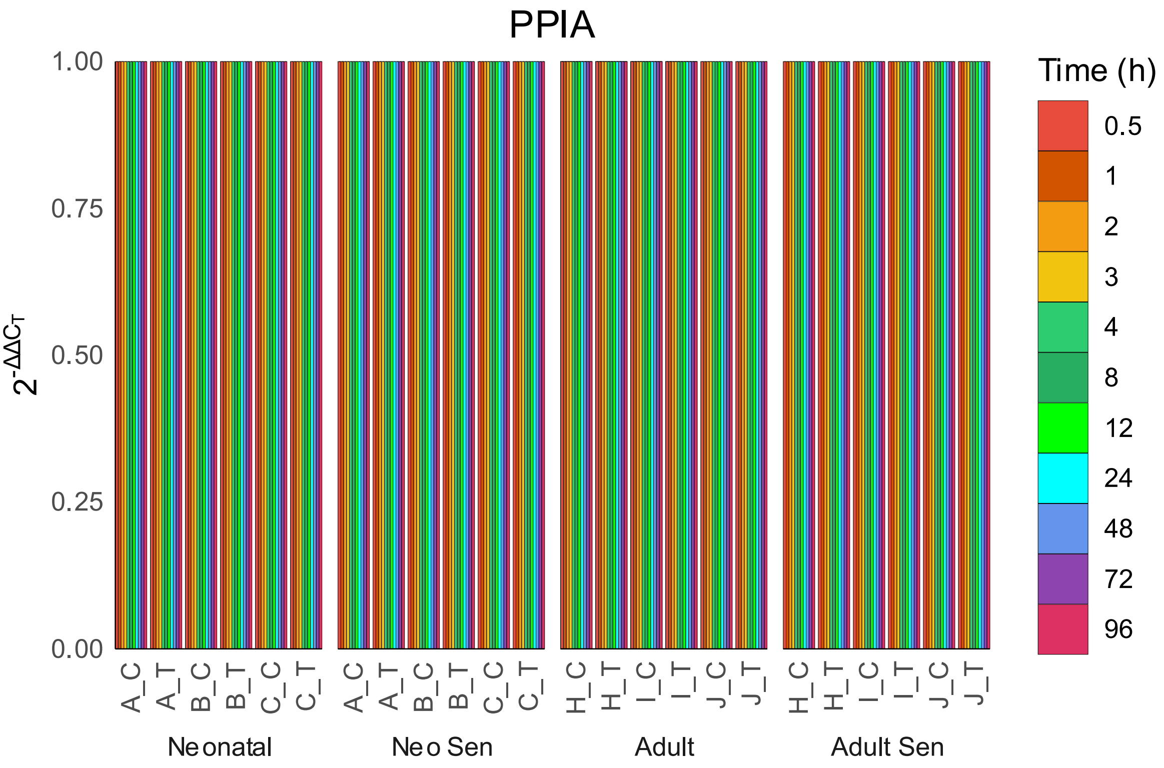
PPIA
Time (h)
0.5
1
2
3
4
8
12
24
48
72
96
1.00
0.75
0.50
0.25
0.00
A_C
A_T
B_C
B_T
C_C
C_T
H_C
H_T
I_C
I_T
J_C
J_T
Neonatal
Neo Sen
Adult
Adult Sen

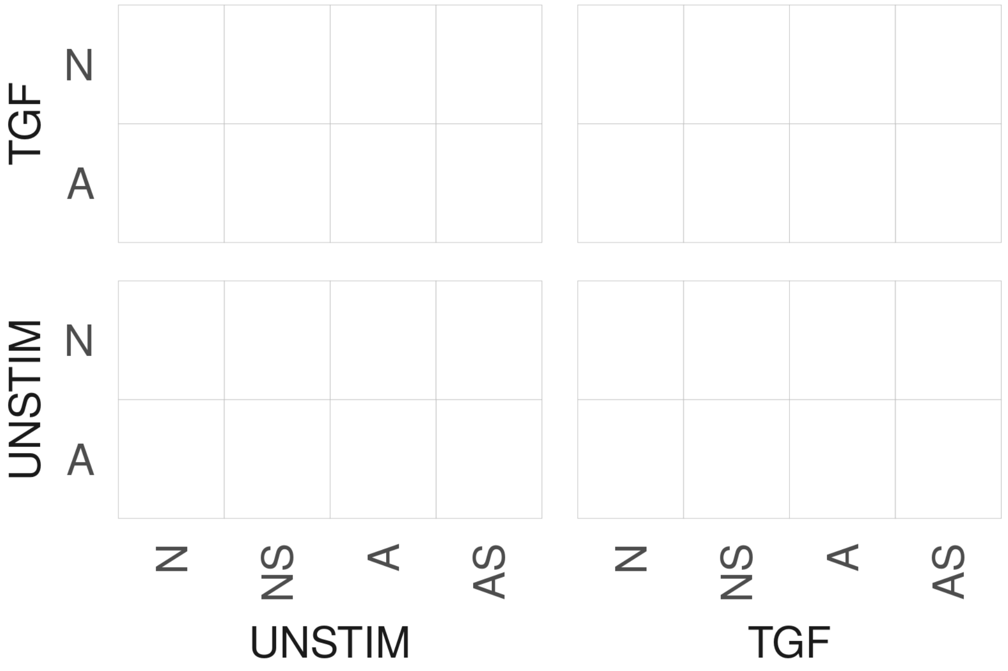
TGF
UNSTIM
N
A
N
NS
A
AS
UNSTIM
TGF

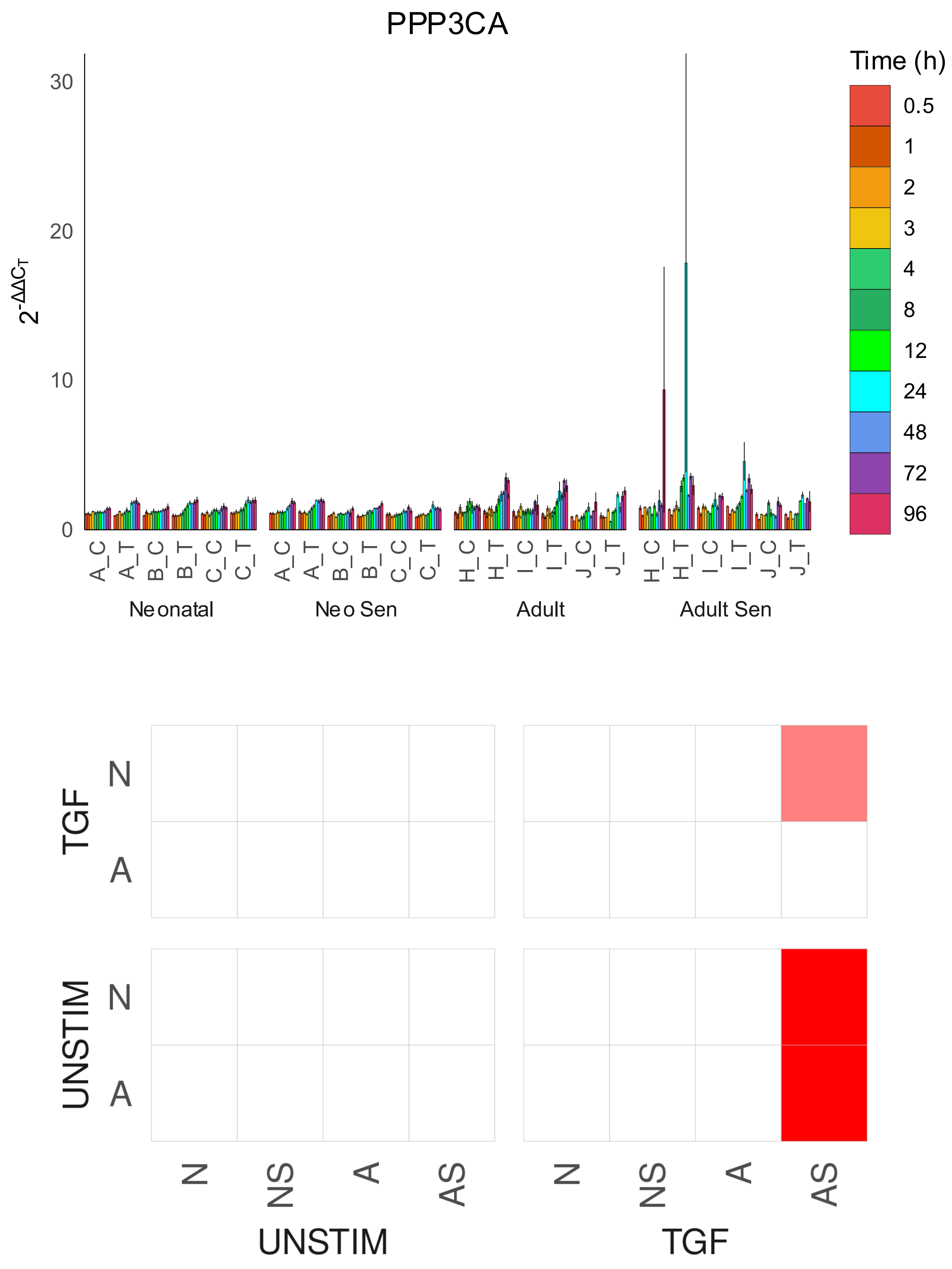

PPP3CA
Time (h)
0.5
1
2
3
4
8
12
24
48
72
96
$2^{-\Delta\Delta C_T}$
30
20
10
0
A_C
A_T
B_C
B_T
C_C
C_T
H_C
H_T
I_C
I_T
J_C
J_T
Neonatal
Neo Sen
Adult
Adult Sen
TGF
UNSTIM
N
A
N
NS
A
AS
UNSTIM
TGF

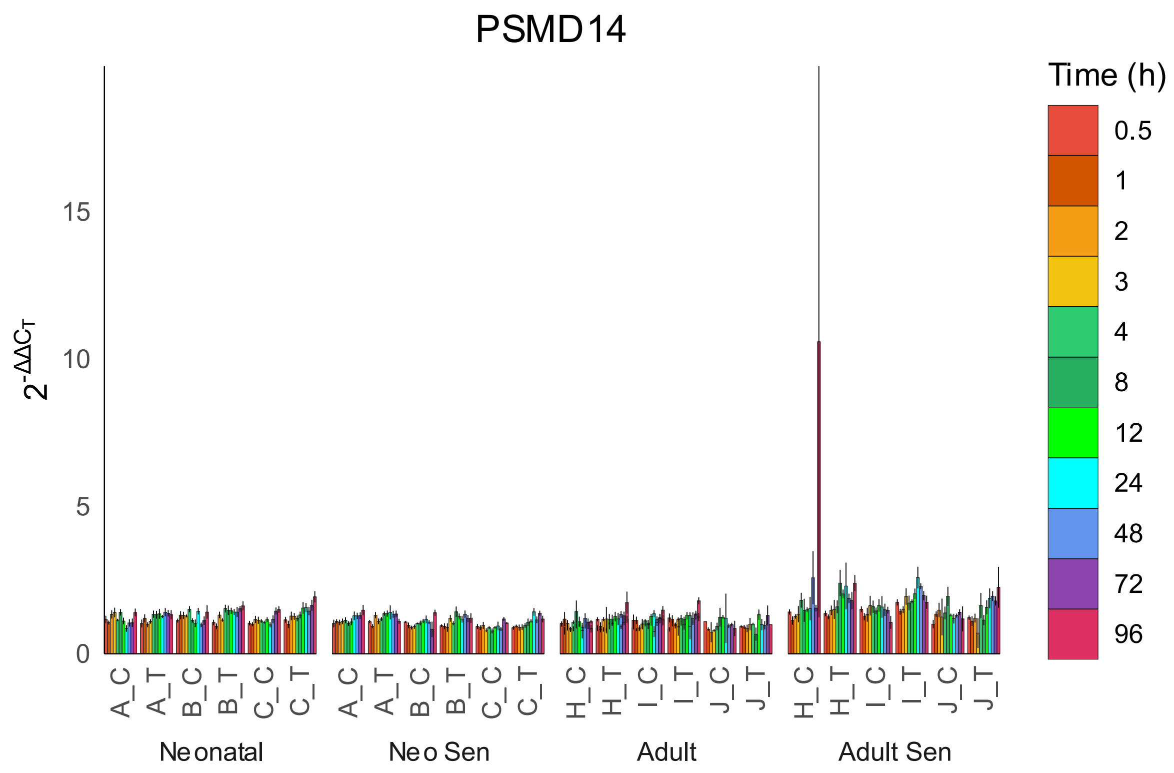
PSMD14
$2^{-\Delta\Delta C_T}$
0
5
10
15
A_C
A_T
B_C
B_T
C_C
C_T
H_C
H_T
I_C
I_T
J_C
J_T
Neonatal
Neo Sen
Adult
Adult Sen
Time (h)
0.5
1
2
3
4
8
12
24
48
72
96

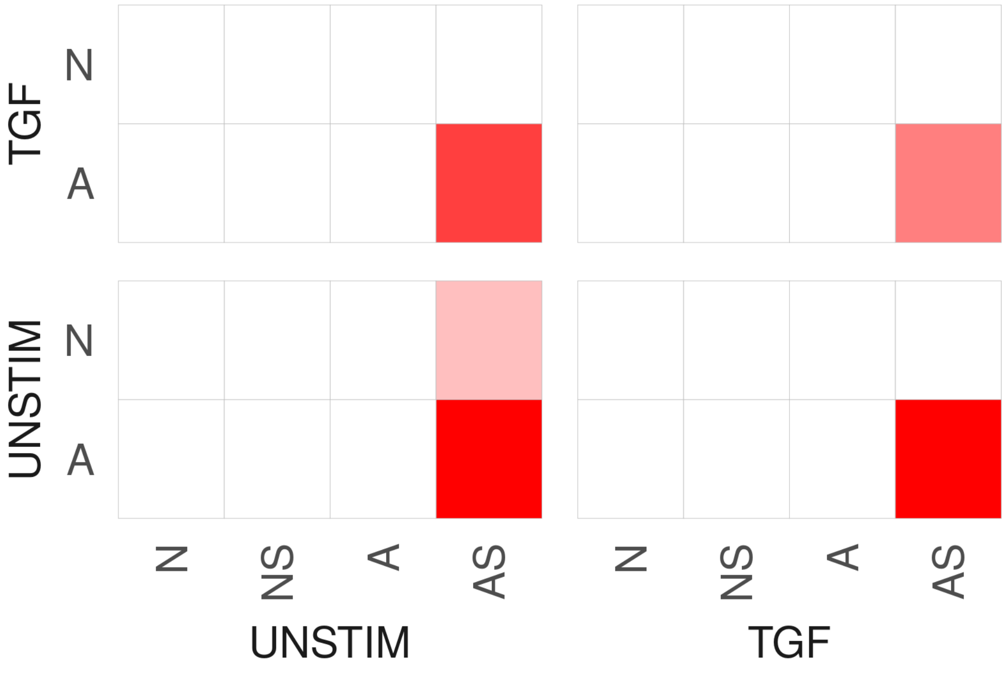
TGF
UNSTIM
N
A
N
NS
A
AS
UNSTIM
TGF

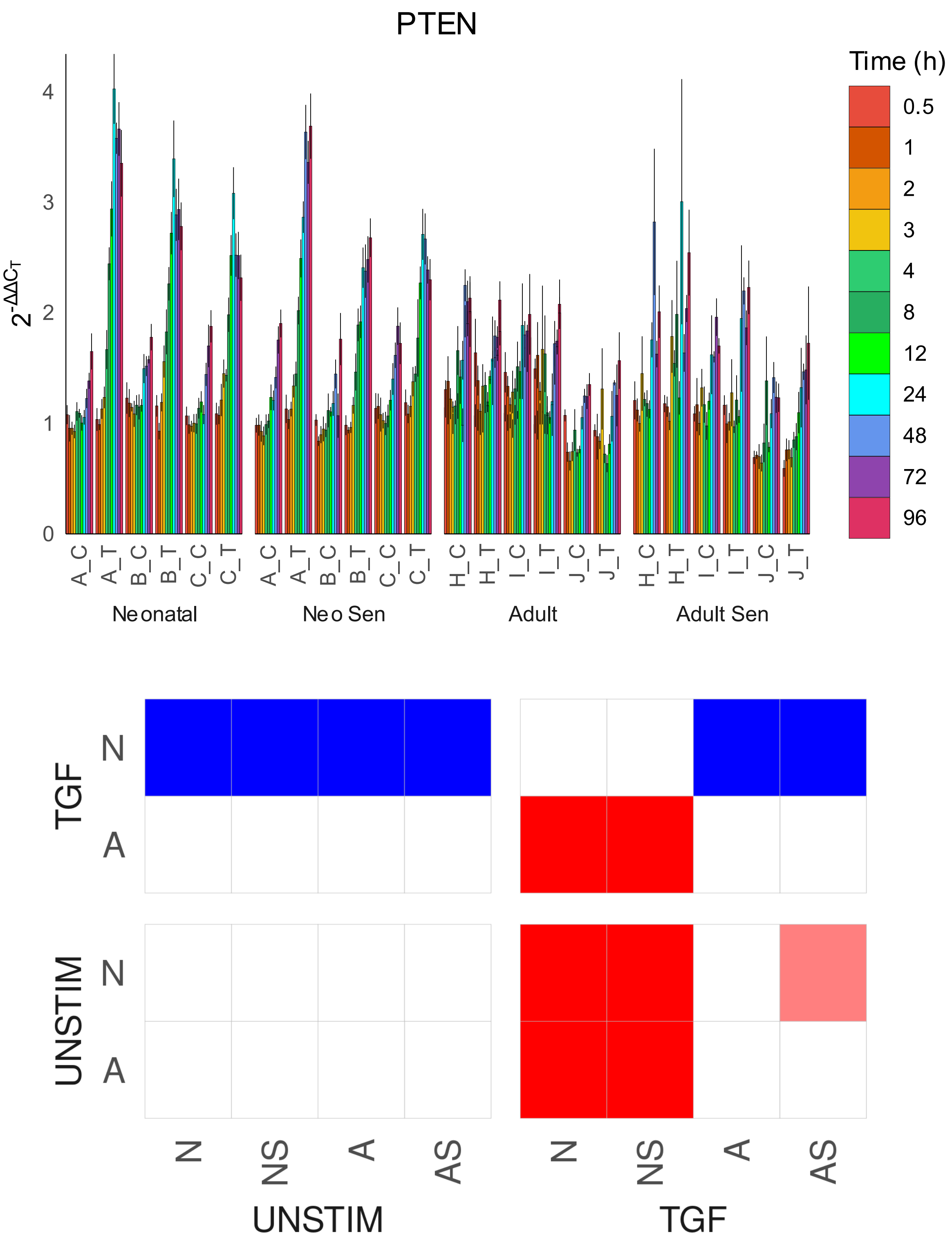
PTEN
Time (h)
0.5
1
2
3
4
8
12
24
48
72
96
$2^{-\Delta\Delta C_T}$
0
1
2
3
4
A_C
A_T
B_C
B_T
C_C
C_T
H_C
H_T
I_C
I_T
J_C
J_T
Neonatal
Neo Sen
Adult
Adult Sen
TGF
UNSTIM
N
A
N
NS
A
AS
UNSTIM
TGF

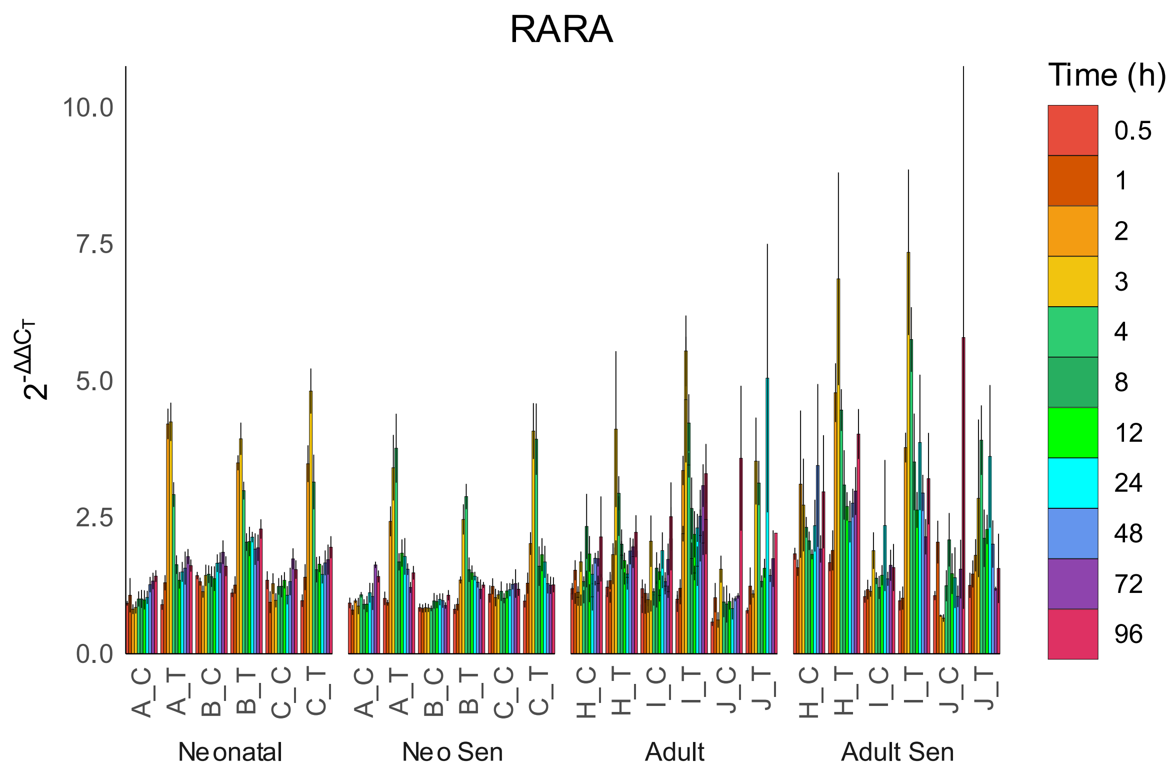
RARA
Time (h)
0.5
1
2
3
4
8
12
24
48
72
96
10.0
7.5
5.0
2.5
0.0
A_C
A_T
B_C
B_T
C_C
C_T
H_C
H_T
I_C
I_T
J_C
J_T
Neonatal
Neo Sen
Adult
Adult Sen

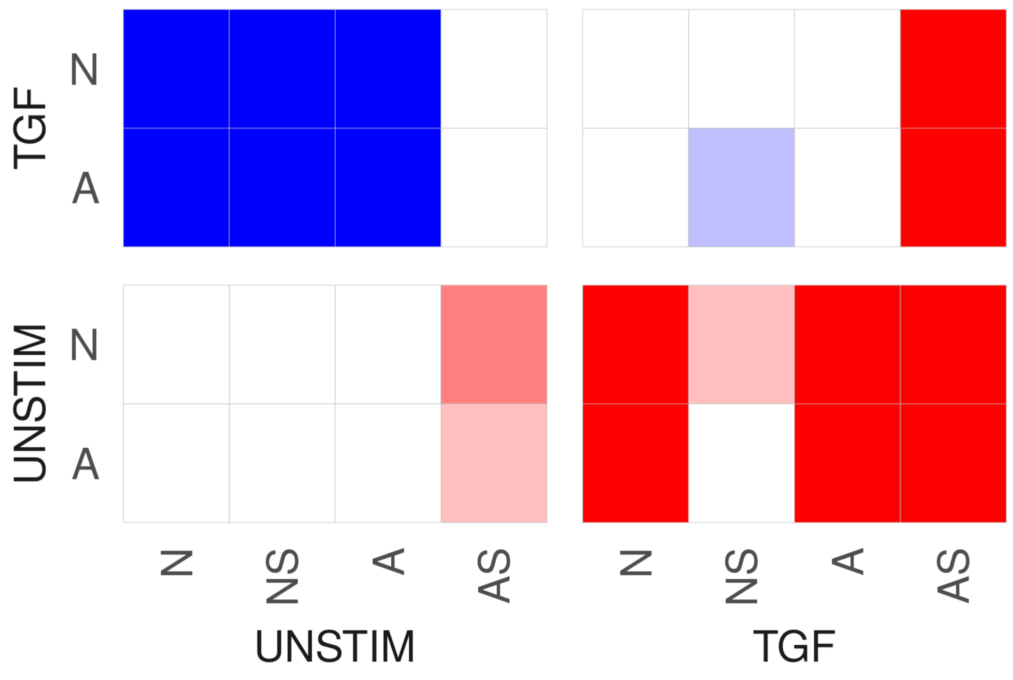
TGF
UNSTIM
N
A
N
NS
A
AS
UNSTIM
TGF

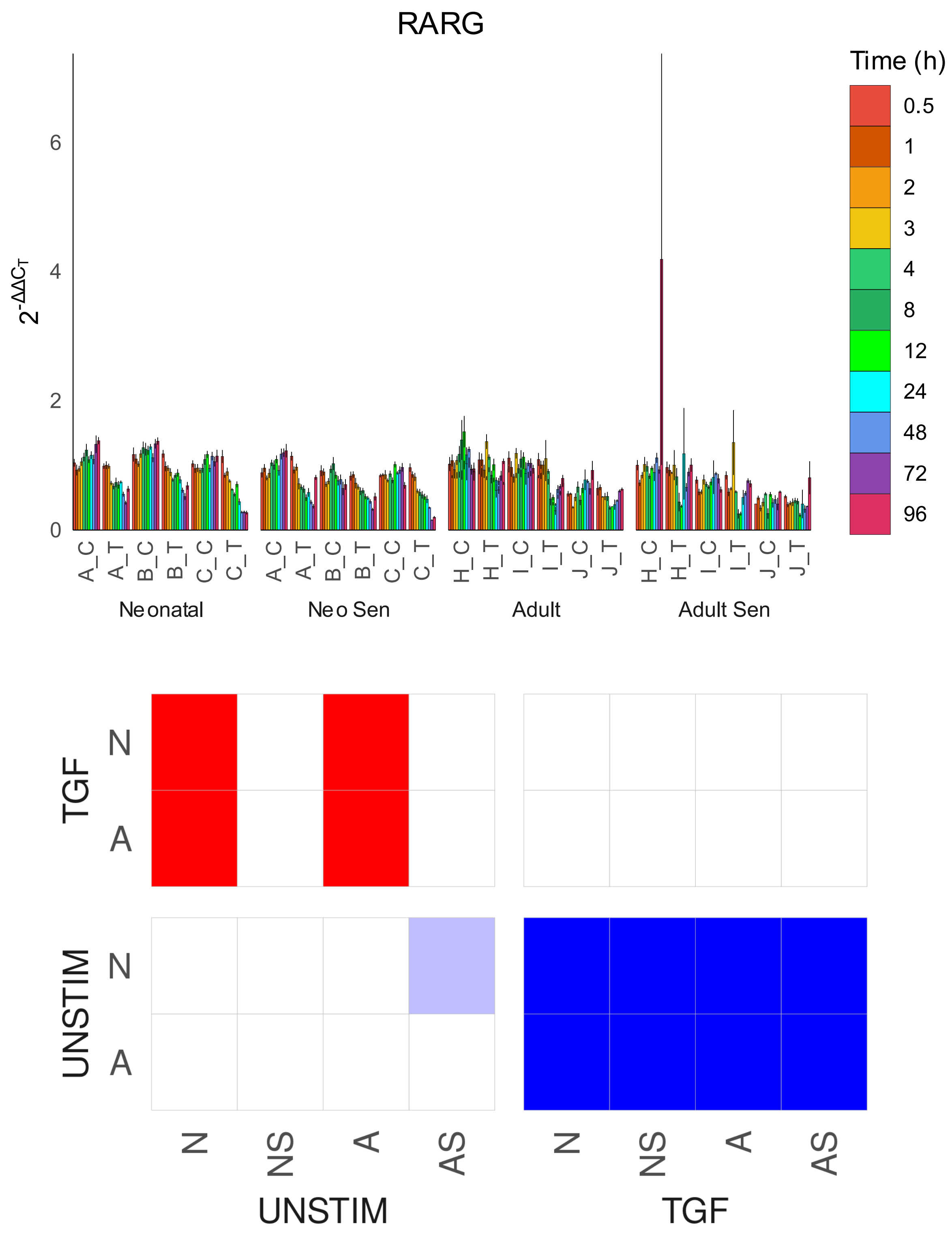

RARG
Time (h)
0.5
1
2
3
4
8
12
24
48
72
96
2-ΔΔCT
0
2
4
6
A_C
A_T
B_C
B_T
C_C
C_T
H_C
H_T
I_C
I_T
J_C
J_T
Neonatal
Neo Sen
Adult
Adult Sen
TGF
UNSTIM
N
A
N
NS
A
AS
UNSTIM
TGF

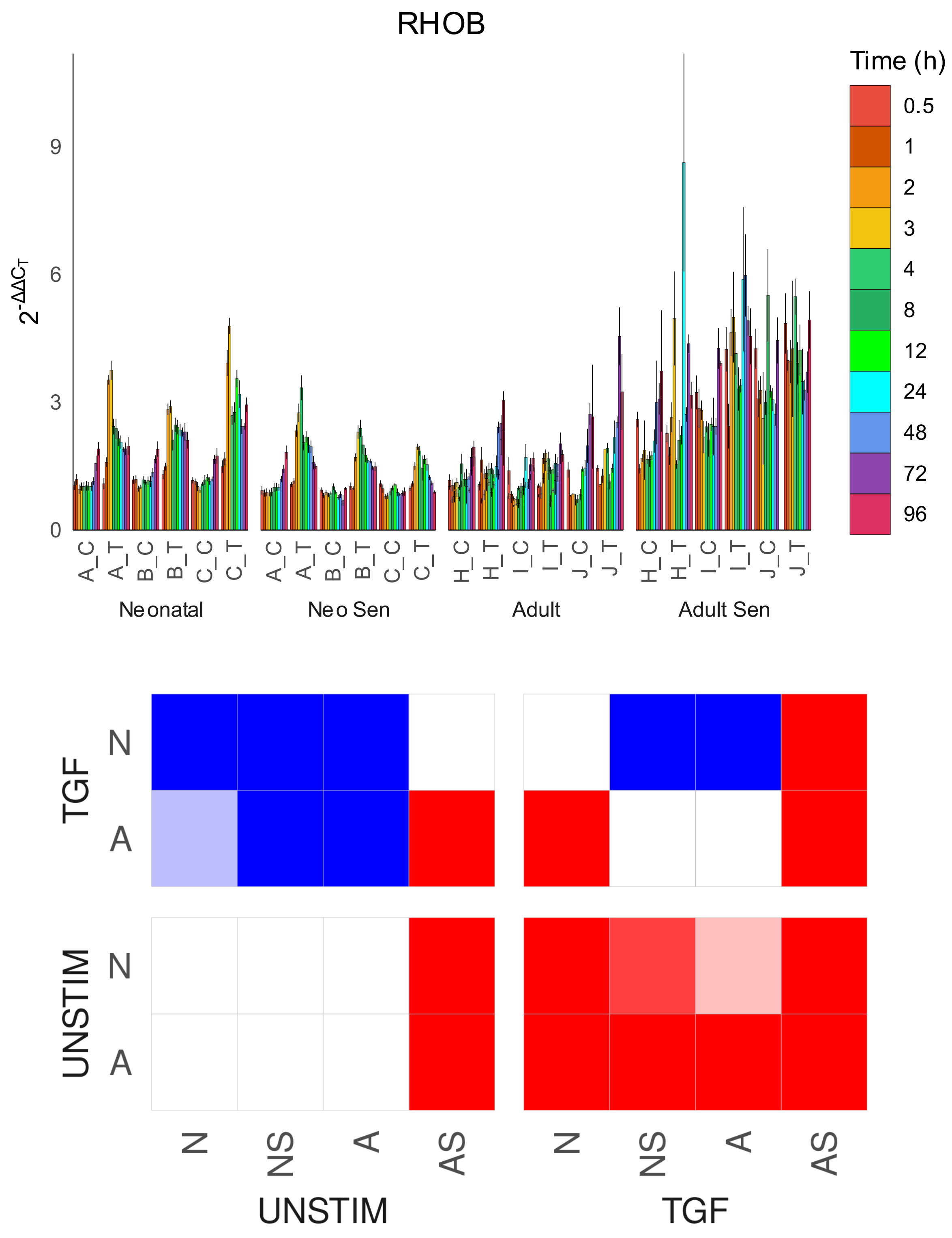

RHOB
Time (h)
0.5
1
2
3
4
8
12
24
48
72
96
$2^{-\Delta\Delta C_T}$
0
3
6
9
A_C
A_T
B_C
B_T
C_C
C_T
H_C
H_T
I_C
I_T
J_C
J_T
Neonatal
Neo Sen
Adult
Adult Sen
TGF
UNSTIM
N
A
N
NS
A
AS
UNSTIM
TGF

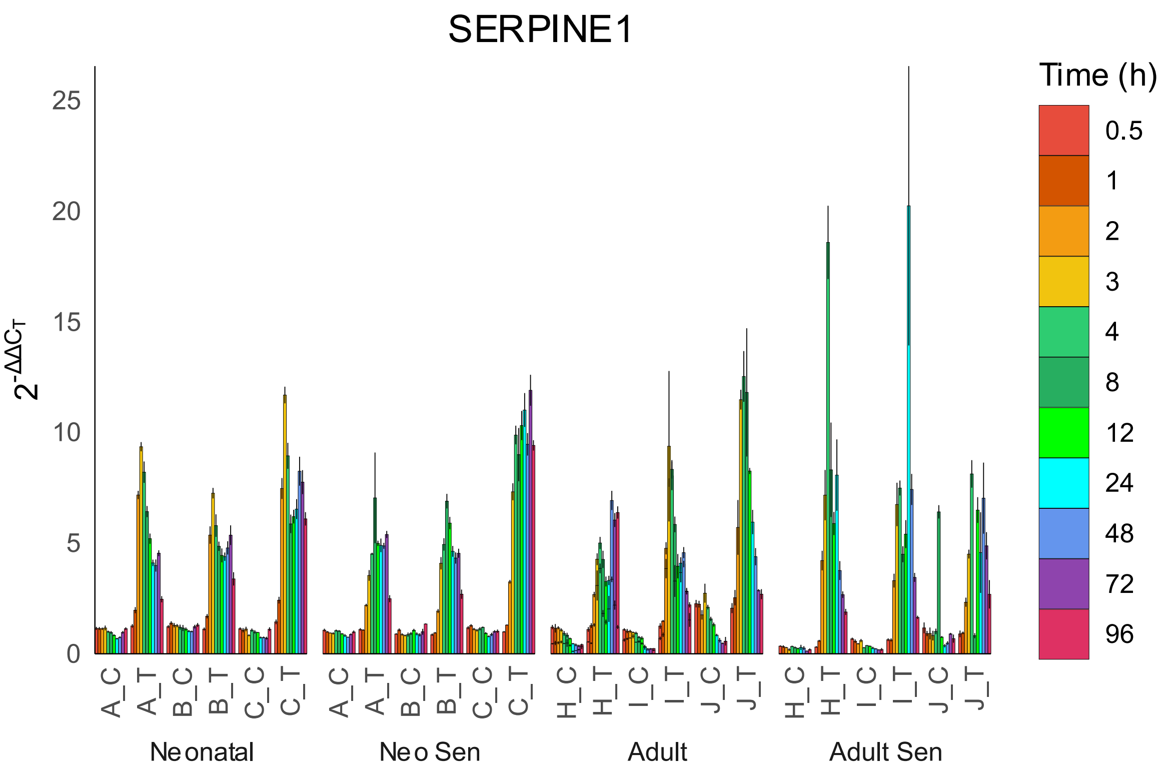

SERPINE1
Time (h)
0.5
1
2
3
4
8
12
24
48
72
96
$2^{-\Delta\Delta C_T}$
25
20
15
10
5
0
A_C
A_T
B_C
B_T
C_C
C_T
H_C
H_T
I_C
I_T
J_C
J_T
Neonatal
Neo Sen
Adult
Adult Sen


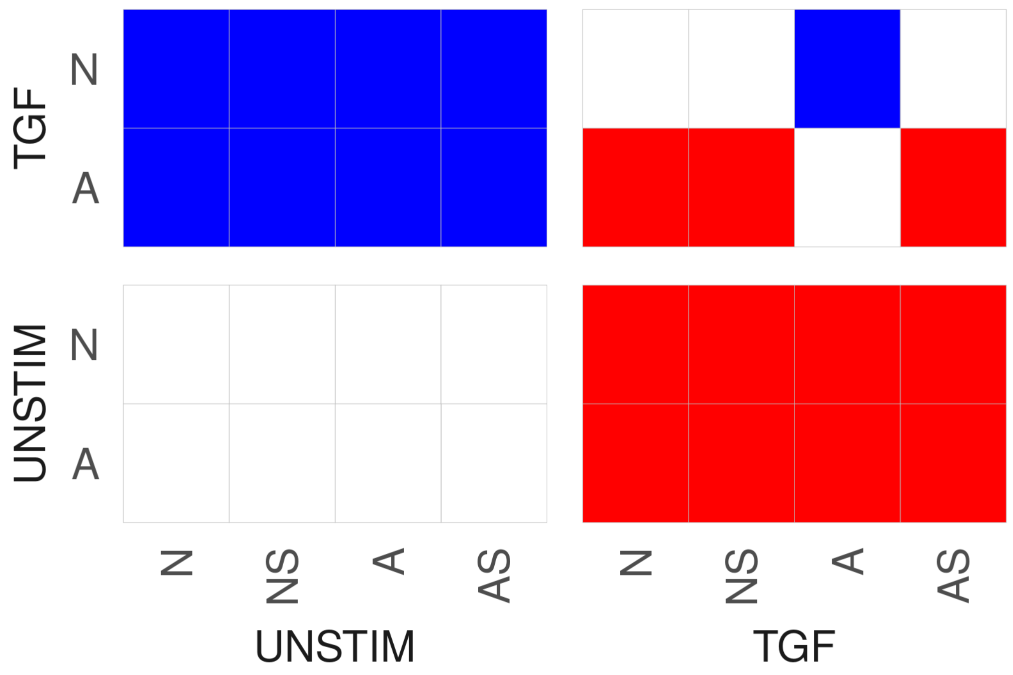

TGF
N
A
UNSTIM
N
A
N
NS
A
AS
UNSTIM
N
NS
A
AS
TGF

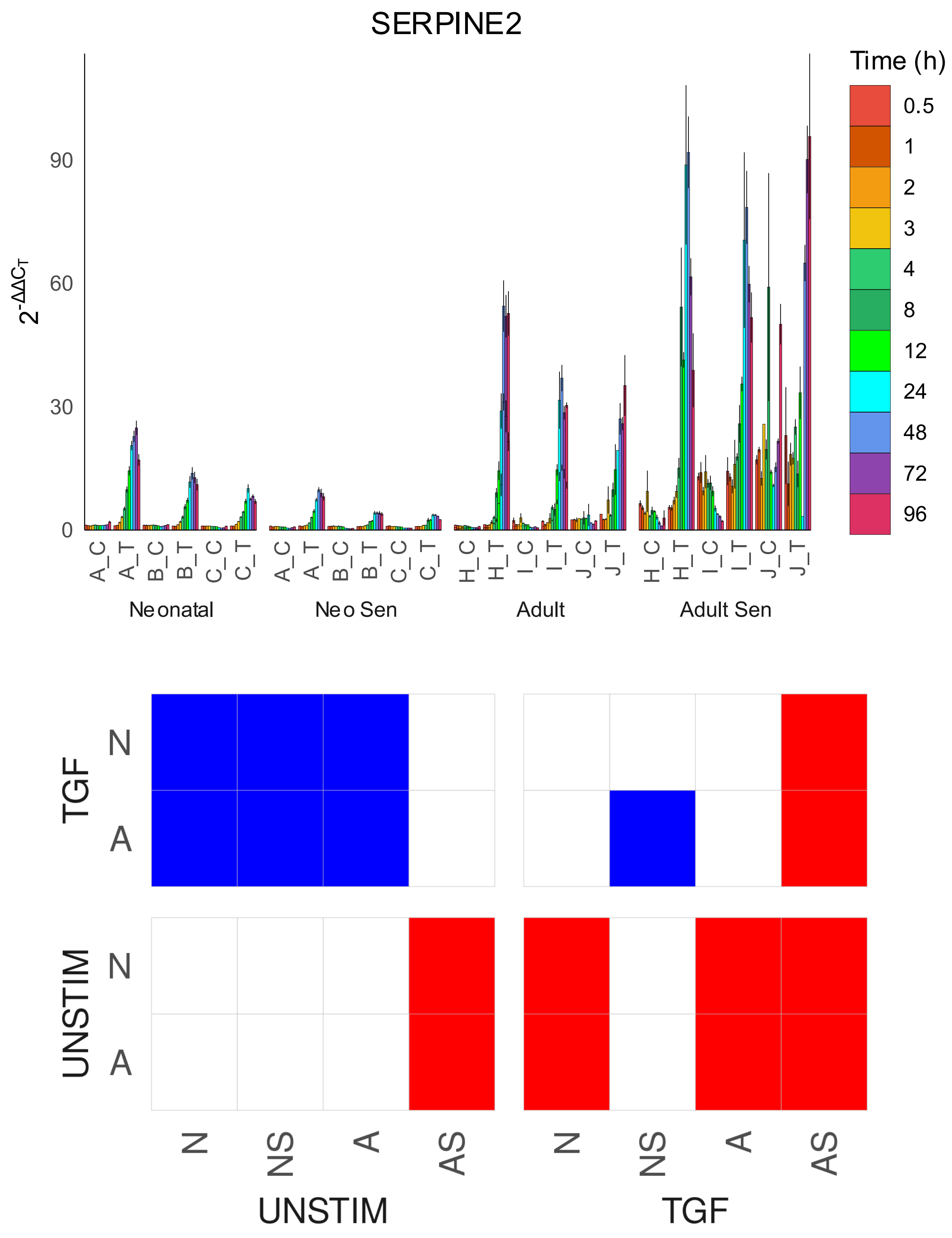
SERPINE2
Time (h)
0.5
1
2
3
4
8
12
24
48
72
96
$2^{-\Delta\Delta C_T}$
0
30
60
90
A_C
A_T
B_C
B_T
C_C
C_T
H_C
H_T
I_C
I_T
J_C
J_T
Neonatal
Neo Sen
Adult
Adult Sen
TGF
UNSTIM
N
A
N
NS
A
AS
UNSTIM
TGF

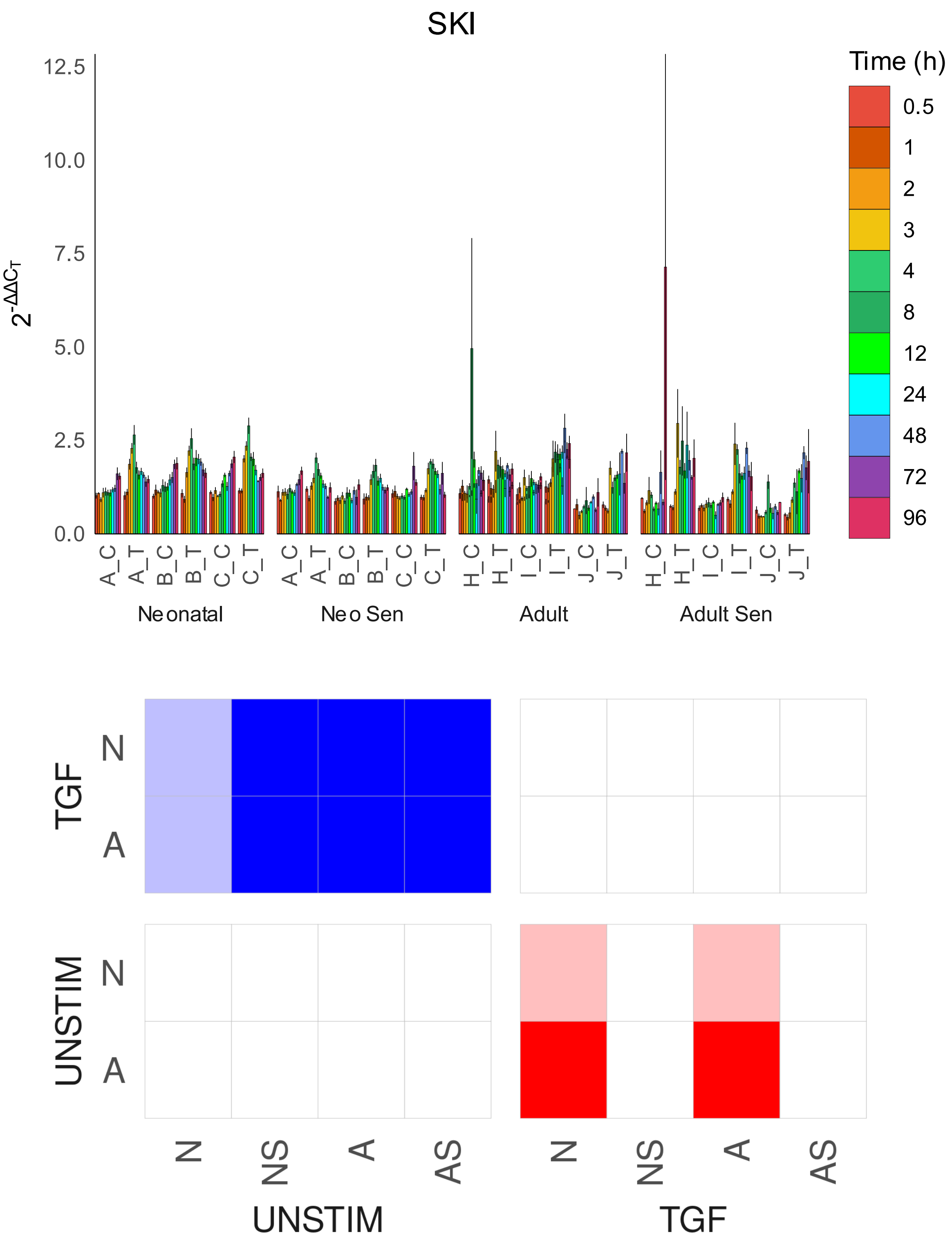
SKI
Time (h)
0.5
1
2
3
4
8
12
24
48
72
96
12.5
10.0
7.5
5.0
2.5
0.0
$2^{-\Delta\Delta C_T}$
A_C A_T B_C B_T C_C C_T
A_C A_T B_C B_T C_C C_T
H_C H_T I_C I_T J_C J_T
H_C H_T I_C I_T J_C J_T
Neonatal
Neo Sen
Adult
Adult Sen
TGF
UNSTIM
N
A
N
A
N NS A AS
N NS A AS
UNSTIM
TGF

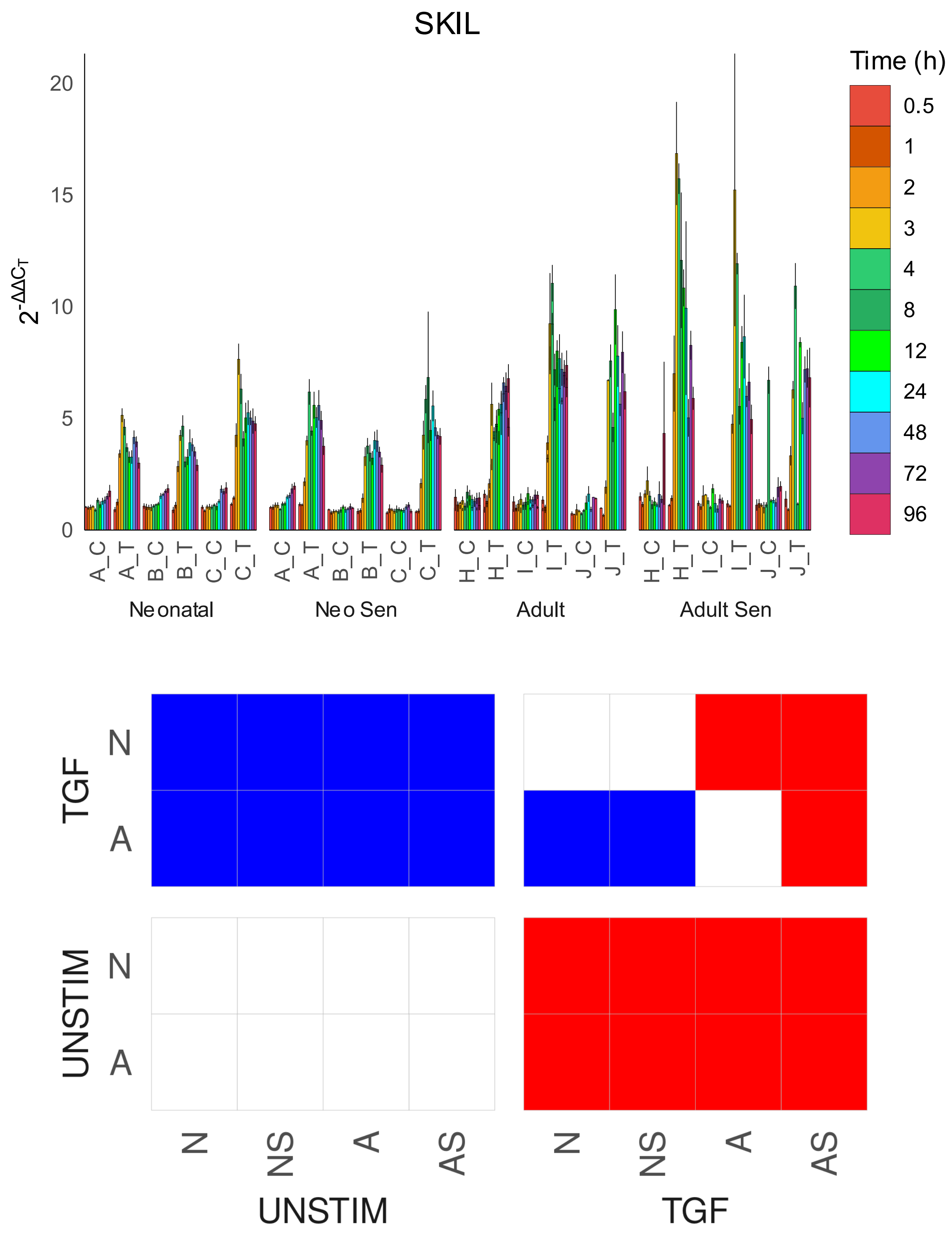
SKIL
Time (h)
0.5
1
2
3
4
8
12
24
48
72
96
$2^{-\Delta\Delta C_T}$
0
5
10
15
20
A_C A_T B_C B_T C_C C_T
A_C A_T B_C B_T C_C C_T
H_C H_T I_C I_T J_C J_T
H_C H_T I_C I_T J_C J_T
Neonatal
Neo Sen
Adult
Adult Sen
TGF
UNSTIM
N
A
N
A
N NS A AS
N NS A AS
UNSTIM
TGF

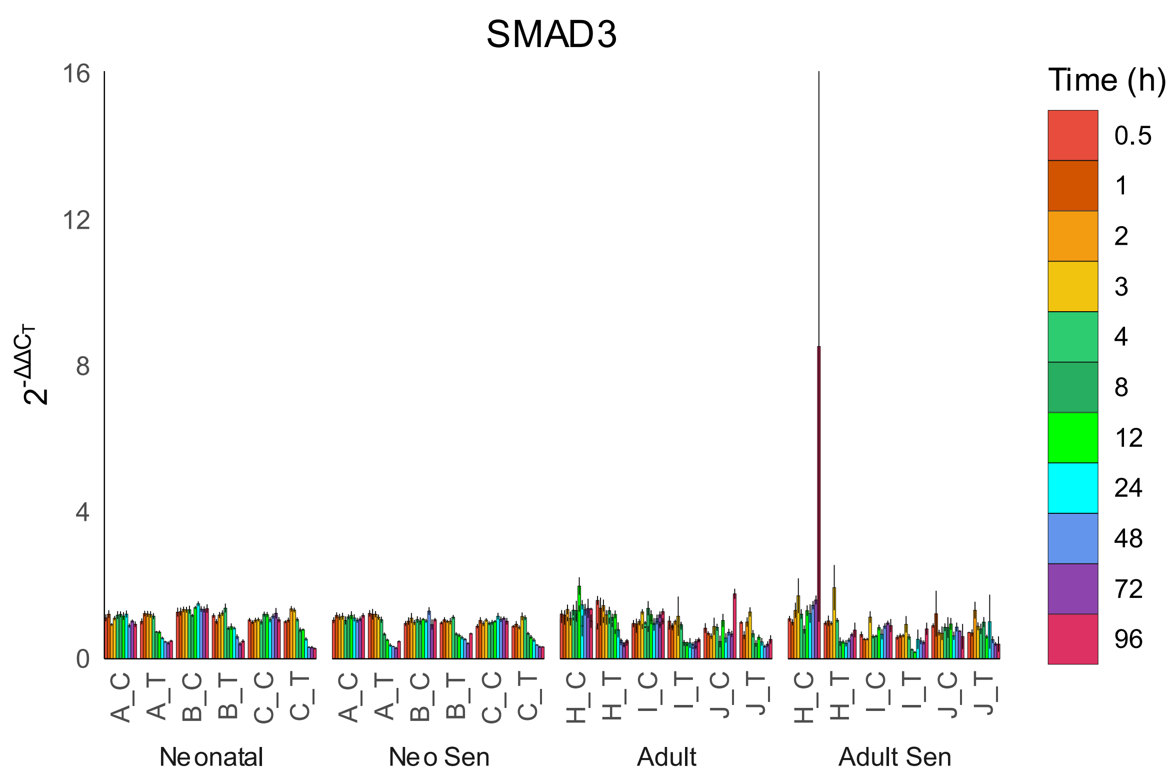
SMAD3
16
12
8
4
0
$2^{-\Delta\Delta C_T}$
Time (h)
0.5
1
2
3
4
8
12
24
48
72
96
A_C
A_T
B_C
B_T
C_C
C_T
H_C
H_T
I_C
I_T
J_C
J_T
Neonatal
Neo Sen
Adult
Adult Sen

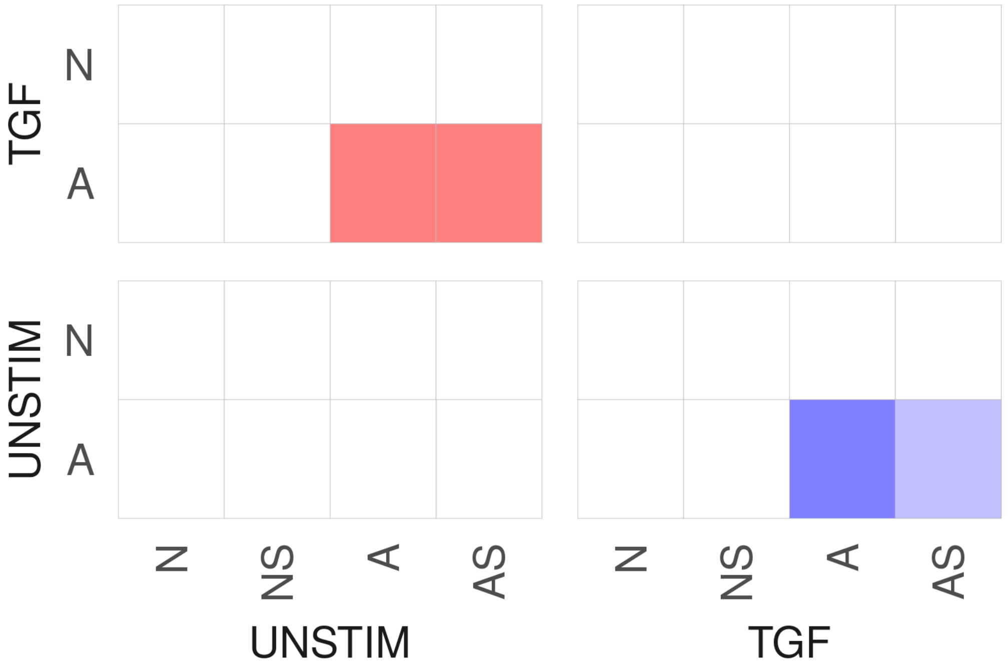
TGF
UNSTIM
N
A
N
NS
A
AS
UNSTIM
TGF

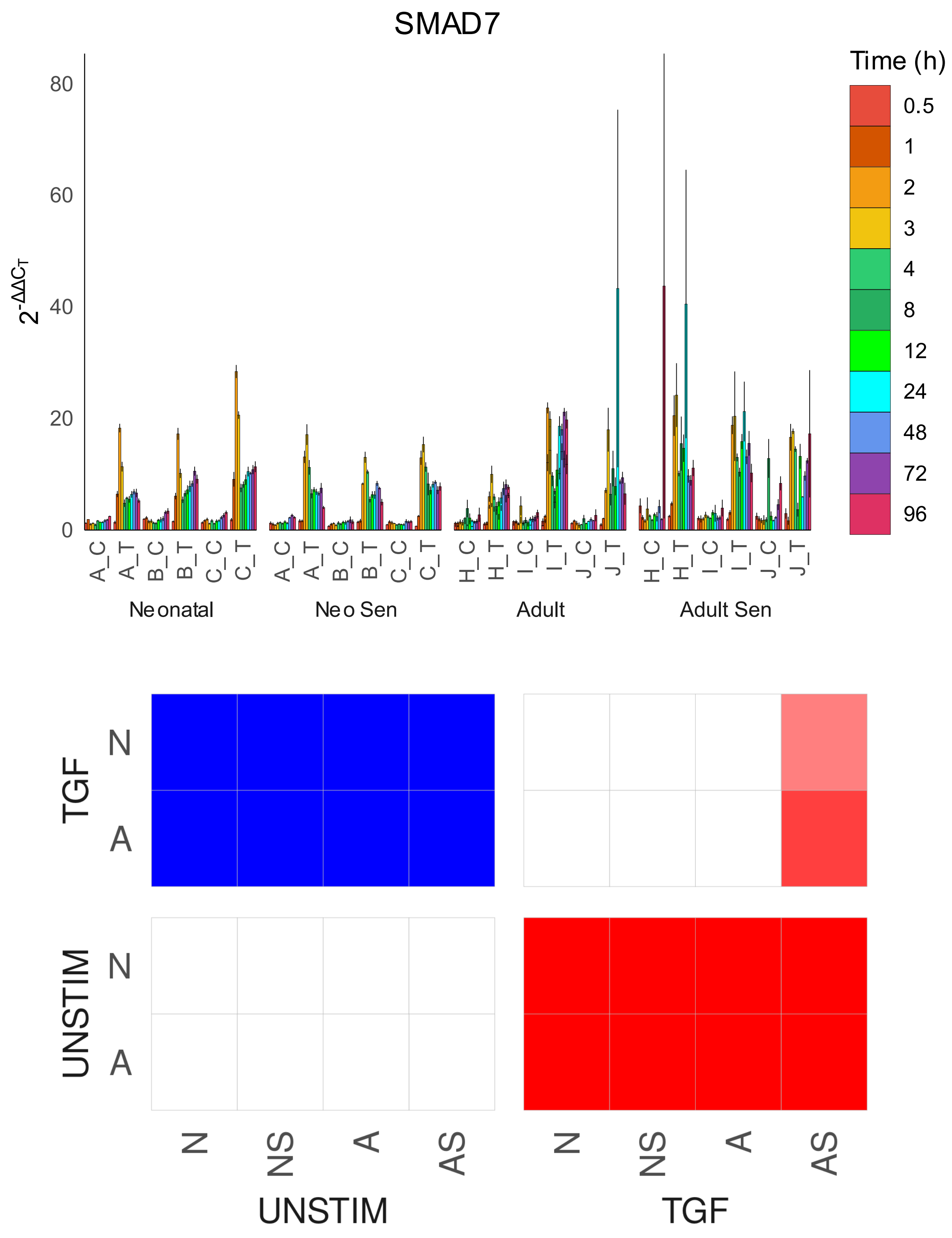

SMAD7
Time (h)
0.5
1
2
3
4
8
12
24
48
72
96
$2^{-\Delta\Delta C_T}$
80
60
40
20
0
A_C
A_T
B_C
B_T
C_C
C_T
H_C
H_T
I_C
I_T
J_C
J_T
Neonatal
Neo Sen
Adult
Adult Sen
TGF
UNSTIM
N
A
N
NS
A
AS
UNSTIM
TGF

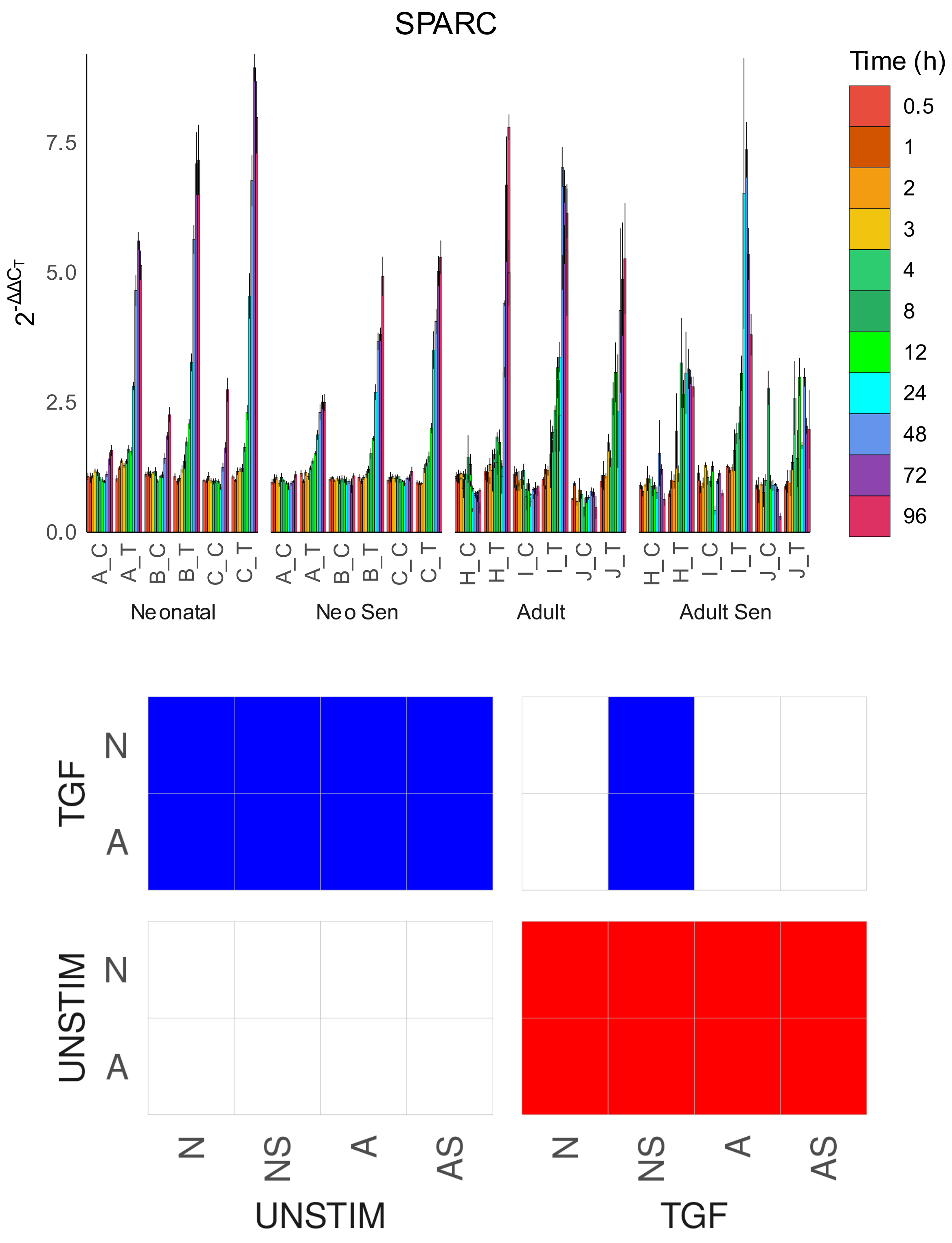
SPARC
Time (h)
0.5
1
2
3
4
8
12
24
48
72
96
$2^{-\Delta\Delta C_T}$
7.5
5.0
2.5
0.0
A_C A_T B_C B_T C_C C_T
Neonatal
A_C A_T B_C B_T C_C C_T
Neo Sen
H_C H_T I_C I_T J_C J_T
Adult
H_C H_T I_C I_T J_C J_T
Adult Sen
TGF
N
A
UNSTIM
N
A
N NS A AS
UNSTIM
N NS A AS
TGF

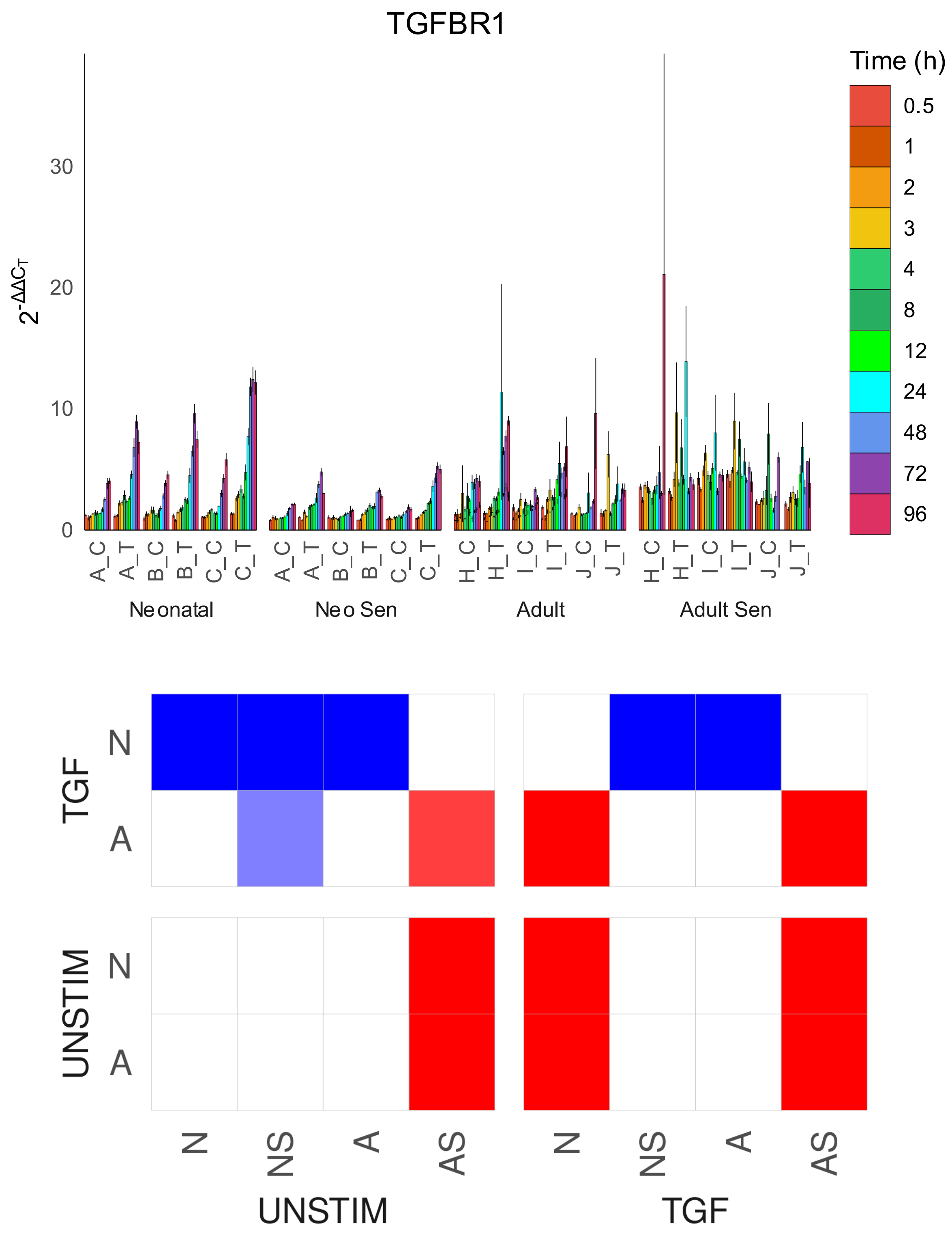
TGFBR1
$2^{-\Delta\Delta C_T}$
0
10
20
30
A_C
A_T
B_C
B_T
C_C
C_T
H_C
H_T
I_C
I_T
J_C
J_T
Neonatal
Neo Sen
Adult
Adult Sen
Time (h)
0.5
1
2
3
4
8
12
24
48
72
96
TGF
UNSTIM
N
A
N
NS
A
AS
UNSTIM
TGF

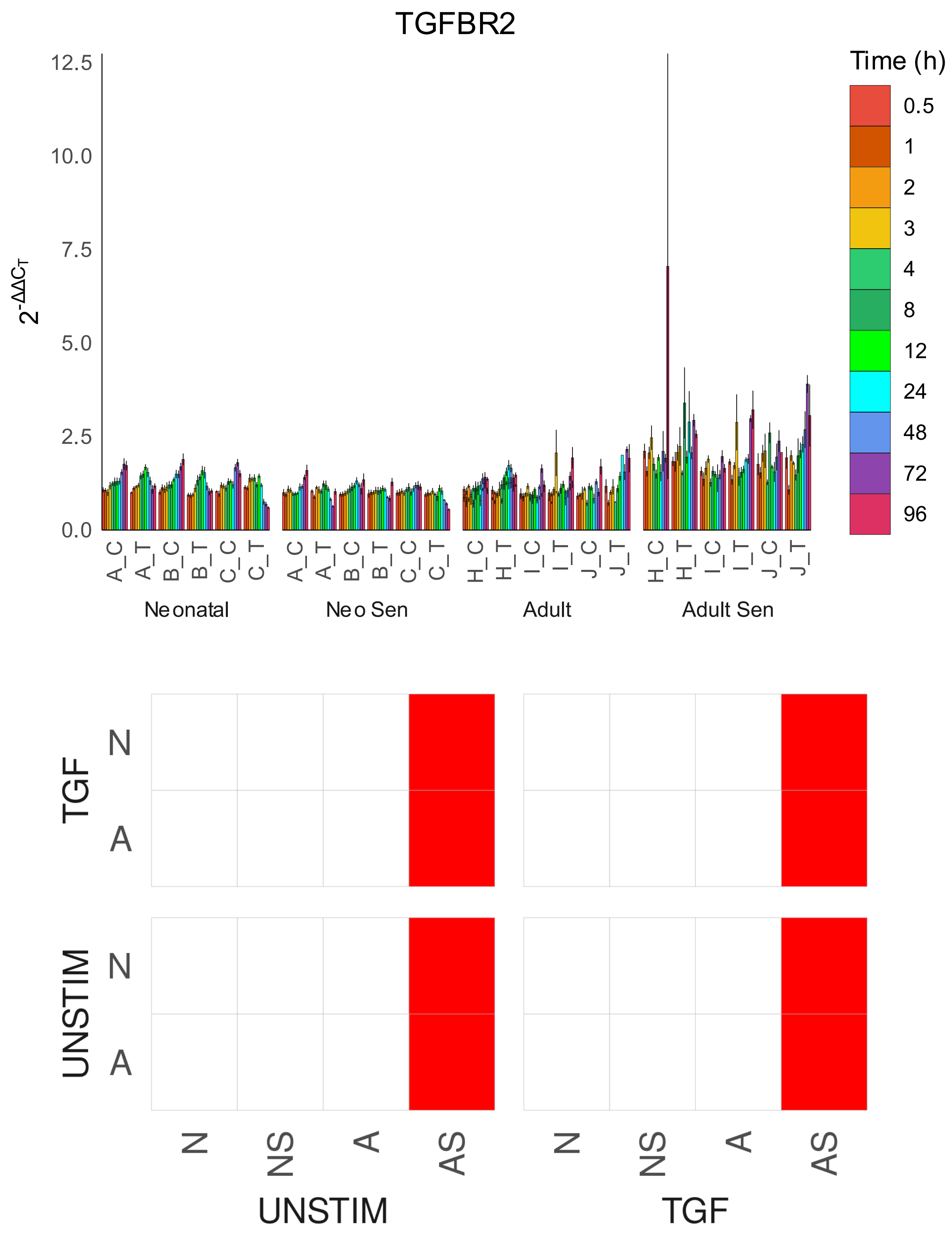

TGFBR2
Time (h)
0.5
1
2
3
4
8
12
24
48
72
96
12.5
10.0
7.5
5.0
2.5
0.0
$2^{-\Delta\Delta C_T}$
A_C
A_T
B_C
B_T
C_C
C_T
H_C
H_T
I_C
I_T
J_C
J_T
Neonatal
Neo Sen
Adult
Adult Sen
TGF
UNSTIM
N
A
N
NS
A
AS
UNSTIM
TGF

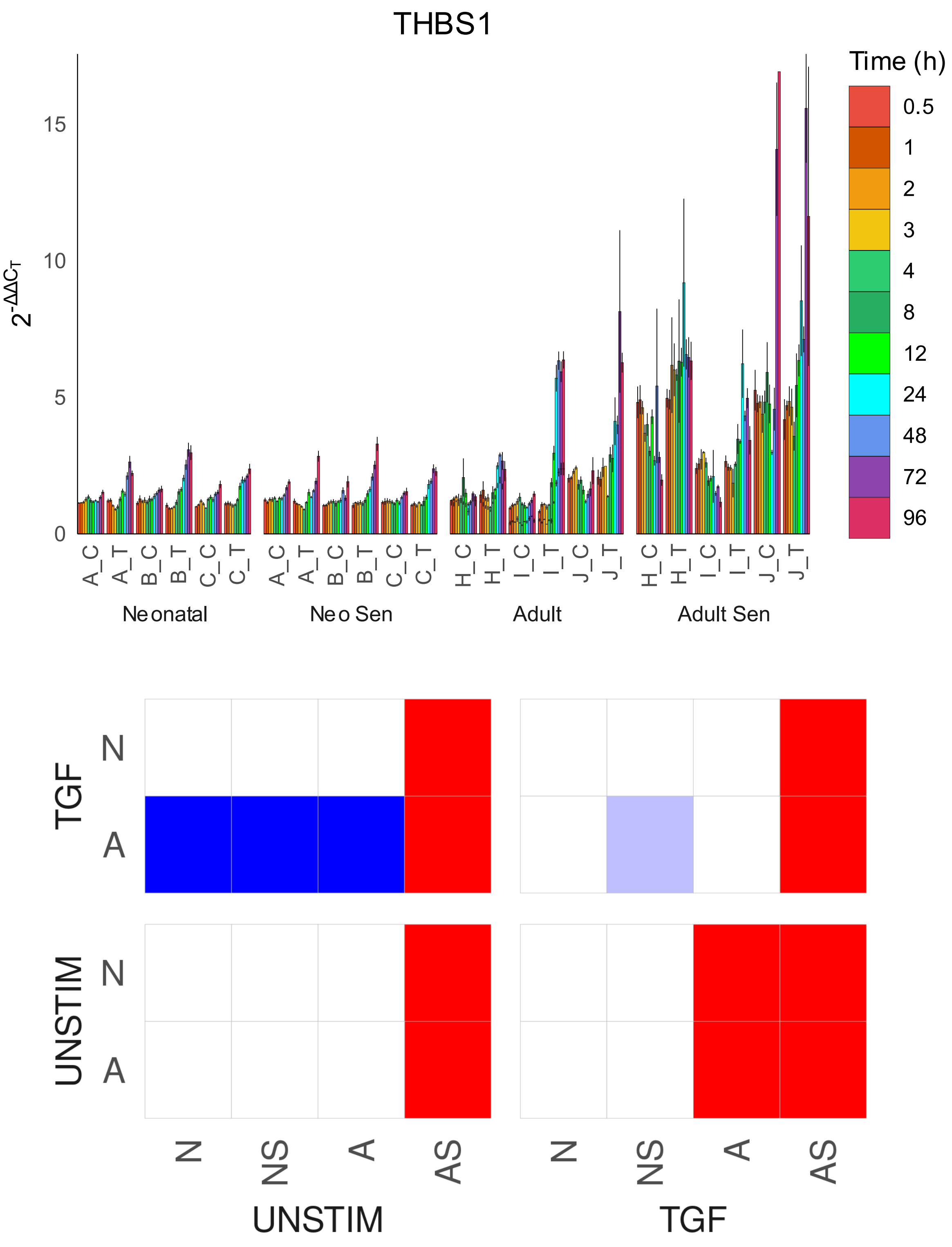

THBS1
$2^{-\Delta\Delta C_T}$
Time (h)
0.5
1
2
3
4
8
12
24
48
72
96
Neonatal
Neo Sen
Adult
Adult Sen
TGF
UNSTIM
N
A
N
NS
A
AS
UNSTIM
TGF

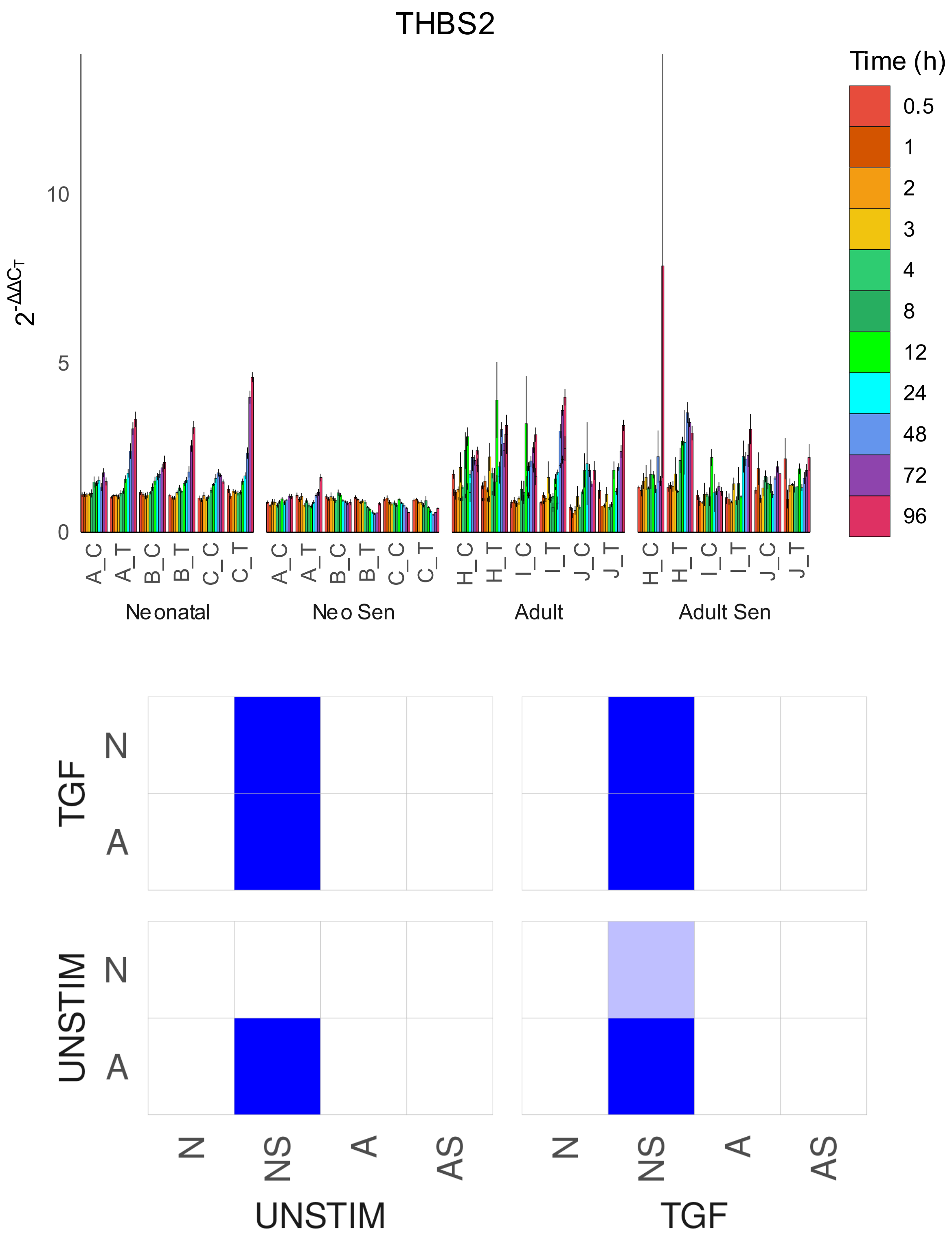
THBS2
Time (h)
0.5
1
2
3
4
8
12
24
48
72
96
$2^{-\Delta\Delta C_T}$
0
5
10
A_C
A_T
B_C
B_T
C_C
C_T
H_C
H_T
I_C
I_T
J_C
J_T
Neonatal
Neo Sen
Adult
Adult Sen
TGF
UNSTIM
N
A
N
NS
A
AS
UNSTIM
TGF

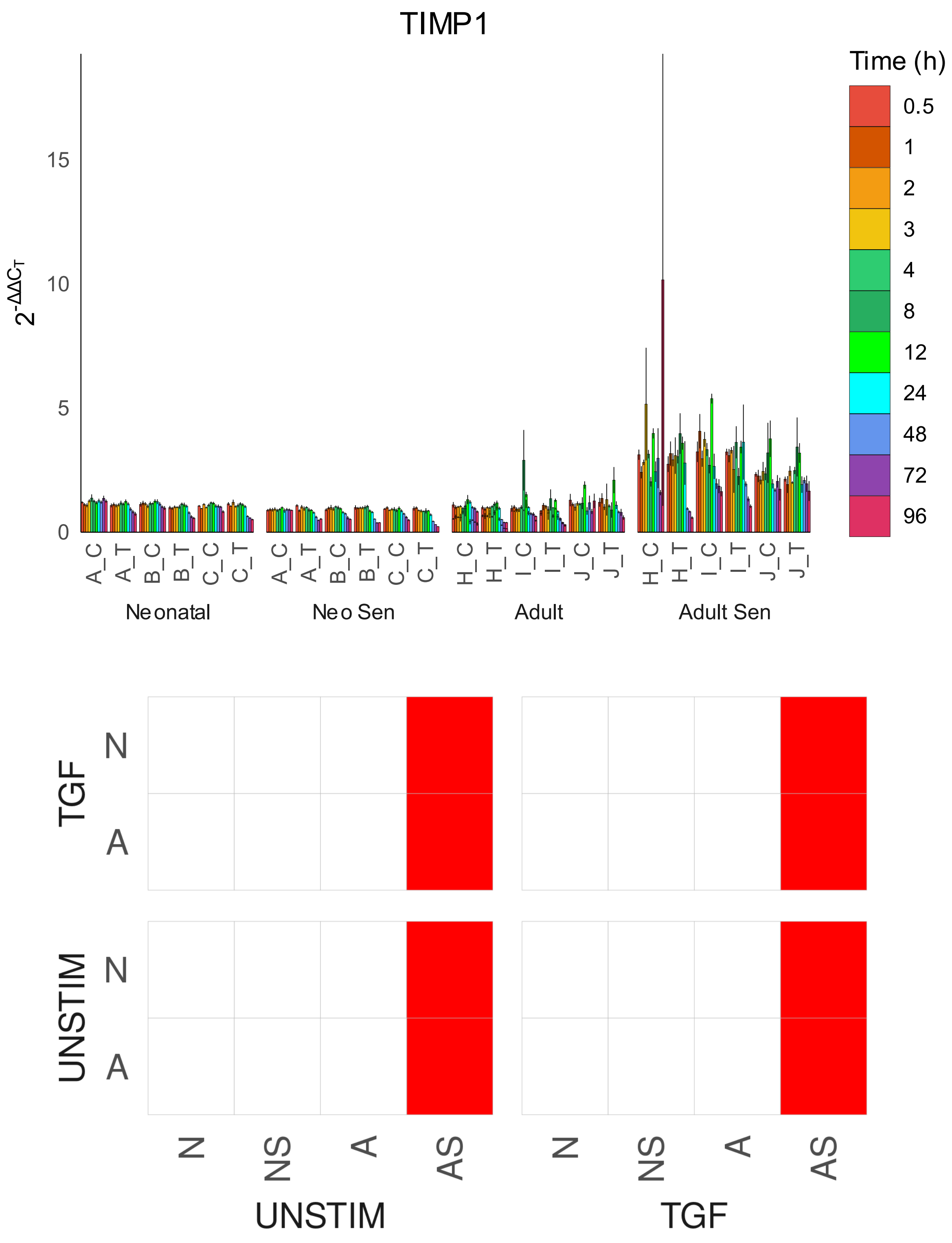

TIMP1
Time (h)
0.5
1
2
3
4
8
12
24
48
72
96
15
10
5
0
A_C
A_T
B_C
B_T
C_C
C_T
H_C
H_T
I_C
I_T
J_C
J_T
Neonatal
Neo Sen
Adult
Adult Sen
TGF
UNSTIM
N
A
N
NS
A
AS
UNSTIM
TGF

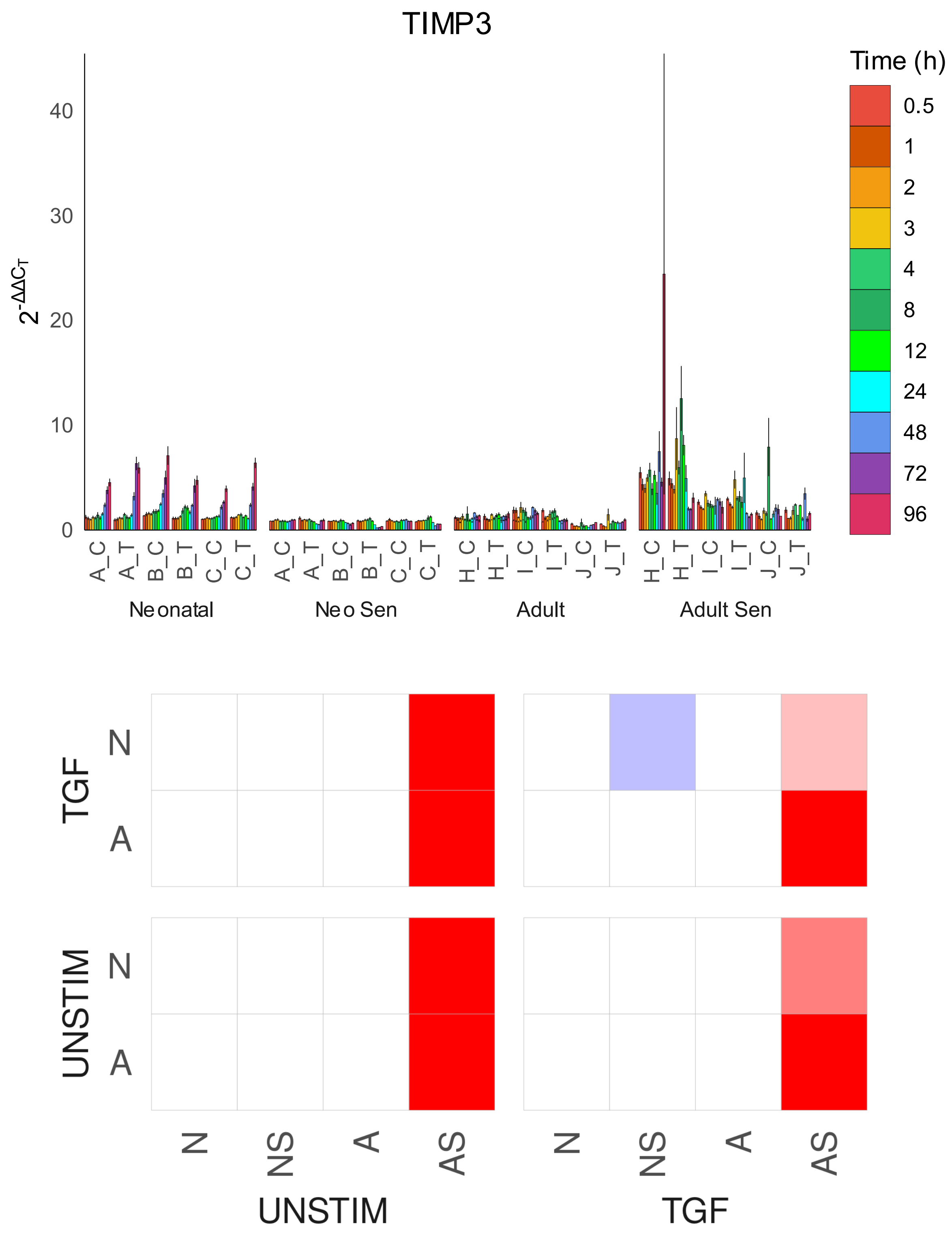
TIMP3
Time (h)
0.5
1
2
3
4
8
12
24
48
72
96
$2^{-\Delta\Delta C_T}$
0
10
20
30
40
A_C
A_T
B_C
B_T
C_C
C_T
H_C
H_T
I_C
I_T
J_C
J_T
Neonatal
Neo Sen
Adult
Adult Sen
TGF
UNSTIM
N
A
N
NS
A
AS
UNSTIM
TGF

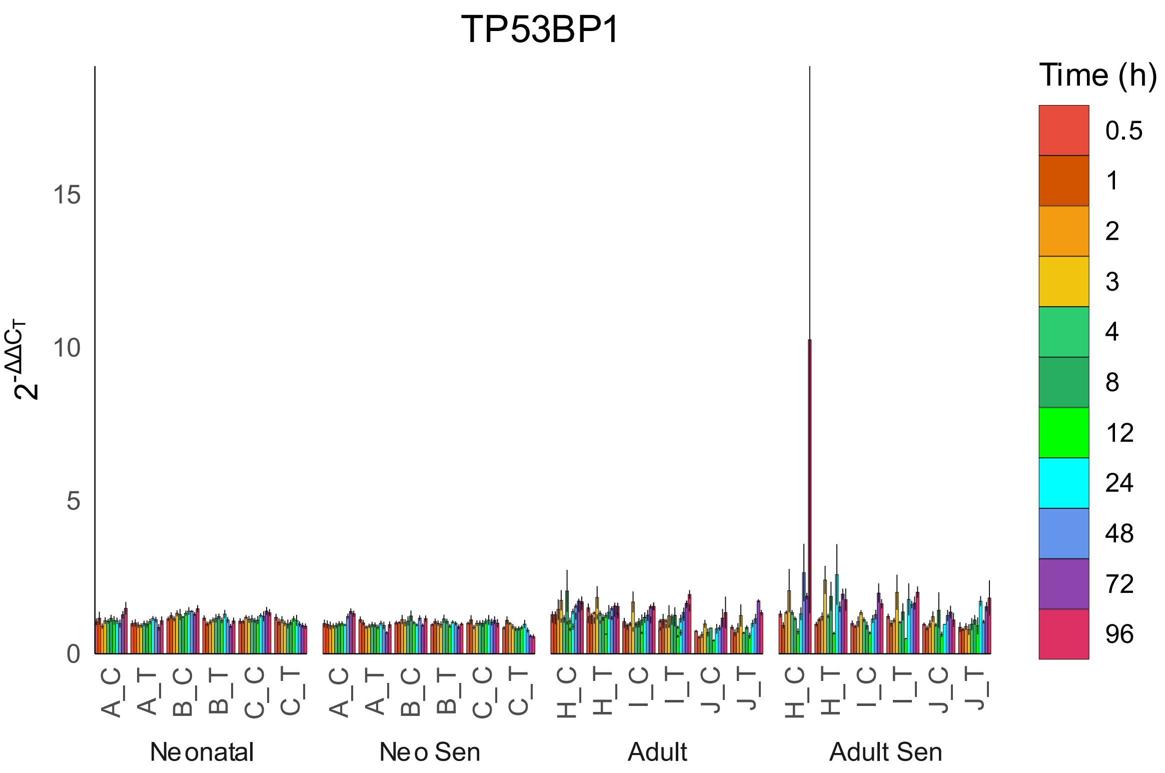

TP53BP1
Time (h)
0.5
1
2
3
4
8
12
24
48
72
96
$2^{-\Delta\Delta C_T}$
15
10
5
0
A_C
A_T
B_C
B_T
C_C
C_T
H_C
H_T
I_C
I_T
J_C
J_T
Neonatal
Neo Sen
Adult
Adult Sen


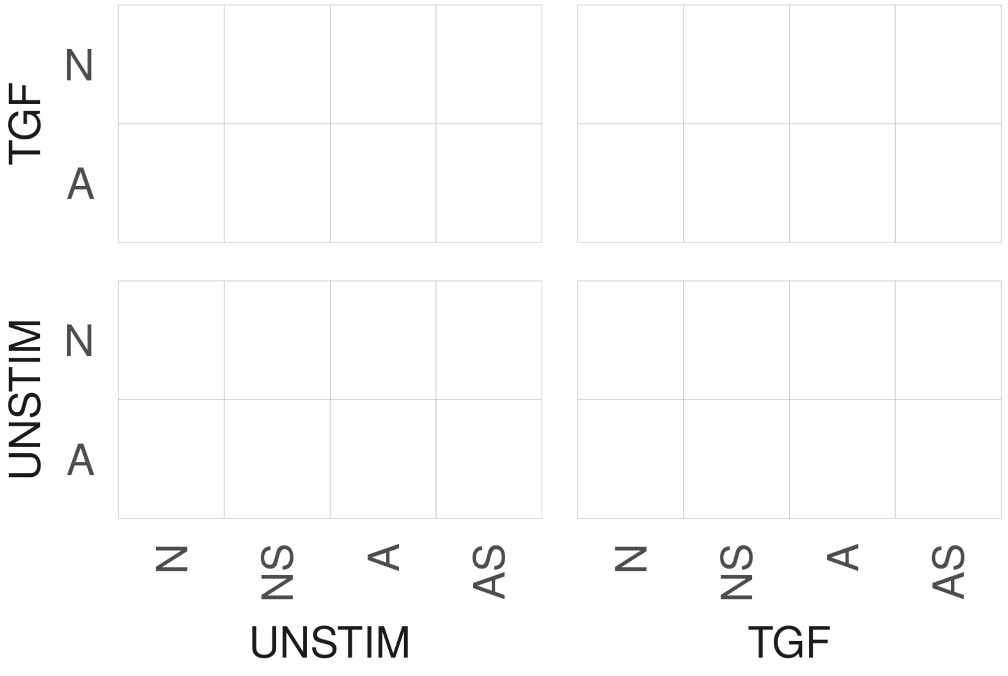

TGF
UNSTIM
N
A
N
NS
A
AS
UNSTIM
TGF

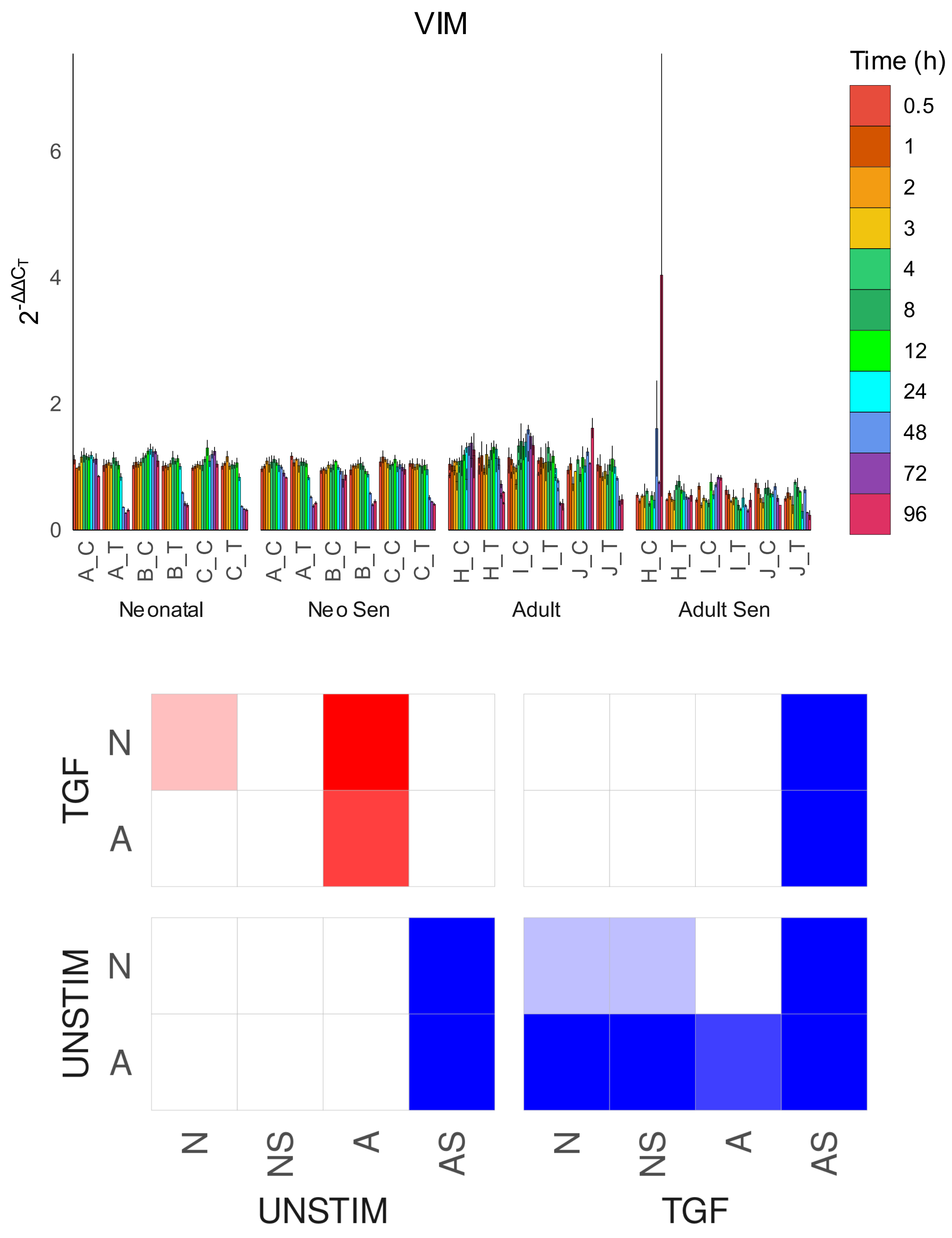

VIM
Time (h)
0.5
1
2
3
4
8
12
24
48
72
96
$2^{-\Delta\Delta C_T}$
0
2
4
6
A_C
A_T
B_C
B_T
C_C
C_T
H_C
H_T
I_C
I_T
J_C
J_T
Neonatal
Neo Sen
Adult
Adult Sen
TGF
UNSTIM
N
A
N
NS
A
AS
UNSTIM
TGF
