## Supplementary Figures for "Cell Senescence-Independent Ageing of Human Skin"

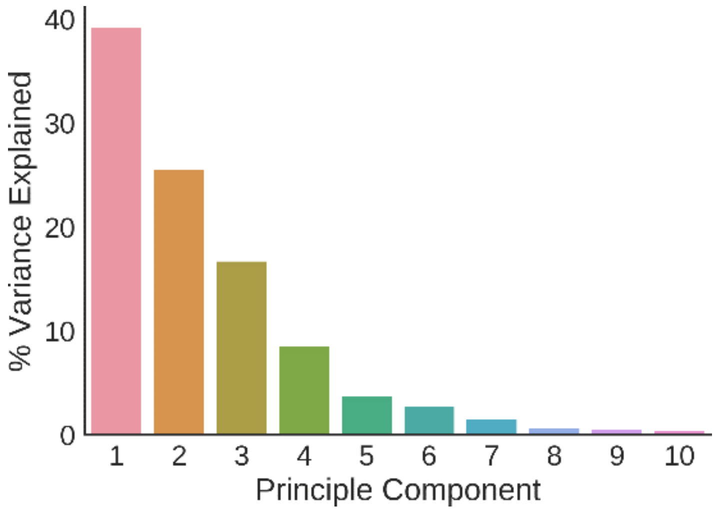


*Figure 1 | Plot showing the percent of variance explained for the first 10 principal components.*

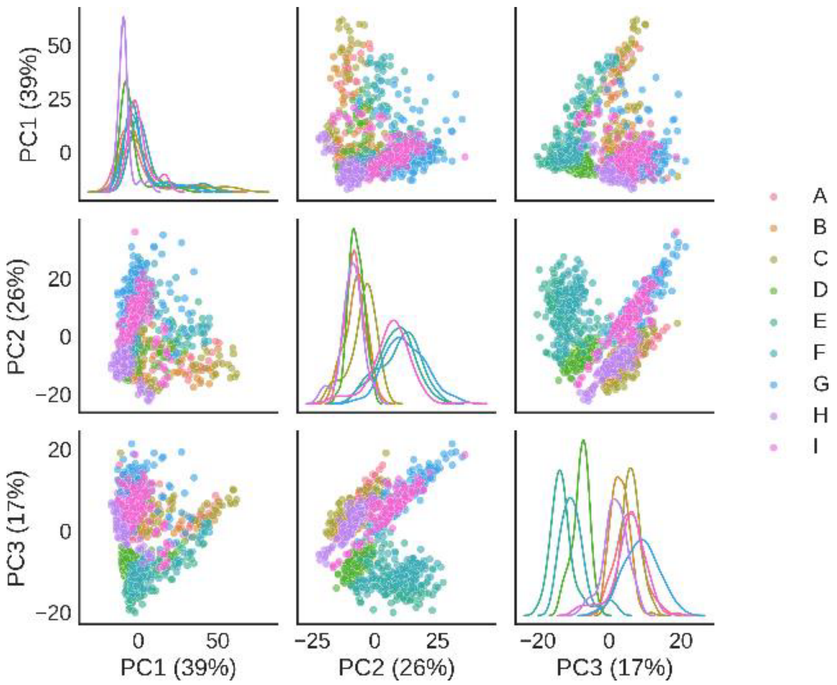


*Figure 2 |Scatter matrix of first three principle components coloured by cell id.*

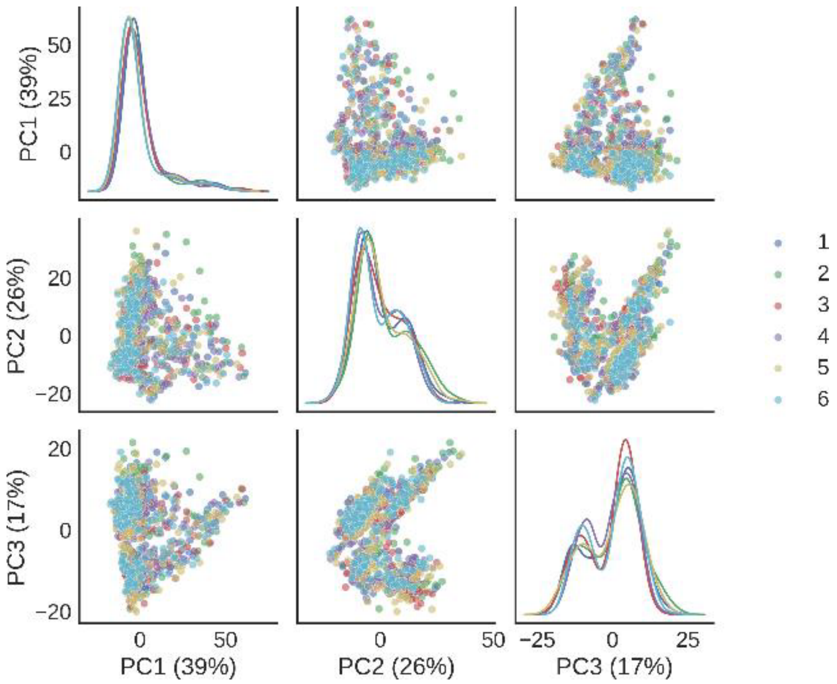


*Figure 3 | Scatter matrix of first three principle components coloured by replicate.*

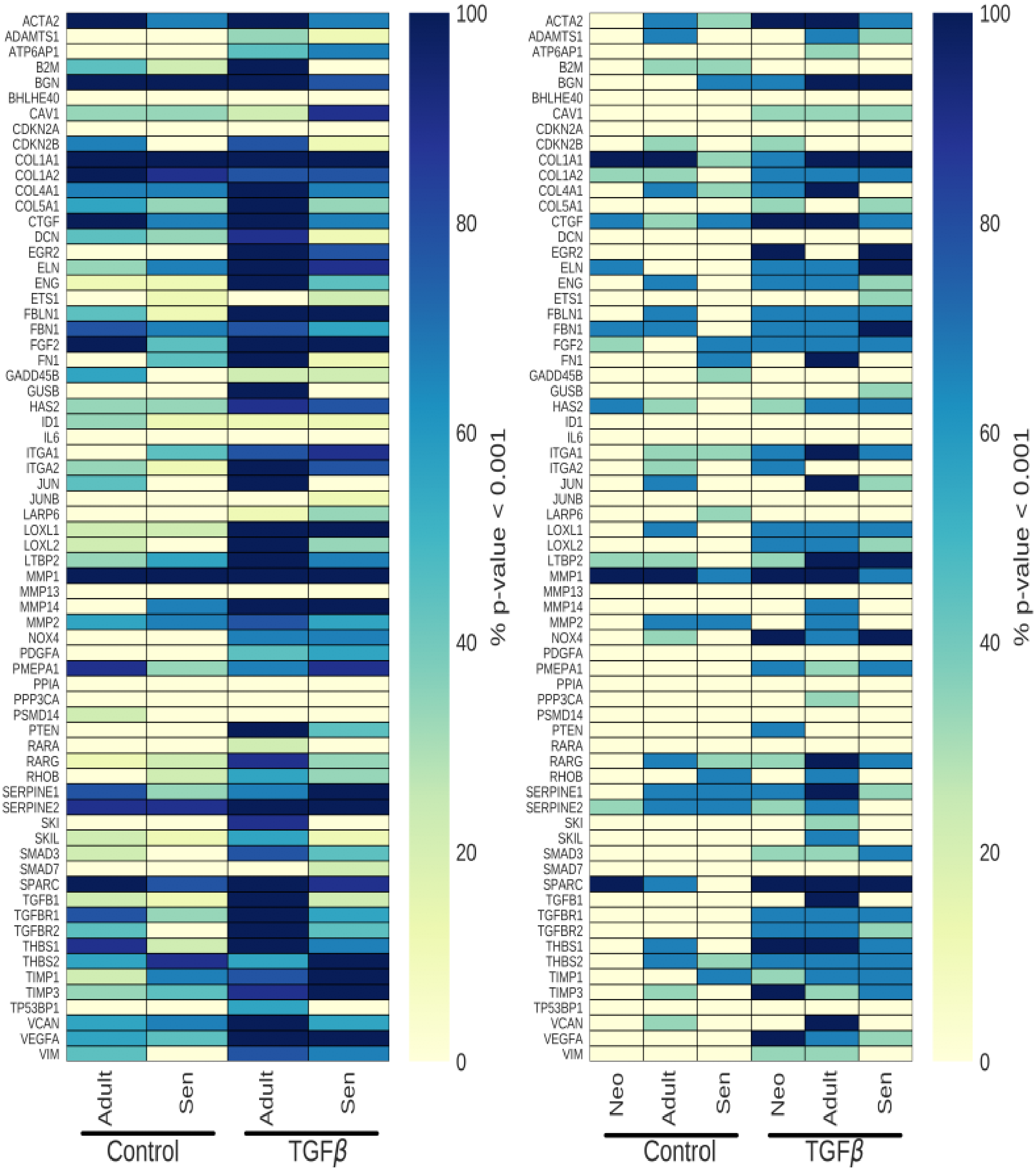


*Figure 4 | Differential expression analysis for (a) between groups comparisons and (b) within groups comparisons. Colours indicate the percentage of the time of all combinations (9 for between groups and 3 for within groups) of LIMMA analysis conducted that a time series was differentially expressed with a FDR corrected p-value<0.001.*

| Comparison | Control | TGF-β |
|---|---|---|
| Adult Vs neonatal | 16 | 46 |
| Senescent Vs neonatal | 16 | 30 |
| Neonatal Vs neonatal | 7 | 28 |
| Adult Vs adult | 17 | 37 |
| Senescent Vs senescent | 10 | 25 |

*Table 1 | Table summarising the number of genes per comparison that were differentially expressed with a (FDR corrected) p-value <0.001 in over 60% of LIMMA analyses per comparison.*
